# Systematically testing human HMBS missense variants to reveal mechanism and pathogenic variation

**DOI:** 10.1101/2023.02.06.527353

**Authors:** Warren van Loggerenberg, Shahin Sowlati-Hashjin, Jochen Weile, Rayna Hamilton, Aditya Chawla, Marinella Gebbia, Nishka Kishore, Laure Frésard, Sami Mustajoki, Elena Pischik, Elena Di Pierro, Michela Barbaro, Ylva Floderus, Caroline Schmitt, Laurent Gouya, Alexandre Colavin, Robert Nussbaum, Edith C. H. Friesema, Raili Kauppinen, Jordi To-Figueras, Aasne K. Aarsand, Robert J. Desnick, Michael Garton, Frederick P. Roth

## Abstract

Defects in hydroxymethylbilane synthase (HMBS) can cause Acute Intermittent Porphyria (AIP), an acute neurological disease. Although sequencing-based diagnosis can be definitive, ~⅓ of clinical HMBS variants are missense variants, and most clinically-reported HMBS missense variants are designated as “variants of uncertain significance” (VUS). Using saturation mutagenesis, *en masse* selection, and sequencing, we applied a multiplexed validated assay to both the erythroid-specific and ubiquitous isoforms of HMBS, obtaining confident functional impact scores for >84% of all possible amino-acid substitutions. The resulting variant effect maps generally agreed with biochemical expectation. However, the maps showed variants at the dimerization interface to be unexpectedly well tolerated, and suggested residue roles in active site dynamics that were supported by molecular dynamics simulations. Most importantly, these HMBS variant effect maps can help discriminate pathogenic from benign variants, proactively providing evidence even for yet-to-be-observed clinical missense variants.

## Introduction

Acute Intermittent Porphyria (AIP; MIM# 176000) is caused by deficiency in the heme biosynthetic enzyme hydroxymethylbilane synthase (HMBS; MIM# 609806, EC 4.3.1.8; also known as porphobilinogen deaminase (PBGD))^1^. AIP, the most frequent acute hepatic porphyria, is an autosomal dominant disorder with an estimated prevalence of 1 in ~1,700 ^2^. The clinical penetrance of AIP is low (1-38%), with ~65% of AIP heterozygotes remaining asymptomatic (i.e. having ‘latent AIP’) throughout their lives ^2–4^. AIP is characterized by potentially life-threatening acute attacks, precipitated by genetic and environmental factors that induce aminolevulinate synthase 1 (ALAS1; MIM# 125290), the first and rate-limiting enzyme in heme synthesis. The *HMBS* gene encodes both erythroid-specific and ubiquitous (housekeeping) isozymes, differing in that the erythroid isoform has a later translational start site that eliminates 17 amino acids at the N-terminus ^5,6^. In classical AIP, both erythroid-specific and ubiquitous HMBS isoforms are deficient. However, ~5% of AIP cases are non-erythroid, caused by a variant that affects only the ubiquitous isoform of HMBS ^7^. In cases of increased heme demand in the liver, the combination of ALAS1 induction with HMBS deficiency results in the accumulation of the porphyrin precursor porphobilinogen (PBG), as well as δ-aminolevulinic acid (ALA),^1,8^ which is likely neurotoxic.

The fact that AIP is a rare disorder, with attacks that are typically episodic and have nonspecific neurovisceral symptoms, can delay clinical recognition and intervention ^3,9,10^. In a symptomatic patient, biochemical diagnosis of AIP is based on demonstrating highly elevated plasma or urine levels of ALA and PBG, after excluding other acute porphyrias by analysis of porphyrin markers in urine and feces. Where a suspected AIP attack is not reported to clinicians in a timely fashion, however, biochemical diagnosis may be more complicated. Both in the latter scenario and more generally, sequencing the three acute porphyria genes *(HMBS, PPOX, CPOX)* to identify a causative variant in a patient with an acute porphyria can be useful. Sequencing has further utility in confirming diagnosis and identifying the *HMBS* variant causing AIP, which facilitates screening of healthy family members to identify individuals who are at-risk for AIP (i.e. have latent AIP). Those with identified latent AIP can then be recommended lifestyle and medication measures to reduce the risk of acute attacks, and be monitored for development of long-term complications such as primary liver cancer ^11^. In recent years, HMBS has been included in gene panels both for inborn error of metabolism and familial liver cancer and has also been suggested to be a tumor suppressor gene ^12^.

Out of the 356 clinical HMBS variants reported in ClinVar, 100(28%) have been annotated as a “variant of uncertain significance” (VUS), with most (66%) of these VUS being missense variants ^13^. The increasing role of sequencing-based diagnostics underlines the importance of providing better tools for variant classification, especially for missense variants. Functional assays can provide strong evidence for clinical variant interpretation but, where they are performed at all, they are typically resource intensive. Moreover, functional assays are typically ‘reactive’, performed only after (and often years after) the first clinical presentation of a variant. By contrast, computational methods can predict the impact of all missense variants ‘proactively’, in advance of the first clinical presentation. Although computational predictors are steadily improving ^14^, this type of evidence is considered weak at best under current American College of Medical Genetics and Genomics and Association for Molecular Pathology (ACMG/AMP) guidelines ^15^.

The functional impact of essentially all possible single variants in a given target protein can be revealed by multiplexed assays of variant effect ^16^. Variant effect maps can provide accurate and proactive identification of dysfunctional alleles ^17,18^; for example, analysis of variant effect maps for four cancer-related genes yielded reclassification for 25%-73% of clinical missense VUS ^19–22^.

Model organism assays, e.g., measuring the ability of a human protein variant to complement loss of the corresponding ortholog’s activity in that model organism, can enable inference of human variant pathogenicity ^23–25^. Here, we employ a multiplexed yeast-based assay of the human HMBS protein to proactively and systematically measure missense variant impacts for both erythroid and ubiquitous HMBS isoforms. We find that the resulting impact scores correspond well with prior knowledge about the atomic structure of human HMBS, and with known patterns of mutational tolerance. We use the map, together with molecular dynamics (MD) simulations, to implicate residues in the control of backbone flexibility and in motions of a ‘lid’ over the active site. Finally, we demonstrate that variant effect map scores can reliably identify pathogenic HMBS alleles.

## Results

### A scalable functional assay for HMBS missense variants

We implemented a scalable yeast-based functional complementation assay, based on the previous observations^26^ that a strain bearing a temperature-sensitive (ts) mutation in the essential yeast ortholog of *HMBS* (*HEM3*) exhibits reduced growth at the non-permissive temperature, and that expression of the human HMBS protein rescues this phenotype. The complementation relationship was confirmed (Figure S1; see Material and Methods), and we validated the assay (Figure S1) for an HMBS variant set which included four missense variants having a stringent “pathogenic” annotation in ClinVar and three ‘proxy-benign’ (not known to be disease associated) missense variants having allele frequencies that roughly matched those of the pathogenic variants. To assess the functional impact of each variant in the yeast *hem3* ts strain, we again assessed growth at the non-permissive temperature (see Material and Methods). Each variant was assayed alongside strains expressing either the wild-type human protein or an empty vector, respectively serving as positive and negative controls for variant functionality. Yeast complementation assays for a variety of genes were previously shown to detect ~60% of pathogenic variants, at a stringency at which 90% of variants observed to be damaging were pathogenic (i.e., 60% recall at 90% precision) ^23^. In line with this expectation, we observed 50% recall (lack of complementation for two of four pathogenic variants) with 100% precision (complementation for all non-pathogenic variants) (Figure S1).

### Systematic maps of HMBS missense variant functional impact

By coupling an efficient functional complementation assay with the TileSeq framework for multiplexed assays of variant effect, we sought to measure the functional consequences of all possible missense HMBS variants ^27^. First, we constructed libraries of HMBS variants (for both erythroid-specific and ubiquitous isoforms) using our previously described POPCode mutagenesis method ^27^. To balance the objective of having substantial representation of each variant in the library against the objective of having roughly one amino acid substitution per clone, we generated two separate full-length libraries, with mutagenesis targeted to the N- and C-terminal halves of the protein, respectively. Variant libraries were initially generated as a pool of amplicons. Large-scale sequencing showed that mutagenesis was relatively even across each of the libraries for both erythroid and ubiquitous HMBS isoforms, with an average of 1.7 amino acid changes per clone (Figure S2A). Amplicons were transferred *en masse* into the appropriate yeast expression vector via two steps of recombinational subcloning (see Methods). The resulting mutagenized expression libraries were then transformed *en masse* into the appropriate ts yeast strain, yielding ~2 million independent yeast transformants for each library.

To assess the functional impact of many HMBS variants in parallel, pools of yeast HMBS mutant strains were grown competitively on synthetic medium at the non-permissive temperature. The frequencies of each HMBS variant within this laboratory strain population were then determined, before and after selection, using the TileSeq framework (Figure 1A)^27^. Briefly, we designed a set of amplicon ‘tiles’ that collectively span the complete coding region. These tiles are sufficiently short (~150bp) to enable sequencing of both strands (“duplex sequencing”), with variants called only where they are detected on both strands. Each nucleotide position was covered by ~2 million duplex reads. We considered variants appearing frequently enough in the pre-selection library (above 10 counts per million reads sequenced) to be well measured. This criterion was satisfied by >88% of all possible missense variants, and by >95% of the amino-acid substitutions that can be achieved via a single-nucleotide variant (SNV), for both the erythroid-specific and ubiquitous isoforms (Figure S2B). Comparing post- to pre-selection variant frequencies, we calculated a functional impact score (see Methods), in both protein isoforms, for nearly all HMBS amino acid substitutions.

**Figure 1.**
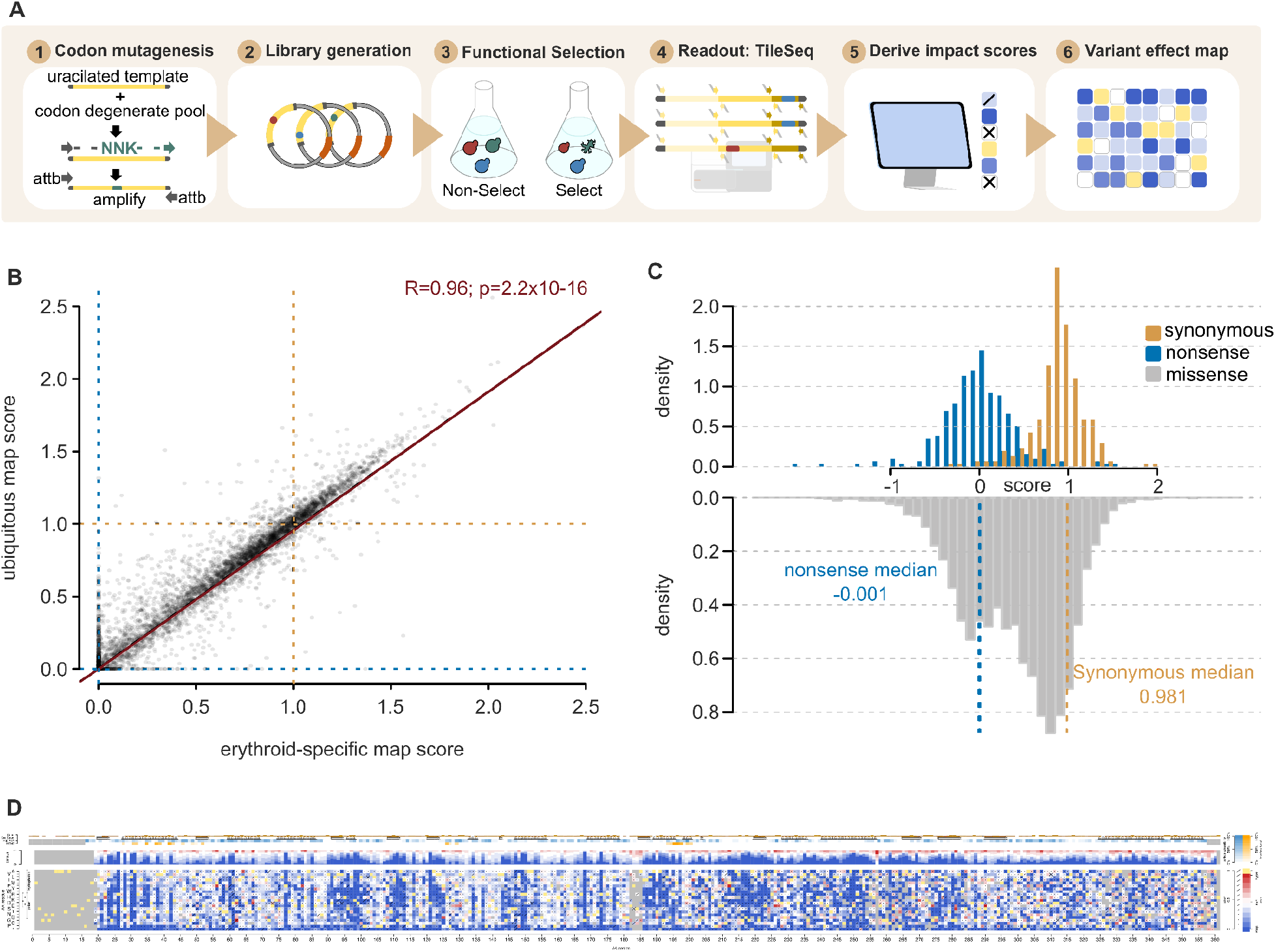
Generating and evaluating HMBS variant effect maps (A) Workflow for generating HMBS variant effect maps. (B) Correspondence between erythroid-specific and ubiquitous HMBS isoform functional scores. For reference, null- and WT-like scores are indicated with dashed red or green lines, respectively, while the blue line corresponds to a linear regression fit (R=0.96; P = 2.2×10^−16^). (C) Distributions of functional impact scores of nonsense (red), synonymous (green), and missense variants (gray) from the combined erythroid-specific and ubiquitous HMBS map (left). (D) Preview of full-sized combined HMBS map

We estimate uncertainty (standard error) for each functional impact score, based both on the agreement between two replicates and on trends in the behavior of replicates for other variants with similar pre-selection variant frequency (see Methods)^27,28^. Impact scores with an estimated standard error above 0.3 were removed, yielding measurements of functional impact for >6,000 missense variants for each isoform. Thus, we obtained high-confidence functional impact scores for 87% and 84% of all possible amino acid substitutions in the erythroid and ubiquitous HMBS isoforms, respectively (Figure S2B). These included 93% (for the erythroid isoform) and 90% (for the ubiquitous isoform) of the amino-acid substitutions accessible by a single-nucleotide change.

Impact scores for erythroid and ubiquitous maps were highly correlated (Pearson’s r = 0.96, Figure 1B). Indeed, where scores were available from both maps, no convincing difference between the maps was observed in any segment of HMBS. We therefore calculated a weighted average score for each variant (see Methods) to generate a single combined map. All scores for the erythroid, ubiquitous and combined variant effect maps are publicly available (MaveDB^29^ accession urn:mavedb:00000108-a).

For each individual map, as well as the combined map, the impact score distributions of synonymous and nonsense variants were well separated (Figure S2C, Figure 1C). Missense variants from each map showed a bimodal distribution, suggesting that variants tended to either have strong or neutral functional impacts, as opposed to having intermediate effects. A small fraction (2.5%) of missense variants exhibited ‘hyper-complementation’, such that yeast growth upon expression of the HMBS variant exceeded that observed for WT human HMBS.

### Hyper-complementing HMBS variants are likely deleterious in humans

It has been previously reported that, for SUMO and UBE2I (the human SUMO E2 conjugase), hyper-complementing variants displaying increased fitness in yeast assays may in fact be disadvantageous^27^. We explored this idea for HMBS using a quantitative phylogenetic approach ^30,31^ that compares three hypotheses about the effects of hyper-complementing variants: 1) variants that confer an advantage in our maps will also do so in humans and related species; 2) hyper-complementing variants are equal in function to wild-type; and 3) hyper-complementing variants are deleterious in humans and related species (a model in which the functional score in humans is modeled as the reciprocal of the observed score in yeast). We found the third (deleterious) model to be the best-performing for HMBS (Table S1), suggesting that hyper-complementing variants in our yeast assay are best treated as deleterious to humans.

### Functional scores captured known roles for many HMBS missense variants

Several features of HMBS biochemistry were recapitulated in our variant effect maps. HMBS activity begins with condensation of two porphobilinogen (PBG) molecules to assemble dipyrromethane (DPM), to which four additional units of PBG are subsequently condensed (and then released by hydrolysis) to generate hydroxymethylbilane (HMB)^32,33^. Importantly, DPM is bound covalently at C261 and, as expected, our maps found this critical cysteine to be intolerant to mutation (Figure 2A)^34^.

**Figure 2.**
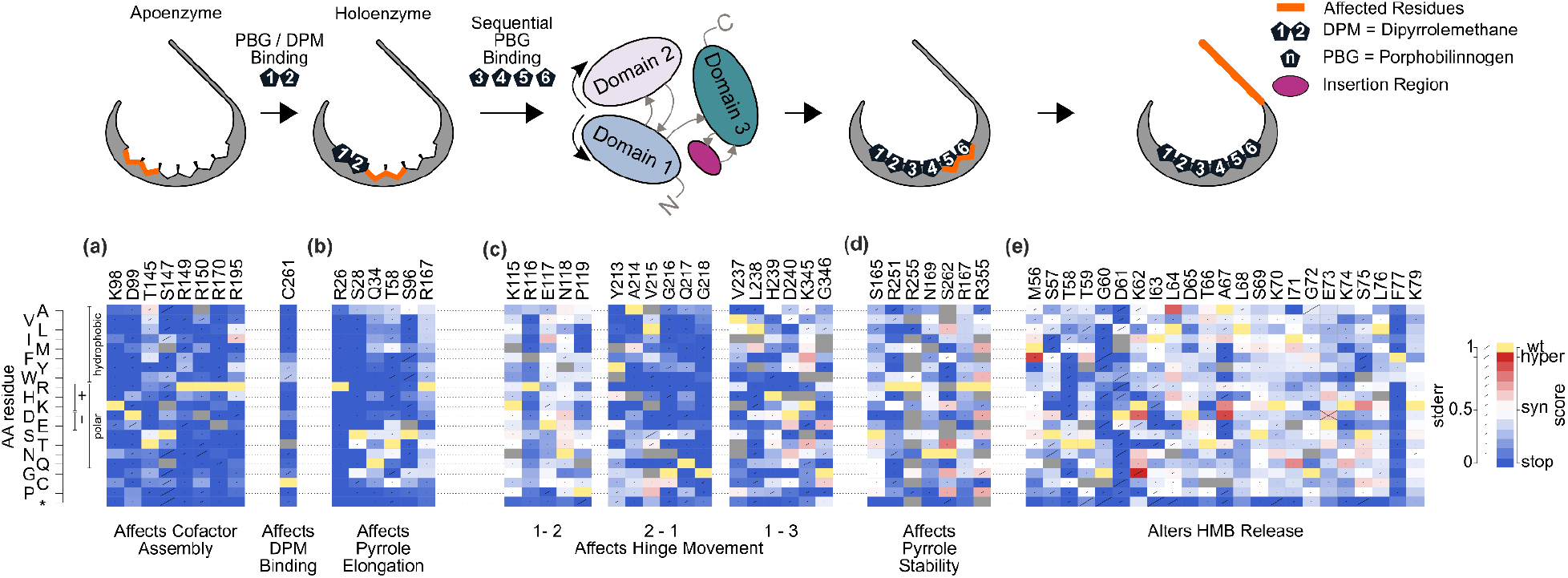
Identifying patterns of mutational tolerance Functional scores for each possible substituted amino acid (*y*-axis) at each active-site residue position (*x*-axis) responsible for: (a) altering cofactor binding; (b) PBG binding for pyrrole chain elongation; (c) hinge flexibility; (d) pyrrole stability; and (e) HMB release. For each substitution, diagonal bar sizes convey estimated measurement error in the corresponding functional score. Box color either indicates the wild-type residue (yellow), a substitution with damaging (blue), tolerated (white), or above-wildtype (‘hyper-complementing’, red) functional score, or missing data (gray).

Several residues are known to be important for DPM assembly, enzyme kinetics and conformational stability, including K98, D99, T145, S147, R149, R150, R173, and R195 ^35,36^. Our maps found all of these critical positions to be generally intolerant to mutation (Figure 2A). Five additional active site residues involved in polypyrrole chain elongation – R26, S28, Q34, T58, and S96 – were each found to be important but slightly more tolerant of variation than those that interact with DPM (Figure 2B)^35^.

Another six residues (S165, N169, R251, R255, S262 and R355) have been predicted to play a key role in stabilizing the growing pyrrole chain ^35,37^. Of these, our map surprisingly only implicated R251 and S262 as being essential for HMBS function (Figure 2D). Also unexpectedly, we found R167, a residue reported to play a dual role in both catalysis and HMB release ^35^, to be highly tolerant to mutation (Figure 2B). R167 is unquestionably important in humans ^2^, with five R167 missense variants having either ‘pathogenic’ and/or ‘likely pathogenic’ classification ^13^. Where a protein has multiple functions and only a subset of these are required to provide growth rescue in a complementation assay, the assay will only detect impacts of a variant on the subset of functions required for rescue. That we find R167 (as well as S165, N169, R255, and R355) to be tolerant to substitution could potentially be explained if the addition of the final two of six PBG monomers were not required to rescue the *hem3 ts* mutation in yeast. One scenario for this is that either a) production of the tetrapyrrole form of HMB is sufficient to sustain growth of the yeast *hem3 ts* strain, or b) if residual activity of the *HEM3* ts mutant can extend the tetrapyrrole form to generate the full hexapyrrole.

The active site loop (residues 56-76), with residues G60-I71 adopting α-helical secondary structure, is known to contribute to the recruitment of PBG and chain elongation ^37^. Within this loop, residues T58, D61, S69 and K70 have been noted as important by some studies, with other studies also implicating residues K74 and K79 ^35,37,38^. We found that substitutions in residues K70, K74 and K79 previously reported as important for stabilizing DPM, were generally tolerated (Figure 2E)^37^. Our data also supports an important role for T58, G60 and D61 residues in enzyme function (Figure 2E).

### Functional impact scores point to key residues modulating HMBS structural fluctuations

The HMBS active site cleft is at the interface of domain 1 (residues 1-114, 219-236) and domain 2 (residues 120-212) ^39^. Structural studies have suggested that movement of HMBS domains 1, 2, and 3 (residues 241-361), facilitated by flexible inter-domain hinge regions, helps accommodate substrates of various sizes during PBG chain elongation ^38,40^. (Here we refer to: “hinge 1-2”, connecting the N-terminal segment of domain 1 with domain 2; “hinge 2-1”, connecting domain 2 with the C-terminal segment of domain 1; and “hinge 1-3”, connecting the C-terminal segment of domain 1 with domain 3.) Consistent with this model, our maps showed severe fitness defects for mutations within hinge 2-1 and hinge 1-3. In addition, G346, positioned within a “hinge 3-3” preceding the C-terminal helix in domain 3, is intolerant to variation (Figure 2C). Surprisingly however, we found hinge 1-2 residues to be highly tolerant to mutation (Figure 2C).

MD simulations have previously suggested that the movement of HMBS domains 1 and 2 relative to domain 3 is constrained by an “insertion region” (residues 296–324) which is absent in bacterial HMBS orthologs ^35^. We performed MD simulations (see Material and Methods) which confirmed both this and the previous suggestion that accommodation of the elongating polypyrrole is assisted by movement of the HMBS cofactor-binding loop in concert with the active-site loop and insertion region (Figure S3)^37,38^. Our maps find the insertion region to be generally tolerant of variation (Figure S4), consistent with a role for the insertion region as a volume-filling ‘wedge’ which separates domain 3 from domains 1 and 2, allowing room for the elongating polypyrrole (a role which does not depend strongly on the precise biochemical nature of specific insertion region residues). Interestingly, a strong functional impact was observed for a set of mutations at the interface of domain 1 and 3: T109, I110, and I113 in domain 1; and G317, I318, T319 and A320 in domain 3 (Figure S4), suggesting that coupling of the mobility of domains 1 and 3 may be more important than previously appreciated.

Mutating the above-mentioned glycine in hinge 3-3 to a proline (G346P) can potentially restrict the flexibility of the backbone and consequently can affect the enzymatic function. The hydrogen bonding pattern of residues at the C-terminal 3-3 hinge region was considerably impacted by the introduction of G346P mutation (Table S4). Our MD simulations showed more persistent interactions between R355 and L257, G259, and D352 (Table S5), and correspondingly reduced flexibility along the C-terminal helix (Figure S5; Table S5). Increased rigidity of the C-terminal helix inhibited mobility of the cofactor-binding loop, which presumably affects accommodation and stabilization of the substrate. Indeed, in the G346P variant the D99-substrate interaction was significantly reduced relative to WT (Table S3).

### Further interrogating key active site loop residues via simulated molecular dynamics

The active site loop, in addition to its roles in recruitment of PBG and chain elongation, has been implicated in two of the three pathways proposed for HMB’s exit from the active site ^35^. Here we used MD simulation to explore two additional hypotheses related to the active site loop.

First, based on analysis of the known structure, we hypothesized that a salt bridge between D61 and K27 controls flexibility and the positioning of the active site loop. Three HMBS variants – D61N, D61A, and the double mutant D61K K27D (representing a ‘swap’ of the amino acids at these two positions) – were investigated virtually using a 500-ns MD simulation. To quantify the relationship between the formation of the D61-K27 salt bridge and the position of the active site loop, the distances between residue pairs G60-R26 and G60-Q34 were measured, and we considered the D61-K27 salt bridge to be present (i.e., in the “closed” conformation) if we observed a D61-K27 distance was equal or below 4 Å, and absent (the “open” conformation) if above 4 Å.

Our simulation showed that the WT apo-enzyme tends to maintain the D61-K27 salt bridge, with a mean distance of 3.5 ± 1.8 Å between D61 and K27 residues, spending ~70% of simulation time in the closed state (Figure S6, Figure 3). Simulations of the D61N, D61A, and D61K K27D variant structures showed that, for each variant, the distance between salt bridge residues increased (10.7 ± 2.2, 8.4 ± 1.6 Å, and 5.4 ± 2.4 Å respectively; Figure S6). The average G60-R26 and G60-Q34 distances also increased (Table S6), with the active site loop of the single mutants remaining in the “open” state (both for ~100% of simulation time), while the double mutant (D61K K27D) was in the closed state ~30% of the time. Because the D61K K27D amino acid swap by itself should not have significantly affected salt bridge formation, the observation that the swap induces the open state suggests other roles for at least one of these residues. One possibility is that D61 has an alternative salt bridge partner, R26 (adjacent to K27), with hydrogen-bonding to the backbone of position 27. This idea that a D61-R26 salt bridge can substitute for that of D61-K27 is supported by our observation that, while K27 is quite tolerant, D61 is generally sensitive to variation. Although the average distance between D61 and R26 for the WT protein is ~4 Å greater than that of D61 and K27 (3.5 Å), the electrostatic interaction between D61 and R26 side chains is noticeable. To further explore this hypothesis, we simulated the effects of an R26P variant which should disrupt a D61-R26 salt bridge, and found that it greatly increased the average D61-K27 distance in the R26P variant (10.2 Å ± 3.1 Å) compared to that of WT (3.5 ± 1.8 Å). A D61 role in active site loop conformation may explain its sensitivity to variation, while the presence of R26 can mitigate the impact of variation at K27 by providing an alternative salt bridge partner for D61.

**Figure 3.**
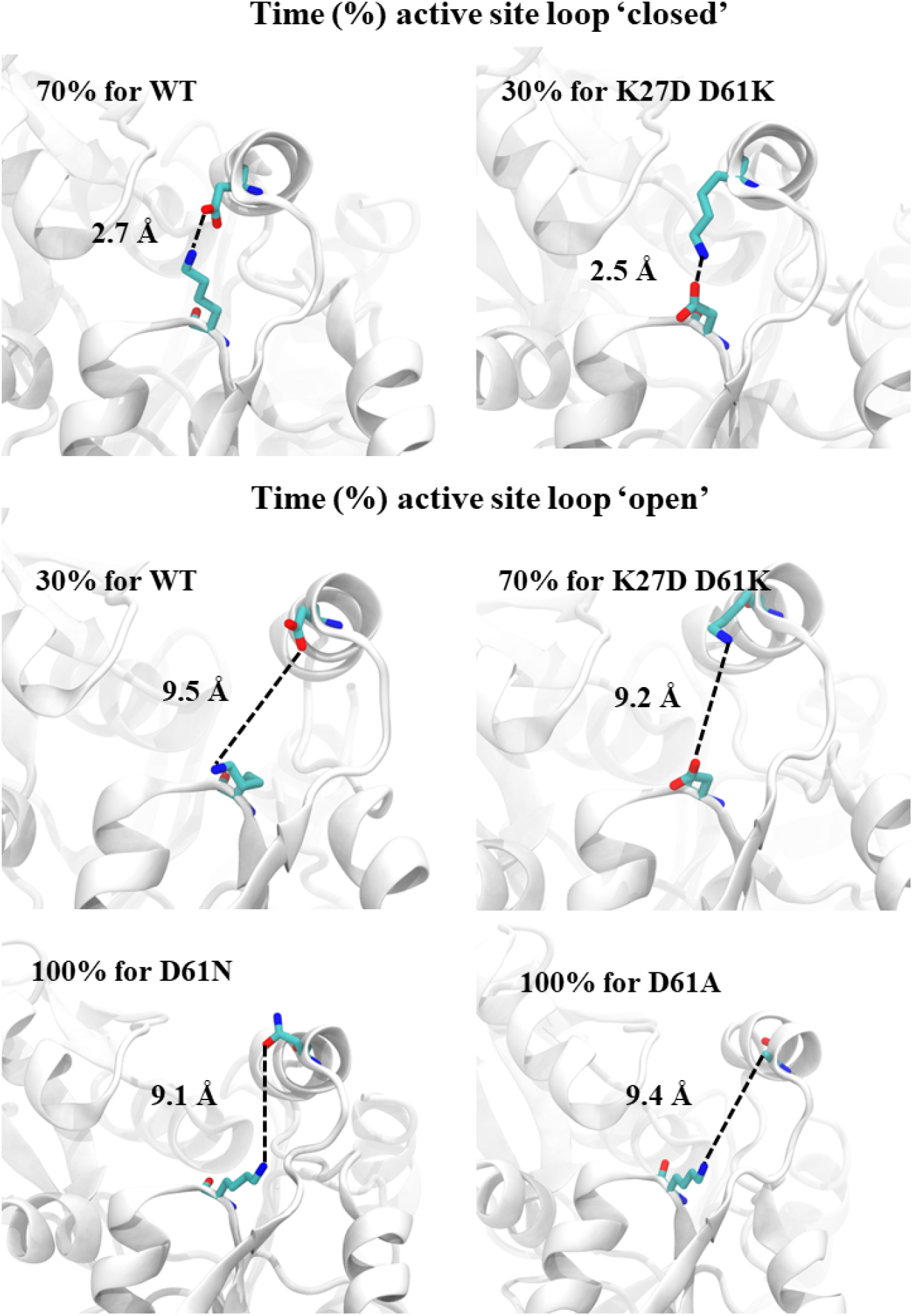
The “closed” and “open” conformations of the active site loop in WT HMBS and three HMBS variants – D61N, D61A, and the double mutant D61K K27D. The average distance (Å) between protein position 27 and 61 is shown, along with time (expressed as a percentage) in each conformation.

Our second hypothesis was that active site loop residue G60, which was striking in its intolerance to mutation, is important due to the backbone flexibility it provides. To explore this hypothesis, we simulated the impact of a G60P substitution that would be expected to constrain backbone motion. These simulations showed that G60P inhibits the D61-K27 salt bridge, likely via the induced rigidity of the loop that hinders the orientation of D61 needed for the K27 salt bridge (Figure S7). Furthermore, a D65-K27 salt bridge was observed for 91% of simulation time for the G60P mutant (Figure S6), which serves to constrain the active site loop to the “open” state (Table S6).

Taken together, the MD simulation results suggest that K27, G60 and D61 each play an important role in the flexibility of the active site loop, thereby impacting the uptake of PBG subunits and/or exit of the processed substrate. Distances between D61 and other residues observed in simulation are summarized in Table S6.

### Comparing measured functional impacts with predicted protein stability effects

It can be instructive to compare variant impacts on stability (ΔΔG) as opposed to overall functionality. For example, it has been shown that mutations having an impact on function but not on stability are more likely to be active site residues ^41,42^. We therefore performed moving window analyses enabling comparison of map scores and predicted ΔΔG values at different protein positions. As expected, these profiles were correlated with one another: Protein positions with predicted-destabilizing substitutions tended to have deleterious functional impact scores (Figure 4A and Figure S8B). Surface residues (here defined by solvent-accessible surface area (ASA) < 20%) tended to have neither predicted stability effects nor strong functional impacts. In contrast, residues with high functional impact that were not predicted to have strong stability effects were restricted to active site residues involved in polypyrrole elongation (R26, T145, S147, R149 and R150), and also to positions 316-319. Positions 316-319, located outside the active site at the interface of domain 3 and domain 1, are packed closely and exhibit low mobility in our MD simulations (Figure S3). An impact of changes in residues 316-319 to function but not stability, coupled with their involvement in inter-domain residue-residue interactions (including T109-T319, I113-I318, and I110-I318), suggests that they help restrain structural fluctuations that would otherwise reduce enzymatic activity.

**Figure 4.**
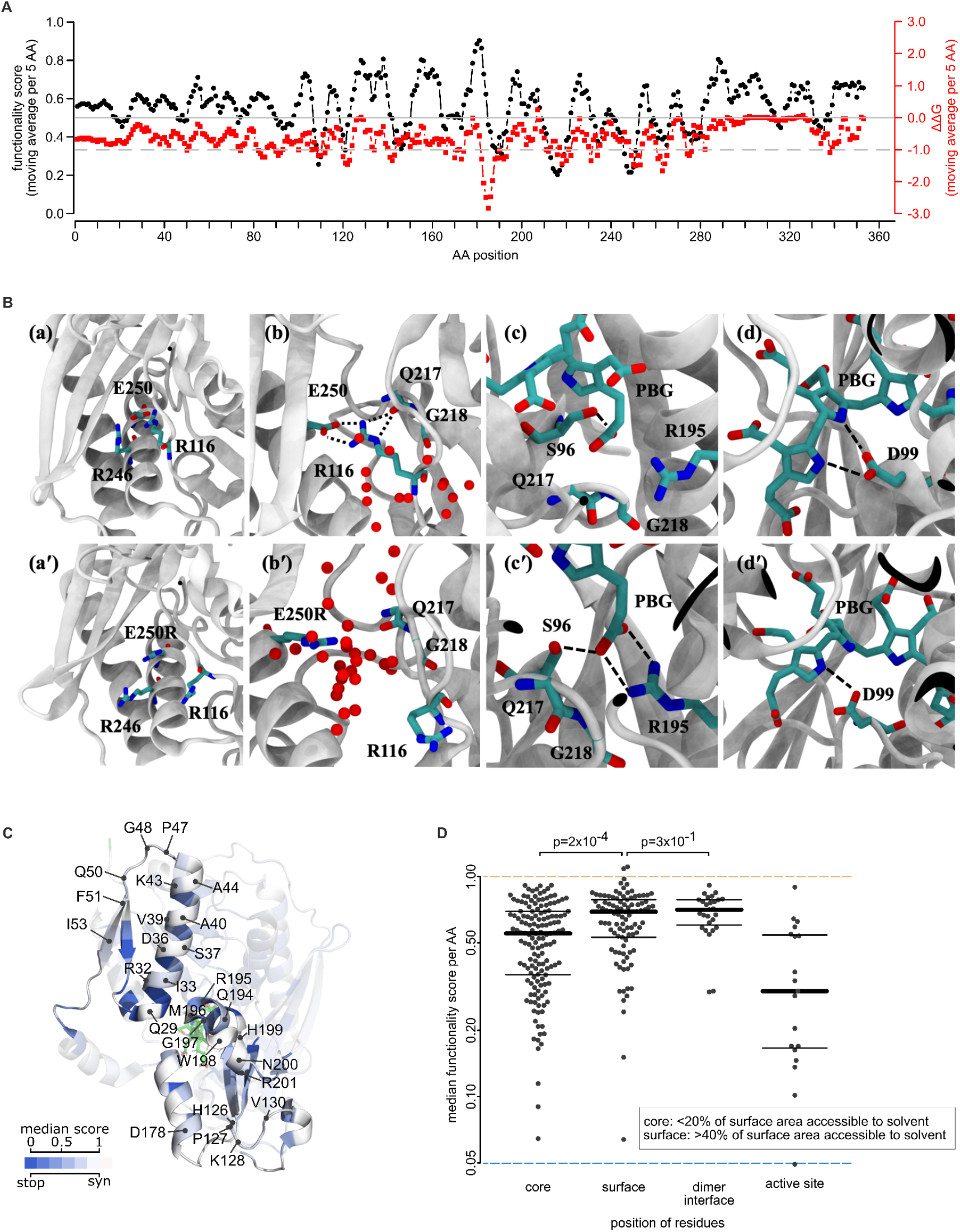
Modeling the effects of HMBS missense variants on protein stability and structure (A) Comparison of functional impact scores (black) and predicted free energy change (ΔΔG; red) values of HMBS missense variants. Plotted values are averages within windows of five amino acid (AA) positions. (B) WT (top) and E250R variant (bottom) comparison. The E250R substitution repels R99 and opens a channel, which is exposed to solvent. PBG-G218 interaction is lost and replaced by a salt bridge between PBG and R195, which in turn disrupts D99-pyrrol interactions. For clarity, hydrogen atoms are not shown. Water molecules within 7 Å of E250 (or R250) and Q217 with larger than 50% occupancy are shown. (C) Structural model of HMBS; colored according to the median functionality score of substitutions at each position, along with a wireframe model of the tetrapyrrole (green), and noting residues located at the dimer interface. (D) Median functionality scores of variants at amino acid positions which were: (1) below 20% accessible surface area (ASA); (2) above 40% ASA; (3) at the dimerization interface with a threshold ΔASA of 1.0; and (4) active site residues required for polypyrrole assembly. P values were calculated by Mann-Whitney U test. Thick and thin bars correspond to median, upper, and lower quartiles, respectively. Light green and dark blue dashed lines correspond to WT-like and null-like scores, respectively.

To further explore the importance of inter-domain residue interactions on structural dynamics, we used MD simulations to examine the variant T319Q, which our map found to be damaging. Root mean square fluctuations of the Cɑ atoms of all amino acids throughout the simulation time (RMSF) were examined, both in the context of the wild-type and T319Q structures. As expected, RMSF values tended to be higher for T319Q in all three domains, with a significant increase for residues in the active-site loop and the insertion regions (Figure S4). Interestingly, the T319Q structure exhibited lower RMSF values within the cofactor-binding loop. Because the cofactor-binding loop (residues in positions 257–263) normally moves during catalysis to accommodate shifting of the initial DPM substrate as the polypyrrole elongates ^35,37^, we conclude that the T319Q variant hinders cofactor-binding loop movement that would otherwise accommodate polypyrrole chain elongation.

We next assessed whether salt bridges outside the active site may be important for protein folding and/or stabilization. Mutations predicted to disrupt the formation of each of several salt bridges – R116-E250, D121-R149, D121-R150, and R225-D91 – appeared damaging in our maps (Figure S4). Because both the R116 and E250 positions are associated with pathogenic variation (R116W, R116Q and E250K), we focused our attention on the corresponding R116-E250 salt bridge. In addition to this salt bridge, hydrogen-bonding of R116 to the Q217 backbone keeps Q217 in place and aligns the G218 side chain to effectively interact with a PBG carbonyl group; Table S3; and Figure 4B). To further evaluate the importance of the R116-E250 salt bridge, we performed MD simulations for variant E250R. Here we observed the E250R mutation to eliminate the E250-R116 salt bridge, with the R250 mutant residue opening a water-filled channel between the helix (224-240) and the loop (198-202) (Figure 4B) and repelling R116. In the E250R simulation, we also observed water molecules hydrogen bonded to both R116 and Q217, thus replacing the R116-Q217 interaction and eliminating the G218-PBG interaction. We attribute this effect to increased active site loop flexibility caused by E250R (Figure 4B). The simulation also suggested that loss of the E250-R116 salt bridge drives a new salt bridge between PBG and R195, in turn altering the position of PBG and weakening hydrogen bonding between D99 and pyrrole groups (Figure 4B). Thus, the combination of our map and simulation analysis suggests that E250R not only disrupts R116-E250 salt bridge and other local structure around position 250, but also causes a cascade of changes in backbone and side chain conformations, altering residue interactions in the active site to impact both protein stability and position of the tetrapyrrole substrate.

### Evaluating HMBS missense impacts in the context of the homodimeric HMBS structure

Examining whether protein core residues are more sensitive to mutation than surface residues, we found buried residues (those with <20% ASA), to have a lower median score than surface residues (>40% ASA; Δmedian=0.13; p=0.0002 Mann-Whitney U test; Figure 4D). Interestingly, interface residues exhibited scores that were similar to other surface residues (Δmedian=0.04; p=0.3; Mann-Whitney U test; Figure 4D).

HMBS is reported to function as two monomers in an asymmetric unit with a weak dimer interface ^39,43^, however the relationship between dimerization and the monomeric enzyme’s stability and activity remains unclear ^44^. To visualize missense functional impact scores in the context of the homodimeric HMBS structure, we colored each residue in the structure according to the median fitness of substitutions at that residue. Residues at the dimer interface—especially those at the center (W198 and H199) — appeared highly tolerant to variation (Figure 4C). One exception was residue F77 for which all variants scored as damaging. The sensitivity of F77 to variation (Figure 2E) can be explained by the interaction between residues F77 and R26, which helps maintain the active-site loop in a closed conformation through the first stage of chain elongation (Figure S7) ^35,45^. In fact, our MD simulation data indicate that eliminating the cation-π interaction in the F77A variant increased the D61-R26 and D61-K27 distances (both by ~ 3 Å) and exposed the active-site, confirming the importance of R26-F77 interaction. Thus, R26 is not only involved in the salt bridge mentioned above, but also provides structural integrity via interaction with F77. Given that the only residue at the dimer interface exhibiting substantial sensitivity to variation can be explained without dimerization, our results support a previous hypothesis that HMBS dimerization is not critical for its stability ^44^.

### Limited correlation between functional impact scores and disease severity

For genetic diseases broadly, penetrance and expressivity can depend on the extent to which a variant impacts the function of the causal gene ^25^, or have other explanations (e.g., genetic variation outside the causal gene or environmental differences). For AIP, the low penetrance and variable expressivity of AIP even amongst individuals carrying a shared HMBS missense variant (e.g. p.R167, p.R173, p.R225 and p.R325)^3^ suggests that extragenic variation and environmental differences are a more likely explanation. Nonetheless, we sought to examine the correlation between our functional impact scores and the age of onset or severity of AIP. Although there is no accepted framework for classifying AIP severity, one study has categorized patients according to AIP severity, reporting that phenotypic severity correlated with a variant’s position relative to the active site ^46^. We adopted these assignments for our analysis of AIP severity but are compelled to note that no objective criteria for these assignments were described. The observation that our functional impact scores correlated poorly with age of disease onset and also with previously-classified disease severity (Figures S8C and S8D), argues against strong dependence of AIP severity on quantitative differences in variant functional impact.

### HMBS functional impact scores identify pathogenic variants

Beyond understanding sequence-structure-function relationships, variant effect maps offer functional evidence in support of clinical variant interpretation. To assess the correspondence of our HMBS maps with pathogenicity, we first assembled a positive reference set of 53 likely pathogenic or pathogenic missense variants and a negative reference set of 13 missense variants that were benign, likely benign, or ‘proxy benign’ (see Methods). We then evaluated the tradeoff between precision (fraction of variants below a given threshold score that have been annotated as either pathogenic or likely pathogenic) and recall (fraction of all variants annotated as pathogenic or likely pathogenic that scored below the threshold). Because precision depends on the proportions of pathogenic and benign variants in the reference set (which may not accurately reflect the prior probability that any given clinical variant is pathogenic or benign), we transformed each empirical precision vs recall curve to the corresponding ‘balanced precision’ vs recall curve where the prior is balanced (with a 50% probability of pathogenicity; Figure 5A). We also measured the recall at a stringent balanced precision threshold of 90% (R90BP)^47^. Both erythroid-specific and ubiquitous maps captured 86% of scored pathogenic variants at high stringency (i.e. R90BP was 86%), while the combined map had an R90BP of 88% (Figure 5A). Interestingly, functional impact scores of annotated pathogenic and benign variants were generally well separated from one another except within a small region (residues 160-215) in the HMBS map where known pathogenic variants exhibited limited functional impact (Figure S9A). Agreement with the map seen for individual yeast complementation assays of variants in this region suggest that the map accurately reflects the yeast-based assay, and is consistent with the hypothesis discussed above – that growth of the polypyrrole chain beyond tetrapyrrole may not be required to rescue the *HEM3-ts* phenotype (Figure S1). In summary, our HMBS variant effect map corresponds well with pathogenicity, with the important caveat that the results should not be taken to infer benignity for residues in positions 160 to 215. After excluding these positions, the combined map had an R90BP performance of 93% (Figure S9B).

**Figure 5.**
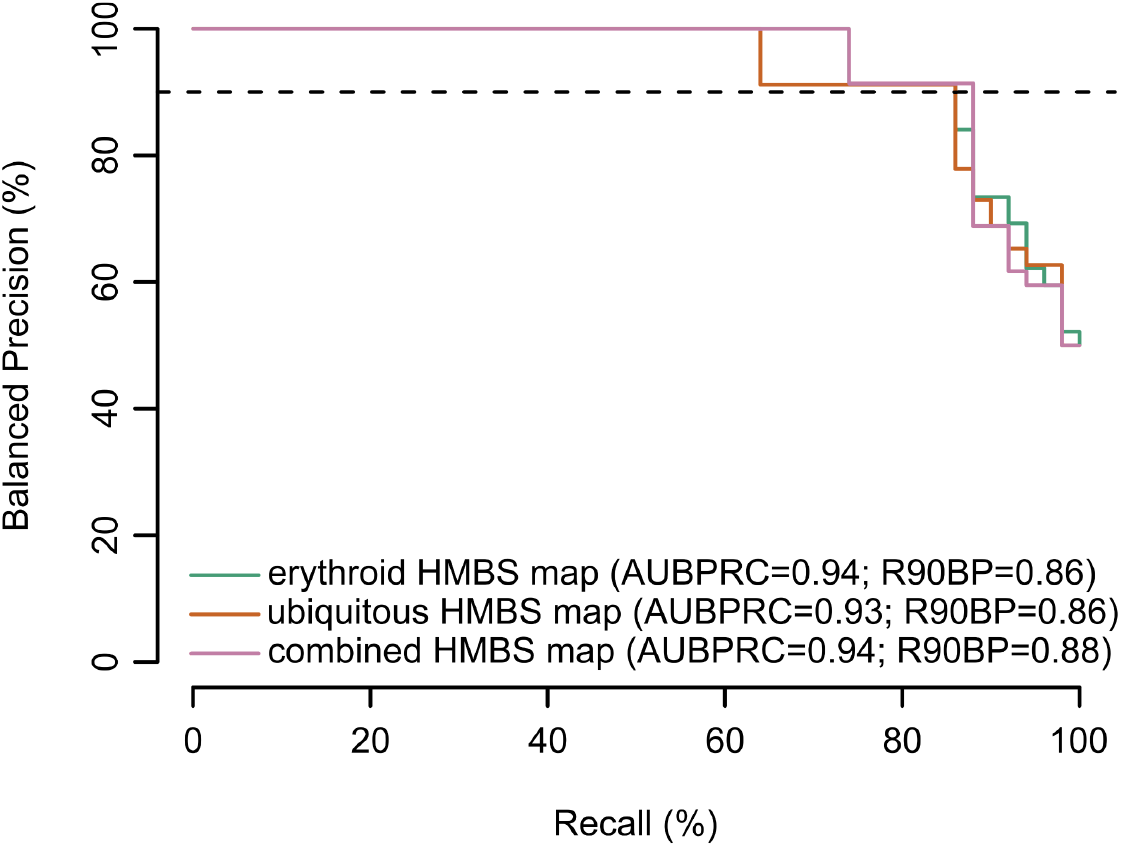
Performance of variant effect maps in distinguishing pathogenic from benign reference variants. Evaluation of precision (fraction of variants scoring below each threshold functionality score that are in the positive reference set containing pathogenic variants) vs recall (fraction of positive reference variants with functionality scores below threshold). Here, precision has been balanced to reflect performance in a balanced test setting where positive and negative sets contain the same number of variants. Balanced precision-recall curves are shown for erythroid-specific (green line), ubiquitous (orange) and combined maps (pink). Performance is also described in terms of area under the balanced precision vs recall curve (AUBPRC) and recall at a balanced precision of 90% (R90BP).

Experimental and computational sources of evidence about variant pathogenicity are complementary, in the sense that current ACMG/AMP guidelines for variant interpretation consider these evidence types to be independent of one another. However, it is interesting to compare the performance of computational predictors with our variant effect map (after again excluding region 160-215). In keeping with previous observations that found experimental functional assays to offer higher sensitivity than most computational predictors ^23–25,27^, our ubiquitous and combined maps each exhibited better performance than PROVEAN ^48^ (5% R90BP) and SIFT ^49^ (85% R90BP). A more recently described computational predictor, VARITY_R_LOO^50^, performed on par with our experimental map (95% R90BP) (Figure S9B).

Finally, to enhance the accuracy and sensitivity of HMBS missense variant interpretation, we re-stated each variant’s fitness score in terms of a likelihood ratio of pathogenicity (LLRp). The LLRp value estimates the likelihood of obtaining the observed score in the positive reference set relative to the corresponding likelihood in the negative reference set, which can also be translated into discrete evidence strengths in the ACMG/AMP framework ^51^. This analysis suggests that LLRp scores above 0.64 should be treated as moderate evidence of pathogenicity, LLRp scores between 0.64 and 0.32 as supporting evidence of pathogenicity, and LLRp scores below −0.32 as supporting evidence of benignity (Figure S9C).

Based on this calibration of LLRp scores to ACMG/AMP evidence strength categories, our map provides new functional evidence for variant classification for 54 (83%) of the 65 HMBS missense VUS reported in ClinVar. Of these 54 variants, 44 (81%) received scores corresponding to supporting evidence of benignity (Table S8), while 1 (2%), 2 (4%) and 7 (13%) received LLRp scores corresponding to supporting, moderate or strong evidence of pathogenicity, respectively.

## Discussion

By combining codon-randomizing mutagenesis and large-scale multiplexed functional assays, we proactively assessed the functional impact of missense variants for the human *HMBS* gene for a large fraction of missense variants in both the erythroid-specific and ubiquitous HMBS isoforms.

Our functional impact scores agreed closely with known sequence-structure-function relationships, with some exceptions within the active site and active site loop, and at the hinges controlling flexibility and dynamics of HMBS domains. For example, at positions important for DPM formation, binding, or chain elongation, we found missense variation to be damaging as expected. However, the maps showed some variants at positions involved in chain elongation (e.g. residues S165, N169, R255, and R355) to be surprisingly tolerated. These were generally at positions important for stabilizing the negatively charged polypyrrole, which is consistent with previous reports that water molecules can compensate for loss of charge stabilization ^35^. Some variants near the active site or within the active-site loop were also tolerated, and our results suggest that a critical determinant of mutational tolerance at these positions is the relative distance from the active site.

Based on our map scores and MD simulations, residue interaction at the interface of domain 1 and 3 modulates the dynamic behavior of HMBS during catalysis. Furthermore, our results demonstrate that a salt bridge network (R26, D61, T58 and K27) and cation-π interaction between F77 (part of the active-site loop) and R26 ensures the active-site loop remains in a “closed” conformation during cofactor assembly and the first addition of PBG^35^. When examining the general contribution of salt bridges (G250-R116, D121-R150, R225-D91, and R149-D121) towards overall protein stability, the maps showed variation at such positions to be damaging, resulting in large-scale structural rearrangements in the backbone conformation and side chains’ geometrical orientation that would impair catalytic activity.

The HMBS variant effect maps revealed other patterns of mutational tolerance. For example, variants at the dimerization interface were generally tolerated, suggesting HMBS does not require dimerization for its function. Moreover, variants at non-dimerization-interface surface residues tended to be strongly damaging, perhaps due to introduction of hydrophobic residues that favour aberrant folding. Both of these findings are consistent with suggestions by Chen and colleagues based on functional assays of eleven missense variants ^44^.

One caveat of our map is that our functional complementation assays are based on the expression of mature cDNA, so that any impact of variants on splicing will necessarily have been missed. Because introns are important for nonsense-mediated decay in mammalian cells, it is possible that nonsense variants seen as tolerated in a yeast-expressed cDNA would be damaging in the endogenous human context. Nevertheless, nearly all nonsense variants were found in our yeast cDNA assay to be highly damaging (Figure 1C).

An important limitation of our maps is that it measures total activity, not specific activity. Thus, we cannot distinguish functional impacts on protein abundance, e.g. due to misfolding that accelerates protein degradation, from impacts on specific activity that do not affect abundance. Our observation that the correlation between functional impact scores and *in vitro* measurements of specific activity was significant but moderate (Spearman’s ϱ *=* 0.54, p= 4 ×10^−9^, Figure S8A), together with a similar observation for correlation of functional impact scores with predicted impacts on thermodynamic stability (Spearman’s ϱ*=* 0.52, p= 2 ×10^−16^, Figure S8B), suggests that both stability and specific-activity effects are at play. For the purpose of understanding sequence-structure-function relationships, it would be interesting to determine whether impacts on total activity arose via impact on steady-state protein level (as could be measured for example by the VAMP-Seq method ^52^) or via impact on specific activity. However, it is not clear that knowing whether total HMBS activity is lost due to reduced expression levels, as opposed to specific activity, would have clinical value.

Another important caveat is that our measurements were necessarily subject to both systematic and random error. We used previously-described methods ^27^ to estimate random error for each experimental score, reflecting the estimated magnitude of random experimental error. Systematic errors could have arisen from many sources. For example, any errors made in recalibrating the score range for each region (see Methods) could cause scores from one region to be systematically higher or lower than another. Some systematic errors may also have arisen from the nature of our assay. For example, we observed some residues implicated in the addition of the final two PBG monomers to be surprisingly well-tolerated in our yeast-based assay, which could be explained if residual activity of the yeast *HEM3* ts mutant can extend (but not generate) HMB. Because many pathogenic variants found tolerated by our assay fell within positions 160-215, which contains many residues important for coupling of the final two PBG monomers, we suggest that positions 160-215 in the map be excluded when inferring benignity. However, this map region could remain useful for inferring pathogenicity.

Despite these limitations, we found that our variant effect maps could reliably discriminate pathogenic or likely-pathogenic from benign, likely-benign, or ‘proxy benign’ variants, with the best performance coming from the combined map. When comparing our maps with computational predictors at thresholds providing equally-stringent (90%) precision, the number of pathogenic variants captured by our maps was higher than two widely used computational predictors, PROVEAN and SIFT, and was on par with VARITY. Although the performance of each map was improved further by excluding residues 160-215, the overall conclusions in the comparison with computational methods did not change. This comparison should not be taken to suggest that there must be a competition between computational and experimental evidence. Indeed, treating these evidence sources independently per ACMG/AMP guidelines means that these evidence sources can be synergistic, albeit with greater evidence strength afforded to experimental assays of variant function ^15^.

An important caveat of variant interpretation is that variants determined to be pathogenic (whether via variant effect maps or otherwise) may not cause disease in every patient. This caveat is particularly pronounced for pathogenic HMBS variants which generally have low penetrance. Our study was also limited by the number of publicly available rare missense variants that have been annotated as “likely benign” or “benign” in ClinVar. We used HMBS variants in the gnomAD database as a negative reference set after excluding those reported to be pathogenic or likely pathogenic. While we cannot exclude the possibility that some of these individuals have AIP (despite the rarity of this condition), variants in this negative reference set can at least be expected to be strongly depleted for pathogenicity.

Some variants in the C-terminal region of HMBS appeared to ‘hyper-complement’, i.e. grow faster than the wildtype control in the complementation assay, especially in the ubiquitous isoform map. Hyper-complementation could indicate increased activity in humans, e.g., possibly arising from increased stability or enhanced conformational flexibility and concomitant increase in catalytic activity. However, previous analyses of a missense variant effect map for the protein *UBE2I* (also based on a yeast assay) suggested both that *UBE2I* hyper-complementing variants tend to be deleterious in humans and that they result from changes that are specifically adaptive in the yeast cellular context ^27^. Using a similar quantitative phylogenetic approach, we found the best-performing models for HMBS penalize hypercomplementing variants, indicating that these variants are best treated as deleterious in nature.

Although we observed no significant correlation between functional impact scores and either age of disease onset or disease severity, we do not wish to suggest that this question is closed, absent more data and an accepted objective framework for classifying AIP severity. It would be interesting to investigate whether the functional impact scores are more predictive of age of onset or severity after stratifying by the presence or absence of known triggers ^1^.

The HMBS variant effects maps we provide could have immediate value in several clinical scenarios. First and most importantly, where a patient has been diagnosed with AIP based on clinical and biochemical data but has an *HMBS* missense variant that would otherwise be classified as VUS, a more definitive classification of the variant could enable cascade screening to identify family members with latent AIP. Second, where a patient has highly elevated ALA and PBG, but does not have access to a laboratory capable of ruling out the two other acute porphyrias, identification of a definitively classified HMBS variant can establish the correct diagnosis. Third, where a patient has a clinical history consistent with AIP, but timely measurements of porphyrin precursors in urine or plasma were not obtained or were inconclusive. Fourth, where an *HMBS* missense variant is revealed, e.g., through direct-to-consumer genome sequencing, and the client wishes to know if they should be vigilant for symptoms of AIP, or avoid its known triggers. In each of these scenarios, a resulting diagnosis of AIP or latent AIP could have the cascading benefit of helping to identify additional cases of AIP or latent AIP in at-risk relatives, thereby increasing the number of patients for whom vigilance, prevention or therapy is supported.

In addition to providing a resource for the understanding of HMBS variation, this study also provides proof of principle for broader application of variant effect mapping to other genes associated with acute hepatic porphyria.

In conclusion, we strengthen the evidence that systematic proactive experimental evaluation of missense variant effects on human enzymes can reveal sequence-structure-function relationships and yield clinically relevant insights, with potential to guide personalized clinical decisions.

### Material and Methods Strains and plasmids

The *Saccharomyces cerevisiae* strain with which we assayed *HMBS* variant libraries (*MATα ts-hem3::KanR his3Δ1 leu2Δ0 ura3Δ0*) was kindly provided by Drs. Guihong Tan, Charles Boone and Brenda Andrews. For yeast expression, we used the Gateway-compatible destination vector pHYC-Dest2 (CEN/ARS-based, *ADH1*promoter, and *LEU2* marker) ^23^. The *HMBS* open reading frame (ORF) clone (Ensembl:, GenBank:. corresponding to UniprotKB accession P08397) was obtained from the Human ORFeome v8.1 library ^53^.

Wild-type reference or mutated disease-associated versions of the HMBS ORFs were transferred into pHYCDest by Gateway LR reactions. After confirmation of ORF identity and expected mutations by Sanger sequencing, the expression clones were transformed into the appropriate yeast ts strain in parallel with an ‘empty’ expression vector control (bearing the counterselectable ccdB marker controlled by a bacterial promoter).

### HMBS assay validation

For yeast ts mutants transformed with vectors expressing HMBS cDNAs, cells were grown to saturation at 30°C. Each culture was then adjusted to an OD_600_ of 1.0 and serially diluted by factors of 5^−1^, 5^−2^, 5^−3^, 5^−4^, and 5^−5^. These cultures (5 μL of each) were then spotted on SC−LEU plates as appropriate to maintain the plasmid and incubated at either 30°C or 35°C for 48 hrs. After imaging, results were interpreted by comparing the growth difference between the yeast strains expressing human genes and the corresponding empty vector control (Figure S1). Two independent cultures were grown and assayed.

### Constructing codon randomized HMBS variant libraries

Libraries of *HMBS* variants were constructed using an oligo-directed codon-randomizing mutagenesis (Precision Oligo-Pool based Code Alteration or POPCode) method ^27^. Mutagenesis was targeted to each of two equal-length regions, so that two full-length mutagenized libraries were generated for each HMBS isoform. Briefly, we designed a ~35nt oligonucleotide corresponding to each codon along the entire length of the *HMBS* ORF, each with a central NNK degeneracy targeting that codon for randomization, using the POPcode oligo suite tool ^27^. From the 360 oligos synthesized, oligos for each region were combined to produce two regional pools and phosphorylated. For each isoform, uracilated full-length WT *HMBS* was used as the template and 2 separate annealing reactions (using Kapa HiFi Uracil+ DNA polymerase (KapaBiosystems) and a dNTP/dUTP mix were set up with either oligo pool. After annealing phosphorylated oligos, KAPA HiFi Uracil+ DNA polymerase (Kapa Biosystems) was used to fill in gaps, and Taq DNA ligase (NEB) applied to seal the nicks. Treatment with Uracil-DNA-glycosylase (UDG) degraded the original uracilated template, and the newly synthesized mutagenized strand was amplified using primers containing attB sites. The mutagenized attB-PCR products were then transferred *en masse* into the entry vector pDONR223 using Gateway BP reactions. These Gateway-entry clone libraries were transferred to a pHYC-Dest2 expression vector via *en masse* Gateway LR reactions to enable yeast expression. Both Gateway-entry and -host libraries were transformed into NEB5α *E. coli* cells (NEB) and selected on LB agar plates containing spectinomycin and ampicillin, respectively. Next, host libraries were transformed into the *S. cerevisiae hem3-ts* mutant strain using the EZ Kit Yeast Transformation kit (Zymo Research). To retain maximal library complexity, plasmids were purified from >350,000 clones at each transfer step, and ~ 1,000,000 yeast transformants were pooled to form the host library.

### Multiplexed assay for HMBS variant function

The underlying yeast-based functional complementation assay of HMBS was previously established ^26^. High-throughput complementation screening was carried out as follows: Yeast transformants were grown at 30°C in synthetic complete (SC) media, using glucose as carbon source, without leucine (USBiological) to ensure plasmid retention (the non-selective condition). For each region, two plasmid pools were prepared from 10 ODU of cells (defined here as the number of yeast cells found in 10mls of a 1 OD_600_ culture, which is typically ~10^8^ cells) and used as templates for the downstream tiling PCR. To allow multiplexed functional analysis of HMBS variants despite cell-non-autonomous effects, two replicates of ~800,000 cells from each of the regional transformant pools were washed three times to remove exogenous porphyrins and heme, plated on selective SC(-Leu) media and grown at restrictive temperature (selective condition; 35°C) for 48 hrs. After pooling the colonies of each replicate, plasmids were extracted from 10 ODU of cells and used as template for downstream tiling PCR. In parallel, the *hem3-ts* mutant strain was transformed with the WT ORF, grown alongside the regional pools, and the plasmid extracted from two of 10 ODU of cells for each condition for use as a control.

### Scoring functional impact of variants using TileSeq

Measuring variant effects using a pooled *en masse* selection strategy was done according to the previously described TileSeq approach ^27^. Briefly, for each of the plasmid libraries from non-selective and selective pools, short template amplicons (~150bp) that tile the ORF (within the context of each regional pool) were amplified with primers carrying a binding site for Illumina sequencing adaptors. Both regional pools consisted of five tiles. In the second-round PCR, Illumina sequencing adaptors with index tags were added to the first-step amplicons. Paired-end sequencing was then performed on the tiled regions across the ORF, thus dramatically reducing base-calling error and enabling accurate detection of very low (parts-per-million) variant frequencies. Separate sequencing runs performed for each isoform using an Illumina NextSeq 500 via a NextSeq 500/550 High Output Kit v2 achieved an average sequencing depth of ~2,000,000 paired-end reads per tile.

Sequencing reads were demultiplexed with bcl2fastq v2.17 (Illumina) and processed using the previously described TileSeq strategy ^27^. In brief, variant frequencies in each condition were determined with custom Python scripts (https://github.com/RyogaLi/tileseq_mutcount). These scripts incorporated Bowtie2 as part of the pipeline to align the sequence of each read pair to the reference template. Following alignment, mutations were called when the resulting pairing for each tile agreed on its presence. Read counts were then normalized based on sequencing depth to yield variant frequency data for each condition and replicate.

Processing of raw read count data was carried out using the tileseqMave R package (https://github.com/jweile/tileseqMave)^25,27^. Briefly, to account for PCR and sequencing error and the possibility of a bottleneck effect from non-selective pool sampling, variants having selective or non-selective frequencies below a level of three standard deviations above the corresponding wildtype frequencies were filtered out. Additionally, the effect of sequencing errors and amplification biases in the libraries generated were minimized by subtracting wildtype from variant frequencies in the non-selective and selective pools. Next, an enrichment ratio (Φ) was calculated for each variant based on the adjusted frequencies. A functional impact score (FS_MUT_) for each variant was calculated as ln(ΦMUT/ΦSTOP)/ln(ΦSYN/ΦSTOP), where ΦMUT is the enrichment ratio calculated for each variant, ΦSTOP is the median enrichment ratio of all nonsense variants, and ΦSYN is the median enrichment ratio of all synonymous variants. Additionally, truncations occurring close to the C-terminus are less likely to be of functional significance, so nonsense variants were excluded from the last 14 amino acids of each HMBS ORF for ΦSTOP. Functional impact scores of each isoform were first rescaled for each region separately, such that FS_MUT_ = 0 when ΦMUT = ΦSTOP and FS_MUT_= 1 when ΦMUT = ΦSYN. However, differences in the stringency of selection for the two isoforms introduced non-linear changes in scale that differ between isoform maps. Scores for the erythroid isoform were therefore rescaled to minimize the average Euclidean distance between scores in the two maps. Filtering further for variants with high quality measurements, we removed variants which had standard error >0.3 or frequency below <0.005% in the corresponding non-select library.

A “delta score” for each variant was calculated as the difference between isoform functional impact scores. Manual inspection revealed no significant differences (Figure S10). Finally, for the combined variant effect map, a simple weighted average was calculated for each variant passing high quality filtering using the equation (FS_1_/σ^2^_1_ +FS_2_/σ^2^_2_)/(1/σ^2^_1_+1/σ^2^_2_). A single combined estimate of measurement errors for scores in the combined map was derived by σ^2^ = 1/(1/(σ^2^_1_+σ^2^_2_)).

### Phylogenetic comparison of different models for hyper-complementation

To assess whether variants exhibiting hyper-complementing behavior in the yeast complementation assay are likely to be advantageous, deleterious or neutral in humans, we performed phylogenetic analysis as described previously ^27,54^. Briefly, we first normalized each variant’s score relative to the wild-type score for that position to avoid penalizing the wild-type variant itself in cases where its score is slightly greater than 1. Additional processing and imputation steps removed low-quality data and imputed likely values for missing data points, respectively. Three sets of scores (*s*) were generated from this dataset, each testing a different way of relating variant score for amino acid *a* at site *r* (*s*_*r,a*_) to the preference for amino acid *a* (defined as 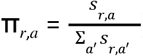) across a set of aligned homologous sequences. In the first (“advantageous”) model, π_*r,a*_ = *s*_*r,a*_. In the second (“neutral”) model, if *s*_*r,a*_ > 1 then *s*_*r,a*_ = 1. In the third (“damaging”) model, if *s*_*r,a*_ >1 we transformed it to a value of 1/*s*_*r,a*_, otherwise π_*r,a*_ = *s*_*r,a*_. For a set of 60 Ensembl homologs having at least 85% sequence identity to the human protein (Table S11), we applied the phydms software package (https://github.com/jbloomlab/phydms) and determined the quality of fit to the phylogeny (as measured by the Akaike information criterion) under each of the three preference models.

### Reference set of disease- and non-disease-associated variants

To assess the ability of variant effect maps to identify pathogenic variants, we used a ‘positive’ set of 53 variants identified as disease-associated collected by European expert centres. As a ‘negative’ set, we used 7 variants identified as non-disease-associated collected by European and American expert centres and 4 variants annotated as either benign or likely benign on ClinVar that had been submitted with review criteria, and which did not have conflicting interpretations. We augmented the negative set with rare variants (MAF < 0.0005) from gnomAD v2.1.1, requiring that they had been observed to be homozygous in at least one individual, and had no pathogenicity annotations. This identified an additional 2 ‘proxy-benign’ variants, yielding a negative set with 13 variants total.

### Evaluating strength of evidence provided by functional impact scores for variant classification

To aid in clinical variant interpretation, we determined a quantitative Bayesian evidence weight for each variant within our maps by translating the scores to log-likelihood ratios of pathogenicity (LLRp). To this end, we fitted normal distributions to the histograms of the scores of damaging and tolerated variants by maximum likelihood parameter estimation in order to obtain estimated probability density functions for pathogenic and benign variants: ρ(S|+) and ρ(S|−), respectively). The pathogenic:benign likelihood ratio for a variant with a given functional impact score, *f*, was calculated as the ratio of the estimated probability density functions evaluated at *f*: *Λ*(ρ(S|+) : ρ(S|−)| *f*) = ρ(*S*| +)(*f*)/ ρ(S|−)(*f*).

To then assess the posterior odds of a variant being pathogenic, O(ρ(S|+) : ρ(S|−) | *f*), we used the Odds form of Bayes’ rule to combine the prior probability of pathogenicity with the likelihood ratio. In other words:

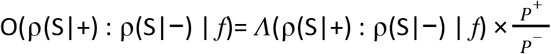

Where *P*^*+*^ is the prior probability that the variant is pathogenic, and *P*^−^ is the prior probability that the variant is tolerated such that *P*^−^ = 1 − *P*^*+*^. A spreadsheet with the pathogenicity log likelihood ratios can be found in Table S8. The code used for these analyses is located at https://github.com/jweile/maveLLR.

We next calibrated the relationship between LLR values and evidence strength categories within the ACMG/AMP framework, using an approach adapted from Tavtigian et al. (2018). Briefly, LLRs for descending evidence levels are modeled to decrease by a factor of 2 at each level starting from a fixed LLR the “Pathogenic Very Strong” (PVSt) level which is chosen such that all ACMG/AMP evidence combination rulesets result in posterior probability of pathogenicity that satisfy the following constraints: >99% for all P rules, >90% for all LP rules, <10% for all LB rules and <1% for all B rules. The most conservative LLR thresholds fulfilling these constraints were found to be 2.54 for PVSt, 1.27 for PSt, 0.63 for PM, 0.31 for PSu, −0.31 for BSu and −1.27 for BSt.

### Computational details (MD Simulation Setup)

To examine variant-specific impacts on the dynamic behavior of HMBS and its movements during catalysis, we next performed MD simulations. Starting structures were obtained by applying necessary changes to the high-resolution X-ray crystal structure (Protein Data Bank (PDB) ID: 5M6R) ^38^. Specifically, to study the apo-enzyme system, the co-factor (DMP) and substrate (PBG units) were removed. Mutations were introduced by PyMol (The PyMOL molecular graphics system, version 1.3, Schrödinger, LLC.), and the protonation states of ionizable amino acids at pH = 7.4 were checked using H++ server ^55^.

For simulations, we used AMBER parm14SB ^56^ parameters and the TIP3P model for solvent ^57^. The nonstandard PBG structure was first energy minimized using B3LYP DFT functional in combination with the 6-31G(d) basis set ^58–60^. Partial atomic charges were obtained from the electrostatic potential using the Gaussian 16 (Revision C.01)^61^. The remaining parameters were obtained from standard parm10 and GAFF2 parameter files, via the ANTECHAMBER module of the Amber18 software suite^62^. All missing heavy atoms and hydrogen atoms were added using the LEaP module of Amber18^62^.

All systems were first neutralized with Na^+^ ions and solvated in an explicit TIP3P water box with at least 1.0 nm from the edge of the enzyme ^57^. Covalent bonds involving hydrogen atoms were constrained using the SHAKE algorithm ^63^ and the particle mesh Ewald^64,65^ algorithm was used for long-range electrostatic interactions. The water molecules and ions were relaxed using 1000 steps of steepest descent and 2000 steps of conjugate gradient minimization, while the protein was constrained using a 500.0 kcal mol^−1^ Å^−2^ force constant. The entire system was then energy minimized using 1000 steps of unrestrained steepest descent, followed by 2500 steps of unrestrained conjugate gradient minimization. Subsequently, the system was heated from 0 to 300 K over 200 ps with restraints on the solute (10.0 kcal mol^−1^ Å^−2^). Each system was then equilibrated for 500 ps under a constant number of particles, volume, and temperature (NVT) condition. All simulations were carried out under a constant number of particles, pressure, and temperature (NPT) using Berendsen thermostat and barostat. Periodic boundary conditions were employed for all MD simulations, which were carried out for 500 ns for apo-enzyme WT, G60P, D61N, D61A, F77A mutants and for the D61K K27D double-mutant. Simulations of 1 μs duration were performed for WT, G346P, and E250R mutant enzymes for their intermediate (ES2) enzyme-substrate conformations, which have two additional substrate pyrrole rings covalently bound to the DPM cofactor.

System stability was measured throughout each simulation with root-mean square deviations of backbone atoms (Table S2). Analyses were carried out using CC-PTRAJ of AMBER 18 ^66^. Hierarchical agglomerative algorithm clustering was completed for the last 500 ns of the trajectories for ES2 systems based on the conformations of DPM-PBG (substrate). The geometrical and interaction analyses were carried out for the structures in the highest occupied cluster. Hydrogen bonds were defined with a 120° angle (donor-hydrogen…acceptor) threshold and a 3.4 Å distance threshold between the donor and acceptor heavy atoms.

### Thermostability calculations

We calculated protein thermostability (ΔΔG) values to relate these to our map scores and thereby identified functionally important HMBS residue positions. Calculations of ΔΔG were carried out using DDGun3D version 0.0.2 (https://github.com/biofold/ddgun), as previously described ^67^. The PDB entry 3ECR for HMBS satisfied the following conditions: a x-ray determined structure with resolution 2.2 Å or better; monomeric structure; and no missing or non-standard residues ^68^. We defined stabilizing amino acid substitutions as those for which ΔΔG ≥ −1, and a destabilizing substitution for ΔΔG <-1.

### Structure modeling and protein positional features

We used the Pymol software to place map scores in the context of a solved HMBS crystal structure (PDB ID: 5M6R). Using the InterfaceResidues.py script (https://pymolwiki.org/index.php/InterfaceResidues), we defined interfacial residues as a change in accessible surface area (ΔASA) greater than 1 Å^2^ between the complex and single-chain structures. We used the FreeSASA program (https://freesasa.github.io/) to calculate relative solvent exposure. A residue position was considered exposed if its solvent accessible surface area value was >40%, intermediate if between 20% and 40%, and buried if <20%.

## Data availability

Custom scripts for all downstream analyses are publicly available https://github.com/wvanlogg/HMBSModel. Functional impact scores for the erythroid, ubiquitous and combined variant effect maps have been deposited on MaveDB^29^ under accession number urn:mavedb:00000108-a. Genotypes and phenotypes for AIP cases drawn from EPNET and literature-curation, respectively, can be found in Table S7.

## Supplemental Figures

**Supplemental Figure S1.**
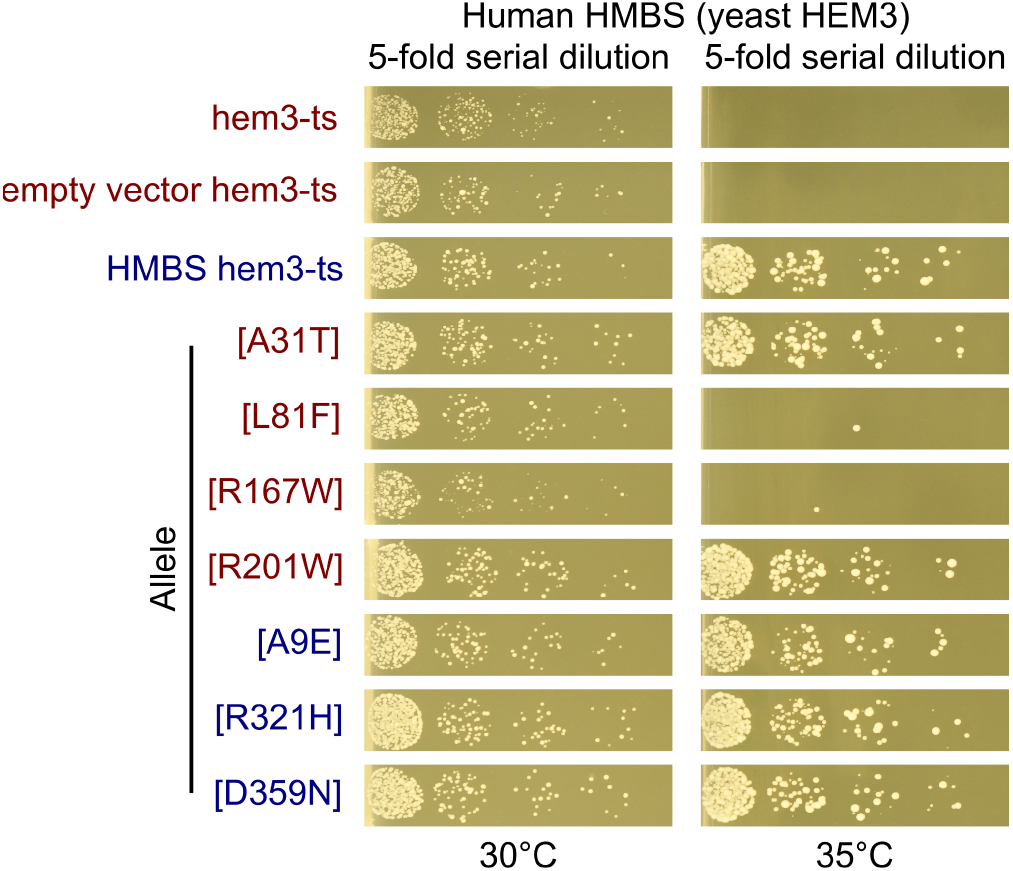
Functional complementation assay results showing whether expression of human HMBS protein variants can rescue growth of a yeast strain bearing a temperature-sensitive mutation in the essential gene *HEM3*. Pathogenic variants A31T, L81F, R167W and R201W and negative controls are indicated in red text, while benign variants A9E, R321H and D359N and a wild-type HMBS control are indicated in blue text. Fivefold serial dilutions of yeast cells were spotted onto plates, with growth evaluation after 48 hours of incubation at either permissive (30°C) or non-permissive (35°C) temperature.

**Supplemental Figure S2.**
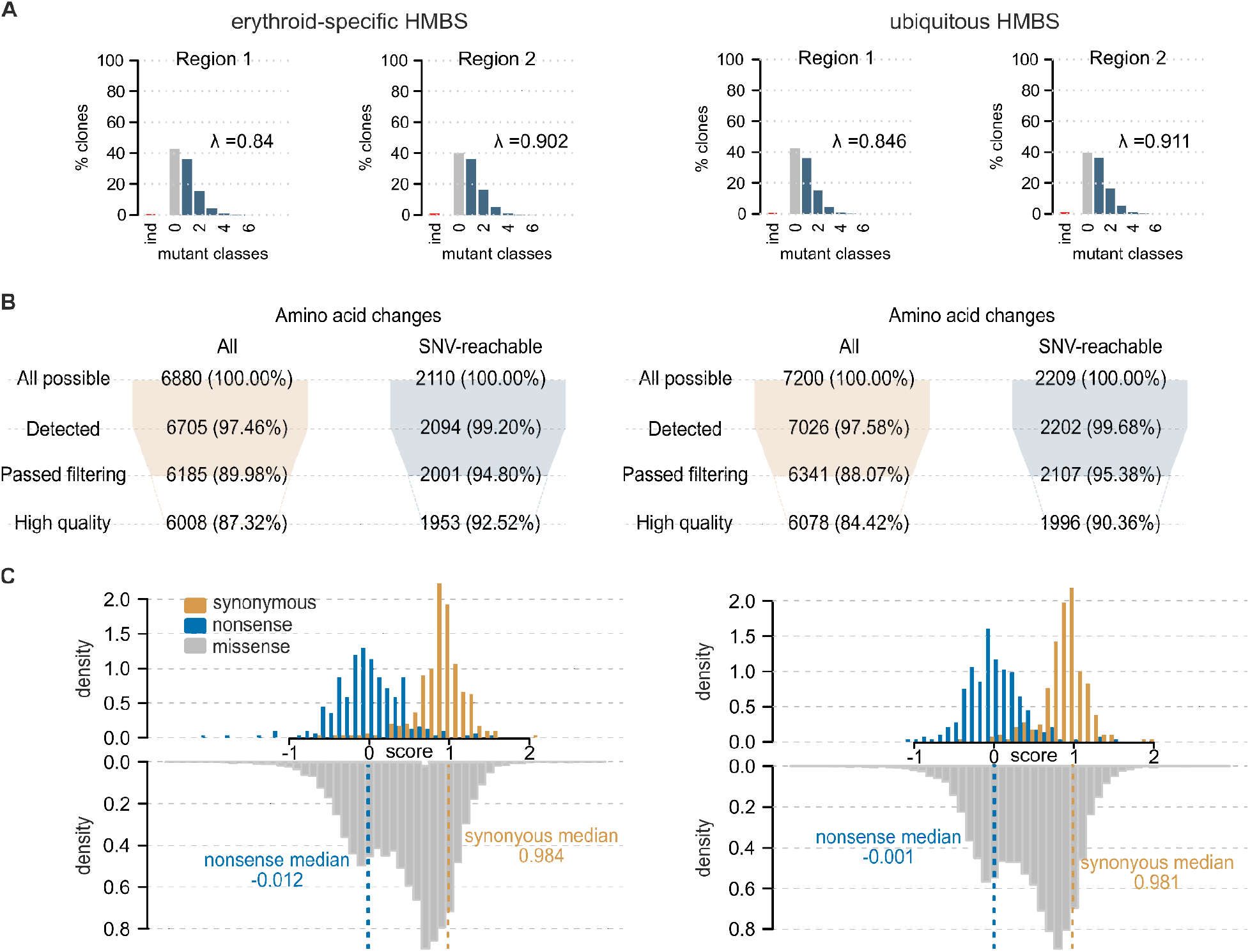
HMBS variant libraries underlying the combined variant effect map (A) Distribution of the number of missense variants in clones from the erythroid-specific (left) and ubiquitous (right) HMBS mutagenized libraries, and the fraction of clones carrying small indels (“ind”). The average number of amino acid changes per clone (λ) is also estimated (see Methods). (B) Fractions of variants detected and passing after each of two levels of quality control are shown for each HMBS isoform. For each isoform, the number of synonymous, nonsense, and missense substitutions (up to 19 possible) are shown across all residue positions, both before (left) and after (right) restricting to substitutions that are possible given a single nucleotide change. The four rows correspond to: 1) theoretically possible substitutions; 2) substitutions detected in the non-selective condition; 3) substitutions above a threshold pre-selection frequency; and 4) substitutions with a sufficiently low regularized standard error. (C) Distributions of measured functional impact scores for nonsense, synonymous, and missense variants for the erythroid-specific (left) and ubiquitous (right) HMBS isoforms.

**Supplemental Figure S3.**
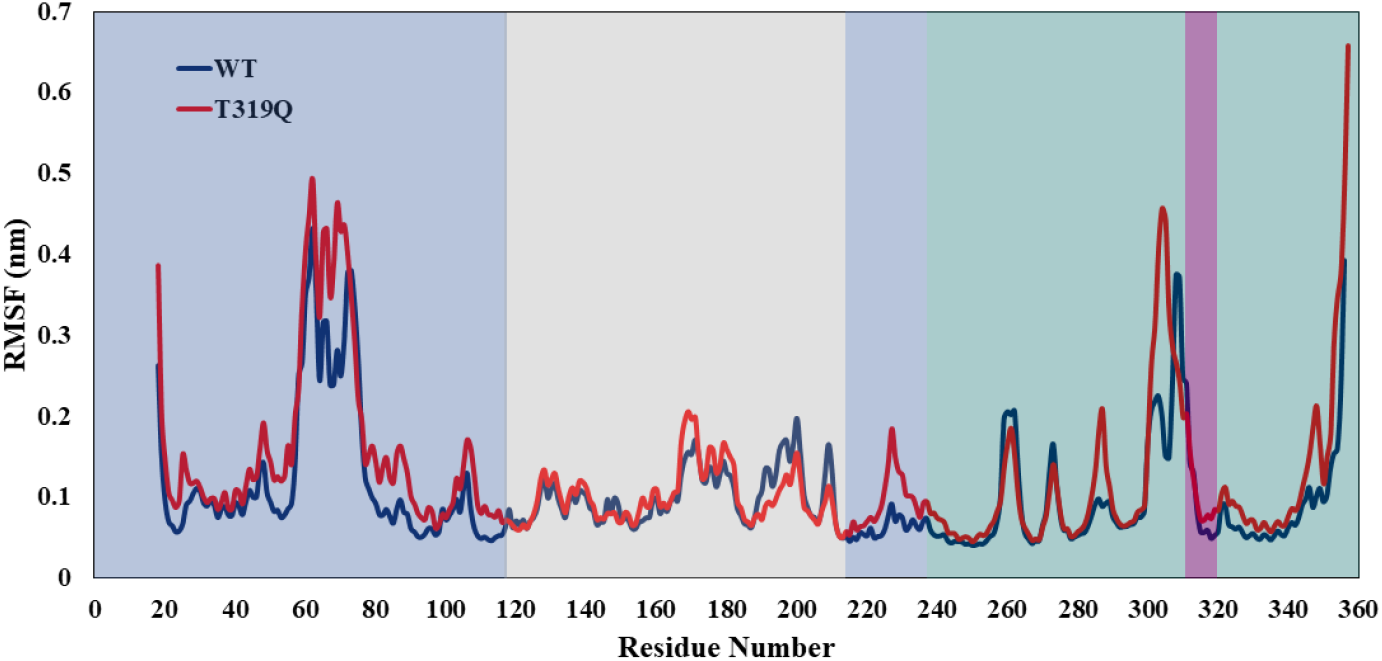
Root-mean-square fluctuation (RMSF) of Cɑ atoms of WT HMBS and the T319Q variant. RMSF reveals the average deviation of atoms throughout the simulation with respect to the initial structure. Domains 1, 2, and 3 are highlighted in light blue, gray, and green, respectively. The 316-319 positions at the interface of domains 1 and 3 are highlighted in purple.

**Supplemental Figure S4.**
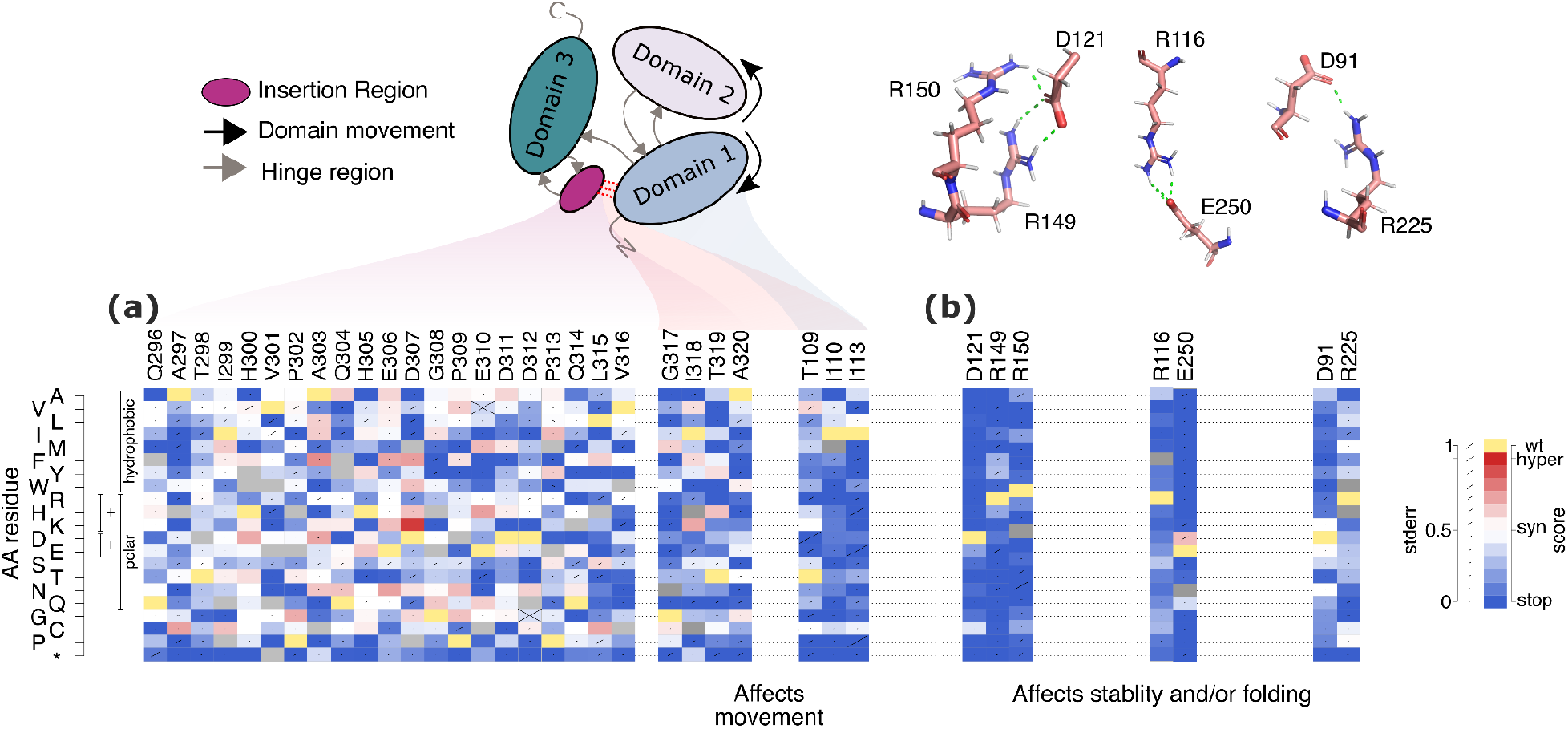
Placing functional impact scores in the context of residue roles in moderating HMBS fluctuation, protein stability or folding. Functional scores for each possible substituted amino acid (*y*-axis) at each residue position outside the active site (*x*-axis) participating in: (a) constraining HMBS domain movement, or (b) salt bridges. For each substitution, diagonal bar sizes convey estimated measurement error in the corresponding functional score. Box color either indicates the wild-type residue (yellow), or a substitution with damaging (blue), tolerated (white), or above-wildtype (‘hyper-complementing’, red) functional score, or missing data (gray).

**Supplemental Figure S5.**
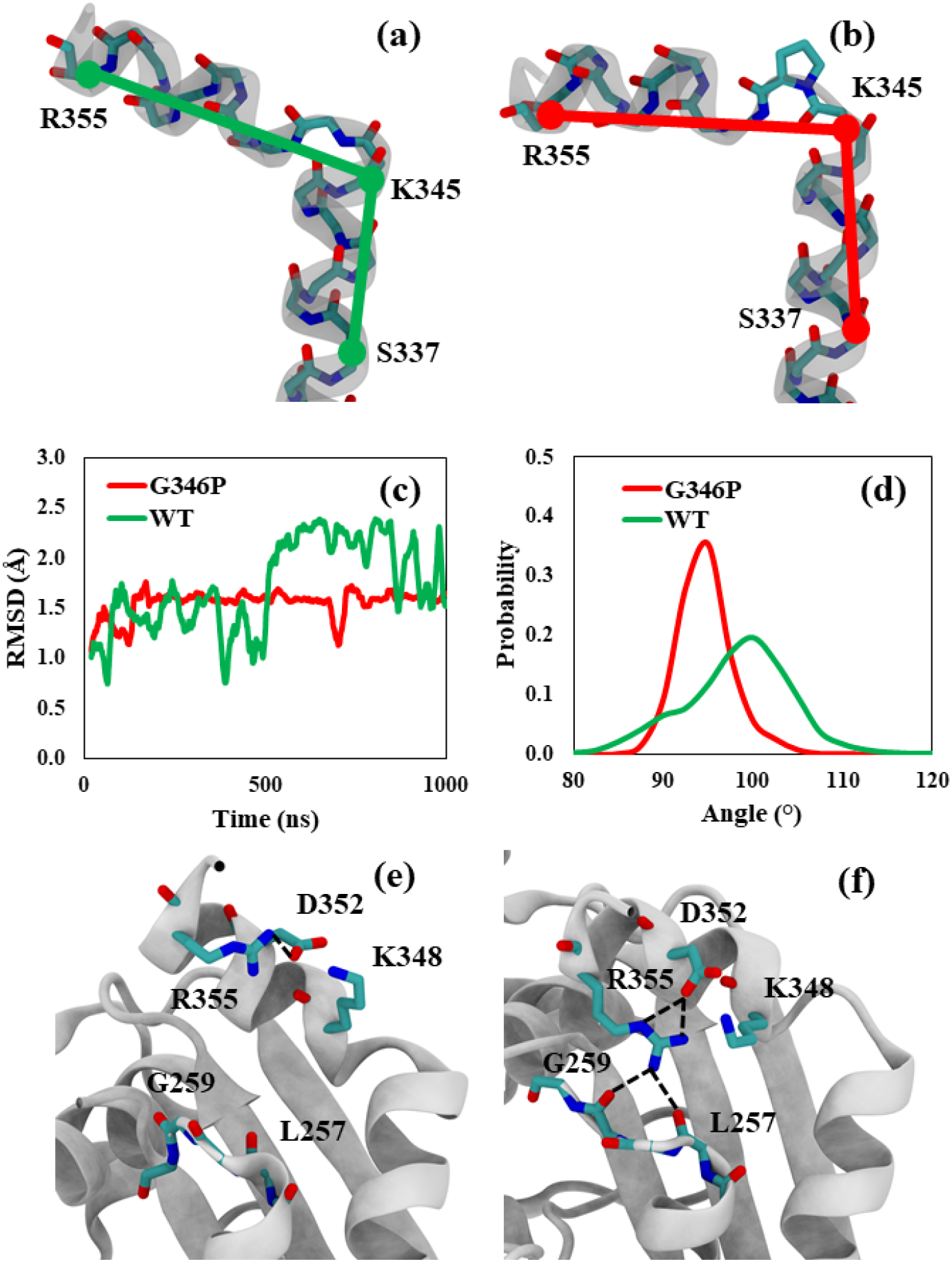
Comparison between WT and the G346P clinical variant showing a widespread impact on protein flexibility. Backbone atoms of S337, K345, and R355 were selected to evaluate the angle between the two helices in (a) WT and (b) G346P. For clarity only the C-terminal helices are shown. (c) The roost-mean-squared deviation (RMSD) of the backbone atoms in the C-terminal helix (K345-L357). (d) Angle distribution for WT (average: 100° ± 6) and G346 (average: 96° ± 3). Interactions of R355 with neighboring residues in (e) WT and (f) G346P

**Supplemental Figure S6.**
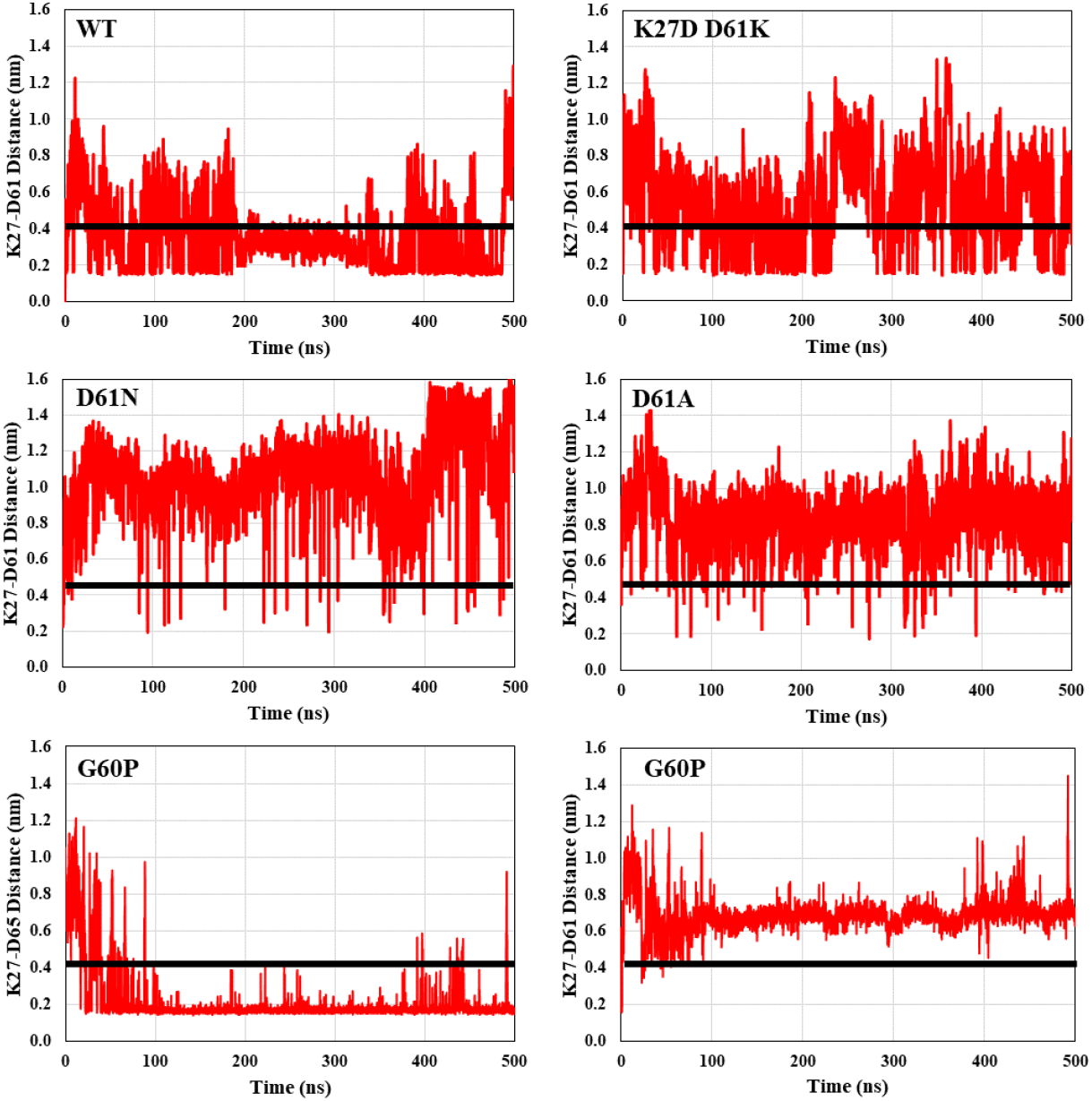
Monitoring open vs. closed active site loop status based on average distance between salt bridge residues 27 and 61 over time: WT (top left), K27D D61K (top right), D61N (mid left), D61A (mid right), and G60P (bottom). Bottom right shows the average distance between an alternative salt bridge between residues 27 and 65 in the G60P mutant. The salt bridge formation threshold (4 Å) is shown by a black solid line.

**Supplemental Figure S7.**
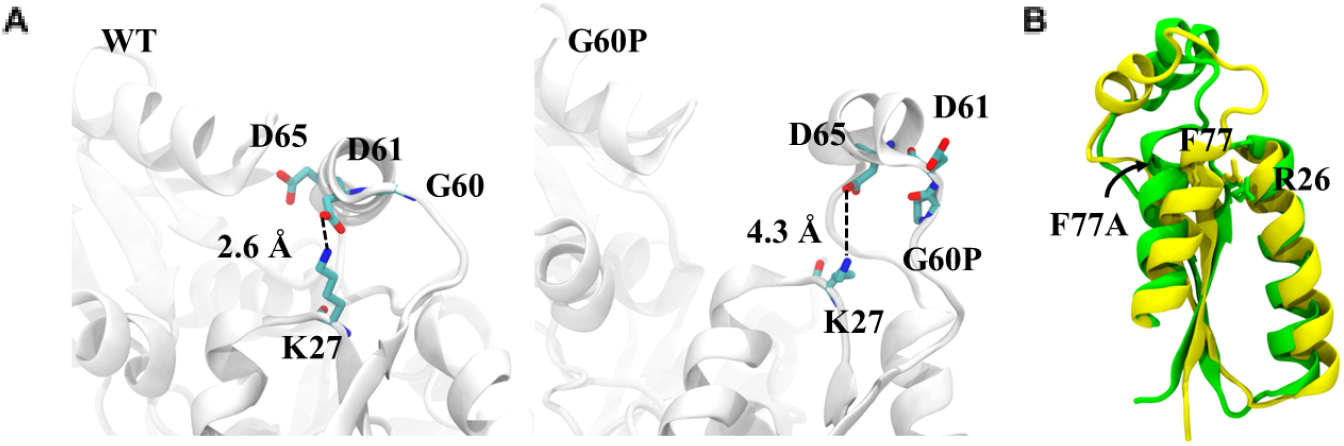
Modeling the effect of G60P and F77A variants on active site loop conformation. (A) Comparison of salt bridge distances (Å) in WT HMBS (left) and the G60P variant (right). The D61-K27 interaction is lost with the G60P variant, and instead replaced by a D65-K27 salt bridge that retains the active site loop in the “open” state. (B) The F77A variant (green) results in a dislocation of the active-site loop compared to the WT (yellow), exposing the active site.

**Supplemental Figure S8.**
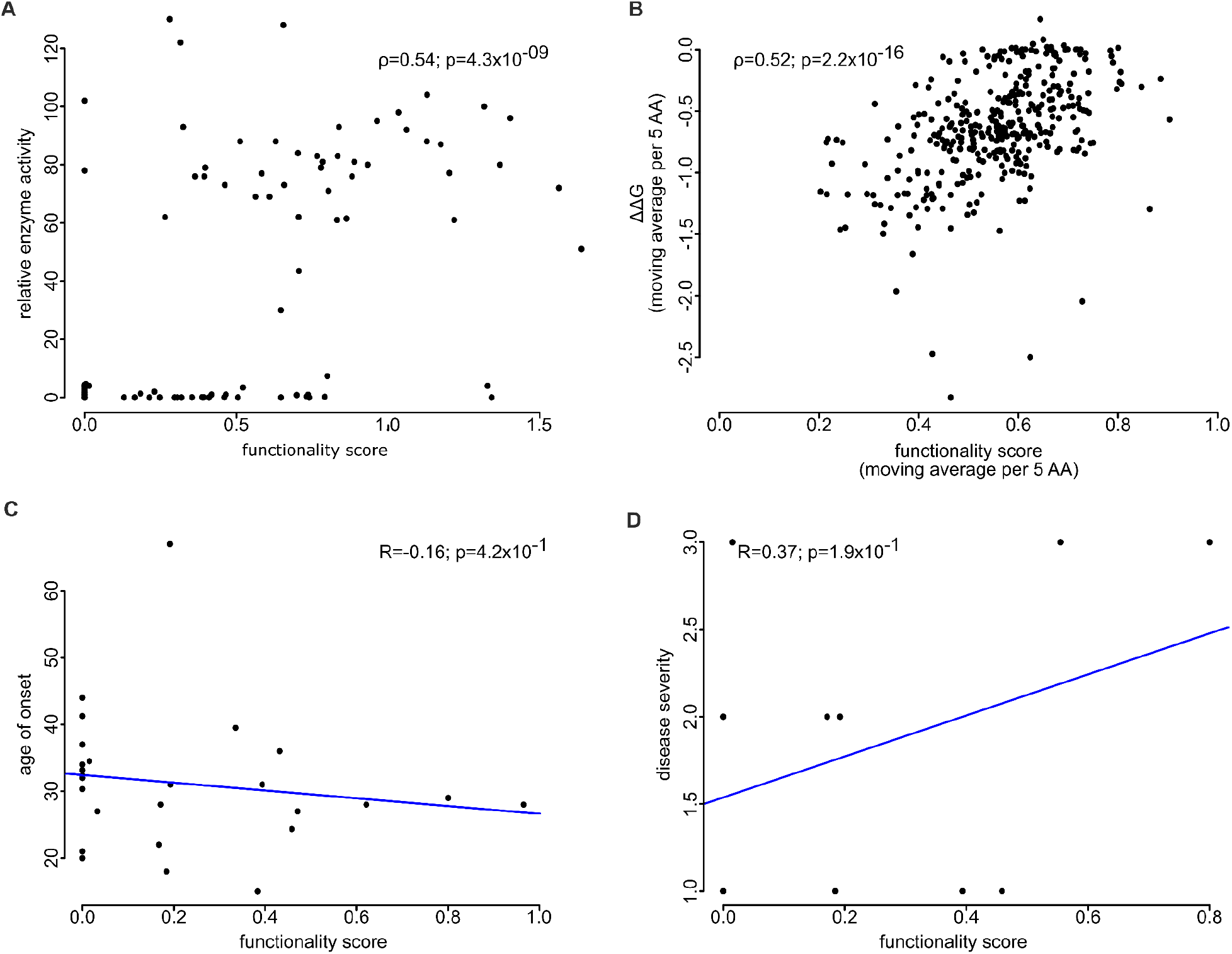
Correlation of functional impact scores from the combined map with other variant properties. Either Spearman’s rank correlation (ρ) with significance (p) of correlation, or Pearson correlation (R) with significance (p) are shown. (A) Correlation (ϱ *=* 0.54, p= 4 ×10^−9^) with HMBS relative enzyme activity (variant activity divided by wild type activity). (B) Correlation (ϱ*=* 0.52, p= 0.2) of running medians (interval size of 5 amino acids) of the combined map scores and predicted folding free energy change (ΔΔG) values. (C) Correlation (R=-0.16; P = 0.4) with age of AIP onset. (D) Correlation (R=0.37; P = 0.2) with AIP severity scores (3 = mild disease, 2 = moderate disease, 1 = severe disease).

**Supplemental Figure S9.**
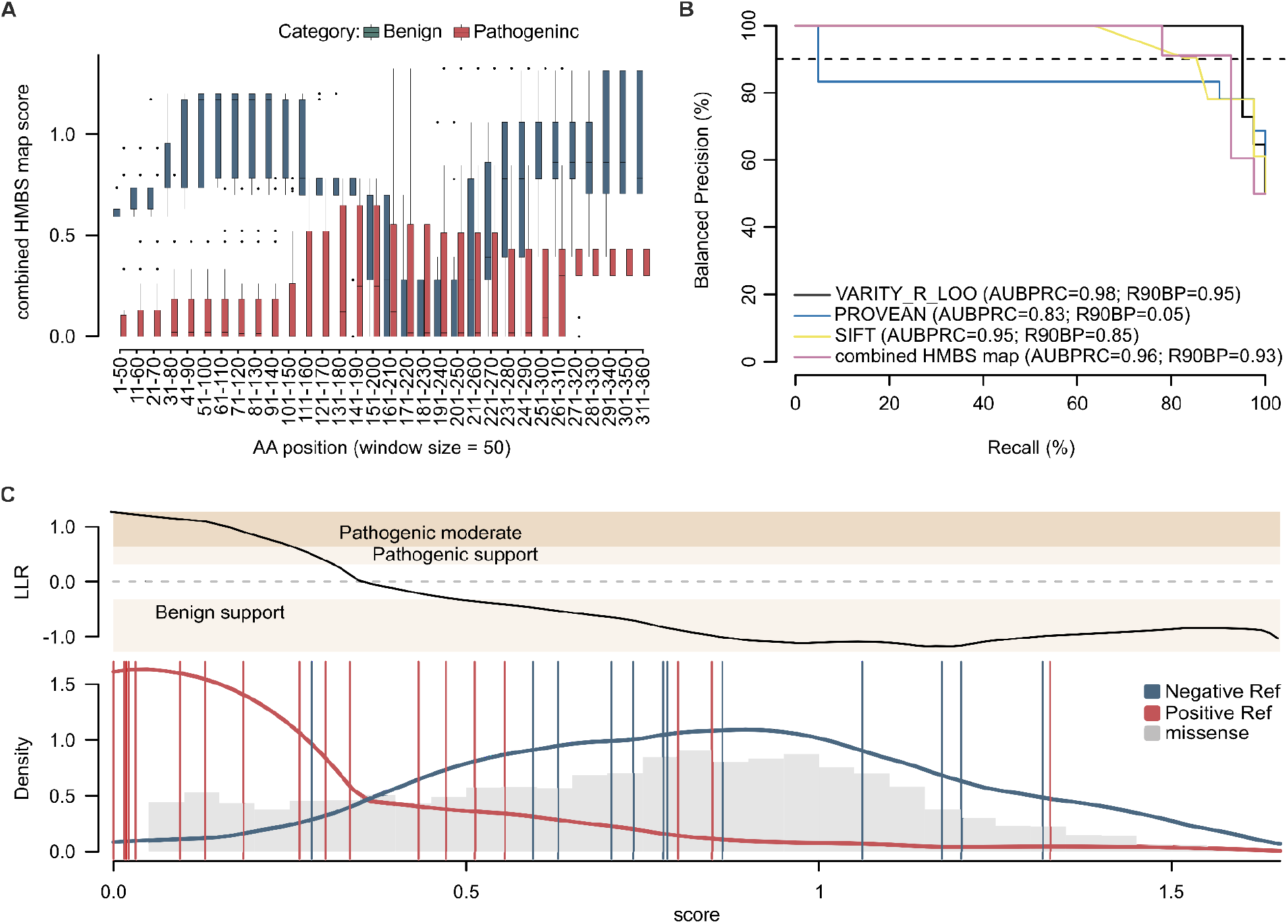
Evaluating the ability of the combined HMBS map, as well as computational predictors, to distinguish positive from random reference variants and interpret clinically-relevant missense variants from the ClinVar database. (A) Moving window analysis of scores from the combined map, as part of the positive (red) or negative (blue) reference variant sets. Plotted values are averages within windows of 50 amino acid (AA) positions. (B) Evaluation of precision (fraction of variants scoring below each threshold functional impact score that are in the positive reference set containing pathogenic variants) vs recall (fraction of positive reference variants with functionality scores below threshold). Here, precision has been balanced to reflect performance in a balanced test setting where positive and negative sets contain the same number of variants, excluding all at residue positions 160-215. Balanced precision-recall curves are shown for the combined map (pink), computational predictors PROVEAN (green), SIFT (yellow), and VARITY (black). Performance is also described in terms of area under the balanced precision vs recall curve (AUBPRC) and recall at a balanced precision of 90% (R90BP). (C) Transformation functions represent variant effects in terms of the strength of evidence for and against pathogenicity, that is, a log-likelihood ratio (LLR) of pathogenicity. The functions (top) express the log ratio between the likelihood of observing a given score in the score distribution of the positive reference (red) set as opposed to that of the negative reference set (blue). Gray histogram bars show the distribution of missense variants for comparison.

Supplemental Figure S10. Full-sized HMBS variant effect maps.

(A) Complete functional maps for erythroid-specific and ubiquitous HMBS isoforms, and the weighted average score values. Colors and labels are as in Figure 2.

(B) Delta map, measuring ubiquitous − erythroid-specific functional impact. Substitutions were coloured red if the score is positive and blue if negative.

## Supplemental Tables

**Table S1.** Comparing different models for effects of activity enhancing mutations.

| Model | ΔAIC relative to best model |
| --- | --- |
| Penalize enhancing mutations | 0 |
| Cap score of enhancing mutations at wild type level | 24.1 |
| Enhancing mutations are beneficial | 38.0 |

**Table S2.**
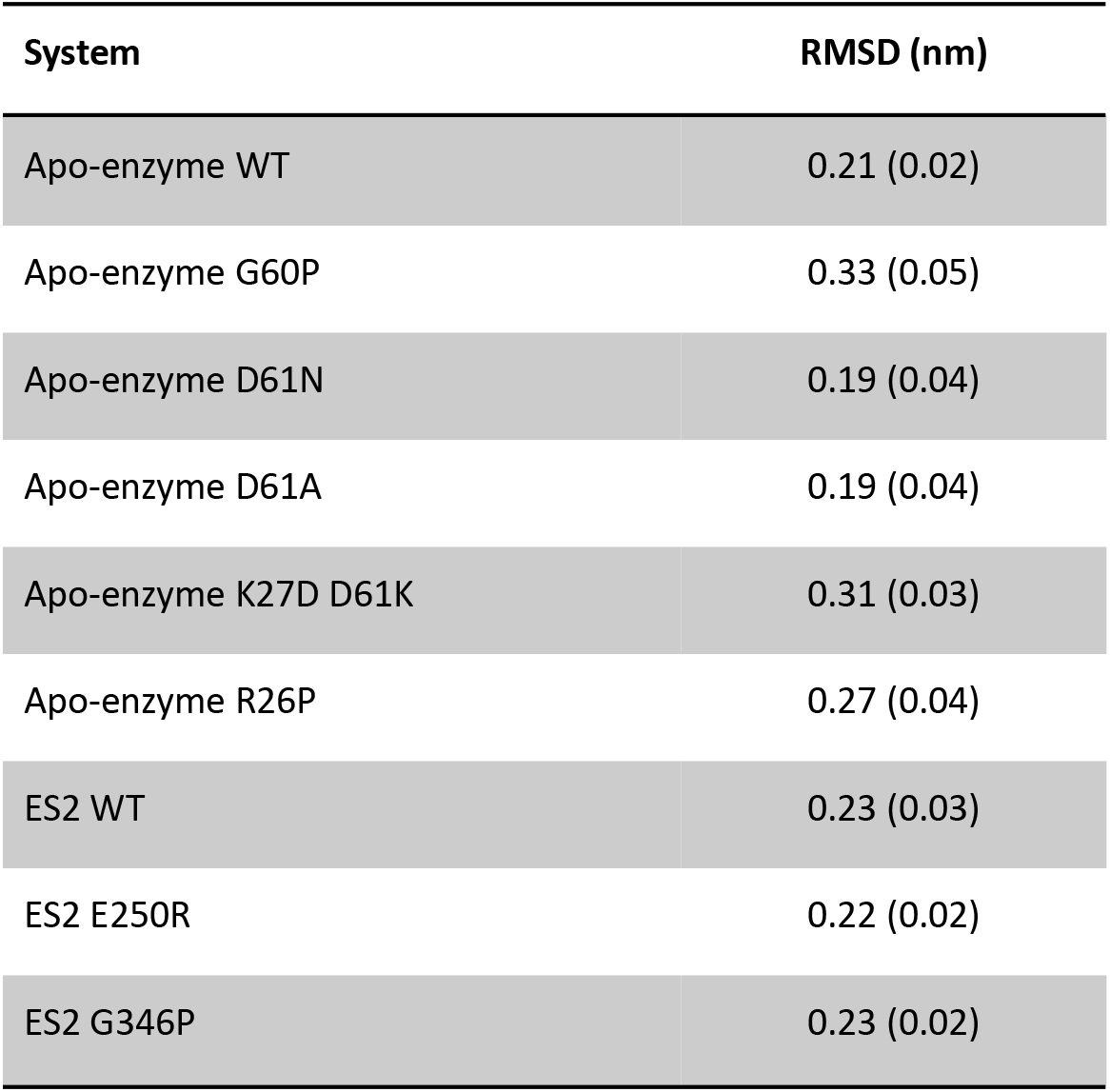
Average RMSD (standard deviation in parentheses) of backbone atoms for the wild-type (WT) or enzyme substrate complex 2 (ES2) structures.

| System | RMSD (nm) |
| --- | --- |
| Apo-enzyme WT | 0.21 (0.02) |
| Apo-enzyme G60P | 0.33 (0.05) |
| Apo-enzyme D61N | 0.19 (0.04) |
| Apo-enzyme D61A | 0.19 (0.04) |
| Apo-enzyme K27D D61K | 0.31 (0.03) |
| Apo-enzyme R26P | 0.27 (0.04) |
| ES2 WT | 0.23 (0.03) |
| ES2 E250R | 0.22 (0.02) |
| ES2 G346P | 0.23 (0.02) |

**Table S3.** Hydrogen bond occupancy (%) between HMBS important residues and the substrate in different systems. For residues with more than one interaction with PBG, only the more persistent interaction is reported.

| DPM-PBG |  |  |  |
| --- | --- | --- | --- |
| Residue | WT | E250R | G346P |
| R149 | 100 | 82 | 87 |
| R150 | 100 | 85 | 100 |
| G218 | 100 | - | 95 |
| S147 | 99 | 57 | 65 |
| S96 | 99 | 55 | 65 |
| A189 | 98 | 56 | 73 |
| R173 | 98 | 95 | 42 |
| T102 | 100 | 58 | 70 |
| L148 | 72 | - | - |
| K98 | 100 | 100 | 100 |
| S146 | 68 | 99 | 54 |
| R195 | 57 | 76 | 50 |
| R117 | - | 93 | 86 |
| T145 | 68 | 85 | 38 |
| D99* | 94, 80 | 20, 4 | 7, 6 |

\*Numbers correspond to hydrogen bonds with the 4th (second PBG) and 3rd (first PBG) pyrrole ring, respectively.

**Table S4.** Hydrogen bond occupancy (%) in the C-terminal helix region for WT and the G346P variant.

|  |  | Occupancy (%) |  |
| --- | --- | --- | --- |
| Backbone Hydrogen Bonds |  | G346P | WT |
| L342 (O) | K345 (NH) | 80 | - |
| L342 (O) | A347 (NH) | - | 87 |
| L342 (O) | G346 (NH) | - | 57 |
| L342 (NH) | L338 (O) | 99 | 100 |
| L343(O) | G346 (O) | - | 46 |
| L343 (NH) | A339 (O) | 92 | 97 |
| S344 (NH) | L341 (O) | 62 | 20 |
| S344 (NH) | N340 (O) | - | 78 |
| K345 (O) | A347 (NH) | 91 | - |
| K345 (NH) | L342 (O) | 80 | - |
| K345 (NH) | L341 (O) | - | 90 |
| G346 (O) | N349 (NH) | - | 55 |
| G346 (NH) | L342 (O) | - | 57 |
| G346 (NH) | L343 (O) | - | 46 |
| P346 (O) | N349 (NH) | 47 | - |
| P346 (O) | I350 (NH) | 32 | - |
| A347 (O) | I350 (NH) | 49 | 65 |
| A347 (O) | L351 (NH) | 92 | 87 |
| A347 (NH) | K345 (O) | 91 | - |
| A347 (NH) | L342 (O) | - | 87 |

**Table S5.** Hydrogen bond occupancy (%) for R355 in WT and the G346P variant.

| G346P |  |  |
| --- | --- | --- |
| Hydrogen Bond |  | Occupancy (%) |
| R355 (NH1) | G259 (O) | 82 |
| R355 (NH2) | L257 (O) | 63 |
| R355 (NH22) | D352 (Oδ2) | 59 |
| R355 (NH21) | D352 (Oδ1) | 32 |
| WT |  |  |
| Hydrogen Bond |  | Occupancy (%) |
| R355 (NH1) | D352 (Oδ1) | 19 |
| R355 (NH) | D352 (O) | 16 |

**Table S6.** Average distance (Å) between selected residue pairs in WT and variants.

|  | WT | K27D D61K | D61N | D61A | G60P | R26P |
| --- | --- | --- | --- | --- | --- | --- |
| R26-D61 | 7.6 | 8.6 | 12 | 11 | 14 | 13 |
| K27-D61 | 3.5 | 5.4 | 11 | 8.5 | 6.9 | 10 |
| G60-Q34 | 17 | 18 | 20 | 20 | 21 | 20 |
| G60-R26 | 9.1 | 8.9 | 10 | 11 | 13 | 12 |

**Table S7**. Curated reference variant sets for validation

**Table S8**. HMBS missense variants and corresponding scores, log likelihood ratios of pathogenicity and calibrated evidence strength labels

**Table S9**. List of TileSeq primers and POPcode mutagenesis oligos

**Table S10**. Ensembl homologs having at least 85% sequence identity to the human HMBS protein

## Acknowledgements

We gratefully acknowledge funding for this project from Alnylam Pharmaceuticals. We further acknowledge the National Institutes of Health National Human Genome Research Institute (NIH/NHGRI) Center of Excellence in Genomic Science Initiative (HG010461), the NIH/NHGRI Impact of Genomic Variation on Function (IGVF) Initiative (UM1HG011989), the Canada Excellence Research Chairs Program, and a Canadian Institutes of Health Research Foundation Grant to F.R.. Computational resources for the molecular dynamics simulations were provided by Compute Canada and SharcNet. We acknowledge Sharon D Whatley for the provision of variant data.

## References

1. Bissell, D. M., Anderson, K. E. & Bonkovsky, H. L. Porphyria. N. Engl. J. Med. 377, 862–872 (2017).

2. Chen, B. et al. Acute Intermittent Porphyria: Predicted Pathogenicity of HMBS Variants Indicates Extremely Low Penetrance of the Autosomal Dominant Disease. Hum. Mutat. 37, 1215–1222 (2016).

3. Baumann, K. & Kauppinen, R. Penetrance and predictive value of genetic screening in acute porphyria. Mol. Genet. Metab. 130, 87–99 (2020).

4. Lenglet, H. et al. From a dominant to an oligogenic model of inheritance with environmental modifiers in acute intermittent porphyria. Hum. Mol. Genet. 27, 1164–1173 (2018).

5. Grandchamp, B. et al. Tissue-specific expression of porphobilinogen deaminase. Two isoenzymes from a single gene. Eur. J. Biochem. 162, 105–110 (1987).

6. Chretien, S. et al. Alternative transcription and splicing of the human porphobilinogen deaminase gene result either in tissue-specific or in housekeeping expression. Proc. Natl. Acad. Sci. U. S. A. 85, 6–10 (1988).

7. Whatley, S. D. et al. Non-erythroid form of acute intermittent porphyria caused by promoter and frameshift mutations distant from the coding sequence of exon 1 of the HMBS gene. Hum. Genet. 107, 243–248 (2000).

8. San Juan, I. et al. ALAD Inhibition by Porphobilinogen Rationalizes the Accumulation of δ-Aminolevulinate in Acute Porphyrias. Biochemistry 61, 2409–2416 (2022).

9. Simon, A. et al. Patient Perspective on Acute Intermittent Porphyria with Frequent Attacks: A Disease with Intermittent and Chronic Manifestations. Patient 11, 527–537 (2018).

10. Elder, G., Harper, P., Badminton, M., Sandberg, S. & Deybach, J.-C. The incidence of inherited porphyrias in Europe. J. Inherit. Metab. Dis. 36, 849–857 (2013).

11. Stein, P. E., Badminton, M. N. & Rees, D. C. Update review of the acute porphyrias. Br. J. Haematol. 176, 527–538 (2017).

12. Molina, L. et al. Bi-allelic hydroxymethylbilane synthase inactivation defines a homogenous clinico-molecular subtype of hepatocellular carcinoma. J. Hepatol. 77, 1038–1046 (2022).

13. Landrum, M. J. et al. ClinVar: public archive of relationships among sequence variation and human phenotype. Nucleic Acids Res. 42, D980–5 (2014).

14. Pejaver, V. et al. Evidence-based calibration of computational tools for missense variant pathogenicity classification and ClinGen recommendations for clinical use of PP3/BP4 criteria. bioRxiv 2022.03.17.484479 (2022) doi:10.1101/2022.03.17.484479.

15. Richards, S. et al. Standards and guidelines for the interpretation of sequence variants: a joint consensus recommendation of the American College of Medical Genetics and Genomics and the Association for Molecular Pathology. Genet. Med. 17, 405–424 (2015).

16. Tabet, D., Parikh, V., Mali, P., Roth, F. P. & Claussnitzer, M. Scalable Functional Assays for the Interpretation of Human Genetic Variation. Annu. Rev. Genet. 56, 441–465 (2022).

17. Floyd, B. et al. Proactive variant effect mapping to accelerate genetic diagnosis for pediatric cardiac arrest. Preprints (2022) doi:10.20944/preprints202201.0177.v1.

18. Fayer, S. et al. Closing the gap: Systematic integration of multiplexed functional data resolves variants of uncertain significance in BRCA1, TP53, and PTEN. Am. J. Hum. Genet. 108, 2248–2258 (2021).

19. Giacomelli, A. O. et al. Mutational processes shape the landscape of TP53 mutations in human cancer. Nat. Genet. 50, 1381–1387 (2018).

20. Jia, X. et al. Massively parallel functional testing of MSH2 missense variants conferring Lynch syndrome risk. Am. J. Hum. Genet. 108, 163–175 (2021).

21. Starita, L. M. et al. Massively Parallel Functional Analysis of BRCA1 RING Domain Variants. Genetics 200, 413–422 (2015).

22. Mighell, T. L., Evans-Dutson, S. & O’Roak, B. J. A Saturation Mutagenesis Approach to Understanding PTEN Lipid Phosphatase Activity and Genotype-Phenotype Relationships. Am. J. Hum. Genet. 102, 943–955 (2018).

23. Sun, S. et al. An extended set of yeast-based functional assays accurately identifies human disease mutations. Genome Res. 26, 670–680 (2016).

24. Weile, J. et al. Shifting landscapes of human MTHFR missense-variant effects. Am. J. Hum. Genet. 108, 1283–1300 (2021).

25. Sun, S. et al. A proactive genotype-to-patient-phenotype map for cystathionine beta-synthase. Genome Med. 12, 13 (2020).

26. Kachroo, A. H. et al. Systematic bacterialization of yeast genes identifies a near-universally swappable pathway. Elife 6, (2017).

27. Weile, J. et al. A framework for exhaustively mapping functional missense variants. Mol. Syst. Biol. 13, 957 (2017).

28. Baldi, P. & Long, A. D. A Bayesian framework for the analysis of microarray expression data: regularized t-test and statistical inferences of gene changes. Bioinformatics 17, 509–519 (2001).

29. Esposito, D. et al. MaveDB: an open-source platform to distribute and interpret data from multiplexed assays of variant effect. Genome Biol. 20, 223 (2019).

30. Bloom, J. D. An experimentally determined evolutionary model dramatically improves phylogenetic fit. Mol. Biol. Evol. 31, 1956–1978 (2014).

31. Bloom, J. D. Identification of positive selection in genes is greatly improved by using experimentally informed site-specific models. Biol. Direct 12, 1 (2017).

32. Bogorad, L. The enzymatic synthesis of porphyrins from porphobilinogen. I. Uroporphyrin I. J. Biol. Chem. 233, 501–509 (1958).

33. Anderson, P. M. & Desnick, R. J. Purification and properties of uroporphyrinogen I synthase from human erythrocytes. Identification of stable enzyme-substrate intermediates. J. Biol. Chem. 255, 1993–1999 (1980).

34. Jordan, P. M., Thomas, S. D. & Warren, M. J. Purification, crystallization and properties of porphobilinogen deaminase from a recombinant strain of Escherichia coli K12. Biochem. J 254, 427–435 (1988).

35. Bung, N. et al. Human hydroxymethylbilane synthase: Molecular dynamics of the pyrrole chain elongation identifies step-specific residues that cause AIP. Proc. Natl. Acad. Sci. U. S. A. 115, E4071–E4080 (2018).

36. Bustad, H. J. et al. Characterization of porphobilinogen deaminase mutants reveals that arginine-173 is crucial for polypyrrole elongation mechanism. iScience 24, 102152 (2021).

37. Sato, H. et al. Crystal structures of hydroxymethylbilane synthase complexed with a substrate analog: a single substrate-binding site for four consecutive condensation steps. Biochem. J 478, 1023–1042 (2021).

38. Pluta, P. et al. Structural basis of pyrrole polymerization in human porphobilinogen deaminase. Biochim. Biophys. Acta Gen. Subj. 1862, 1948–1955 (2018).

39. Gill, R. et al. Structure of human porphobilinogen deaminase at 2.8 A: the molecular basis of acute intermittent porphyria. Biochem. J 420, 17–25 (2009).

40. Louie, G. V. Porphobilinogen deaminase and its structural similarity to the bidomain binding proteins. Curr. Opin. Struct. Biol. 3, 401–408 (1993).

41. Høie, M. H., Cagiada, M., Beck Frederiksen, A. H., Stein, A. & Lindorff-Larsen, K. Predicting and interpreting large-scale mutagenesis data using analyses of protein stability and conservation. Cell Rep. 38, 110207 (2022).

42. Cagiada, M. et al. Understanding the Origins of Loss of Protein Function by Analyzing the Effects of Thousands of Variants on Activity and Abundance. Mol. Biol. Evol. 38, 3235–3246 (2021).

43. Medlock, A. E. et al. Identification of the Mitochondrial Heme Metabolism Complex. PLoS One 10, e0135896 (2015).

44. Chen, B. et al. Identification and characterization of 40 novel hydroxymethylbilane synthase mutations that cause acute intermittent porphyria. J. Inherit. Metab. Dis. 42, 186–194 (2019).

45. Christie, M. S., Laitaoja, M., Aarsand, A. K., Kallio, J. P. & Bustad, H. J. Characterisation of a common hotspot variant in acute intermittent porphyria sheds light on the mechanism of hydroxymethylbilane synthase function. FEBS Open Bio 12, 2136–2146 (2022).

46. Fu, Y. et al. Systematically Analyzing the Pathogenic Variations for Acute Intermittent Porphyria. Front. Pharmacol. 10, 1018 (2019).

47. Wu, Y. et al. Improved pathogenicity prediction for rare human missense variants. Am. J. Hum. Genet. 108, 2389 (2021).

48. Choi, Y., Sims, G. E., Murphy, S., Miller, J. R. & Chan, A. P. Predicting the functional effect of amino acid substitutions and indels. PLoS One 7, e46688 (2012).

49. Sim, N.-L. et al. SIFT web server: predicting effects of amino acid substitutions on proteins. Nucleic Acids Res. 40, W452–7 (2012).

50. Wu, Y., Li, R., Sun, S., Weile, J. & Roth, F. P. Improved pathogenicity prediction for rare human missense variants. Am. J. Hum. Genet. 108, 1891–1906 (2021).

51. Tavtigian, S. V. et al. Modeling the ACMG/AMP variant classification guidelines as a Bayesian classification framework. Genet. Med. 20, 1054–1060 (2018).

52. Matreyek, K. A. et al. Multiplex assessment of protein variant abundance by massively parallel sequencing. Nat. Genet. 50, 874–882 (2018).

53. Yang, X. et al. A public genome-scale lentiviral expression library of human ORFs. Nat. Methods 8, 659–661 (2011).

54. Hilton, S. K., Doud, M. B. & Bloom, J. D. phydms: software for phylogenetic analyses informed by deep mutational scanning. PeerJ 5, e3657 (2017).

55. Anandakrishnan, R., Aguilar, B. & Onufriev, A. V. H++ 3.0: automating pK prediction and the preparation of biomolecular structures for atomistic molecular modeling and simulations. Nucleic Acids Res. 40, W537–41 (2012).

56. Maier, J. A. et al. ff14SB: Improving the Accuracy of Protein Side Chain and Backbone Parameters from ff99SB. J. Chem. Theory Comput. 11, 3696–3713 (2015).

57. Jorgensen, W. L., Chandrasekhar, J., Madura, J. D., Impey, R. W. & Klein, M. L. Comparison of simple potential functions for simulating liquid water. J. Chem. Phys. 79, 926–935 (1983).

58. Becke, A. D. Density-functional thermochemistry. III. The role of exact exchange. J. Chem. Phys. 98, 5648–5652 (1993).

59. Lee, C., Yang, W. & Parr, R. G. Development of the Colle-Salvetti correlation-energy formula into a functional of the electron density. Phys. Rev. B Condens. Matter 37, 785–789 (1988).

60. Vosko, S. H., Wilk, L. & Nusair, M. Accurate spin-dependent electron liquid correlation energies for local spin density calculations: a critical analysis. Can. J. Phys. (1980).

61. Frisch, M. J., Trucks, G. W., Schlegel, H. B. & Scuseria, G. E. Gaussian 16, Revision C. 01. Gaussian, Inc., Wallingford CT. 2016. Google Scholar There is no.

62. Case, D. A., Ben-Shalom, I. Y., Brozell, S. R. & Cerutti, D. S. AMBER; University of California: San Francisco, 2018. is no corresponding record for this ….

63. Ryckaert, J.-P., Ciccotti, G. & Berendsen, H. J. C. Numerical integration of the cartesian equations of motion of a system with constraints: molecular dynamics of n-alkanes. J. Comput. Phys. 23, 327–341 (1977).

64. Darden, T., York, D. & Pedersen, L. Particle mesh Ewald: An N·log(N) method for Ewald sums in large systems. J. Chem. Phys. 98, 10089–10092 (1993).

65. Essmann, U. et al. A smooth particle mesh Ewald method. J. Chem. Phys. 103, 8577–8593 (1995).

66. Roe, D. R. & Cheatham, T. E., 3rd. PTRAJ and CPPTRAJ: Software for Processing and Analysis of Molecular Dynamics Trajectory Data. J. Chem. Theory Comput. 9, 3084–3095 (2013).

67. Montanucci, L., Capriotti, E., Frank, Y., Ben-Tal, N. & Fariselli, P. DDGun: an untrained method for the prediction of protein stability changes upon single and multiple point variations. BMC Bioinformatics 20, 335 (2019).

68. Song, G. et al. Structure of human porphobilinogen deaminase. Preprint at https://doi.org/10.2210/pdb3ecr/pdb (2008).

