## Supplementary figures and images for "Systematically testing human HMBS missense variants to reveal mechanism and pathogenic variation"

### Supplemental Figure S10

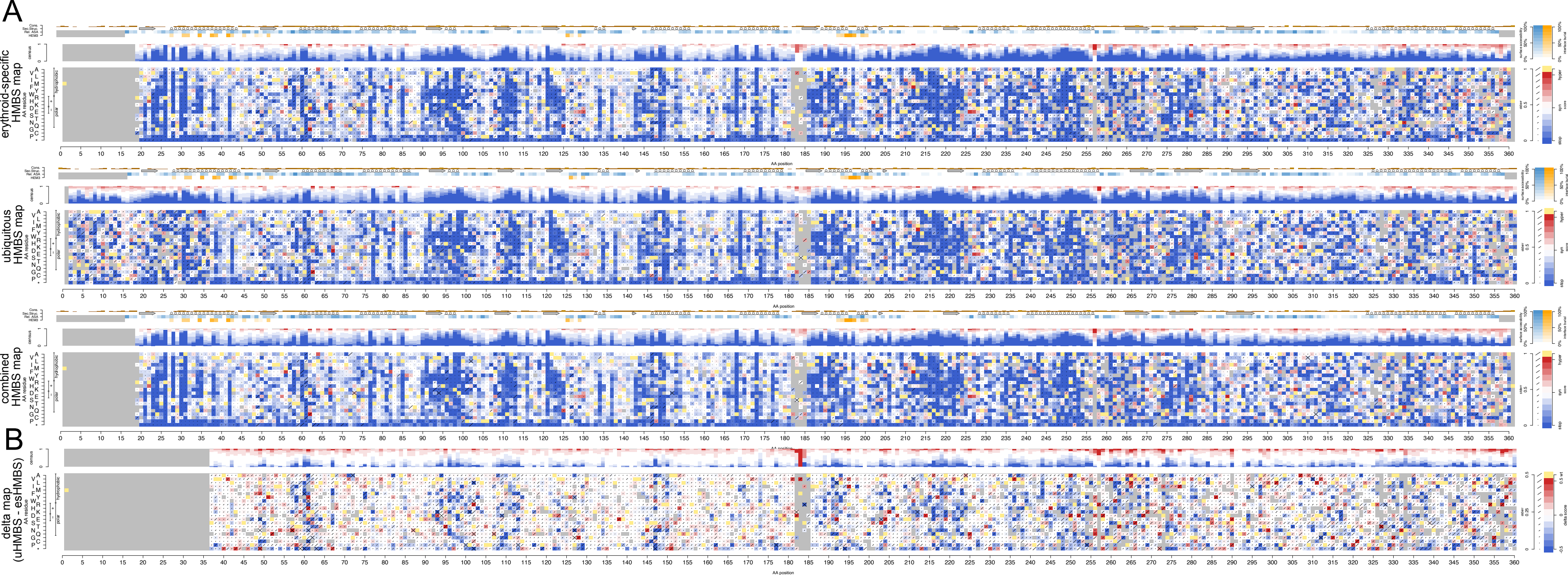
