## Supplemental Table S7 for "Systematically testing human HMBS missense variants to reveal mechanism and pathogenic variation"

Table S7: Curated reference set of disease- and non-disease-associated variants

| protein consequence | nucleotide change | wt aa | pos | mut aa |
|---|---|---|---|---|
| p.Ala219Asp | c.0656C>A | Ala | 219 | Asp |
| p.Ala31Pro | c.0091G>C | Ala | 31 | Pro |
| p.Ala31Thr | c.0091G>A | Ala | 31 | Thr |
| p.Arg116Gln | c.0347G>A | Arg | 116 | Gln |
| p.Arg116Trp | c.0346C>T | Arg | 116 | Trp |
| p.Arg149Gln | c.0446G>A | Arg | 149 | Gln |
| p.Arg149Leu | c.0446G>T | Arg | 149 | Leu |
| p.Arg167Gln | c.0500G>A | Arg | 167 | Gln |
| p.Arg167Trp | c.0499C>T | Arg | 167 | Trp |
| p.Arg173Gln | c.0518G>A; c.0578G>A | Arg | 173 | Gln |
| p.Arg173Trp | c.0517C>T | Arg | 173 | Trp |
| p.Arg195Cys | c.0583C>T | Arg | 195 | Cys |
| p.Arg195His | c.0584G>A | Arg | 195 | His |
| p.Arg201Trp | c.0601C>T | Arg | 201 | Trp |
| p.Arg225Gln | c.0674G>A | Arg | 225 | Gln |
| p.Arg246Cys | c.0736C>T | Arg | 246 | Cys |
| p.Arg255Ser | c.0765G>C | Arg | 255 | Ser |
| p.Arg26Cys | c.0076C>T | Arg | 26 | Cys |
| p.Arg26His | c.0077G>A | Arg | 26 | His |
| p.Arg321His | c.0962G>A | Arg | 321 | His |
| p.Arg32His | c.0095G>A | Arg | 32 | His |
| p.Asp230Tyr | c.0688G>T | Asp | 230 | Tyr |
| p.Asp359Asn | c.1075G>A | Asp | 359 | Asn |
| p.Asp61Tyr | c.0181G>A | Asp | 61 | Tyr |
| p.Asp99Gly | c.0296A>G | Asp | 99 | Gly |
| p.Cys247Arg | c.0739T>C | Cys | 247 | Arg |
| p.Gln296Arg | c.0887A>G | Gln | 296 | Arg |
| p.Glu250Ala | c.0749A>C | Glu | 250 | Ala |
| p.Glu250Lys | c.0748G>A | Glu | 250 | Lys |
| p.Glu86Val | c.0257A>T | Glu | 86 | Val |
| p.Gly111Arg | c.0331G>A | Gly | 111 | Arg |
| p.Gly168Arg | c.0502G>A | Gly | 168 | Arg |
| p.Gly216Asp | c.0647G>A | Gly | 216 | Asp |
| p.Gly280Glu | c.0839G>A | Gly | 280 | Glu |
| p.Gly335Asp | c.1004G>A | Gly | 335 | Asp |
| p.Ile113Thr | c.0338T>C | Ile | 113 | Thr |
| p.Leu177Arg | c.0530T>G | Leu | 177 | Arg |
| p.Leu234Pro | c.0701T>C | Leu | 234 | Pro |
| p.Leu238Pro | c.0713T>C | Leu | 238 | Pro |
| p.Leu254Pro | c.0761T>C | Leu | 254 | Pro |
| p.Leu276Arg | c.0827T>G | Leu | 276 | Arg |
| p.Leu278Arg | c.0833T>G | Leu | 278 | Arg |
| p.Leu30Ile | c.0088C>A | Leu | 30 | Ile |

| | | | | |
|---|---|---|---|---|
| p.Leu30Phe | c.0088C>T | Leu | 30 | Phe |
| p.Leu30Pro | c.0089T>C | Leu | 30 | Pro |
| p.Leu343Pro | c.1028T>C | Leu | 343 | Pro |
| p.Leu42Ser | c.0125T>C | Leu | 42 | Ser |
| p.Leu85Arg | c.0254T>G | Leu | 85 | Arg |
| p.Leu92Pro | c.0275T>C | Leu | 92 | Pro |
| p.Lys132Asn | c.0396G>C | Lys | 132 | Asn |
| p.Lys348Arg | c.1043A>G | Lys | 348 | Arg |
| p.Pro119Leu | c.0356C>T | Pro | 119 | Leu |
| p.Pro127Gln | c.0380C>A | Pro | 127 | Gln |
| p.Pro159Leu | c.0476C>T | Pro | 159 | Leu |
| p.Pro309Ser | c.0925C>T | Pro | 309 | Ser |
| p.Ser147Pro | c.0439T>C | Ser | 147 | Pro |
| p.Ser28Asn | c.0083G>A | Ser | 28 | Asn |
| p.Ser45Leu | c.0134C>T | Ser | 45 | Leu |
| p.Ser96Phe | c.0287C>T | Ser | 96 | Phe |
| p.Thr269Ile | c.0806C>T | Thr | 269 | Ile |
| p.Thr35Met | c.0104C>T | Thr | 35 | Met |
| p.Val124Asp | c.0371T>A | Val | 124 | Asp |
| p.Val215Glu | c.0644T>A | Val | 215 | Glu |
| p.Val267Leu | c.0799G>C | Val | 267 | Leu |
| p.Val267Met | c.0799G>A | Val | 267 | Met |
| p.Val90Gly | c.0269T>G | Val | 90 | Gly |

| clinical annotation | combined map | eHMBS map | uHMBS map | VARITY | PROVEAN |
|---|---|---|---|---|---|
| Pathogenic | 0.000 | 0.000 | 0.371 | 0.993 | -5.48 |
| Pathogenic | 0.000 | 0.000 | 0.581 | 0.994 | -4.6 |
| Pathogenic | 0.000 | 0.000 | 0.000 | 0.978 | -3.68 |
| Pathogenic | 0.264 | NA | 0.264 | 0.984 | -3.71 |
| Pathogenic | 0.184 | 0.161 | 0.202 | 0.97 | -7.35 |
| Pathogenic | 0.004 | 0.002 | 0.005 | 0.981 | -3.72 |
| Pathogenic | 0.000 | 0.145 | 0.100 | 0.986 | -6.5 |
| Pathogenic | 0.733 | 0.726 | 0.750 | 0.961 | -3.85 |
| Pathogenic | 0.522 | 0.519 | 0.576 | 0.955 | -7.68 |
| Pathogenic | 0.000 | 0.000 | 0.000 | 0.976 | -3.85 |
| Pathogenic | 0.000 | 0.000 | 0.000 | 0.963 | -7.72 |
| Pathogenic | 0.248 | 0.149 | 0.543 | 0.982 | -7.72 |
| Pathogenic | 0.783 | 0.719 | 0.854 | 0.989 | -4.82 |
| Pathogenic | 0.719 | 0.714 | 0.724 | 0.961 | -7.74 |
| Pathogenic | 0.000 | 0.585 | 0.000 | 0.913 | -3.42 |
| Benign | 0.281 | 0.306 | 0.272 | 0.824 | -5.75 |
| Benign | 0.883 | 0.954 | 0.841 | 0.765 | -2.98 |
| Pathogenic | 0.000 | 0.000 | 0.000 | 0.967 | -7.03 |
| Pathogenic | 0.000 | 0.000 | 0.287 | 0.936 | -4.39 |
| Benign | 0.863 | NA | 0.863 | 0.115 | -0.82 |
| Benign | 0.595 | 0.595 | 1.005 | 0.862 | -1.85 |
| Pathogenic | 0.512 | 0.459 | 0.581 | 0.458 | -2.08 |
| Benign | 0.706 | 0.723 | 0.650 | 0.377 | -1.51 |
| Pathogenic | 0.335 | 0.617 | 0.083 | 0.976 | -8.33 |
| Pathogenic | 0.130 | 0.121 | 0.289 | 0.989 | -6.5 |
| Pathogenic | 0.000 | 0.025 | 0.159 | 0.997 | -8.65 |
| Benign | 1.061 | 1.073 | 1.045 | 0.846 | -1.64 |
| Pathogenic | 0.000 | 0.526 | 0.000 | 0.983 | -5.36 |
| Pathogenic | 0.018 | 0.153 | 0.104 | 0.985 | -3.57 |
| Benign | 0.737 | 0.737 | 0.737 | 0.455 | -1.47 |
| Pathogenic | 0.015 | 0.093 | 0.131 | 0.976 | -6.26 |
| Pathogenic | 0.000 | 0.049 | 0.009 | 0.987 | -7.72 |
| Pathogenic | 0.000 | 0.000 | 0.390 | 0.992 | -6.1 |
| Pathogenic | 0.000 | 0.053 | 0.016 | 0.993 | -7.14 |
| Pathogenic | 0.432 | 0.219 | 0.522 | 0.968 | -5.69 |
| Pathogenic | 0.471 | 0.434 | 0.561 | 0.777 | -3.83 |
| Pathogenic | 0.121 | 0.108 | 0.144 | 0.987 | -5.77 |
| Pathogenic | 0.000 | 0.000 | 0.201 | 0.99 | -5.15 |
| Pathogenic | 0.555 | NA | 0.555 | 0.99 | -5.75 |
| Pathogenic | 0.432 | 0.000 | 0.445 | 0.998 | -5.71 |
| Pathogenic | 0.000 | 0.060 | 0.005 | 0.986 | -5.04 |
| Pathogenic | 0.094 | 0.106 | 0.083 | 0.994 | -5.45 |
| Pathogenic | 0.000 | 0.000 | 0.675 | 0.957 | -1.84 |

| | | | | | |
|---|---|---|---|---|---|
| Pathogenic | 0.000 | 0.000 | 0.000 | 0.972 | -3.68 |
| Pathogenic | 0.000 | 0.000 | 0.000 | 0.992 | -6.45 |
| Pathogenic | 0.300 | 0.331 | 0.211 | 0.915 | -3.71 |
| Pathogenic | 0.000 | 0.000 | 0.000 | 0.985 | -5.53 |
| Pathogenic | 0.000 | 0.025 | 0.028 | 0.992 | -4.96 |
| Pathogenic | 0.000 | 0.000 | 0.019 | 0.982 | -5.12 |
| Benign | 1.202 | 1.227 | 1.186 | 0.665 | -2.71 |
| Benign | 1.317 | 1.364 | 1.230 | 0.355 | -1.03 |
| Pathogenic | 0.021 | 0.097 | 0.128 | 0.973 | -9.28 |
| Benign | 1.174 | 1.190 | 1.162 | 0.571 | -1.96 |
| Benign | 0.785 | 0.727 | 0.828 | 0.835 | -8.55 |
| Benign | 0.779 | 0.807 | 0.751 | 0.244 | -1.44 |
| Pathogenic | 0.000 | 0.000 | 0.000 | 0.994 | -4.64 |
| Pathogenic | 0.000 | 0.000 | 0.000 | 0.928 | -2.73 |
| Benign | 0.630 | 0.621 | 0.647 | 0.083 | 1.89 |
| Pathogenic | 0.000 | 0.026 | 0.000 | 0.971 | -5.56 |
| Pathogenic | 0.000 | 0.048 | 0.026 | 0.957 | -4.28 |
| Pathogenic | 0.800 | 0.800 | 0.853 | 0.938 | -5.53 |
| Pathogenic | 0.000 | 0.000 | 0.000 | 0.989 | -6.33 |
| Pathogenic | 0.647 | 0.647 | NA | 0.993 | -5.83 |
| Pathogenic | 0.848 | 0.879 | 0.779 | 0.949 | -2.61 |
| Pathogenic | 1.328 | 1.344 | 1.247 | 0.921 | -2.65 |
| Pathogenic | 0.032 | 0.165 | 0.029 | 0.972 | -6.07 |

| SIFT | Investigator(s) or Source |
|---|---|
| 0 | Sh. Wh. |
| 0 | Sh. Wh. |
| 0 | Ed.Fr. |
| 0 | Sh. Wh. |
| 0 | Ed.Fr.; JTF; Ca.Sc; Sh. Wh.; AKA |
| 0 | Ca.Sc |
| 0 | Ca.Sc |
| 0 | Sh. Wh.; Ed.Fr. |
| 0 | Sh. Wh.; AKA; Ed.Fr. |
| 0 | JTF; Sh. Wh.; Ca.Sc |
| 0 | JTF; Sh. Wh.; AKA |
| 0 | Sh. Wh. |
| 0 | Sh. Wh. |
| 0 | Sh. Wh. |
| 0.008 | Yl.Fl |
| 0.097 | ClinVar |
| 0.016 | C.R |
| 0 | E.Di.Pi.; Ca.Sc; Sh. Wh. |
| 0.001 | E.Di.Pi.; JTF; Ca.Sc; Sh. Wh. |
| 0.341 | Yl.Fl |
| 0.003 | gnomAD v2.1.1 |
| 0.266 | Sh. Wh. |
| 0.003 | Clinvar |
| 0 | Sh. Wh. |
| 0 | Yl.Fl |
| 0.001 | Sh. Wh. |
| 0.336 | RJ.D |
| 0 | Ca.Sc |
| 0 | Sh. Wh. |
| 0.223 | Yl.Fl |
| 0.001 | E.Di.Pi.; JTF; Ca.Sc; Sh. Wh. |
| 0 | Sh. Wh. |
| 0.001 | Sh. Wh.; Yl.Fl |
| 0 | Sh. Wh. |
| 0 | Ca.Sc |
| 0.001 | Yl.Fl |
| 0 | Sh. Wh. |
| 0.001 | Sh. Wh. |
| 0 | JTF; Sh. Wh. |
| 0 | Sh. Wh. |
| 0 | Sh. Wh. |
| 0 | JTF |
| 0 | Sh. Wh. |

| | |
|---|---|
| 0 | Ca.Sc |
| 0 | AKA |
| 0.01 | Yl.Fl |
| 0 | Sh. Wh. |
| 0 | Sh. Wh. |
| 0.003 | Yl.Fl |
| 0.269 | AKA |
| 0.399 | Yl.Fl |
| 0.001 | Yl.Fl |
| 0.186 | ClinVar |
| 0.001 | gnomAD v2.1.1 |
| 0.151 | RJ.D |
| 0 | Sh. Wh. |
| 0.001 | Ca.Sc |
| 1 | ClinVar |
| 0 | Ca.Sc |
| 0.005 | Sh. Wh. |
| 0 | Ca.Sc |
| 0 | Ca.Sc |
| 0 | AKA |
| 0.005 | Sh. Wh. |
| 0.002 | Ca.Sc |
| 0 | Sh. Wh. |

Table S7: Curated percentages of enzymatic activity expressed as a ratio between variant anc

| HGVS | wt aa | pos | mut aa | activity relative to wt (%) |
|---|---|---|---|---|
| p.Arg26Cys | Arg | 26 | Cys | 5 |
| p.Arg26His | Arg | 26 | His | 3 |
| p.Ser28Asn | Ser | 28 | Asn | 4 |
| p.Leu30Phe | Leu | 30 | Phe | 3 |
| p.Thr35Met | Thr | 35 | Met | 11 |
| p.Ser96Phe | Ser | 96 | Phe | 3 |
| p.Gly111Arg | Gly | 111 | Arg | 4 |
| p.Arg116Trp | Arg | 116 | Trp | 4 |
| p.Val124Asp | Val | 124 | Asp | 4 |
| p.Arg149Gln | Arg | 149 | Gln | 5 |
| p.Arg149Leu | Arg | 149 | Leu | 5 |
| p.Arg173Gln | Arg | 173 | Gln | 4 |
| p.Glu250Ala | Glu | 250 | Ala | 4 |
| p.Val267Met | Val | 267 | Met | 4 |
| p.Val215Glu | Val | 215 | Glu | 30 |
| p.Arg173Trp | Arg | 173 | Trp | 0.6 |
| p.Arg116Trp | Arg | 116 | Trp | 0.5 |
| p.Arg167Trp | Arg | 167 | Trp | 4.2 |
| p.Leu30Pro | Leu | 30 | Pro | 0.2 |
| p.Lys132Asn | Lys | 132 | Asn | 95 |
| p.Lys132Asn | Lys | 132 | Asn | 0.66 |
| p.Gly24Ser | Gly | 24 | Ser | 0.07 |
| p.Arg26Cys | Arg | 26 | Cys | 0 |
| p.Arg26His | Arg | 26 | His | 0 |
| p.Ser28Asn | Ser | 28 | Asn | 0.01 |
| p.Leu30Pro | Leu | 30 | Pro | 0 |
| p.Ala31Pro | Ala | 31 | Pro | 0.01 |
| p.Arg32Pro | Arg | 32 | Pro | 1 |
| p.Gln34Arg | Gln | 34 | Arg | 0.01 |
| p.Gln34Lys | Gln | 34 | Lys | 0 |
| p.Gln34Pro | Gln | 34 | Pro | 0.01 |
| p.Thr35Met | Thr | 35 | Met | 0.04 |
| p.Thr59Ile | Thr | 59 | Ile | 81 |
| p.Phe77Leu | Phe | 77 | Leu | 0 |
| p.Thr78Pro | Thr | 78 | Pro | 0.03 |
| p.Glu80Gly | Glu | 80 | Gly | 0.02 |
| p.Glu86Val | Glu | 86 | Val | 0.93 |
| p.Val93Phe | Val | 93 | Phe | 0.01 |
| p.Ser96Phe | Ser | 96 | Phe | 0.01 |
| p.Lys98Asn | Lys | 98 | Asn | 1 |
| p.Lys98Arg | Lys | 98 | Arg | 0.01 |
| p.Asp99Gly | Asp | 99 | Gly | 0.03 |
| p.Asp99His | Asp | 99 | His | 0.03 |

| | | | | |
|---|---|---|---|---|
| p.Ala112Pro | Ala | 112 | Pro | 0 |
| p.Arg116Trp | Arg | 116 | Trp | 0.01 |
| p.Ala122Asp | Ala | 122 | Asp | 0 |
| p.Lys132Asn | Lys | 132 | Asn | 0.97 |
| p.Thr145Ile | Thr | 145 | Ile | 0 |
| p.Ser147Pro | Ser | 147 | Pro | 0 |
| p.Arg149Leu | Arg | 149 | Leu | 0 |
| p.Arg149Gln | Arg | 149 | Gln | 0.01 |
| p.Arg149Leu | Arg | 149 | Leu | 0.05 |
| p.Leu154Pro | Leu | 154 | Pro | 0 |
| p.Arg167Gln | Arg | 167 | Gln | 0.01 |
| p.Arg167Trp | Arg | 167 | Trp | 0.02 |
| p.Leu170Pro | Leu | 170 | Pro | 1 |
| p.Arg173Gln | Arg | 173 | Gln | 0.01 |
| p.Arg173Gln | Arg | 173 | Gln | 0 |
| p.Asp178Asn | Asp | 178 | Asn | 0.81 |
| p.Arg195Cys | Arg | 195 | Cys | 0.03 |
| p.Arg195Cys | Arg | 195 | Cys | 0 |
| p.Met212Val | Met | 212 | Val | 0.02 |
| p.Val215Met | Val | 215 | Met | 0.19 |
| p.Gln217His | Gln | 217 | His | 0.05 |
| p.Gly218Arg | Gly | 218 | Arg | 0 |
| p.Val224Glu | Val | 224 | Glu | 0.01 |
| p.Val235Glu | Val | 235 | Glu | 0.03 |
| p.Asp240Gly | Asp | 240 | Gly | 0.05 |
| p.Cys247Arg | Cys | 247 | Arg | 0.02 |
| p.Cys247Phe | Cys | 247 | Phe | 0.11 |
| p.Glu250Asp | Glu | 250 | Asp | 0.01 |
| p.Glu250Gln | Glu | 250 | Gln | 0.02 |
| p.His256Tyr | His | 256 | Tyr | 0.05 |
| p.leu278Pro | leu | 278 | Pro | 1.22 |
| p.Arg321His | Arg | 321 | His | 0.96 |
| p.Gly335Ser | Gly | 335 | Ser | 0.03 |
| p.Ala347Pro | Ala | 347 | Pro | 51 |
| p.Asp359Asn | Asp | 359 | Asn | 0.93 |
| p.Met1Ile | Met | 1 | Ile | 1 |
| p.Arg167Gln | Arg | 167 | Gln | 1 |
| p.Arg167Trp | Arg | 167 | Trp | 3 |
| p.Arg195Cysys | Arg | 195 | Cysys | 3 |
| p.Ala252Val | Ala | 252 | Val | 61 |
| p.Ala331Val | Ala | 331 | Val | 62 |
| p.Thr59Ile | Thr | 59 | Ile | 77 |
| p.Asp230Tyr | Asp | 230 | Tyr | 88 |
| p.Asp359Asn | Asp | 359 | Asn | 86 |
| p.Glylu86Val | Glylu | 86 | Val | 94 |

| | | | | |
|---|---|---|---|---|
| p.Arg225Gln | Arg | 225 | Gln | 102 |
| p.Arg321His | Arg | 321 | His | 122 |
| p.Leu161Pro | Leu | 161 | Pro | 1 |
| p.Arg251Ser | Arg | 251 | Ser | 1 |
| p.Ala122Pro | Ala | 122 | Pro | 2 |
| p.Asn118Lys | Asn | 118 | Lys | 73 |
| p.Gly3Val | Gly | 3 | Val | 34 |
| p.Arg164Gly | Arg | 164 | Gly | 61 |
| p.Ile71Thr | Ile | 71 | Thr | 62 |
| p.Gly72Arg | Gly | 72 | Arg | 69 |
| p.Gly221Ser | Gly | 221 | Ser | 69 |
| p.Ile54Leu | Ile | 54 | Leu | 71 |
| p.Gly274Ala | Gly | 274 | Ala | 72 |
| p.Val282Leu | Val | 282 | Leu | 73 |
| p.Lys115Gln | Lys | 115 | Gln | 76 |
| p.Gly111Ala | Gly | 111 | Ala | 76 |
| p.Ala297Ser | Ala | 297 | Ser | 76 |
| p.Arg321Cys | Arg | 321 | Cys | 77 |
| p.Asp273Gly | Asp | 273 | Gly | 78 |
| p.Ala9Glylu | Ala | 9 | Glylu | 79 |
| p.Pro309Ser | Pro | 309 | Ser | 79 |
| p.Thr10Met | Thr | 10 | Met | 79 |
| p.Arg355Trp | Arg | 355 | Trp | 80 |
| p.Arg22Cys | Arg | 22 | Cys | 80 |
| p.Arg164Ser | Arg | 164 | Ser | 81 |
| p.Pro159Leu | Pro | 159 | Leu | 81 |
| p.Ile205Met | Ile | 205 | Met | 83 |
| p.Glu242Lys | Glu | 242 | Lys | 83 |
| p.Pro127Ala | Pro | 127 | Ala | 84 |
| p.Thr10Lys | Thr | 10 | Lys | 86 |
| p.Pro127Gln | Pro | 127 | Gln | 87 |
| p.Lys132Asn | Lys | 132 | Asn | 87 |
| p.Ser45Leu | Ser | 45 | Leu | 88 |
| p.Arg325Gln | Arg | 325 | Gln | 88 |
| p.Gln296Arg | Gln | 296 | Arg | 92 |
| p.Asp65His | Asp | 65 | His | 93 |
| p.His256Gln | His | 256 | Gln | 93 |
| p.His120Asn | His | 120 | Asn | 95 |
| p.Ser262Asn | Ser | 262 | Asn | 96 |
| p.Arg32Cysys | Arg | 32 | Cysys | 98 |
| p.Arg355Gln | Arg | 355 | Gln | 98 |
| p.Lys348Arg | Lys | 348 | Arg | 100 |
| p.Glu86Lys | Glu | 86 | Lys | 104 |
| p.Glu12Ala | Glu | 12 | Ala | 106 |
| p.Arg246His | Arg | 246 | His | 122 |

| | | | | |
|---|---|---|---|---|
| p.Val237Met | Val | 237 | Met | 128 |
| p.Arg246Cys | Arg | 246 | Cys | 130 |
| p.Gly3Ser | Gly | 3 | Ser | 211 |

d wild-type HMBS activity

**Sources and References**


Lenglet, H. et al. From a dominant to an oligogenic model of inheritance with environmental
Lenglet, H. et al. From a dominant to an oligogenic model of inheritance with environmental
Lenglet, H. et al. From a dominant to an oligogenic model of inheritance with environmental
Lenglet, H. et al. From a dominant to an oligogenic model of inheritance with environmental
Lenglet, H. et al. From a dominant to an oligogenic model of inheritance with environmental
Lenglet, H. et al. From a dominant to an oligogenic model of inheritance with environmental
Lenglet, H. et al. From a dominant to an oligogenic model of inheritance with environmental
Lenglet, H. et al. From a dominant to an oligogenic model of inheritance with environmental
Lenglet, H. et al. From a dominant to an oligogenic model of inheritance with environmental
Lenglet, H. et al. From a dominant to an oligogenic model of inheritance with environmental
Lenglet, H. et al. From a dominant to an oligogenic model of inheritance with environmental
Bung, N. et al. Human hydroxymethylbilane synthase: Molecular dynamics of the pyrrole chai
Lenglet, H. et al. From a dominant to an oligogenic model of inheritance with environmental
Lenglet, H. et al. From a dominant to an oligogenic model of inheritance with environmental
Bustad, H.J., et al. Conformational stability and activity analysis of two hydroxymethylbilane s
Bustad, H.J., et al. Conformational stability and activity analysis of two hydroxymethylbilane s
Bustad, H.J., et al. Conformational stability and activity analysis of two hydroxymethylbilane s
Bustad, H.J., et al. Conformational stability and activity analysis of two hydroxymethylbilane s
Chen, B. et al. Identification and characterization of 40 novel hydroxymethylbilane synthase r
Bustad, H.J., et al. Conformational stability and activity analysis of two hydroxymethylbilane s
Bustad, H.J., et al. Conformational stability and activity analysis of two hydroxymethylbilane s
Bustad, H.J. et al. Acute intermittent porphyria: an overview of therapy developments and fu
Bustad, H.J. et al. Acute intermittent porphyria: an overview of therapy developments and fu
Bustad, H.J. et al. Acute intermittent porphyria: an overview of therapy developments and fu
Bustad, H.J. et al. Acute intermittent porphyria: an overview of therapy developments and fu
Bustad, H.J. et al. Acute intermittent porphyria: an overview of therapy developments and fu
Bustad, H.J. et al. Acute intermittent porphyria: an overview of therapy developments and fu
Bustad, H.J. et al. Acute intermittent porphyria: an overview of therapy developments and fu
Bustad, H.J. et al. Acute intermittent porphyria: an overview of therapy developments and fu
Bustad, H.J. et al. Acute intermittent porphyria: an overview of therapy developments and fu
Bustad, H.J. et al. Acute intermittent porphyria: an overview of therapy developments and fu
Bustad, H.J. et al. Acute intermittent porphyria: an overview of therapy developments and fu
Bustad, H.J. et al. Acute intermittent porphyria: an overview of therapy developments and fu
Bustad, H.J. et al. Acute intermittent porphyria: an overview of therapy developments and fu
Bustad, H.J. et al. Acute intermittent porphyria: an overview of therapy developments and fu
Bustad, H.J. et al. Acute intermittent porphyria: an overview of therapy developments and fu
Bustad, H.J. et al. Acute intermittent porphyria: an overview of therapy developments and fu
Bustad, H.J. et al. Acute intermittent porphyria: an overview of therapy developments and fu
Bustad, H.J. et al. Acute intermittent porphyria: an overview of therapy developments and fu
Bustad, H.J. et al. Acute intermittent porphyria: an overview of therapy developments and fu
Bustad, H.J. et al. Acute intermittent porphyria: an overview of therapy developments and fu
Bustad, H.J. et al. Acute intermittent porphyria: an overview of therapy developments and fu
Bustad, H.J. et al. Acute intermittent porphyria: an overview of therapy developments and fu

Bustad, H.J. et al. Acute intermittent porphyria: an overview of therapy developments and fu
Bustad, H.J. et al. Acute intermittent porphyria: an overview of therapy developments and fu
Bustad, H.J. et al. Acute intermittent porphyria: an overview of therapy developments and fu
Bustad, H.J. et al. Acute intermittent porphyria: an overview of therapy developments and fu
Bustad, H.J. et al. Acute intermittent porphyria: an overview of therapy developments and fu
Bustad, H.J. et al. Acute intermittent porphyria: an overview of therapy developments and fu
Bustad, H.J. et al. Acute intermittent porphyria: an overview of therapy developments and fu
Bustad, H.J. et al. Acute intermittent porphyria: an overview of therapy developments and fu
Bustad, H.J. et al. Acute intermittent porphyria: an overview of therapy developments and fu
Bustad, H.J. et al. Acute intermittent porphyria: an overview of therapy developments and fu
Bustad, H.J. et al. Acute intermittent porphyria: an overview of therapy developments and fu
Bustad, H.J. et al. Acute intermittent porphyria: an overview of therapy developments and fu
Bustad, H.J. et al. Acute intermittent porphyria: an overview of therapy developments and fu
Bustad, H.J. et al. Acute intermittent porphyria: an overview of therapy developments and fu
Bustad, H.J. et al. Acute intermittent porphyria: an overview of therapy developments and fu
Bustad, H.J. et al. Acute intermittent porphyria: an overview of therapy developments and fu
Bustad, H.J. et al. Acute intermittent porphyria: an overview of therapy developments and fu
Bustad, H.J. et al. Acute intermittent porphyria: an overview of therapy developments and fu
Bustad, H.J. et al. Acute intermittent porphyria: an overview of therapy developments and fu
Bustad, H.J. et al. Acute intermittent porphyria: an overview of therapy developments and fu
Bustad, H.J. et al. Acute intermittent porphyria: an overview of therapy developments and fu
Bustad, H.J. et al. Acute intermittent porphyria: an overview of therapy developments and fu
Bustad, H.J. et al. Acute intermittent porphyria: an overview of therapy developments and fu
Bustad, H.J. et al. Acute intermittent porphyria: an overview of therapy developments and fu
Bustad, H.J. et al. Acute intermittent porphyria: an overview of therapy developments and fu
Bustad, H.J. et al. Acute intermittent porphyria: an overview of therapy developments and fu
Bustad, H.J. et al. Acute intermittent porphyria: an overview of therapy developments and fu
Bustad, H.J. et al. Acute intermittent porphyria: an overview of therapy developments and fu
Bustad, H.J. et al. Acute intermittent porphyria: an overview of therapy developments and fu
Bustad, H.J. et al. Acute intermittent porphyria: an overview of therapy developments and fu
Bustad, H.J. et al. Acute intermittent porphyria: an overview of therapy developments and fu
Bustad, H.J. et al. Acute intermittent porphyria: an overview of therapy developments and fu
Bustad, H.J. et al. Acute intermittent porphyria: an overview of therapy developments and fu
Bustad, H.J. et al. Acute intermittent porphyria: an overview of therapy developments and fu
Bustad, H.J. et al. Acute intermittent porphyria: an overview of therapy developments and fu
Chen, B. et al. Acute Intermittent Porphyria: Predicted Pathogenicity of HMBS Variants Indica
Chen, B. et al. Acute Intermittent Porphyria: Predicted Pathogenicity of HMBS Variants Indica
Chen, B. et al. Acute Intermittent Porphyria: Predicted Pathogenicity of HMBS Variants Indica
Chen, B. et al. Acute Intermittent Porphyria: Predicted Pathogenicity of HMBS Variants Indica
Chen, B. et al. Acute Intermittent Porphyria: Predicted Pathogenicity of HMBS Variants Indica
Chen, B. et al. Acute Intermittent Porphyria: Predicted Pathogenicity of HMBS Variants Indica
Chen, B. et al. Acute Intermittent Porphyria: Predicted Pathogenicity of HMBS Variants Indica
Chen, B. et al. Acute Intermittent Porphyria: Predicted Pathogenicity of HMBS Variants Indica
Chen, B. et al. Acute Intermittent Porphyria: Predicted Pathogenicity of HMBS Variants Indica
Chen, B. et al. Acute Intermittent Porphyria: Predicted Pathogenicity of HMBS Variants Indica

Chen, B. et al. Acute Intermittent Porphyria: Predicted Pathogenicity of HMBS Variants Indica
Chen, B. et al. Acute Intermittent Porphyria: Predicted Pathogenicity of HMBS Variants Indica
Chen, B. et al. Acute Intermittent Porphyria: Predicted Pathogenicity of HMBS Variants Indica
Chen, B. et al. Acute Intermittent Porphyria: Predicted Pathogenicity of HMBS Variants Indica
Chen, B. et al. Acute Intermittent Porphyria: Predicted Pathogenicity of HMBS Variants Indica
Chen, B. et al. Acute Intermittent Porphyria: Predicted Pathogenicity of HMBS Variants Indica
Chen, B. et al. Acute Intermittent Porphyria: Predicted Pathogenicity of HMBS Variants Indica
Chen, B. et al. Acute Intermittent Porphyria: Predicted Pathogenicity of HMBS Variants Indica
Chen, B. et al. Acute Intermittent Porphyria: Predicted Pathogenicity of HMBS Variants Indica
Chen, B. et al. Acute Intermittent Porphyria: Predicted Pathogenicity of HMBS Variants Indica
Chen, B. et al. Acute Intermittent Porphyria: Predicted Pathogenicity of HMBS Variants Indica
Chen, B. et al. Acute Intermittent Porphyria: Predicted Pathogenicity of HMBS Variants Indica
Chen, B. et al. Acute Intermittent Porphyria: Predicted Pathogenicity of HMBS Variants Indica
Chen, B. et al. Acute Intermittent Porphyria: Predicted Pathogenicity of HMBS Variants Indica
Chen, B. et al. Acute Intermittent Porphyria: Predicted Pathogenicity of HMBS Variants Indica
Chen, B. et al. Acute Intermittent Porphyria: Predicted Pathogenicity of HMBS Variants Indica
Chen, B. et al. Acute Intermittent Porphyria: Predicted Pathogenicity of HMBS Variants Indica
Chen, B. et al. Acute Intermittent Porphyria: Predicted Pathogenicity of HMBS Variants Indica
Chen, B. et al. Acute Intermittent Porphyria: Predicted Pathogenicity of HMBS Variants Indica
Chen, B. et al. Acute Intermittent Porphyria: Predicted Pathogenicity of HMBS Variants Indica
Chen, B. et al. Acute Intermittent Porphyria: Predicted Pathogenicity of HMBS Variants Indica
Chen, B. et al. Acute Intermittent Porphyria: Predicted Pathogenicity of HMBS Variants Indica
Chen, B. et al. Acute Intermittent Porphyria: Predicted Pathogenicity of HMBS Variants Indica
Chen, B. et al. Acute Intermittent Porphyria: Predicted Pathogenicity of HMBS Variants Indica
Chen, B. et al. Acute Intermittent Porphyria: Predicted Pathogenicity of HMBS Variants Indica
Chen, B. et al. Acute Intermittent Porphyria: Predicted Pathogenicity of HMBS Variants Indica
Chen, B. et al. Acute Intermittent Porphyria: Predicted Pathogenicity of HMBS Variants Indica
Chen, B. et al. Acute Intermittent Porphyria: Predicted Pathogenicity of HMBS Variants Indica
Chen, B. et al. Acute Intermittent Porphyria: Predicted Pathogenicity of HMBS Variants Indica
Chen, B. et al. Acute Intermittent Porphyria: Predicted Pathogenicity of HMBS Variants Indica
Chen, B. et al. Acute Intermittent Porphyria: Predicted Pathogenicity of HMBS Variants Indica
Chen, B. et al. Acute Intermittent Porphyria: Predicted Pathogenicity of HMBS Variants Indica
Chen, B. et al. Acute Intermittent Porphyria: Predicted Pathogenicity of HMBS Variants Indica
Chen, B. et al. Acute Intermittent Porphyria: Predicted Pathogenicity of HMBS Variants Indica
Chen, B. et al. Acute Intermittent Porphyria: Predicted Pathogenicity of HMBS Variants Indica
Chen, B. et al. Acute Intermittent Porphyria: Predicted Pathogenicity of HMBS Variants Indica
Chen, B. et al. Acute Intermittent Porphyria: Predicted Pathogenicity of HMBS Variants Indica
Chen, B. et al. Acute Intermittent Porphyria: Predicted Pathogenicity of HMBS Variants Indica
Chen, B. et al. Acute Intermittent Porphyria: Predicted Pathogenicity of HMBS Variants Indica
Chen, B. et al. Acute Intermittent Porphyria: Predicted Pathogenicity of HMBS Variants Indica
Chen, B. et al. Acute Intermittent Porphyria: Predicted Pathogenicity of HMBS Variants Indica
Chen, B. et al. Acute Intermittent Porphyria: Predicted Pathogenicity of HMBS Variants Indica
Chen, B. et al. Acute Intermittent Porphyria: Predicted Pathogenicity of HMBS Variants Indica
Chen, B. et al. Acute Intermittent Porphyria: Predicted Pathogenicity of HMBS Variants Indica
Chen, B. et al. Acute Intermittent Porphyria: Predicted Pathogenicity of HMBS Variants Indica

Chen, B. et al. Acute Intermittent Porphyria: Predicted Pathogenicity of HMBS Variants Indica
Chen, B. et al. Acute Intermittent Porphyria: Predicted Pathogenicity of HMBS Variants Indica
Chen, B. et al. Acute Intermittent Porphyria: Predicted Pathogenicity of HMBS Variants Indica

modifiers in acute intermittent porphyria. Hum. Mol. Genet. 27, 1164-1173 (2018)
modifiers in acute intermittent porphyria. Hum. Mol. Genet. 27, 1164-1173 (2018)
modifiers in acute intermittent porphyria. Hum. Mol. Genet. 27, 1164-1173 (2018)
modifiers in acute intermittent porphyria. Hum. Mol. Genet. 27, 1164-1173 (2018)
modifiers in acute intermittent porphyria. Hum. Mol. Genet. 27, 1164-1173 (2018)
modifiers in acute intermittent porphyria. Hum. Mol. Genet. 27, 1164-1173 (2018)
modifiers in acute intermittent porphyria. Hum. Mol. Genet. 27, 1164-1173 (2018)
modifiers in acute intermittent porphyria. Hum. Mol. Genet. 27, 1164-1173 (2018)
modifiers in acute intermittent porphyria. Hum. Mol. Genet. 27, 1164-1173 (2018)
modifiers in acute intermittent porphyria. Hum. Mol. Genet. 27, 1164-1173 (2018)
modifiers in acute intermittent porphyria. Hum. Mol. Genet. 27, 1164-1173 (2018)
n elongation identifies step-specific residues that cause AIP. Proc. Natl. Acad. Sci. U.S.A. 115,
modifiers in acute intermittent porphyria. Hum. Mol. Genet. 27, 1164-1173 (2018)
modifiers in acute intermittent porphyria. Hum. Mol. Genet. 27, 1164-1173 (2018)
synthase mutants, K132N and V215E, with different phenotypic association with acute interm
synthase mutants, K132N and V215E, with different phenotypic association with acute interm
synthase mutants, K132N and V215E, with different phenotypic association with acute interm
synthase mutants, K132N and V215E, with different phenotypic association with acute interm
nutations that cause acute intermittent porphyria. J. Inherit. Metab. Dis. 42, 186,Äì194 (2019)
synthase mutants, K132N and V215E, with different phenotypic association with acute interm
synthase mutants, K132N and V215E, with different phenotypic association with acute interm
ture perspectives focusing on stabilisation of HMBS and proteostasis regulators. Int. J. Mol. 2
ture perspectives focusing on stabilisation of HMBS and proteostasis regulators. Int. J. Mol. 2
ture perspectives focusing on stabilisation of HMBS and proteostasis regulators. Int. J. Mol. 2
ture perspectives focusing on stabilisation of HMBS and proteostasis regulators. Int. J. Mol. 2
ture perspectives focusing on stabilisation of HMBS and proteostasis regulators. Int. J. Mol. 2
ture perspectives focusing on stabilisation of HMBS and proteostasis regulators. Int. J. Mol. 2
ture perspectives focusing on stabilisation of HMBS and proteostasis regulators. Int. J. Mol. 2
ture perspectives focusing on stabilisation of HMBS and proteostasis regulators. Int. J. Mol. 2
ture perspectives focusing on stabilisation of HMBS and proteostasis regulators. Int. J. Mol. 2
ture perspectives focusing on stabilisation of HMBS and proteostasis regulators. Int. J. Mol. 2
ture perspectives focusing on stabilisation of HMBS and proteostasis regulators. Int. J. Mol. 2
ture perspectives focusing on stabilisation of HMBS and proteostasis regulators. Int. J. Mol. 2
ture perspectives focusing on stabilisation of HMBS and proteostasis regulators. Int. J. Mol. 2
ture perspectives focusing on stabilisation of HMBS and proteostasis regulators. Int. J. Mol. 2
ture perspectives focusing on stabilisation of HMBS and proteostasis regulators. Int. J. Mol. 2
ture perspectives focusing on stabilisation of HMBS and proteostasis regulators. Int. J. Mol. 2
ture perspectives focusing on stabilisation of HMBS and proteostasis regulators. Int. J. Mol. 2
ture perspectives focusing on stabilisation of HMBS and proteostasis regulators. Int. J. Mol. 2
ture perspectives focusing on stabilisation of HMBS and proteostasis regulators. Int. J. Mol. 2
ture perspectives focusing on stabilisation of HMBS and proteostasis regulators. Int. J. Mol. 2
ture perspectives focusing on stabilisation of HMBS and proteostasis regulators. Int. J. Mol. 2
ture perspectives focusing on stabilisation of HMBS and proteostasis regulators. Int. J. Mol. 2

ture perspectives focusing on stabilisation of HMBS and proteostasis regulators. Int. J. Mol. 2
ture perspectives focusing on stabilisation of HMBS and proteostasis regulators. Int. J. Mol. 2
ture perspectives focusing on stabilisation of HMBS and proteostasis regulators. Int. J. Mol. 2
ture perspectives focusing on stabilisation of HMBS and proteostasis regulators. Int. J. Mol. 2
ture perspectives focusing on stabilisation of HMBS and proteostasis regulators. Int. J. Mol. 2
ture perspectives focusing on stabilisation of HMBS and proteostasis regulators. Int. J. Mol. 2
ture perspectives focusing on stabilisation of HMBS and proteostasis regulators. Int. J. Mol. 2
ture perspectives focusing on stabilisation of HMBS and proteostasis regulators. Int. J. Mol. 2
ture perspectives focusing on stabilisation of HMBS and proteostasis regulators. Int. J. Mol. 2
ture perspectives focusing on stabilisation of HMBS and proteostasis regulators. Int. J. Mol. 2
ture perspectives focusing on stabilisation of HMBS and proteostasis regulators. Int. J. Mol. 2
ture perspectives focusing on stabilisation of HMBS and proteostasis regulators. Int. J. Mol. 2
ture perspectives focusing on stabilisation of HMBS and proteostasis regulators. Int. J. Mol. 2
ture perspectives focusing on stabilisation of HMBS and proteostasis regulators. Int. J. Mol. 2
ture perspectives focusing on stabilisation of HMBS and proteostasis regulators. Int. J. Mol. 2
ture perspectives focusing on stabilisation of HMBS and proteostasis regulators. Int. J. Mol. 2
ture perspectives focusing on stabilisation of HMBS and proteostasis regulators. Int. J. Mol. 2
ture perspectives focusing on stabilisation of HMBS and proteostasis regulators. Int. J. Mol. 2
ture perspectives focusing on stabilisation of HMBS and proteostasis regulators. Int. J. Mol. 2
ture perspectives focusing on stabilisation of HMBS and proteostasis regulators. Int. J. Mol. 2
ture perspectives focusing on stabilisation of HMBS and proteostasis regulators. Int. J. Mol. 2
ture perspectives focusing on stabilisation of HMBS and proteostasis regulators. Int. J. Mol. 2
ture perspectives focusing on stabilisation of HMBS and proteostasis regulators. Int. J. Mol. 2
ture perspectives focusing on stabilisation of HMBS and proteostasis regulators. Int. J. Mol. 2
ture perspectives focusing on stabilisation of HMBS and proteostasis regulators. Int. J. Mol. 2
ture perspectives focusing on stabilisation of HMBS and proteostasis regulators. Int. J. Mol. 2
ture perspectives focusing on stabilisation of HMBS and proteostasis regulators. Int. J. Mol. 2
ture perspectives focusing on stabilisation of HMBS and proteostasis regulators. Int. J. Mol. 2
ture perspectives focusing on stabilisation of HMBS and proteostasis regulators. Int. J. Mol. 2
ture perspectives focusing on stabilisation of HMBS and proteostasis regulators. Int. J. Mol. 2
ture perspectives focusing on stabilisation of HMBS and proteostasis regulators. Int. J. Mol. 2
ture perspectives focusing on stabilisation of HMBS and proteostasis regulators. Int. J. Mol. 2
ture perspectives focusing on stabilisation of HMBS and proteostasis regulators. Int. J. Mol. 2
ture perspectives focusing on stabilisation of HMBS and proteostasis regulators. Int. J. Mol. 2
ture perspectives focusing on stabilisation of HMBS and proteostasis regulators. Int. J. Mol. 2
tes Extremely Low Penetrance of the Autosomal Dominant Disease. Hum. Mutat. 37, 1215,Äì
tes Extremely Low Penetrance of the Autosomal Dominant Disease. Hum. Mutat. 37, 1215,Äì
tes Extremely Low Penetrance of the Autosomal Dominant Disease. Hum. Mutat. 37, 1215,Äì
tes Extremely Low Penetrance of the Autosomal Dominant Disease. Hum. Mutat. 37, 1215,Äì
tes Extremely Low Penetrance of the Autosomal Dominant Disease. Hum. Mutat. 37, 1215,Äì
tes Extremely Low Penetrance of the Autosomal Dominant Disease. Hum. Mutat. 37, 1215,Äì
tes Extremely Low Penetrance of the Autosomal Dominant Disease. Hum. Mutat. 37, 1215,Äì
tes Extremely Low Penetrance of the Autosomal Dominant Disease. Hum. Mutat. 37, 1215,Äì
tes Extremely Low Penetrance of the Autosomal Dominant Disease. Hum. Mutat. 37, 1215,Äì
tes Extremely Low Penetrance of the Autosomal Dominant Disease. Hum. Mutat. 37, 1215,Äì

tes Extremely Low Penetrance of the Autosomal Dominant Disease. Hum. Mutat. 37, 1215,Äì
tes Extremely Low Penetrance of the Autosomal Dominant Disease. Hum. Mutat. 37, 1215,Äì
tes Extremely Low Penetrance of the Autosomal Dominant Disease. Hum. Mutat. 37, 1215,Äì
tes Extremely Low Penetrance of the Autosomal Dominant Disease. Hum. Mutat. 37, 1215,Äì
tes Extremely Low Penetrance of the Autosomal Dominant Disease. Hum. Mutat. 37, 1215,Äì
tes Extremely Low Penetrance of the Autosomal Dominant Disease. Hum. Mutat. 37, 1215,Äì
tes Extremely Low Penetrance of the Autosomal Dominant Disease. Hum. Mutat. 37, 1215,Äì
tes Extremely Low Penetrance of the Autosomal Dominant Disease. Hum. Mutat. 37, 1215,Äì
tes Extremely Low Penetrance of the Autosomal Dominant Disease. Hum. Mutat. 37, 1215,Äì
tes Extremely Low Penetrance of the Autosomal Dominant Disease. Hum. Mutat. 37, 1215,Äì
tes Extremely Low Penetrance of the Autosomal Dominant Disease. Hum. Mutat. 37, 1215,Äì
tes Extremely Low Penetrance of the Autosomal Dominant Disease. Hum. Mutat. 37, 1215,Äì
tes Extremely Low Penetrance of the Autosomal Dominant Disease. Hum. Mutat. 37, 1215,Äì
tes Extremely Low Penetrance of the Autosomal Dominant Disease. Hum. Mutat. 37, 1215,Äì
tes Extremely Low Penetrance of the Autosomal Dominant Disease. Hum. Mutat. 37, 1215,Äì
tes Extremely Low Penetrance of the Autosomal Dominant Disease. Hum. Mutat. 37, 1215,Äì
tes Extremely Low Penetrance of the Autosomal Dominant Disease. Hum. Mutat. 37, 1215,Äì
tes Extremely Low Penetrance of the Autosomal Dominant Disease. Hum. Mutat. 37, 1215,Äì
tes Extremely Low Penetrance of the Autosomal Dominant Disease. Hum. Mutat. 37, 1215,Äì
tes Extremely Low Penetrance of the Autosomal Dominant Disease. Hum. Mutat. 37, 1215,Äì
tes Extremely Low Penetrance of the Autosomal Dominant Disease. Hum. Mutat. 37, 1215,Äì
tes Extremely Low Penetrance of the Autosomal Dominant Disease. Hum. Mutat. 37, 1215,Äì
tes Extremely Low Penetrance of the Autosomal Dominant Disease. Hum. Mutat. 37, 1215,Äì
tes Extremely Low Penetrance of the Autosomal Dominant Disease. Hum. Mutat. 37, 1215,Äì
tes Extremely Low Penetrance of the Autosomal Dominant Disease. Hum. Mutat. 37, 1215,Äì
tes Extremely Low Penetrance of the Autosomal Dominant Disease. Hum. Mutat. 37, 1215,Äì
tes Extremely Low Penetrance of the Autosomal Dominant Disease. Hum. Mutat. 37, 1215,Äì
tes Extremely Low Penetrance of the Autosomal Dominant Disease. Hum. Mutat. 37, 1215,Äì
tes Extremely Low Penetrance of the Autosomal Dominant Disease. Hum. Mutat. 37, 1215,Äì
tes Extremely Low Penetrance of the Autosomal Dominant Disease. Hum. Mutat. 37, 1215,Äì
tes Extremely Low Penetrance of the Autosomal Dominant Disease. Hum. Mutat. 37, 1215,Äì
tes Extremely Low Penetrance of the Autosomal Dominant Disease. Hum. Mutat. 37, 1215,Äì
tes Extremely Low Penetrance of the Autosomal Dominant Disease. Hum. Mutat. 37, 1215,Äì
tes Extremely Low Penetrance of the Autosomal Dominant Disease. Hum. Mutat. 37, 1215,Äì
tes Extremely Low Penetrance of the Autosomal Dominant Disease. Hum. Mutat. 37, 1215,Äì
tes Extremely Low Penetrance of the Autosomal Dominant Disease. Hum. Mutat. 37, 1215,Äì
tes Extremely Low Penetrance of the Autosomal Dominant Disease. Hum. Mutat. 37, 1215,Äì
tes Extremely Low Penetrance of the Autosomal Dominant Disease. Hum. Mutat. 37, 1215,Äì
tes Extremely Low Penetrance of the Autosomal Dominant Disease. Hum. Mutat. 37, 1215,Äì
tes Extremely Low Penetrance of the Autosomal Dominant Disease. Hum. Mutat. 37, 1215,Äì
tes Extremely Low Penetrance of the Autosomal Dominant Disease. Hum. Mutat. 37, 1215,Äì
tes Extremely Low Penetrance of the Autosomal Dominant Disease. Hum. Mutat. 37, 1215,Äì
tes Extremely Low Penetrance of the Autosomal Dominant Disease. Hum. Mutat. 37, 1215,Äì
tes Extremely Low Penetrance of the Autosomal Dominant Disease. Hum. Mutat. 37, 1215,Äì
tes Extremely Low Penetrance of the Autosomal Dominant Disease. Hum. Mutat. 37, 1215,Äì

tes Extremely Low Penetrance of the Autosomal Dominant Disease. Hum. Mutat. 37, 1215,Äì
tes Extremely Low Penetrance of the Autosomal Dominant Disease. Hum. Mutat. 37, 1215,Äì
tes Extremely Low Penetrance of the Autosomal Dominant Disease. Hum. Mutat. 37, 1215,Äì

, E4071,ÄìE4080 (2018)

ıittent porphyria. Bioscience reports. 33 (2013)
ıittent porphyria. Bioscience reports. 33 (2013)
ıittent porphyria. Bioscience reports. 33 (2013)
ıittent porphyria. Bioscience reports. 33 (2013)
)
ıittent porphyria. Bioscience reports. 33 (2013)
ıittent porphyria. Bioscience reports. 33 (2013)
2, 675 (2021)
2, 675 (2021)
2, 675 (2021)
2, 675 (2021)
2, 675 (2021)
2, 675 (2021)
2, 675 (2021)
2, 675 (2021)
2, 675 (2021)
2, 675 (2021)
2, 675 (2021)
2, 675 (2021)
2, 675 (2021)
2, 675 (2021)
2, 675 (2021)
2, 675 (2021)
2, 675 (2021)
2, 675 (2021)
2, 675 (2021)
2, 675 (2021)
2, 675 (2021)
2, 675 (2021)

2, 675 (2021)
2, 675 (2021)
2, 675 (2021)
2, 675 (2021)
2, 675 (2021)
2, 675 (2021)
2, 675 (2021)
2, 675 (2021)
2, 675 (2021)
2, 675 (2021)
2, 675 (2021)
2, 675 (2021)
2, 675 (2021)
2, 675 (2021)
2, 675 (2021)
2, 675 (2021)
2, 675 (2021)
2, 675 (2021)
2, 675 (2021)
2, 675 (2021)
2, 675 (2021)
2, 675 (2021)
2, 675 (2021)
2, 675 (2021)
2, 675 (2021)
2, 675 (2021)
2, 675 (2021)
2, 675 (2021)
2, 675 (2021)
2, 675 (2021)
2, 675 (2021)
2, 675 (2021)
2, 675 (2021)
2, 675 (2021)
2, 675 (2021)
1222 (2016)
1222 (2016)
1222 (2016)
1222 (2016)
1222 (2016)
1222 (2016)
1222 (2016)
1222 (2016)
1222 (2016)
1222 (2016)

1222 (2016)
1222 (2016)
1222 (2016)
1222 (2016)
1222 (2016)
1222 (2016)
1222 (2016)
1222 (2016)
1222 (2016)
1222 (2016)
1222 (2016)
1222 (2016)
1222 (2016)
1222 (2016)
1222 (2016)
1222 (2016)
1222 (2016)
1222 (2016)
1222 (2016)
1222 (2016)
1222 (2016)
1222 (2016)
1222 (2016)
1222 (2016)
1222 (2016)
1222 (2016)
1222 (2016)
1222 (2016)
1222 (2016)
1222 (2016)
1222 (2016)
1222 (2016)
1222 (2016)
1222 (2016)
1222 (2016)
1222 (2016)
1222 (2016)
1222 (2016)
1222 (2016)
1222 (2016)
1222 (2016)
1222 (2016)
1222 (2016)
1222 (2016)
1222 (2016)

1222 (2016)
1222 (2016)
1222 (2016)
