## Supplemental Table S8 for "Systematically testing human HMBS missense variants to reveal mechanism and pathogenic variation"

Table S8: Log likelihood ratios of pathogenicity derived from the combine HMBS map.

| Variant | | | | | Combined Map | |
|---|---|---|---|---|---|---|
| **HGVS** | **type** | **pos** | **wt aa** | **mut aa** | **score** | **stderr** |
| p.Ala112= | synonymous | 112 | Ala | NA | 1.015 | 0.003 |
| p.Ala112Arg | substitution | 112 | Ala | Arg | 0.000 | 0.151 |
| p.Ala112Asn | substitution | 112 | Ala | Asn | 0.000 | 0.542 |
| p.Ala112Asp | substitution | 112 | Ala | Asp | 0.122 | 0.048 |
| p.Ala112Cys | substitution | 112 | Ala | Cys | 1.107 | 0.020 |
| p.Ala112Gln | substitution | 112 | Ala | Gln | 0.548 | 0.021 |
| p.Ala112Glu | substitution | 112 | Ala | Glu | 0.000 | 0.077 |
| p.Ala112Gly | substitution | 112 | Ala | Gly | 0.597 | 0.035 |
| p.Ala112His | substitution | 112 | Ala | His | 0.000 | 0.030 |
| p.Ala112Ile | substitution | 112 | Ala | Ile | 0.000 | 0.024 |
| p.Ala112Leu | substitution | 112 | Ala | Leu | 0.598 | 0.064 |
| p.Ala112Lys | substitution | 112 | Ala | Lys | 0.000 | 0.040 |
| p.Ala112Met | substitution | 112 | Ala | Met | 0.000 | 0.027 |
| p.Ala112Phe | substitution | 112 | Ala | Phe | 0.000 | 0.028 |
| p.Ala112Pro | substitution | 112 | Ala | Pro | 0.319 | 0.058 |
| p.Ala112Ser | substitution | 112 | Ala | Ser | 0.737 | 0.028 |
| p.Ala112Ter | nonsense | 112 | Ala | Ter | 0.318 | 0.018 |
| p.Ala112Thr | substitution | 112 | Ala | Thr | 0.760 | 0.020 |
| p.Ala112Trp | substitution | 112 | Ala | Trp | 0.000 | 0.034 |
| p.Ala112Tyr | substitution | 112 | Ala | Tyr | 0.227 | 0.032 |
| p.Ala112Val | substitution | 112 | Ala | Val | 0.508 | 0.057 |
| p.Ala122= | synonymous | 122 | Ala | NA | 1.115 | 0.022 |
| p.Ala122Arg | substitution | 122 | Ala | Arg | 0.000 | 0.027 |
| p.Ala122Asn | substitution | 122 | Ala | Asn | 0.875 | 0.015 |
| p.Ala122Asp | substitution | 122 | Ala | Asp | 0.306 | 0.009 |
| p.Ala122Gln | substitution | 122 | Ala | Gln | 0.396 | 0.020 |
| p.Ala122Glu | substitution | 122 | Ala | Glu | 0.277 | 0.024 |
| p.Ala122Gly | substitution | 122 | Ala | Gly | 0.854 | 0.008 |
| p.Ala122His | substitution | 122 | Ala | His | 0.205 | 0.012 |
| p.Ala122Ile | substitution | 122 | Ala | Ile | 0.106 | 0.031 |
| p.Ala122Leu | substitution | 122 | Ala | Leu | 0.254 | 0.036 |
| p.Ala122Lys | substitution | 122 | Ala | Lys | 0.109 | 0.034 |
| p.Ala122Met | substitution | 122 | Ala | Met | 0.609 | 0.022 |
| p.Ala122Phe | substitution | 122 | Ala | Phe | 0.059 | 0.029 |
| p.Ala122Pro | substitution | 122 | Ala | Pro | 0.231 | 0.029 |
| p.Ala122Ser | substitution | 122 | Ala | Ser | 0.783 | 0.060 |
| p.Ala122Ter | nonsense | 122 | Ala | Ter | 0.000 | 0.024 |
| p.Ala122Thr | substitution | 122 | Ala | Thr | 0.800 | 0.011 |
| p.Ala122Trp | substitution | 122 | Ala | Trp | 0.000 | 0.021 |
| p.Ala122Tyr | substitution | 122 | Ala | Tyr | 0.083 | 0.015 |

| | | | | | | | |
|---|---|---|---|---|---|---|---|
| p.Ala122Val | substitution | 122 | Ala | Val | | 0.997 | 0.003 |
| p.Ala151= | synonymous | 151 | Ala | NA | | 0.795 | 0.008 |
| p.Ala151Arg | substitution | 151 | Ala | Arg | | 0.898 | 0.008 |
| p.Ala151Asn | substitution | 151 | Ala | Asn | | 0.946 | 0.011 |
| p.Ala151Asp | substitution | 151 | Ala | Asp | | 0.947 | 0.009 |
| p.Ala151Cys | substitution | 151 | Ala | Cys | | 0.849 | 0.008 |
| p.Ala151Gln | substitution | 151 | Ala | Gln | | 1.170 | 0.014 |
| p.Ala151Glu | substitution | 151 | Ala | Glu | | 0.000 | 0.067 |
| p.Ala151Gly | substitution | 151 | Ala | Gly | | 0.900 | 0.009 |
| p.Ala151His | substitution | 151 | Ala | His | | 0.880 | 0.007 |
| p.Ala151Ile | substitution | 151 | Ala | Ile | | 0.358 | 0.131 |
| p.Ala151Leu | substitution | 151 | Ala | Leu | | 0.907 | 0.022 |
| p.Ala151Lys | substitution | 151 | Ala | Lys | | 0.642 | 0.010 |
| p.Ala151Met | substitution | 151 | Ala | Met | | 0.113 | 0.033 |
| p.Ala151Phe | substitution | 151 | Ala | Phe | | 0.773 | 0.023 |
| p.Ala151Pro | substitution | 151 | Ala | Pro | | 0.180 | 0.093 |
| p.Ala151Ser | substitution | 151 | Ala | Ser | | 0.785 | 0.061 |
| p.Ala151Ter | nonsense | 151 | Ala | Ter | | 0.000 | 0.099 |
| p.Ala151Thr | substitution | 151 | Ala | Thr | | 0.658 | 0.019 |
| p.Ala151Trp | substitution | 151 | Ala | Trp | | 0.884 | 0.009 |
| p.Ala151Tyr | substitution | 151 | Ala | Tyr | | 0.728 | 0.008 |
| p.Ala151Val | substitution | 151 | Ala | Val | | 0.849 | 0.004 |
| p.Ala152= | synonymous | 152 | Ala | NA | | 0.907 | 0.015 |
| p.Ala152Arg | substitution | 152 | Ala | Arg | | 0.903 | 0.029 |
| p.Ala152Asn | substitution | 152 | Ala | Asn | | 0.527 | 0.010 |
| p.Ala152Asp | substitution | 152 | Ala | Asp | | 0.193 | 1.069 |
| p.Ala152Cys | substitution | 152 | Ala | Cys | | 0.811 | 0.028 |
| p.Ala152Gln | substitution | 152 | Ala | Gln | | 0.219 | 0.012 |
| p.Ala152Glu | substitution | 152 | Ala | Glu | | 0.297 | 0.015 |
| p.Ala152Gly | substitution | 152 | Ala | Gly | | 0.774 | 0.021 |
| p.Ala152His | substitution | 152 | Ala | His | | 1.161 | 0.013 |
| p.Ala152Ile | substitution | 152 | Ala | Ile | | 0.969 | 0.011 |
| p.Ala152Leu | substitution | 152 | Ala | Leu | | 0.996 | 0.016 |
| p.Ala152Lys | substitution | 152 | Ala | Lys | | 0.598 | 0.043 |
| p.Ala152Met | substitution | 152 | Ala | Met | | 0.795 | 0.014 |
| p.Ala152Phe | substitution | 152 | Ala | Phe | | 0.155 | 0.025 |
| p.Ala152Pro | substitution | 152 | Ala | Pro | | 0.527 | 0.045 |
| p.Ala152Ser | substitution | 152 | Ala | Ser | | 0.523 | 0.089 |
| p.Ala152Thr | substitution | 152 | Ala | Thr | | 1.055 | 0.033 |
| p.Ala152Trp | substitution | 152 | Ala | Trp | | 0.801 | 0.006 |
| p.Ala152Tyr | substitution | 152 | Ala | Tyr | | 0.708 | 0.026 |
| p.Ala152Val | substitution | 152 | Ala | Val | | 1.054 | 0.030 |
| p.Ala185= | synonymous | 185 | Ala | NA | | 1.357 | 0.094 |

| | | | | | | | |
|---|---|---|---|---|---|---|---|
| p.Ala185Asp | substitution | 185 | Ala | Asp | | 0.455 | 0.127 |
| p.Ala185Gly | substitution | 185 | Ala | Gly | | 1.508 | 0.185 |
| p.Ala185Ser | substitution | 185 | Ala | Ser | | 0.372 | 0.110 |
| p.Ala185Thr | substitution | 185 | Ala | Thr | | 1.150 | 0.025 |
| p.Ala185Val | substitution | 185 | Ala | Val | | 0.748 | 0.193 |
| p.Ala189= | synonymous | 189 | Ala | NA | | 0.981 | 0.112 |
| p.Ala189Arg | substitution | 189 | Ala | Arg | | 0.000 | 0.023 |
| p.Ala189Asn | substitution | 189 | Ala | Asn | | 0.028 | 0.027 |
| p.Ala189Asp | substitution | 189 | Ala | Asp | | 0.000 | 0.028 |
| p.Ala189Cys | substitution | 189 | Ala | Cys | | 0.800 | 0.007 |
| p.Ala189Gln | substitution | 189 | Ala | Gln | | 0.036 | 0.061 |
| p.Ala189Glu | substitution | 189 | Ala | Glu | | 0.045 | 0.063 |
| p.Ala189Gly | substitution | 189 | Ala | Gly | | 0.740 | 0.015 |
| p.Ala189His | substitution | 189 | Ala | His | | 0.000 | 0.017 |
| p.Ala189Ile | substitution | 189 | Ala | Ile | | 0.000 | 0.090 |
| p.Ala189Leu | substitution | 189 | Ala | Leu | | 0.000 | 0.031 |
| p.Ala189Lys | substitution | 189 | Ala | Lys | | 0.312 | 0.026 |
| p.Ala189Met | substitution | 189 | Ala | Met | | 0.047 | 0.030 |
| p.Ala189Phe | substitution | 189 | Ala | Phe | | 0.000 | 0.072 |
| p.Ala189Pro | substitution | 189 | Ala | Pro | | 0.077 | 0.087 |
| p.Ala189Ser | substitution | 189 | Ala | Ser | | 0.652 | 0.130 |
| p.Ala189Ter | nonsense | 189 | Ala | Ter | | 0.328 | 0.202 |
| p.Ala189Thr | substitution | 189 | Ala | Thr | | 0.333 | 0.088 |
| p.Ala189Trp | substitution | 189 | Ala | Trp | | 0.000 | 0.127 |
| p.Ala189Tyr | substitution | 189 | Ala | Tyr | | 0.000 | 0.019 |
| p.Ala189Val | substitution | 189 | Ala | Val | | 0.000 | 0.010 |
| p.Ala191= | synonymous | 191 | Ala | NA | | 0.802 | 0.039 |
| p.Ala191Arg | substitution | 191 | Ala | Arg | | 0.000 | 0.057 |
| p.Ala191Asp | substitution | 191 | Ala | Asp | | 0.201 | 0.021 |
| p.Ala191Cys | substitution | 191 | Ala | Cys | | 0.194 | 0.031 |
| p.Ala191Gln | substitution | 191 | Ala | Gln | | 0.000 | 0.231 |
| p.Ala191Glu | substitution | 191 | Ala | Glu | | 0.000 | 0.049 |
| p.Ala191Gly | substitution | 191 | Ala | Gly | | 0.416 | 0.008 |
| p.Ala191His | substitution | 191 | Ala | His | | 0.000 | 0.217 |
| p.Ala191Ile | substitution | 191 | Ala | Ile | | 0.000 | 0.031 |
| p.Ala191Leu | substitution | 191 | Ala | Leu | | 0.000 | 0.019 |
| p.Ala191Lys | substitution | 191 | Ala | Lys | | 0.198 | 0.084 |
| p.Ala191Met | substitution | 191 | Ala | Met | | 0.000 | 0.021 |
| p.Ala191Phe | substitution | 191 | Ala | Phe | | 0.133 | 0.014 |
| p.Ala191Pro | substitution | 191 | Ala | Pro | | 0.000 | 0.033 |
| p.Ala191Ser | substitution | 191 | Ala | Ser | | 0.656 | 0.043 |
| p.Ala191Thr | substitution | 191 | Ala | Thr | | 0.711 | 0.005 |
| p.Ala191Trp | substitution | 191 | Ala | Trp | | 0.000 | 0.050 |

| | | | | | | | |
|---|---|---|---|---|---|---|---|
| p.Ala191Tyr | substitution | 191 | Ala | Tyr | | 0.125 | 0.087 |
| p.Ala191Val | substitution | 191 | Ala | Val | | 0.000 | 0.066 |
| p.Ala214= | synonymous | 214 | Ala | NA | | 0.707 | 0.022 |
| p.Ala214Arg | substitution | 214 | Ala | Arg | | 0.000 | 0.012 |
| p.Ala214Asn | substitution | 214 | Ala | Asn | | 0.426 | 0.005 |
| p.Ala214Asp | substitution | 214 | Ala | Asp | | 0.427 | 0.012 |
| p.Ala214Cys | substitution | 214 | Ala | Cys | | 0.783 | 0.007 |
| p.Ala214Gln | substitution | 214 | Ala | Gln | | 0.000 | 0.038 |
| p.Ala214Glu | substitution | 214 | Ala | Glu | | 0.254 | 0.005 |
| p.Ala214Gly | substitution | 214 | Ala | Gly | | 0.675 | 0.008 |
| p.Ala214Ile | substitution | 214 | Ala | Ile | | 0.114 | 0.007 |
| p.Ala214Leu | substitution | 214 | Ala | Leu | | 0.002 | 0.017 |
| p.Ala214Lys | substitution | 214 | Ala | Lys | | 0.000 | 0.012 |
| p.Ala214Met | substitution | 214 | Ala | Met | | 1.168 | 0.001 |
| p.Ala214Phe | substitution | 214 | Ala | Phe | | 0.364 | 0.011 |
| p.Ala214Pro | substitution | 214 | Ala | Pro | | 0.692 | 0.011 |
| p.Ala214Ser | substitution | 214 | Ala | Ser | | 0.516 | 0.063 |
| p.Ala214Ter | nonsense | 214 | Ala | Ter | | 0.000 | 0.018 |
| p.Ala214Thr | substitution | 214 | Ala | Thr | | 0.610 | 0.002 |
| p.Ala214Trp | substitution | 214 | Ala | Trp | | 0.000 | 0.028 |
| p.Ala214Tyr | substitution | 214 | Ala | Tyr | | 0.000 | 0.017 |
| p.Ala214Val | substitution | 214 | Ala | Val | | 0.000 | 0.007 |
| p.Ala219= | synonymous | 219 | Ala | NA | | 0.850 | 0.017 |
| p.Ala219Arg | substitution | 219 | Ala | Arg | | 0.034 | 0.137 |
| p.Ala219Asn | substitution | 219 | Ala | Asn | | 0.000 | 0.030 |
| p.Ala219Asp | substitution | 219 | Ala | Asp | | 0.000 | 0.099 |
| p.Ala219Cys | substitution | 219 | Ala | Cys | | 0.000 | 0.041 |
| p.Ala219Glu | substitution | 219 | Ala | Glu | | 0.307 | 0.079 |
| p.Ala219Gly | substitution | 219 | Ala | Gly | | 0.693 | 0.088 |
| p.Ala219His | substitution | 219 | Ala | His | | 0.000 | 0.042 |
| p.Ala219Leu | substitution | 219 | Ala | Leu | | 0.000 | 0.011 |
| p.Ala219Lys | substitution | 219 | Ala | Lys | | 0.000 | 0.215 |
| p.Ala219Phe | substitution | 219 | Ala | Phe | | 0.000 | 0.172 |
| p.Ala219Pro | substitution | 219 | Ala | Pro | | 0.000 | 0.021 |
| p.Ala219Ser | substitution | 219 | Ala | Ser | | 0.704 | 0.068 |
| p.Ala219Thr | substitution | 219 | Ala | Thr | | 0.492 | 0.016 |
| p.Ala219Tyr | substitution | 219 | Ala | Tyr | | 0.197 | 0.051 |
| p.Ala219Val | substitution | 219 | Ala | Val | | 1.027 | 0.156 |
| p.Ala226= | synonymous | 226 | Ala | NA | | 1.084 | 0.002 |
| p.Ala226Arg | substitution | 226 | Ala | Arg | | 0.587 | 0.015 |
| p.Ala226Asn | substitution | 226 | Ala | Asn | | 0.074 | 0.016 |
| p.Ala226Asp | substitution | 226 | Ala | Asp | | 0.863 | 0.013 |
| p.Ala226Cys | substitution | 226 | Ala | Cys | | 0.947 | 0.017 |

| | | | | | | |
|---|---|---|---|---|---|---|
| p.Ala226Gln | substitution | 226 | Ala | Gln | 0.963 | 0.004 |
| p.Ala226Glu | substitution | 226 | Ala | Glu | 1.907 | 0.001 |
| p.Ala226Gly | substitution | 226 | Ala | Gly | 0.906 | 0.012 |
| p.Ala226His | substitution | 226 | Ala | His | 0.964 | 0.009 |
| p.Ala226Ile | substitution | 226 | Ala | Ile | 0.562 | 0.005 |
| p.Ala226Leu | substitution | 226 | Ala | Leu | 0.675 | 0.025 |
| p.Ala226Lys | substitution | 226 | Ala | Lys | 1.417 | 0.150 |
| p.Ala226Met | substitution | 226 | Ala | Met | 1.290 | 0.008 |
| p.Ala226Phe | substitution | 226 | Ala | Phe | 0.791 | 0.010 |
| p.Ala226Pro | substitution | 226 | Ala | Pro | 0.503 | 0.021 |
| p.Ala226Ser | substitution | 226 | Ala | Ser | 0.663 | 0.015 |
| p.Ala226Ter | nonsense | 226 | Ala | Ter | 0.028 | 0.018 |
| p.Ala226Thr | substitution | 226 | Ala | Thr | 0.707 | 0.049 |
| p.Ala226Trp | substitution | 226 | Ala | Trp | 1.563 | 0.001 |
| p.Ala226Tyr | substitution | 226 | Ala | Tyr | 0.996 | 0.057 |
| p.Ala226Val | substitution | 226 | Ala | Val | 0.982 | 0.020 |
| p.Ala249= | synonymous | 249 | Ala | NA | 1.317 | 0.018 |
| p.Ala249Arg | substitution | 249 | Ala | Arg | 0.000 | 0.023 |
| p.Ala249Asn | substitution | 249 | Ala | Asn | 0.000 | 0.075 |
| p.Ala249Asp | substitution | 249 | Ala | Asp | 0.140 | 0.014 |
| p.Ala249Gln | substitution | 249 | Ala | Gln | 0.000 | 0.039 |
| p.Ala249Glu | substitution | 249 | Ala | Glu | 0.030 | 0.064 |
| p.Ala249Gly | substitution | 249 | Ala | Gly | 0.000 | 0.010 |
| p.Ala249His | substitution | 249 | Ala | His | 0.000 | 0.027 |
| p.Ala249Ile | substitution | 249 | Ala | Ile | 0.000 | 0.075 |
| p.Ala249Leu | substitution | 249 | Ala | Leu | 0.199 | 0.043 |
| p.Ala249Lys | substitution | 249 | Ala | Lys | 0.166 | 0.019 |
| p.Ala249Met | substitution | 249 | Ala | Met | 0.000 | 0.041 |
| p.Ala249Phe | substitution | 249 | Ala | Phe | 0.000 | 0.018 |
| p.Ala249Pro | substitution | 249 | Ala | Pro | 0.191 | 0.084 |
| p.Ala249Ser | substitution | 249 | Ala | Ser | 1.076 | 0.030 |
| p.Ala249Ter | nonsense | 249 | Ala | Ter | 0.000 | 0.080 |
| p.Ala249Thr | substitution | 249 | Ala | Thr | 1.199 | 0.012 |
| p.Ala249Trp | substitution | 249 | Ala | Trp | 0.000 | 0.029 |
| p.Ala249Tyr | substitution | 249 | Ala | Tyr | 0.000 | 0.015 |
| p.Ala249Val | substitution | 249 | Ala | Val | 0.546 | 0.023 |
| p.Ala252= | synonymous | 252 | Ala | NA | 0.952 | 0.060 |
| p.Ala252Arg | substitution | 252 | Ala | Arg | 0.085 | 0.074 |
| p.Ala252Asn | substitution | 252 | Ala | Asn | 0.000 | 0.013 |
| p.Ala252Asp | substitution | 252 | Ala | Asp | 0.000 | 0.093 |
| p.Ala252Cys | substitution | 252 | Ala | Cys | 0.000 | 0.226 |
| p.Ala252Glu | substitution | 252 | Ala | Glu | 0.668 | 0.006 |
| p.Ala252Gly | substitution | 252 | Ala | Gly | 0.419 | 0.137 |

| | | | | | | |
|---|---|---|---|---|---|---|
| p.Ala252His | substitution | 252 | Ala | His | 0.000 | 0.184 |
| p.Ala252Leu | substitution | 252 | Ala | Leu | 0.060 | 0.026 |
| p.Ala252Phe | substitution | 252 | Ala | Phe | 0.000 | 0.091 |
| p.Ala252Pro | substitution | 252 | Ala | Pro | 0.000 | 0.072 |
| p.Ala252Ser | substitution | 252 | Ala | Ser | 0.000 | 0.041 |
| p.Ala252Ter | nonsense | 252 | Ala | Ter | 1.334 | 0.304 |
| p.Ala252Thr | substitution | 252 | Ala | Thr | 1.196 | 0.105 |
| p.Ala252Trp | substitution | 252 | Ala | Trp | 0.000 | 0.097 |
| p.Ala252Tyr | substitution | 252 | Ala | Tyr | 0.000 | 0.037 |
| p.Ala252Val | substitution | 252 | Ala | Val | 1.218 | 0.026 |
| p.Ala266= | synonymous | 266 | Ala | NA | 0.844 | 0.045 |
| p.Ala266Arg | substitution | 266 | Ala | Arg | 0.118 | 0.011 |
| p.Ala266Asp | substitution | 266 | Ala | Asp | 0.701 | 0.026 |
| p.Ala266Cys | substitution | 266 | Ala | Cys | 0.000 | 0.129 |
| p.Ala266Gln | substitution | 266 | Ala | Gln | 0.000 | 0.102 |
| p.Ala266Glu | substitution | 266 | Ala | Glu | 0.222 | 0.056 |
| p.Ala266Gly | substitution | 266 | Ala | Gly | 0.175 | 0.077 |
| p.Ala266Ile | substitution | 266 | Ala | Ile | 0.131 | 0.214 |
| p.Ala266Leu | substitution | 266 | Ala | Leu | 0.050 | 0.040 |
| p.Ala266Lys | substitution | 266 | Ala | Lys | 0.627 | 0.008 |
| p.Ala266Pro | substitution | 266 | Ala | Pro | 0.000 | 0.119 |
| p.Ala266Ser | substitution | 266 | Ala | Ser | 0.000 | 0.192 |
| p.Ala266Ter | nonsense | 266 | Ala | Ter | 0.060 | 0.122 |
| p.Ala266Thr | substitution | 266 | Ala | Thr | 0.514 | 0.509 |
| p.Ala266Val | substitution | 266 | Ala | Val | 1.316 | 0.079 |
| p.Ala270= | synonymous | 270 | Ala | NA | 0.329 | 0.008 |
| p.Ala270Arg | substitution | 270 | Ala | Arg | 1.004 | 0.032 |
| p.Ala270Asn | substitution | 270 | Ala | Asn | 0.091 | 0.014 |
| p.Ala270Asp | substitution | 270 | Ala | Asp | 0.951 | 0.042 |
| p.Ala270Gln | substitution | 270 | Ala | Gln | 1.070 | 0.005 |
| p.Ala270Glu | substitution | 270 | Ala | Glu | 1.187 | 0.006 |
| p.Ala270Gly | substitution | 270 | Ala | Gly | 0.399 | 0.033 |
| p.Ala270His | substitution | 270 | Ala | His | 0.857 | 0.004 |
| p.Ala270Ile | substitution | 270 | Ala | Ile | 0.558 | 0.006 |
| p.Ala270Leu | substitution | 270 | Ala | Leu | 0.000 | 0.004 |
| p.Ala270Lys | substitution | 270 | Ala | Lys | 1.756 | 0.004 |
| p.Ala270Met | substitution | 270 | Ala | Met | 1.207 | 0.005 |
| p.Ala270Phe | substitution | 270 | Ala | Phe | 1.138 | 0.006 |
| p.Ala270Pro | substitution | 270 | Ala | Pro | 0.000 | 0.053 |
| p.Ala270Ser | substitution | 270 | Ala | Ser | 0.534 | 0.163 |
| p.Ala270Ter | nonsense | 270 | Ala | Ter | 0.436 | 0.034 |
| p.Ala270Thr | substitution | 270 | Ala | Thr | 0.516 | 0.015 |
| p.Ala270Trp | substitution | 270 | Ala | Trp | 0.519 | 0.008 |

| | | | | | | | |
|---|---|---|---|---|---|---|---|
| p.Ala270Tyr | substitution | 270 | Ala | Tyr | | 0.386 | 0.008 |
| p.Ala270Val | substitution | 270 | Ala | Val | | 0.965 | 0.006 |
| p.Ala297= | synonymous | 297 | Ala | NA | | 1.039 | 0.018 |
| p.Ala297Arg | substitution | 297 | Ala | Arg | | 0.000 | 0.055 |
| p.Ala297Asn | substitution | 297 | Ala | Asn | | 0.000 | 0.079 |
| p.Ala297Asp | substitution | 297 | Ala | Asp | | 0.844 | 0.114 |
| p.Ala297Cys | substitution | 297 | Ala | Cys | | 1.461 | 0.006 |
| p.Ala297Gln | substitution | 297 | Ala | Gln | | 0.245 | 0.013 |
| p.Ala297Glu | substitution | 297 | Ala | Glu | | 0.272 | 0.013 |
| p.Ala297Gly | substitution | 297 | Ala | Gly | | 0.815 | 0.124 |
| p.Ala297Ile | substitution | 297 | Ala | Ile | | 0.000 | 0.125 |
| p.Ala297Leu | substitution | 297 | Ala | Leu | | 0.299 | 0.134 |
| p.Ala297Lys | substitution | 297 | Ala | Lys | | 0.000 | 0.012 |
| p.Ala297Met | substitution | 297 | Ala | Met | | 0.000 | 0.221 |
| p.Ala297Phe | substitution | 297 | Ala | Phe | | 0.000 | 0.040 |
| p.Ala297Pro | substitution | 297 | Ala | Pro | | 0.000 | 0.235 |
| p.Ala297Ser | substitution | 297 | Ala | Ser | | 0.395 | 0.076 |
| p.Ala297Ter | nonsense | 297 | Ala | Ter | | 0.000 | 0.016 |
| p.Ala297Thr | substitution | 297 | Ala | Thr | | 0.929 | 0.040 |
| p.Ala297Trp | substitution | 297 | Ala | Trp | | 0.912 | 0.007 |
| p.Ala297Tyr | substitution | 297 | Ala | Tyr | | 0.184 | 0.062 |
| p.Ala297Val | substitution | 297 | Ala | Val | | 0.527 | 0.223 |
| p.Ala303= | synonymous | 303 | Ala | NA | | 0.698 | 0.011 |
| p.Ala303Arg | substitution | 303 | Ala | Arg | | 1.014 | 0.130 |
| p.Ala303Asn | substitution | 303 | Ala | Asn | | 1.306 | 0.077 |
| p.Ala303Asp | substitution | 303 | Ala | Asp | | 1.442 | 0.139 |
| p.Ala303Gln | substitution | 303 | Ala | Gln | | 0.000 | 0.024 |
| p.Ala303Glu | substitution | 303 | Ala | Glu | | 0.788 | 0.003 |
| p.Ala303Gly | substitution | 303 | Ala | Gly | | 0.119 | 0.041 |
| p.Ala303His | substitution | 303 | Ala | His | | 0.016 | 0.075 |
| p.Ala303Ile | substitution | 303 | Ala | Ile | | 1.272 | 0.004 |
| p.Ala303Leu | substitution | 303 | Ala | Leu | | 1.263 | 0.051 |
| p.Ala303Lys | substitution | 303 | Ala | Lys | | 1.136 | 0.003 |
| p.Ala303Met | substitution | 303 | Ala | Met | | 0.101 | 0.219 |
| p.Ala303Phe | substitution | 303 | Ala | Phe | | 1.547 | 0.002 |
| p.Ala303Pro | substitution | 303 | Ala | Pro | | 0.000 | 0.053 |
| p.Ala303Ser | substitution | 303 | Ala | Ser | | 0.796 | 0.098 |
| p.Ala303Ter | nonsense | 303 | Ala | Ter | | 0.732 | 0.034 |
| p.Ala303Thr | substitution | 303 | Ala | Thr | | 0.961 | 0.021 |
| p.Ala303Trp | substitution | 303 | Ala | Trp | | 0.000 | 0.022 |
| p.Ala303Tyr | substitution | 303 | Ala | Tyr | | 0.418 | 0.048 |
| p.Ala303Val | substitution | 303 | Ala | Val | | 0.245 | 0.053 |
| p.Ala31= | synonymous | 31 | Ala | NA | | 0.634 | 0.011 |

| | | | | | | |
|---|---|---|---|---|---|---|
| p.Ala31Arg | substitution | 31 | Ala | Arg | 0.000 | 0.006 |
| p.Ala31Asn | substitution | 31 | Ala | Asn | 0.000 | 0.008 |
| p.Ala31Asp | substitution | 31 | Ala | Asp | 0.000 | 0.003 |
| p.Ala31Cys | substitution | 31 | Ala | Cys | 0.000 | 0.002 |
| p.Ala31Gln | substitution | 31 | Ala | Gln | 0.000 | 0.009 |
| p.Ala31Glu | substitution | 31 | Ala | Glu | 0.000 | 0.001 |
| p.Ala31Gly | substitution | 31 | Ala | Gly | 0.664 | 0.000 |
| p.Ala31His | substitution | 31 | Ala | His | 0.000 | 0.005 |
| p.Ala31Ile | substitution | 31 | Ala | Ile | 0.000 | 0.003 |
| p.Ala31Leu | substitution | 31 | Ala | Leu | 0.000 | 0.008 |
| p.Ala31Lys | substitution | 31 | Ala | Lys | 0.000 | 0.006 |
| p.Ala31Met | substitution | 31 | Ala | Met | 0.000 | 0.021 |
| p.Ala31Phe | substitution | 31 | Ala | Phe | 0.000 | 0.003 |
| p.Ala31Pro | substitution | 31 | Ala | Pro | 0.000 | 0.013 |
| p.Ala31Ser | substitution | 31 | Ala | Ser | 0.846 | 0.012 |
| p.Ala31Ter | nonsense | 31 | Ala | Ter | 0.000 | 0.009 |
| p.Ala31Thr | substitution | 31 | Ala | Thr | 0.000 | 0.094 |
| p.Ala31Trp | substitution | 31 | Ala | Trp | 0.000 | 0.003 |
| p.Ala31Tyr | substitution | 31 | Ala | Tyr | 0.000 | 0.006 |
| p.Ala31Val | substitution | 31 | Ala | Val | 0.000 | 0.015 |
| p.Ala320= | synonymous | 320 | Ala | NA | 0.820 | 0.028 |
| p.Ala320Arg | substitution | 320 | Ala | Arg | 0.110 | 0.021 |
| p.Ala320Asn | substitution | 320 | Ala | Asn | 0.000 | 0.138 |
| p.Ala320Asp | substitution | 320 | Ala | Asp | 0.723 | 0.009 |
| p.Ala320Cys | substitution | 320 | Ala | Cys | 0.468 | 0.006 |
| p.Ala320Gln | substitution | 320 | Ala | Gln | 0.042 | 0.189 |
| p.Ala320Glu | substitution | 320 | Ala | Glu | 0.880 | 0.002 |
| p.Ala320Gly | substitution | 320 | Ala | Gly | 1.157 | 0.020 |
| p.Ala320His | substitution | 320 | Ala | His | 0.000 | 0.046 |
| p.Ala320Ile | substitution | 320 | Ala | Ile | 0.000 | 0.038 |
| p.Ala320Leu | substitution | 320 | Ala | Leu | 0.643 | 0.237 |
| p.Ala320Lys | substitution | 320 | Ala | Lys | 0.228 | 0.007 |
| p.Ala320Met | substitution | 320 | Ala | Met | 0.257 | 0.232 |
| p.Ala320Phe | substitution | 320 | Ala | Phe | 0.009 | 0.013 |
| p.Ala320Pro | substitution | 320 | Ala | Pro | 0.170 | 0.139 |
| p.Ala320Ser | substitution | 320 | Ala | Ser | 0.000 | 0.034 |
| p.Ala320Ter | nonsense | 320 | Ala | Ter | 0.000 | 0.222 |
| p.Ala320Thr | substitution | 320 | Ala | Thr | 0.962 | 0.137 |
| p.Ala320Trp | substitution | 320 | Ala | Trp | 1.118 | 0.012 |
| p.Ala320Tyr | substitution | 320 | Ala | Tyr | 0.000 | 0.054 |
| p.Ala320Val | substitution | 320 | Ala | Val | 0.829 | 0.003 |
| p.Ala330= | synonymous | 330 | Ala | NA | 0.484 | 0.095 |
| p.Ala330Arg | substitution | 330 | Ala | Arg | 1.237 | 0.001 |

| | | | | | | |
|---|---|---|---|---|---|---|
| p.Ala330Asn | substitution | 330 | Ala | Asn | 1.052 | 0.004 |
| p.Ala330Asp | substitution | 330 | Ala | Asp | 0.931 | 0.006 |
| p.Ala330Cys | substitution | 330 | Ala | Cys | 1.699 | 0.006 |
| p.Ala330Gly | substitution | 330 | Ala | Gly | 0.742 | 0.405 |
| p.Ala330His | substitution | 330 | Ala | His | 0.023 | 0.076 |
| p.Ala330Ile | substitution | 330 | Ala | Ile | 0.000 | 0.106 |
| p.Ala330Leu | substitution | 330 | Ala | Leu | 1.325 | 0.129 |
| p.Ala330Met | substitution | 330 | Ala | Met | 0.000 | 0.089 |
| p.Ala330Phe | substitution | 330 | Ala | Phe | 0.544 | 0.028 |
| p.Ala330Pro | substitution | 330 | Ala | Pro | 0.459 | 0.049 |
| p.Ala330Ser | substitution | 330 | Ala | Ser | 0.614 | 0.146 |
| p.Ala330Thr | substitution | 330 | Ala | Thr | 0.794 | 0.156 |
| p.Ala330Tyr | substitution | 330 | Ala | Tyr | 0.000 | 0.090 |
| p.Ala330Val | substitution | 330 | Ala | Val | 1.198 | 0.009 |
| p.Ala331= | synonymous | 331 | Ala | NA | 0.810 | 0.806 |
| p.Ala331Arg | substitution | 331 | Ala | Arg | 0.000 | 0.148 |
| p.Ala331Asn | substitution | 331 | Ala | Asn | 0.000 | 0.121 |
| p.Ala331Asp | substitution | 331 | Ala | Asp | 0.548 | 0.221 |
| p.Ala331Cys | substitution | 331 | Ala | Cys | 1.009 | 0.008 |
| p.Ala331Glu | substitution | 331 | Ala | Glu | 0.125 | 0.015 |
| p.Ala331Gly | substitution | 331 | Ala | Gly | 1.589 | 0.047 |
| p.Ala331Leu | substitution | 331 | Ala | Leu | 0.000 | 0.028 |
| p.Ala331Met | substitution | 331 | Ala | Met | 0.099 | 0.012 |
| p.Ala331Pro | substitution | 331 | Ala | Pro | 0.000 | 0.019 |
| p.Ala331Ser | substitution | 331 | Ala | Ser | 0.199 | 0.068 |
| p.Ala331Ter | nonsense | 331 | Ala | Ter | 0.177 | 0.014 |
| p.Ala331Thr | substitution | 331 | Ala | Thr | 0.686 | 0.116 |
| p.Ala331Val | substitution | 331 | Ala | Val | 0.265 | 0.040 |
| p.Ala339= | synonymous | 339 | Ala | NA | 0.913 | 0.013 |
| p.Ala339Arg | substitution | 339 | Ala | Arg | 0.128 | 0.092 |
| p.Ala339Asp | substitution | 339 | Ala | Asp | 0.489 | 0.033 |
| p.Ala339Cys | substitution | 339 | Ala | Cys | 0.000 | 0.018 |
| p.Ala339Glu | substitution | 339 | Ala | Glu | 0.000 | 0.111 |
| p.Ala339Gly | substitution | 339 | Ala | Gly | 1.296 | 0.025 |
| p.Ala339Ile | substitution | 339 | Ala | Ile | 0.000 | 0.035 |
| p.Ala339Met | substitution | 339 | Ala | Met | 0.000 | 0.186 |
| p.Ala339Pro | substitution | 339 | Ala | Pro | 0.000 | 0.188 |
| p.Ala339Ser | substitution | 339 | Ala | Ser | 0.523 | 0.585 |
| p.Ala339Ter | nonsense | 339 | Ala | Ter | 0.000 | 0.257 |
| p.Ala339Thr | substitution | 339 | Ala | Thr | 1.049 | 0.011 |
| p.Ala339Tyr | substitution | 339 | Ala | Tyr | 0.183 | 0.200 |
| p.Ala339Val | substitution | 339 | Ala | Val | 0.839 | 0.074 |
| p.Ala347= | synonymous | 347 | Ala | NA | 1.194 | 0.018 |

| | | | | | | | |
|---|---|---|---|---|---|---|---|
| p.Ala347Arg | substitution | 347 | Ala | Arg | | 1.450 | 0.021 |
| p.Ala347Asp | substitution | 347 | Ala | Asp | | 0.000 | 0.128 |
| p.Ala347Cys | substitution | 347 | Ala | Cys | | 0.132 | 0.143 |
| p.Ala347Gln | substitution | 347 | Ala | Gln | | 1.076 | 0.046 |
| p.Ala347Gly | substitution | 347 | Ala | Gly | | 0.000 | 0.078 |
| p.Ala347His | substitution | 347 | Ala | His | | 0.255 | 0.099 |
| p.Ala347Ile | substitution | 347 | Ala | Ile | | 0.011 | 0.258 |
| p.Ala347Lys | substitution | 347 | Ala | Lys | | 0.000 | 0.268 |
| p.Ala347Met | substitution | 347 | Ala | Met | | 0.374 | 0.053 |
| p.Ala347Phe | substitution | 347 | Ala | Phe | | 0.023 | 0.073 |
| p.Ala347Pro | substitution | 347 | Ala | Pro | | 1.638 | 0.117 |
| p.Ala347Ser | substitution | 347 | Ala | Ser | | 0.918 | 0.080 |
| p.Ala347Thr | substitution | 347 | Ala | Thr | | 1.147 | 0.063 |
| p.Ala347Trp | substitution | 347 | Ala | Trp | | 0.000 | 0.058 |
| p.Ala347Tyr | substitution | 347 | Ala | Tyr | | 1.499 | 0.010 |
| p.Ala347Val | substitution | 347 | Ala | Val | | 0.013 | 0.030 |
| p.Ala354= | synonymous | 354 | Ala | NA | | 1.566 | 0.245 |
| p.Ala354Arg | substitution | 354 | Ala | Arg | | 0.034 | 0.267 |
| p.Ala354Asn | substitution | 354 | Ala | Asn | | 0.000 | 0.199 |
| p.Ala354Asp | substitution | 354 | Ala | Asp | | 1.285 | 0.006 |
| p.Ala354Glu | substitution | 354 | Ala | Glu | | 1.001 | 0.080 |
| p.Ala354Gly | substitution | 354 | Ala | Gly | | 1.246 | 0.389 |
| p.Ala354Ile | substitution | 354 | Ala | Ile | | 0.000 | 0.127 |
| p.Ala354Leu | substitution | 354 | Ala | Leu | | 0.121 | 0.028 |
| p.Ala354Met | substitution | 354 | Ala | Met | | 1.060 | 0.006 |
| p.Ala354Pro | substitution | 354 | Ala | Pro | | 1.173 | 0.018 |
| p.Ala354Ser | substitution | 354 | Ala | Ser | | 0.203 | 0.054 |
| p.Ala354Thr | substitution | 354 | Ala | Thr | | 1.096 | 0.102 |
| p.Ala354Tyr | substitution | 354 | Ala | Tyr | | 0.000 | 0.078 |
| p.Ala354Val | substitution | 354 | Ala | Val | | 0.313 | 0.060 |
| p.Ala360= | synonymous | 360 | Ala | NA | | 1.173 | 0.032 |
| p.Ala360Arg | substitution | 360 | Ala | Arg | | 0.380 | 0.051 |
| p.Ala360Asn | substitution | 360 | Ala | Asn | | 0.000 | 0.072 |
| p.Ala360Asp | substitution | 360 | Ala | Asp | | 0.930 | 0.015 |
| p.Ala360Cys | substitution | 360 | Ala | Cys | | 0.000 | 0.219 |
| p.Ala360Gln | substitution | 360 | Ala | Gln | | 0.000 | 0.026 |
| p.Ala360Gly | substitution | 360 | Ala | Gly | | 1.113 | 0.085 |
| p.Ala360Ile | substitution | 360 | Ala | Ile | | 0.266 | 0.154 |
| p.Ala360Leu | substitution | 360 | Ala | Leu | | 0.534 | 0.083 |
| p.Ala360Lys | substitution | 360 | Ala | Lys | | 0.000 | 0.021 |
| p.Ala360Met | substitution | 360 | Ala | Met | | 0.000 | 0.053 |
| p.Ala360Pro | substitution | 360 | Ala | Pro | | 0.000 | 0.123 |
| p.Ala360Ser | substitution | 360 | Ala | Ser | | 0.910 | 0.022 |

| | | | | | | | |
|---|---|---|---|---|---|---|---|
| p.Ala360Thr | substitution | 360 | Ala | Thr | | 1.064 | 0.154 |
| p.Ala360Trp | substitution | 360 | Ala | Trp | | 0.000 | 0.027 |
| p.Ala360Val | substitution | 360 | Ala | Val | | 1.311 | 0.009 |
| p.Ala40= | synonymous | 40 | Ala | NA | | 0.959 | 0.019 |
| p.Ala40Arg | substitution | 40 | Ala | Arg | | 0.586 | 0.022 |
| p.Ala40Asn | substitution | 40 | Ala | Asn | | 0.558 | 0.008 |
| p.Ala40Asp | substitution | 40 | Ala | Asp | | 0.891 | 0.007 |
| p.Ala40Cys | substitution | 40 | Ala | Cys | | 1.208 | 0.004 |
| p.Ala40Gln | substitution | 40 | Ala | Gln | | 0.591 | 0.125 |
| p.Ala40Glu | substitution | 40 | Ala | Glu | | 0.800 | 0.007 |
| p.Ala40Gly | substitution | 40 | Ala | Gly | | 0.944 | 0.002 |
| p.Ala40His | substitution | 40 | Ala | His | | 0.658 | 0.006 |
| p.Ala40Ile | substitution | 40 | Ala | Ile | | 0.615 | 0.010 |
| p.Ala40Leu | substitution | 40 | Ala | Leu | | 0.728 | 0.011 |
| p.Ala40Lys | substitution | 40 | Ala | Lys | | 1.049 | 0.012 |
| p.Ala40Met | substitution | 40 | Ala | Met | | 1.085 | 0.006 |
| p.Ala40Phe | substitution | 40 | Ala | Phe | | 0.554 | 0.007 |
| p.Ala40Pro | substitution | 40 | Ala | Pro | | 0.061 | 0.077 |
| p.Ala40Ser | substitution | 40 | Ala | Ser | | 0.775 | 0.017 |
| p.Ala40Ter | nonsense | 40 | Ala | Ter | | 0.248 | 0.077 |
| p.Ala40Thr | substitution | 40 | Ala | Thr | | 0.945 | 0.030 |
| p.Ala40Trp | substitution | 40 | Ala | Trp | | 0.820 | 0.006 |
| p.Ala40Tyr | substitution | 40 | Ala | Tyr | | 1.097 | 0.004 |
| p.Ala40Val | substitution | 40 | Ala | Val | | 0.967 | 0.021 |
| p.Ala44= | synonymous | 44 | Ala | NA | | 0.977 | 0.002 |
| p.Ala44Arg | substitution | 44 | Ala | Arg | | 1.006 | 0.008 |
| p.Ala44Asn | substitution | 44 | Ala | Asn | | 1.035 | 0.013 |
| p.Ala44Asp | substitution | 44 | Ala | Asp | | 0.726 | 0.005 |
| p.Ala44Cys | substitution | 44 | Ala | Cys | | 0.333 | 0.031 |
| p.Ala44Gln | substitution | 44 | Ala | Gln | | 0.789 | 0.005 |
| p.Ala44Glu | substitution | 44 | Ala | Glu | | 0.987 | 0.009 |
| p.Ala44Gly | substitution | 44 | Ala | Gly | | 0.904 | 0.009 |
| p.Ala44His | substitution | 44 | Ala | His | | 0.767 | 0.018 |
| p.Ala44Ile | substitution | 44 | Ala | Ile | | 0.754 | 0.163 |
| p.Ala44Leu | substitution | 44 | Ala | Leu | | 0.792 | 0.012 |
| p.Ala44Lys | substitution | 44 | Ala | Lys | | 1.073 | 0.004 |
| p.Ala44Met | substitution | 44 | Ala | Met | | 0.110 | 0.010 |
| p.Ala44Phe | substitution | 44 | Ala | Phe | | 0.980 | 0.015 |
| p.Ala44Pro | substitution | 44 | Ala | Pro | | 0.464 | 0.014 |
| p.Ala44Ser | substitution | 44 | Ala | Ser | | 0.934 | 0.007 |
| p.Ala44Ter | nonsense | 44 | Ala | Ter | | 0.000 | 0.086 |
| p.Ala44Thr | substitution | 44 | Ala | Thr | | 0.964 | 0.015 |
| p.Ala44Trp | substitution | 44 | Ala | Trp | | 0.796 | 0.002 |

| | | | | | | | |
|---|---|---|---|---|---|---|---|
| p.Ala44Tyr | substitution | 44 | Ala | Tyr | | 1.015 | 0.014 |
| p.Ala44Val | substitution | 44 | Ala | Val | | 1.059 | 0.022 |
| p.Ala55= | synonymous | 55 | Ala | NA | | 1.114 | 0.020 |
| p.Ala55Arg | substitution | 55 | Ala | Arg | | 0.807 | 0.024 |
| p.Ala55Asn | substitution | 55 | Ala | Asn | | 0.595 | 0.008 |
| p.Ala55Asp | substitution | 55 | Ala | Asp | | 0.884 | 0.003 |
| p.Ala55Cys | substitution | 55 | Ala | Cys | | 0.630 | 0.009 |
| p.Ala55Gln | substitution | 55 | Ala | Gln | | 0.819 | 0.017 |
| p.Ala55Glu | substitution | 55 | Ala | Glu | | 1.611 | 0.004 |
| p.Ala55Gly | substitution | 55 | Ala | Gly | | 0.686 | 0.003 |
| p.Ala55His | substitution | 55 | Ala | His | | 0.965 | 0.008 |
| p.Ala55Ile | substitution | 55 | Ala | Ile | | 0.670 | 0.005 |
| p.Ala55Leu | substitution | 55 | Ala | Leu | | 0.823 | 0.004 |
| p.Ala55Lys | substitution | 55 | Ala | Lys | | 0.584 | 0.007 |
| p.Ala55Met | substitution | 55 | Ala | Met | | 0.578 | 0.009 |
| p.Ala55Phe | substitution | 55 | Ala | Phe | | 1.213 | 0.007 |
| p.Ala55Pro | substitution | 55 | Ala | Pro | | 0.896 | 0.018 |
| p.Ala55Ser | substitution | 55 | Ala | Ser | | 0.966 | 0.009 |
| p.Ala55Ter | nonsense | 55 | Ala | Ter | | 0.053 | 0.017 |
| p.Ala55Thr | substitution | 55 | Ala | Thr | | 0.683 | 0.046 |
| p.Ala55Trp | substitution | 55 | Ala | Trp | | 0.680 | 0.010 |
| p.Ala55Tyr | substitution | 55 | Ala | Tyr | | 0.892 | 0.006 |
| p.Ala55Val | substitution | 55 | Ala | Val | | 0.939 | 0.010 |
| p.Ala67= | synonymous | 67 | Ala | NA | | 1.199 | 0.972 |
| p.Ala67Arg | substitution | 67 | Ala | Arg | | 1.126 | 0.121 |
| p.Ala67Asn | substitution | 67 | Ala | Asn | | 0.230 | 0.090 |
| p.Ala67Asp | substitution | 67 | Ala | Asp | | 1.857 | 0.037 |
| p.Ala67Gln | substitution | 67 | Ala | Gln | | 0.665 | 0.068 |
| p.Ala67Glu | substitution | 67 | Ala | Glu | | 0.000 | 0.374 |
| p.Ala67Gly | substitution | 67 | Ala | Gly | | 0.759 | 0.047 |
| p.Ala67His | substitution | 67 | Ala | His | | 0.109 | 0.390 |
| p.Ala67Ile | substitution | 67 | Ala | Ile | | 1.271 | 0.013 |
| p.Ala67Leu | substitution | 67 | Ala | Leu | | 0.708 | 0.195 |
| p.Ala67Lys | substitution | 67 | Ala | Lys | | 1.575 | 0.029 |
| p.Ala67Met | substitution | 67 | Ala | Met | | 0.000 | 0.058 |
| p.Ala67Phe | substitution | 67 | Ala | Phe | | 0.000 | 0.160 |
| p.Ala67Pro | substitution | 67 | Ala | Pro | | 0.100 | 0.374 |
| p.Ala67Ser | substitution | 67 | Ala | Ser | | 0.000 | 0.091 |
| p.Ala67Ter | nonsense | 67 | Ala | Ter | | 0.000 | 0.307 |
| p.Ala67Thr | substitution | 67 | Ala | Thr | | 1.139 | 0.022 |
| p.Ala67Trp | substitution | 67 | Ala | Trp | | 0.000 | 0.042 |
| p.Ala67Val | substitution | 67 | Ala | Val | | 0.845 | 0.094 |
| p.Ala84= | synonymous | 84 | Ala | NA | | 0.816 | 0.007 |

| | | | | | | |
|---|---|---|---|---|---|---|
| p.Ala84Arg | substitution | 84 | Ala | Arg | 0.377 | 0.023 |
| p.Ala84Asn | substitution | 84 | Ala | Asn | 0.557 | 0.015 |
| p.Ala84Asp | substitution | 84 | Ala | Asp | 0.637 | 0.007 |
| p.Ala84Cys | substitution | 84 | Ala | Cys | 0.596 | 0.018 |
| p.Ala84Gln | substitution | 84 | Ala | Gln | 0.445 | 0.012 |
| p.Ala84Glu | substitution | 84 | Ala | Glu | 0.505 | 0.007 |
| p.Ala84Gly | substitution | 84 | Ala | Gly | 0.565 | 0.019 |
| p.Ala84His | substitution | 84 | Ala | His | 0.680 | 0.035 |
| p.Ala84Ile | substitution | 84 | Ala | Ile | 0.354 | 0.084 |
| p.Ala84Leu | substitution | 84 | Ala | Leu | 0.507 | 0.071 |
| p.Ala84Lys | substitution | 84 | Ala | Lys | 0.677 | 0.016 |
| p.Ala84Met | substitution | 84 | Ala | Met | 1.114 | 0.024 |
| p.Ala84Phe | substitution | 84 | Ala | Phe | 0.904 | 0.028 |
| p.Ala84Pro | substitution | 84 | Ala | Pro | 0.445 | 0.100 |
| p.Ala84Ser | substitution | 84 | Ala | Ser | 0.763 | 0.051 |
| p.Ala84Ter | nonsense | 84 | Ala | Ter | 0.055 | 0.077 |
| p.Ala84Thr | substitution | 84 | Ala | Thr | 0.895 | 0.038 |
| p.Ala84Trp | substitution | 84 | Ala | Trp | 0.410 | 0.021 |
| p.Ala84Tyr | substitution | 84 | Ala | Tyr | 0.680 | 0.590 |
| p.Ala84Val | substitution | 84 | Ala | Val | 0.597 | 0.017 |
| p.Arg116= | synonymous | 116 | Arg | NA | 1.080 | 0.020 |
| p.Arg116Ala | substitution | 116 | Arg | Ala | 0.624 | 0.004 |
| p.Arg116Asn | substitution | 116 | Arg | Asn | 0.125 | 0.008 |
| p.Arg116Asp | substitution | 116 | Arg | Asp | 0.180 | 0.006 |
| p.Arg116Cys | substitution | 116 | Arg | Cys | 0.524 | 0.004 |
| p.Arg116Gln | substitution | 116 | Arg | Gln | 0.264 | 0.016 |
| p.Arg116Glu | substitution | 116 | Arg | Glu | 0.315 | 0.010 |
| p.Arg116Gly | substitution | 116 | Arg | Gly | 0.470 | 0.014 |
| p.Arg116His | substitution | 116 | Arg | His | 0.146 | 0.010 |
| p.Arg116Ile | substitution | 116 | Arg | Ile | 0.000 | 0.006 |
| p.Arg116Leu | substitution | 116 | Arg | Leu | 0.107 | 0.005 |
| p.Arg116Lys | substitution | 116 | Arg | Lys | 0.059 | 0.005 |
| p.Arg116Met | substitution | 116 | Arg | Met | 0.032 | 0.009 |
| p.Arg116Pro | substitution | 116 | Arg | Pro | 0.163 | 0.027 |
| p.Arg116Ser | substitution | 116 | Arg | Ser | 0.611 | 0.020 |
| p.Arg116Ter | nonsense | 116 | Arg | Ter | 0.103 | 0.041 |
| p.Arg116Thr | substitution | 116 | Arg | Thr | 0.203 | 0.004 |
| p.Arg116Trp | substitution | 116 | Arg | Trp | 0.184 | 0.005 |
| p.Arg116Tyr | substitution | 116 | Arg | Tyr | 0.169 | 0.006 |
| p.Arg116Val | substitution | 116 | Arg | Val | 0.000 | 0.007 |
| p.Arg149= | synonymous | 149 | Arg | NA | 0.967 | 0.021 |
| p.Arg149Ala | substitution | 149 | Arg | Ala | 0.000 | 0.030 |
| p.Arg149Asn | substitution | 149 | Arg | Asn | 0.000 | 0.157 |

| | | | | | | | |
|---|---|---|---|---|---|---|---|
| p.Arg149Asp | substitution | 149 | Arg | Asp | | 0.150 | 0.021 |
| p.Arg149Cys | substitution | 149 | Arg | Cys | | 0.018 | 0.020 |
| p.Arg149Gln | substitution | 149 | Arg | Gln | | 0.004 | 0.014 |
| p.Arg149Glu | substitution | 149 | Arg | Glu | | 0.000 | 0.302 |
| p.Arg149Gly | substitution | 149 | Arg | Gly | | 0.000 | 0.036 |
| p.Arg149His | substitution | 149 | Arg | His | | 0.433 | 0.010 |
| p.Arg149Ile | substitution | 149 | Arg | Ile | | 0.149 | 0.180 |
| p.Arg149Leu | substitution | 149 | Arg | Leu | | 0.000 | 0.031 |
| p.Arg149Lys | substitution | 149 | Arg | Lys | | 0.029 | 0.018 |
| p.Arg149Met | substitution | 149 | Arg | Met | | 0.000 | 0.140 |
| p.Arg149Phe | substitution | 149 | Arg | Phe | | 0.501 | 0.136 |
| p.Arg149Pro | substitution | 149 | Arg | Pro | | 0.000 | 0.028 |
| p.Arg149Ser | substitution | 149 | Arg | Ser | | 0.000 | 0.014 |
| p.Arg149Ter | nonsense | 149 | Arg | Ter | | 0.000 | 0.104 |
| p.Arg149Thr | substitution | 149 | Arg | Thr | | 0.000 | 0.031 |
| p.Arg149Trp | substitution | 149 | Arg | Trp | | 0.047 | 0.013 |
| p.Arg149Tyr | substitution | 149 | Arg | Tyr | | 0.565 | 0.236 |
| p.Arg149Val | substitution | 149 | Arg | Val | | 0.000 | 0.136 |
| p.Arg150= | synonymous | 150 | Arg | NA | | 0.849 | 0.012 |
| p.Arg150Asn | substitution | 150 | Arg | Asn | | 0.000 | 0.542 |
| p.Arg150Cys | substitution | 150 | Arg | Cys | | 0.437 | 0.011 |
| p.Arg150Gln | substitution | 150 | Arg | Gln | | 0.000 | 0.105 |
| p.Arg150Glu | substitution | 150 | Arg | Glu | | 0.000 | 0.016 |
| p.Arg150Gly | substitution | 150 | Arg | Gly | | 0.163 | 0.067 |
| p.Arg150His | substitution | 150 | Arg | His | | 0.266 | 0.028 |
| p.Arg150Ile | substitution | 150 | Arg | Ile | | 0.000 | 0.045 |
| p.Arg150Leu | substitution | 150 | Arg | Leu | | 0.000 | 0.028 |
| p.Arg150Lys | substitution | 150 | Arg | Lys | | 0.000 | 0.024 |
| p.Arg150Met | substitution | 150 | Arg | Met | | 0.374 | 0.014 |
| p.Arg150Phe | substitution | 150 | Arg | Phe | | 0.000 | 0.034 |
| p.Arg150Pro | substitution | 150 | Arg | Pro | | 0.073 | 0.137 |
| p.Arg150Ser | substitution | 150 | Arg | Ser | | 0.090 | 0.043 |
| p.Arg150Ter | nonsense | 150 | Arg | Ter | | 0.032 | 0.185 |
| p.Arg150Thr | substitution | 150 | Arg | Thr | | 0.000 | 0.201 |
| p.Arg150Trp | substitution | 150 | Arg | Trp | | 0.093 | 0.017 |
| p.Arg150Tyr | substitution | 150 | Arg | Tyr | | 0.000 | 0.014 |
| p.Arg150Val | substitution | 150 | Arg | Val | | 0.398 | 0.290 |
| p.Arg156= | synonymous | 156 | Arg | NA | | 1.009 | 0.016 |
| p.Arg156Ala | substitution | 156 | Arg | Ala | | 1.087 | 0.007 |
| p.Arg156Asn | substitution | 156 | Arg | Asn | | 0.933 | 0.017 |
| p.Arg156Asp | substitution | 156 | Arg | Asp | | 0.556 | 0.025 |
| p.Arg156Cys | substitution | 156 | Arg | Cys | | 0.724 | 0.006 |
| p.Arg156Gln | substitution | 156 | Arg | Gln | | 0.918 | 0.018 |

| | | | | | | | |
|---|---|---|---|---|---|---|---|
| p.Arg156Glu | substitution | 156 | Arg | Glu | | 0.861 | 0.008 |
| p.Arg156Gly | substitution | 156 | Arg | Gly | | 0.614 | 0.008 |
| p.Arg156His | substitution | 156 | Arg | His | | 1.067 | 0.006 |
| p.Arg156Ile | substitution | 156 | Arg | Ile | | 1.131 | 0.011 |
| p.Arg156Leu | substitution | 156 | Arg | Leu | | 0.763 | 0.017 |
| p.Arg156Lys | substitution | 156 | Arg | Lys | | 0.968 | 0.053 |
| p.Arg156Met | substitution | 156 | Arg | Met | | 1.122 | 0.008 |
| p.Arg156Phe | substitution | 156 | Arg | Phe | | 0.257 | 0.008 |
| p.Arg156Pro | substitution | 156 | Arg | Pro | | 0.996 | 0.030 |
| p.Arg156Ser | substitution | 156 | Arg | Ser | | 1.048 | 0.019 |
| p.Arg156Ter | nonsense | 156 | Arg | Ter | | 0.026 | 0.037 |
| p.Arg156Thr | substitution | 156 | Arg | Thr | | 0.977 | 0.024 |
| p.Arg156Trp | substitution | 156 | Arg | Trp | | 0.901 | 0.005 |
| p.Arg156Tyr | substitution | 156 | Arg | Tyr | | 0.372 | 0.013 |
| p.Arg156Val | substitution | 156 | Arg | Val | | 0.961 | 0.007 |
| p.Arg164= | synonymous | 164 | Arg | NA | | 1.118 | 0.010 |
| p.Arg164Ala | substitution | 164 | Arg | Ala | | 0.902 | 0.017 |
| p.Arg164Asn | substitution | 164 | Arg | Asn | | 0.881 | 0.004 |
| p.Arg164Asp | substitution | 164 | Arg | Asp | | 0.968 | 0.004 |
| p.Arg164Cys | substitution | 164 | Arg | Cys | | 0.821 | 0.004 |
| p.Arg164Gln | substitution | 164 | Arg | Gln | | 0.697 | 0.005 |
| p.Arg164Glu | substitution | 164 | Arg | Glu | | 1.129 | 0.004 |
| p.Arg164Gly | substitution | 164 | Arg | Gly | | 0.832 | 0.004 |
| p.Arg164His | substitution | 164 | Arg | His | | 1.078 | 0.006 |
| p.Arg164Ile | substitution | 164 | Arg | Ile | | 0.747 | 0.006 |
| p.Arg164Leu | substitution | 164 | Arg | Leu | | 0.839 | 0.005 |
| p.Arg164Lys | substitution | 164 | Arg | Lys | | 0.985 | 0.005 |
| p.Arg164Met | substitution | 164 | Arg | Met | | 0.921 | 0.011 |
| p.Arg164Phe | substitution | 164 | Arg | Phe | | 0.702 | 0.004 |
| p.Arg164Pro | substitution | 164 | Arg | Pro | | 0.531 | 0.025 |
| p.Arg164Ser | substitution | 164 | Arg | Ser | | 0.889 | 0.006 |
| p.Arg164Ter | nonsense | 164 | Arg | Ter | | 0.000 | 0.015 |
| p.Arg164Thr | substitution | 164 | Arg | Thr | | 0.954 | 0.005 |
| p.Arg164Trp | substitution | 164 | Arg | Trp | | 0.901 | 0.005 |
| p.Arg164Tyr | substitution | 164 | Arg | Tyr | | 0.542 | 0.006 |
| p.Arg164Val | substitution | 164 | Arg | Val | | 0.811 | 0.007 |
| p.Arg167= | synonymous | 167 | Arg | NA | | 1.064 | 0.024 |
| p.Arg167Ala | substitution | 167 | Arg | Ala | | 0.800 | 0.005 |
| p.Arg167Asn | substitution | 167 | Arg | Asn | | 0.625 | 0.008 |
| p.Arg167Asp | substitution | 167 | Arg | Asp | | 0.286 | 0.008 |
| p.Arg167Cys | substitution | 167 | Arg | Cys | | 0.223 | 0.006 |
| p.Arg167Gln | substitution | 167 | Arg | Gln | | 0.733 | 0.016 |
| p.Arg167Glu | substitution | 167 | Arg | Glu | | 0.000 | 0.009 |

| | | | | | | |
|---|---|---|---|---|---|---|
| p.Arg167Gly | substitution | 167 | Arg | Gly | 0.771 | 0.045 |
| p.Arg167His | substitution | 167 | Arg | His | 0.543 | 0.008 |
| p.Arg167Ile | substitution | 167 | Arg | Ile | 0.683 | 0.007 |
| p.Arg167Leu | substitution | 167 | Arg | Leu | 0.702 | 0.007 |
| p.Arg167Lys | substitution | 167 | Arg | Lys | 0.448 | 0.013 |
| p.Arg167Me | substitution | 167 | Arg | Met | 0.532 | 0.006 |
| p.Arg167Phe | substitution | 167 | Arg | Phe | 0.325 | 0.008 |
| p.Arg167Pro | substitution | 167 | Arg | Pro | 0.136 | 0.006 |
| p.Arg167Ser | substitution | 167 | Arg | Ser | 0.662 | 0.011 |
| p.Arg167Ter | nonsense | 167 | Arg | Ter | 0.122 | 0.008 |
| p.Arg167Thr | substitution | 167 | Arg | Thr | 0.919 | 0.006 |
| p.Arg167Trp | substitution | 167 | Arg | Trp | 0.522 | 0.088 |
| p.Arg167Tyr | substitution | 167 | Arg | Tyr | 0.700 | 0.003 |
| p.Arg167Val | substitution | 167 | Arg | Val | 0.693 | 0.008 |
| p.Arg173= | synonymous | 173 | Arg | NA | 0.983 | 0.109 |
| p.Arg173Ala | substitution | 173 | Arg | Ala | 0.000 | 0.013 |
| p.Arg173Asn | substitution | 173 | Arg | Asn | 0.048 | 0.009 |
| p.Arg173Asp | substitution | 173 | Arg | Asp | 0.000 | 0.046 |
| p.Arg173Cys | substitution | 173 | Arg | Cys | 0.000 | 0.011 |
| p.Arg173Gln | substitution | 173 | Arg | Gln | 0.000 | 0.300 |
| p.Arg173Gly | substitution | 173 | Arg | Gly | 0.000 | 0.008 |
| p.Arg173His | substitution | 173 | Arg | His | 0.063 | 0.012 |
| p.Arg173Ile | substitution | 173 | Arg | Ile | 0.000 | 0.024 |
| p.Arg173Leu | substitution | 173 | Arg | Leu | 0.046 | 0.024 |
| p.Arg173Lys | substitution | 173 | Arg | Lys | 0.152 | 0.013 |
| p.Arg173Me | substitution | 173 | Arg | Met | 0.000 | 0.010 |
| p.Arg173Phe | substitution | 173 | Arg | Phe | 0.000 | 0.012 |
| p.Arg173Pro | substitution | 173 | Arg | Pro | 0.000 | 0.022 |
| p.Arg173Ser | substitution | 173 | Arg | Ser | 0.000 | 0.020 |
| p.Arg173Ter | nonsense | 173 | Arg | Ter | 0.000 | 0.039 |
| p.Arg173Thr | substitution | 173 | Arg | Thr | 0.000 | 0.030 |
| p.Arg173Trp | substitution | 173 | Arg | Trp | 0.000 | 0.100 |
| p.Arg173Tyr | substitution | 173 | Arg | Tyr | 0.000 | 0.029 |
| p.Arg173Val | substitution | 173 | Arg | Val | 0.000 | 0.011 |
| p.Arg175= | synonymous | 175 | Arg | NA | 0.883 | 0.011 |
| p.Arg175Ala | substitution | 175 | Arg | Ala | 0.882 | 0.021 |
| p.Arg175Asn | substitution | 175 | Arg | Asn | 0.872 | 0.014 |
| p.Arg175Asp | substitution | 175 | Arg | Asp | 0.509 | 0.045 |
| p.Arg175Cys | substitution | 175 | Arg | Cys | 0.909 | 0.012 |
| p.Arg175Gln | substitution | 175 | Arg | Gln | 1.048 | 0.015 |
| p.Arg175Glu | substitution | 175 | Arg | Glu | 0.378 | 0.008 |
| p.Arg175Gly | substitution | 175 | Arg | Gly | 0.793 | 0.029 |
| p.Arg175His | substitution | 175 | Arg | His | 0.342 | 0.021 |

| | | | | | | | |
|---|---|---|---|---|---|---|---|
| p.Arg175Leu | substitution | 175 | Arg | Leu | | 0.756 | 0.010 |
| p.Arg175Lys | substitution | 175 | Arg | Lys | | 1.012 | 0.009 |
| p.Arg175Met | substitution | 175 | Arg | Met | | 1.032 | 0.006 |
| p.Arg175Phe | substitution | 175 | Arg | Phe | | 0.077 | 0.013 |
| p.Arg175Pro | substitution | 175 | Arg | Pro | | 0.160 | 0.106 |
| p.Arg175Ser | substitution | 175 | Arg | Ser | | 0.875 | 0.070 |
| p.Arg175Ter | nonsense | 175 | Arg | Ter | | 0.092 | 0.008 |
| p.Arg175Thr | substitution | 175 | Arg | Thr | | 0.688 | 0.045 |
| p.Arg175Trp | substitution | 175 | Arg | Trp | | 0.897 | 0.025 |
| p.Arg175Tyr | substitution | 175 | Arg | Tyr | | 0.429 | 0.030 |
| p.Arg175Val | substitution | 175 | Arg | Val | | 1.018 | 0.032 |
| p.Arg195= | synonymous | 195 | Arg | NA | | 1.065 | 0.032 |
| p.Arg195Ala | substitution | 195 | Arg | Ala | | 0.100 | 0.018 |
| p.Arg195Asn | substitution | 195 | Arg | Asn | | 0.000 | 0.016 |
| p.Arg195Asp | substitution | 195 | Arg | Asp | | 0.000 | 0.246 |
| p.Arg195Cys | substitution | 195 | Arg | Cys | | 0.248 | 0.093 |
| p.Arg195Gln | substitution | 195 | Arg | Gln | | 0.152 | 0.007 |
| p.Arg195Glu | substitution | 195 | Arg | Glu | | 0.112 | 0.018 |
| p.Arg195Gly | substitution | 195 | Arg | Gly | | 0.122 | 0.043 |
| p.Arg195His | substitution | 195 | Arg | His | | 0.783 | 0.058 |
| p.Arg195Ile | substitution | 195 | Arg | Ile | | 1.280 | 0.149 |
| p.Arg195Leu | substitution | 195 | Arg | Leu | | 0.868 | 0.043 |
| p.Arg195Lys | substitution | 195 | Arg | Lys | | 0.005 | 0.029 |
| p.Arg195Met | substitution | 195 | Arg | Met | | 0.000 | 0.036 |
| p.Arg195Phe | substitution | 195 | Arg | Phe | | 0.197 | 0.059 |
| p.Arg195Pro | substitution | 195 | Arg | Pro | | 0.117 | 0.022 |
| p.Arg195Ser | substitution | 195 | Arg | Ser | | 0.000 | 0.105 |
| p.Arg195Ter | nonsense | 195 | Arg | Ter | | 0.000 | 0.013 |
| p.Arg195Thr | substitution | 195 | Arg | Thr | | 0.000 | 0.019 |
| p.Arg195Trp | substitution | 195 | Arg | Trp | | 0.328 | 0.009 |
| p.Arg195Tyr | substitution | 195 | Arg | Tyr | | 0.805 | 0.013 |
| p.Arg195Val | substitution | 195 | Arg | Val | | 0.151 | 0.177 |
| p.Arg19= | synonymous | 19 | Arg | NA | | 0.687 | 0.000 |
| p.Arg19Cys | substitution | 19 | Arg | Cys | | 0.222 | 0.363 |
| p.Arg19Gly | substitution | 19 | Arg | Gly | | 1.024 | 0.066 |
| p.Arg19Ile | substitution | 19 | Arg | Ile | | 0.949 | 0.005 |
| p.Arg19Lys | substitution | 19 | Arg | Lys | | 1.015 | 0.008 |
| p.Arg19Ser | substitution | 19 | Arg | Ser | | 0.631 | 0.000 |
| p.Arg19Thr | substitution | 19 | Arg | Thr | | 0.810 | 0.056 |
| p.Arg201= | synonymous | 201 | Arg | NA | | 1.087 | 0.014 |
| p.Arg201Ala | substitution | 201 | Arg | Ala | | 0.000 | 0.006 |
| p.Arg201Asn | substitution | 201 | Arg | Asn | | 0.520 | 0.010 |
| p.Arg201Asp | substitution | 201 | Arg | Asp | | 0.952 | 0.005 |

| | | | | | | |
|---|---|---|---|---|---|---|
| p.Arg201Cys | substitution | 201 | Arg | Cys | 0.540 | 0.005 |
| p.Arg201Gln | substitution | 201 | Arg | Gln | 0.724 | 0.015 |
| p.Arg201Glu | substitution | 201 | Arg | Glu | 0.000 | 0.023 |
| p.Arg201Gly | substitution | 201 | Arg | Gly | 0.673 | 0.031 |
| p.Arg201His | substitution | 201 | Arg | His | 0.000 | 0.027 |
| p.Arg201Ile | substitution | 201 | Arg | Ile | 1.755 | 0.012 |
| p.Arg201Leu | substitution | 201 | Arg | Leu | 1.092 | 0.007 |
| p.Arg201Lys | substitution | 201 | Arg | Lys | 0.755 | 0.004 |
| p.Arg201Met | substitution | 201 | Arg | Met | 0.000 | 0.136 |
| p.Arg201Phe | substitution | 201 | Arg | Phe | 0.231 | 0.007 |
| p.Arg201Pro | substitution | 201 | Arg | Pro | 0.205 | 0.044 |
| p.Arg201Ser | substitution | 201 | Arg | Ser | 0.000 | 0.110 |
| p.Arg201Ter | nonsense | 201 | Arg | Ter | 0.043 | 0.041 |
| p.Arg201Thr | substitution | 201 | Arg | Thr | 0.473 | 0.023 |
| p.Arg201Trp | substitution | 201 | Arg | Trp | 0.719 | 0.004 |
| p.Arg201Tyr | substitution | 201 | Arg | Tyr | 1.345 | 0.003 |
| p.Arg201Val | substitution | 201 | Arg | Val | 0.798 | 0.018 |
| p.Arg225= | synonymous | 225 | Arg | NA | 0.935 | 0.020 |
| p.Arg225Ala | substitution | 225 | Arg | Ala | 0.028 | 0.048 |
| p.Arg225Asn | substitution | 225 | Arg | Asn | 0.096 | 0.004 |
| p.Arg225Asp | substitution | 225 | Arg | Asp | 0.496 | 0.002 |
| p.Arg225Cys | substitution | 225 | Arg | Cys | 0.859 | 0.001 |
| p.Arg225Gln | substitution | 225 | Arg | Gln | 0.000 | 0.137 |
| p.Arg225Glu | substitution | 225 | Arg | Glu | 0.607 | 0.007 |
| p.Arg225Gly | substitution | 225 | Arg | Gly | 0.000 | 0.093 |
| p.Arg225Ile | substitution | 225 | Arg | Ile | 0.245 | 0.159 |
| p.Arg225Leu | substitution | 225 | Arg | Leu | 0.758 | 0.009 |
| p.Arg225Lys | substitution | 225 | Arg | Lys | 0.133 | 0.038 |
| p.Arg225Met | substitution | 225 | Arg | Met | 0.985 | 0.001 |
| p.Arg225Phe | substitution | 225 | Arg | Phe | 0.311 | 0.003 |
| p.Arg225Pro | substitution | 225 | Arg | Pro | 1.063 | 0.030 |
| p.Arg225Ser | substitution | 225 | Arg | Ser | 0.633 | 0.032 |
| p.Arg225Ter | nonsense | 225 | Arg | Ter | 0.000 | 0.104 |
| p.Arg225Thr | substitution | 225 | Arg | Thr | 0.541 | 0.015 |
| p.Arg225Tyr | substitution | 225 | Arg | Tyr | 0.712 | 0.002 |
| p.Arg225Val | substitution | 225 | Arg | Val | 0.577 | 0.017 |
| p.Arg22= | synonymous | 22 | Arg | NA | 1.200 | 0.012 |
| p.Arg22Ala | substitution | 22 | Arg | Ala | 0.836 | 0.009 |
| p.Arg22Asn | substitution | 22 | Arg | Asn | 0.830 | 0.008 |
| p.Arg22Asp | substitution | 22 | Arg | Asp | 0.297 | 0.002 |
| p.Arg22Cys | substitution | 22 | Arg | Cys | 0.933 | 0.009 |
| p.Arg22Gln | substitution | 22 | Arg | Gln | 0.839 | 0.001 |
| p.Arg22Glu | substitution | 22 | Arg | Glu | 0.786 | 0.002 |

| | | | | | | |
|---|---|---|---|---|---|---|
| p.Arg22Gly | substitution | 22 | Arg | Gly | 0.631 | 0.003 |
| p.Arg22His | substitution | 22 | Arg | His | 1.713 | 0.021 |
| p.Arg22Ile | substitution | 22 | Arg | Ile | 1.159 | 0.003 |
| p.Arg22Leu | substitution | 22 | Arg | Leu | 1.161 | 0.001 |
| p.Arg22Lys | substitution | 22 | Arg | Lys | 1.006 | 0.002 |
| p.Arg22Met | substitution | 22 | Arg | Met | 0.239 | 0.003 |
| p.Arg22Phe | substitution | 22 | Arg | Phe | 0.783 | 0.002 |
| p.Arg22Pro | substitution | 22 | Arg | Pro | 0.379 | 0.002 |
| p.Arg22Ser | substitution | 22 | Arg | Ser | 0.914 | 0.002 |
| p.Arg22Ter | nonsense | 22 | Arg | Ter | 0.000 | 0.023 |
| p.Arg22Thr | substitution | 22 | Arg | Thr | 0.872 | 0.013 |
| p.Arg22Trp | substitution | 22 | Arg | Trp | 1.210 | 0.001 |
| p.Arg22Tyr | substitution | 22 | Arg | Tyr | 0.711 | 0.005 |
| p.Arg22Val | substitution | 22 | Arg | Val | 1.123 | 0.007 |
| p.Arg246= | synonymous | 246 | Arg | NA | 1.054 | 0.006 |
| p.Arg246Ala | substitution | 246 | Arg | Ala | 1.189 | 0.054 |
| p.Arg246Asn | substitution | 246 | Arg | Asn | 0.606 | 0.018 |
| p.Arg246Asp | substitution | 246 | Arg | Asp | 0.000 | 0.017 |
| p.Arg246Cys | substitution | 246 | Arg | Cys | 0.281 | 0.032 |
| p.Arg246Gln | substitution | 246 | Arg | Gln | 1.152 | 0.001 |
| p.Arg246Glu | substitution | 246 | Arg | Glu | 0.000 | 0.104 |
| p.Arg246Gly | substitution | 246 | Arg | Gly | 0.804 | 0.152 |
| p.Arg246His | substitution | 246 | Arg | His | 0.316 | 0.012 |
| p.Arg246Ile | substitution | 246 | Arg | Ile | 0.428 | 0.029 |
| p.Arg246Leu | substitution | 246 | Arg | Leu | 0.799 | 0.030 |
| p.Arg246Lys | substitution | 246 | Arg | Lys | 0.405 | 0.009 |
| p.Arg246Met | substitution | 246 | Arg | Met | 0.718 | 0.001 |
| p.Arg246Phe | substitution | 246 | Arg | Phe | 0.240 | 0.068 |
| p.Arg246Pro | substitution | 246 | Arg | Pro | 0.283 | 0.049 |
| p.Arg246Ser | substitution | 246 | Arg | Ser | 0.998 | 0.047 |
| p.Arg246Ter | nonsense | 246 | Arg | Ter | 0.000 | 0.038 |
| p.Arg246Thr | substitution | 246 | Arg | Thr | 0.647 | 0.004 |
| p.Arg246Trp | substitution | 246 | Arg | Trp | 0.387 | 0.004 |
| p.Arg246Tyr | substitution | 246 | Arg | Tyr | 0.388 | 0.013 |
| p.Arg246Val | substitution | 246 | Arg | Val | 1.016 | 0.124 |
| p.Arg251= | synonymous | 251 | Arg | NA | 0.902 | 0.039 |
| p.Arg251Ala | substitution | 251 | Arg | Ala | 0.000 | 0.217 |
| p.Arg251Asn | substitution | 251 | Arg | Asn | 0.332 | 0.365 |
| p.Arg251Asp | substitution | 251 | Arg | Asp | 0.000 | 0.013 |
| p.Arg251Cys | substitution | 251 | Arg | Cys | 0.176 | 0.238 |
| p.Arg251Gln | substitution | 251 | Arg | Gln | 0.000 | 0.119 |
| p.Arg251Glu | substitution | 251 | Arg | Glu | 0.072 | 0.020 |
| p.Arg251Gly | substitution | 251 | Arg | Gly | 0.000 | 0.029 |

| | | | | | | |
|---|---|---|---|---|---|---|
| p.Arg251His | substitution | 251 | Arg | His | 0.000 | 0.044 |
| p.Arg251Ile | substitution | 251 | Arg | Ile | 0.053 | 0.022 |
| p.Arg251Leu | substitution | 251 | Arg | Leu | 0.000 | 0.008 |
| p.Arg251Lys | substitution | 251 | Arg | Lys | 0.953 | 0.002 |
| p.Arg251Met | substitution | 251 | Arg | Met | 0.835 | 0.006 |
| p.Arg251Phe | substitution | 251 | Arg | Phe | 0.032 | 0.150 |
| p.Arg251Pro | substitution | 251 | Arg | Pro | 0.363 | 0.028 |
| p.Arg251Ser | substitution | 251 | Arg | Ser | 0.418 | 0.090 |
| p.Arg251Ter | nonsense | 251 | Arg | Ter | 0.000 | 0.011 |
| p.Arg251Thr | substitution | 251 | Arg | Thr | 0.423 | 0.082 |
| p.Arg251Trp | substitution | 251 | Arg | Trp | 0.827 | 0.005 |
| p.Arg251Tyr | substitution | 251 | Arg | Tyr | 0.000 | 0.007 |
| p.Arg251Val | substitution | 251 | Arg | Val | 0.641 | 0.252 |
| p.Arg255= | synonymous | 255 | Arg | NA | 0.549 | 0.066 |
| p.Arg255Ala | substitution | 255 | Arg | Ala | 0.311 | 0.099 |
| p.Arg255Asn | substitution | 255 | Arg | Asn | 0.113 | 0.013 |
| p.Arg255Gly | substitution | 255 | Arg | Gly | 1.094 | 0.010 |
| p.Arg255Leu | substitution | 255 | Arg | Leu | 0.000 | 0.217 |
| p.Arg255Lys | substitution | 255 | Arg | Lys | 0.370 | 0.176 |
| p.Arg255Met | substitution | 255 | Arg | Met | 0.996 | 0.003 |
| p.Arg255Pro | substitution | 255 | Arg | Pro | 0.076 | 0.214 |
| p.Arg255Ser | substitution | 255 | Arg | Ser | 0.883 | 0.036 |
| p.Arg255Trp | substitution | 255 | Arg | Trp | 0.973 | 0.168 |
| p.Arg26= | synonymous | 26 | Arg | NA | 0.916 | 0.004 |
| p.Arg26Ala | substitution | 26 | Arg | Ala | 0.121 | 0.039 |
| p.Arg26Asn | substitution | 26 | Arg | Asn | 0.000 | 0.002 |
| p.Arg26Asp | substitution | 26 | Arg | Asp | 0.000 | 0.002 |
| p.Arg26Cys | substitution | 26 | Arg | Cys | 0.000 | 0.001 |
| p.Arg26Gln | substitution | 26 | Arg | Gln | 0.000 | 0.018 |
| p.Arg26Glu | substitution | 26 | Arg | Glu | 0.000 | 0.048 |
| p.Arg26Gly | substitution | 26 | Arg | Gly | 0.000 | 0.022 |
| p.Arg26His | substitution | 26 | Arg | His | 0.000 | 0.005 |
| p.Arg26Ile | substitution | 26 | Arg | Ile | 0.000 | 0.017 |
| p.Arg26Leu | substitution | 26 | Arg | Leu | 0.000 | 0.004 |
| p.Arg26Lys | substitution | 26 | Arg | Lys | 0.000 | 0.005 |
| p.Arg26Met | substitution | 26 | Arg | Met | 0.000 | 0.050 |
| p.Arg26Phe | substitution | 26 | Arg | Phe | 0.000 | 0.005 |
| p.Arg26Pro | substitution | 26 | Arg | Pro | 0.000 | 0.008 |
| p.Arg26Ser | substitution | 26 | Arg | Ser | 0.000 | 0.013 |
| p.Arg26Ter | nonsense | 26 | Arg | Ter | 0.000 | 0.004 |
| p.Arg26Thr | substitution | 26 | Arg | Thr | 0.000 | 0.026 |
| p.Arg26Trp | substitution | 26 | Arg | Trp | 0.000 | 0.004 |
| p.Arg26Tyr | substitution | 26 | Arg | Tyr | 0.000 | 0.006 |

| | | | | | | |
|---|---|---|---|---|---|---|
| p.Arg26Val | substitution | 26 | Arg | Val | 0.000 | 0.026 |
| p.Arg321= | synonymous | 321 | Arg | NA | 1.074 | 0.025 |
| p.Arg321Ala | substitution | 321 | Arg | Ala | 0.955 | 0.044 |
| p.Arg321Asn | substitution | 321 | Arg | Asn | 0.431 | 0.007 |
| p.Arg321Asp | substitution | 321 | Arg | Asp | 0.189 | 0.006 |
| p.Arg321Cys | substitution | 321 | Arg | Cys | 0.583 | 0.469 |
| p.Arg321Gln | substitution | 321 | Arg | Gln | 0.440 | 0.004 |
| p.Arg321Gly | substitution | 321 | Arg | Gly | 0.993 | 0.018 |
| p.Arg321His | substitution | 321 | Arg | His | 0.863 | 0.005 |
| p.Arg321Ile | substitution | 321 | Arg | Ile | 0.818 | 0.016 |
| p.Arg321Leu | substitution | 321 | Arg | Leu | 0.886 | 0.005 |
| p.Arg321Lys | substitution | 321 | Arg | Lys | 1.366 | 0.003 |
| p.Arg321Met | substitution | 321 | Arg | Met | 0.537 | 0.005 |
| p.Arg321Phe | substitution | 321 | Arg | Phe | 1.127 | 0.001 |
| p.Arg321Pro | substitution | 321 | Arg | Pro | 0.546 | 0.223 |
| p.Arg321Ser | substitution | 321 | Arg | Ser | 0.962 | 0.045 |
| p.Arg321Ter | nonsense | 321 | Arg | Ter | 0.023 | 0.014 |
| p.Arg321Thr | substitution | 321 | Arg | Thr | 0.752 | 0.160 |
| p.Arg321Trp | substitution | 321 | Arg | Trp | 0.000 | 0.005 |
| p.Arg321Val | substitution | 321 | Arg | Val | 0.052 | 0.030 |
| p.Arg325= | synonymous | 325 | Arg | NA | 0.819 | 0.068 |
| p.Arg325Ala | substitution | 325 | Arg | Ala | 0.731 | 0.060 |
| p.Arg325Asn | substitution | 325 | Arg | Asn | 0.294 | 0.015 |
| p.Arg325Asp | substitution | 325 | Arg | Asp | 0.936 | 0.002 |
| p.Arg325Cys | substitution | 325 | Arg | Cys | 0.491 | 0.049 |
| p.Arg325Gln | substitution | 325 | Arg | Gln | 1.128 | 0.042 |
| p.Arg325Glu | substitution | 325 | Arg | Glu | 0.886 | 0.008 |
| p.Arg325Gly | substitution | 325 | Arg | Gly | 1.070 | 0.004 |
| p.Arg325His | substitution | 325 | Arg | His | 1.049 | 0.003 |
| p.Arg325Ile | substitution | 325 | Arg | Ile | 1.240 | 0.052 |
| p.Arg325Leu | substitution | 325 | Arg | Leu | 1.210 | 0.007 |
| p.Arg325Lys | substitution | 325 | Arg | Lys | 0.622 | 0.007 |
| p.Arg325Met | substitution | 325 | Arg | Met | 0.000 | 0.152 |
| p.Arg325Phe | substitution | 325 | Arg | Phe | 1.134 | 0.001 |
| p.Arg325Pro | substitution | 325 | Arg | Pro | 1.168 | 0.047 |
| p.Arg325Ser | substitution | 325 | Arg | Ser | 0.724 | 0.050 |
| p.Arg325Ter | nonsense | 325 | Arg | Ter | 0.033 | 0.048 |
| p.Arg325Thr | substitution | 325 | Arg | Thr | 0.693 | 0.012 |
| p.Arg325Trp | substitution | 325 | Arg | Trp | 1.010 | 0.016 |
| p.Arg325Tyr | substitution | 325 | Arg | Tyr | 0.567 | 0.004 |
| p.Arg325Val | substitution | 325 | Arg | Val | 0.022 | 0.071 |
| p.Arg32= | synonymous | 32 | Arg | NA | 0.937 | 0.001 |
| p.Arg32Ala | substitution | 32 | Arg | Ala | 0.660 | 0.003 |

| | | | | | | |
|---|---|---|---|---|---|---|
| p.Arg32Asn | substitution | 32 | Arg | Asn | 1.049 | 0.001 |
| p.Arg32Asp | substitution | 32 | Arg | Asp | 0.264 | 0.001 |
| p.Arg32Cys | substitution | 32 | Arg | Cys | 0.908 | 0.008 |
| p.Arg32Gln | substitution | 32 | Arg | Gln | 1.140 | 0.000 |
| p.Arg32Glu | substitution | 32 | Arg | Glu | 0.586 | 0.001 |
| p.Arg32Gly | substitution | 32 | Arg | Gly | 0.896 | 0.000 |
| p.Arg32His | substitution | 32 | Arg | His | 0.595 | 0.004 |
| p.Arg32Ile | substitution | 32 | Arg | Ile | 1.087 | 0.004 |
| p.Arg32Leu | substitution | 32 | Arg | Leu | 0.689 | 0.009 |
| p.Arg32Lys | substitution | 32 | Arg | Lys | 0.673 | 0.001 |
| p.Arg32Met | substitution | 32 | Arg | Met | 0.944 | 0.001 |
| p.Arg32Phe | substitution | 32 | Arg | Phe | 0.555 | 0.001 |
| p.Arg32Pro | substitution | 32 | Arg | Pro | 0.000 | 0.010 |
| p.Arg32Ser | substitution | 32 | Arg | Ser | 0.691 | 0.008 |
| p.Arg32Ter | nonsense | 32 | Arg | Ter | 0.000 | 0.002 |
| p.Arg32Thr | substitution | 32 | Arg | Thr | 0.507 | 0.006 |
| p.Arg32Trp | substitution | 32 | Arg | Trp | 0.747 | 0.000 |
| p.Arg32Tyr | substitution | 32 | Arg | Tyr | 1.143 | 0.001 |
| p.Arg32Val | substitution | 32 | Arg | Val | 0.754 | 0.011 |
| p.Arg355= | synonymous | 355 | Arg | NA | 1.042 | 0.015 |
| p.Arg355Ala | substitution | 355 | Arg | Ala | 0.104 | 0.213 |
| p.Arg355Asp | substitution | 355 | Arg | Asp | 1.423 | 0.079 |
| p.Arg355Cys | substitution | 355 | Arg | Cys | 0.000 | 0.128 |
| p.Arg355Gln | substitution | 355 | Arg | Gln | 1.034 | 0.007 |
| p.Arg355Gly | substitution | 355 | Arg | Gly | 1.242 | 0.357 |
| p.Arg355Leu | substitution | 355 | Arg | Leu | 0.815 | 0.009 |
| p.Arg355Lys | substitution | 355 | Arg | Lys | 0.347 | 0.166 |
| p.Arg355Met | substitution | 355 | Arg | Met | 1.396 | 0.003 |
| p.Arg355Pro | substitution | 355 | Arg | Pro | 1.415 | 0.148 |
| p.Arg355Ser | substitution | 355 | Arg | Ser | 1.266 | 0.013 |
| p.Arg355Thr | substitution | 355 | Arg | Thr | 0.267 | 0.139 |
| p.Arg355Trp | substitution | 355 | Arg | Trp | 1.369 | 0.004 |
| p.Arg355Tyr | substitution | 355 | Arg | Tyr | 0.630 | 0.012 |
| p.Arg355Val | substitution | 355 | Arg | Val | 1.050 | 0.084 |
| p.Asn118= | synonymous | 118 | Asn | NA | 0.980 | 0.004 |
| p.Asn118Ala | substitution | 118 | Asn | Ala | 1.112 | 0.014 |
| p.Asn118Arg | substitution | 118 | Asn | Arg | 0.890 | 0.013 |
| p.Asn118Asp | substitution | 118 | Asn | Asp | 1.249 | 0.029 |
| p.Asn118Cys | substitution | 118 | Asn | Cys | 1.067 | 0.012 |
| p.Asn118Gln | substitution | 118 | Asn | Gln | 0.710 | 0.015 |
| p.Asn118Glu | substitution | 118 | Asn | Glu | 1.160 | 0.013 |
| p.Asn118Gly | substitution | 118 | Asn | Gly | 0.210 | 0.035 |
| p.Asn118His | substitution | 118 | Asn | His | 0.916 | 0.071 |

| | | | | | | |
|---|---|---|---|---|---|---|
| p.Asn118Ile | substitution | 118 | Asn | Ile | 1.000 | 0.069 |
| p.Asn118Leu | substitution | 118 | Asn | Leu | 0.459 | 0.103 |
| p.Asn118Lys | substitution | 118 | Asn | Lys | 0.658 | 0.022 |
| p.Asn118Me | substitution | 118 | Asn | Met | 0.910 | 0.010 |
| p.Asn118Phe | substitution | 118 | Asn | Phe | 0.777 | 0.128 |
| p.Asn118Pro | substitution | 118 | Asn | Pro | 0.850 | 0.046 |
| p.Asn118Ser | substitution | 118 | Asn | Ser | 1.121 | 0.027 |
| p.Asn118Ter | nonsense | 118 | Asn | Ter | 0.158 | 0.018 |
| p.Asn118Thr | substitution | 118 | Asn | Thr | 1.173 | 0.009 |
| p.Asn118Trp | substitution | 118 | Asn | Trp | 0.812 | 0.010 |
| p.Asn118Val | substitution | 118 | Asn | Val | 0.948 | 0.007 |
| p.Asn169= | synonymous | 169 | Asn | NA | 1.080 | 0.007 |
| p.Asn169Ala | substitution | 169 | Asn | Ala | 0.445 | 0.012 |
| p.Asn169Arg | substitution | 169 | Asn | Arg | 0.841 | 0.006 |
| p.Asn169Asp | substitution | 169 | Asn | Asp | 0.471 | 0.017 |
| p.Asn169Cys | substitution | 169 | Asn | Cys | 0.239 | 0.032 |
| p.Asn169Gln | substitution | 169 | Asn | Gln | 0.269 | 0.018 |
| p.Asn169Glu | substitution | 169 | Asn | Glu | 0.375 | 0.006 |
| p.Asn169Gly | substitution | 169 | Asn | Gly | 0.372 | 0.028 |
| p.Asn169His | substitution | 169 | Asn | His | 0.906 | 0.031 |
| p.Asn169Ile | substitution | 169 | Asn | Ile | 0.100 | 0.099 |
| p.Asn169Leu | substitution | 169 | Asn | Leu | 0.733 | 0.034 |
| p.Asn169Lys | substitution | 169 | Asn | Lys | 0.893 | 0.027 |
| p.Asn169Phe | substitution | 169 | Asn | Phe | 0.363 | 0.083 |
| p.Asn169Pro | substitution | 169 | Asn | Pro | 0.000 | 0.067 |
| p.Asn169Ser | substitution | 169 | Asn | Ser | 0.920 | 0.010 |
| p.Asn169Ter | nonsense | 169 | Asn | Ter | 0.001 | 0.008 |
| p.Asn169Thr | substitution | 169 | Asn | Thr | 0.673 | 0.015 |
| p.Asn169Trp | substitution | 169 | Asn | Trp | 0.309 | 0.015 |
| p.Asn169Tyr | substitution | 169 | Asn | Tyr | 0.753 | 0.050 |
| p.Asn169Val | substitution | 169 | Asn | Val | 0.944 | 0.016 |
| p.Asn171= | synonymous | 171 | Asn | NA | 1.041 | 0.006 |
| p.Asn171Ala | substitution | 171 | Asn | Ala | 0.792 | 0.016 |
| p.Asn171Arg | substitution | 171 | Asn | Arg | 0.689 | 0.009 |
| p.Asn171Asp | substitution | 171 | Asn | Asp | 1.177 | 0.053 |
| p.Asn171Gln | substitution | 171 | Asn | Gln | 0.946 | 0.009 |
| p.Asn171Glu | substitution | 171 | Asn | Glu | 1.002 | 0.005 |
| p.Asn171Gly | substitution | 171 | Asn | Gly | 0.869 | 0.011 |
| p.Asn171His | substitution | 171 | Asn | His | 0.829 | 0.052 |
| p.Asn171Ile | substitution | 171 | Asn | Ile | 0.890 | 0.025 |
| p.Asn171Leu | substitution | 171 | Asn | Leu | 0.988 | 0.010 |
| p.Asn171Lys | substitution | 171 | Asn | Lys | 1.038 | 0.019 |
| p.Asn171Me | substitution | 171 | Asn | Met | 0.853 | 0.016 |

| | | | | | | | |
|---|---|---|---|---|---|---|---|
| p.Asn171Phe | substitution | 171 | Asn | Phe | | 0.203 | 0.012 |
| p.Asn171Pro | substitution | 171 | Asn | Pro | | 0.516 | 0.129 |
| p.Asn171Ser | substitution | 171 | Asn | Ser | | 0.932 | 0.006 |
| p.Asn171Ter | nonsense | 171 | Asn | Ter | | 0.000 | 0.008 |
| p.Asn171Thr | substitution | 171 | Asn | Thr | | 0.918 | 0.014 |
| p.Asn171Trp | substitution | 171 | Asn | Trp | | 1.101 | 0.007 |
| p.Asn171Tyr | substitution | 171 | Asn | Tyr | | 1.065 | 0.010 |
| p.Asn171Val | substitution | 171 | Asn | Val | | 0.550 | 0.323 |
| p.Asn200= | synonymous | 200 | Asn | NA | | 0.789 | 0.002 |
| p.Asn200Ala | substitution | 200 | Asn | Ala | | 0.914 | 0.022 |
| p.Asn200Arg | substitution | 200 | Asn | Arg | | 0.101 | 0.017 |
| p.Asn200Asp | substitution | 200 | Asn | Asp | | 1.153 | 0.001 |
| p.Asn200Cys | substitution | 200 | Asn | Cys | | 0.950 | 0.032 |
| p.Asn200Gln | substitution | 200 | Asn | Gln | | 0.000 | 0.064 |
| p.Asn200Glu | substitution | 200 | Asn | Glu | | 0.586 | 0.003 |
| p.Asn200Gly | substitution | 200 | Asn | Gly | | 0.731 | 0.033 |
| p.Asn200His | substitution | 200 | Asn | His | | 1.187 | 0.046 |
| p.Asn200Ile | substitution | 200 | Asn | Ile | | 0.998 | 0.021 |
| p.Asn200Leu | substitution | 200 | Asn | Leu | | 0.288 | 0.022 |
| p.Asn200Lys | substitution | 200 | Asn | Lys | | 1.110 | 0.002 |
| p.Asn200Me | substitution | 200 | Asn | Met | | 0.837 | 0.003 |
| p.Asn200Phe | substitution | 200 | Asn | Phe | | 1.281 | 0.010 |
| p.Asn200Ser | substitution | 200 | Asn | Ser | | 0.744 | 0.073 |
| p.Asn200Ter | nonsense | 200 | Asn | Ter | | 0.000 | 0.014 |
| p.Asn200Thr | substitution | 200 | Asn | Thr | | 1.043 | 0.107 |
| p.Asn200Trp | substitution | 200 | Asn | Trp | | 1.001 | 0.008 |
| p.Asn200Tyr | substitution | 200 | Asn | Tyr | | 0.673 | 0.034 |
| p.Asn200Val | substitution | 200 | Asn | Val | | 0.494 | 0.033 |
| p.Asn322= | synonymous | 322 | Asn | NA | | 1.204 | 0.002 |
| p.Asn322Ala | substitution | 322 | Asn | Ala | | 0.000 | 0.074 |
| p.Asn322Arg | substitution | 322 | Asn | Arg | | 0.997 | 0.195 |
| p.Asn322Asp | substitution | 322 | Asn | Asp | | 1.645 | 0.148 |
| p.Asn322Cys | substitution | 322 | Asn | Cys | | 1.251 | 0.011 |
| p.Asn322Gln | substitution | 322 | Asn | Gln | | 0.804 | 0.005 |
| p.Asn322Glu | substitution | 322 | Asn | Glu | | 1.013 | 0.003 |
| p.Asn322Gly | substitution | 322 | Asn | Gly | | 0.000 | 0.066 |
| p.Asn322His | substitution | 322 | Asn | His | | 0.711 | 0.221 |
| p.Asn322Ile | substitution | 322 | Asn | Ile | | 0.689 | 0.045 |
| p.Asn322Leu | substitution | 322 | Asn | Leu | | 0.440 | 0.071 |
| p.Asn322Lys | substitution | 322 | Asn | Lys | | 0.777 | 0.013 |
| p.Asn322Me | substitution | 322 | Asn | Met | | 0.786 | 0.004 |
| p.Asn322Phe | substitution | 322 | Asn | Phe | | 1.117 | 0.006 |
| p.Asn322Pro | substitution | 322 | Asn | Pro | | 0.694 | 0.008 |

| | | | | | | |
|---|---|---|---|---|---|---|
| p.Asn322Ser | substitution | 322 | Asn | Ser | 0.536 | 0.036 |
| p.Asn322Ter | nonsense | 322 | Asn | Ter | 0.000 | 0.062 |
| p.Asn322Thr | substitution | 322 | Asn | Thr | 0.519 | 0.109 |
| p.Asn322Trp | substitution | 322 | Asn | Trp | 1.152 | 0.003 |
| p.Asn322Tyr | substitution | 322 | Asn | Tyr | 0.676 | 0.018 |
| p.Asn322Val | substitution | 322 | Asn | Val | 0.865 | 0.008 |
| p.Asn333= | synonymous | 333 | Asn | NA | 1.431 | 0.008 |
| p.Asn333Asp | substitution | 333 | Asn | Asp | 0.000 | 0.063 |
| p.Asn333Cys | substitution | 333 | Asn | Cys | 0.000 | 0.035 |
| p.Asn333His | substitution | 333 | Asn | His | 1.178 | 0.143 |
| p.Asn333Ile | substitution | 333 | Asn | Ile | 1.239 | 0.006 |
| p.Asn333Lys | substitution | 333 | Asn | Lys | 1.202 | 0.022 |
| p.Asn333Ser | substitution | 333 | Asn | Ser | 0.000 | 0.126 |
| p.Asn333Thr | substitution | 333 | Asn | Thr | 0.000 | 0.224 |
| p.Asn333Tyr | substitution | 333 | Asn | Tyr | 0.301 | 0.167 |
| p.Asn340= | synonymous | 340 | Asn | NA | 0.944 | 0.005 |
| p.Asn340Ala | substitution | 340 | Asn | Ala | 0.138 | 0.019 |
| p.Asn340Arg | substitution | 340 | Asn | Arg | 0.625 | 0.136 |
| p.Asn340Asp | substitution | 340 | Asn | Asp | 0.953 | 0.220 |
| p.Asn340Cys | substitution | 340 | Asn | Cys | 0.000 | 0.027 |
| p.Asn340Gln | substitution | 340 | Asn | Gln | 1.254 | 0.003 |
| p.Asn340Glu | substitution | 340 | Asn | Glu | 0.000 | 0.143 |
| p.Asn340Gly | substitution | 340 | Asn | Gly | 0.906 | 0.086 |
| p.Asn340His | substitution | 340 | Asn | His | 0.000 | 0.036 |
| p.Asn340Ile | substitution | 340 | Asn | Ile | 1.290 | 0.005 |
| p.Asn340Leu | substitution | 340 | Asn | Leu | 0.675 | 0.038 |
| p.Asn340Lys | substitution | 340 | Asn | Lys | 0.974 | 0.005 |
| p.Asn340Me | substitution | 340 | Asn | Met | 1.346 | 0.053 |
| p.Asn340Phe | substitution | 340 | Asn | Phe | 0.556 | 0.014 |
| p.Asn340Pro | substitution | 340 | Asn | Pro | 0.000 | 0.021 |
| p.Asn340Ser | substitution | 340 | Asn | Ser | 0.720 | 0.330 |
| p.Asn340Ter | nonsense | 340 | Asn | Ter | 1.508 | 0.033 |
| p.Asn340Thr | substitution | 340 | Asn | Thr | 0.800 | 0.293 |
| p.Asn340Trp | substitution | 340 | Asn | Trp | 1.480 | 0.002 |
| p.Asn340Tyr | substitution | 340 | Asn | Tyr | 0.880 | 0.059 |
| p.Asn340Val | substitution | 340 | Asn | Val | 0.114 | 0.045 |
| p.Asn349= | synonymous | 349 | Asn | NA | 1.337 | 0.003 |
| p.Asn349Ala | substitution | 349 | Asn | Ala | 1.353 | 0.023 |
| p.Asn349Arg | substitution | 349 | Asn | Arg | 1.198 | 0.075 |
| p.Asn349Asp | substitution | 349 | Asn | Asp | 0.951 | 0.009 |
| p.Asn349Cys | substitution | 349 | Asn | Cys | 0.467 | 0.082 |
| p.Asn349Gln | substitution | 349 | Asn | Gln | 1.172 | 0.003 |
| p.Asn349Glu | substitution | 349 | Asn | Glu | 0.764 | 0.004 |

| | | | | | | |
|---|---|---|---|---|---|---|
| p.Asn349Gly | substitution | 349 | Asn | Gly | 0.900 | 0.285 |
| p.Asn349His | substitution | 349 | Asn | His | 0.236 | 0.113 |
| p.Asn349Ile | substitution | 349 | Asn | Ile | 0.430 | 0.023 |
| p.Asn349Leu | substitution | 349 | Asn | Leu | 1.017 | 0.012 |
| p.Asn349Lys | substitution | 349 | Asn | Lys | 1.048 | 0.024 |
| p.Asn349Me | substitution | 349 | Asn | Met | 0.894 | 0.009 |
| p.Asn349Pro | substitution | 349 | Asn | Pro | 0.820 | 0.285 |
| p.Asn349Ser | substitution | 349 | Asn | Ser | 0.517 | 0.123 |
| p.Asn349Ter | nonsense | 349 | Asn | Ter | 0.997 | 0.003 |
| p.Asn349Thr | substitution | 349 | Asn | Thr | 0.948 | 0.007 |
| p.Asn349Trp | substitution | 349 | Asn | Trp | 1.671 | 0.008 |
| p.Asn349Tyr | substitution | 349 | Asn | Tyr | 0.439 | 0.018 |
| p.Asn349Val | substitution | 349 | Asn | Val | 0.777 | 0.156 |
| p.Asn358= | synonymous | 358 | Asn | NA | 1.102 | 0.021 |
| p.Asn358Ala | substitution | 358 | Asn | Ala | 1.305 | 0.022 |
| p.Asn358Arg | substitution | 358 | Asn | Arg | 1.239 | 0.017 |
| p.Asn358Asp | substitution | 358 | Asn | Asp | 0.000 | 0.288 |
| p.Asn358Gly | substitution | 358 | Asn | Gly | 0.120 | 0.017 |
| p.Asn358Ile | substitution | 358 | Asn | Ile | 1.300 | 0.065 |
| p.Asn358Lys | substitution | 358 | Asn | Lys | 1.413 | 0.003 |
| p.Asn358Me | substitution | 358 | Asn | Met | 1.456 | 0.008 |
| p.Asn358Pro | substitution | 358 | Asn | Pro | 1.271 | 0.003 |
| p.Asn358Ser | substitution | 358 | Asn | Ser | 1.219 | 0.062 |
| p.Asn358Ter | nonsense | 358 | Asn | Ter | 1.424 | 0.046 |
| p.Asn358Thr | substitution | 358 | Asn | Thr | 1.403 | 0.077 |
| p.Asn358Tyr | substitution | 358 | Asn | Tyr | 1.123 | 0.350 |
| p.Asn88Ala | substitution | 88 | Asn | Ala | 0.806 | 0.022 |
| p.Asn88Arg | substitution | 88 | Asn | Arg | 0.885 | 0.009 |
| p.Asn88Asp | substitution | 88 | Asn | Asp | 0.499 | 0.009 |
| p.Asn88Cys | substitution | 88 | Asn | Cys | 1.054 | 0.005 |
| p.Asn88Glu | substitution | 88 | Asn | Glu | 0.671 | 0.006 |
| p.Asn88Gly | substitution | 88 | Asn | Gly | 0.700 | 0.043 |
| p.Asn88His | substitution | 88 | Asn | His | 0.841 | 0.006 |
| p.Asn88Ile | substitution | 88 | Asn | Ile | 0.879 | 0.003 |
| p.Asn88Leu | substitution | 88 | Asn | Leu | 0.955 | 0.015 |
| p.Asn88Lys | substitution | 88 | Asn | Lys | 1.130 | 0.004 |
| p.Asn88Met | substitution | 88 | Asn | Met | 1.004 | 0.006 |
| p.Asn88Phe | substitution | 88 | Asn | Phe | 0.944 | 0.002 |
| p.Asn88Pro | substitution | 88 | Asn | Pro | 0.548 | 0.012 |
| p.Asn88Ser | substitution | 88 | Asn | Ser | 0.900 | 0.010 |
| p.Asn88Ter | nonsense | 88 | Asn | Ter | 0.000 | 0.026 |
| p.Asn88Thr | substitution | 88 | Asn | Thr | 1.044 | 0.032 |
| p.Asn88Trp | substitution | 88 | Asn | Trp | 1.046 | 0.003 |

| | | | | | | |
|---|---|---|---|---|---|---|
| p.Asn88Tyr | substitution | 88 | Asn | Tyr | 0.985 | 0.003 |
| p.Asn88Val | substitution | 88 | Asn | Val | 0.476 | 0.019 |
| p.Asp121Ala | substitution | 121 | Asp | Ala | 0.000 | 0.009 |
| p.Asp121Arg | substitution | 121 | Asp | Arg | 0.000 | 0.007 |
| p.Asp121Asn | substitution | 121 | Asp | Asn | 0.091 | 0.008 |
| p.Asp121Cys | substitution | 121 | Asp | Cys | 0.789 | 0.003 |
| p.Asp121Gln | substitution | 121 | Asp | Gln | 0.000 | 0.017 |
| p.Asp121Glu | substitution | 121 | Asp | Glu | 0.406 | 0.038 |
| p.Asp121Gly | substitution | 121 | Asp | Gly | 0.127 | 0.009 |
| p.Asp121His | substitution | 121 | Asp | His | 0.009 | 0.011 |
| p.Asp121Ile | substitution | 121 | Asp | Ile | 0.000 | 0.007 |
| p.Asp121Leu | substitution | 121 | Asp | Leu | 0.034 | 0.004 |
| p.Asp121Lys | substitution | 121 | Asp | Lys | 0.000 | 0.011 |
| p.Asp121Me | substitution | 121 | Asp | Met | 0.000 | 0.006 |
| p.Asp121Phe | substitution | 121 | Asp | Phe | 0.000 | 0.010 |
| p.Asp121Pro | substitution | 121 | Asp | Pro | 0.011 | 0.012 |
| p.Asp121Ser | substitution | 121 | Asp | Ser | 0.103 | 0.024 |
| p.Asp121Ter | nonsense | 121 | Asp | Ter | 0.002 | 0.017 |
| p.Asp121Thr | substitution | 121 | Asp | Thr | 0.000 | 0.011 |
| p.Asp121Trp | substitution | 121 | Asp | Trp | 0.000 | 0.009 |
| p.Asp121Tyr | substitution | 121 | Asp | Tyr | 0.039 | 0.008 |
| p.Asp121Val | substitution | 121 | Asp | Val | 0.000 | 0.024 |
| p.Asp178= | synonymous | 178 | Asp | NA | 0.756 | 0.015 |
| p.Asp178Ala | substitution | 178 | Asp | Ala | 0.502 | 0.058 |
| p.Asp178Arg | substitution | 178 | Asp | Arg | 0.050 | 0.027 |
| p.Asp178Asn | substitution | 178 | Asp | Asn | 0.699 | 0.021 |
| p.Asp178Cys | substitution | 178 | Asp | Cys | 1.084 | 0.007 |
| p.Asp178Gln | substitution | 178 | Asp | Gln | 0.391 | 0.021 |
| p.Asp178Glu | substitution | 178 | Asp | Glu | 0.967 | 0.009 |
| p.Asp178Gly | substitution | 178 | Asp | Gly | 0.995 | 0.019 |
| p.Asp178His | substitution | 178 | Asp | His | 0.935 | 0.007 |
| p.Asp178Ile | substitution | 178 | Asp | Ile | 0.000 | 0.034 |
| p.Asp178Leu | substitution | 178 | Asp | Leu | 0.836 | 0.038 |
| p.Asp178Lys | substitution | 178 | Asp | Lys | 0.190 | 0.034 |
| p.Asp178Me | substitution | 178 | Asp | Met | 0.899 | 0.007 |
| p.Asp178Phe | substitution | 178 | Asp | Phe | 0.383 | 0.011 |
| p.Asp178Pro | substitution | 178 | Asp | Pro | 0.000 | 0.168 |
| p.Asp178Ser | substitution | 178 | Asp | Ser | 0.793 | 0.026 |
| p.Asp178Ter | nonsense | 178 | Asp | Ter | 0.000 | 0.088 |
| p.Asp178Thr | substitution | 178 | Asp | Thr | 0.560 | 0.014 |
| p.Asp178Trp | substitution | 178 | Asp | Trp | 0.396 | 0.010 |
| p.Asp178Tyr | substitution | 178 | Asp | Tyr | 1.001 | 0.011 |
| p.Asp178Val | substitution | 178 | Asp | Val | 0.838 | 0.008 |

| | | | | | | |
|---|---|---|---|---|---|---|
| p.Asp228= | synonymous | 228 | Asp | NA | 0.740 | 0.002 |
| p.Asp228Ala | substitution | 228 | Asp | Ala | 0.932 | 0.149 |
| p.Asp228Arg | substitution | 228 | Asp | Arg | 1.064 | 0.047 |
| p.Asp228Asn | substitution | 228 | Asp | Asn | 1.061 | 0.165 |
| p.Asp228Cys | substitution | 228 | Asp | Cys | 0.960 | 0.004 |
| p.Asp228Gln | substitution | 228 | Asp | Gln | 1.363 | 0.006 |
| p.Asp228Glu | substitution | 228 | Asp | Glu | 1.268 | 0.002 |
| p.Asp228Gly | substitution | 228 | Asp | Gly | 0.935 | 0.099 |
| p.Asp228His | substitution | 228 | Asp | His | 0.000 | 0.092 |
| p.Asp228Ile | substitution | 228 | Asp | Ile | 0.000 | 0.007 |
| p.Asp228Leu | substitution | 228 | Asp | Leu | 0.081 | 0.009 |
| p.Asp228Lys | substitution | 228 | Asp | Lys | 0.037 | 0.025 |
| p.Asp228Pro | substitution | 228 | Asp | Pro | 1.525 | 0.033 |
| p.Asp228Ser | substitution | 228 | Asp | Ser | 0.689 | 0.104 |
| p.Asp228Ter | nonsense | 228 | Asp | Ter | 0.000 | 0.013 |
| p.Asp228Thr | substitution | 228 | Asp | Thr | 0.041 | 0.039 |
| p.Asp228Trp | substitution | 228 | Asp | Trp | 0.000 | 0.189 |
| p.Asp228Tyr | substitution | 228 | Asp | Tyr | 0.000 | 0.042 |
| p.Asp228Val | substitution | 228 | Asp | Val | 0.166 | 0.012 |
| p.Asp230= | synonymous | 230 | Asp | NA | 0.149 | 0.447 |
| p.Asp230Ala | substitution | 230 | Asp | Ala | 0.317 | 0.029 |
| p.Asp230Arg | substitution | 230 | Asp | Arg | 0.994 | 0.009 |
| p.Asp230Asn | substitution | 230 | Asp | Asn | 0.853 | 0.143 |
| p.Asp230Cys | substitution | 230 | Asp | Cys | 0.000 | 0.083 |
| p.Asp230Gln | substitution | 230 | Asp | Gln | 1.379 | 0.001 |
| p.Asp230Glu | substitution | 230 | Asp | Glu | 1.044 | 0.006 |
| p.Asp230Gly | substitution | 230 | Asp | Gly | 0.505 | 0.039 |
| p.Asp230His | substitution | 230 | Asp | His | 0.000 | 0.197 |
| p.Asp230Ile | substitution | 230 | Asp | Ile | 1.325 | 0.004 |
| p.Asp230Leu | substitution | 230 | Asp | Leu | 0.000 | 0.026 |
| p.Asp230Lys | substitution | 230 | Asp | Lys | 0.715 | 0.003 |
| p.Asp230Me | substitution | 230 | Asp | Met | 1.483 | 0.003 |
| p.Asp230Phe | substitution | 230 | Asp | Phe | 0.000 | 0.120 |
| p.Asp230Pro | substitution | 230 | Asp | Pro | 0.885 | 0.164 |
| p.Asp230Ser | substitution | 230 | Asp | Ser | 1.074 | 0.023 |
| p.Asp230Ter | nonsense | 230 | Asp | Ter | 0.227 | 0.014 |
| p.Asp230Thr | substitution | 230 | Asp | Thr | 0.000 | 0.030 |
| p.Asp230Trp | substitution | 230 | Asp | Trp | 1.010 | 0.003 |
| p.Asp230Tyr | substitution | 230 | Asp | Tyr | 0.512 | 0.038 |
| p.Asp230Val | substitution | 230 | Asp | Val | 0.908 | 0.014 |
| p.Asp233Ala | substitution | 233 | Asp | Ala | 0.532 | 0.021 |
| p.Asp233Arg | substitution | 233 | Asp | Arg | 0.693 | 0.008 |
| p.Asp233Asn | substitution | 233 | Asp | Asn | 1.066 | 0.003 |

| | | | | | | |
|---|---|---|---|---|---|---|
| p.Asp233Cys | substitution | 233 | Asp | Cys | 0.856 | 0.003 |
| p.Asp233Gln | substitution | 233 | Asp | Gln | 0.703 | 0.002 |
| p.Asp233Glu | substitution | 233 | Asp | Glu | 0.908 | 0.002 |
| p.Asp233Gly | substitution | 233 | Asp | Gly | 0.537 | 0.010 |
| p.Asp233His | substitution | 233 | Asp | His | 1.003 | 0.003 |
| p.Asp233Ile | substitution | 233 | Asp | Ile | 0.722 | 0.002 |
| p.Asp233Leu | substitution | 233 | Asp | Leu | 0.054 | 0.043 |
| p.Asp233Lys | substitution | 233 | Asp | Lys | 0.776 | 0.004 |
| p.Asp233Me | substitution | 233 | Asp | Met | 0.556 | 0.013 |
| p.Asp233Pro | substitution | 233 | Asp | Pro | 0.387 | 0.094 |
| p.Asp233Ser | substitution | 233 | Asp | Ser | 0.864 | 0.106 |
| p.Asp233Ter | nonsense | 233 | Asp | Ter | 0.000 | 0.006 |
| p.Asp233Thr | substitution | 233 | Asp | Thr | 0.676 | 0.002 |
| p.Asp233Trp | substitution | 233 | Asp | Trp | 0.029 | 0.006 |
| p.Asp233Tyr | substitution | 233 | Asp | Tyr | 0.597 | 0.001 |
| p.Asp233Val | substitution | 233 | Asp | Val | 0.561 | 0.076 |
| p.Asp240Ala | substitution | 240 | Asp | Ala | 0.627 | 0.090 |
| p.Asp240Arg | substitution | 240 | Asp | Arg | 0.000 | 0.066 |
| p.Asp240Asn | substitution | 240 | Asp | Asn | 1.283 | 0.004 |
| p.Asp240Cys | substitution | 240 | Asp | Cys | 0.968 | 0.002 |
| p.Asp240Gln | substitution | 240 | Asp | Gln | 1.083 | 0.004 |
| p.Asp240Glu | substitution | 240 | Asp | Glu | 1.271 | 0.001 |
| p.Asp240Gly | substitution | 240 | Asp | Gly | 0.000 | 0.032 |
| p.Asp240Ile | substitution | 240 | Asp | Ile | 0.425 | 0.005 |
| p.Asp240Leu | substitution | 240 | Asp | Leu | 0.000 | 0.033 |
| p.Asp240Lys | substitution | 240 | Asp | Lys | 0.853 | 0.011 |
| p.Asp240Me | substitution | 240 | Asp | Met | 0.145 | 0.019 |
| p.Asp240Phe | substitution | 240 | Asp | Phe | 0.810 | 0.015 |
| p.Asp240Pro | substitution | 240 | Asp | Pro | 0.134 | 0.049 |
| p.Asp240Ser | substitution | 240 | Asp | Ser | 0.000 | 0.035 |
| p.Asp240Ter | nonsense | 240 | Asp | Ter | 0.030 | 0.077 |
| p.Asp240Thr | substitution | 240 | Asp | Thr | 0.421 | 0.008 |
| p.Asp240Trp | substitution | 240 | Asp | Trp | 0.000 | 0.022 |
| p.Asp240Tyr | substitution | 240 | Asp | Tyr | 1.072 | 0.006 |
| p.Asp240Val | substitution | 240 | Asp | Val | 0.930 | 0.191 |
| p.Asp273Ala | substitution | 273 | Asp | Ala | 0.146 | 0.244 |
| p.Asp273Arg | substitution | 273 | Asp | Arg | 1.126 | 0.033 |
| p.Asp273Asn | substitution | 273 | Asp | Asn | 1.436 | 0.013 |
| p.Asp273Glu | substitution | 273 | Asp | Glu | 0.904 | 0.026 |
| p.Asp273Gly | substitution | 273 | Asp | Gly | 0.000 | 0.532 |
| p.Asp273His | substitution | 273 | Asp | His | 0.469 | 0.344 |
| p.Asp273Ile | substitution | 273 | Asp | Ile | 0.381 | 0.042 |
| p.Asp273Phe | substitution | 273 | Asp | Phe | 0.000 | 0.028 |

| | | | | | | |
|---|---|---|---|---|---|---|
| p.Asp273Ser | substitution | 273 | Asp | Ser | 0.158 | 0.039 |
| p.Asp273Tyr | substitution | 273 | Asp | Tyr | 0.891 | 0.009 |
| p.Asp286= | synonymous | 286 | Asp | NA | 0.925 | 0.007 |
| p.Asp286Ala | substitution | 286 | Asp | Ala | 0.285 | 0.033 |
| p.Asp286Arg | substitution | 286 | Asp | Arg | 0.922 | 0.005 |
| p.Asp286Asn | substitution | 286 | Asp | Asn | 1.098 | 0.010 |
| p.Asp286Cys | substitution | 286 | Asp | Cys | 0.591 | 0.009 |
| p.Asp286Gln | substitution | 286 | Asp | Gln | 0.884 | 0.004 |
| p.Asp286Glu | substitution | 286 | Asp | Glu | 1.066 | 0.011 |
| p.Asp286Gly | substitution | 286 | Asp | Gly | 0.556 | 0.006 |
| p.Asp286His | substitution | 286 | Asp | His | 0.708 | 0.020 |
| p.Asp286Ile | substitution | 286 | Asp | Ile | 1.520 | 0.033 |
| p.Asp286Leu | substitution | 286 | Asp | Leu | 0.656 | 0.006 |
| p.Asp286Lys | substitution | 286 | Asp | Lys | 0.366 | 0.006 |
| p.Asp286Phe | substitution | 286 | Asp | Phe | 0.506 | 0.020 |
| p.Asp286Pro | substitution | 286 | Asp | Pro | 0.652 | 0.069 |
| p.Asp286Ser | substitution | 286 | Asp | Ser | 0.621 | 0.073 |
| p.Asp286Ter | nonsense | 286 | Asp | Ter | 0.000 | 0.097 |
| p.Asp286Thr | substitution | 286 | Asp | Thr | 0.613 | 0.031 |
| p.Asp286Trp | substitution | 286 | Asp | Trp | 0.729 | 0.005 |
| p.Asp286Tyr | substitution | 286 | Asp | Tyr | 0.227 | 0.070 |
| p.Asp286Val | substitution | 286 | Asp | Val | 0.898 | 0.014 |
| p.Asp289Ala | substitution | 289 | Asp | Ala | 1.043 | 0.004 |
| p.Asp289Arg | substitution | 289 | Asp | Arg | 0.758 | 0.028 |
| p.Asp289Asn | substitution | 289 | Asp | Asn | 0.448 | 0.036 |
| p.Asp289Cys | substitution | 289 | Asp | Cys | 0.181 | 0.018 |
| p.Asp289Gln | substitution | 289 | Asp | Gln | 1.423 | 0.015 |
| p.Asp289Glu | substitution | 289 | Asp | Glu | 0.752 | 0.004 |
| p.Asp289Gly | substitution | 289 | Asp | Gly | 0.619 | 0.078 |
| p.Asp289Ile | substitution | 289 | Asp | Ile | 1.398 | 0.006 |
| p.Asp289Leu | substitution | 289 | Asp | Leu | 0.685 | 0.228 |
| p.Asp289Lys | substitution | 289 | Asp | Lys | 0.151 | 0.011 |
| p.Asp289Phe | substitution | 289 | Asp | Phe | 1.079 | 0.010 |
| p.Asp289Pro | substitution | 289 | Asp | Pro | 1.523 | 0.042 |
| p.Asp289Ser | substitution | 289 | Asp | Ser | 0.786 | 0.095 |
| p.Asp289Ter | nonsense | 289 | Asp | Ter | 0.000 | 0.013 |
| p.Asp289Thr | substitution | 289 | Asp | Thr | 1.217 | 0.143 |
| p.Asp289Trp | substitution | 289 | Asp | Trp | 1.067 | 0.007 |
| p.Asp289Tyr | substitution | 289 | Asp | Tyr | 0.865 | 0.008 |
| p.Asp289Val | substitution | 289 | Asp | Val | 1.052 | 0.042 |
| p.Asp307Ala | substitution | 307 | Asp | Ala | 0.000 | 0.138 |
| p.Asp307Arg | substitution | 307 | Asp | Arg | 0.291 | 0.157 |
| p.Asp307Asn | substitution | 307 | Asp | Asn | 1.183 | 0.009 |

| | | | | | | |
|---|---|---|---|---|---|---|
| p.Asp307Cys | substitution | 307 | Asp | Cys | 0.812 | 0.003 |
| p.Asp307Gln | substitution | 307 | Asp | Gln | 0.146 | 0.258 |
| p.Asp307Glu | substitution | 307 | Asp | Glu | 0.523 | 0.052 |
| p.Asp307Gly | substitution | 307 | Asp | Gly | 1.223 | 0.027 |
| p.Asp307His | substitution | 307 | Asp | His | 1.323 | 0.003 |
| p.Asp307Ile | substitution | 307 | Asp | Ile | 0.000 | 0.166 |
| p.Asp307Leu | substitution | 307 | Asp | Leu | 0.000 | 0.183 |
| p.Asp307Lys | substitution | 307 | Asp | Lys | 1.907 | 0.007 |
| p.Asp307Phe | substitution | 307 | Asp | Phe | 1.475 | 0.002 |
| p.Asp307Ser | substitution | 307 | Asp | Ser | 0.639 | 0.017 |
| p.Asp307Ter | nonsense | 307 | Asp | Ter | 0.264 | 0.044 |
| p.Asp307Thr | substitution | 307 | Asp | Thr | 0.403 | 0.229 |
| p.Asp307Trp | substitution | 307 | Asp | Trp | 1.323 | 0.002 |
| p.Asp307Tyr | substitution | 307 | Asp | Tyr | 0.808 | 0.004 |
| p.Asp307Val | substitution | 307 | Asp | Val | 0.672 | 0.210 |
| p.Asp311Ala | substitution | 311 | Asp | Ala | 1.232 | 0.062 |
| p.Asp311Arg | substitution | 311 | Asp | Arg | 0.832 | 0.014 |
| p.Asp311Asn | substitution | 311 | Asp | Asn | 0.925 | 0.002 |
| p.Asp311Cys | substitution | 311 | Asp | Cys | 0.000 | 0.022 |
| p.Asp311Gln | substitution | 311 | Asp | Gln | 0.875 | 0.003 |
| p.Asp311Glu | substitution | 311 | Asp | Glu | 1.148 | 0.014 |
| p.Asp311Gly | substitution | 311 | Asp | Gly | 1.087 | 0.064 |
| p.Asp311His | substitution | 311 | Asp | His | 1.129 | 0.013 |
| p.Asp311Ile | substitution | 311 | Asp | Ile | 0.163 | 0.009 |
| p.Asp311Leu | substitution | 311 | Asp | Leu | 0.237 | 0.059 |
| p.Asp311Lys | substitution | 311 | Asp | Lys | 0.000 | 0.024 |
| p.Asp311Me | substitution | 311 | Asp | Met | 1.162 | 0.002 |
| p.Asp311Phe | substitution | 311 | Asp | Phe | 0.012 | 0.015 |
| p.Asp311Pro | substitution | 311 | Asp | Pro | 0.705 | 0.022 |
| p.Asp311Ser | substitution | 311 | Asp | Ser | 0.370 | 0.030 |
| p.Asp311Ter | nonsense | 311 | Asp | Ter | 0.000 | 0.014 |
| p.Asp311Thr | substitution | 311 | Asp | Thr | 0.372 | 0.031 |
| p.Asp311Trp | substitution | 311 | Asp | Trp | 0.680 | 0.008 |
| p.Asp311Tyr | substitution | 311 | Asp | Tyr | 0.523 | 0.005 |
| p.Asp311Val | substitution | 311 | Asp | Val | 0.901 | 0.027 |
| p.Asp312= | synonymous | 312 | Asp | NA | 1.109 | 0.003 |
| p.Asp312Ala | substitution | 312 | Asp | Ala | 0.925 | 0.092 |
| p.Asp312Arg | substitution | 312 | Asp | Arg | 0.957 | 0.074 |
| p.Asp312Asn | substitution | 312 | Asp | Asn | 1.251 | 0.009 |
| p.Asp312Cys | substitution | 312 | Asp | Cys | 1.128 | 0.034 |
| p.Asp312Glu | substitution | 312 | Asp | Glu | 0.804 | 0.004 |
| p.Asp312Gly | substitution | 312 | Asp | Gly | 0.758 | 1.048 |
| p.Asp312His | substitution | 312 | Asp | His | 0.994 | 0.020 |

| | | | | | | |
|---|---|---|---|---|---|---|
| p.Asp312Ile | substitution | 312 | Asp | Ile | 0.000 | 0.029 |
| p.Asp312Leu | substitution | 312 | Asp | Leu | 0.410 | 0.041 |
| p.Asp312Lys | substitution | 312 | Asp | Lys | 0.000 | 0.037 |
| p.Asp312Me | substitution | 312 | Asp | Met | 0.610 | 0.011 |
| p.Asp312Phe | substitution | 312 | Asp | Phe | 0.367 | 0.108 |
| p.Asp312Pro | substitution | 312 | Asp | Pro | 0.032 | 0.060 |
| p.Asp312Ser | substitution | 312 | Asp | Ser | 0.000 | 0.035 |
| p.Asp312Ter | nonsense | 312 | Asp | Ter | 0.000 | 0.020 |
| p.Asp312Thr | substitution | 312 | Asp | Thr | 0.000 | 0.010 |
| p.Asp312Trp | substitution | 312 | Asp | Trp | 0.363 | 0.008 |
| p.Asp312Tyr | substitution | 312 | Asp | Tyr | 0.919 | 0.009 |
| p.Asp312Val | substitution | 312 | Asp | Val | 0.427 | 0.063 |
| p.Asp352Ala | substitution | 352 | Asp | Ala | 0.976 | 0.012 |
| p.Asp352Arg | substitution | 352 | Asp | Arg | 0.833 | 0.012 |
| p.Asp352Asn | substitution | 352 | Asp | Asn | 0.440 | 0.014 |
| p.Asp352Cys | substitution | 352 | Asp | Cys | 0.000 | 0.029 |
| p.Asp352Gln | substitution | 352 | Asp | Gln | 0.868 | 0.012 |
| p.Asp352Glu | substitution | 352 | Asp | Glu | 0.193 | 0.026 |
| p.Asp352Gly | substitution | 352 | Asp | Gly | 0.000 | 0.574 |
| p.Asp352His | substitution | 352 | Asp | His | 1.225 | 0.004 |
| p.Asp352Ile | substitution | 352 | Asp | Ile | 0.000 | 0.036 |
| p.Asp352Leu | substitution | 352 | Asp | Leu | 0.935 | 0.015 |
| p.Asp352Lys | substitution | 352 | Asp | Lys | 0.782 | 0.004 |
| p.Asp352Me | substitution | 352 | Asp | Met | 0.723 | 0.005 |
| p.Asp352Phe | substitution | 352 | Asp | Phe | 1.159 | 0.002 |
| p.Asp352Pro | substitution | 352 | Asp | Pro | 1.119 | 0.021 |
| p.Asp352Ser | substitution | 352 | Asp | Ser | 0.773 | 0.003 |
| p.Asp352Ter | nonsense | 352 | Asp | Ter | 0.796 | 0.062 |
| p.Asp352Thr | substitution | 352 | Asp | Thr | 0.528 | 0.100 |
| p.Asp352Trp | substitution | 352 | Asp | Trp | 0.729 | 0.003 |
| p.Asp352Tyr | substitution | 352 | Asp | Tyr | 1.081 | 0.004 |
| p.Asp352Val | substitution | 352 | Asp | Val | 0.659 | 0.012 |
| p.Asp359Ala | substitution | 359 | Asp | Ala | 1.528 | 0.007 |
| p.Asp359Arg | substitution | 359 | Asp | Arg | 1.316 | 0.004 |
| p.Asp359Asn | substitution | 359 | Asp | Asn | 0.706 | 0.036 |
| p.Asp359Cys | substitution | 359 | Asp | Cys | 1.114 | 0.006 |
| p.Asp359Gln | substitution | 359 | Asp | Gln | 0.256 | 0.068 |
| p.Asp359Glu | substitution | 359 | Asp | Glu | 0.884 | 0.048 |
| p.Asp359Gly | substitution | 359 | Asp | Gly | 1.009 | 0.046 |
| p.Asp359His | substitution | 359 | Asp | His | 1.048 | 0.006 |
| p.Asp359Leu | substitution | 359 | Asp | Leu | 0.425 | 0.059 |
| p.Asp359Lys | substitution | 359 | Asp | Lys | 0.212 | 0.012 |
| p.Asp359Me | substitution | 359 | Asp | Met | 0.000 | 0.106 |

| | | | | | | | |
|---|---|---|---|---|---|---|---|
| p.Asp359Phe | substitution | 359 | Asp | Phe | | 0.564 | 0.029 |
| p.Asp359Ser | substitution | 359 | Asp | Ser | | 0.050 | 0.021 |
| p.Asp359Ter | nonsense | 359 | Asp | Ter | | 1.328 | 0.003 |
| p.Asp359Trp | substitution | 359 | Asp | Trp | | 1.041 | 0.008 |
| p.Asp359Tyr | substitution | 359 | Asp | Tyr | | 1.067 | 0.013 |
| p.Asp359Val | substitution | 359 | Asp | Val | | 0.080 | 0.124 |
| p.Asp36= | synonymous | 36 | Asp | NA | | 0.930 | 0.001 |
| p.Asp36Ala | substitution | 36 | Asp | Ala | | 0.755 | 0.026 |
| p.Asp36Arg | substitution | 36 | Asp | Arg | | 0.574 | 0.012 |
| p.Asp36Asn | substitution | 36 | Asp | Asn | | 0.836 | 0.017 |
| p.Asp36Cys | substitution | 36 | Asp | Cys | | 0.502 | 0.002 |
| p.Asp36Gln | substitution | 36 | Asp | Gln | | 1.333 | 0.003 |
| p.Asp36Glu | substitution | 36 | Asp | Glu | | 0.868 | 0.002 |
| p.Asp36Gly | substitution | 36 | Asp | Gly | | 1.036 | 0.010 |
| p.Asp36His | substitution | 36 | Asp | His | | 1.048 | 0.003 |
| p.Asp36Ile | substitution | 36 | Asp | Ile | | 0.527 | 0.003 |
| p.Asp36Leu | substitution | 36 | Asp | Leu | | 0.714 | 0.040 |
| p.Asp36Lys | substitution | 36 | Asp | Lys | | 1.108 | 0.004 |
| p.Asp36Met | substitution | 36 | Asp | Met | | 1.169 | 0.002 |
| p.Asp36Pro | substitution | 36 | Asp | Pro | | 0.000 | 0.063 |
| p.Asp36Ser | substitution | 36 | Asp | Ser | | 0.954 | 0.018 |
| p.Asp36Ter | nonsense | 36 | Asp | Ter | | 0.141 | 0.006 |
| p.Asp36Thr | substitution | 36 | Asp | Thr | | 0.903 | 0.002 |
| p.Asp36Trp | substitution | 36 | Asp | Trp | | 0.562 | 0.002 |
| p.Asp36Tyr | substitution | 36 | Asp | Tyr | | 0.835 | 0.012 |
| p.Asp36Val | substitution | 36 | Asp | Val | | 0.985 | 0.008 |
| p.Asp61= | synonymous | 61 | Asp | NA | | 0.874 | 0.042 |
| p.Asp61Ala | substitution | 61 | Asp | Ala | | 0.000 | 0.484 |
| p.Asp61Arg | substitution | 61 | Asp | Arg | | 0.000 | 0.880 |
| p.Asp61Asn | substitution | 61 | Asp | Asn | | 0.000 | 0.579 |
| p.Asp61Cys | substitution | 61 | Asp | Cys | | 0.000 | 0.772 |
| p.Asp61Gln | substitution | 61 | Asp | Gln | | 0.000 | 0.159 |
| p.Asp61Glu | substitution | 61 | Asp | Glu | | 1.091 | 0.241 |
| p.Asp61Gly | substitution | 61 | Asp | Gly | | 0.040 | 0.203 |
| p.Asp61His | substitution | 61 | Asp | His | | 0.256 | 0.497 |
| p.Asp61Leu | substitution | 61 | Asp | Leu | | 0.897 | 0.447 |
| p.Asp61Lys | substitution | 61 | Asp | Lys | | 0.358 | 0.044 |
| p.Asp61Phe | substitution | 61 | Asp | Phe | | 0.000 | 0.038 |
| p.Asp61Pro | substitution | 61 | Asp | Pro | | 0.061 | 0.243 |
| p.Asp61Ser | substitution | 61 | Asp | Ser | | 0.000 | 0.340 |
| p.Asp61Ter | nonsense | 61 | Asp | Ter | | 0.000 | 0.179 |
| p.Asp61Thr | substitution | 61 | Asp | Thr | | 0.000 | 0.135 |
| p.Asp61Trp | substitution | 61 | Asp | Trp | | 0.427 | 0.831 |

| | | | | | | |
|---|---|---|---|---|---|---|
| p.Asp61Tyr | substitution | 61 | Asp | Tyr | 0.335 | 0.127 |
| p.Asp61Val | substitution | 61 | Asp | Val | 0.000 | 0.245 |
| p.Asp65Ala | substitution | 65 | Asp | Ala | 0.528 | 0.004 |
| p.Asp65Arg | substitution | 65 | Asp | Arg | 0.987 | 0.002 |
| p.Asp65Asn | substitution | 65 | Asp | Asn | 0.316 | 0.012 |
| p.Asp65Cys | substitution | 65 | Asp | Cys | 0.391 | 0.008 |
| p.Asp65Gln | substitution | 65 | Asp | Gln | 0.585 | 0.022 |
| p.Asp65Glu | substitution | 65 | Asp | Glu | 1.210 | 0.011 |
| p.Asp65Gly | substitution | 65 | Asp | Gly | 0.710 | 0.017 |
| p.Asp65His | substitution | 65 | Asp | His | 0.838 | 0.011 |
| p.Asp65Ile | substitution | 65 | Asp | Ile | 0.645 | 0.013 |
| p.Asp65Leu | substitution | 65 | Asp | Leu | 0.171 | 0.015 |
| p.Asp65Lys | substitution | 65 | Asp | Lys | 1.134 | 0.004 |
| p.Asp65Met | substitution | 65 | Asp | Met | 0.121 | 0.011 |
| p.Asp65Phe | substitution | 65 | Asp | Phe | 0.784 | 0.004 |
| p.Asp65Pro | substitution | 65 | Asp | Pro | 0.761 | 0.053 |
| p.Asp65Ser | substitution | 65 | Asp | Ser | 0.824 | 0.030 |
| p.Asp65Ter | nonsense | 65 | Asp | Ter | 0.000 | 0.015 |
| p.Asp65Thr | substitution | 65 | Asp | Thr | 0.763 | 0.014 |
| p.Asp65Trp | substitution | 65 | Asp | Trp | 0.655 | 0.005 |
| p.Asp65Tyr | substitution | 65 | Asp | Tyr | 0.778 | 0.004 |
| p.Asp65Val | substitution | 65 | Asp | Val | 0.767 | 0.010 |
| p.Asp91= | synonymous | 91 | Asp | NA | 1.333 | 0.002 |
| p.Asp91Ala | substitution | 91 | Asp | Ala | 0.616 | 0.014 |
| p.Asp91Arg | substitution | 91 | Asp | Arg | 0.126 | 0.132 |
| p.Asp91Asn | substitution | 91 | Asp | Asn | 0.956 | 0.010 |
| p.Asp91Cys | substitution | 91 | Asp | Cys | 0.303 | 0.021 |
| p.Asp91Gln | substitution | 91 | Asp | Gln | 0.464 | 0.017 |
| p.Asp91Glu | substitution | 91 | Asp | Glu | 1.065 | 0.039 |
| p.Asp91Gly | substitution | 91 | Asp | Gly | 0.312 | 0.013 |
| p.Asp91His | substitution | 91 | Asp | His | 0.147 | 0.082 |
| p.Asp91Ile | substitution | 91 | Asp | Ile | 0.133 | 0.055 |
| p.Asp91Leu | substitution | 91 | Asp | Leu | 0.029 | 0.054 |
| p.Asp91Lys | substitution | 91 | Asp | Lys | 0.965 | 0.015 |
| p.Asp91Met | substitution | 91 | Asp | Met | 0.060 | 0.012 |
| p.Asp91Phe | substitution | 91 | Asp | Phe | 0.000 | 0.046 |
| p.Asp91Pro | substitution | 91 | Asp | Pro | 0.235 | 0.084 |
| p.Asp91Ser | substitution | 91 | Asp | Ser | 0.711 | 0.027 |
| p.Asp91Ter | nonsense | 91 | Asp | Ter | 0.000 | 0.033 |
| p.Asp91Thr | substitution | 91 | Asp | Thr | 0.045 | 0.017 |
| p.Asp91Trp | substitution | 91 | Asp | Trp | 0.000 | 0.044 |
| p.Asp91Tyr | substitution | 91 | Asp | Tyr | 0.248 | 0.120 |
| p.Asp91Val | substitution | 91 | Asp | Val | 0.199 | 0.007 |

| | | | | | | |
|---|---|---|---|---|---|---|
| p.Asp99= | synonymous | 99 | Asp | NA | 1.127 | 0.012 |
| p.Asp99Ala | substitution | 99 | Asp | Ala | 0.125 | 0.045 |
| p.Asp99Arg | substitution | 99 | Asp | Arg | 0.000 | 0.033 |
| p.Asp99Asn | substitution | 99 | Asp | Asn | 0.192 | 0.092 |
| p.Asp99Cys | substitution | 99 | Asp | Cys | 0.000 | 0.037 |
| p.Asp99Gln | substitution | 99 | Asp | Gln | 0.000 | 0.114 |
| p.Asp99Glu | substitution | 99 | Asp | Glu | 0.473 | 0.433 |
| p.Asp99Gly | substitution | 99 | Asp | Gly | 0.130 | 0.057 |
| p.Asp99His | substitution | 99 | Asp | His | 0.505 | 0.063 |
| p.Asp99Ile | substitution | 99 | Asp | Ile | 0.000 | 0.095 |
| p.Asp99Leu | substitution | 99 | Asp | Leu | 0.000 | 0.023 |
| p.Asp99Lys | substitution | 99 | Asp | Lys | 0.000 | 0.165 |
| p.Asp99Met | substitution | 99 | Asp | Met | 0.188 | 0.021 |
| p.Asp99Phe | substitution | 99 | Asp | Phe | 0.000 | 0.155 |
| p.Asp99Pro | substitution | 99 | Asp | Pro | 0.329 | 0.014 |
| p.Asp99Ser | substitution | 99 | Asp | Ser | 0.000 | 0.026 |
| p.Asp99Ter | nonsense | 99 | Asp | Ter | 0.352 | 0.021 |
| p.Asp99Thr | substitution | 99 | Asp | Thr | 0.000 | 0.050 |
| p.Asp99Trp | substitution | 99 | Asp | Trp | 0.000 | 0.013 |
| p.Asp99Tyr | substitution | 99 | Asp | Tyr | 0.000 | 0.064 |
| p.Asp99Val | substitution | 99 | Asp | Val | 0.000 | 0.056 |
| p.Cys114= | synonymous | 114 | Cys | NA | 0.942 | 0.004 |
| p.Cys114Ala | substitution | 114 | Cys | Ala | 0.362 | 0.084 |
| p.Cys114Arg | substitution | 114 | Cys | Arg | 0.726 | 0.032 |
| p.Cys114Asn | substitution | 114 | Cys | Asn | 0.780 | 0.025 |
| p.Cys114Asp | substitution | 114 | Cys | Asp | 0.813 | 0.022 |
| p.Cys114Gln | substitution | 114 | Cys | Gln | 0.956 | 0.007 |
| p.Cys114Glu | substitution | 114 | Cys | Glu | 0.000 | 0.017 |
| p.Cys114Gly | substitution | 114 | Cys | Gly | 0.561 | 0.021 |
| p.Cys114His | substitution | 114 | Cys | His | 0.270 | 0.078 |
| p.Cys114Ile | substitution | 114 | Cys | Ile | 1.030 | 0.015 |
| p.Cys114Leu | substitution | 114 | Cys | Leu | 0.796 | 0.005 |
| p.Cys114Lys | substitution | 114 | Cys | Lys | 0.597 | 0.015 |
| p.Cys114Phe | substitution | 114 | Cys | Phe | 0.633 | 0.010 |
| p.Cys114Pro | substitution | 114 | Cys | Pro | 0.836 | 0.117 |
| p.Cys114Ser | substitution | 114 | Cys | Ser | 0.971 | 0.029 |
| p.Cys114Ter | nonsense | 114 | Cys | Ter | 0.010 | 0.064 |
| p.Cys114Thr | substitution | 114 | Cys | Thr | 1.112 | 0.012 |
| p.Cys114Trp | substitution | 114 | Cys | Trp | 0.682 | 0.010 |
| p.Cys114Tyr | substitution | 114 | Cys | Tyr | 0.405 | 0.138 |
| p.Cys114Val | substitution | 114 | Cys | Val | 0.386 | 0.043 |
| p.Cys211= | synonymous | 211 | Cys | NA | 1.073 | 0.004 |
| p.Cys211Ala | substitution | 211 | Cys | Ala | 0.845 | 0.045 |

| | | | | | | | |
|---|---|---|---|---|---|---|---|
| p.Cys211Arg | substitution | 211 | Cys | Arg | | 0.000 | 0.053 |
| p.Cys211Glu | substitution | 211 | Cys | Glu | | 0.000 | 0.014 |
| p.Cys211Gly | substitution | 211 | Cys | Gly | | 0.580 | 0.054 |
| p.Cys211Leu | substitution | 211 | Cys | Leu | | 0.838 | 0.141 |
| p.Cys211Lys | substitution | 211 | Cys | Lys | | 0.000 | 0.032 |
| p.Cys211Met | substitution | 211 | Cys | Met | | 1.436 | 0.003 |
| p.Cys211Phe | substitution | 211 | Cys | Phe | | 1.298 | 0.017 |
| p.Cys211Pro | substitution | 211 | Cys | Pro | | 0.000 | 0.066 |
| p.Cys211Ser | substitution | 211 | Cys | Ser | | 0.908 | 0.086 |
| p.Cys211Ter | nonsense | 211 | Cys | Ter | | 0.057 | 0.169 |
| p.Cys211Thr | substitution | 211 | Cys | Thr | | 0.288 | 0.119 |
| p.Cys211Trp | substitution | 211 | Cys | Trp | | 0.535 | 0.087 |
| p.Cys211Tyr | substitution | 211 | Cys | Tyr | | 0.748 | 0.119 |
| p.Cys211Val | substitution | 211 | Cys | Val | | 0.932 | 0.638 |
| p.Cys247= | synonymous | 247 | Cys | NA | | 1.344 | 0.004 |
| p.Cys247Ala | substitution | 247 | Cys | Ala | | 1.288 | 0.064 |
| p.Cys247Arg | substitution | 247 | Cys | Arg | | 0.000 | 0.035 |
| p.Cys247Asn | substitution | 247 | Cys | Asn | | 0.475 | 0.011 |
| p.Cys247Gln | substitution | 247 | Cys | Gln | | 0.000 | 0.058 |
| p.Cys247Glu | substitution | 247 | Cys | Glu | | 0.225 | 0.009 |
| p.Cys247Gly | substitution | 247 | Cys | Gly | | 0.461 | 0.053 |
| p.Cys247His | substitution | 247 | Cys | His | | 0.000 | 0.189 |
| p.Cys247Ile | substitution | 247 | Cys | Ile | | 0.000 | 0.120 |
| p.Cys247Leu | substitution | 247 | Cys | Leu | | 0.263 | 0.022 |
| p.Cys247Met | substitution | 247 | Cys | Met | | 0.144 | 0.221 |
| p.Cys247Phe | substitution | 247 | Cys | Phe | | 0.387 | 0.134 |
| p.Cys247Ser | substitution | 247 | Cys | Ser | | 0.785 | 0.057 |
| p.Cys247Ter | nonsense | 247 | Cys | Ter | | 0.000 | 0.081 |
| p.Cys247Thr | substitution | 247 | Cys | Thr | | 0.409 | 0.020 |
| p.Cys247Trp | substitution | 247 | Cys | Trp | | 0.128 | 0.009 |
| p.Cys247Tyr | substitution | 247 | Cys | Tyr | | 0.822 | 0.086 |
| p.Cys247Val | substitution | 247 | Cys | Val | | 0.716 | 0.008 |
| p.Cys261= | synonymous | 261 | Cys | NA | | 1.305 | 0.005 |
| p.Cys261Ala | substitution | 261 | Cys | Ala | | 0.000 | 0.054 |
| p.Cys261Arg | substitution | 261 | Cys | Arg | | 0.036 | 0.029 |
| p.Cys261Asn | substitution | 261 | Cys | Asn | | 0.000 | 0.130 |
| p.Cys261Asp | substitution | 261 | Cys | Asp | | 0.274 | 0.031 |
| p.Cys261Gln | substitution | 261 | Cys | Gln | | 0.057 | 0.128 |
| p.Cys261Glu | substitution | 261 | Cys | Glu | | 0.000 | 0.015 |
| p.Cys261Gly | substitution | 261 | Cys | Gly | | 0.218 | 0.315 |
| p.Cys261His | substitution | 261 | Cys | His | | 0.000 | 0.027 |
| p.Cys261Ile | substitution | 261 | Cys | Ile | | 0.000 | 0.062 |
| p.Cys261Leu | substitution | 261 | Cys | Leu | | 0.000 | 0.105 |

| | | | | | | |
|---|---|---|---|---|---|---|
| p.Cys261Lys | substitution | 261 | Cys | Lys | 0.000 | 0.030 |
| p.Cys261Met | substitution | 261 | Cys | Met | 0.000 | 0.114 |
| p.Cys261Phe | substitution | 261 | Cys | Phe | 0.265 | 0.035 |
| p.Cys261Pro | substitution | 261 | Cys | Pro | 0.000 | 0.098 |
| p.Cys261Ser | substitution | 261 | Cys | Ser | 0.000 | 0.090 |
| p.Cys261Ter | nonsense | 261 | Cys | Ter | 0.061 | 0.100 |
| p.Cys261Trp | substitution | 261 | Cys | Trp | 0.600 | 0.402 |
| p.Cys261Tyr | substitution | 261 | Cys | Tyr | 0.576 | 0.149 |
| p.Cys261Val | substitution | 261 | Cys | Val | 0.000 | 0.011 |
| p.Gln153= | synonymous | 153 | Gln | NA | 1.072 | 0.009 |
| p.Gln153Ala | substitution | 153 | Gln | Ala | 0.276 | 0.269 |
| p.Gln153Arg | substitution | 153 | Gln | Arg | 0.061 | 0.023 |
| p.Gln153Asn | substitution | 153 | Gln | Asn | 0.000 | 0.042 |
| p.Gln153Asp | substitution | 153 | Gln | Asp | 0.000 | 0.035 |
| p.Gln153Cys | substitution | 153 | Gln | Cys | 0.592 | 0.013 |
| p.Gln153Glu | substitution | 153 | Gln | Glu | 0.288 | 0.069 |
| p.Gln153Gly | substitution | 153 | Gln | Gly | 0.605 | 0.006 |
| p.Gln153His | substitution | 153 | Gln | His | 0.447 | 0.048 |
| p.Gln153Ile | substitution | 153 | Gln | Ile | 0.075 | 0.019 |
| p.Gln153Leu | substitution | 153 | Gln | Leu | 0.548 | 0.022 |
| p.Gln153Lys | substitution | 153 | Gln | Lys | 0.123 | 0.025 |
| p.Gln153Met | substitution | 153 | Gln | Met | 0.552 | 0.008 |
| p.Gln153Phe | substitution | 153 | Gln | Phe | 0.000 | 0.313 |
| p.Gln153Pro | substitution | 153 | Gln | Pro | 0.884 | 0.015 |
| p.Gln153Ser | substitution | 153 | Gln | Ser | 0.671 | 0.007 |
| p.Gln153Ter | nonsense | 153 | Gln | Ter | 0.088 | 0.315 |
| p.Gln153Thr | substitution | 153 | Gln | Thr | 0.271 | 0.136 |
| p.Gln153Tyr | substitution | 153 | Gln | Tyr | 0.981 | 0.012 |
| p.Gln153Val | substitution | 153 | Gln | Val | 0.388 | 0.223 |
| p.Gln155= | synonymous | 155 | Gln | NA | 1.403 | 0.029 |
| p.Gln155Ala | substitution | 155 | Gln | Ala | 1.112 | 0.004 |
| p.Gln155Arg | substitution | 155 | Gln | Arg | 0.880 | 0.011 |
| p.Gln155Asp | substitution | 155 | Gln | Asp | 0.496 | 0.014 |
| p.Gln155Cys | substitution | 155 | Gln | Cys | 1.029 | 0.140 |
| p.Gln155Glu | substitution | 155 | Gln | Glu | 0.813 | 0.007 |
| p.Gln155Gly | substitution | 155 | Gln | Gly | 0.860 | 0.005 |
| p.Gln155His | substitution | 155 | Gln | His | 0.959 | 0.009 |
| p.Gln155Ile | substitution | 155 | Gln | Ile | 1.263 | 0.005 |
| p.Gln155Leu | substitution | 155 | Gln | Leu | 0.808 | 0.010 |
| p.Gln155Lys | substitution | 155 | Gln | Lys | 1.030 | 0.007 |
| p.Gln155Met | substitution | 155 | Gln | Met | 0.864 | 0.005 |
| p.Gln155Phe | substitution | 155 | Gln | Phe | 0.879 | 0.009 |
| p.Gln155Pro | substitution | 155 | Gln | Pro | 0.797 | 0.046 |

| | | | | | | |
|---|---|---|---|---|---|---|
| p.Gln155Ser | substitution | 155 | Gln | Ser | 1.054 | 0.011 |
| p.Gln155Ter | nonsense | 155 | Gln | Ter | 0.000 | 0.028 |
| p.Gln155Thr | substitution | 155 | Gln | Thr | 0.859 | 0.008 |
| p.Gln155Trp | substitution | 155 | Gln | Trp | 0.859 | 0.005 |
| p.Gln155Tyr | substitution | 155 | Gln | Tyr | 0.704 | 0.007 |
| p.Gln155Val | substitution | 155 | Gln | Val | 0.803 | 0.006 |
| p.Gln180= | synonymous | 180 | Gln | NA | 0.823 | 0.015 |
| p.Gln180Ala | substitution | 180 | Gln | Ala | 1.151 | 0.007 |
| p.Gln180Arg | substitution | 180 | Gln | Arg | 1.048 | 0.009 |
| p.Gln180Asn | substitution | 180 | Gln | Asn | 1.507 | 0.010 |
| p.Gln180Asp | substitution | 180 | Gln | Asp | 0.623 | 0.019 |
| p.Gln180Cys | substitution | 180 | Gln | Cys | 1.308 | 0.009 |
| p.Gln180Gly | substitution | 180 | Gln | Gly | 0.805 | 0.006 |
| p.Gln180His | substitution | 180 | Gln | His | 0.917 | 0.033 |
| p.Gln180Ile | substitution | 180 | Gln | Ile | 0.095 | 0.012 |
| p.Gln180Leu | substitution | 180 | Gln | Leu | 0.914 | 0.019 |
| p.Gln180Lys | substitution | 180 | Gln | Lys | 1.070 | 0.009 |
| p.Gln180Met | substitution | 180 | Gln | Met | 0.806 | 0.011 |
| p.Gln180Phe | substitution | 180 | Gln | Phe | 0.068 | 0.018 |
| p.Gln180Pro | substitution | 180 | Gln | Pro | 0.750 | 0.011 |
| p.Gln180Ser | substitution | 180 | Gln | Ser | 0.885 | 0.079 |
| p.Gln180Ter | nonsense | 180 | Gln | Ter | 0.268 | 0.019 |
| p.Gln180Thr | substitution | 180 | Gln | Thr | 0.835 | 0.005 |
| p.Gln180Trp | substitution | 180 | Gln | Trp | 0.435 | 0.019 |
| p.Gln180Tyr | substitution | 180 | Gln | Tyr | 0.897 | 0.017 |
| p.Gln180Val | substitution | 180 | Gln | Val | 0.733 | 0.010 |
| p.Gln181= | synonymous | 181 | Gln | NA | 0.660 | 0.057 |
| p.Gln181Ala | substitution | 181 | Gln | Ala | 0.589 | 0.031 |
| p.Gln181Arg | substitution | 181 | Gln | Arg | 1.029 | 0.018 |
| p.Gln181Asn | substitution | 181 | Gln | Asn | 0.430 | 0.008 |
| p.Gln181Asp | substitution | 181 | Gln | Asp | 0.614 | 0.008 |
| p.Gln181Cys | substitution | 181 | Gln | Cys | 0.000 | 0.047 |
| p.Gln181Glu | substitution | 181 | Gln | Glu | 1.183 | 0.005 |
| p.Gln181Gly | substitution | 181 | Gln | Gly | 0.880 | 0.087 |
| p.Gln181His | substitution | 181 | Gln | His | 0.721 | 0.011 |
| p.Gln181Ile | substitution | 181 | Gln | Ile | 0.595 | 0.006 |
| p.Gln181Leu | substitution | 181 | Gln | Leu | 0.735 | 0.014 |
| p.Gln181Lys | substitution | 181 | Gln | Lys | 0.913 | 0.009 |
| p.Gln181Met | substitution | 181 | Gln | Met | 0.803 | 0.006 |
| p.Gln181Phe | substitution | 181 | Gln | Phe | 0.802 | 0.006 |
| p.Gln181Pro | substitution | 181 | Gln | Pro | 0.374 | 0.038 |
| p.Gln181Ser | substitution | 181 | Gln | Ser | 0.801 | 0.015 |
| p.Gln181Ter | nonsense | 181 | Gln | Ter | 0.055 | 0.027 |

| | | | | | | |
|---|---|---|---|---|---|---|
| p.Gln181Thr | substitution | 181 | Gln | Thr | 1.016 | 0.005 |
| p.Gln181Trp | substitution | 181 | Gln | Trp | 0.807 | 0.007 |
| p.Gln181Tyr | substitution | 181 | Gln | Tyr | 1.164 | 0.005 |
| p.Gln181Val | substitution | 181 | Gln | Val | 0.442 | 0.055 |
| p.Gln194= | synonymous | 194 | Gln | NA | 1.117 | 0.011 |
| p.Gln194Ala | substitution | 194 | Gln | Ala | 1.000 | 0.019 |
| p.Gln194Arg | substitution | 194 | Gln | Arg | 1.065 | 0.098 |
| p.Gln194Asn | substitution | 194 | Gln | Asn | 0.458 | 0.033 |
| p.Gln194Asp | substitution | 194 | Gln | Asp | 0.801 | 0.005 |
| p.Gln194Cys | substitution | 194 | Gln | Cys | 0.083 | 0.018 |
| p.Gln194Glu | substitution | 194 | Gln | Glu | 0.389 | 0.007 |
| p.Gln194Gly | substitution | 194 | Gln | Gly | 0.375 | 0.012 |
| p.Gln194His | substitution | 194 | Gln | His | 0.350 | 0.033 |
| p.Gln194Ile | substitution | 194 | Gln | Ile | 0.000 | 0.251 |
| p.Gln194Leu | substitution | 194 | Gln | Leu | 0.769 | 0.038 |
| p.Gln194Lys | substitution | 194 | Gln | Lys | 1.001 | 0.003 |
| p.Gln194Met | substitution | 194 | Gln | Met | 1.360 | 0.006 |
| p.Gln194Phe | substitution | 194 | Gln | Phe | 0.000 | 0.036 |
| p.Gln194Pro | substitution | 194 | Gln | Pro | 0.780 | 0.199 |
| p.Gln194Ser | substitution | 194 | Gln | Ser | 0.120 | 0.019 |
| p.Gln194Ter | nonsense | 194 | Gln | Ter | 0.193 | 0.833 |
| p.Gln194Thr | substitution | 194 | Gln | Thr | 0.111 | 0.042 |
| p.Gln194Tyr | substitution | 194 | Gln | Tyr | 0.264 | 0.049 |
| p.Gln194Val | substitution | 194 | Gln | Val | 1.123 | 0.095 |
| p.Gln204= | synonymous | 204 | Gln | NA | 0.830 | 0.036 |
| p.Gln204Ala | substitution | 204 | Gln | Ala | 0.147 | 0.023 |
| p.Gln204Arg | substitution | 204 | Gln | Arg | 0.722 | 0.061 |
| p.Gln204Asn | substitution | 204 | Gln | Asn | 0.000 | 0.065 |
| p.Gln204Asp | substitution | 204 | Gln | Asp | 0.044 | 0.013 |
| p.Gln204Cys | substitution | 204 | Gln | Cys | 0.000 | 0.038 |
| p.Gln204Glu | substitution | 204 | Gln | Glu | 0.745 | 0.160 |
| p.Gln204Gly | substitution | 204 | Gln | Gly | 0.512 | 0.193 |
| p.Gln204His | substitution | 204 | Gln | His | 0.987 | 0.003 |
| p.Gln204Ile | substitution | 204 | Gln | Ile | 0.000 | 0.035 |
| p.Gln204Leu | substitution | 204 | Gln | Leu | 1.160 | 0.055 |
| p.Gln204Lys | substitution | 204 | Gln | Lys | 0.586 | 0.006 |
| p.Gln204Met | substitution | 204 | Gln | Met | 0.000 | 0.115 |
| p.Gln204Pro | substitution | 204 | Gln | Pro | 0.641 | 0.010 |
| p.Gln204Ser | substitution | 204 | Gln | Ser | 1.590 | 0.177 |
| p.Gln204Ter | nonsense | 204 | Gln | Ter | 0.306 | 0.010 |
| p.Gln204Thr | substitution | 204 | Gln | Thr | 0.000 | 0.051 |
| p.Gln204Trp | substitution | 204 | Gln | Trp | 0.000 | 0.008 |
| p.Gln204Tyr | substitution | 204 | Gln | Tyr | 0.629 | 0.005 |

| | | | | | | | |
|---|---|---|---|---|---|---|---|
| p.Gln204Val | substitution | 204 | Gln | Val | | 0.995 | 0.005 |
| p.Gln217= | synonymous | 217 | Gln | NA | | 0.386 | 0.743 |
| p.Gln217Ala | substitution | 217 | Gln | Ala | | 0.427 | 0.099 |
| p.Gln217Arg | substitution | 217 | Gln | Arg | | 0.000 | 0.081 |
| p.Gln217Asn | substitution | 217 | Gln | Asn | | 0.000 | 0.039 |
| p.Gln217Asp | substitution | 217 | Gln | Asp | | 0.000 | 0.264 |
| p.Gln217Cys | substitution | 217 | Gln | Cys | | 0.034 | 0.018 |
| p.Gln217Glu | substitution | 217 | Gln | Glu | | 0.119 | 0.022 |
| p.Gln217Gly | substitution | 217 | Gln | Gly | | 0.000 | 0.023 |
| p.Gln217His | substitution | 217 | Gln | His | | 0.394 | 0.007 |
| p.Gln217Ile | substitution | 217 | Gln | Ile | | 0.035 | 0.008 |
| p.Gln217Leu | substitution | 217 | Gln | Leu | | 0.000 | 0.066 |
| p.Gln217Lys | substitution | 217 | Gln | Lys | | 0.215 | 0.016 |
| p.Gln217Met | substitution | 217 | Gln | Met | | 0.000 | 0.031 |
| p.Gln217Phe | substitution | 217 | Gln | Phe | | 0.000 | 0.050 |
| p.Gln217Pro | substitution | 217 | Gln | Pro | | 0.585 | 0.153 |
| p.Gln217Ser | substitution | 217 | Gln | Ser | | 0.712 | 0.193 |
| p.Gln217Ter | nonsense | 217 | Gln | Ter | | 0.124 | 0.218 |
| p.Gln217Thr | substitution | 217 | Gln | Thr | | 0.000 | 0.011 |
| p.Gln217Trp | substitution | 217 | Gln | Trp | | 0.000 | 0.008 |
| p.Gln217Tyr | substitution | 217 | Gln | Tyr | | 0.441 | 0.097 |
| p.Gln217Val | substitution | 217 | Gln | Val | | 0.000 | 0.017 |
| p.Gln229Ala | substitution | 229 | Gln | Ala | | 0.501 | 0.031 |
| p.Gln229Arg | substitution | 229 | Gln | Arg | | 0.949 | 0.017 |
| p.Gln229Asn | substitution | 229 | Gln | Asn | | 1.076 | 0.002 |
| p.Gln229Asp | substitution | 229 | Gln | Asp | | 1.072 | 0.002 |
| p.Gln229Cys | substitution | 229 | Gln | Cys | | 1.199 | 0.002 |
| p.Gln229Glu | substitution | 229 | Gln | Glu | | 1.059 | 0.002 |
| p.Gln229Gly | substitution | 229 | Gln | Gly | | 1.101 | 0.011 |
| p.Gln229His | substitution | 229 | Gln | His | | 0.912 | 0.002 |
| p.Gln229Ile | substitution | 229 | Gln | Ile | | 0.252 | 0.028 |
| p.Gln229Leu | substitution | 229 | Gln | Leu | | 0.707 | 0.005 |
| p.Gln229Lys | substitution | 229 | Gln | Lys | | 0.646 | 0.003 |
| p.Gln229Met | substitution | 229 | Gln | Met | | 0.924 | 0.002 |
| p.Gln229Phe | substitution | 229 | Gln | Phe | | 1.042 | 0.001 |
| p.Gln229Pro | substitution | 229 | Gln | Pro | | 0.502 | 0.074 |
| p.Gln229Ser | substitution | 229 | Gln | Ser | | 0.082 | 0.033 |
| p.Gln229Ter | nonsense | 229 | Gln | Ter | | 0.000 | 0.289 |
| p.Gln229Thr | substitution | 229 | Gln | Thr | | 0.581 | 0.046 |
| p.Gln229Tyr | substitution | 229 | Gln | Tyr | | 1.086 | 0.002 |
| p.Gln229Val | substitution | 229 | Gln | Val | | 0.383 | 0.019 |
| p.Gln275= | synonymous | 275 | Gln | NA | | 0.756 | 0.312 |
| p.Gln275Ala | substitution | 275 | Gln | Ala | | 0.000 | 0.024 |

| | | | | | | |
|---|---|---|---|---|---|---|
| p.Gln275Arg | substitution | 275 | Gln | Arg | 1.085 | 0.041 |
| p.Gln275Asn | substitution | 275 | Gln | Asn | 0.083 | 0.007 |
| p.Gln275Asp | substitution | 275 | Gln | Asp | 0.008 | 0.049 |
| p.Gln275Cys | substitution | 275 | Gln | Cys | 0.000 | 0.033 |
| p.Gln275Glu | substitution | 275 | Gln | Glu | 0.304 | 0.074 |
| p.Gln275Gly | substitution | 275 | Gln | Gly | 1.539 | 0.007 |
| p.Gln275His | substitution | 275 | Gln | His | 1.149 | 0.077 |
| p.Gln275Ile | substitution | 275 | Gln | Ile | 0.299 | 0.094 |
| p.Gln275Leu | substitution | 275 | Gln | Leu | 0.000 | 0.052 |
| p.Gln275Lys | substitution | 275 | Gln | Lys | 0.990 | 0.030 |
| p.Gln275Met | substitution | 275 | Gln | Met | 1.434 | 0.004 |
| p.Gln275Phe | substitution | 275 | Gln | Phe | 0.402 | 0.013 |
| p.Gln275Pro | substitution | 275 | Gln | Pro | 0.601 | 0.230 |
| p.Gln275Ser | substitution | 275 | Gln | Ser | 0.000 | 0.009 |
| p.Gln275Ter | nonsense | 275 | Gln | Ter | 0.000 | 0.073 |
| p.Gln275Thr | substitution | 275 | Gln | Thr | 0.000 | 0.017 |
| p.Gln275Trp | substitution | 275 | Gln | Trp | 0.000 | 0.006 |
| p.Gln275Tyr | substitution | 275 | Gln | Tyr | 0.000 | 0.073 |
| p.Gln275Val | substitution | 275 | Gln | Val | 1.000 | 0.080 |
| p.Gln292= | synonymous | 292 | Gln | NA | 1.255 | 0.004 |
| p.Gln292Ala | substitution | 292 | Gln | Ala | 0.621 | 0.043 |
| p.Gln292Arg | substitution | 292 | Gln | Arg | 0.761 | 0.024 |
| p.Gln292Asn | substitution | 292 | Gln | Asn | 0.851 | 0.004 |
| p.Gln292Asp | substitution | 292 | Gln | Asp | 0.791 | 0.010 |
| p.Gln292Cys | substitution | 292 | Gln | Cys | 0.951 | 0.004 |
| p.Gln292Glu | substitution | 292 | Gln | Glu | 1.224 | 0.012 |
| p.Gln292Gly | substitution | 292 | Gln | Gly | 0.653 | 0.015 |
| p.Gln292His | substitution | 292 | Gln | His | 1.149 | 0.013 |
| p.Gln292Ile | substitution | 292 | Gln | Ile | 1.447 | 0.003 |
| p.Gln292Leu | substitution | 292 | Gln | Leu | 1.001 | 0.009 |
| p.Gln292Lys | substitution | 292 | Gln | Lys | 0.300 | 0.003 |
| p.Gln292Met | substitution | 292 | Gln | Met | 1.033 | 0.005 |
| p.Gln292Phe | substitution | 292 | Gln | Phe | 0.705 | 0.004 |
| p.Gln292Pro | substitution | 292 | Gln | Pro | 0.519 | 0.005 |
| p.Gln292Ser | substitution | 292 | Gln | Ser | 1.134 | 0.027 |
| p.Gln292Ter | nonsense | 292 | Gln | Ter | 0.384 | 0.062 |
| p.Gln292Thr | substitution | 292 | Gln | Thr | 0.815 | 0.031 |
| p.Gln292Trp | substitution | 292 | Gln | Trp | 0.476 | 0.005 |
| p.Gln292Tyr | substitution | 292 | Gln | Tyr | 0.392 | 0.004 |
| p.Gln292Val | substitution | 292 | Gln | Val | 0.552 | 0.017 |
| p.Gln296= | synonymous | 296 | Gln | NA | 1.067 | 0.042 |
| p.Gln296Ala | substitution | 296 | Gln | Ala | 0.000 | 0.010 |
| p.Gln296Arg | substitution | 296 | Gln | Arg | 1.061 | 0.003 |

| | | | | | | | |
|---|---|---|---|---|---|---|---|
| p.Gln296Asn | substitution | 296 | Gln | Asn | | 0.941 | 0.005 |
| p.Gln296Asp | substitution | 296 | Gln | Asp | | 0.122 | 0.009 |
| p.Gln296Cys | substitution | 296 | Gln | Cys | | 0.000 | 0.011 |
| p.Gln296Glu | substitution | 296 | Gln | Glu | | 0.865 | 0.004 |
| p.Gln296Gly | substitution | 296 | Gln | Gly | | 0.973 | 0.004 |
| p.Gln296His | substitution | 296 | Gln | His | | 1.110 | 0.041 |
| p.Gln296Ile | substitution | 296 | Gln | Ile | | 0.506 | 0.007 |
| p.Gln296Leu | substitution | 296 | Gln | Leu | | 1.020 | 0.050 |
| p.Gln296Lys | substitution | 296 | Gln | Lys | | 0.957 | 0.010 |
| p.Gln296Met | substitution | 296 | Gln | Met | | 0.000 | 0.008 |
| p.Gln296Pro | substitution | 296 | Gln | Pro | | 0.576 | 0.040 |
| p.Gln296Ser | substitution | 296 | Gln | Ser | | 0.632 | 0.030 |
| p.Gln296Ter | nonsense | 296 | Gln | Ter | | 0.152 | 0.363 |
| p.Gln296Thr | substitution | 296 | Gln | Thr | | 0.367 | 0.061 |
| p.Gln296Trp | substitution | 296 | Gln | Trp | | 0.547 | 0.010 |
| p.Gln296Tyr | substitution | 296 | Gln | Tyr | | 0.814 | 0.012 |
| p.Gln296Val | substitution | 296 | Gln | Val | | 0.971 | 0.013 |
| p.Gln29= | synonymous | 29 | Gln | NA | | 1.197 | 0.001 |
| p.Gln29Ala | substitution | 29 | Gln | Ala | | 0.971 | 0.012 |
| p.Gln29Arg | substitution | 29 | Gln | Arg | | 1.066 | 0.000 |
| p.Gln29Asn | substitution | 29 | Gln | Asn | | 1.353 | 0.001 |
| p.Gln29Asp | substitution | 29 | Gln | Asp | | 0.945 | 0.001 |
| p.Gln29Cys | substitution | 29 | Gln | Cys | | 1.202 | 0.000 |
| p.Gln29Glu | substitution | 29 | Gln | Glu | | 0.511 | 0.002 |
| p.Gln29Gly | substitution | 29 | Gln | Gly | | 1.161 | 0.001 |
| p.Gln29His | substitution | 29 | Gln | His | | 0.818 | 0.001 |
| p.Gln29Ile | substitution | 29 | Gln | Ile | | 0.503 | 0.001 |
| p.Gln29Leu | substitution | 29 | Gln | Leu | | 0.874 | 0.002 |
| p.Gln29Lys | substitution | 29 | Gln | Lys | | 0.951 | 0.001 |
| p.Gln29Met | substitution | 29 | Gln | Met | | 1.026 | 0.000 |
| p.Gln29Phe | substitution | 29 | Gln | Phe | | 0.499 | 0.003 |
| p.Gln29Pro | substitution | 29 | Gln | Pro | | 0.826 | 0.001 |
| p.Gln29Ser | substitution | 29 | Gln | Ser | | 0.980 | 0.005 |
| p.Gln29Ter | nonsense | 29 | Gln | Ter | | 0.000 | 0.008 |
| p.Gln29Thr | substitution | 29 | Gln | Thr | | 0.770 | 0.005 |
| p.Gln29Trp | substitution | 29 | Gln | Trp | | 0.850 | 0.000 |
| p.Gln29Tyr | substitution | 29 | Gln | Tyr | | 0.668 | 0.001 |
| p.Gln29Val | substitution | 29 | Gln | Val | | 0.522 | 0.001 |
| p.Gln304= | synonymous | 304 | Gln | NA | | 1.188 | 0.004 |
| p.Gln304Ala | substitution | 304 | Gln | Ala | | 1.270 | 0.016 |
| p.Gln304Arg | substitution | 304 | Gln | Arg | | 1.020 | 0.010 |
| p.Gln304Asn | substitution | 304 | Gln | Asn | | 1.315 | 0.059 |
| p.Gln304Asp | substitution | 304 | Gln | Asp | | 0.000 | 0.025 |

| | | | | | | |
|---|---|---|---|---|---|---|
| p.Gln304Cys | substitution | 304 | Gln | Cys | 1.038 | 0.002 |
| p.Gln304Glu | substitution | 304 | Gln | Glu | 1.071 | 0.001 |
| p.Gln304Gly | substitution | 304 | Gln | Gly | 0.452 | 0.044 |
| p.Gln304His | substitution | 304 | Gln | His | 0.686 | 0.027 |
| p.Gln304Ile | substitution | 304 | Gln | Ile | 0.053 | 0.159 |
| p.Gln304Leu | substitution | 304 | Gln | Leu | 0.269 | 0.081 |
| p.Gln304Lys | substitution | 304 | Gln | Lys | 1.134 | 0.002 |
| p.Gln304Met | substitution | 304 | Gln | Met | 0.471 | 0.006 |
| p.Gln304Pro | substitution | 304 | Gln | Pro | 0.377 | 0.084 |
| p.Gln304Ser | substitution | 304 | Gln | Ser | 1.094 | 0.073 |
| p.Gln304Ter | nonsense | 304 | Gln | Ter | 0.014 | 0.123 |
| p.Gln304Thr | substitution | 304 | Gln | Thr | 0.520 | 0.008 |
| p.Gln304Trp | substitution | 304 | Gln | Trp | 0.458 | 0.004 |
| p.Gln304Val | substitution | 304 | Gln | Val | 0.000 | 0.057 |
| p.Gln314= | synonymous | 314 | Gln | NA | 1.253 | 0.388 |
| p.Gln314Ala | substitution | 314 | Gln | Ala | 0.171 | 0.013 |
| p.Gln314Arg | substitution | 314 | Gln | Arg | 0.000 | 0.020 |
| p.Gln314Asn | substitution | 314 | Gln | Asn | 1.140 | 0.006 |
| p.Gln314Asp | substitution | 314 | Gln | Asp | 0.000 | 0.017 |
| p.Gln314Cys | substitution | 314 | Gln | Cys | 1.012 | 0.002 |
| p.Gln314Glu | substitution | 314 | Gln | Glu | 1.109 | 0.003 |
| p.Gln314Gly | substitution | 314 | Gln | Gly | 0.000 | 0.026 |
| p.Gln314His | substitution | 314 | Gln | His | 1.031 | 0.037 |
| p.Gln314Ile | substitution | 314 | Gln | Ile | 0.499 | 0.115 |
| p.Gln314Leu | substitution | 314 | Gln | Leu | 1.002 | 0.032 |
| p.Gln314Met | substitution | 314 | Gln | Met | 0.000 | 0.162 |
| p.Gln314Phe | substitution | 314 | Gln | Phe | 1.066 | 0.003 |
| p.Gln314Pro | substitution | 314 | Gln | Pro | 0.854 | 0.094 |
| p.Gln314Ser | substitution | 314 | Gln | Ser | 0.314 | 0.421 |
| p.Gln314Ter | nonsense | 314 | Gln | Ter | 0.207 | 0.022 |
| p.Gln314Thr | substitution | 314 | Gln | Thr | 0.809 | 0.224 |
| p.Gln314Trp | substitution | 314 | Gln | Trp | 0.810 | 0.003 |
| p.Gln314Tyr | substitution | 314 | Gln | Tyr | 0.000 | 0.016 |
| p.Gln314Val | substitution | 314 | Gln | Val | 0.850 | 0.022 |
| p.Gln328= | synonymous | 328 | Gln | NA | 1.337 | 0.460 |
| p.Gln328Ala | substitution | 328 | Gln | Ala | 0.462 | 0.007 |
| p.Gln328Arg | substitution | 328 | Gln | Arg | 0.000 | 0.794 |
| p.Gln328Asn | substitution | 328 | Gln | Asn | 0.000 | 0.230 |
| p.Gln328Glu | substitution | 328 | Gln | Glu | 1.044 | 0.010 |
| p.Gln328Gly | substitution | 328 | Gln | Gly | 0.700 | 0.003 |
| p.Gln328His | substitution | 328 | Gln | His | 0.913 | 0.042 |
| p.Gln328Leu | substitution | 328 | Gln | Leu | 0.691 | 0.075 |
| p.Gln328Lys | substitution | 328 | Gln | Lys | 1.041 | 0.005 |

| | | | | | | |
|---|---|---|---|---|---|---|
| p.Gln328Met | substitution | 328 | Gln | Met | 1.301 | 0.004 |
| p.Gln328Pro | substitution | 328 | Gln | Pro | 0.808 | 0.019 |
| p.Gln328Ser | substitution | 328 | Gln | Ser | 1.511 | 0.142 |
| p.Gln328Ter | nonsense | 328 | Gln | Ter | 0.072 | 0.016 |
| p.Gln328Trp | substitution | 328 | Gln | Trp | 1.134 | 0.005 |
| p.Gln328Val | substitution | 328 | Gln | Val | 0.302 | 0.056 |
| p.Gln332= | synonymous | 332 | Gln | NA | 0.000 | 1.175 |
| p.Gln332Arg | substitution | 332 | Gln | Arg | 0.857 | 0.200 |
| p.Gln332Asn | substitution | 332 | Gln | Asn | 0.455 | 0.040 |
| p.Gln332Asp | substitution | 332 | Gln | Asp | 0.584 | 0.044 |
| p.Gln332Cys | substitution | 332 | Gln | Cys | 0.000 | 0.226 |
| p.Gln332Glu | substitution | 332 | Gln | Glu | 0.980 | 0.078 |
| p.Gln332Gly | substitution | 332 | Gln | Gly | 0.632 | 0.012 |
| p.Gln332His | substitution | 332 | Gln | His | 1.252 | 0.021 |
| p.Gln332Ile | substitution | 332 | Gln | Ile | 0.000 | 0.082 |
| p.Gln332Leu | substitution | 332 | Gln | Leu | 0.662 | 0.173 |
| p.Gln332Lys | substitution | 332 | Gln | Lys | 1.241 | 0.002 |
| p.Gln332Met | substitution | 332 | Gln | Met | 1.367 | 0.002 |
| p.Gln332Phe | substitution | 332 | Gln | Phe | 0.271 | 0.006 |
| p.Gln332Pro | substitution | 332 | Gln | Pro | 0.362 | 0.070 |
| p.Gln332Ser | substitution | 332 | Gln | Ser | 1.294 | 0.068 |
| p.Gln332Ter | nonsense | 332 | Gln | Ter | 0.121 | 0.320 |
| p.Gln332Thr | substitution | 332 | Gln | Thr | 1.061 | 0.112 |
| p.Gln332Trp | substitution | 332 | Gln | Trp | 1.056 | 0.007 |
| p.Gln332Tyr | substitution | 332 | Gln | Tyr | 1.074 | 0.006 |
| p.Gln34= | synonymous | 34 | Gln | NA | 1.146 | 0.029 |
| p.Gln34Ala | substitution | 34 | Gln | Ala | 0.623 | 0.024 |
| p.Gln34Arg | substitution | 34 | Gln | Arg | 0.000 | 0.001 |
| p.Gln34Asn | substitution | 34 | Gln | Asn | 0.755 | 0.001 |
| p.Gln34Asp | substitution | 34 | Gln | Asp | 0.086 | 0.001 |
| p.Gln34Cys | substitution | 34 | Gln | Cys | 0.387 | 0.001 |
| p.Gln34Glu | substitution | 34 | Gln | Glu | 0.000 | 0.002 |
| p.Gln34Gly | substitution | 34 | Gln | Gly | 0.567 | 0.001 |
| p.Gln34His | substitution | 34 | Gln | His | 0.332 | 0.001 |
| p.Gln34Ile | substitution | 34 | Gln | Ile | 0.424 | 0.001 |
| p.Gln34Leu | substitution | 34 | Gln | Leu | 0.281 | 0.002 |
| p.Gln34Lys | substitution | 34 | Gln | Lys | 0.000 | 0.010 |
| p.Gln34Met | substitution | 34 | Gln | Met | 0.614 | 0.001 |
| p.Gln34Phe | substitution | 34 | Gln | Phe | 0.183 | 0.001 |
| p.Gln34Pro | substitution | 34 | Gln | Pro | 0.000 | 0.063 |
| p.Gln34Ser | substitution | 34 | Gln | Ser | 0.825 | 0.001 |
| p.Gln34Ter | nonsense | 34 | Gln | Ter | 0.000 | 0.025 |
| p.Gln34Thr | substitution | 34 | Gln | Thr | 0.334 | 0.002 |

| | | | | | | | |
|---|---|---|---|---|---|---|---|
| p.Gln34Trp | substitution | 34 | Gln | Trp | | 0.000 | 0.002 |
| p.Gln34Tyr | substitution | 34 | Gln | Tyr | | 0.079 | 0.001 |
| p.Gln34Val | substitution | 34 | Gln | Val | | 0.419 | 0.018 |
| p.Gln356Asn | substitution | 356 | Gln | Asn | | 1.544 | 0.011 |
| p.Gln356Cys | substitution | 356 | Gln | Cys | | 0.000 | 0.202 |
| p.Gln356Glu | substitution | 356 | Gln | Glu | | 1.074 | 0.201 |
| p.Gln356His | substitution | 356 | Gln | His | | 1.069 | 0.021 |
| p.Gln356Ile | substitution | 356 | Gln | Ile | | 0.000 | 0.253 |
| p.Gln356Leu | substitution | 356 | Gln | Leu | | 0.772 | 0.142 |
| p.Gln356Lys | substitution | 356 | Gln | Lys | | 0.903 | 0.154 |
| p.Gln356Met | substitution | 356 | Gln | Met | | 0.000 | 0.123 |
| p.Gln356Phe | substitution | 356 | Gln | Phe | | 0.083 | 0.078 |
| p.Gln356Pro | substitution | 356 | Gln | Pro | | 1.471 | 0.003 |
| p.Gln356Ter | nonsense | 356 | Gln | Ter | | 0.672 | 0.016 |
| p.Gln356Thr | substitution | 356 | Gln | Thr | | 0.036 | 0.082 |
| p.Gln356Tyr | substitution | 356 | Gln | Tyr | | 0.350 | 0.053 |
| p.Gln50= | synonymous | 50 | Gln | NA | | 1.337 | 0.014 |
| p.Gln50Arg | substitution | 50 | Gln | Arg | | 0.853 | 0.087 |
| p.Gln50Asp | substitution | 50 | Gln | Asp | | 0.044 | 0.208 |
| p.Gln50Cys | substitution | 50 | Gln | Cys | | 0.000 | 0.174 |
| p.Gln50Glu | substitution | 50 | Gln | Glu | | 1.358 | 0.004 |
| p.Gln50Gly | substitution | 50 | Gln | Gly | | 0.375 | 0.102 |
| p.Gln50His | substitution | 50 | Gln | His | | 0.953 | 0.068 |
| p.Gln50Ile | substitution | 50 | Gln | Ile | | 0.681 | 0.081 |
| p.Gln50Leu | substitution | 50 | Gln | Leu | | 1.009 | 0.013 |
| p.Gln50Lys | substitution | 50 | Gln | Lys | | 1.187 | 0.041 |
| p.Gln50Met | substitution | 50 | Gln | Met | | 1.266 | 0.022 |
| p.Gln50Phe | substitution | 50 | Gln | Phe | | 0.661 | 0.030 |
| p.Gln50Pro | substitution | 50 | Gln | Pro | | 1.306 | 0.072 |
| p.Gln50Ser | substitution | 50 | Gln | Ser | | 0.000 | 0.300 |
| p.Gln50Ter | nonsense | 50 | Gln | Ter | | 0.091 | 0.009 |
| p.Gln50Thr | substitution | 50 | Gln | Thr | | 0.785 | 0.061 |
| p.Gln50Trp | substitution | 50 | Gln | Trp | | 0.000 | 0.159 |
| p.Gln50Tyr | substitution | 50 | Gln | Tyr | | 0.986 | 0.036 |
| p.Gln50Val | substitution | 50 | Gln | Val | | 0.000 | 0.126 |
| p.Glu117= | synonymous | 117 | Glu | NA | | 0.814 | 0.017 |
| p.Glu117Ala | substitution | 117 | Glu | Ala | | 0.513 | 0.077 |
| p.Glu117Arg | substitution | 117 | Glu | Arg | | 0.505 | 0.054 |
| p.Glu117Asn | substitution | 117 | Glu | Asn | | 0.390 | 0.028 |
| p.Glu117Asp | substitution | 117 | Glu | Asp | | 0.971 | 0.036 |
| p.Glu117Cys | substitution | 117 | Glu | Cys | | 1.026 | 0.008 |
| p.Glu117Gln | substitution | 117 | Glu | Gln | | 0.764 | 0.130 |
| p.Glu117Gly | substitution | 117 | Glu | Gly | | 0.897 | 0.012 |

| | | | | | | |
|---|---|---|---|---|---|---|
| p.Glu117His | substitution | 117 | Glu | His | 0.876 | 0.014 |
| p.Glu117Ile | substitution | 117 | Glu | Ile | 1.342 | 0.012 |
| p.Glu117Leu | substitution | 117 | Glu | Leu | 0.783 | 0.026 |
| p.Glu117Lys | substitution | 117 | Glu | Lys | 0.817 | 0.235 |
| p.Glu117Met | substitution | 117 | Glu | Met | 0.449 | 0.019 |
| p.Glu117Phe | substitution | 117 | Glu | Phe | 0.964 | 0.011 |
| p.Glu117Pro | substitution | 117 | Glu | Pro | 0.000 | 0.245 |
| p.Glu117Ser | substitution | 117 | Glu | Ser | 0.404 | 0.033 |
| p.Glu117Ter | nonsense | 117 | Glu | Ter | 0.074 | 0.029 |
| p.Glu117Thr | substitution | 117 | Glu | Thr | 0.421 | 0.018 |
| p.Glu117Trp | substitution | 117 | Glu | Trp | 0.887 | 0.039 |
| p.Glu117Tyr | substitution | 117 | Glu | Tyr | 0.896 | 0.015 |
| p.Glu117Val | substitution | 117 | Glu | Val | 1.108 | 0.033 |
| p.Glu135= | synonymous | 135 | Glu | NA | 0.833 | 0.005 |
| p.Glu135Ala | substitution | 135 | Glu | Ala | 1.043 | 0.010 |
| p.Glu135Arg | substitution | 135 | Glu | Arg | 1.039 | 0.017 |
| p.Glu135Asn | substitution | 135 | Glu | Asn | 0.610 | 0.006 |
| p.Glu135Asp | substitution | 135 | Glu | Asp | 0.916 | 0.011 |
| p.Glu135Cys | substitution | 135 | Glu | Cys | 0.717 | 0.007 |
| p.Glu135Gln | substitution | 135 | Glu | Gln | 1.118 | 0.043 |
| p.Glu135Gly | substitution | 135 | Glu | Gly | 0.707 | 0.020 |
| p.Glu135His | substitution | 135 | Glu | His | 0.133 | 0.022 |
| p.Glu135Ile | substitution | 135 | Glu | Ile | 0.873 | 0.015 |
| p.Glu135Leu | substitution | 135 | Glu | Leu | 0.992 | 0.033 |
| p.Glu135Lys | substitution | 135 | Glu | Lys | 0.226 | 0.022 |
| p.Glu135Met | substitution | 135 | Glu | Met | 1.113 | 0.007 |
| p.Glu135Phe | substitution | 135 | Glu | Phe | 0.871 | 0.005 |
| p.Glu135Pro | substitution | 135 | Glu | Pro | 0.865 | 0.015 |
| p.Glu135Ser | substitution | 135 | Glu | Ser | 0.630 | 0.013 |
| p.Glu135Ter | nonsense | 135 | Glu | Ter | 0.246 | 0.053 |
| p.Glu135Thr | substitution | 135 | Glu | Thr | 0.843 | 0.097 |
| p.Glu135Trp | substitution | 135 | Glu | Trp | 0.882 | 0.006 |
| p.Glu135Tyr | substitution | 135 | Glu | Tyr | 0.718 | 0.007 |
| p.Glu135Val | substitution | 135 | Glu | Val | 1.110 | 0.003 |
| p.Glu139= | synonymous | 139 | Glu | NA | 0.010 | 3.475 |
| p.Glu139Ala | substitution | 139 | Glu | Ala | 1.061 | 0.005 |
| p.Glu139Arg | substitution | 139 | Glu | Arg | 0.627 | 0.020 |
| p.Glu139Asn | substitution | 139 | Glu | Asn | 1.035 | 0.007 |
| p.Glu139Asp | substitution | 139 | Glu | Asp | 1.043 | 0.025 |
| p.Glu139Cys | substitution | 139 | Glu | Cys | 1.086 | 0.005 |
| p.Glu139Gln | substitution | 139 | Glu | Gln | 1.260 | 0.010 |
| p.Glu139Gly | substitution | 139 | Glu | Gly | 0.816 | 0.003 |
| p.Glu139His | substitution | 139 | Glu | His | 1.175 | 0.021 |

| | | | | | | | |
|---|---|---|---|---|---|---|---|
| p.Glu139Ile | substitution | 139 | Glu | Ile | | 1.174 | 0.006 |
| p.Glu139Leu | substitution | 139 | Glu | Leu | | 1.123 | 0.009 |
| p.Glu139Lys | substitution | 139 | Glu | Lys | | 1.215 | 0.004 |
| p.Glu139Met | substitution | 139 | Glu | Met | | 0.663 | 0.005 |
| p.Glu139Phe | substitution | 139 | Glu | Phe | | 1.091 | 0.004 |
| p.Glu139Pro | substitution | 139 | Glu | Pro | | 0.000 | 0.097 |
| p.Glu139Ser | substitution | 139 | Glu | Ser | | 0.753 | 0.004 |
| p.Glu139Ter | nonsense | 139 | Glu | Ter | | 0.422 | 0.008 |
| p.Glu139Thr | substitution | 139 | Glu | Thr | | 0.614 | 0.013 |
| p.Glu139Trp | substitution | 139 | Glu | Trp | | 1.206 | 0.005 |
| p.Glu139Tyr | substitution | 139 | Glu | Tyr | | 0.982 | 0.005 |
| p.Glu139Val | substitution | 139 | Glu | Val | | 0.621 | 0.005 |
| p.Glu162= | synonymous | 162 | Glu | NA | | 1.175 | 0.024 |
| p.Glu162Ala | substitution | 162 | Glu | Ala | | 0.560 | 0.022 |
| p.Glu162Arg | substitution | 162 | Glu | Arg | | 0.929 | 0.072 |
| p.Glu162Asn | substitution | 162 | Glu | Asn | | 0.346 | 0.018 |
| p.Glu162Asp | substitution | 162 | Glu | Asp | | 0.842 | 0.026 |
| p.Glu162Cys | substitution | 162 | Glu | Cys | | 1.387 | 0.015 |
| p.Glu162Gln | substitution | 162 | Glu | Gln | | 0.502 | 0.048 |
| p.Glu162Gly | substitution | 162 | Glu | Gly | | 0.694 | 0.163 |
| p.Glu162His | substitution | 162 | Glu | His | | 0.394 | 0.216 |
| p.Glu162Ile | substitution | 162 | Glu | Ile | | 0.914 | 0.010 |
| p.Glu162Leu | substitution | 162 | Glu | Leu | | 0.985 | 0.042 |
| p.Glu162Lys | substitution | 162 | Glu | Lys | | 1.117 | 0.017 |
| p.Glu162Met | substitution | 162 | Glu | Met | | 0.887 | 0.014 |
| p.Glu162Phe | substitution | 162 | Glu | Phe | | 1.165 | 0.009 |
| p.Glu162Pro | substitution | 162 | Glu | Pro | | 0.000 | 0.273 |
| p.Glu162Ser | substitution | 162 | Glu | Ser | | 0.409 | 0.032 |
| p.Glu162Ter | nonsense | 162 | Glu | Ter | | 0.540 | 0.020 |
| p.Glu162Thr | substitution | 162 | Glu | Thr | | 0.237 | 0.308 |
| p.Glu162Trp | substitution | 162 | Glu | Trp | | 0.060 | 0.018 |
| p.Glu162Tyr | substitution | 162 | Glu | Tyr | | 1.362 | 0.007 |
| p.Glu162Val | substitution | 162 | Glu | Val | | 1.096 | 0.009 |
| p.Glu179= | synonymous | 179 | Glu | NA | | 1.193 | 0.020 |
| p.Glu179Ala | substitution | 179 | Glu | Ala | | 1.094 | 0.026 |
| p.Glu179Arg | substitution | 179 | Glu | Arg | | 0.722 | 0.036 |
| p.Glu179Asn | substitution | 179 | Glu | Asn | | 0.947 | 0.007 |
| p.Glu179Asp | substitution | 179 | Glu | Asp | | 0.556 | 0.035 |
| p.Glu179Cys | substitution | 179 | Glu | Cys | | 0.811 | 0.005 |
| p.Glu179Gln | substitution | 179 | Glu | Gln | | 0.954 | 0.007 |
| p.Glu179Gly | substitution | 179 | Glu | Gly | | 0.613 | 0.008 |
| p.Glu179His | substitution | 179 | Glu | His | | 0.718 | 0.009 |
| p.Glu179Ile | substitution | 179 | Glu | Ile | | 1.267 | 0.007 |

| | | | | | | |
|---|---|---|---|---|---|---|
| p.Glu179Leu | substitution | 179 | Glu | Leu | 0.705 | 0.022 |
| p.Glu179Lys | substitution | 179 | Glu | Lys | 0.691 | 0.031 |
| p.Glu179Met | substitution | 179 | Glu | Met | 0.796 | 0.008 |
| p.Glu179Phe | substitution | 179 | Glu | Phe | 0.529 | 0.011 |
| p.Glu179Pro | substitution | 179 | Glu | Pro | 0.459 | 0.018 |
| p.Glu179Ser | substitution | 179 | Glu | Ser | 0.865 | 0.054 |
| p.Glu179Ter | nonsense | 179 | Glu | Ter | 0.363 | 0.014 |
| p.Glu179Thr | substitution | 179 | Glu | Thr | 0.864 | 0.011 |
| p.Glu179Trp | substitution | 179 | Glu | Trp | 0.758 | 0.008 |
| p.Glu179Tyr | substitution | 179 | Glu | Tyr | 0.812 | 0.012 |
| p.Glu179Val | substitution | 179 | Glu | Val | 1.074 | 0.007 |
| p.Glu182= | synonymous | 182 | Glu | NA | 0.909 | 0.022 |
| p.Glu182Ala | substitution | 182 | Glu | Ala | 1.006 | 0.039 |
| p.Glu182Asp | substitution | 182 | Glu | Asp | 0.720 | 0.082 |
| p.Glu182Gly | substitution | 182 | Glu | Gly | 1.529 | 0.039 |
| p.Glu182Lys | substitution | 182 | Glu | Lys | 0.509 | 0.532 |
| p.Glu182Ter | nonsense | 182 | Glu | Ter | 0.333 | 0.068 |
| p.Glu182Val | substitution | 182 | Glu | Val | 1.440 | 0.385 |
| p.Glu209= | synonymous | 209 | Glu | NA | 0.116 | 0.266 |
| p.Glu209Ala | substitution | 209 | Glu | Ala | 0.024 | 0.145 |
| p.Glu209Arg | substitution | 209 | Glu | Arg | 0.748 | 0.197 |
| p.Glu209Asn | substitution | 209 | Glu | Asn | 0.000 | 0.009 |
| p.Glu209Asp | substitution | 209 | Glu | Asp | 0.751 | 0.048 |
| p.Glu209Cys | substitution | 209 | Glu | Cys | 1.109 | 0.007 |
| p.Glu209Gln | substitution | 209 | Glu | Gln | 1.269 | 0.012 |
| p.Glu209Gly | substitution | 209 | Glu | Gly | 0.604 | 0.097 |
| p.Glu209His | substitution | 209 | Glu | His | 0.766 | 0.006 |
| p.Glu209Ile | substitution | 209 | Glu | Ile | 0.000 | 0.137 |
| p.Glu209Leu | substitution | 209 | Glu | Leu | 0.274 | 0.095 |
| p.Glu209Met | substitution | 209 | Glu | Met | 0.316 | 0.310 |
| p.Glu209Phe | substitution | 209 | Glu | Phe | 1.316 | 0.001 |
| p.Glu209Pro | substitution | 209 | Glu | Pro | 1.451 | 0.019 |
| p.Glu209Ser | substitution | 209 | Glu | Ser | 0.362 | 0.076 |
| p.Glu209Ter | nonsense | 209 | Glu | Ter | 0.113 | 0.017 |
| p.Glu209Thr | substitution | 209 | Glu | Thr | 0.063 | 0.056 |
| p.Glu209Trp | substitution | 209 | Glu | Trp | 0.000 | 0.010 |
| p.Glu209Tyr | substitution | 209 | Glu | Tyr | 1.108 | 0.063 |
| p.Glu209Val | substitution | 209 | Glu | Val | 0.064 | 0.039 |
| p.Glu210= | synonymous | 210 | Glu | NA | 0.713 | 0.268 |
| p.Glu210Ala | substitution | 210 | Glu | Ala | 0.587 | 0.058 |
| p.Glu210Arg | substitution | 210 | Glu | Arg | 1.004 | 0.029 |
| p.Glu210Asn | substitution | 210 | Glu | Asn | 0.994 | 0.013 |
| p.Glu210Asp | substitution | 210 | Glu | Asp | 0.702 | 0.106 |

| | | | | | | | |
|---|---|---|---|---|---|---|---|
| p.Glu210Cys | substitution | 210 | Glu | Cys | | 1.325 | 0.002 |
| p.Glu210Gln | substitution | 210 | Glu | Gln | | 0.335 | 0.041 |
| p.Glu210Gly | substitution | 210 | Glu | Gly | | 0.160 | 0.079 |
| p.Glu210His | substitution | 210 | Glu | His | | 0.000 | 0.062 |
| p.Glu210Ile | substitution | 210 | Glu | Ile | | 1.627 | 0.007 |
| p.Glu210Leu | substitution | 210 | Glu | Leu | | 1.064 | 0.015 |
| p.Glu210Lys | substitution | 210 | Glu | Lys | | 0.000 | 0.011 |
| p.Glu210Met | substitution | 210 | Glu | Met | | 0.000 | 0.015 |
| p.Glu210Pro | substitution | 210 | Glu | Pro | | 0.000 | 0.015 |
| p.Glu210Ser | substitution | 210 | Glu | Ser | | 0.264 | 0.027 |
| p.Glu210Ter | nonsense | 210 | Glu | Ter | | 0.284 | 0.106 |
| p.Glu210Thr | substitution | 210 | Glu | Thr | | 0.599 | 0.124 |
| p.Glu210Trp | substitution | 210 | Glu | Trp | | 1.317 | 0.005 |
| p.Glu210Tyr | substitution | 210 | Glu | Tyr | | 0.331 | 0.074 |
| p.Glu210Val | substitution | 210 | Glu | Val | | 0.909 | 0.004 |
| p.Glu223= | synonymous | 223 | Glu | NA | | 0.504 | 0.525 |
| p.Glu223Ala | substitution | 223 | Glu | Ala | | 0.072 | 0.130 |
| p.Glu223Arg | substitution | 223 | Glu | Arg | | 0.000 | 0.045 |
| p.Glu223Asn | substitution | 223 | Glu | Asn | | 0.870 | 0.005 |
| p.Glu223Asp | substitution | 223 | Glu | Asp | | 0.595 | 0.419 |
| p.Glu223Cys | substitution | 223 | Glu | Cys | | 1.209 | 0.006 |
| p.Glu223Gln | substitution | 223 | Glu | Gln | | 0.054 | 0.298 |
| p.Glu223Gly | substitution | 223 | Glu | Gly | | 0.568 | 0.103 |
| p.Glu223His | substitution | 223 | Glu | His | | 0.463 | 0.078 |
| p.Glu223Ile | substitution | 223 | Glu | Ile | | 0.053 | 0.039 |
| p.Glu223Leu | substitution | 223 | Glu | Leu | | 0.832 | 0.107 |
| p.Glu223Lys | substitution | 223 | Glu | Lys | | 0.000 | 0.224 |
| p.Glu223Met | substitution | 223 | Glu | Met | | 0.000 | 0.194 |
| p.Glu223Phe | substitution | 223 | Glu | Phe | | 0.000 | 0.074 |
| p.Glu223Pro | substitution | 223 | Glu | Pro | | 0.138 | 0.044 |
| p.Glu223Ser | substitution | 223 | Glu | Ser | | 0.067 | 0.041 |
| p.Glu223Ter | nonsense | 223 | Glu | Ter | | 0.452 | 0.194 |
| p.Glu223Thr | substitution | 223 | Glu | Thr | | 0.568 | 0.138 |
| p.Glu223Trp | substitution | 223 | Glu | Trp | | 0.000 | 0.013 |
| p.Glu223Tyr | substitution | 223 | Glu | Tyr | | 0.000 | 0.013 |
| p.Glu223Val | substitution | 223 | Glu | Val | | 0.280 | 0.181 |
| p.Glu242Ala | substitution | 242 | Glu | Ala | | 1.100 | 0.026 |
| p.Glu242Arg | substitution | 242 | Glu | Arg | | 0.728 | 0.125 |
| p.Glu242Asn | substitution | 242 | Glu | Asn | | 0.000 | 0.182 |
| p.Glu242Asp | substitution | 242 | Glu | Asp | | 0.958 | 0.003 |
| p.Glu242Cys | substitution | 242 | Glu | Cys | | 0.009 | 0.079 |
| p.Glu242Gln | substitution | 242 | Glu | Gln | | 0.794 | 0.004 |
| p.Glu242Gly | substitution | 242 | Glu | Gly | | 0.025 | 0.024 |

| | | | | | | | |
|---|---|---|---|---|---|---|---|
| p.Glu242His | substitution | 242 | Glu | His | | 1.335 | 0.003 |
| p.Glu242Ile | substitution | 242 | Glu | Ile | | 0.000 | 0.011 |
| p.Glu242Leu | substitution | 242 | Glu | Leu | | 0.856 | 0.330 |
| p.Glu242Lys | substitution | 242 | Glu | Lys | | 0.834 | 0.004 |
| p.Glu242Met | substitution | 242 | Glu | Met | | 0.302 | 0.005 |
| p.Glu242Phe | substitution | 242 | Glu | Phe | | 1.484 | 0.008 |
| p.Glu242Pro | substitution | 242 | Glu | Pro | | 1.076 | 0.223 |
| p.Glu242Ser | substitution | 242 | Glu | Ser | | 0.134 | 0.026 |
| p.Glu242Ter | nonsense | 242 | Glu | Ter | | 0.273 | 0.005 |
| p.Glu242Thr | substitution | 242 | Glu | Thr | | 0.607 | 0.164 |
| p.Glu242Tyr | substitution | 242 | Glu | Tyr | | 0.000 | 0.027 |
| p.Glu242Val | substitution | 242 | Glu | Val | | 1.107 | 0.005 |
| p.Glu250= | synonymous | 250 | Glu | NA | | 1.296 | 0.003 |
| p.Glu250Ala | substitution | 250 | Glu | Ala | | 0.000 | 0.202 |
| p.Glu250Arg | substitution | 250 | Glu | Arg | | 0.000 | 0.017 |
| p.Glu250Asp | substitution | 250 | Glu | Asp | | 1.342 | 0.306 |
| p.Glu250Cys | substitution | 250 | Glu | Cys | | 0.000 | 0.019 |
| p.Glu250Gln | substitution | 250 | Glu | Gln | | 0.740 | 0.017 |
| p.Glu250Gly | substitution | 250 | Glu | Gly | | 0.000 | 0.013 |
| p.Glu250His | substitution | 250 | Glu | His | | 0.000 | 0.018 |
| p.Glu250Ile | substitution | 250 | Glu | Ile | | 0.000 | 0.040 |
| p.Glu250Leu | substitution | 250 | Glu | Leu | | 0.000 | 0.040 |
| p.Glu250Lys | substitution | 250 | Glu | Lys | | 0.018 | 0.204 |
| p.Glu250Met | substitution | 250 | Glu | Met | | 0.000 | 0.032 |
| p.Glu250Phe | substitution | 250 | Glu | Phe | | 0.000 | 0.086 |
| p.Glu250Pro | substitution | 250 | Glu | Pro | | 0.000 | 0.031 |
| p.Glu250Ser | substitution | 250 | Glu | Ser | | 0.000 | 0.023 |
| p.Glu250Ter | nonsense | 250 | Glu | Ter | | 0.000 | 0.066 |
| p.Glu250Thr | substitution | 250 | Glu | Thr | | 0.068 | 0.069 |
| p.Glu250Trp | substitution | 250 | Glu | Trp | | 0.000 | 0.008 |
| p.Glu250Tyr | substitution | 250 | Glu | Tyr | | 0.000 | 0.042 |
| p.Glu250Val | substitution | 250 | Glu | Val | | 0.000 | 0.064 |
| p.Glu258= | synonymous | 258 | Glu | NA | | 0.703 | 0.007 |
| p.Glu258Ala | substitution | 258 | Glu | Ala | | 0.119 | 0.069 |
| p.Glu258Arg | substitution | 258 | Glu | Arg | | 0.000 | 0.146 |
| p.Glu258Asn | substitution | 258 | Glu | Asn | | 0.000 | 0.023 |
| p.Glu258Asp | substitution | 258 | Glu | Asp | | 1.263 | 0.360 |
| p.Glu258Cys | substitution | 258 | Glu | Cys | | 1.326 | 0.002 |
| p.Glu258Gly | substitution | 258 | Glu | Gly | | 1.303 | 0.048 |
| p.Glu258Ile | substitution | 258 | Glu | Ile | | 1.007 | 0.009 |
| p.Glu258Leu | substitution | 258 | Glu | Leu | | 0.027 | 0.072 |
| p.Glu258Lys | substitution | 258 | Glu | Lys | | 1.169 | 0.124 |
| p.Glu258Met | substitution | 258 | Glu | Met | | 0.007 | 0.010 |

| | | | | | | |
|---|---|---|---|---|---|---|
| p.Glu258Phe | substitution | 258 | Glu | Phe | 0.475 | 0.013 |
| p.Glu258Pro | substitution | 258 | Glu | Pro | 0.391 | 0.038 |
| p.Glu258Ser | substitution | 258 | Glu | Ser | 0.580 | 0.045 |
| p.Glu258Ter | nonsense | 258 | Glu | Ter | 0.313 | 0.081 |
| p.Glu258Thr | substitution | 258 | Glu | Thr | 0.000 | 0.033 |
| p.Glu258Trp | substitution | 258 | Glu | Trp | 0.061 | 0.232 |
| p.Glu258Tyr | substitution | 258 | Glu | Tyr | 1.053 | 0.005 |
| p.Glu258Val | substitution | 258 | Glu | Val | 1.326 | 0.229 |
| p.Glu293= | synonymous | 293 | Glu | NA | 1.412 | 0.161 |
| p.Glu293Ala | substitution | 293 | Glu | Ala | 0.858 | 0.007 |
| p.Glu293Arg | substitution | 293 | Glu | Arg | 0.570 | 0.021 |
| p.Glu293Asn | substitution | 293 | Glu | Asn | 0.688 | 0.004 |
| p.Glu293Asp | substitution | 293 | Glu | Asp | 1.116 | 0.014 |
| p.Glu293Cys | substitution | 293 | Glu | Cys | 0.597 | 0.003 |
| p.Glu293Gln | substitution | 293 | Glu | Gln | 1.140 | 0.009 |
| p.Glu293Gly | substitution | 293 | Glu | Gly | 1.099 | 0.004 |
| p.Glu293His | substitution | 293 | Glu | His | 1.059 | 0.005 |
| p.Glu293Ile | substitution | 293 | Glu | Ile | 0.728 | 0.004 |
| p.Glu293Leu | substitution | 293 | Glu | Leu | 0.445 | 0.017 |
| p.Glu293Lys | substitution | 293 | Glu | Lys | 0.889 | 0.005 |
| p.Glu293Phe | substitution | 293 | Glu | Phe | 0.588 | 0.004 |
| p.Glu293Pro | substitution | 293 | Glu | Pro | 0.000 | 0.015 |
| p.Glu293Ser | substitution | 293 | Glu | Ser | 0.942 | 0.008 |
| p.Glu293Ter | nonsense | 293 | Glu | Ter | 0.203 | 0.011 |
| p.Glu293Thr | substitution | 293 | Glu | Thr | 0.079 | 0.037 |
| p.Glu293Trp | substitution | 293 | Glu | Trp | 1.136 | 0.004 |
| p.Glu293Tyr | substitution | 293 | Glu | Tyr | 0.540 | 0.005 |
| p.Glu293Val | substitution | 293 | Glu | Val | 0.535 | 0.005 |
| p.Glu306= | synonymous | 306 | Glu | NA | 1.123 | 0.003 |
| p.Glu306Ala | substitution | 306 | Glu | Ala | 1.235 | 0.005 |
| p.Glu306Arg | substitution | 306 | Glu | Arg | 0.951 | 0.017 |
| p.Glu306Asn | substitution | 306 | Glu | Asn | 1.459 | 0.002 |
| p.Glu306Asp | substitution | 306 | Glu | Asp | 0.524 | 0.003 |
| p.Glu306Cys | substitution | 306 | Glu | Cys | 0.135 | 0.015 |
| p.Glu306Gln | substitution | 306 | Glu | Gln | 1.006 | 0.003 |
| p.Glu306Gly | substitution | 306 | Glu | Gly | 0.498 | 0.057 |
| p.Glu306His | substitution | 306 | Glu | His | 0.953 | 0.005 |
| p.Glu306Ile | substitution | 306 | Glu | Ile | 0.912 | 0.006 |
| p.Glu306Leu | substitution | 306 | Glu | Leu | 1.114 | 0.022 |
| p.Glu306Lys | substitution | 306 | Glu | Lys | 1.330 | 0.015 |
| p.Glu306Met | substitution | 306 | Glu | Met | 0.172 | 0.008 |
| p.Glu306Phe | substitution | 306 | Glu | Phe | 1.441 | 0.003 |
| p.Glu306Pro | substitution | 306 | Glu | Pro | 0.524 | 0.066 |

| | | | | | | |
|---|---|---|---|---|---|---|
| p.Glu306Ser | substitution | 306 | Glu | Ser | 0.844 | 0.165 |
| p.Glu306Ter | nonsense | 306 | Glu | Ter | 0.461 | 0.063 |
| p.Glu306Thr | substitution | 306 | Glu | Thr | 0.033 | 0.009 |
| p.Glu306Trp | substitution | 306 | Glu | Trp | 0.903 | 0.003 |
| p.Glu306Tyr | substitution | 306 | Glu | Tyr | 0.307 | 0.013 |
| p.Glu306Val | substitution | 306 | Glu | Val | 1.133 | 0.006 |
| p.Glu310Ala | substitution | 310 | Glu | Ala | 1.064 | 0.120 |
| p.Glu310Arg | substitution | 310 | Glu | Arg | 1.020 | 0.044 |
| p.Glu310Asn | substitution | 310 | Glu | Asn | 0.000 | 0.047 |
| p.Glu310Asp | substitution | 310 | Glu | Asp | 0.408 | 0.062 |
| p.Glu310Gln | substitution | 310 | Glu | Gln | 1.264 | 0.003 |
| p.Glu310Gly | substitution | 310 | Glu | Gly | 0.959 | 0.057 |
| p.Glu310His | substitution | 310 | Glu | His | 1.466 | 0.001 |
| p.Glu310Ile | substitution | 310 | Glu | Ile | 0.000 | 0.018 |
| p.Glu310Leu | substitution | 310 | Glu | Leu | 0.775 | 0.056 |
| p.Glu310Lys | substitution | 310 | Glu | Lys | 0.697 | 0.005 |
| p.Glu310Met | substitution | 310 | Glu | Met | 1.358 | 0.008 |
| p.Glu310Phe | substitution | 310 | Glu | Phe | 0.000 | 0.044 |
| p.Glu310Pro | substitution | 310 | Glu | Pro | 0.066 | 0.030 |
| p.Glu310Ser | substitution | 310 | Glu | Ser | 0.254 | 0.207 |
| p.Glu310Ter | nonsense | 310 | Glu | Ter | 0.425 | 0.004 |
| p.Glu310Thr | substitution | 310 | Glu | Thr | 0.000 | 0.316 |
| p.Glu310Trp | substitution | 310 | Glu | Trp | 0.087 | 0.009 |
| p.Glu310Tyr | substitution | 310 | Glu | Tyr | 0.000 | 0.020 |
| p.Glu310Val | substitution | 310 | Glu | Val | 0.714 | 1.008 |
| p.Glu52= | synonymous | 52 | Glu | NA | 0.525 | 0.029 |
| p.Glu52Ala | substitution | 52 | Glu | Ala | 1.080 | 0.037 |
| p.Glu52Arg | substitution | 52 | Glu | Arg | 0.989 | 0.044 |
| p.Glu52Asn | substitution | 52 | Glu | Asn | 0.184 | 0.014 |
| p.Glu52Asp | substitution | 52 | Glu | Asp | 0.904 | 0.004 |
| p.Glu52Cys | substitution | 52 | Glu | Cys | 1.045 | 0.011 |
| p.Glu52Gln | substitution | 52 | Glu | Gln | 0.000 | 0.293 |
| p.Glu52Gly | substitution | 52 | Glu | Gly | 0.921 | 0.012 |
| p.Glu52His | substitution | 52 | Glu | His | 0.267 | 0.058 |
| p.Glu52Ile | substitution | 52 | Glu | Ile | 0.992 | 0.008 |
| p.Glu52Leu | substitution | 52 | Glu | Leu | 0.259 | 0.048 |
| p.Glu52Lys | substitution | 52 | Glu | Lys | 0.342 | 0.026 |
| p.Glu52Met | substitution | 52 | Glu | Met | 0.000 | 0.024 |
| p.Glu52Phe | substitution | 52 | Glu | Phe | 0.892 | 0.025 |
| p.Glu52Pro | substitution | 52 | Glu | Pro | 0.561 | 0.071 |
| p.Glu52Ser | substitution | 52 | Glu | Ser | 0.910 | 0.101 |
| p.Glu52Ter | nonsense | 52 | Glu | Ter | 0.000 | 0.070 |
| p.Glu52Thr | substitution | 52 | Glu | Thr | 0.878 | 0.034 |

| | | | | | | | |
|---|---|---|---|---|---|---|---|
| p.Glu52Trp | substitution | 52 | Glu | Trp | | 0.228 | 0.015 |
| p.Glu52Tyr | substitution | 52 | Glu | Tyr | | 0.803 | 0.020 |
| p.Glu52Val | substitution | 52 | Glu | Val | | 0.946 | 0.051 |
| p.Glu73= | synonymous | 73 | Glu | NA | | 0.359 | 1.589 |
| p.Glu73Ala | substitution | 73 | Glu | Ala | | 0.733 | 0.005 |
| p.Glu73Arg | substitution | 73 | Glu | Arg | | 0.857 | 0.006 |
| p.Glu73Asn | substitution | 73 | Glu | Asn | | 0.498 | 0.016 |
| p.Glu73Asp | substitution | 73 | Glu | Asp | | 1.310 | 1.035 |
| p.Glu73Cys | substitution | 73 | Glu | Cys | | 0.549 | 0.020 |
| p.Glu73Gln | substitution | 73 | Glu | Gln | | 0.719 | 0.017 |
| p.Glu73Gly | substitution | 73 | Glu | Gly | | 0.469 | 0.009 |
| p.Glu73His | substitution | 73 | Glu | His | | 1.290 | 0.006 |
| p.Glu73Ile | substitution | 73 | Glu | Ile | | 0.517 | 0.015 |
| p.Glu73Leu | substitution | 73 | Glu | Leu | | 0.489 | 0.143 |
| p.Glu73Lys | substitution | 73 | Glu | Lys | | 0.780 | 0.011 |
| p.Glu73Met | substitution | 73 | Glu | Met | | 0.651 | 0.009 |
| p.Glu73Phe | substitution | 73 | Glu | Phe | | 0.486 | 0.007 |
| p.Glu73Pro | substitution | 73 | Glu | Pro | | 0.597 | 0.013 |
| p.Glu73Ser | substitution | 73 | Glu | Ser | | 0.909 | 0.018 |
| p.Glu73Ter | nonsense | 73 | Glu | Ter | | 0.236 | 0.009 |
| p.Glu73Thr | substitution | 73 | Glu | Thr | | 1.172 | 0.004 |
| p.Glu73Trp | substitution | 73 | Glu | Trp | | 0.186 | 0.010 |
| p.Glu73Tyr | substitution | 73 | Glu | Tyr | | 0.819 | 0.005 |
| p.Glu73Val | substitution | 73 | Glu | Val | | 0.727 | 0.018 |
| p.Glu80= | synonymous | 80 | Glu | NA | | 0.631 | 0.017 |
| p.Glu80Ala | substitution | 80 | Glu | Ala | | 0.869 | 0.002 |
| p.Glu80Arg | substitution | 80 | Glu | Arg | | 0.730 | 0.004 |
| p.Glu80Asn | substitution | 80 | Glu | Asn | | 0.438 | 0.006 |
| p.Glu80Asp | substitution | 80 | Glu | Asp | | 1.128 | 0.012 |
| p.Glu80Cys | substitution | 80 | Glu | Cys | | 0.893 | 0.002 |
| p.Glu80Gln | substitution | 80 | Glu | Gln | | 0.772 | 0.003 |
| p.Glu80Gly | substitution | 80 | Glu | Gly | | 0.647 | 0.002 |
| p.Glu80His | substitution | 80 | Glu | His | | 0.568 | 0.018 |
| p.Glu80Ile | substitution | 80 | Glu | Ile | | 0.703 | 0.005 |
| p.Glu80Leu | substitution | 80 | Glu | Leu | | 0.490 | 0.009 |
| p.Glu80Lys | substitution | 80 | Glu | Lys | | 0.739 | 0.004 |
| p.Glu80Met | substitution | 80 | Glu | Met | | 0.816 | 0.003 |
| p.Glu80Phe | substitution | 80 | Glu | Phe | | 0.000 | 0.043 |
| p.Glu80Pro | substitution | 80 | Glu | Pro | | 0.625 | 0.005 |
| p.Glu80Ser | substitution | 80 | Glu | Ser | | 0.757 | 0.014 |
| p.Glu80Ter | nonsense | 80 | Glu | Ter | | 0.015 | 0.154 |
| p.Glu80Thr | substitution | 80 | Glu | Thr | | 0.898 | 0.025 |
| p.Glu80Trp | substitution | 80 | Glu | Trp | | 0.583 | 0.003 |

| | | | | | | | |
|---|---|---|---|---|---|---|---|
| p.Glu80Tyr | substitution | 80 | Glu | Tyr | | 0.000 | 0.009 |
| p.Glu80Val | substitution | 80 | Glu | Val | | 0.585 | 0.002 |
| p.Glu82= | synonymous | 82 | Glu | NA | | 0.895 | 0.016 |
| p.Glu82Ala | substitution | 82 | Glu | Ala | | 0.931 | 0.228 |
| p.Glu82Arg | substitution | 82 | Glu | Arg | | 0.624 | 0.023 |
| p.Glu82Asn | substitution | 82 | Glu | Asn | | 0.804 | 0.018 |
| p.Glu82Asp | substitution | 82 | Glu | Asp | | 0.818 | 0.030 |
| p.Glu82Cys | substitution | 82 | Glu | Cys | | 0.479 | 0.019 |
| p.Glu82Gln | substitution | 82 | Glu | Gln | | 0.461 | 0.034 |
| p.Glu82Gly | substitution | 82 | Glu | Gly | | 0.132 | 0.077 |
| p.Glu82His | substitution | 82 | Glu | His | | 0.060 | 0.022 |
| p.Glu82Ile | substitution | 82 | Glu | Ile | | 0.000 | 0.019 |
| p.Glu82Leu | substitution | 82 | Glu | Leu | | 0.603 | 0.032 |
| p.Glu82Lys | substitution | 82 | Glu | Lys | | 0.455 | 0.017 |
| p.Glu82Met | substitution | 82 | Glu | Met | | 0.000 | 0.022 |
| p.Glu82Phe | substitution | 82 | Glu | Phe | | 1.792 | 0.006 |
| p.Glu82Pro | substitution | 82 | Glu | Pro | | 0.000 | 0.033 |
| p.Glu82Ser | substitution | 82 | Glu | Ser | | 0.821 | 0.044 |
| p.Glu82Ter | nonsense | 82 | Glu | Ter | | 0.298 | 0.077 |
| p.Glu82Thr | substitution | 82 | Glu | Thr | | 0.602 | 0.245 |
| p.Glu82Trp | substitution | 82 | Glu | Trp | | 0.413 | 0.013 |
| p.Glu82Tyr | substitution | 82 | Glu | Tyr | | 0.151 | 0.016 |
| p.Glu82Val | substitution | 82 | Glu | Val | | 0.417 | 0.097 |
| p.Glu86= | synonymous | 86 | Glu | NA | | 1.415 | 0.020 |
| p.Glu86Ala | substitution | 86 | Glu | Ala | | 0.874 | 0.020 |
| p.Glu86Arg | substitution | 86 | Glu | Arg | | 1.022 | 0.012 |
| p.Glu86Asn | substitution | 86 | Glu | Asn | | 0.426 | 0.012 |
| p.Glu86Asp | substitution | 86 | Glu | Asp | | 0.969 | 0.008 |
| p.Glu86Gln | substitution | 86 | Glu | Gln | | 0.916 | 0.009 |
| p.Glu86Gly | substitution | 86 | Glu | Gly | | 0.613 | 0.010 |
| p.Glu86His | substitution | 86 | Glu | His | | 1.039 | 0.008 |
| p.Glu86Ile | substitution | 86 | Glu | Ile | | 1.012 | 0.006 |
| p.Glu86Leu | substitution | 86 | Glu | Leu | | 0.806 | 0.032 |
| p.Glu86Lys | substitution | 86 | Glu | Lys | | 1.129 | 0.014 |
| p.Glu86Met | substitution | 86 | Glu | Met | | 0.786 | 0.016 |
| p.Glu86Phe | substitution | 86 | Glu | Phe | | 0.694 | 0.012 |
| p.Glu86Pro | substitution | 86 | Glu | Pro | | 0.000 | 0.057 |
| p.Glu86Ser | substitution | 86 | Glu | Ser | | 0.884 | 0.032 |
| p.Glu86Ter | nonsense | 86 | Glu | Ter | | 0.484 | 0.008 |
| p.Glu86Thr | substitution | 86 | Glu | Thr | | 0.806 | 0.022 |
| p.Glu86Trp | substitution | 86 | Glu | Trp | | 0.187 | 0.018 |
| p.Glu86Tyr | substitution | 86 | Glu | Tyr | | 1.545 | 0.008 |
| p.Glu86Val | substitution | 86 | Glu | Val | | 0.737 | 0.020 |

| | | | | | | | |
|---|---|---|---|---|---|---|---|
| p.Glu89= | synonymous | 89 | Glu | NA | | 0.781 | 0.004 |
| p.Glu89Ala | substitution | 89 | Glu | Ala | | 0.833 | 0.006 |
| p.Glu89Arg | substitution | 89 | Glu | Arg | | 1.042 | 0.006 |
| p.Glu89Asn | substitution | 89 | Glu | Asn | | 1.002 | 0.002 |
| p.Glu89Asp | substitution | 89 | Glu | Asp | | 0.945 | 0.019 |
| p.Glu89Cys | substitution | 89 | Glu | Cys | | 0.666 | 0.003 |
| p.Glu89Gln | substitution | 89 | Glu | Gln | | 1.124 | 0.054 |
| p.Glu89Gly | substitution | 89 | Glu | Gly | | 0.679 | 0.014 |
| p.Glu89His | substitution | 89 | Glu | His | | 0.690 | 0.004 |
| p.Glu89Ile | substitution | 89 | Glu | Ile | | 1.033 | 0.009 |
| p.Glu89Leu | substitution | 89 | Glu | Leu | | 0.827 | 0.007 |
| p.Glu89Lys | substitution | 89 | Glu | Lys | | 0.738 | 0.007 |
| p.Glu89Met | substitution | 89 | Glu | Met | | 0.688 | 0.006 |
| p.Glu89Phe | substitution | 89 | Glu | Phe | | 0.992 | 0.003 |
| p.Glu89Pro | substitution | 89 | Glu | Pro | | 0.000 | 0.027 |
| p.Glu89Ser | substitution | 89 | Glu | Ser | | 1.026 | 0.009 |
| p.Glu89Ter | nonsense | 89 | Glu | Ter | | 0.332 | 0.043 |
| p.Glu89Thr | substitution | 89 | Glu | Thr | | 0.829 | 0.007 |
| p.Glu89Trp | substitution | 89 | Glu | Trp | | 0.631 | 0.004 |
| p.Glu89Tyr | substitution | 89 | Glu | Tyr | | 1.020 | 0.003 |
| p.Glu89Val | substitution | 89 | Glu | Val | | 0.906 | 0.019 |
| p.Gly107= | synonymous | 107 | Gly | NA | | 0.958 | 0.004 |
| p.Gly107Ala | substitution | 107 | Gly | Ala | | 0.754 | 0.027 |
| p.Gly107Arg | substitution | 107 | Gly | Arg | | 0.754 | 0.029 |
| p.Gly107Asn | substitution | 107 | Gly | Asn | | 0.563 | 0.021 |
| p.Gly107Asp | substitution | 107 | Gly | Asp | | 0.947 | 0.062 |
| p.Gly107Cys | substitution | 107 | Gly | Cys | | 0.962 | 0.008 |
| p.Gly107Gln | substitution | 107 | Gly | Gln | | 1.120 | 0.511 |
| p.Gly107Glu | substitution | 107 | Gly | Glu | | 0.808 | 0.021 |
| p.Gly107His | substitution | 107 | Gly | His | | 0.684 | 0.052 |
| p.Gly107Ile | substitution | 107 | Gly | Ile | | 0.000 | 0.323 |
| p.Gly107Leu | substitution | 107 | Gly | Leu | | 0.736 | 0.069 |
| p.Gly107Lys | substitution | 107 | Gly | Lys | | 0.571 | 0.016 |
| p.Gly107Met | substitution | 107 | Gly | Met | | 1.064 | 0.015 |
| p.Gly107Phe | substitution | 107 | Gly | Phe | | 1.048 | 0.008 |
| p.Gly107Pro | substitution | 107 | Gly | Pro | | 0.031 | 0.245 |
| p.Gly107Ser | substitution | 107 | Gly | Ser | | 0.820 | 0.020 |
| p.Gly107Ter | nonsense | 107 | Gly | Ter | | 0.000 | 0.021 |
| p.Gly107Thr | substitution | 107 | Gly | Thr | | 0.000 | 0.165 |
| p.Gly107Trp | substitution | 107 | Gly | Trp | | 1.360 | 0.006 |
| p.Gly107Tyr | substitution | 107 | Gly | Tyr | | 0.000 | 0.061 |
| p.Gly107Val | substitution | 107 | Gly | Val | | 0.726 | 0.012 |
| p.Gly111= | synonymous | 111 | Gly | NA | | 0.800 | 0.016 |

| | | | | | | |
|---|---|---|---|---|---|---|
| p.Gly111Ala | substitution | 111 | Gly | Ala | 0.364 | 0.033 |
| p.Gly111Arg | substitution | 111 | Gly | Arg | 0.015 | 0.040 |
| p.Gly111Asn | substitution | 111 | Gly | Asn | 0.000 | 0.011 |
| p.Gly111Asp | substitution | 111 | Gly | Asp | 0.011 | 0.012 |
| p.Gly111Cys | substitution | 111 | Gly | Cys | 0.000 | 0.008 |
| p.Gly111Gln | substitution | 111 | Gly | Gln | 0.000 | 0.042 |
| p.Gly111Glu | substitution | 111 | Gly | Glu | 0.064 | 0.008 |
| p.Gly111His | substitution | 111 | Gly | His | 0.000 | 0.022 |
| p.Gly111Ile | substitution | 111 | Gly | Ile | 0.000 | 0.054 |
| p.Gly111Leu | substitution | 111 | Gly | Leu | 0.000 | 0.017 |
| p.Gly111Lys | substitution | 111 | Gly | Lys | 0.182 | 0.063 |
| p.Gly111Met | substitution | 111 | Gly | Met | 0.000 | 0.015 |
| p.Gly111Phe | substitution | 111 | Gly | Phe | 0.087 | 0.015 |
| p.Gly111Pro | substitution | 111 | Gly | Pro | 0.000 | 0.032 |
| p.Gly111Ser | substitution | 111 | Gly | Ser | 0.550 | 0.006 |
| p.Gly111Ter | nonsense | 111 | Gly | Ter | 0.119 | 0.024 |
| p.Gly111Thr | substitution | 111 | Gly | Thr | 0.037 | 0.014 |
| p.Gly111Trp | substitution | 111 | Gly | Trp | 0.000 | 0.009 |
| p.Gly111Tyr | substitution | 111 | Gly | Tyr | 0.000 | 0.024 |
| p.Gly111Val | substitution | 111 | Gly | Val | 0.179 | 0.007 |
| p.Gly131= | synonymous | 131 | Gly | NA | 0.922 | 0.038 |
| p.Gly131Ala | substitution | 131 | Gly | Ala | 0.956 | 0.012 |
| p.Gly131Arg | substitution | 131 | Gly | Arg | 0.821 | 0.006 |
| p.Gly131Asn | substitution | 131 | Gly | Asn | 0.750 | 0.008 |
| p.Gly131Asp | substitution | 131 | Gly | Asp | 0.786 | 0.007 |
| p.Gly131Cys | substitution | 131 | Gly | Cys | 0.505 | 0.010 |
| p.Gly131Gln | substitution | 131 | Gly | Gln | 1.092 | 0.009 |
| p.Gly131Glu | substitution | 131 | Gly | Glu | 0.725 | 0.012 |
| p.Gly131His | substitution | 131 | Gly | His | 1.061 | 0.008 |
| p.Gly131Ile | substitution | 131 | Gly | Ile | 0.229 | 0.007 |
| p.Gly131Leu | substitution | 131 | Gly | Leu | 0.634 | 0.007 |
| p.Gly131Lys | substitution | 131 | Gly | Lys | 0.667 | 0.014 |
| p.Gly131Phe | substitution | 131 | Gly | Phe | 0.805 | 0.006 |
| p.Gly131Pro | substitution | 131 | Gly | Pro | 0.416 | 0.021 |
| p.Gly131Ser | substitution | 131 | Gly | Ser | 0.877 | 0.011 |
| p.Gly131Ter | nonsense | 131 | Gly | Ter | 0.000 | 0.011 |
| p.Gly131Thr | substitution | 131 | Gly | Thr | 0.674 | 0.055 |
| p.Gly131Trp | substitution | 131 | Gly | Trp | 0.932 | 0.003 |
| p.Gly131Tyr | substitution | 131 | Gly | Tyr | 0.846 | 0.008 |
| p.Gly131Val | substitution | 131 | Gly | Val | 0.932 | 0.006 |
| p.Gly144= | synonymous | 144 | Gly | NA | 0.984 | 0.006 |
| p.Gly144Ala | substitution | 144 | Gly | Ala | 0.889 | 0.043 |
| p.Gly144Arg | substitution | 144 | Gly | Arg | 0.550 | 0.029 |

| | | | | | | |
|---|---|---|---|---|---|---|
| p.Gly144Asn | substitution | 144 | Gly | Asn | 0.000 | 0.043 |
| p.Gly144Asp | substitution | 144 | Gly | Asp | 0.109 | 0.050 |
| p.Gly144Cys | substitution | 144 | Gly | Cys | 0.640 | 0.017 |
| p.Gly144Gln | substitution | 144 | Gly | Gln | 0.000 | 0.066 |
| p.Gly144Glu | substitution | 144 | Gly | Glu | 0.259 | 0.012 |
| p.Gly144His | substitution | 144 | Gly | His | 0.000 | 0.044 |
| p.Gly144Ile | substitution | 144 | Gly | Ile | 0.000 | 0.027 |
| p.Gly144Leu | substitution | 144 | Gly | Leu | 0.398 | 0.026 |
| p.Gly144Lys | substitution | 144 | Gly | Lys | 0.679 | 0.015 |
| p.Gly144Met | substitution | 144 | Gly | Met | 0.931 | 0.008 |
| p.Gly144Phe | substitution | 144 | Gly | Phe | 0.000 | 0.019 |
| p.Gly144Pro | substitution | 144 | Gly | Pro | 0.000 | 0.103 |
| p.Gly144Ser | substitution | 144 | Gly | Ser | 0.408 | 0.009 |
| p.Gly144Ter | nonsense | 144 | Gly | Ter | 0.402 | 0.096 |
| p.Gly144Thr | substitution | 144 | Gly | Thr | 0.762 | 0.016 |
| p.Gly144Trp | substitution | 144 | Gly | Trp | 0.101 | 0.006 |
| p.Gly144Val | substitution | 144 | Gly | Val | 0.346 | 0.012 |
| p.Gly168= | synonymous | 168 | Gly | NA | 0.902 | 0.024 |
| p.Gly168Ala | substitution | 168 | Gly | Ala | 0.000 | 0.037 |
| p.Gly168Arg | substitution | 168 | Gly | Arg | 0.000 | 0.022 |
| p.Gly168Asn | substitution | 168 | Gly | Asn | 0.137 | 0.016 |
| p.Gly168Asp | substitution | 168 | Gly | Asp | 0.000 | 0.009 |
| p.Gly168Cys | substitution | 168 | Gly | Cys | 0.000 | 0.012 |
| p.Gly168Gln | substitution | 168 | Gly | Gln | 0.000 | 0.026 |
| p.Gly168Glu | substitution | 168 | Gly | Glu | 0.000 | 0.036 |
| p.Gly168His | substitution | 168 | Gly | His | 0.024 | 0.010 |
| p.Gly168Ile | substitution | 168 | Gly | Ile | 0.000 | 0.042 |
| p.Gly168Leu | substitution | 168 | Gly | Leu | 0.110 | 0.010 |
| p.Gly168Lys | substitution | 168 | Gly | Lys | 0.000 | 0.055 |
| p.Gly168Met | substitution | 168 | Gly | Met | 0.000 | 0.024 |
| p.Gly168Phe | substitution | 168 | Gly | Phe | 0.000 | 0.015 |
| p.Gly168Pro | substitution | 168 | Gly | Pro | 0.000 | 0.031 |
| p.Gly168Ser | substitution | 168 | Gly | Ser | 0.000 | 0.016 |
| p.Gly168Ter | nonsense | 168 | Gly | Ter | 0.171 | 0.049 |
| p.Gly168Thr | substitution | 168 | Gly | Thr | 0.000 | 0.052 |
| p.Gly168Trp | substitution | 168 | Gly | Trp | 0.000 | 0.008 |
| p.Gly168Tyr | substitution | 168 | Gly | Tyr | 0.000 | 0.008 |
| p.Gly168Val | substitution | 168 | Gly | Val | 0.000 | 0.055 |
| p.Gly192= | synonymous | 192 | Gly | NA | 1.066 | 0.016 |
| p.Gly192Ala | substitution | 192 | Gly | Ala | 0.377 | 0.023 |
| p.Gly192Arg | substitution | 192 | Gly | Arg | 0.000 | 0.116 |
| p.Gly192Asn | substitution | 192 | Gly | Asn | 0.000 | 0.059 |
| p.Gly192Asp | substitution | 192 | Gly | Asp | 0.117 | 0.044 |

| | | | | | | | |
|---|---|---|---|---|---|---|---|
| p.Gly192Cys | substitution | 192 | Gly | Cys | | 0.885 | 0.247 |
| p.Gly192Gln | substitution | 192 | Gly | Gln | | 0.000 | 0.155 |
| p.Gly192Glu | substitution | 192 | Gly | Glu | | 0.284 | 0.171 |
| p.Gly192His | substitution | 192 | Gly | His | | 0.000 | 0.126 |
| p.Gly192Leu | substitution | 192 | Gly | Leu | | 0.000 | 0.848 |
| p.Gly192Met | substitution | 192 | Gly | Met | | 0.311 | 0.207 |
| p.Gly192Phe | substitution | 192 | Gly | Phe | | 0.000 | 0.168 |
| p.Gly192Ser | substitution | 192 | Gly | Ser | | 0.947 | 0.558 |
| p.Gly192Ter | nonsense | 192 | Gly | Ter | | 0.000 | 0.223 |
| p.Gly192Thr | substitution | 192 | Gly | Thr | | 0.150 | 0.022 |
| p.Gly192Trp | substitution | 192 | Gly | Trp | | 0.000 | 0.037 |
| p.Gly192Val | substitution | 192 | Gly | Val | | 0.507 | 0.088 |
| p.Gly197= | synonymous | 197 | Gly | NA | | 0.518 | 0.003 |
| p.Gly197Ala | substitution | 197 | Gly | Ala | | 0.217 | 0.031 |
| p.Gly197Arg | substitution | 197 | Gly | Arg | | 0.775 | 0.061 |
| p.Gly197Asn | substitution | 197 | Gly | Asn | | 0.176 | 0.014 |
| p.Gly197Asp | substitution | 197 | Gly | Asp | | 0.025 | 0.055 |
| p.Gly197Cys | substitution | 197 | Gly | Cys | | 1.134 | 0.018 |
| p.Gly197Gln | substitution | 197 | Gly | Gln | | 0.000 | 0.046 |
| p.Gly197Glu | substitution | 197 | Gly | Glu | | 0.258 | 0.003 |
| p.Gly197His | substitution | 197 | Gly | His | | 0.000 | 0.006 |
| p.Gly197Ile | substitution | 197 | Gly | Ile | | 0.000 | 0.107 |
| p.Gly197Leu | substitution | 197 | Gly | Leu | | 0.175 | 0.062 |
| p.Gly197Lys | substitution | 197 | Gly | Lys | | 0.071 | 0.034 |
| p.Gly197Met | substitution | 197 | Gly | Met | | 0.000 | 0.029 |
| p.Gly197Phe | substitution | 197 | Gly | Phe | | 0.526 | 0.021 |
| p.Gly197Pro | substitution | 197 | Gly | Pro | | 0.145 | 0.142 |
| p.Gly197Ser | substitution | 197 | Gly | Ser | | 0.983 | 0.100 |
| p.Gly197Ter | nonsense | 197 | Gly | Ter | | 0.000 | 0.014 |
| p.Gly197Thr | substitution | 197 | Gly | Thr | | 0.099 | 0.184 |
| p.Gly197Trp | substitution | 197 | Gly | Trp | | 0.229 | 0.004 |
| p.Gly197Tyr | substitution | 197 | Gly | Tyr | | 0.504 | 0.032 |
| p.Gly197Val | substitution | 197 | Gly | Val | | 0.441 | 0.021 |
| p.Gly203= | synonymous | 203 | Gly | NA | | 1.318 | 0.019 |
| p.Gly203Ala | substitution | 203 | Gly | Ala | | 1.023 | 0.192 |
| p.Gly203Arg | substitution | 203 | Gly | Arg | | 0.168 | 0.174 |
| p.Gly203Cys | substitution | 203 | Gly | Cys | | 1.056 | 0.096 |
| p.Gly203Glu | substitution | 203 | Gly | Glu | | 0.276 | 0.671 |
| p.Gly203His | substitution | 203 | Gly | His | | 0.000 | 0.178 |
| p.Gly203Ile | substitution | 203 | Gly | Ile | | 0.000 | 0.019 |
| p.Gly203Leu | substitution | 203 | Gly | Leu | | 0.000 | 0.053 |
| p.Gly203Met | substitution | 203 | Gly | Met | | 0.000 | 0.030 |
| p.Gly203Phe | substitution | 203 | Gly | Phe | | 0.000 | 0.031 |

| | | | | | | |
|---|---|---|---|---|---|---|
| p.Gly203Pro | substitution | 203 | Gly | Pro | 0.000 | 0.077 |
| p.Gly203Ser | substitution | 203 | Gly | Ser | 1.344 | 0.014 |
| p.Gly203Ter | nonsense | 203 | Gly | Ter | 0.000 | 0.220 |
| p.Gly203Thr | substitution | 203 | Gly | Thr | 0.000 | 0.100 |
| p.Gly203Trp | substitution | 203 | Gly | Trp | 1.142 | 0.002 |
| p.Gly203Val | substitution | 203 | Gly | Val | 1.119 | 0.003 |
| p.Gly216= | synonymous | 216 | Gly | NA | 0.888 | 0.005 |
| p.Gly216Ala | substitution | 216 | Gly | Ala | 0.184 | 0.130 |
| p.Gly216Arg | substitution | 216 | Gly | Arg | 0.000 | 0.068 |
| p.Gly216Asp | substitution | 216 | Gly | Asp | 0.000 | 0.141 |
| p.Gly216Cys | substitution | 216 | Gly | Cys | 1.179 | 0.009 |
| p.Gly216Gln | substitution | 216 | Gly | Gln | 0.000 | 0.229 |
| p.Gly216Glu | substitution | 216 | Gly | Glu | 0.000 | 0.129 |
| p.Gly216His | substitution | 216 | Gly | His | 0.000 | 0.230 |
| p.Gly216Ile | substitution | 216 | Gly | Ile | 0.000 | 0.072 |
| p.Gly216Leu | substitution | 216 | Gly | Leu | 0.000 | 0.053 |
| p.Gly216Lys | substitution | 216 | Gly | Lys | 0.000 | 0.041 |
| p.Gly216Met | substitution | 216 | Gly | Met | 0.040 | 0.055 |
| p.Gly216Phe | substitution | 216 | Gly | Phe | 0.276 | 0.008 |
| p.Gly216Pro | substitution | 216 | Gly | Pro | 0.000 | 0.097 |
| p.Gly216Ser | substitution | 216 | Gly | Ser | 0.942 | 0.124 |
| p.Gly216Ter | nonsense | 216 | Gly | Ter | 0.000 | 0.090 |
| p.Gly216Thr | substitution | 216 | Gly | Thr | 0.000 | 0.025 |
| p.Gly216Trp | substitution | 216 | Gly | Trp | 0.000 | 0.016 |
| p.Gly216Tyr | substitution | 216 | Gly | Tyr | 0.339 | 0.278 |
| p.Gly216Val | substitution | 216 | Gly | Val | 0.000 | 0.075 |
| p.Gly218= | synonymous | 218 | Gly | NA | 0.735 | 0.111 |
| p.Gly218Ala | substitution | 218 | Gly | Ala | 0.361 | 0.056 |
| p.Gly218Arg | substitution | 218 | Gly | Arg | 0.000 | 0.017 |
| p.Gly218Asn | substitution | 218 | Gly | Asn | 0.000 | 0.034 |
| p.Gly218Asp | substitution | 218 | Gly | Asp | 0.000 | 0.016 |
| p.Gly218Cys | substitution | 218 | Gly | Cys | 0.000 | 0.138 |
| p.Gly218Gln | substitution | 218 | Gly | Gln | 0.000 | 0.029 |
| p.Gly218Glu | substitution | 218 | Gly | Glu | 0.122 | 0.046 |
| p.Gly218His | substitution | 218 | Gly | His | 0.000 | 0.018 |
| p.Gly218Ile | substitution | 218 | Gly | Ile | 0.000 | 0.068 |
| p.Gly218Leu | substitution | 218 | Gly | Leu | 0.141 | 0.044 |
| p.Gly218Lys | substitution | 218 | Gly | Lys | 0.000 | 0.065 |
| p.Gly218Met | substitution | 218 | Gly | Met | 0.000 | 0.222 |
| p.Gly218Phe | substitution | 218 | Gly | Phe | 0.000 | 0.173 |
| p.Gly218Pro | substitution | 218 | Gly | Pro | 0.000 | 0.113 |
| p.Gly218Ser | substitution | 218 | Gly | Ser | 0.000 | 0.019 |
| p.Gly218Ter | nonsense | 218 | Gly | Ter | 0.194 | 0.080 |

| | | | | | | | | |
|---|---|---|---|---|---|---|---|---|
| p.Gly218Thr | substitution | 218 | Gly | Thr | | | 0.000 | 0.075 |
| p.Gly218Trp | substitution | 218 | Gly | Trp | | | 0.094 | 0.006 |
| p.Gly218Tyr | substitution | 218 | Gly | Tyr | | | 0.000 | 0.011 |
| p.Gly218Val | substitution | 218 | Gly | Val | | | 0.247 | 0.022 |
| p.Gly221= | synonymous | 221 | Gly | NA | | | 0.508 | 0.056 |
| p.Gly221Ala | substitution | 221 | Gly | Ala | | | 0.541 | 0.074 |
| p.Gly221Arg | substitution | 221 | Gly | Arg | | | 0.000 | 0.010 |
| p.Gly221Asn | substitution | 221 | Gly | Asn | | | 0.000 | 0.075 |
| p.Gly221Asp | substitution | 221 | Gly | Asp | | | 0.000 | 0.026 |
| p.Gly221Cys | substitution | 221 | Gly | Cys | | | 0.840 | 0.152 |
| p.Gly221Glu | substitution | 221 | Gly | Glu | | | 0.000 | 0.016 |
| p.Gly221His | substitution | 221 | Gly | His | | | 0.380 | 0.012 |
| p.Gly221Ile | substitution | 221 | Gly | Ile | | | 1.066 | 0.003 |
| p.Gly221Leu | substitution | 221 | Gly | Leu | | | 0.231 | 0.019 |
| p.Gly221Lys | substitution | 221 | Gly | Lys | | | 0.000 | 0.069 |
| p.Gly221Met | substitution | 221 | Gly | Met | | | 0.000 | 0.019 |
| p.Gly221Phe | substitution | 221 | Gly | Phe | | | 0.072 | 0.058 |
| p.Gly221Pro | substitution | 221 | Gly | Pro | | | 0.202 | 0.042 |
| p.Gly221Ser | substitution | 221 | Gly | Ser | | | 0.564 | 0.065 |
| p.Gly221Ter | nonsense | 221 | Gly | Ter | | | 0.000 | 0.104 |
| p.Gly221Thr | substitution | 221 | Gly | Thr | | | 0.482 | 0.009 |
| p.Gly221Trp | substitution | 221 | Gly | Trp | | | 0.000 | 0.014 |
| p.Gly221Tyr | substitution | 221 | Gly | Tyr | | | 0.243 | 0.023 |
| p.Gly221Val | substitution | 221 | Gly | Val | | | 0.057 | 0.042 |
| p.Gly236= | synonymous | 236 | Gly | NA | | | 0.895 | 0.002 |
| p.Gly236Ala | substitution | 236 | Gly | Ala | | | 1.119 | 0.264 |
| p.Gly236Arg | substitution | 236 | Gly | Arg | | | 1.030 | 0.183 |
| p.Gly236Asn | substitution | 236 | Gly | Asn | | | 0.000 | 0.252 |
| p.Gly236Asp | substitution | 236 | Gly | Asp | | | 1.257 | 0.006 |
| p.Gly236Cys | substitution | 236 | Gly | Cys | | | 1.372 | 0.002 |
| p.Gly236Glu | substitution | 236 | Gly | Glu | | | 0.000 | 0.047 |
| p.Gly236Leu | substitution | 236 | Gly | Leu | | | 0.228 | 0.049 |
| p.Gly236Met | substitution | 236 | Gly | Met | | | 0.000 | 0.206 |
| p.Gly236Phe | substitution | 236 | Gly | Phe | | | 0.000 | 0.010 |
| p.Gly236Ser | substitution | 236 | Gly | Ser | | | 0.950 | 0.008 |
| p.Gly236Ter | nonsense | 236 | Gly | Ter | | | 0.000 | 0.012 |
| p.Gly236Thr | substitution | 236 | Gly | Thr | | | 0.770 | 0.236 |
| p.Gly236Trp | substitution | 236 | Gly | Trp | | | 0.000 | 0.032 |
| p.Gly236Tyr | substitution | 236 | Gly | Tyr | | | 0.000 | 0.067 |
| p.Gly236Val | substitution | 236 | Gly | Val | | | 0.129 | 0.065 |
| p.Gly24= | synonymous | 24 | Gly | NA | | | 0.964 | 0.000 |
| p.Gly24Ala | substitution | 24 | Gly | Ala | | | 0.771 | 0.002 |
| p.Gly24Arg | substitution | 24 | Gly | Arg | | | 0.000 | 0.003 |

| | | | | | | |
|---|---|---|---|---|---|---|
| p.Gly24Asn | substitution | 24 | Gly | Asn | 0.000 | 0.004 |
| p.Gly24Asp | substitution | 24 | Gly | Asp | 0.000 | 0.029 |
| p.Gly24Cys | substitution | 24 | Gly | Cys | 0.706 | 0.000 |
| p.Gly24Gln | substitution | 24 | Gly | Gln | 0.000 | 0.002 |
| p.Gly24Glu | substitution | 24 | Gly | Glu | 0.000 | 0.001 |
| p.Gly24His | substitution | 24 | Gly | His | 0.000 | 0.002 |
| p.Gly24Ile | substitution | 24 | Gly | Ile | 0.158 | 0.002 |
| p.Gly24Leu | substitution | 24 | Gly | Leu | 0.751 | 0.002 |
| p.Gly24Lys | substitution | 24 | Gly | Lys | 0.000 | 0.002 |
| p.Gly24Met | substitution | 24 | Gly | Met | 0.485 | 0.001 |
| p.Gly24Phe | substitution | 24 | Gly | Phe | 0.000 | 0.002 |
| p.Gly24Pro | substitution | 24 | Gly | Pro | 0.000 | 0.020 |
| p.Gly24Ser | substitution | 24 | Gly | Ser | 0.461 | 0.008 |
| p.Gly24Ter | nonsense | 24 | Gly | Ter | 0.000 | 0.001 |
| p.Gly24Thr | substitution | 24 | Gly | Thr | 0.080 | 0.003 |
| p.Gly24Trp | substitution | 24 | Gly | Trp | 0.000 | 0.001 |
| p.Gly24Tyr | substitution | 24 | Gly | Tyr | 0.000 | 0.001 |
| p.Gly24Val | substitution | 24 | Gly | Val | 0.692 | 0.002 |
| p.Gly259= | synonymous | 259 | Gly | NA | 0.925 | 0.056 |
| p.Gly259Ala | substitution | 259 | Gly | Ala | 0.850 | 0.028 |
| p.Gly259Arg | substitution | 259 | Gly | Arg | 0.447 | 0.063 |
| p.Gly259Asn | substitution | 259 | Gly | Asn | 0.000 | 0.137 |
| p.Gly259Asp | substitution | 259 | Gly | Asp | 0.895 | 0.005 |
| p.Gly259Cys | substitution | 259 | Gly | Cys | 0.780 | 0.012 |
| p.Gly259Gln | substitution | 259 | Gly | Gln | 0.000 | 0.027 |
| p.Gly259Glu | substitution | 259 | Gly | Glu | 0.850 | 0.009 |
| p.Gly259His | substitution | 259 | Gly | His | 0.000 | 0.031 |
| p.Gly259Leu | substitution | 259 | Gly | Leu | 0.924 | 0.055 |
| p.Gly259Lys | substitution | 259 | Gly | Lys | 1.236 | 0.003 |
| p.Gly259Phe | substitution | 259 | Gly | Phe | 0.900 | 0.006 |
| p.Gly259Pro | substitution | 259 | Gly | Pro | 0.000 | 0.142 |
| p.Gly259Ser | substitution | 259 | Gly | Ser | 0.864 | 0.042 |
| p.Gly259Ter | nonsense | 259 | Gly | Ter | 0.552 | 0.273 |
| p.Gly259Thr | substitution | 259 | Gly | Thr | 1.059 | 0.054 |
| p.Gly259Trp | substitution | 259 | Gly | Trp | 0.000 | 0.095 |
| p.Gly259Tyr | substitution | 259 | Gly | Tyr | 0.330 | 0.010 |
| p.Gly259Val | substitution | 259 | Gly | Val | 0.356 | 0.089 |
| p.Gly260= | synonymous | 260 | Gly | NA | 0.429 | 0.042 |
| p.Gly260Ala | substitution | 260 | Gly | Ala | 0.210 | 0.045 |
| p.Gly260Arg | substitution | 260 | Gly | Arg | 0.000 | 0.118 |
| p.Gly260Asp | substitution | 260 | Gly | Asp | 1.387 | 0.003 |
| p.Gly260Cys | substitution | 260 | Gly | Cys | 1.258 | 0.005 |
| p.Gly260Ser | substitution | 260 | Gly | Ser | 0.000 | 0.085 |

| | | | | | | | |
|---|---|---|---|---|---|---|---|
| p.Gly260Trp | substitution | 260 | Gly | Trp | | 0.256 | 0.201 |
| p.Gly260Val | substitution | 260 | Gly | Val | | 1.211 | 0.335 |
| p.Gly274= | synonymous | 274 | Gly | NA | | 1.201 | 0.014 |
| p.Gly274Ala | substitution | 274 | Gly | Ala | | 1.563 | 0.264 |
| p.Gly274Arg | substitution | 274 | Gly | Arg | | 0.273 | 0.130 |
| p.Gly274Asn | substitution | 274 | Gly | Asn | | 0.000 | 0.011 |
| p.Gly274Cys | substitution | 274 | Gly | Cys | | 0.191 | 0.047 |
| p.Gly274Glu | substitution | 274 | Gly | Glu | | 1.747 | 0.088 |
| p.Gly274His | substitution | 274 | Gly | His | | 0.278 | 0.015 |
| p.Gly274Ile | substitution | 274 | Gly | Ile | | 0.000 | 0.057 |
| p.Gly274Leu | substitution | 274 | Gly | Leu | | 0.710 | 0.040 |
| p.Gly274Lys | substitution | 274 | Gly | Lys | | 0.652 | 0.065 |
| p.Gly274Ser | substitution | 274 | Gly | Ser | | 0.000 | 0.077 |
| p.Gly274Ter | nonsense | 274 | Gly | Ter | | 0.000 | 0.034 |
| p.Gly274Thr | substitution | 274 | Gly | Thr | | 0.000 | 0.174 |
| p.Gly274Trp | substitution | 274 | Gly | Trp | | 0.784 | 0.009 |
| p.Gly274Tyr | substitution | 274 | Gly | Tyr | | 0.000 | 0.029 |
| p.Gly274Val | substitution | 274 | Gly | Val | | 0.970 | 0.032 |
| p.Gly280= | synonymous | 280 | Gly | NA | | 0.565 | 0.063 |
| p.Gly280Ala | substitution | 280 | Gly | Ala | | 1.174 | 0.004 |
| p.Gly280Arg | substitution | 280 | Gly | Arg | | 0.063 | 0.078 |
| p.Gly280Asn | substitution | 280 | Gly | Asn | | 0.000 | 0.006 |
| p.Gly280Asp | substitution | 280 | Gly | Asp | | 0.139 | 0.022 |
| p.Gly280Cys | substitution | 280 | Gly | Cys | | 0.000 | 0.084 |
| p.Gly280Gln | substitution | 280 | Gly | Gln | | 0.000 | 0.010 |
| p.Gly280Glu | substitution | 280 | Gly | Glu | | 0.000 | 0.098 |
| p.Gly280Ile | substitution | 280 | Gly | Ile | | 0.000 | 0.130 |
| p.Gly280Leu | substitution | 280 | Gly | Leu | | 0.075 | 0.040 |
| p.Gly280Lys | substitution | 280 | Gly | Lys | | 0.715 | 0.011 |
| p.Gly280Met | substitution | 280 | Gly | Met | | 0.000 | 0.008 |
| p.Gly280Phe | substitution | 280 | Gly | Phe | | 0.000 | 0.008 |
| p.Gly280Pro | substitution | 280 | Gly | Pro | | 0.000 | 0.036 |
| p.Gly280Ser | substitution | 280 | Gly | Ser | | 0.184 | 0.041 |
| p.Gly280Ter | nonsense | 280 | Gly | Ter | | 0.090 | 0.072 |
| p.Gly280Thr | substitution | 280 | Gly | Thr | | 0.000 | 0.013 |
| p.Gly280Trp | substitution | 280 | Gly | Trp | | 0.000 | 0.007 |
| p.Gly280Tyr | substitution | 280 | Gly | Tyr | | 0.000 | 0.012 |
| p.Gly280Val | substitution | 280 | Gly | Val | | 0.514 | 0.060 |
| p.Gly281= | synonymous | 281 | Gly | NA | | 1.227 | 0.019 |
| p.Gly281Ala | substitution | 281 | Gly | Ala | | 0.929 | 0.107 |
| p.Gly281Arg | substitution | 281 | Gly | Arg | | 0.679 | 0.132 |
| p.Gly281Asn | substitution | 281 | Gly | Asn | | 0.000 | 0.020 |
| p.Gly281Asp | substitution | 281 | Gly | Asp | | 0.000 | 0.056 |

| | | | | | | |
|---|---|---|---|---|---|---|
| p.Gly281Cys | substitution | 281 | Gly | Cys | 0.795 | 0.092 |
| p.Gly281Glu | substitution | 281 | Gly | Glu | 1.608 | 0.018 |
| p.Gly281His | substitution | 281 | Gly | His | 0.251 | 0.021 |
| p.Gly281Ile | substitution | 281 | Gly | Ile | 0.000 | 0.137 |
| p.Gly281Leu | substitution | 281 | Gly | Leu | 0.625 | 0.087 |
| p.Gly281Lys | substitution | 281 | Gly | Lys | 0.000 | 0.024 |
| p.Gly281Met | substitution | 281 | Gly | Met | 0.000 | 0.016 |
| p.Gly281Pro | substitution | 281 | Gly | Pro | 0.000 | 0.030 |
| p.Gly281Ser | substitution | 281 | Gly | Ser | 1.915 | 0.008 |
| p.Gly281Ter | nonsense | 281 | Gly | Ter | 0.304 | 0.067 |
| p.Gly281Trp | substitution | 281 | Gly | Trp | 0.816 | 0.057 |
| p.Gly281Tyr | substitution | 281 | Gly | Tyr | 0.000 | 0.058 |
| p.Gly281Val | substitution | 281 | Gly | Val | 0.792 | 0.065 |
| p.Gly287= | synonymous | 287 | Gly | NA | 1.030 | 0.005 |
| p.Gly287Ala | substitution | 287 | Gly | Ala | 0.333 | 0.068 |
| p.Gly287Arg | substitution | 287 | Gly | Arg | 0.503 | 0.081 |
| p.Gly287Asn | substitution | 287 | Gly | Asn | 0.029 | 0.015 |
| p.Gly287Asp | substitution | 287 | Gly | Asp | 0.464 | 0.012 |
| p.Gly287Cys | substitution | 287 | Gly | Cys | 0.345 | 0.014 |
| p.Gly287Gln | substitution | 287 | Gly | Gln | 0.000 | 0.032 |
| p.Gly287Glu | substitution | 287 | Gly | Glu | 0.000 | 0.014 |
| p.Gly287His | substitution | 287 | Gly | His | 0.260 | 0.104 |
| p.Gly287Ile | substitution | 287 | Gly | Ile | 0.250 | 0.030 |
| p.Gly287Leu | substitution | 287 | Gly | Leu | 0.348 | 0.067 |
| p.Gly287Lys | substitution | 287 | Gly | Lys | 0.000 | 0.076 |
| p.Gly287Met | substitution | 287 | Gly | Met | 1.094 | 0.005 |
| p.Gly287Phe | substitution | 287 | Gly | Phe | 0.208 | 0.051 |
| p.Gly287Pro | substitution | 287 | Gly | Pro | 0.000 | 0.013 |
| p.Gly287Ser | substitution | 287 | Gly | Ser | 0.574 | 0.061 |
| p.Gly287Ter | nonsense | 287 | Gly | Ter | 0.071 | 0.048 |
| p.Gly287Thr | substitution | 287 | Gly | Thr | 0.000 | 0.023 |
| p.Gly287Trp | substitution | 287 | Gly | Trp | 0.000 | 0.083 |
| p.Gly287Tyr | substitution | 287 | Gly | Tyr | 0.399 | 0.062 |
| p.Gly287Val | substitution | 287 | Gly | Val | 1.290 | 0.016 |
| p.Gly308= | synonymous | 308 | Gly | NA | 0.779 | 0.040 |
| p.Gly308Ala | substitution | 308 | Gly | Ala | 0.905 | 0.008 |
| p.Gly308Arg | substitution | 308 | Gly | Arg | 0.567 | 0.013 |
| p.Gly308Asn | substitution | 308 | Gly | Asn | 1.073 | 0.006 |
| p.Gly308Asp | substitution | 308 | Gly | Asp | 1.049 | 0.029 |
| p.Gly308Cys | substitution | 308 | Gly | Cys | 0.834 | 0.003 |
| p.Gly308Gln | substitution | 308 | Gly | Gln | 1.292 | 0.003 |
| p.Gly308Ile | substitution | 308 | Gly | Ile | 1.185 | 0.010 |
| p.Gly308Leu | substitution | 308 | Gly | Leu | 0.890 | 0.012 |

| | | | | | | |
|---|---|---|---|---|---|---|
| p.Gly308Lys | substitution | 308 | Gly | Lys | 0.000 | 0.048 |
| p.Gly308Met | substitution | 308 | Gly | Met | 0.097 | 0.007 |
| p.Gly308Phe | substitution | 308 | Gly | Phe | 0.000 | 0.184 |
| p.Gly308Pro | substitution | 308 | Gly | Pro | 0.316 | 0.036 |
| p.Gly308Ser | substitution | 308 | Gly | Ser | 0.764 | 0.070 |
| p.Gly308Ter | nonsense | 308 | Gly | Ter | 0.105 | 0.008 |
| p.Gly308Thr | substitution | 308 | Gly | Thr | 0.658 | 0.006 |
| p.Gly308Trp | substitution | 308 | Gly | Trp | 0.895 | 0.001 |
| p.Gly308Tyr | substitution | 308 | Gly | Tyr | 0.000 | 0.104 |
| p.Gly308Val | substitution | 308 | Gly | Val | 1.018 | 0.007 |
| p.Gly317= | synonymous | 317 | Gly | NA | 1.013 | 0.024 |
| p.Gly317Ala | substitution | 317 | Gly | Ala | 0.000 | 0.088 |
| p.Gly317Arg | substitution | 317 | Gly | Arg | 0.079 | 0.043 |
| p.Gly317Asp | substitution | 317 | Gly | Asp | 0.296 | 0.133 |
| p.Gly317Cys | substitution | 317 | Gly | Cys | 1.159 | 0.021 |
| p.Gly317Gln | substitution | 317 | Gly | Gln | 0.000 | 0.018 |
| p.Gly317Glu | substitution | 317 | Gly | Glu | 0.000 | 0.061 |
| p.Gly317His | substitution | 317 | Gly | His | 0.000 | 0.033 |
| p.Gly317Ile | substitution | 317 | Gly | Ile | 0.729 | 0.047 |
| p.Gly317Leu | substitution | 317 | Gly | Leu | 0.016 | 0.023 |
| p.Gly317Lys | substitution | 317 | Gly | Lys | 0.000 | 0.157 |
| p.Gly317Met | substitution | 317 | Gly | Met | 1.257 | 0.001 |
| p.Gly317Phe | substitution | 317 | Gly | Phe | 0.000 | 0.078 |
| p.Gly317Pro | substitution | 317 | Gly | Pro | 0.848 | 0.013 |
| p.Gly317Ser | substitution | 317 | Gly | Ser | 0.173 | 0.060 |
| p.Gly317Ter | nonsense | 317 | Gly | Ter | 0.000 | 0.187 |
| p.Gly317Thr | substitution | 317 | Gly | Thr | 0.531 | 0.053 |
| p.Gly317Trp | substitution | 317 | Gly | Trp | 0.937 | 0.003 |
| p.Gly317Tyr | substitution | 317 | Gly | Tyr | 0.519 | 0.023 |
| p.Gly317Val | substitution | 317 | Gly | Val | 0.457 | 0.056 |
| p.Gly326= | synonymous | 326 | Gly | NA | 0.306 | 0.613 |
| p.Gly326Ala | substitution | 326 | Gly | Ala | 1.265 | 0.004 |
| p.Gly326Arg | substitution | 326 | Gly | Arg | 0.653 | 0.046 |
| p.Gly326Cys | substitution | 326 | Gly | Cys | 1.042 | 0.098 |
| p.Gly326Gln | substitution | 326 | Gly | Gln | 0.000 | 0.087 |
| p.Gly326Glu | substitution | 326 | Gly | Glu | 1.072 | 0.002 |
| p.Gly326Leu | substitution | 326 | Gly | Leu | 0.239 | 0.124 |
| p.Gly326Phe | substitution | 326 | Gly | Phe | 0.000 | 0.077 |
| p.Gly326Pro | substitution | 326 | Gly | Pro | 0.900 | 0.054 |
| p.Gly326Ser | substitution | 326 | Gly | Ser | 0.864 | 0.032 |
| p.Gly326Ter | nonsense | 326 | Gly | Ter | 0.054 | 0.198 |
| p.Gly326Trp | substitution | 326 | Gly | Trp | 0.799 | 0.003 |
| p.Gly326Tyr | substitution | 326 | Gly | Tyr | 0.942 | 0.028 |

| | | | | | | |
|---|---|---|---|---|---|---|
| p.Gly326Val | substitution | 326 | Gly | Val | 0.786 | 0.003 |
| p.Gly335= | synonymous | 335 | Gly | NA | 0.965 | 0.054 |
| p.Gly335Ala | substitution | 335 | Gly | Ala | 0.023 | 0.182 |
| p.Gly335Arg | substitution | 335 | Gly | Arg | 0.000 | 0.130 |
| p.Gly335Asp | substitution | 335 | Gly | Asp | 0.432 | 0.061 |
| p.Gly335Cys | substitution | 335 | Gly | Cys | 0.566 | 0.148 |
| p.Gly335Gln | substitution | 335 | Gly | Gln | 0.306 | 0.250 |
| p.Gly335Glu | substitution | 335 | Gly | Glu | 0.036 | 0.130 |
| p.Gly335Ile | substitution | 335 | Gly | Ile | 0.000 | 0.187 |
| p.Gly335Leu | substitution | 335 | Gly | Leu | 0.000 | 0.031 |
| p.Gly335Lys | substitution | 335 | Gly | Lys | 0.020 | 0.054 |
| p.Gly335Met | substitution | 335 | Gly | Met | 0.192 | 0.042 |
| p.Gly335Phe | substitution | 335 | Gly | Phe | 0.802 | 0.048 |
| p.Gly335Pro | substitution | 335 | Gly | Pro | 0.000 | 0.265 |
| p.Gly335Ser | substitution | 335 | Gly | Ser | 0.000 | 0.065 |
| p.Gly335Ter | nonsense | 335 | Gly | Ter | 0.438 | 0.050 |
| p.Gly335Thr | substitution | 335 | Gly | Thr | 0.113 | 0.014 |
| p.Gly335Trp | substitution | 335 | Gly | Trp | 0.064 | 0.012 |
| p.Gly335Tyr | substitution | 335 | Gly | Tyr | 0.000 | 0.077 |
| p.Gly335Val | substitution | 335 | Gly | Val | 0.000 | 0.089 |
| p.Gly346= | synonymous | 346 | Gly | NA | 0.037 | 0.503 |
| p.Gly346Ala | substitution | 346 | Gly | Ala | 0.361 | 0.057 |
| p.Gly346Arg | substitution | 346 | Gly | Arg | 0.863 | 0.018 |
| p.Gly346Asn | substitution | 346 | Gly | Asn | 0.577 | 0.204 |
| p.Gly346Asp | substitution | 346 | Gly | Asp | 0.054 | 0.123 |
| p.Gly346Cys | substitution | 346 | Gly | Cys | 1.266 | 0.003 |
| p.Gly346Gln | substitution | 346 | Gly | Gln | 0.000 | 0.066 |
| p.Gly346Glu | substitution | 346 | Gly | Glu | 1.334 | 0.063 |
| p.Gly346His | substitution | 346 | Gly | His | 0.271 | 0.023 |
| p.Gly346Leu | substitution | 346 | Gly | Leu | 0.704 | 0.173 |
| p.Gly346Lys | substitution | 346 | Gly | Lys | 0.000 | 0.021 |
| p.Gly346Phe | substitution | 346 | Gly | Phe | 0.511 | 0.005 |
| p.Gly346Pro | substitution | 346 | Gly | Pro | 0.000 | 0.113 |
| p.Gly346Ser | substitution | 346 | Gly | Ser | 0.000 | 0.075 |
| p.Gly346Ter | nonsense | 346 | Gly | Ter | 0.848 | 0.015 |
| p.Gly346Thr | substitution | 346 | Gly | Thr | 0.667 | 0.011 |
| p.Gly346Tyr | substitution | 346 | Gly | Tyr | 0.000 | 0.056 |
| p.Gly346Val | substitution | 346 | Gly | Val | 0.593 | 0.025 |
| p.Gly48= | synonymous | 48 | Gly | NA | 0.974 | 0.009 |
| p.Gly48Ala | substitution | 48 | Gly | Ala | 1.003 | 0.021 |
| p.Gly48Arg | substitution | 48 | Gly | Arg | 0.856 | 0.019 |
| p.Gly48Asn | substitution | 48 | Gly | Asn | 1.325 | 0.010 |
| p.Gly48Asp | substitution | 48 | Gly | Asp | 1.008 | 0.004 |

| | | | | | | | |
|---|---|---|---|---|---|---|---|
| p.Gly48Cys | substitution | 48 | Gly | Cys | | 0.903 | 0.009 |
| p.Gly48Gln | substitution | 48 | Gly | Gln | | 1.158 | 0.007 |
| p.Gly48Glu | substitution | 48 | Gly | Glu | | 0.995 | 0.003 |
| p.Gly48His | substitution | 48 | Gly | His | | 0.164 | 0.084 |
| p.Gly48Ile | substitution | 48 | Gly | Ile | | 0.699 | 0.111 |
| p.Gly48Leu | substitution | 48 | Gly | Leu | | 1.004 | 0.029 |
| p.Gly48Lys | substitution | 48 | Gly | Lys | | 1.101 | 0.009 |
| p.Gly48Met | substitution | 48 | Gly | Met | | 0.492 | 0.007 |
| p.Gly48Phe | substitution | 48 | Gly | Phe | | 0.592 | 0.057 |
| p.Gly48Pro | substitution | 48 | Gly | Pro | | 0.368 | 0.027 |
| p.Gly48Ser | substitution | 48 | Gly | Ser | | 0.914 | 0.021 |
| p.Gly48Ter | nonsense | 48 | Gly | Ter | | 0.000 | 0.016 |
| p.Gly48Thr | substitution | 48 | Gly | Thr | | 0.947 | 0.020 |
| p.Gly48Trp | substitution | 48 | Gly | Trp | | 0.818 | 0.005 |
| p.Gly48Tyr | substitution | 48 | Gly | Tyr | | 0.721 | 0.015 |
| p.Gly48Val | substitution | 48 | Gly | Val | | 0.844 | 0.018 |
| p.Gly60= | synonymous | 60 | Gly | NA | | 1.223 | 0.052 |
| p.Gly60Ala | substitution | 60 | Gly | Ala | | 0.204 | 0.137 |
| p.Gly60Arg | substitution | 60 | Gly | Arg | | 0.231 | 0.056 |
| p.Gly60Asn | substitution | 60 | Gly | Asn | | 0.632 | 0.040 |
| p.Gly60Asp | substitution | 60 | Gly | Asp | | 0.099 | 0.100 |
| p.Gly60Gln | substitution | 60 | Gly | Gln | | 0.594 | 0.055 |
| p.Gly60Glu | substitution | 60 | Gly | Glu | | 0.000 | 0.631 |
| p.Gly60His | substitution | 60 | Gly | His | | 0.040 | 0.488 |
| p.Gly60Ile | substitution | 60 | Gly | Ile | | 0.000 | 0.160 |
| p.Gly60Leu | substitution | 60 | Gly | Leu | | 0.000 | 0.123 |
| p.Gly60Lys | substitution | 60 | Gly | Lys | | 0.000 | 0.061 |
| p.Gly60Met | substitution | 60 | Gly | Met | | 0.000 | 0.115 |
| p.Gly60Phe | substitution | 60 | Gly | Phe | | 0.000 | 0.869 |
| p.Gly60Pro | substitution | 60 | Gly | Pro | | 0.000 | 0.084 |
| p.Gly60Ser | substitution | 60 | Gly | Ser | | 0.000 | 0.077 |
| p.Gly60Ter | nonsense | 60 | Gly | Ter | | 0.000 | 0.884 |
| p.Gly60Thr | substitution | 60 | Gly | Thr | | 0.542 | 0.035 |
| p.Gly60Trp | substitution | 60 | Gly | Trp | | 0.530 | 0.014 |
| p.Gly60Tyr | substitution | 60 | Gly | Tyr | | 0.000 | 0.446 |
| p.Gly60Val | substitution | 60 | Gly | Val | | 0.871 | 0.008 |
| p.Gly72= | synonymous | 72 | Gly | NA | | 0.978 | 0.028 |
| p.Gly72Ala | substitution | 72 | Gly | Ala | | 0.821 | 0.939 |
| p.Gly72Arg | substitution | 72 | Gly | Arg | | 0.610 | 0.050 |
| p.Gly72Asp | substitution | 72 | Gly | Asp | | 0.496 | 0.015 |
| p.Gly72Cys | substitution | 72 | Gly | Cys | | 0.402 | 0.028 |
| p.Gly72Gln | substitution | 72 | Gly | Gln | | 0.152 | 0.023 |
| p.Gly72Glu | substitution | 72 | Gly | Glu | | 1.015 | 0.006 |

| | | | | | | |
|---|---|---|---|---|---|---|
| p.Gly72His | substitution | 72 | Gly | His | 0.709 | 0.009 |
| p.Gly72Ile | substitution | 72 | Gly | Ile | 1.045 | 0.041 |
| p.Gly72Leu | substitution | 72 | Gly | Leu | 0.582 | 0.018 |
| p.Gly72Lys | substitution | 72 | Gly | Lys | 0.088 | 0.030 |
| p.Gly72Met | substitution | 72 | Gly | Met | 0.755 | 0.014 |
| p.Gly72Phe | substitution | 72 | Gly | Phe | 1.145 | 0.005 |
| p.Gly72Pro | substitution | 72 | Gly | Pro | 1.061 | 0.055 |
| p.Gly72Ser | substitution | 72 | Gly | Ser | 1.019 | 0.014 |
| p.Gly72Ter | nonsense | 72 | Gly | Ter | 0.703 | 0.039 |
| p.Gly72Thr | substitution | 72 | Gly | Thr | 0.716 | 0.010 |
| p.Gly72Trp | substitution | 72 | Gly | Trp | 0.380 | 0.013 |
| p.Gly72Tyr | substitution | 72 | Gly | Tyr | 0.837 | 0.003 |
| p.Gly72Val | substitution | 72 | Gly | Val | 0.742 | 0.009 |
| p.His120Ala | substitution | 120 | His | Ala | 0.307 | 0.102 |
| p.His120Arg | substitution | 120 | His | Arg | 1.012 | 0.008 |
| p.His120Asn | substitution | 120 | His | Asn | 0.964 | 0.003 |
| p.His120Asp | substitution | 120 | His | Asp | 0.781 | 0.004 |
| p.His120Cys | substitution | 120 | His | Cys | 0.962 | 0.003 |
| p.His120Gln | substitution | 120 | His | Gln | 0.712 | 0.041 |
| p.His120Glu | substitution | 120 | His | Glu | 1.090 | 0.004 |
| p.His120Gly | substitution | 120 | His | Gly | 0.742 | 0.005 |
| p.His120Ile | substitution | 120 | His | Ile | 0.396 | 0.006 |
| p.His120Leu | substitution | 120 | His | Leu | 1.061 | 0.008 |
| p.His120Met | substitution | 120 | His | Met | 0.989 | 0.005 |
| p.His120Phe | substitution | 120 | His | Phe | 1.235 | 0.003 |
| p.His120Pro | substitution | 120 | His | Pro | 0.121 | 0.006 |
| p.His120Ser | substitution | 120 | His | Ser | 0.966 | 0.005 |
| p.His120Ter | nonsense | 120 | His | Ter | 0.047 | 0.013 |
| p.His120Thr | substitution | 120 | His | Thr | 1.199 | 0.002 |
| p.His120Trp | substitution | 120 | His | Trp | 0.999 | 0.004 |
| p.His120Tyr | substitution | 120 | His | Tyr | 1.056 | 0.006 |
| p.His120Val | substitution | 120 | His | Val | 0.700 | 0.012 |
| p.His126= | synonymous | 126 | His | NA | 0.728 | 0.019 |
| p.His126Ala | substitution | 126 | His | Ala | 0.808 | 0.014 |
| p.His126Arg | substitution | 126 | His | Arg | 0.650 | 0.022 |
| p.His126Asn | substitution | 126 | His | Asn | 0.856 | 0.055 |
| p.His126Asp | substitution | 126 | His | Asp | 0.768 | 0.067 |
| p.His126Cys | substitution | 126 | His | Cys | 1.032 | 0.024 |
| p.His126Gln | substitution | 126 | His | Gln | 1.011 | 0.011 |
| p.His126Glu | substitution | 126 | His | Glu | 1.047 | 0.183 |
| p.His126Gly | substitution | 126 | His | Gly | 0.878 | 0.010 |
| p.His126Ile | substitution | 126 | His | Ile | 1.310 | 0.018 |
| p.His126Leu | substitution | 126 | His | Leu | 0.671 | 0.040 |

| | | | | | | |
|---|---|---|---|---|---|---|
| p.His126Lys | substitution | 126 | His | Lys | 0.999 | 0.033 |
| p.His126Met | substitution | 126 | His | Met | 0.607 | 0.007 |
| p.His126Phe | substitution | 126 | His | Phe | 0.602 | 0.009 |
| p.His126Pro | substitution | 126 | His | Pro | 0.904 | 0.031 |
| p.His126Ser | substitution | 126 | His | Ser | 1.100 | 0.007 |
| p.His126Ter | nonsense | 126 | His | Ter | 0.168 | 0.023 |
| p.His126Thr | substitution | 126 | His | Thr | 0.679 | 0.037 |
| p.His126Trp | substitution | 126 | His | Trp | 0.341 | 0.017 |
| p.His126Tyr | substitution | 126 | His | Tyr | 0.396 | 0.042 |
| p.His126Val | substitution | 126 | His | Val | 0.543 | 0.016 |
| p.His160Ala | substitution | 160 | His | Ala | 0.359 | 0.034 |
| p.His160Arg | substitution | 160 | His | Arg | 1.097 | 0.012 |
| p.His160Asn | substitution | 160 | His | Asn | 0.986 | 0.009 |
| p.His160Asp | substitution | 160 | His | Asp | 0.956 | 0.008 |
| p.His160Gln | substitution | 160 | His | Gln | 1.046 | 0.004 |
| p.His160Glu | substitution | 160 | His | Glu | 0.980 | 0.005 |
| p.His160Gly | substitution | 160 | His | Gly | 0.891 | 0.021 |
| p.His160Ile | substitution | 160 | His | Ile | 0.821 | 0.006 |
| p.His160Leu | substitution | 160 | His | Leu | 0.839 | 0.005 |
| p.His160Lys | substitution | 160 | His | Lys | 1.125 | 0.006 |
| p.His160Met | substitution | 160 | His | Met | 1.186 | 0.010 |
| p.His160Pro | substitution | 160 | His | Pro | 0.834 | 0.010 |
| p.His160Ser | substitution | 160 | His | Ser | 0.826 | 0.033 |
| p.His160Ter | nonsense | 160 | His | Ter | 0.000 | 0.014 |
| p.His160Thr | substitution | 160 | His | Thr | 1.073 | 0.016 |
| p.His160Trp | substitution | 160 | His | Trp | 0.802 | 0.009 |
| p.His160Val | substitution | 160 | His | Val | 0.258 | 0.013 |
| p.His199= | synonymous | 199 | His | NA | 0.852 | 0.001 |
| p.His199Ala | substitution | 199 | His | Ala | 0.804 | 0.048 |
| p.His199Arg | substitution | 199 | His | Arg | 0.900 | 0.029 |
| p.His199Asn | substitution | 199 | His | Asn | 1.119 | 0.002 |
| p.His199Asp | substitution | 199 | His | Asp | 1.444 | 0.037 |
| p.His199Gln | substitution | 199 | His | Gln | 0.387 | 0.012 |
| p.His199Glu | substitution | 199 | His | Glu | 0.787 | 0.004 |
| p.His199Gly | substitution | 199 | His | Gly | 0.766 | 0.021 |
| p.His199Ile | substitution | 199 | His | Ile | 0.970 | 0.013 |
| p.His199Leu | substitution | 199 | His | Leu | 0.755 | 0.019 |
| p.His199Lys | substitution | 199 | His | Lys | 1.083 | 0.007 |
| p.His199Met | substitution | 199 | His | Met | 1.052 | 0.002 |
| p.His199Phe | substitution | 199 | His | Phe | 1.274 | 0.005 |
| p.His199Pro | substitution | 199 | His | Pro | 1.263 | 0.008 |
| p.His199Ser | substitution | 199 | His | Ser | 1.054 | 0.038 |
| p.His199Ter | nonsense | 199 | His | Ter | 0.290 | 0.234 |

| | | | | | | |
|---|---|---|---|---|---|---|
| p.His199Thr | substitution | 199 | His | Thr | 0.417 | 0.017 |
| p.His199Trp | substitution | 199 | His | Trp | 0.807 | 0.008 |
| p.His199Tyr | substitution | 199 | His | Tyr | 1.076 | 0.004 |
| p.His199Val | substitution | 199 | His | Val | 1.126 | 0.004 |
| p.His207= | synonymous | 207 | His | NA | 0.206 | 0.080 |
| p.His207Ala | substitution | 207 | His | Ala | 1.087 | 0.100 |
| p.His207Arg | substitution | 207 | His | Arg | 0.616 | 0.104 |
| p.His207Asn | substitution | 207 | His | Asn | 0.346 | 0.006 |
| p.His207Asp | substitution | 207 | His | Asp | 0.315 | 0.153 |
| p.His207Cys | substitution | 207 | His | Cys | 0.900 | 0.010 |
| p.His207Gln | substitution | 207 | His | Gln | 0.974 | 0.009 |
| p.His207Glu | substitution | 207 | His | Glu | 1.133 | 0.002 |
| p.His207Gly | substitution | 207 | His | Gly | 0.849 | 0.013 |
| p.His207Ile | substitution | 207 | His | Ile | 0.525 | 0.012 |
| p.His207Leu | substitution | 207 | His | Leu | 0.855 | 0.011 |
| p.His207Lys | substitution | 207 | His | Lys | 1.039 | 0.002 |
| p.His207Phe | substitution | 207 | His | Phe | 0.620 | 0.014 |
| p.His207Pro | substitution | 207 | His | Pro | 0.905 | 0.032 |
| p.His207Ser | substitution | 207 | His | Ser | 1.036 | 0.023 |
| p.His207Ter | nonsense | 207 | His | Ter | 0.037 | 0.035 |
| p.His207Thr | substitution | 207 | His | Thr | 1.010 | 0.008 |
| p.His207Trp | substitution | 207 | His | Trp | 1.123 | 0.004 |
| p.His207Tyr | substitution | 207 | His | Tyr | 1.149 | 0.040 |
| p.His207Val | substitution | 207 | His | Val | 0.808 | 0.010 |
| p.His239= | synonymous | 239 | His | NA | 1.236 | 0.001 |
| p.His239Ala | substitution | 239 | His | Ala | 0.980 | 0.006 |
| p.His239Arg | substitution | 239 | His | Arg | 0.000 | 0.033 |
| p.His239Asn | substitution | 239 | His | Asn | 0.515 | 0.028 |
| p.His239Asp | substitution | 239 | His | Asp | 1.087 | 0.030 |
| p.His239Cys | substitution | 239 | His | Cys | 0.794 | 0.006 |
| p.His239Gln | substitution | 239 | His | Gln | 0.586 | 0.014 |
| p.His239Glu | substitution | 239 | His | Glu | 0.250 | 0.008 |
| p.His239Gly | substitution | 239 | His | Gly | 0.000 | 0.033 |
| p.His239Ile | substitution | 239 | His | Ile | 0.815 | 0.019 |
| p.His239Leu | substitution | 239 | His | Leu | 0.350 | 0.017 |
| p.His239Lys | substitution | 239 | His | Lys | 0.366 | 0.006 |
| p.His239Met | substitution | 239 | His | Met | 0.313 | 0.022 |
| p.His239Phe | substitution | 239 | His | Phe | 0.000 | 0.035 |
| p.His239Pro | substitution | 239 | His | Pro | 0.521 | 0.226 |
| p.His239Ser | substitution | 239 | His | Ser | 0.932 | 0.056 |
| p.His239Ter | nonsense | 239 | His | Ter | 0.000 | 0.015 |
| p.His239Thr | substitution | 239 | His | Thr | 1.104 | 0.090 |
| p.His239Trp | substitution | 239 | His | Trp | 0.000 | 0.222 |

| | | | | | | |
|---|---|---|---|---|---|---|
| p.His239Tyr | substitution | 239 | His | Tyr | 0.387 | 0.059 |
| p.His239Val | substitution | 239 | His | Val | 0.084 | 0.032 |
| p.His256= | synonymous | 256 | His | NA | 0.906 | 0.006 |
| p.His256Ala | substitution | 256 | His | Ala | 1.403 | 0.010 |
| p.His256Arg | substitution | 256 | His | Arg | 0.897 | 0.012 |
| p.His256Asn | substitution | 256 | His | Asn | 0.273 | 0.014 |
| p.His256Asp | substitution | 256 | His | Asp | 0.075 | 0.062 |
| p.His256Cys | substitution | 256 | His | Cys | 0.937 | 0.006 |
| p.His256Gln | substitution | 256 | His | Gln | 0.325 | 0.014 |
| p.His256Glu | substitution | 256 | His | Glu | 1.371 | 0.002 |
| p.His256Gly | substitution | 256 | His | Gly | 1.253 | 0.264 |
| p.His256Ile | substitution | 256 | His | Ile | 0.163 | 0.209 |
| p.His256Leu | substitution | 256 | His | Leu | 0.754 | 0.054 |
| p.His256Lys | substitution | 256 | His | Lys | 0.371 | 0.006 |
| p.His256Phe | substitution | 256 | His | Phe | 1.213 | 0.006 |
| p.His256Pro | substitution | 256 | His | Pro | 0.276 | 0.036 |
| p.His256Ser | substitution | 256 | His | Ser | 0.918 | 0.212 |
| p.His256Ter | nonsense | 256 | His | Ter | 0.000 | 0.037 |
| p.His256Thr | substitution | 256 | His | Thr | 1.088 | 0.021 |
| p.His256Trp | substitution | 256 | His | Trp | 1.247 | 0.026 |
| p.His256Tyr | substitution | 256 | His | Tyr | 0.354 | 0.092 |
| p.His256Val | substitution | 256 | His | Val | 0.731 | 0.012 |
| p.His268Ala | substitution | 268 | His | Ala | 1.076 | 0.042 |
| p.His268Arg | substitution | 268 | His | Arg | 1.371 | 0.005 |
| p.His268Asn | substitution | 268 | His | Asn | 1.228 | 0.127 |
| p.His268Asp | substitution | 268 | His | Asp | 0.766 | 0.272 |
| p.His268Cys | substitution | 268 | His | Cys | 0.575 | 0.021 |
| p.His268Gln | substitution | 268 | His | Gln | 1.197 | 0.005 |
| p.His268Glu | substitution | 268 | His | Glu | 1.239 | 0.006 |
| p.His268Gly | substitution | 268 | His | Gly | 0.000 | 0.016 |
| p.His268Ile | substitution | 268 | His | Ile | 0.087 | 0.026 |
| p.His268Leu | substitution | 268 | His | Leu | 0.700 | 0.151 |
| p.His268Lys | substitution | 268 | His | Lys | 0.688 | 0.064 |
| p.His268Met | substitution | 268 | His | Met | 1.297 | 0.008 |
| p.His268Phe | substitution | 268 | His | Phe | 1.032 | 0.005 |
| p.His268Pro | substitution | 268 | His | Pro | 0.164 | 0.049 |
| p.His268Ser | substitution | 268 | His | Ser | 0.163 | 0.032 |
| p.His268Thr | substitution | 268 | His | Thr | 0.000 | 0.019 |
| p.His268Tyr | substitution | 268 | His | Tyr | 1.100 | 0.010 |
| p.His268Val | substitution | 268 | His | Val | 1.190 | 0.255 |
| p.His300Ala | substitution | 300 | His | Ala | 0.000 | 0.129 |
| p.His300Arg | substitution | 300 | His | Arg | 0.278 | 0.037 |
| p.His300Asn | substitution | 300 | His | Asn | 0.965 | 0.004 |

| | | | | | | | |
|---|---|---|---|---|---|---|---|
| p.His300Asp | substitution | 300 | His | Asp | | 1.416 | 0.002 |
| p.His300Cys | substitution | 300 | His | Cys | | 1.034 | 0.006 |
| p.His300Gln | substitution | 300 | His | Gln | | 0.852 | 0.033 |
| p.His300Glu | substitution | 300 | His | Glu | | 0.913 | 0.016 |
| p.His300Gly | substitution | 300 | His | Gly | | 0.998 | 0.007 |
| p.His300Ile | substitution | 300 | His | Ile | | 0.841 | 0.058 |
| p.His300Leu | substitution | 300 | His | Leu | | 0.935 | 0.040 |
| p.His300Lys | substitution | 300 | His | Lys | | 1.146 | 0.003 |
| p.His300Met | substitution | 300 | His | Met | | 0.000 | 0.043 |
| p.His300Phe | substitution | 300 | His | Phe | | 1.177 | 0.008 |
| p.His300Pro | substitution | 300 | His | Pro | | 0.817 | 0.055 |
| p.His300Ser | substitution | 300 | His | Ser | | 0.609 | 0.092 |
| p.His300Ter | nonsense | 300 | His | Ter | | 0.000 | 0.018 |
| p.His300Thr | substitution | 300 | His | Thr | | 1.201 | 0.002 |
| p.His300Val | substitution | 300 | His | Val | | 1.097 | 0.266 |
| p.His305= | synonymous | 305 | His | NA | | 1.175 | 0.686 |
| p.His305Ala | substitution | 305 | His | Ala | | 0.000 | 0.022 |
| p.His305Arg | substitution | 305 | His | Arg | | 0.310 | 0.056 |
| p.His305Asn | substitution | 305 | His | Asn | | 1.026 | 0.003 |
| p.His305Asp | substitution | 305 | His | Asp | | 1.125 | 0.002 |
| p.His305Cys | substitution | 305 | His | Cys | | 1.132 | 0.003 |
| p.His305Gln | substitution | 305 | His | Gln | | 1.025 | 0.045 |
| p.His305Glu | substitution | 305 | His | Glu | | 1.295 | 0.004 |
| p.His305Gly | substitution | 305 | His | Gly | | 1.083 | 0.007 |
| p.His305Ile | substitution | 305 | His | Ile | | 1.104 | 0.002 |
| p.His305Leu | substitution | 305 | His | Leu | | 0.835 | 0.158 |
| p.His305Lys | substitution | 305 | His | Lys | | 0.038 | 0.015 |
| p.His305Met | substitution | 305 | His | Met | | 0.548 | 0.003 |
| p.His305Phe | substitution | 305 | His | Phe | | 0.000 | 0.021 |
| p.His305Pro | substitution | 305 | His | Pro | | 0.000 | 0.015 |
| p.His305Ser | substitution | 305 | His | Ser | | 0.439 | 0.071 |
| p.His305Ter | nonsense | 305 | His | Ter | | 0.003 | 0.058 |
| p.His305Thr | substitution | 305 | His | Thr | | 0.000 | 0.123 |
| p.His305Trp | substitution | 305 | His | Trp | | 0.708 | 0.005 |
| p.His305Tyr | substitution | 305 | His | Tyr | | 1.204 | 0.003 |
| p.His305Val | substitution | 305 | His | Val | | 0.791 | 0.087 |
| p.His83= | synonymous | 83 | His | NA | | 1.992 | 0.045 |
| p.His83Ala | substitution | 83 | His | Ala | | 0.978 | 0.007 |
| p.His83Arg | substitution | 83 | His | Arg | | 1.026 | 0.001 |
| p.His83Asn | substitution | 83 | His | Asn | | 1.004 | 0.003 |
| p.His83Asp | substitution | 83 | His | Asp | | 0.970 | 0.003 |
| p.His83Cys | substitution | 83 | His | Cys | | 1.048 | 0.003 |
| p.His83Gln | substitution | 83 | His | Gln | | 1.053 | 0.006 |

| | | | | | | |
|---|---|---|---|---|---|---|
| p.His83Glu | substitution | 83 | His | Glu | 1.014 | 0.003 |
| p.His83Gly | substitution | 83 | His | Gly | 0.810 | 0.003 |
| p.His83Ile | substitution | 83 | His | Ile | 0.690 | 0.005 |
| p.His83Leu | substitution | 83 | His | Leu | 1.125 | 0.006 |
| p.His83Lys | substitution | 83 | His | Lys | 0.920 | 0.012 |
| p.His83Met | substitution | 83 | His | Met | 0.984 | 0.011 |
| p.His83Phe | substitution | 83 | His | Phe | 0.906 | 0.007 |
| p.His83Pro | substitution | 83 | His | Pro | 0.386 | 0.005 |
| p.His83Ser | substitution | 83 | His | Ser | 0.961 | 0.009 |
| p.His83Ter | nonsense | 83 | His | Ter | 0.040 | 0.016 |
| p.His83Thr | substitution | 83 | His | Thr | 1.004 | 0.029 |
| p.His83Trp | substitution | 83 | His | Trp | 1.076 | 0.004 |
| p.His83Tyr | substitution | 83 | His | Tyr | 1.129 | 0.003 |
| p.His83Val | substitution | 83 | His | Val | 0.847 | 0.002 |
| p.His95= | synonymous | 95 | His | NA | 0.908 | 0.004 |
| p.His95Ala | substitution | 95 | His | Ala | 0.699 | 0.011 |
| p.His95Arg | substitution | 95 | His | Arg | 0.000 | 0.018 |
| p.His95Asn | substitution | 95 | His | Asn | 1.065 | 0.004 |
| p.His95Asp | substitution | 95 | His | Asp | 0.144 | 0.069 |
| p.His95Gln | substitution | 95 | His | Gln | 0.849 | 0.052 |
| p.His95Glu | substitution | 95 | His | Glu | 0.090 | 0.019 |
| p.His95Gly | substitution | 95 | His | Gly | 0.211 | 0.013 |
| p.His95Ile | substitution | 95 | His | Ile | 0.000 | 0.068 |
| p.His95Leu | substitution | 95 | His | Leu | 0.704 | 0.027 |
| p.His95Lys | substitution | 95 | His | Lys | 0.000 | 0.045 |
| p.His95Met | substitution | 95 | His | Met | 0.423 | 0.007 |
| p.His95Phe | substitution | 95 | His | Phe | 0.947 | 0.023 |
| p.His95Pro | substitution | 95 | His | Pro | 0.000 | 0.015 |
| p.His95Ser | substitution | 95 | His | Ser | 0.284 | 0.017 |
| p.His95Ter | nonsense | 95 | His | Ter | 0.178 | 0.008 |
| p.His95Thr | substitution | 95 | His | Thr | 0.525 | 0.271 |
| p.His95Trp | substitution | 95 | His | Trp | 0.000 | 0.013 |
| p.His95Tyr | substitution | 95 | His | Tyr | 0.026 | 0.054 |
| p.His95Val | substitution | 95 | His | Val | 0.739 | 0.027 |
| p.Ile110= | synonymous | 110 | Ile | NA | 0.966 | 0.009 |
| p.Ile110Ala | substitution | 110 | Ile | Ala | 0.000 | 0.051 |
| p.Ile110Arg | substitution | 110 | Ile | Arg | 0.000 | 0.017 |
| p.Ile110Asn | substitution | 110 | Ile | Asn | 0.211 | 0.022 |
| p.Ile110Asp | substitution | 110 | Ile | Asp | 0.175 | 0.043 |
| p.Ile110Cys | substitution | 110 | Ile | Cys | 0.728 | 0.010 |
| p.Ile110Gln | substitution | 110 | Ile | Gln | 0.000 | 0.031 |
| p.Ile110Glu | substitution | 110 | Ile | Glu | 0.124 | 0.027 |
| p.Ile110Gly | substitution | 110 | Ile | Gly | 0.000 | 0.072 |

| | | | | | | | |
|---|---|---|---|---|---|---|---|
| p.Ile110His | substitution | 110 | Ile | His | | 0.159 | 0.015 |
| p.Ile110Leu | substitution | 110 | Ile | Leu | | 0.771 | 0.013 |
| p.Ile110Lys | substitution | 110 | Ile | Lys | | 0.183 | 0.025 |
| p.Ile110Phe | substitution | 110 | Ile | Phe | | 0.309 | 0.170 |
| p.Ile110Pro | substitution | 110 | Ile | Pro | | 0.000 | 0.045 |
| p.Ile110Ser | substitution | 110 | Ile | Ser | | 0.000 | 0.024 |
| p.Ile110Ter | nonsense | 110 | Ile | Ter | | 0.000 | 0.014 |
| p.Ile110Thr | substitution | 110 | Ile | Thr | | 0.311 | 0.017 |
| p.Ile110Trp | substitution | 110 | Ile | Trp | | 0.000 | 0.017 |
| p.Ile110Tyr | substitution | 110 | Ile | Tyr | | 0.095 | 0.033 |
| p.Ile110Val | substitution | 110 | Ile | Val | | 0.824 | 0.065 |
| p.Ile113= | synonymous | 113 | Ile | NA | | 1.089 | 0.010 |
| p.Ile113Ala | substitution | 113 | Ile | Ala | | 0.759 | 0.102 |
| p.Ile113Arg | substitution | 113 | Ile | Arg | | 0.000 | 0.126 |
| p.Ile113Asn | substitution | 113 | Ile | Asn | | 0.000 | 0.051 |
| p.Ile113Asp | substitution | 113 | Ile | Asp | | 0.186 | 0.039 |
| p.Ile113Cys | substitution | 113 | Ile | Cys | | 0.867 | 0.025 |
| p.Ile113Gln | substitution | 113 | Ile | Gln | | 0.235 | 0.110 |
| p.Ile113Glu | substitution | 113 | Ile | Glu | | 0.000 | 0.861 |
| p.Ile113Gly | substitution | 113 | Ile | Gly | | 0.073 | 0.122 |
| p.Ile113His | substitution | 113 | Ile | His | | 0.000 | 0.564 |
| p.Ile113Leu | substitution | 113 | Ile | Leu | | 1.062 | 0.016 |
| p.Ile113Lys | substitution | 113 | Ile | Lys | | 0.000 | 0.022 |
| p.Ile113Met | substitution | 113 | Ile | Met | | 0.000 | 0.053 |
| p.Ile113Phe | substitution | 113 | Ile | Phe | | 0.000 | 0.249 |
| p.Ile113Pro | substitution | 113 | Ile | Pro | | 0.000 | 0.849 |
| p.Ile113Ser | substitution | 113 | Ile | Ser | | 0.075 | 0.050 |
| p.Ile113Ter | nonsense | 113 | Ile | Ter | | 0.000 | 0.109 |
| p.Ile113Thr | substitution | 113 | Ile | Thr | | 0.471 | 0.116 |
| p.Ile113Trp | substitution | 113 | Ile | Trp | | 0.322 | 0.327 |
| p.Ile113Tyr | substitution | 113 | Ile | Tyr | | 0.000 | 0.039 |
| p.Ile113Val | substitution | 113 | Ile | Val | | 0.000 | 0.162 |
| p.Ile166= | synonymous | 166 | Ile | NA | | 1.145 | 1.899 |
| p.Ile166Ala | substitution | 166 | Ile | Ala | | 0.670 | 0.091 |
| p.Ile166Arg | substitution | 166 | Ile | Arg | | 0.211 | 0.026 |
| p.Ile166Asn | substitution | 166 | Ile | Asn | | 0.631 | 0.012 |
| p.Ile166Asp | substitution | 166 | Ile | Asp | | 0.000 | 0.015 |
| p.Ile166Cys | substitution | 166 | Ile | Cys | | 0.989 | 0.006 |
| p.Ile166Gln | substitution | 166 | Ile | Gln | | 0.378 | 0.027 |
| p.Ile166Glu | substitution | 166 | Ile | Glu | | 0.000 | 0.024 |
| p.Ile166Gly | substitution | 166 | Ile | Gly | | 0.210 | 0.009 |
| p.Ile166His | substitution | 166 | Ile | His | | 0.717 | 0.007 |
| p.Ile166Leu | substitution | 166 | Ile | Leu | | 0.860 | 0.017 |

| | | | | | | |
|---|---|---|---|---|---|---|
| p.Ile166Lys | substitution | 166 | Ile | Lys | 0.135 | 0.011 |
| p.Ile166Met | substitution | 166 | Ile | Met | 1.057 | 0.007 |
| p.Ile166Phe | substitution | 166 | Ile | Phe | 1.014 | 0.007 |
| p.Ile166Pro | substitution | 166 | Ile | Pro | 0.000 | 0.017 |
| p.Ile166Ser | substitution | 166 | Ile | Ser | 0.521 | 0.015 |
| p.Ile166Ter | nonsense | 166 | Ile | Ter | 0.050 | 0.015 |
| p.Ile166Thr | substitution | 166 | Ile | Thr | 0.418 | 0.048 |
| p.Ile166Trp | substitution | 166 | Ile | Trp | 0.955 | 0.012 |
| p.Ile166Tyr | substitution | 166 | Ile | Tyr | 0.561 | 0.016 |
| p.Ile166Val | substitution | 166 | Ile | Val | 0.483 | 0.086 |
| p.Ile186= | synonymous | 186 | Ile | NA | 0.903 | 0.002 |
| p.Ile186Ala | substitution | 186 | Ile | Ala | 0.877 | 0.015 |
| p.Ile186Arg | substitution | 186 | Ile | Arg | 0.000 | 0.019 |
| p.Ile186Asn | substitution | 186 | Ile | Asn | 0.421 | 0.002 |
| p.Ile186Asp | substitution | 186 | Ile | Asp | 0.199 | 0.078 |
| p.Ile186Cys | substitution | 186 | Ile | Cys | 0.735 | 0.005 |
| p.Ile186Gln | substitution | 186 | Ile | Gln | 0.943 | 0.002 |
| p.Ile186Glu | substitution | 186 | Ile | Glu | 1.240 | 0.001 |
| p.Ile186Gly | substitution | 186 | Ile | Gly | 0.360 | 0.025 |
| p.Ile186His | substitution | 186 | Ile | His | 0.574 | 0.013 |
| p.Ile186Leu | substitution | 186 | Ile | Leu | 0.688 | 0.013 |
| p.Ile186Lys | substitution | 186 | Ile | Lys | 0.085 | 0.014 |
| p.Ile186Phe | substitution | 186 | Ile | Phe | 0.596 | 0.002 |
| p.Ile186Pro | substitution | 186 | Ile | Pro | 0.000 | 0.012 |
| p.Ile186Ser | substitution | 186 | Ile | Ser | 0.404 | 0.014 |
| p.Ile186Ter | nonsense | 186 | Ile | Ter | 0.103 | 0.024 |
| p.Ile186Thr | substitution | 186 | Ile | Thr | 1.097 | 0.005 |
| p.Ile186Trp | substitution | 186 | Ile | Trp | 0.000 | 0.008 |
| p.Ile186Tyr | substitution | 186 | Ile | Tyr | 0.074 | 0.013 |
| p.Ile186Val | substitution | 186 | Ile | Val | 0.881 | 0.008 |
| p.Ile187= | synonymous | 187 | Ile | NA | 1.333 | 0.009 |
| p.Ile187Ala | substitution | 187 | Ile | Ala | 0.637 | 0.006 |
| p.Ile187Arg | substitution | 187 | Ile | Arg | 0.000 | 0.027 |
| p.Ile187Asn | substitution | 187 | Ile | Asn | 0.316 | 0.025 |
| p.Ile187Asp | substitution | 187 | Ile | Asp | 0.000 | 0.051 |
| p.Ile187Cys | substitution | 187 | Ile | Cys | 0.454 | 0.013 |
| p.Ile187Gln | substitution | 187 | Ile | Gln | 0.145 | 0.010 |
| p.Ile187Glu | substitution | 187 | Ile | Glu | 0.012 | 0.006 |
| p.Ile187Gly | substitution | 187 | Ile | Gly | 0.486 | 0.004 |
| p.Ile187His | substitution | 187 | Ile | His | 0.000 | 0.020 |
| p.Ile187Leu | substitution | 187 | Ile | Leu | 0.727 | 0.011 |
| p.Ile187Lys | substitution | 187 | Ile | Lys | 0.000 | 0.006 |
| p.Ile187Met | substitution | 187 | Ile | Met | 0.645 | 0.001 |

| | | | | | | |
|---|---|---|---|---|---|---|
| p.Ile187Phe | substitution | 187 | Ile | Phe | 0.747 | 0.007 |
| p.Ile187Pro | substitution | 187 | Ile | Pro | 0.000 | 0.008 |
| p.Ile187Ser | substitution | 187 | Ile | Ser | 0.507 | 0.020 |
| p.Ile187Ter | nonsense | 187 | Ile | Ter | 0.004 | 0.005 |
| p.Ile187Thr | substitution | 187 | Ile | Thr | 0.270 | 0.142 |
| p.Ile187Trp | substitution | 187 | Ile | Trp | 0.000 | 0.010 |
| p.Ile187Tyr | substitution | 187 | Ile | Tyr | 0.000 | 0.018 |
| p.Ile187Val | substitution | 187 | Ile | Val | 0.730 | 0.015 |
| p.Ile205= | synonymous | 205 | Ile | NA | 1.307 | 0.007 |
| p.Ile205Ala | substitution | 205 | Ile | Ala | 0.638 | 0.040 |
| p.Ile205Arg | substitution | 205 | Ile | Arg | 0.530 | 0.058 |
| p.Ile205Asn | substitution | 205 | Ile | Asn | 0.783 | 0.026 |
| p.Ile205Asp | substitution | 205 | Ile | Asp | 1.052 | 0.007 |
| p.Ile205Cys | substitution | 205 | Ile | Cys | 1.031 | 0.009 |
| p.Ile205Gln | substitution | 205 | Ile | Gln | 1.221 | 0.012 |
| p.Ile205Glu | substitution | 205 | Ile | Glu | 1.163 | 0.001 |
| p.Ile205Gly | substitution | 205 | Ile | Gly | 0.992 | 0.005 |
| p.Ile205Leu | substitution | 205 | Ile | Leu | 1.114 | 0.020 |
| p.Ile205Lys | substitution | 205 | Ile | Lys | 1.114 | 0.143 |
| p.Ile205Met | substitution | 205 | Ile | Met | 0.767 | 0.002 |
| p.Ile205Phe | substitution | 205 | Ile | Phe | 0.000 | 0.218 |
| p.Ile205Pro | substitution | 205 | Ile | Pro | 0.000 | 0.063 |
| p.Ile205Ser | substitution | 205 | Ile | Ser | 0.451 | 0.077 |
| p.Ile205Ter | nonsense | 205 | Ile | Ter | 0.270 | 0.016 |
| p.Ile205Thr | substitution | 205 | Ile | Thr | 0.848 | 0.053 |
| p.Ile205Trp | substitution | 205 | Ile | Trp | 0.000 | 0.249 |
| p.Ile205Tyr | substitution | 205 | Ile | Tyr | 1.474 | 0.006 |
| p.Ile205Val | substitution | 205 | Ile | Val | 0.954 | 0.005 |
| p.Ile21Ala | substitution | 21 | Ile | Ala | 1.032 | 0.000 |
| p.Ile21Arg | substitution | 21 | Ile | Arg | 0.405 | 0.001 |
| p.Ile21Asn | substitution | 21 | Ile | Asn | 0.017 | 0.005 |
| p.Ile21Asp | substitution | 21 | Ile | Asp | 0.000 | 0.006 |
| p.Ile21Cys | substitution | 21 | Ile | Cys | 0.296 | 0.001 |
| p.Ile21Gln | substitution | 21 | Ile | Gln | 0.000 | 0.005 |
| p.Ile21Glu | substitution | 21 | Ile | Glu | 0.000 | 0.021 |
| p.Ile21Gly | substitution | 21 | Ile | Gly | 0.000 | 0.037 |
| p.Ile21His | substitution | 21 | Ile | His | 0.567 | 0.001 |
| p.Ile21Leu | substitution | 21 | Ile | Leu | 1.261 | 0.002 |
| p.Ile21Lys | substitution | 21 | Ile | Lys | 0.000 | 0.002 |
| p.Ile21Met | substitution | 21 | Ile | Met | 1.016 | 0.000 |
| p.Ile21Phe | substitution | 21 | Ile | Phe | 0.755 | 0.001 |
| p.Ile21Pro | substitution | 21 | Ile | Pro | 0.280 | 0.002 |
| p.Ile21Ser | substitution | 21 | Ile | Ser | 0.000 | 0.077 |

| | | | | | | |
|---|---|---|---|---|---|---|
| p.Ile21Ter | nonsense | 21 | Ile | Ter | 0.000 | 0.059 |
| p.Ile21Thr | substitution | 21 | Ile | Thr | 0.702 | 0.004 |
| p.Ile21Trp | substitution | 21 | Ile | Trp | 0.819 | 0.000 |
| p.Ile21Tyr | substitution | 21 | Ile | Tyr | 0.747 | 0.001 |
| p.Ile21Val | substitution | 21 | Ile | Val | 1.010 | 0.002 |
| p.Ile231= | synonymous | 231 | Ile | NA | 0.751 | 0.009 |
| p.Ile231Ala | substitution | 231 | Ile | Ala | 1.340 | 0.135 |
| p.Ile231Arg | substitution | 231 | Ile | Arg | 0.161 | 0.050 |
| p.Ile231Asn | substitution | 231 | Ile | Asn | 0.516 | 0.074 |
| p.Ile231Asp | substitution | 231 | Ile | Asp | 0.965 | 0.005 |
| p.Ile231Cys | substitution | 231 | Ile | Cys | 0.004 | 0.071 |
| p.Ile231Gln | substitution | 231 | Ile | Gln | 0.000 | 0.039 |
| p.Ile231Glu | substitution | 231 | Ile | Glu | 0.000 | 0.015 |
| p.Ile231Gly | substitution | 231 | Ile | Gly | 0.473 | 0.017 |
| p.Ile231His | substitution | 231 | Ile | His | 0.000 | 0.020 |
| p.Ile231Leu | substitution | 231 | Ile | Leu | 0.144 | 0.021 |
| p.Ile231Lys | substitution | 231 | Ile | Lys | 0.283 | 0.009 |
| p.Ile231Met | substitution | 231 | Ile | Met | 1.043 | 0.014 |
| p.Ile231Phe | substitution | 231 | Ile | Phe | 0.000 | 0.038 |
| p.Ile231Pro | substitution | 231 | Ile | Pro | 0.000 | 0.021 |
| p.Ile231Ser | substitution | 231 | Ile | Ser | 0.787 | 0.026 |
| p.Ile231Ter | nonsense | 231 | Ile | Ter | 0.000 | 0.071 |
| p.Ile231Thr | substitution | 231 | Ile | Thr | 0.598 | 0.035 |
| p.Ile231Trp | substitution | 231 | Ile | Trp | 0.570 | 0.003 |
| p.Ile231Val | substitution | 231 | Ile | Val | 1.057 | 0.002 |
| p.Ile248= | synonymous | 248 | Ile | NA | 0.875 | 0.014 |
| p.Ile248Ala | substitution | 248 | Ile | Ala | 0.032 | 0.045 |
| p.Ile248Arg | substitution | 248 | Ile | Arg | 0.166 | 0.016 |
| p.Ile248Asn | substitution | 248 | Ile | Asn | 0.456 | 0.021 |
| p.Ile248Cys | substitution | 248 | Ile | Cys | 0.000 | 0.035 |
| p.Ile248Glu | substitution | 248 | Ile | Glu | 0.000 | 0.259 |
| p.Ile248Gly | substitution | 248 | Ile | Gly | 0.000 | 0.099 |
| p.Ile248His | substitution | 248 | Ile | His | 0.000 | 0.087 |
| p.Ile248Leu | substitution | 248 | Ile | Leu | 0.864 | 0.173 |
| p.Ile248Lys | substitution | 248 | Ile | Lys | 0.000 | 0.156 |
| p.Ile248Met | substitution | 248 | Ile | Met | 0.263 | 0.242 |
| p.Ile248Phe | substitution | 248 | Ile | Phe | 0.888 | 0.176 |
| p.Ile248Pro | substitution | 248 | Ile | Pro | 0.000 | 0.119 |
| p.Ile248Ser | substitution | 248 | Ile | Ser | 0.628 | 0.212 |
| p.Ile248Ter | nonsense | 248 | Ile | Ter | 0.000 | 0.202 |
| p.Ile248Thr | substitution | 248 | Ile | Thr | 0.272 | 0.097 |
| p.Ile248Trp | substitution | 248 | Ile | Trp | 0.287 | 0.034 |
| p.Ile248Tyr | substitution | 248 | Ile | Tyr | 0.000 | 0.025 |

| | | | | | | |
|---|---|---|---|---|---|---|
| p.Ile248Val | substitution | 248 | Ile | Val | 0.576 | 0.069 |
| p.Ile291= | synonymous | 291 | Ile | NA | 1.103 | 0.005 |
| p.Ile291Ala | substitution | 291 | Ile | Ala | 1.018 | 0.008 |
| p.Ile291Arg | substitution | 291 | Ile | Arg | 1.384 | 0.015 |
| p.Ile291Asn | substitution | 291 | Ile | Asn | 0.744 | 0.009 |
| p.Ile291Gln | substitution | 291 | Ile | Gln | 1.013 | 0.007 |
| p.Ile291Glu | substitution | 291 | Ile | Glu | 1.013 | 0.008 |
| p.Ile291Gly | substitution | 291 | Ile | Gly | 0.590 | 0.020 |
| p.Ile291His | substitution | 291 | Ile | His | 0.798 | 0.005 |
| p.Ile291Leu | substitution | 291 | Ile | Leu | 0.876 | 0.010 |
| p.Ile291Lys | substitution | 291 | Ile | Lys | 1.059 | 0.029 |
| p.Ile291Met | substitution | 291 | Ile | Met | 1.067 | 0.005 |
| p.Ile291Pro | substitution | 291 | Ile | Pro | 0.751 | 0.008 |
| p.Ile291Ser | substitution | 291 | Ile | Ser | 0.556 | 0.026 |
| p.Ile291Ter | nonsense | 291 | Ile | Ter | 0.470 | 0.028 |
| p.Ile291Thr | substitution | 291 | Ile | Thr | 0.707 | 0.004 |
| p.Ile291Trp | substitution | 291 | Ile | Trp | 0.651 | 0.004 |
| p.Ile291Val | substitution | 291 | Ile | Val | 0.539 | 0.014 |
| p.Ile299= | synonymous | 299 | Ile | NA | 0.844 | 0.043 |
| p.Ile299Ala | substitution | 299 | Ile | Ala | 0.901 | 0.085 |
| p.Ile299Arg | substitution | 299 | Ile | Arg | 0.630 | 0.006 |
| p.Ile299Asn | substitution | 299 | Ile | Asn | 0.659 | 0.029 |
| p.Ile299Asp | substitution | 299 | Ile | Asp | 0.942 | 0.010 |
| p.Ile299Cys | substitution | 299 | Ile | Cys | 1.327 | 0.004 |
| p.Ile299Glu | substitution | 299 | Ile | Glu | 0.874 | 0.005 |
| p.Ile299Gly | substitution | 299 | Ile | Gly | 0.000 | 0.034 |
| p.Ile299His | substitution | 299 | Ile | His | 0.769 | 0.004 |
| p.Ile299Leu | substitution | 299 | Ile | Leu | 0.795 | 0.105 |
| p.Ile299Lys | substitution | 299 | Ile | Lys | 0.343 | 0.004 |
| p.Ile299Met | substitution | 299 | Ile | Met | 1.280 | 0.005 |
| p.Ile299Phe | substitution | 299 | Ile | Phe | 1.130 | 0.014 |
| p.Ile299Ser | substitution | 299 | Ile | Ser | 0.479 | 0.108 |
| p.Ile299Ter | nonsense | 299 | Ile | Ter | 0.000 | 0.010 |
| p.Ile299Thr | substitution | 299 | Ile | Thr | 1.121 | 0.008 |
| p.Ile299Trp | substitution | 299 | Ile | Trp | 0.223 | 0.029 |
| p.Ile299Tyr | substitution | 299 | Ile | Tyr | 0.966 | 0.011 |
| p.Ile299Val | substitution | 299 | Ile | Val | 0.961 | 0.112 |
| p.Ile318= | synonymous | 318 | Ile | NA | 1.028 | 0.048 |
| p.Ile318Ala | substitution | 318 | Ile | Ala | 0.000 | 0.174 |
| p.Ile318Arg | substitution | 318 | Ile | Arg | 0.281 | 0.106 |
| p.Ile318Asn | substitution | 318 | Ile | Asn | 0.444 | 0.103 |
| p.Ile318Asp | substitution | 318 | Ile | Asp | 0.131 | 0.027 |
| p.Ile318Gln | substitution | 318 | Ile | Gln | 0.000 | 0.139 |

| | | | | | | |
|---|---|---|---|---|---|---|
| p.Ile318Glu | substitution | 318 | Ile | Glu | 0.246 | 0.163 |
| p.Ile318Gly | substitution | 318 | Ile | Gly | 0.577 | 0.014 |
| p.Ile318Leu | substitution | 318 | Ile | Leu | 0.354 | 0.026 |
| p.Ile318Lys | substitution | 318 | Ile | Lys | 1.430 | 0.003 |
| p.Ile318Met | substitution | 318 | Ile | Met | 0.658 | 0.009 |
| p.Ile318Phe | substitution | 318 | Ile | Phe | 1.218 | 0.013 |
| p.Ile318Pro | substitution | 318 | Ile | Pro | 0.000 | 0.133 |
| p.Ile318Ser | substitution | 318 | Ile | Ser | 0.843 | 0.076 |
| p.Ile318Ter | nonsense | 318 | Ile | Ter | 0.702 | 0.051 |
| p.Ile318Thr | substitution | 318 | Ile | Thr | 0.514 | 0.048 |
| p.Ile318Trp | substitution | 318 | Ile | Trp | 0.000 | 0.011 |
| p.Ile318Tyr | substitution | 318 | Ile | Tyr | 0.835 | 0.004 |
| p.Ile318Val | substitution | 318 | Ile | Val | 1.196 | 0.018 |
| p.Ile323= | synonymous | 323 | Ile | NA | 1.203 | 0.185 |
| p.Ile323Ala | substitution | 323 | Ile | Ala | 0.076 | 0.009 |
| p.Ile323Arg | substitution | 323 | Ile | Arg | 0.564 | 0.109 |
| p.Ile323Asn | substitution | 323 | Ile | Asn | 0.728 | 0.002 |
| p.Ile323Asp | substitution | 323 | Ile | Asp | 0.548 | 0.003 |
| p.Ile323Cys | substitution | 323 | Ile | Cys | 0.883 | 0.002 |
| p.Ile323Gln | substitution | 323 | Ile | Gln | 1.335 | 0.004 |
| p.Ile323Glu | substitution | 323 | Ile | Glu | 0.466 | 0.006 |
| p.Ile323Gly | substitution | 323 | Ile | Gly | 0.129 | 0.016 |
| p.Ile323His | substitution | 323 | Ile | His | 0.581 | 0.002 |
| p.Ile323Leu | substitution | 323 | Ile | Leu | 1.186 | 0.019 |
| p.Ile323Lys | substitution | 323 | Ile | Lys | 0.016 | 0.066 |
| p.Ile323Met | substitution | 323 | Ile | Met | 0.265 | 0.004 |
| p.Ile323Phe | substitution | 323 | Ile | Phe | 0.810 | 0.002 |
| p.Ile323Pro | substitution | 323 | Ile | Pro | 0.000 | 0.045 |
| p.Ile323Ser | substitution | 323 | Ile | Ser | 0.812 | 0.025 |
| p.Ile323Ter | nonsense | 323 | Ile | Ter | 0.229 | 0.030 |
| p.Ile323Thr | substitution | 323 | Ile | Thr | 0.656 | 0.004 |
| p.Ile323Tyr | substitution | 323 | Ile | Tyr | 0.820 | 0.002 |
| p.Ile323Val | substitution | 323 | Ile | Val | 0.902 | 0.002 |
| p.Ile336= | synonymous | 336 | Ile | NA | 1.224 | 0.003 |
| p.Ile336Ala | substitution | 336 | Ile | Ala | 0.592 | 0.004 |
| p.Ile336Arg | substitution | 336 | Ile | Arg | 1.351 | 0.005 |
| p.Ile336Asp | substitution | 336 | Ile | Asp | 0.000 | 0.126 |
| p.Ile336Cys | substitution | 336 | Ile | Cys | 0.000 | 0.037 |
| p.Ile336Gly | substitution | 336 | Ile | Gly | 0.000 | 0.070 |
| p.Ile336Leu | substitution | 336 | Ile | Leu | 0.271 | 0.014 |
| p.Ile336Lys | substitution | 336 | Ile | Lys | 0.000 | 0.038 |
| p.Ile336Met | substitution | 336 | Ile | Met | 0.230 | 0.017 |
| p.Ile336Phe | substitution | 336 | Ile | Phe | 0.818 | 0.005 |

| | | | | | | |
|---|---|---|---|---|---|---|
| p.Ile336Ser | substitution | 336 | Ile | Ser | 1.318 | 0.015 |
| p.Ile336Ter | nonsense | 336 | Ile | Ter | 0.000 | 0.156 |
| p.Ile336Thr | substitution | 336 | Ile | Thr | 0.000 | 0.044 |
| p.Ile336Trp | substitution | 336 | Ile | Trp | 1.031 | 0.009 |
| p.Ile336Tyr | substitution | 336 | Ile | Tyr | 0.000 | 0.101 |
| p.Ile336Val | substitution | 336 | Ile | Val | 0.362 | 0.039 |
| p.Ile33= | synonymous | 33 | Ile | NA | 0.376 | 0.003 |
| p.Ile33Ala | substitution | 33 | Ile | Ala | 1.185 | 0.001 |
| p.Ile33Arg | substitution | 33 | Ile | Arg | 0.824 | 0.004 |
| p.Ile33Asn | substitution | 33 | Ile | Asn | 0.866 | 0.006 |
| p.Ile33Asp | substitution | 33 | Ile | Asp | 0.893 | 0.002 |
| p.Ile33Gln | substitution | 33 | Ile | Gln | 0.664 | 0.002 |
| p.Ile33Glu | substitution | 33 | Ile | Glu | 0.017 | 0.008 |
| p.Ile33Gly | substitution | 33 | Ile | Gly | 0.582 | 0.023 |
| p.Ile33His | substitution | 33 | Ile | His | 0.547 | 0.004 |
| p.Ile33Leu | substitution | 33 | Ile | Leu | 0.866 | 0.011 |
| p.Ile33Lys | substitution | 33 | Ile | Lys | 0.000 | 0.019 |
| p.Ile33Met | substitution | 33 | Ile | Met | 0.639 | 0.004 |
| p.Ile33Phe | substitution | 33 | Ile | Phe | 0.468 | 0.001 |
| p.Ile33Pro | substitution | 33 | Ile | Pro | 0.000 | 0.015 |
| p.Ile33Ser | substitution | 33 | Ile | Ser | 0.875 | 0.075 |
| p.Ile33Ter | nonsense | 33 | Ile | Ter | 0.000 | 0.012 |
| p.Ile33Thr | substitution | 33 | Ile | Thr | 0.956 | 0.018 |
| p.Ile33Trp | substitution | 33 | Ile | Trp | 0.944 | 0.006 |
| p.Ile33Tyr | substitution | 33 | Ile | Tyr | 1.331 | 0.001 |
| p.Ile33Val | substitution | 33 | Ile | Val | 0.678 | 0.001 |
| p.Ile350= | synonymous | 350 | Ile | NA | 0.996 | 0.008 |
| p.Ile350Ala | substitution | 350 | Ile | Ala | 0.242 | 0.053 |
| p.Ile350Arg | substitution | 350 | Ile | Arg | 0.512 | 0.224 |
| p.Ile350Asn | substitution | 350 | Ile | Asn | 0.393 | 0.160 |
| p.Ile350Asp | substitution | 350 | Ile | Asp | 0.188 | 0.062 |
| p.Ile350Cys | substitution | 350 | Ile | Cys | 0.388 | 0.027 |
| p.Ile350Gln | substitution | 350 | Ile | Gln | 0.180 | 0.063 |
| p.Ile350Glu | substitution | 350 | Ile | Glu | 0.000 | 0.199 |
| p.Ile350Gly | substitution | 350 | Ile | Gly | 0.000 | 0.192 |
| p.Ile350His | substitution | 350 | Ile | His | 1.469 | 0.004 |
| p.Ile350Leu | substitution | 350 | Ile | Leu | 0.130 | 0.052 |
| p.Ile350Lys | substitution | 350 | Ile | Lys | 0.000 | 0.115 |
| p.Ile350Met | substitution | 350 | Ile | Met | 0.711 | 0.251 |
| p.Ile350Phe | substitution | 350 | Ile | Phe | 1.199 | 0.040 |
| p.Ile350Pro | substitution | 350 | Ile | Pro | 0.000 | 0.082 |
| p.Ile350Ser | substitution | 350 | Ile | Ser | 0.142 | 0.027 |
| p.Ile350Ter | nonsense | 350 | Ile | Ter | 0.000 | 0.022 |

| | | | | | | | |
|---|---|---|---|---|---|---|---|
| p.Ile350Thr | substitution | 350 | Ile | Thr | | 0.000 | 0.031 |
| p.Ile350Trp | substitution | 350 | Ile | Trp | | 1.342 | 0.007 |
| p.Ile350Tyr | substitution | 350 | Ile | Tyr | | 1.564 | 0.005 |
| p.Ile350Val | substitution | 350 | Ile | Val | | 1.058 | 0.065 |
| p.Ile53= | synonymous | 53 | Ile | NA | | 0.920 | 0.011 |
| p.Ile53Ala | substitution | 53 | Ile | Ala | | 0.460 | 0.022 |
| p.Ile53Arg | substitution | 53 | Ile | Arg | | 0.666 | 0.077 |
| p.Ile53Asn | substitution | 53 | Ile | Asn | | 0.451 | 0.046 |
| p.Ile53Asp | substitution | 53 | Ile | Asp | | 0.770 | 0.032 |
| p.Ile53Cys | substitution | 53 | Ile | Cys | | 0.598 | 0.011 |
| p.Ile53Gln | substitution | 53 | Ile | Gln | | 0.609 | 0.476 |
| p.Ile53Glu | substitution | 53 | Ile | Glu | | 0.578 | 0.025 |
| p.Ile53Gly | substitution | 53 | Ile | Gly | | 0.254 | 0.010 |
| p.Ile53His | substitution | 53 | Ile | His | | 0.000 | 0.097 |
| p.Ile53Leu | substitution | 53 | Ile | Leu | | 0.928 | 0.018 |
| p.Ile53Lys | substitution | 53 | Ile | Lys | | 0.863 | 0.007 |
| p.Ile53Met | substitution | 53 | Ile | Met | | 0.688 | 0.005 |
| p.Ile53Phe | substitution | 53 | Ile | Phe | | 1.025 | 0.029 |
| p.Ile53Pro | substitution | 53 | Ile | Pro | | 1.207 | 0.069 |
| p.Ile53Ser | substitution | 53 | Ile | Ser | | 0.642 | 0.011 |
| p.Ile53Ter | nonsense | 53 | Ile | Ter | | 0.456 | 0.011 |
| p.Ile53Thr | substitution | 53 | Ile | Thr | | 0.573 | 0.011 |
| p.Ile53Trp | substitution | 53 | Ile | Trp | | 0.353 | 0.010 |
| p.Ile53Tyr | substitution | 53 | Ile | Tyr | | 1.056 | 0.005 |
| p.Ile53Val | substitution | 53 | Ile | Val | | 0.899 | 0.060 |
| p.Ile54= | synonymous | 54 | Ile | NA | | 1.039 | 1.346 |
| p.Ile54Ala | substitution | 54 | Ile | Ala | | 0.537 | 0.024 |
| p.Ile54Arg | substitution | 54 | Ile | Arg | | 0.707 | 0.005 |
| p.Ile54Asn | substitution | 54 | Ile | Asn | | 0.738 | 0.009 |
| p.Ile54Asp | substitution | 54 | Ile | Asp | | 0.429 | 0.009 |
| p.Ile54Cys | substitution | 54 | Ile | Cys | | 0.060 | 0.022 |
| p.Ile54Gln | substitution | 54 | Ile | Gln | | 0.031 | 0.014 |
| p.Ile54Glu | substitution | 54 | Ile | Glu | | 1.124 | 0.004 |
| p.Ile54Gly | substitution | 54 | Ile | Gly | | 0.383 | 0.009 |
| p.Ile54His | substitution | 54 | Ile | His | | 0.300 | 0.009 |
| p.Ile54Leu | substitution | 54 | Ile | Leu | | 0.803 | 0.007 |
| p.Ile54Lys | substitution | 54 | Ile | Lys | | 0.909 | 0.007 |
| p.Ile54Met | substitution | 54 | Ile | Met | | 0.710 | 0.006 |
| p.Ile54Phe | substitution | 54 | Ile | Phe | | 0.979 | 0.004 |
| p.Ile54Pro | substitution | 54 | Ile | Pro | | 0.000 | 0.012 |
| p.Ile54Ser | substitution | 54 | Ile | Ser | | 0.596 | 0.006 |
| p.Ile54Ter | nonsense | 54 | Ile | Ter | | 0.000 | 0.017 |
| p.Ile54Thr | substitution | 54 | Ile | Thr | | 0.719 | 0.010 |

| | | | | | | | |
|---|---|---|---|---|---|---|---|
| p.Ile54Trp | substitution | 54 | Ile | Trp | | 0.208 | 0.009 |
| p.Ile54Tyr | substitution | 54 | Ile | Tyr | | 0.904 | 0.006 |
| p.Ile54Val | substitution | 54 | Ile | Val | | 0.786 | 0.006 |
| p.Ile63Ala | substitution | 63 | Ile | Ala | | 1.053 | 0.035 |
| p.Ile63Arg | substitution | 63 | Ile | Arg | | 0.781 | 0.337 |
| p.Ile63Asn | substitution | 63 | Ile | Asn | | 1.084 | 0.018 |
| p.Ile63Asp | substitution | 63 | Ile | Asp | | 0.000 | 0.088 |
| p.Ile63Cys | substitution | 63 | Ile | Cys | | 0.849 | 0.048 |
| p.Ile63Gln | substitution | 63 | Ile | Gln | | 0.397 | 0.046 |
| p.Ile63Glu | substitution | 63 | Ile | Glu | | 0.000 | 0.175 |
| p.Ile63Gly | substitution | 63 | Ile | Gly | | 0.909 | 0.019 |
| p.Ile63His | substitution | 63 | Ile | His | | 0.401 | 0.096 |
| p.Ile63Leu | substitution | 63 | Ile | Leu | | 1.029 | 0.027 |
| p.Ile63Lys | substitution | 63 | Ile | Lys | | 0.194 | 0.479 |
| p.Ile63Met | substitution | 63 | Ile | Met | | 1.004 | 0.010 |
| p.Ile63Phe | substitution | 63 | Ile | Phe | | 1.004 | 0.014 |
| p.Ile63Pro | substitution | 63 | Ile | Pro | | 0.936 | 0.241 |
| p.Ile63Ser | substitution | 63 | Ile | Ser | | 0.785 | 0.116 |
| p.Ile63Ter | nonsense | 63 | Ile | Ter | | 0.000 | 0.928 |
| p.Ile63Thr | substitution | 63 | Ile | Thr | | 0.000 | 0.275 |
| p.Ile63Trp | substitution | 63 | Ile | Trp | | 0.685 | 0.029 |
| p.Ile63Tyr | substitution | 63 | Ile | Tyr | | 0.372 | 0.053 |
| p.Ile63Val | substitution | 63 | Ile | Val | | 0.633 | 0.137 |
| p.Ile71= | synonymous | 71 | Ile | NA | | 1.315 | 0.159 |
| p.Ile71Ala | substitution | 71 | Ile | Ala | | 0.000 | 0.044 |
| p.Ile71Arg | substitution | 71 | Ile | Arg | | 0.289 | 0.055 |
| p.Ile71Asn | substitution | 71 | Ile | Asn | | 0.821 | 0.009 |
| p.Ile71Asp | substitution | 71 | Ile | Asp | | 0.547 | 0.023 |
| p.Ile71Cys | substitution | 71 | Ile | Cys | | 0.769 | 0.009 |
| p.Ile71Gln | substitution | 71 | Ile | Gln | | 1.561 | 0.013 |
| p.Ile71Gly | substitution | 71 | Ile | Gly | | 0.720 | 0.009 |
| p.Ile71His | substitution | 71 | Ile | His | | 0.657 | 0.012 |
| p.Ile71Leu | substitution | 71 | Ile | Leu | | 0.979 | 0.011 |
| p.Ile71Lys | substitution | 71 | Ile | Lys | | 0.929 | 0.015 |
| p.Ile71Met | substitution | 71 | Ile | Met | | 1.248 | 0.007 |
| p.Ile71Phe | substitution | 71 | Ile | Phe | | 0.861 | 0.021 |
| p.Ile71Pro | substitution | 71 | Ile | Pro | | 0.671 | 0.015 |
| p.Ile71Ser | substitution | 71 | Ile | Ser | | 0.745 | 0.009 |
| p.Ile71Ter | nonsense | 71 | Ile | Ter | | 0.000 | 0.138 |
| p.Ile71Thr | substitution | 71 | Ile | Thr | | 0.705 | 0.027 |
| p.Ile71Trp | substitution | 71 | Ile | Trp | | 0.505 | 0.014 |
| p.Ile71Tyr | substitution | 71 | Ile | Tyr | | 0.850 | 0.008 |
| p.Ile71Val | substitution | 71 | Ile | Val | | 0.891 | 0.079 |

| | | | | | | |
|---|---|---|---|---|---|---|
| p.Leu100= | synonymous | 100 | Leu | NA | 0.974 | 0.008 |
| p.Leu100Ala | substitution | 100 | Leu | Ala | 0.103 | 0.019 |
| p.Leu100Arg | substitution | 100 | Leu | Arg | 0.000 | 0.055 |
| p.Leu100Asn | substitution | 100 | Leu | Asn | 0.000 | 0.043 |
| p.Leu100Asp | substitution | 100 | Leu | Asp | 0.000 | 0.057 |
| p.Leu100Cys | substitution | 100 | Leu | Cys | 1.421 | 0.006 |
| p.Leu100Gln | substitution | 100 | Leu | Gln | 0.736 | 0.118 |
| p.Leu100Glu | substitution | 100 | Leu | Glu | 1.017 | 0.014 |
| p.Leu100Gly | substitution | 100 | Leu | Gly | 0.000 | 0.030 |
| p.Leu100His | substitution | 100 | Leu | His | 0.000 | 0.049 |
| p.Leu100Ile | substitution | 100 | Leu | Ile | 1.144 | 0.027 |
| p.Leu100Lys | substitution | 100 | Leu | Lys | 0.000 | 0.062 |
| p.Leu100Met | substitution | 100 | Leu | Met | 0.763 | 0.004 |
| p.Leu100Phe | substitution | 100 | Leu | Phe | 0.000 | 0.032 |
| p.Leu100Pro | substitution | 100 | Leu | Pro | 0.303 | 0.547 |
| p.Leu100Ser | substitution | 100 | Leu | Ser | 0.459 | 0.122 |
| p.Leu100Ter | nonsense | 100 | Leu | Ter | 0.000 | 0.095 |
| p.Leu100Thr | substitution | 100 | Leu | Thr | 0.487 | 0.093 |
| p.Leu100Trp | substitution | 100 | Leu | Trp | 0.304 | 0.020 |
| p.Leu100Tyr | substitution | 100 | Leu | Tyr | 0.970 | 0.024 |
| p.Leu100Val | substitution | 100 | Leu | Val | 0.638 | 0.088 |
| p.Leu104= | synonymous | 104 | Leu | NA | 0.700 | 0.006 |
| p.Leu104Ala | substitution | 104 | Leu | Ala | 0.685 | 0.082 |
| p.Leu104Arg | substitution | 104 | Leu | Arg | 0.536 | 0.037 |
| p.Leu104Asn | substitution | 104 | Leu | Asn | 0.023 | 0.021 |
| p.Leu104Asp | substitution | 104 | Leu | Asp | 0.324 | 0.012 |
| p.Leu104Cys | substitution | 104 | Leu | Cys | 0.706 | 0.006 |
| p.Leu104Gln | substitution | 104 | Leu | Gln | 0.635 | 0.006 |
| p.Leu104Glu | substitution | 104 | Leu | Glu | 0.281 | 0.012 |
| p.Leu104Gly | substitution | 104 | Leu | Gly | 0.649 | 0.018 |
| p.Leu104His | substitution | 104 | Leu | His | 0.808 | 0.008 |
| p.Leu104Ile | substitution | 104 | Leu | Ile | 1.006 | 0.002 |
| p.Leu104Lys | substitution | 104 | Leu | Lys | 0.000 | 0.087 |
| p.Leu104Met | substitution | 104 | Leu | Met | 1.059 | 0.006 |
| p.Leu104Phe | substitution | 104 | Leu | Phe | 0.782 | 0.003 |
| p.Leu104Pro | substitution | 104 | Leu | Pro | 0.664 | 0.014 |
| p.Leu104Ser | substitution | 104 | Leu | Ser | 0.781 | 0.014 |
| p.Leu104Ter | nonsense | 104 | Leu | Ter | 0.000 | 0.014 |
| p.Leu104Thr | substitution | 104 | Leu | Thr | 0.407 | 0.011 |
| p.Leu104Tyr | substitution | 104 | Leu | Tyr | 0.754 | 0.007 |
| p.Leu104Val | substitution | 104 | Leu | Val | 0.765 | 0.008 |
| p.Leu134= | synonymous | 134 | Leu | NA | 0.986 | 0.011 |
| p.Leu134Ala | substitution | 134 | Leu | Ala | 0.601 | 0.054 |

| | | | | | | |
|---|---|---|---|---|---|---|
| p.Leu134Arg | substitution | 134 | Leu | Arg | 0.042 | 0.016 |
| p.Leu134Asn | substitution | 134 | Leu | Asn | 0.000 | 0.023 |
| p.Leu134Asp | substitution | 134 | Leu | Asp | 0.000 | 0.022 |
| p.Leu134Cys | substitution | 134 | Leu | Cys | 0.603 | 0.007 |
| p.Leu134Gln | substitution | 134 | Leu | Gln | 0.000 | 0.087 |
| p.Leu134Glu | substitution | 134 | Leu | Glu | 0.104 | 0.050 |
| p.Leu134Gly | substitution | 134 | Leu | Gly | 0.188 | 0.039 |
| p.Leu134His | substitution | 134 | Leu | His | 0.000 | 0.012 |
| p.Leu134Ile | substitution | 134 | Leu | Ile | 0.698 | 0.013 |
| p.Leu134Lys | substitution | 134 | Leu | Lys | 0.000 | 0.061 |
| p.Leu134Met | substitution | 134 | Leu | Met | 0.781 | 0.006 |
| p.Leu134Phe | substitution | 134 | Leu | Phe | 0.529 | 0.005 |
| p.Leu134Pro | substitution | 134 | Leu | Pro | 0.161 | 0.055 |
| p.Leu134Ser | substitution | 134 | Leu | Ser | 0.000 | 0.034 |
| p.Leu134Ter | nonsense | 134 | Leu | Ter | 0.325 | 0.045 |
| p.Leu134Thr | substitution | 134 | Leu | Thr | 0.352 | 0.024 |
| p.Leu134Trp | substitution | 134 | Leu | Trp | 0.408 | 0.006 |
| p.Leu134Tyr | substitution | 134 | Leu | Tyr | 0.000 | 0.016 |
| p.Leu134Val | substitution | 134 | Leu | Val | 0.719 | 0.591 |
| p.Leu137= | synonymous | 137 | Leu | NA | 0.990 | 0.010 |
| p.Leu137Ala | substitution | 137 | Leu | Ala | 0.414 | 0.023 |
| p.Leu137Arg | substitution | 137 | Leu | Arg | 0.261 | 0.031 |
| p.Leu137Asn | substitution | 137 | Leu | Asn | 0.210 | 0.006 |
| p.Leu137Asp | substitution | 137 | Leu | Asp | 0.073 | 0.013 |
| p.Leu137Cys | substitution | 137 | Leu | Cys | 0.460 | 0.003 |
| p.Leu137Gln | substitution | 137 | Leu | Gln | 0.364 | 0.004 |
| p.Leu137Glu | substitution | 137 | Leu | Glu | 0.009 | 0.010 |
| p.Leu137Gly | substitution | 137 | Leu | Gly | 0.042 | 0.002 |
| p.Leu137His | substitution | 137 | Leu | His | 0.314 | 0.009 |
| p.Leu137Ile | substitution | 137 | Leu | Ile | 1.178 | 0.003 |
| p.Leu137Lys | substitution | 137 | Leu | Lys | 0.000 | 0.009 |
| p.Leu137Met | substitution | 137 | Leu | Met | 0.894 | 0.002 |
| p.Leu137Phe | substitution | 137 | Leu | Phe | 0.553 | 0.003 |
| p.Leu137Pro | substitution | 137 | Leu | Pro | 0.000 | 0.010 |
| p.Leu137Ser | substitution | 137 | Leu | Ser | 0.486 | 0.005 |
| p.Leu137Ter | nonsense | 137 | Leu | Ter | 0.000 | 0.006 |
| p.Leu137Thr | substitution | 137 | Leu | Thr | 0.213 | 0.007 |
| p.Leu137Trp | substitution | 137 | Leu | Trp | 0.000 | 0.005 |
| p.Leu137Tyr | substitution | 137 | Leu | Tyr | 0.209 | 0.005 |
| p.Leu137Val | substitution | 137 | Leu | Val | 0.870 | 0.002 |
| p.Leu148= | synonymous | 148 | Leu | NA | 1.077 | 0.006 |
| p.Leu148Ala | substitution | 148 | Leu | Ala | 0.000 | 0.158 |
| p.Leu148Arg | substitution | 148 | Leu | Arg | 0.743 | 0.052 |

| | | | | | | | |
|---|---|---|---|---|---|---|---|
| p.Leu148Asn | substitution | 148 | Leu | Asn | | 0.000 | 0.770 |
| p.Leu148Asp | substitution | 148 | Leu | Asp | | 0.491 | 0.021 |
| p.Leu148Cys | substitution | 148 | Leu | Cys | | 0.720 | 0.015 |
| p.Leu148Gln | substitution | 148 | Leu | Gln | | 0.893 | 0.012 |
| p.Leu148Glu | substitution | 148 | Leu | Glu | | 0.254 | 0.072 |
| p.Leu148Gly | substitution | 148 | Leu | Gly | | 1.106 | 0.007 |
| p.Leu148His | substitution | 148 | Leu | His | | 0.180 | 0.024 |
| p.Leu148Ile | substitution | 148 | Leu | Ile | | 1.056 | 0.010 |
| p.Leu148Lys | substitution | 148 | Leu | Lys | | 1.139 | 0.008 |
| p.Leu148Met | substitution | 148 | Leu | Met | | 1.195 | 0.008 |
| p.Leu148Phe | substitution | 148 | Leu | Phe | | 1.103 | 0.011 |
| p.Leu148Pro | substitution | 148 | Leu | Pro | | 1.521 | 0.105 |
| p.Leu148Ser | substitution | 148 | Leu | Ser | | 0.492 | 0.135 |
| p.Leu148Ter | nonsense | 148 | Leu | Ter | | 0.000 | 0.017 |
| p.Leu148Thr | substitution | 148 | Leu | Thr | | 0.519 | 0.256 |
| p.Leu148Trp | substitution | 148 | Leu | Trp | | 0.459 | 0.012 |
| p.Leu148Tyr | substitution | 148 | Leu | Tyr | | 0.000 | 0.014 |
| p.Leu148Val | substitution | 148 | Leu | Val | | 0.905 | 0.004 |
| p.Leu154= | synonymous | 154 | Leu | NA | | 0.972 | 0.009 |
| p.Leu154Ala | substitution | 154 | Leu | Ala | | 0.429 | 0.069 |
| p.Leu154Arg | substitution | 154 | Leu | Arg | | 0.197 | 0.012 |
| p.Leu154Asn | substitution | 154 | Leu | Asn | | 0.209 | 0.010 |
| p.Leu154Asp | substitution | 154 | Leu | Asp | | 0.302 | 0.022 |
| p.Leu154Cys | substitution | 154 | Leu | Cys | | 0.988 | 0.005 |
| p.Leu154Gln | substitution | 154 | Leu | Gln | | 0.695 | 0.012 |
| p.Leu154Glu | substitution | 154 | Leu | Glu | | 0.000 | 0.007 |
| p.Leu154Gly | substitution | 154 | Leu | Gly | | 0.804 | 0.003 |
| p.Leu154His | substitution | 154 | Leu | His | | 0.752 | 0.005 |
| p.Leu154Ile | substitution | 154 | Leu | Ile | | 0.335 | 0.021 |
| p.Leu154Lys | substitution | 154 | Leu | Lys | | 0.279 | 0.009 |
| p.Leu154Met | substitution | 154 | Leu | Met | | 0.764 | 0.010 |
| p.Leu154Phe | substitution | 154 | Leu | Phe | | 0.527 | 0.007 |
| p.Leu154Pro | substitution | 154 | Leu | Pro | | 0.000 | 0.047 |
| p.Leu154Ser | substitution | 154 | Leu | Ser | | 0.459 | 0.070 |
| p.Leu154Ter | nonsense | 154 | Leu | Ter | | 0.000 | 0.030 |
| p.Leu154Thr | substitution | 154 | Leu | Thr | | 0.607 | 0.018 |
| p.Leu154Trp | substitution | 154 | Leu | Trp | | 0.513 | 0.007 |
| p.Leu154Tyr | substitution | 154 | Leu | Tyr | | 0.489 | 0.007 |
| p.Leu154Val | substitution | 154 | Leu | Val | | 0.807 | 0.010 |
| p.Leu161= | synonymous | 161 | Leu | NA | | 1.017 | 0.018 |
| p.Leu161Ala | substitution | 161 | Leu | Ala | | 0.348 | 0.013 |
| p.Leu161Arg | substitution | 161 | Leu | Arg | | 0.000 | 0.032 |
| p.Leu161Asn | substitution | 161 | Leu | Asn | | 0.131 | 0.035 |

| | | | | | | |
|---|---|---|---|---|---|---|
| p.Leu161Asp | substitution | 161 | Leu | Asp | 0.000 | 0.045 |
| p.Leu161Cys | substitution | 161 | Leu | Cys | 0.979 | 0.005 |
| p.Leu161Gln | substitution | 161 | Leu | Gln | 0.979 | 0.014 |
| p.Leu161Glu | substitution | 161 | Leu | Glu | 0.000 | 0.019 |
| p.Leu161Gly | substitution | 161 | Leu | Gly | 0.166 | 0.009 |
| p.Leu161His | substitution | 161 | Leu | His | 0.873 | 0.006 |
| p.Leu161Ile | substitution | 161 | Leu | Ile | 0.749 | 0.009 |
| p.Leu161Lys | substitution | 161 | Leu | Lys | 0.696 | 0.007 |
| p.Leu161Met | substitution | 161 | Leu | Met | 0.951 | 0.008 |
| p.Leu161Phe | substitution | 161 | Leu | Phe | 0.552 | 0.010 |
| p.Leu161Pro | substitution | 161 | Leu | Pro | 0.000 | 0.043 |
| p.Leu161Ser | substitution | 161 | Leu | Ser | 0.475 | 0.012 |
| p.Leu161Ter | nonsense | 161 | Leu | Ter | 0.025 | 0.010 |
| p.Leu161Thr | substitution | 161 | Leu | Thr | 0.656 | 0.022 |
| p.Leu161Trp | substitution | 161 | Leu | Trp | 0.519 | 0.007 |
| p.Leu161Tyr | substitution | 161 | Leu | Tyr | 0.902 | 0.006 |
| p.Leu161Val | substitution | 161 | Leu | Val | 0.568 | 0.008 |
| p.Leu170= | synonymous | 170 | Leu | NA | 0.918 | 0.008 |
| p.Leu170Ala | substitution | 170 | Leu | Ala | 0.551 | 0.037 |
| p.Leu170Arg | substitution | 170 | Leu | Arg | 0.000 | 0.018 |
| p.Leu170Asn | substitution | 170 | Leu | Asn | 0.781 | 0.019 |
| p.Leu170Asp | substitution | 170 | Leu | Asp | 0.000 | 0.045 |
| p.Leu170Cys | substitution | 170 | Leu | Cys | 0.691 | 0.014 |
| p.Leu170Gln | substitution | 170 | Leu | Gln | 1.097 | 0.007 |
| p.Leu170Glu | substitution | 170 | Leu | Glu | 0.527 | 0.019 |
| p.Leu170Gly | substitution | 170 | Leu | Gly | 0.958 | 0.031 |
| p.Leu170His | substitution | 170 | Leu | His | 0.720 | 0.040 |
| p.Leu170Ile | substitution | 170 | Leu | Ile | 0.790 | 0.042 |
| p.Leu170Lys | substitution | 170 | Leu | Lys | 0.038 | 0.027 |
| p.Leu170Met | substitution | 170 | Leu | Met | 1.074 | 0.008 |
| p.Leu170Phe | substitution | 170 | Leu | Phe | 0.904 | 0.008 |
| p.Leu170Pro | substitution | 170 | Leu | Pro | 0.000 | 0.014 |
| p.Leu170Ser | substitution | 170 | Leu | Ser | 0.520 | 0.034 |
| p.Leu170Ter | nonsense | 170 | Leu | Ter | 0.042 | 0.009 |
| p.Leu170Thr | substitution | 170 | Leu | Thr | 0.589 | 0.030 |
| p.Leu170Trp | substitution | 170 | Leu | Trp | 0.000 | 0.018 |
| p.Leu170Tyr | substitution | 170 | Leu | Tyr | 1.333 | 0.012 |
| p.Leu170Val | substitution | 170 | Leu | Val | 0.848 | 0.012 |
| p.Leu174= | synonymous | 174 | Leu | NA | 0.987 | 0.010 |
| p.Leu174Ala | substitution | 174 | Leu | Ala | 0.454 | 0.019 |
| p.Leu174Arg | substitution | 174 | Leu | Arg | 0.659 | 0.019 |
| p.Leu174Asn | substitution | 174 | Leu | Asn | 0.208 | 0.007 |
| p.Leu174Asp | substitution | 174 | Leu | Asp | 0.000 | 0.007 |

| | | | | | | |
|---|---|---|---|---|---|---|
| p.Leu174Cys | substitution | 174 | Leu | Cys | 0.960 | 0.004 |
| p.Leu174Gln | substitution | 174 | Leu | Gln | 0.394 | 0.009 |
| p.Leu174Glu | substitution | 174 | Leu | Glu | 0.142 | 0.013 |
| p.Leu174Gly | substitution | 174 | Leu | Gly | 0.215 | 0.014 |
| p.Leu174His | substitution | 174 | Leu | His | 0.688 | 0.007 |
| p.Leu174Ile | substitution | 174 | Leu | Ile | 0.944 | 0.001 |
| p.Leu174Met | substitution | 174 | Leu | Met | 1.127 | 0.005 |
| p.Leu174Phe | substitution | 174 | Leu | Phe | 1.171 | 0.005 |
| p.Leu174Pro | substitution | 174 | Leu | Pro | 0.000 | 0.032 |
| p.Leu174Ser | substitution | 174 | Leu | Ser | 0.415 | 0.007 |
| p.Leu174Ter | nonsense | 174 | Leu | Ter | 0.000 | 0.012 |
| p.Leu174Thr | substitution | 174 | Leu | Thr | 0.588 | 0.049 |
| p.Leu174Trp | substitution | 174 | Leu | Trp | 0.404 | 0.010 |
| p.Leu174Tyr | substitution | 174 | Leu | Tyr | 0.695 | 0.007 |
| p.Leu174Val | substitution | 174 | Leu | Val | 0.814 | 0.017 |
| p.Leu177= | synonymous | 177 | Leu | NA | 1.116 | 0.014 |
| p.Leu177Ala | substitution | 177 | Leu | Ala | 0.294 | 0.007 |
| p.Leu177Arg | substitution | 177 | Leu | Arg | 0.121 | 0.007 |
| p.Leu177Asn | substitution | 177 | Leu | Asn | 0.554 | 0.011 |
| p.Leu177Asp | substitution | 177 | Leu | Asp | 0.169 | 0.035 |
| p.Leu177Cys | substitution | 177 | Leu | Cys | 0.509 | 0.007 |
| p.Leu177Gln | substitution | 177 | Leu | Gln | 0.284 | 0.009 |
| p.Leu177Glu | substitution | 177 | Leu | Glu | 0.000 | 0.009 |
| p.Leu177Gly | substitution | 177 | Leu | Gly | 0.000 | 0.008 |
| p.Leu177His | substitution | 177 | Leu | His | 0.000 | 0.010 |
| p.Leu177Ile | substitution | 177 | Leu | Ile | 0.527 | 0.012 |
| p.Leu177Lys | substitution | 177 | Leu | Lys | 0.000 | 0.015 |
| p.Leu177Met | substitution | 177 | Leu | Met | 0.988 | 0.006 |
| p.Leu177Phe | substitution | 177 | Leu | Phe | 0.354 | 0.005 |
| p.Leu177Pro | substitution | 177 | Leu | Pro | 0.000 | 0.751 |
| p.Leu177Ser | substitution | 177 | Leu | Ser | 0.000 | 0.003 |
| p.Leu177Ter | nonsense | 177 | Leu | Ter | 0.000 | 0.020 |
| p.Leu177Thr | substitution | 177 | Leu | Thr | 0.481 | 0.009 |
| p.Leu177Trp | substitution | 177 | Leu | Trp | 0.000 | 0.023 |
| p.Leu177Tyr | substitution | 177 | Leu | Tyr | 0.000 | 0.014 |
| p.Leu177Val | substitution | 177 | Leu | Val | 0.953 | 0.027 |
| p.Leu188= | synonymous | 188 | Leu | NA | 0.988 | 0.016 |
| p.Leu188Ala | substitution | 188 | Leu | Ala | 0.872 | 0.015 |
| p.Leu188Arg | substitution | 188 | Leu | Arg | 0.000 | 0.003 |
| p.Leu188Asn | substitution | 188 | Leu | Asn | 0.450 | 0.035 |
| p.Leu188Asp | substitution | 188 | Leu | Asp | 0.000 | 0.017 |
| p.Leu188Cys | substitution | 188 | Leu | Cys | 0.893 | 0.002 |
| p.Leu188Gln | substitution | 188 | Leu | Gln | 0.496 | 0.009 |

| | | | | | | |
|---|---|---|---|---|---|---|
| p.Leu188Glu | substitution | 188 | Leu | Glu | 0.000 | 0.009 |
| p.Leu188Gly | substitution | 188 | Leu | Gly | 0.729 | 0.031 |
| p.Leu188His | substitution | 188 | Leu | His | 0.956 | 0.001 |
| p.Leu188Ile | substitution | 188 | Leu | Ile | 0.063 | 0.006 |
| p.Leu188Lys | substitution | 188 | Leu | Lys | 0.000 | 0.013 |
| p.Leu188Met | substitution | 188 | Leu | Met | 0.673 | 0.005 |
| p.Leu188Phe | substitution | 188 | Leu | Phe | 0.649 | 0.004 |
| p.Leu188Pro | substitution | 188 | Leu | Pro | 0.240 | 0.018 |
| p.Leu188Ser | substitution | 188 | Leu | Ser | 0.425 | 0.064 |
| p.Leu188Ter | nonsense | 188 | Leu | Ter | 0.231 | 0.005 |
| p.Leu188Thr | substitution | 188 | Leu | Thr | 0.845 | 0.040 |
| p.Leu188Trp | substitution | 188 | Leu | Trp | 0.000 | 0.008 |
| p.Leu188Tyr | substitution | 188 | Leu | Tyr | 0.000 | 0.018 |
| p.Leu188Val | substitution | 188 | Leu | Val | 0.000 | 0.014 |
| p.Leu193= | synonymous | 193 | Leu | NA | 0.933 | 0.016 |
| p.Leu193Ala | substitution | 193 | Leu | Ala | 0.000 | 0.062 |
| p.Leu193Arg | substitution | 193 | Leu | Arg | 0.000 | 0.004 |
| p.Leu193Asn | substitution | 193 | Leu | Asn | 0.451 | 0.016 |
| p.Leu193Asp | substitution | 193 | Leu | Asp | 0.000 | 0.011 |
| p.Leu193Cys | substitution | 193 | Leu | Cys | 0.000 | 0.019 |
| p.Leu193Gln | substitution | 193 | Leu | Gln | 0.522 | 0.006 |
| p.Leu193Glu | substitution | 193 | Leu | Glu | 0.000 | 0.021 |
| p.Leu193Gly | substitution | 193 | Leu | Gly | 1.029 | 0.002 |
| p.Leu193His | substitution | 193 | Leu | His | 0.123 | 0.008 |
| p.Leu193Ile | substitution | 193 | Leu | Ile | 0.292 | 0.010 |
| p.Leu193Lys | substitution | 193 | Leu | Lys | 0.000 | 0.021 |
| p.Leu193Met | substitution | 193 | Leu | Met | 1.094 | 0.001 |
| p.Leu193Phe | substitution | 193 | Leu | Phe | 0.424 | 0.009 |
| p.Leu193Pro | substitution | 193 | Leu | Pro | 0.622 | 0.170 |
| p.Leu193Ser | substitution | 193 | Leu | Ser | 0.637 | 0.007 |
| p.Leu193Ter | nonsense | 193 | Leu | Ter | 0.033 | 0.056 |
| p.Leu193Thr | substitution | 193 | Leu | Thr | 0.361 | 0.041 |
| p.Leu193Trp | substitution | 193 | Leu | Trp | 0.000 | 0.021 |
| p.Leu193Tyr | substitution | 193 | Leu | Tyr | 0.000 | 0.037 |
| p.Leu193Val | substitution | 193 | Leu | Val | 0.272 | 0.047 |
| p.Leu206= | synonymous | 206 | Leu | NA | 0.708 | 0.017 |
| p.Leu206Ala | substitution | 206 | Leu | Ala | 0.000 | 0.016 |
| p.Leu206Arg | substitution | 206 | Leu | Arg | 0.032 | 0.002 |
| p.Leu206Asn | substitution | 206 | Leu | Asn | 0.000 | 0.062 |
| p.Leu206Asp | substitution | 206 | Leu | Asp | 0.273 | 0.005 |
| p.Leu206Cys | substitution | 206 | Leu | Cys | 0.790 | 0.001 |
| p.Leu206Gln | substitution | 206 | Leu | Gln | 0.059 | 0.014 |
| p.Leu206Glu | substitution | 206 | Leu | Glu | 0.000 | 0.006 |

| | | | | | | | |
|---|---|---|---|---|---|---|---|
| p.Leu206Gly | substitution | 206 | Leu | Gly | | 0.449 | 0.016 |
| p.Leu206His | substitution | 206 | Leu | His | | 0.000 | 0.101 |
| p.Leu206Ile | substitution | 206 | Leu | Ile | | 0.027 | 0.008 |
| p.Leu206Lys | substitution | 206 | Leu | Lys | | 0.000 | 0.007 |
| p.Leu206Met | substitution | 206 | Leu | Met | | 1.129 | 0.002 |
| p.Leu206Phe | substitution | 206 | Leu | Phe | | 0.950 | 0.001 |
| p.Leu206Pro | substitution | 206 | Leu | Pro | | 0.285 | 0.120 |
| p.Leu206Ser | substitution | 206 | Leu | Ser | | 0.176 | 0.018 |
| p.Leu206Ter | nonsense | 206 | Leu | Ter | | 0.000 | 0.022 |
| p.Leu206Thr | substitution | 206 | Leu | Thr | | 0.432 | 0.193 |
| p.Leu206Trp | substitution | 206 | Leu | Trp | | 0.207 | 0.002 |
| p.Leu206Tyr | substitution | 206 | Leu | Tyr | | 0.000 | 0.014 |
| p.Leu206Val | substitution | 206 | Leu | Val | | 0.499 | 0.152 |
| p.Leu220= | synonymous | 220 | Leu | NA | | 0.755 | 0.036 |
| p.Leu220Ala | substitution | 220 | Leu | Ala | | 1.124 | 0.015 |
| p.Leu220Arg | substitution | 220 | Leu | Arg | | 0.042 | 0.014 |
| p.Leu220Asn | substitution | 220 | Leu | Asn | | 0.004 | 0.019 |
| p.Leu220Asp | substitution | 220 | Leu | Asp | | 0.149 | 0.031 |
| p.Leu220Cys | substitution | 220 | Leu | Cys | | 0.762 | 0.004 |
| p.Leu220Gln | substitution | 220 | Leu | Gln | | 0.000 | 0.008 |
| p.Leu220Glu | substitution | 220 | Leu | Glu | | 0.000 | 0.009 |
| p.Leu220Gly | substitution | 220 | Leu | Gly | | 0.088 | 0.049 |
| p.Leu220His | substitution | 220 | Leu | His | | 0.059 | 0.009 |
| p.Leu220Ile | substitution | 220 | Leu | Ile | | 0.000 | 0.060 |
| p.Leu220Met | substitution | 220 | Leu | Met | | 1.001 | 0.003 |
| p.Leu220Phe | substitution | 220 | Leu | Phe | | 0.475 | 0.012 |
| p.Leu220Pro | substitution | 220 | Leu | Pro | | 0.000 | 0.076 |
| p.Leu220Ser | substitution | 220 | Leu | Ser | | 0.399 | 0.144 |
| p.Leu220Ter | nonsense | 220 | Leu | Ter | | 0.000 | 0.014 |
| p.Leu220Thr | substitution | 220 | Leu | Thr | | 0.881 | 0.030 |
| p.Leu220Trp | substitution | 220 | Leu | Trp | | 0.320 | 0.007 |
| p.Leu220Tyr | substitution | 220 | Leu | Tyr | | 0.721 | 0.005 |
| p.Leu220Val | substitution | 220 | Leu | Val | | 0.886 | 0.011 |
| p.Leu232= | synonymous | 232 | Leu | NA | | 1.439 | 0.001 |
| p.Leu232Ala | substitution | 232 | Leu | Ala | | 0.608 | 0.043 |
| p.Leu232Arg | substitution | 232 | Leu | Arg | | 0.218 | 0.058 |
| p.Leu232Asn | substitution | 232 | Leu | Asn | | 1.049 | 0.003 |
| p.Leu232Asp | substitution | 232 | Leu | Asp | | 0.354 | 0.008 |
| p.Leu232Cys | substitution | 232 | Leu | Cys | | 0.889 | 0.003 |
| p.Leu232Gln | substitution | 232 | Leu | Gln | | 0.000 | 0.013 |
| p.Leu232Glu | substitution | 232 | Leu | Glu | | 0.184 | 0.010 |
| p.Leu232Gly | substitution | 232 | Leu | Gly | | 1.006 | 0.004 |
| p.Leu232His | substitution | 232 | Leu | His | | 0.511 | 0.005 |

| | | | | | | |
|---|---|---|---|---|---|---|
| p.Leu232Ile | substitution | 232 | Leu | Ile | 1.100 | 0.004 |
| p.Leu232Lys | substitution | 232 | Leu | Lys | 0.999 | 0.003 |
| p.Leu232Met | substitution | 232 | Leu | Met | 0.910 | 0.003 |
| p.Leu232Phe | substitution | 232 | Leu | Phe | 0.899 | 0.009 |
| p.Leu232Pro | substitution | 232 | Leu | Pro | 0.394 | 0.026 |
| p.Leu232Ser | substitution | 232 | Leu | Ser | 0.139 | 0.052 |
| p.Leu232Ter | nonsense | 232 | Leu | Ter | 0.384 | 0.011 |
| p.Leu232Thr | substitution | 232 | Leu | Thr | 0.980 | 0.001 |
| p.Leu232Trp | substitution | 232 | Leu | Trp | 0.645 | 0.002 |
| p.Leu232Tyr | substitution | 232 | Leu | Tyr | 0.278 | 0.052 |
| p.Leu232Val | substitution | 232 | Leu | Val | 1.133 | 0.033 |
| p.Leu234= | synonymous | 234 | Leu | NA | 0.962 | 0.011 |
| p.Leu234Ala | substitution | 234 | Leu | Ala | 1.135 | 0.002 |
| p.Leu234Arg | substitution | 234 | Leu | Arg | 0.569 | 0.223 |
| p.Leu234Asn | substitution | 234 | Leu | Asn | 1.180 | 0.003 |
| p.Leu234Asp | substitution | 234 | Leu | Asp | 0.226 | 0.019 |
| p.Leu234Cys | substitution | 234 | Leu | Cys | 0.909 | 0.002 |
| p.Leu234Gln | substitution | 234 | Leu | Gln | 0.665 | 0.006 |
| p.Leu234Glu | substitution | 234 | Leu | Glu | 0.000 | 0.010 |
| p.Leu234Gly | substitution | 234 | Leu | Gly | 0.110 | 0.011 |
| p.Leu234His | substitution | 234 | Leu | His | 0.609 | 0.012 |
| p.Leu234Lys | substitution | 234 | Leu | Lys | 0.939 | 0.004 |
| p.Leu234Met | substitution | 234 | Leu | Met | 0.501 | 0.005 |
| p.Leu234Phe | substitution | 234 | Leu | Phe | 0.593 | 0.003 |
| p.Leu234Pro | substitution | 234 | Leu | Pro | 0.000 | 0.068 |
| p.Leu234Ser | substitution | 234 | Leu | Ser | 0.047 | 0.027 |
| p.Leu234Ter | nonsense | 234 | Leu | Ter | 0.245 | 0.074 |
| p.Leu234Thr | substitution | 234 | Leu | Thr | 0.437 | 0.067 |
| p.Leu234Trp | substitution | 234 | Leu | Trp | 0.894 | 0.002 |
| p.Leu234Tyr | substitution | 234 | Leu | Tyr | 0.000 | 0.067 |
| p.Leu234Val | substitution | 234 | Leu | Val | 1.091 | 0.016 |
| p.Leu238= | synonymous | 238 | Leu | NA | 0.919 | 0.008 |
| p.Leu238Ala | substitution | 238 | Leu | Ala | 0.000 | 0.075 |
| p.Leu238Arg | substitution | 238 | Leu | Arg | 0.000 | 0.021 |
| p.Leu238Asn | substitution | 238 | Leu | Asn | 0.000 | 0.117 |
| p.Leu238Asp | substitution | 238 | Leu | Asp | 0.883 | 0.049 |
| p.Leu238Cys | substitution | 238 | Leu | Cys | 0.098 | 0.228 |
| p.Leu238Gln | substitution | 238 | Leu | Gln | 0.565 | 0.043 |
| p.Leu238Glu | substitution | 238 | Leu | Glu | 0.048 | 0.015 |
| p.Leu238Gly | substitution | 238 | Leu | Gly | 0.000 | 0.096 |
| p.Leu238His | substitution | 238 | Leu | His | 0.000 | 0.139 |
| p.Leu238Lys | substitution | 238 | Leu | Lys | 0.000 | 0.257 |
| p.Leu238Met | substitution | 238 | Leu | Met | 1.236 | 0.001 |

| | | | | | | |
|---|---|---|---|---|---|---|
| p.Leu238Phe | substitution | 238 | Leu | Phe | 0.000 | 0.014 |
| p.Leu238Pro | substitution | 238 | Leu | Pro | 0.555 | 0.066 |
| p.Leu238Ser | substitution | 238 | Leu | Ser | 0.127 | 0.013 |
| p.Leu238Ter | nonsense | 238 | Leu | Ter | 0.000 | 0.125 |
| p.Leu238Thr | substitution | 238 | Leu | Thr | 0.000 | 0.145 |
| p.Leu238Trp | substitution | 238 | Leu | Trp | 0.098 | 0.009 |
| p.Leu238Val | substitution | 238 | Leu | Val | 0.583 | 0.047 |
| p.Leu244= | synonymous | 244 | Leu | NA | 0.990 | 0.014 |
| p.Leu244Ala | substitution | 244 | Leu | Ala | 0.123 | 0.031 |
| p.Leu244Arg | substitution | 244 | Leu | Arg | 0.088 | 0.007 |
| p.Leu244Asn | substitution | 244 | Leu | Asn | 0.298 | 0.015 |
| p.Leu244Asp | substitution | 244 | Leu | Asp | 0.000 | 0.200 |
| p.Leu244Cys | substitution | 244 | Leu | Cys | 0.969 | 0.003 |
| p.Leu244Gln | substitution | 244 | Leu | Gln | 1.413 | 0.003 |
| p.Leu244Glu | substitution | 244 | Leu | Glu | 0.000 | 0.078 |
| p.Leu244Gly | substitution | 244 | Leu | Gly | 0.000 | 0.040 |
| p.Leu244His | substitution | 244 | Leu | His | 0.082 | 0.012 |
| p.Leu244Ile | substitution | 244 | Leu | Ile | 0.000 | 0.011 |
| p.Leu244Lys | substitution | 244 | Leu | Lys | 0.000 | 0.021 |
| p.Leu244Me | substitution | 244 | Leu | Met | 0.719 | 0.005 |
| p.Leu244Phe | substitution | 244 | Leu | Phe | 0.000 | 0.006 |
| p.Leu244Pro | substitution | 244 | Leu | Pro | 0.142 | 0.035 |
| p.Leu244Ser | substitution | 244 | Leu | Ser | 1.131 | 0.137 |
| p.Leu244Ter | nonsense | 244 | Leu | Ter | 0.273 | 0.051 |
| p.Leu244Thr | substitution | 244 | Leu | Thr | 1.106 | 0.152 |
| p.Leu244Trp | substitution | 244 | Leu | Trp | 0.000 | 0.014 |
| p.Leu244Tyr | substitution | 244 | Leu | Tyr | 1.182 | 0.002 |
| p.Leu244Val | substitution | 244 | Leu | Val | 0.748 | 0.172 |
| p.Leu245= | synonymous | 245 | Leu | NA | 0.869 | 0.057 |
| p.Leu245Ala | substitution | 245 | Leu | Ala | 1.132 | 0.054 |
| p.Leu245Arg | substitution | 245 | Leu | Arg | 0.031 | 0.026 |
| p.Leu245Asp | substitution | 245 | Leu | Asp | 0.000 | 0.054 |
| p.Leu245Cys | substitution | 245 | Leu | Cys | 0.847 | 0.003 |
| p.Leu245Gln | substitution | 245 | Leu | Gln | 0.726 | 0.003 |
| p.Leu245Glu | substitution | 245 | Leu | Glu | 0.085 | 0.004 |
| p.Leu245Gly | substitution | 245 | Leu | Gly | 0.226 | 0.067 |
| p.Leu245His | substitution | 245 | Leu | His | 0.719 | 0.038 |
| p.Leu245Ile | substitution | 245 | Leu | Ile | 0.963 | 0.002 |
| p.Leu245Lys | substitution | 245 | Leu | Lys | 0.143 | 0.013 |
| p.Leu245Me | substitution | 245 | Leu | Met | 1.159 | 0.007 |
| p.Leu245Phe | substitution | 245 | Leu | Phe | 0.439 | 0.071 |
| p.Leu245Pro | substitution | 245 | Leu | Pro | 0.000 | 0.130 |
| p.Leu245Ser | substitution | 245 | Leu | Ser | 0.000 | 0.030 |

| | | | | | | |
|---|---|---|---|---|---|---|
| p.Leu245Thr | substitution | 245 | Leu | Thr | 0.000 | 0.025 |
| p.Leu245Trp | substitution | 245 | Leu | Trp | 0.807 | 0.003 |
| p.Leu245Tyr | substitution | 245 | Leu | Tyr | 1.185 | 0.014 |
| p.Leu245Val | substitution | 245 | Leu | Val | 0.754 | 0.079 |
| p.Leu254= | synonymous | 254 | Leu | NA | 0.941 | 0.078 |
| p.Leu254Ala | substitution | 254 | Leu | Ala | 0.000 | 0.014 |
| p.Leu254Arg | substitution | 254 | Leu | Arg | 0.069 | 0.221 |
| p.Leu254Cys | substitution | 254 | Leu | Cys | 0.000 | 0.038 |
| p.Leu254Gln | substitution | 254 | Leu | Gln | 0.967 | 0.009 |
| p.Leu254Gly | substitution | 254 | Leu | Gly | 0.000 | 0.082 |
| p.Leu254Ile | substitution | 254 | Leu | Ile | 0.000 | 0.025 |
| p.Leu254Lys | substitution | 254 | Leu | Lys | 0.000 | 0.072 |
| p.Leu254Met | substitution | 254 | Leu | Met | 0.720 | 0.008 |
| p.Leu254Phe | substitution | 254 | Leu | Phe | 0.000 | 0.093 |
| p.Leu254Pro | substitution | 254 | Leu | Pro | 0.432 | 0.664 |
| p.Leu254Ser | substitution | 254 | Leu | Ser | 0.445 | 0.037 |
| p.Leu254Ter | nonsense | 254 | Leu | Ter | 0.000 | 0.132 |
| p.Leu254Trp | substitution | 254 | Leu | Trp | 1.293 | 0.002 |
| p.Leu254Tyr | substitution | 254 | Leu | Tyr | 0.000 | 0.164 |
| p.Leu254Val | substitution | 254 | Leu | Val | 0.845 | 0.158 |
| p.Leu257= | synonymous | 257 | Leu | NA | 2.078 | 0.123 |
| p.Leu257Arg | substitution | 257 | Leu | Arg | 1.340 | 0.162 |
| p.Leu257Met | substitution | 257 | Leu | Met | 0.963 | 0.005 |
| p.Leu257Val | substitution | 257 | Leu | Val | 1.129 | 0.030 |
| p.Leu276= | synonymous | 276 | Leu | NA | 0.935 | 0.034 |
| p.Leu276Ala | substitution | 276 | Leu | Ala | 0.193 | 0.026 |
| p.Leu276Arg | substitution | 276 | Leu | Arg | 0.000 | 0.017 |
| p.Leu276Asn | substitution | 276 | Leu | Asn | 0.000 | 0.145 |
| p.Leu276Asp | substitution | 276 | Leu | Asp | 0.039 | 0.009 |
| p.Leu276Cys | substitution | 276 | Leu | Cys | 0.853 | 0.006 |
| p.Leu276Gln | substitution | 276 | Leu | Gln | 0.207 | 0.239 |
| p.Leu276Glu | substitution | 276 | Leu | Glu | 0.008 | 0.015 |
| p.Leu276Gly | substitution | 276 | Leu | Gly | 0.307 | 0.084 |
| p.Leu276His | substitution | 276 | Leu | His | 0.000 | 0.017 |
| p.Leu276Ile | substitution | 276 | Leu | Ile | 0.316 | 0.042 |
| p.Leu276Lys | substitution | 276 | Leu | Lys | 0.000 | 0.012 |
| p.Leu276Met | substitution | 276 | Leu | Met | 1.189 | 0.005 |
| p.Leu276Phe | substitution | 276 | Leu | Phe | 0.156 | 0.005 |
| p.Leu276Pro | substitution | 276 | Leu | Pro | 0.000 | 0.088 |
| p.Leu276Ser | substitution | 276 | Leu | Ser | 0.014 | 0.012 |
| p.Leu276Ter | nonsense | 276 | Leu | Ter | 0.000 | 0.010 |
| p.Leu276Thr | substitution | 276 | Leu | Thr | 0.000 | 0.084 |
| p.Leu276Trp | substitution | 276 | Leu | Trp | 0.000 | 0.021 |

| | | | | | | | |
|---|---|---|---|---|---|---|---|
| p.Leu276Tyr | substitution | 276 | Leu | Tyr | | 1.034 | 0.004 |
| p.Leu276Val | substitution | 276 | Leu | Val | | 1.066 | 0.004 |
| p.Leu278= | synonymous | 278 | Leu | NA | | 1.046 | 0.016 |
| p.Leu278Ala | substitution | 278 | Leu | Ala | | 0.302 | 0.007 |
| p.Leu278Arg | substitution | 278 | Leu | Arg | | 0.094 | 0.009 |
| p.Leu278Asn | substitution | 278 | Leu | Asn | | 0.000 | 0.016 |
| p.Leu278Asp | substitution | 278 | Leu | Asp | | 0.000 | 0.060 |
| p.Leu278Cys | substitution | 278 | Leu | Cys | | 0.967 | 0.005 |
| p.Leu278Gln | substitution | 278 | Leu | Gln | | 1.371 | 0.010 |
| p.Leu278Glu | substitution | 278 | Leu | Glu | | 0.000 | 0.016 |
| p.Leu278Gly | substitution | 278 | Leu | Gly | | 0.745 | 0.075 |
| p.Leu278His | substitution | 278 | Leu | His | | 0.025 | 0.207 |
| p.Leu278Lys | substitution | 278 | Leu | Lys | | 0.000 | 0.009 |
| p.Leu278Met | substitution | 278 | Leu | Met | | 0.449 | 0.010 |
| p.Leu278Phe | substitution | 278 | Leu | Phe | | 0.765 | 0.013 |
| p.Leu278Pro | substitution | 278 | Leu | Pro | | 0.587 | 0.096 |
| p.Leu278Ser | substitution | 278 | Leu | Ser | | 0.543 | 0.015 |
| p.Leu278Ter | nonsense | 278 | Leu | Ter | | 0.000 | 0.011 |
| p.Leu278Thr | substitution | 278 | Leu | Thr | | 0.483 | 0.115 |
| p.Leu278Trp | substitution | 278 | Leu | Trp | | 0.162 | 0.007 |
| p.Leu278Tyr | substitution | 278 | Leu | Tyr | | 0.000 | 0.018 |
| p.Leu278Val | substitution | 278 | Leu | Val | | 0.381 | 0.041 |
| p.Leu285= | synonymous | 285 | Leu | NA | | 0.932 | 0.011 |
| p.Leu285Ala | substitution | 285 | Leu | Ala | | 1.003 | 0.011 |
| p.Leu285Arg | substitution | 285 | Leu | Arg | | 0.802 | 0.011 |
| p.Leu285Asn | substitution | 285 | Leu | Asn | | 1.418 | 0.004 |
| p.Leu285Asp | substitution | 285 | Leu | Asp | | 1.174 | 0.006 |
| p.Leu285Cys | substitution | 285 | Leu | Cys | | 0.471 | 0.004 |
| p.Leu285Gln | substitution | 285 | Leu | Gln | | 0.595 | 0.040 |
| p.Leu285Glu | substitution | 285 | Leu | Glu | | 0.238 | 0.055 |
| p.Leu285Gly | substitution | 285 | Leu | Gly | | 1.211 | 0.006 |
| p.Leu285His | substitution | 285 | Leu | His | | 0.648 | 0.004 |
| p.Leu285Ile | substitution | 285 | Leu | Ile | | 0.766 | 0.011 |
| p.Leu285Lys | substitution | 285 | Leu | Lys | | 0.739 | 0.042 |
| p.Leu285Met | substitution | 285 | Leu | Met | | 0.962 | 0.004 |
| p.Leu285Phe | substitution | 285 | Leu | Phe | | 1.076 | 0.002 |
| p.Leu285Pro | substitution | 285 | Leu | Pro | | 1.171 | 0.004 |
| p.Leu285Ser | substitution | 285 | Leu | Ser | | 0.000 | 0.048 |
| p.Leu285Ter | nonsense | 285 | Leu | Ter | | 0.101 | 0.096 |
| p.Leu285Thr | substitution | 285 | Leu | Thr | | 0.794 | 0.032 |
| p.Leu285Trp | substitution | 285 | Leu | Trp | | 0.620 | 0.005 |
| p.Leu285Tyr | substitution | 285 | Leu | Tyr | | 1.216 | 0.004 |
| p.Leu285Val | substitution | 285 | Leu | Val | | 0.778 | 0.013 |

| | | | | | | |
|---|---|---|---|---|---|---|
| p.Leu30= | synonymous | 30 | Leu | NA | 0.935 | 0.000 |
| p.Leu30Ala | substitution | 30 | Leu | Ala | 0.000 | 0.002 |
| p.Leu30Arg | substitution | 30 | Leu | Arg | 0.000 | 0.001 |
| p.Leu30Asn | substitution | 30 | Leu | Asn | 0.000 | 0.002 |
| p.Leu30Asp | substitution | 30 | Leu | Asp | 0.000 | 0.003 |
| p.Leu30Cys | substitution | 30 | Leu | Cys | 0.842 | 0.000 |
| p.Leu30Gln | substitution | 30 | Leu | Gln | 0.000 | 0.004 |
| p.Leu30Glu | substitution | 30 | Leu | Glu | 0.000 | 0.001 |
| p.Leu30Gly | substitution | 30 | Leu | Gly | 0.000 | 0.001 |
| p.Leu30His | substitution | 30 | Leu | His | 0.000 | 0.002 |
| p.Leu30Ile | substitution | 30 | Leu | Ile | 0.000 | 0.001 |
| p.Leu30Lys | substitution | 30 | Leu | Lys | 0.000 | 0.002 |
| p.Leu30Met | substitution | 30 | Leu | Met | 0.570 | 0.000 |
| p.Leu30Phe | substitution | 30 | Leu | Phe | 0.000 | 0.002 |
| p.Leu30Pro | substitution | 30 | Leu | Pro | 0.000 | 0.004 |
| p.Leu30Ser | substitution | 30 | Leu | Ser | 0.000 | 0.005 |
| p.Leu30Ter | nonsense | 30 | Leu | Ter | 0.000 | 0.003 |
| p.Leu30Thr | substitution | 30 | Leu | Thr | 0.000 | 0.001 |
| p.Leu30Trp | substitution | 30 | Leu | Trp | 0.000 | 0.001 |
| p.Leu30Tyr | substitution | 30 | Leu | Tyr | 0.208 | 0.001 |
| p.Leu30Val | substitution | 30 | Leu | Val | 0.616 | 0.004 |
| p.Leu315= | synonymous | 315 | Leu | NA | 0.925 | 0.063 |
| p.Leu315Ala | substitution | 315 | Leu | Ala | 0.502 | 0.204 |
| p.Leu315Arg | substitution | 315 | Leu | Arg | 0.161 | 0.308 |
| p.Leu315Asn | substitution | 315 | Leu | Asn | 0.204 | 0.018 |
| p.Leu315Asp | substitution | 315 | Leu | Asp | 0.109 | 0.011 |
| p.Leu315Cys | substitution | 315 | Leu | Cys | 1.428 | 0.004 |
| p.Leu315Gln | substitution | 315 | Leu | Gln | 0.068 | 0.035 |
| p.Leu315Glu | substitution | 315 | Leu | Glu | 0.000 | 0.024 |
| p.Leu315Gly | substitution | 315 | Leu | Gly | 0.264 | 0.104 |
| p.Leu315Ile | substitution | 315 | Leu | Ile | 0.000 | 0.023 |
| p.Leu315Lys | substitution | 315 | Leu | Lys | 0.557 | 0.018 |
| p.Leu315Met | substitution | 315 | Leu | Met | 0.731 | 0.006 |
| p.Leu315Phe | substitution | 315 | Leu | Phe | 1.254 | 0.002 |
| p.Leu315Pro | substitution | 315 | Leu | Pro | 0.754 | 0.245 |
| p.Leu315Ser | substitution | 315 | Leu | Ser | 1.229 | 0.117 |
| p.Leu315Ter | nonsense | 315 | Leu | Ter | 0.219 | 0.134 |
| p.Leu315Thr | substitution | 315 | Leu | Thr | 0.029 | 0.191 |
| p.Leu315Trp | substitution | 315 | Leu | Trp | 0.813 | 0.005 |
| p.Leu315Tyr | substitution | 315 | Leu | Tyr | 0.000 | 0.035 |
| p.Leu315Val | substitution | 315 | Leu | Val | 0.079 | 0.127 |
| p.Leu329= | synonymous | 329 | Leu | NA | 1.310 | 0.080 |
| p.Leu329Arg | substitution | 329 | Leu | Arg | 1.383 | 0.026 |

| | | | | | | | |
|---|---|---|---|---|---|---|---|
| p.Leu329Asp | substitution | 329 | Leu | Asp | | 0.753 | 0.086 |
| p.Leu329Cys | substitution | 329 | Leu | Cys | | 0.000 | 0.080 |
| p.Leu329Gln | substitution | 329 | Leu | Gln | | 1.218 | 0.007 |
| p.Leu329Glu | substitution | 329 | Leu | Glu | | 0.000 | 0.016 |
| p.Leu329Ile | substitution | 329 | Leu | Ile | | 0.987 | 0.189 |
| p.Leu329Met | substitution | 329 | Leu | Met | | 1.493 | 0.038 |
| p.Leu329Phe | substitution | 329 | Leu | Phe | | 0.891 | 0.002 |
| p.Leu329Pro | substitution | 329 | Leu | Pro | | 0.000 | 0.170 |
| p.Leu329Ser | substitution | 329 | Leu | Ser | | 1.011 | 0.022 |
| p.Leu329Ter | nonsense | 329 | Leu | Ter | | 0.934 | 0.305 |
| p.Leu329Thr | substitution | 329 | Leu | Thr | | 1.232 | 0.324 |
| p.Leu329Trp | substitution | 329 | Leu | Trp | | 0.243 | 0.028 |
| p.Leu329Tyr | substitution | 329 | Leu | Tyr | | 0.348 | 0.016 |
| p.Leu329Val | substitution | 329 | Leu | Val | | 1.365 | 0.025 |
| p.Leu334= | synonymous | 334 | Leu | NA | | 1.439 | 0.007 |
| p.Leu334Arg | substitution | 334 | Leu | Arg | | 0.000 | 0.033 |
| p.Leu334Asn | substitution | 334 | Leu | Asn | | 1.903 | 0.010 |
| p.Leu334Cys | substitution | 334 | Leu | Cys | | 1.071 | 0.004 |
| p.Leu334Gln | substitution | 334 | Leu | Gln | | 0.000 | 0.019 |
| p.Leu334Gly | substitution | 334 | Leu | Gly | | 1.296 | 0.011 |
| p.Leu334His | substitution | 334 | Leu | His | | 0.000 | 0.019 |
| p.Leu334Lys | substitution | 334 | Leu | Lys | | 0.000 | 0.016 |
| p.Leu334Met | substitution | 334 | Leu | Met | | 1.058 | 0.005 |
| p.Leu334Phe | substitution | 334 | Leu | Phe | | 0.821 | 0.036 |
| p.Leu334Ser | substitution | 334 | Leu | Ser | | 0.509 | 0.225 |
| p.Leu334Ter | nonsense | 334 | Leu | Ter | | 0.535 | 0.014 |
| p.Leu334Thr | substitution | 334 | Leu | Thr | | 0.000 | 0.143 |
| p.Leu334Trp | substitution | 334 | Leu | Trp | | 0.604 | 0.013 |
| p.Leu334Tyr | substitution | 334 | Leu | Tyr | | 0.340 | 0.020 |
| p.Leu334Val | substitution | 334 | Leu | Val | | 1.547 | 0.014 |
| p.Leu338= | synonymous | 338 | Leu | NA | | 1.025 | 0.024 |
| p.Leu338Ala | substitution | 338 | Leu | Ala | | 1.212 | 0.005 |
| p.Leu338Arg | substitution | 338 | Leu | Arg | | 0.100 | 0.081 |
| p.Leu338Asp | substitution | 338 | Leu | Asp | | 0.000 | 0.104 |
| p.Leu338Cys | substitution | 338 | Leu | Cys | | 0.632 | 0.011 |
| p.Leu338Gln | substitution | 338 | Leu | Gln | | 0.129 | 0.233 |
| p.Leu338Gly | substitution | 338 | Leu | Gly | | 0.000 | 0.022 |
| p.Leu338His | substitution | 338 | Leu | His | | 0.000 | 0.035 |
| p.Leu338Ile | substitution | 338 | Leu | Ile | | 0.052 | 0.101 |
| p.Leu338Lys | substitution | 338 | Leu | Lys | | 0.374 | 0.148 |
| p.Leu338Met | substitution | 338 | Leu | Met | | 0.978 | 0.005 |
| p.Leu338Phe | substitution | 338 | Leu | Phe | | 0.000 | 0.117 |
| p.Leu338Pro | substitution | 338 | Leu | Pro | | 0.770 | 0.159 |

| | | | | | | | |
|---|---|---|---|---|---|---|---|
| p.Leu338Ser | substitution | 338 | Leu | Ser | | 0.918 | 0.003 |
| p.Leu338Ter | nonsense | 338 | Leu | Ter | | 0.000 | 0.134 |
| p.Leu338Thr | substitution | 338 | Leu | Thr | | 0.726 | 0.012 |
| p.Leu338Trp | substitution | 338 | Leu | Trp | | 0.000 | 0.084 |
| p.Leu338Tyr | substitution | 338 | Leu | Tyr | | 0.000 | 0.161 |
| p.Leu338Val | substitution | 338 | Leu | Val | | 0.000 | 0.142 |
| p.Leu341= | synonymous | 341 | Leu | NA | | 0.797 | 0.035 |
| p.Leu341Ala | substitution | 341 | Leu | Ala | | 0.000 | 0.266 |
| p.Leu341Arg | substitution | 341 | Leu | Arg | | 0.603 | 0.048 |
| p.Leu341Asp | substitution | 341 | Leu | Asp | | 1.203 | 0.004 |
| p.Leu341Cys | substitution | 341 | Leu | Cys | | 0.529 | 0.003 |
| p.Leu341Glu | substitution | 341 | Leu | Glu | | 0.873 | 0.008 |
| p.Leu341Gly | substitution | 341 | Leu | Gly | | 0.674 | 0.061 |
| p.Leu341His | substitution | 341 | Leu | His | | 0.427 | 0.008 |
| p.Leu341Ile | substitution | 341 | Leu | Ile | | 0.000 | 0.104 |
| p.Leu341Lys | substitution | 341 | Leu | Lys | | 0.967 | 0.003 |
| p.Leu341Met | substitution | 341 | Leu | Met | | 0.850 | 0.008 |
| p.Leu341Phe | substitution | 341 | Leu | Phe | | 0.523 | 0.014 |
| p.Leu341Pro | substitution | 341 | Leu | Pro | | 0.135 | 0.055 |
| p.Leu341Ser | substitution | 341 | Leu | Ser | | 0.352 | 0.129 |
| p.Leu341Ter | nonsense | 341 | Leu | Ter | | 0.338 | 0.020 |
| p.Leu341Thr | substitution | 341 | Leu | Thr | | 1.096 | 0.009 |
| p.Leu341Trp | substitution | 341 | Leu | Trp | | 0.652 | 0.003 |
| p.Leu341Tyr | substitution | 341 | Leu | Tyr | | 0.570 | 0.023 |
| p.Leu341Val | substitution | 341 | Leu | Val | | 0.832 | 0.085 |
| p.Leu342= | synonymous | 342 | Leu | NA | | 1.069 | 0.008 |
| p.Leu342Arg | substitution | 342 | Leu | Arg | | 0.000 | 0.007 |
| p.Leu342Asp | substitution | 342 | Leu | Asp | | 0.000 | 0.007 |
| p.Leu342Cys | substitution | 342 | Leu | Cys | | 0.307 | 0.003 |
| p.Leu342Gln | substitution | 342 | Leu | Gln | | 0.641 | 0.007 |
| p.Leu342Glu | substitution | 342 | Leu | Glu | | 0.003 | 0.007 |
| p.Leu342Gly | substitution | 342 | Leu | Gly | | 0.375 | 0.009 |
| p.Leu342His | substitution | 342 | Leu | His | | 1.301 | 0.005 |
| p.Leu342Ile | substitution | 342 | Leu | Ile | | 0.495 | 0.005 |
| p.Leu342Phe | substitution | 342 | Leu | Phe | | 1.334 | 0.164 |
| p.Leu342Pro | substitution | 342 | Leu | Pro | | 0.000 | 0.020 |
| p.Leu342Ser | substitution | 342 | Leu | Ser | | 0.630 | 0.013 |
| p.Leu342Ter | nonsense | 342 | Leu | Ter | | 0.174 | 0.009 |
| p.Leu342Thr | substitution | 342 | Leu | Thr | | 0.597 | 0.016 |
| p.Leu342Trp | substitution | 342 | Leu | Trp | | 0.037 | 0.005 |
| p.Leu342Tyr | substitution | 342 | Leu | Tyr | | 1.014 | 0.002 |
| p.Leu342Val | substitution | 342 | Leu | Val | | 1.446 | 0.019 |
| p.Leu343= | synonymous | 343 | Leu | NA | | 1.065 | 0.014 |

| | | | | | | |
|---|---|---|---|---|---|---|
| p.Leu343Ala | substitution | 343 | Leu | Ala | 0.659 | 0.098 |
| p.Leu343Arg | substitution | 343 | Leu | Arg | 1.102 | 0.010 |
| p.Leu343Asn | substitution | 343 | Leu | Asn | 1.219 | 0.002 |
| p.Leu343Asp | substitution | 343 | Leu | Asp | 1.181 | 0.004 |
| p.Leu343Cys | substitution | 343 | Leu | Cys | 0.460 | 0.005 |
| p.Leu343Gln | substitution | 343 | Leu | Gln | 0.520 | 0.008 |
| p.Leu343Glu | substitution | 343 | Leu | Glu | 0.562 | 0.007 |
| p.Leu343Gly | substitution | 343 | Leu | Gly | 0.786 | 0.005 |
| p.Leu343His | substitution | 343 | Leu | His | 1.079 | 0.004 |
| p.Leu343Ile | substitution | 343 | Leu | Ile | 0.000 | 0.011 |
| p.Leu343Lys | substitution | 343 | Leu | Lys | 0.303 | 0.014 |
| p.Leu343Met | substitution | 343 | Leu | Met | 1.218 | 0.005 |
| p.Leu343Phe | substitution | 343 | Leu | Phe | 0.828 | 0.003 |
| p.Leu343Pro | substitution | 343 | Leu | Pro | 0.300 | 0.134 |
| p.Leu343Ser | substitution | 343 | Leu | Ser | 0.449 | 0.043 |
| p.Leu343Ter | nonsense | 343 | Leu | Ter | 0.000 | 0.108 |
| p.Leu343Thr | substitution | 343 | Leu | Thr | 0.762 | 0.246 |
| p.Leu343Trp | substitution | 343 | Leu | Trp | 1.373 | 0.004 |
| p.Leu343Tyr | substitution | 343 | Leu | Tyr | 0.000 | 0.030 |
| p.Leu343Val | substitution | 343 | Leu | Val | 1.369 | 0.011 |
| p.Leu351= | synonymous | 351 | Leu | NA | 0.243 | 0.065 |
| p.Leu351Ala | substitution | 351 | Leu | Ala | 0.000 | 0.149 |
| p.Leu351Arg | substitution | 351 | Leu | Arg | 0.513 | 0.052 |
| p.Leu351Asn | substitution | 351 | Leu | Asn | 0.000 | 0.252 |
| p.Leu351Asp | substitution | 351 | Leu | Asp | 1.099 | 0.013 |
| p.Leu351Cys | substitution | 351 | Leu | Cys | 1.418 | 0.004 |
| p.Leu351Gln | substitution | 351 | Leu | Gln | 1.294 | 0.000 |
| p.Leu351Glu | substitution | 351 | Leu | Glu | 1.125 | 0.003 |
| p.Leu351Gly | substitution | 351 | Leu | Gly | 0.000 | 0.150 |
| p.Leu351His | substitution | 351 | Leu | His | 0.884 | 0.016 |
| p.Leu351Ile | substitution | 351 | Leu | Ile | 0.671 | 0.009 |
| p.Leu351Lys | substitution | 351 | Leu | Lys | 0.000 | 0.051 |
| p.Leu351Met | substitution | 351 | Leu | Met | 1.136 | 0.004 |
| p.Leu351Pro | substitution | 351 | Leu | Pro | 0.746 | 0.016 |
| p.Leu351Ser | substitution | 351 | Leu | Ser | 0.519 | 0.230 |
| p.Leu351Ter | nonsense | 351 | Leu | Ter | 0.063 | 0.021 |
| p.Leu351Thr | substitution | 351 | Leu | Thr | 1.019 | 0.140 |
| p.Leu351Trp | substitution | 351 | Leu | Trp | 0.340 | 0.008 |
| p.Leu351Tyr | substitution | 351 | Leu | Tyr | 0.913 | 0.269 |
| p.Leu351Val | substitution | 351 | Leu | Val | 0.349 | 0.042 |
| p.Leu357= | synonymous | 357 | Leu | NA | 1.267 | 0.297 |
| p.Leu357Arg | substitution | 357 | Leu | Arg | 1.086 | 0.262 |
| p.Leu357Cys | substitution | 357 | Leu | Cys | 0.000 | 0.097 |

| | | | | | | | |
|---|---|---|---|---|---|---|---|
| p.Leu357His | substitution | 357 | Leu | His | | 1.485 | 0.036 |
| p.Leu357Ile | substitution | 357 | Leu | Ile | | 1.112 | 0.110 |
| p.Leu357Lys | substitution | 357 | Leu | Lys | | 0.820 | 0.016 |
| p.Leu357Phe | substitution | 357 | Leu | Phe | | 1.160 | 0.005 |
| p.Leu357Ser | substitution | 357 | Leu | Ser | | 1.376 | 0.029 |
| p.Leu357Trp | substitution | 357 | Leu | Trp | | 0.000 | 0.216 |
| p.Leu357Tyr | substitution | 357 | Leu | Tyr | | 0.865 | 0.004 |
| p.Leu357Val | substitution | 357 | Leu | Val | | 0.796 | 0.122 |
| p.Leu42= | synonymous | 42 | Leu | NA | | 0.902 | 0.004 |
| p.Leu42Ala | substitution | 42 | Leu | Ala | | 0.448 | 0.007 |
| p.Leu42Arg | substitution | 42 | Leu | Arg | | 0.000 | 0.009 |
| p.Leu42Asn | substitution | 42 | Leu | Asn | | 0.000 | 0.019 |
| p.Leu42Asp | substitution | 42 | Leu | Asp | | 0.000 | 0.009 |
| p.Leu42Cys | substitution | 42 | Leu | Cys | | 0.611 | 0.003 |
| p.Leu42Gln | substitution | 42 | Leu | Gln | | 0.000 | 0.022 |
| p.Leu42Glu | substitution | 42 | Leu | Glu | | 0.038 | 0.012 |
| p.Leu42Gly | substitution | 42 | Leu | Gly | | 0.000 | 0.006 |
| p.Leu42His | substitution | 42 | Leu | His | | 0.000 | 0.015 |
| p.Leu42Ile | substitution | 42 | Leu | Ile | | 0.776 | 0.004 |
| p.Leu42Lys | substitution | 42 | Leu | Lys | | 0.000 | 0.012 |
| p.Leu42Met | substitution | 42 | Leu | Met | | 0.677 | 0.004 |
| p.Leu42Phe | substitution | 42 | Leu | Phe | | 0.708 | 0.003 |
| p.Leu42Pro | substitution | 42 | Leu | Pro | | 0.000 | 0.008 |
| p.Leu42Ser | substitution | 42 | Leu | Ser | | 0.000 | 0.007 |
| p.Leu42Ter | nonsense | 42 | Leu | Ter | | 0.109 | 0.014 |
| p.Leu42Thr | substitution | 42 | Leu | Thr | | 0.221 | 0.008 |
| p.Leu42Tyr | substitution | 42 | Leu | Tyr | | 0.182 | 0.005 |
| p.Leu42Val | substitution | 42 | Leu | Val | | 0.523 | 0.004 |
| p.Leu49= | synonymous | 49 | Leu | NA | | 1.065 | 0.006 |
| p.Leu49Ala | substitution | 49 | Leu | Ala | | 1.110 | 0.023 |
| p.Leu49Arg | substitution | 49 | Leu | Arg | | 0.515 | 0.085 |
| p.Leu49Asn | substitution | 49 | Leu | Asn | | 0.060 | 0.669 |
| p.Leu49Asp | substitution | 49 | Leu | Asp | | 0.642 | 0.137 |
| p.Leu49Cys | substitution | 49 | Leu | Cys | | 0.000 | 0.278 |
| p.Leu49Gln | substitution | 49 | Leu | Gln | | 1.019 | 0.013 |
| p.Leu49Glu | substitution | 49 | Leu | Glu | | 0.000 | 0.789 |
| p.Leu49Gly | substitution | 49 | Leu | Gly | | 0.926 | 0.146 |
| p.Leu49His | substitution | 49 | Leu | His | | 0.833 | 0.012 |
| p.Leu49Ile | substitution | 49 | Leu | Ile | | 0.524 | 0.051 |
| p.Leu49Lys | substitution | 49 | Leu | Lys | | 0.581 | 0.021 |
| p.Leu49Met | substitution | 49 | Leu | Met | | 0.858 | 0.008 |
| p.Leu49Phe | substitution | 49 | Leu | Phe | | 0.232 | 0.119 |
| p.Leu49Pro | substitution | 49 | Leu | Pro | | 0.896 | 0.077 |

| | | | | | | | |
|---|---|---|---|---|---|---|---|
| p.Leu49Ser | substitution | 49 | Leu | Ser | | 0.906 | 0.014 |
| p.Leu49Ter | nonsense | 49 | Leu | Ter | | 0.210 | 0.769 |
| p.Leu49Thr | substitution | 49 | Leu | Thr | | 1.007 | 0.034 |
| p.Leu49Trp | substitution | 49 | Leu | Trp | | 0.618 | 0.010 |
| p.Leu49Tyr | substitution | 49 | Leu | Tyr | | 1.683 | 0.033 |
| p.Leu49Val | substitution | 49 | Leu | Val | | 1.046 | 0.033 |
| p.Leu64= | synonymous | 64 | Leu | NA | | 0.900 | 0.099 |
| p.Leu64Ala | substitution | 64 | Leu | Ala | | 1.755 | 0.011 |
| p.Leu64Arg | substitution | 64 | Leu | Arg | | 0.730 | 0.079 |
| p.Leu64Asp | substitution | 64 | Leu | Asp | | 0.641 | 0.016 |
| p.Leu64Cys | substitution | 64 | Leu | Cys | | 0.964 | 0.029 |
| p.Leu64Gln | substitution | 64 | Leu | Gln | | 0.656 | 0.013 |
| p.Leu64Glu | substitution | 64 | Leu | Glu | | 0.000 | 0.160 |
| p.Leu64Gly | substitution | 64 | Leu | Gly | | 1.092 | 0.094 |
| p.Leu64His | substitution | 64 | Leu | His | | 1.300 | 0.012 |
| p.Leu64Ile | substitution | 64 | Leu | Ile | | 0.858 | 0.009 |
| p.Leu64Lys | substitution | 64 | Leu | Lys | | 0.996 | 0.019 |
| p.Leu64Met | substitution | 64 | Leu | Met | | 0.687 | 0.013 |
| p.Leu64Phe | substitution | 64 | Leu | Phe | | 0.759 | 0.006 |
| p.Leu64Pro | substitution | 64 | Leu | Pro | | 0.977 | 0.038 |
| p.Leu64Ser | substitution | 64 | Leu | Ser | | 0.479 | 0.063 |
| p.Leu64Ter | nonsense | 64 | Leu | Ter | | 0.000 | 0.524 |
| p.Leu64Thr | substitution | 64 | Leu | Thr | | 0.933 | 0.218 |
| p.Leu64Trp | substitution | 64 | Leu | Trp | | 1.289 | 0.012 |
| p.Leu64Tyr | substitution | 64 | Leu | Tyr | | 1.104 | 0.009 |
| p.Leu64Val | substitution | 64 | Leu | Val | | 0.315 | 0.099 |
| p.Leu68= | synonymous | 68 | Leu | NA | | 0.897 | 0.006 |
| p.Leu68Ala | substitution | 68 | Leu | Ala | | 0.846 | 0.150 |
| p.Leu68Arg | substitution | 68 | Leu | Arg | | 0.926 | 0.039 |
| p.Leu68Asn | substitution | 68 | Leu | Asn | | 0.419 | 0.012 |
| p.Leu68Asp | substitution | 68 | Leu | Asp | | 0.717 | 0.013 |
| p.Leu68Cys | substitution | 68 | Leu | Cys | | 0.762 | 0.015 |
| p.Leu68Gln | substitution | 68 | Leu | Gln | | 0.392 | 0.016 |
| p.Leu68Glu | substitution | 68 | Leu | Glu | | 0.552 | 0.017 |
| p.Leu68Gly | substitution | 68 | Leu | Gly | | 0.487 | 0.021 |
| p.Leu68His | substitution | 68 | Leu | His | | 0.265 | 0.081 |
| p.Leu68Ile | substitution | 68 | Leu | Ile | | 1.013 | 0.026 |
| p.Leu68Lys | substitution | 68 | Leu | Lys | | 1.039 | 0.007 |
| p.Leu68Met | substitution | 68 | Leu | Met | | 1.089 | 0.010 |
| p.Leu68Phe | substitution | 68 | Leu | Phe | | 0.374 | 0.038 |
| p.Leu68Pro | substitution | 68 | Leu | Pro | | 0.667 | 0.008 |
| p.Leu68Ser | substitution | 68 | Leu | Ser | | 0.788 | 0.013 |
| p.Leu68Ter | nonsense | 68 | Leu | Ter | | 0.305 | 0.018 |

| | | | | | | |
|---|---|---|---|---|---|---|
| p.Leu68Thr | substitution | 68 | Leu | Thr | 0.375 | 0.019 |
| p.Leu68Trp | substitution | 68 | Leu | Trp | 0.588 | 0.016 |
| p.Leu68Tyr | substitution | 68 | Leu | Tyr | 0.852 | 0.012 |
| p.Leu68Val | substitution | 68 | Leu | Val | 0.913 | 0.029 |
| p.Leu76= | synonymous | 76 | Leu | NA | 1.055 | 0.018 |
| p.Leu76Ala | substitution | 76 | Leu | Ala | 0.426 | 0.014 |
| p.Leu76Arg | substitution | 76 | Leu | Arg | 0.904 | 0.007 |
| p.Leu76Asn | substitution | 76 | Leu | Asn | 0.926 | 0.011 |
| p.Leu76Asp | substitution | 76 | Leu | Asp | 0.916 | 0.010 |
| p.Leu76Cys | substitution | 76 | Leu | Cys | 1.159 | 0.005 |
| p.Leu76Gln | substitution | 76 | Leu | Gln | 0.808 | 0.006 |
| p.Leu76Glu | substitution | 76 | Leu | Glu | 0.351 | 0.009 |
| p.Leu76Gly | substitution | 76 | Leu | Gly | 0.771 | 0.069 |
| p.Leu76Ile | substitution | 76 | Leu | Ile | 0.432 | 0.022 |
| p.Leu76Lys | substitution | 76 | Leu | Lys | 1.112 | 0.004 |
| p.Leu76Met | substitution | 76 | Leu | Met | 0.984 | 0.003 |
| p.Leu76Phe | substitution | 76 | Leu | Phe | 0.544 | 0.007 |
| p.Leu76Pro | substitution | 76 | Leu | Pro | 0.777 | 0.009 |
| p.Leu76Ser | substitution | 76 | Leu | Ser | 0.850 | 0.047 |
| p.Leu76Ter | nonsense | 76 | Leu | Ter | 0.000 | 0.014 |
| p.Leu76Thr | substitution | 76 | Leu | Thr | 0.825 | 0.018 |
| p.Leu76Trp | substitution | 76 | Leu | Trp | 0.775 | 0.003 |
| p.Leu76Tyr | substitution | 76 | Leu | Tyr | 0.567 | 0.015 |
| p.Leu76Val | substitution | 76 | Leu | Val | 1.048 | 0.007 |
| p.Leu81= | synonymous | 81 | Leu | NA | 0.913 | 0.092 |
| p.Leu81Ala | substitution | 81 | Leu | Ala | 0.170 | 0.018 |
| p.Leu81Arg | substitution | 81 | Leu | Arg | 0.000 | 0.032 |
| p.Leu81Asn | substitution | 81 | Leu | Asn | 0.000 | 0.032 |
| p.Leu81Asp | substitution | 81 | Leu | Asp | 0.108 | 0.013 |
| p.Leu81Cys | substitution | 81 | Leu | Cys | 0.665 | 0.008 |
| p.Leu81Gln | substitution | 81 | Leu | Gln | 0.000 | 0.018 |
| p.Leu81Glu | substitution | 81 | Leu | Glu | 0.035 | 0.017 |
| p.Leu81Gly | substitution | 81 | Leu | Gly | 0.592 | 0.011 |
| p.Leu81His | substitution | 81 | Leu | His | 0.854 | 0.011 |
| p.Leu81Ile | substitution | 81 | Leu | Ile | 0.949 | 0.003 |
| p.Leu81Lys | substitution | 81 | Leu | Lys | 0.000 | 0.012 |
| p.Leu81Met | substitution | 81 | Leu | Met | 0.618 | 0.004 |
| p.Leu81Phe | substitution | 81 | Leu | Phe | 0.987 | 0.002 |
| p.Leu81Pro | substitution | 81 | Leu | Pro | 0.519 | 0.021 |
| p.Leu81Ser | substitution | 81 | Leu | Ser | 0.039 | 0.013 |
| p.Leu81Ter | nonsense | 81 | Leu | Ter | 0.000 | 0.012 |
| p.Leu81Thr | substitution | 81 | Leu | Thr | 0.778 | 0.013 |
| p.Leu81Trp | substitution | 81 | Leu | Trp | 0.724 | 0.006 |

| | | | | | | | |
|---|---|---|---|---|---|---|---|
| p.Leu81Tyr | substitution | 81 | Leu | Tyr | | 0.100 | 0.020 |
| p.Leu81Val | substitution | 81 | Leu | Val | | 0.655 | 0.007 |
| p.Leu85= | synonymous | 85 | Leu | NA | | 1.125 | 0.010 |
| p.Leu85Ala | substitution | 85 | Leu | Ala | | 0.486 | 0.031 |
| p.Leu85Arg | substitution | 85 | Leu | Arg | | 0.000 | 0.015 |
| p.Leu85Asn | substitution | 85 | Leu | Asn | | 0.000 | 0.043 |
| p.Leu85Asp | substitution | 85 | Leu | Asp | | 0.000 | 0.037 |
| p.Leu85Cys | substitution | 85 | Leu | Cys | | 0.864 | 0.004 |
| p.Leu85Gln | substitution | 85 | Leu | Gln | | 0.436 | 0.005 |
| p.Leu85Glu | substitution | 85 | Leu | Glu | | 0.021 | 0.007 |
| p.Leu85Gly | substitution | 85 | Leu | Gly | | 0.000 | 0.019 |
| p.Leu85His | substitution | 85 | Leu | His | | 0.000 | 0.011 |
| p.Leu85Ile | substitution | 85 | Leu | Ile | | 0.556 | 0.011 |
| p.Leu85Lys | substitution | 85 | Leu | Lys | | 0.066 | 0.024 |
| p.Leu85Met | substitution | 85 | Leu | Met | | 0.850 | 0.003 |
| p.Leu85Phe | substitution | 85 | Leu | Phe | | 0.182 | 0.006 |
| p.Leu85Pro | substitution | 85 | Leu | Pro | | 0.000 | 0.062 |
| p.Leu85Ser | substitution | 85 | Leu | Ser | | 0.000 | 0.006 |
| p.Leu85Ter | nonsense | 85 | Leu | Ter | | 0.279 | 0.016 |
| p.Leu85Thr | substitution | 85 | Leu | Thr | | 0.395 | 0.014 |
| p.Leu85Trp | substitution | 85 | Leu | Trp | | 0.000 | 0.011 |
| p.Leu85Tyr | substitution | 85 | Leu | Tyr | | 0.000 | 0.023 |
| p.Leu85Val | substitution | 85 | Leu | Val | | 0.780 | 0.003 |
| p.Leu92= | synonymous | 92 | Leu | NA | | 1.039 | 0.008 |
| p.Leu92Ala | substitution | 92 | Leu | Ala | | 0.791 | 0.009 |
| p.Leu92Arg | substitution | 92 | Leu | Arg | | 0.000 | 0.015 |
| p.Leu92Asn | substitution | 92 | Leu | Asn | | 0.983 | 0.060 |
| p.Leu92Asp | substitution | 92 | Leu | Asp | | 0.000 | 0.067 |
| p.Leu92Cys | substitution | 92 | Leu | Cys | | 0.988 | 0.012 |
| p.Leu92Gln | substitution | 92 | Leu | Gln | | 0.133 | 0.062 |
| p.Leu92Glu | substitution | 92 | Leu | Glu | | 0.000 | 0.064 |
| p.Leu92Gly | substitution | 92 | Leu | Gly | | 0.430 | 0.015 |
| p.Leu92His | substitution | 92 | Leu | His | | 0.272 | 0.008 |
| p.Leu92Ile | substitution | 92 | Leu | Ile | | 0.932 | 0.007 |
| p.Leu92Lys | substitution | 92 | Leu | Lys | | 0.025 | 0.015 |
| p.Leu92Met | substitution | 92 | Leu | Met | | 0.948 | 0.004 |
| p.Leu92Phe | substitution | 92 | Leu | Phe | | 0.794 | 0.008 |
| p.Leu92Pro | substitution | 92 | Leu | Pro | | 0.000 | 0.144 |
| p.Leu92Ser | substitution | 92 | Leu | Ser | | 0.215 | 0.006 |
| p.Leu92Ter | nonsense | 92 | Leu | Ter | | 0.330 | 0.016 |
| p.Leu92Thr | substitution | 92 | Leu | Thr | | 0.600 | 0.108 |
| p.Leu92Trp | substitution | 92 | Leu | Trp | | 0.000 | 0.039 |
| p.Leu92Tyr | substitution | 92 | Leu | Tyr | | 0.000 | 0.015 |

| | | | | | | | |
|---|---|---|---|---|---|---|---|
| p.Leu92Val | substitution | 92 | Leu | Val | | 0.516 | 0.155 |
| p.Leu97= | synonymous | 97 | Leu | NA | | 1.077 | 0.009 |
| p.Leu97Ala | substitution | 97 | Leu | Ala | | 0.834 | 0.005 |
| p.Leu97Arg | substitution | 97 | Leu | Arg | | 0.000 | 0.024 |
| p.Leu97Asn | substitution | 97 | Leu | Asn | | 0.190 | 0.008 |
| p.Leu97Asp | substitution | 97 | Leu | Asp | | 0.000 | 0.012 |
| p.Leu97Cys | substitution | 97 | Leu | Cys | | 0.527 | 0.015 |
| p.Leu97Gln | substitution | 97 | Leu | Gln | | 0.000 | 0.014 |
| p.Leu97Glu | substitution | 97 | Leu | Glu | | 0.133 | 0.010 |
| p.Leu97Gly | substitution | 97 | Leu | Gly | | 0.000 | 0.009 |
| p.Leu97His | substitution | 97 | Leu | His | | 0.000 | 0.015 |
| p.Leu97Ile | substitution | 97 | Leu | Ile | | 0.635 | 0.005 |
| p.Leu97Lys | substitution | 97 | Leu | Lys | | 0.183 | 0.009 |
| p.Leu97Met | substitution | 97 | Leu | Met | | 0.647 | 0.012 |
| p.Leu97Phe | substitution | 97 | Leu | Phe | | 0.796 | 0.006 |
| p.Leu97Pro | substitution | 97 | Leu | Pro | | 0.000 | 0.028 |
| p.Leu97Ser | substitution | 97 | Leu | Ser | | 0.353 | 0.048 |
| p.Leu97Ter | nonsense | 97 | Leu | Ter | | 0.383 | 0.043 |
| p.Leu97Thr | substitution | 97 | Leu | Thr | | 1.290 | 0.007 |
| p.Leu97Trp | substitution | 97 | Leu | Trp | | 0.000 | 0.025 |
| p.Leu97Tyr | substitution | 97 | Leu | Tyr | | 0.000 | 0.017 |
| p.Leu97Val | substitution | 97 | Leu | Val | | 0.972 | 0.014 |
| p.Lys115= | synonymous | 115 | Lys | NA | | 1.084 | 0.023 |
| p.Lys115Ala | substitution | 115 | Lys | Ala | | 0.523 | 0.021 |
| p.Lys115Arg | substitution | 115 | Lys | Arg | | 0.851 | 0.013 |
| p.Lys115Asp | substitution | 115 | Lys | Asp | | 1.068 | 0.008 |
| p.Lys115Cys | substitution | 115 | Lys | Cys | | 0.631 | 0.012 |
| p.Lys115Gln | substitution | 115 | Lys | Gln | | 0.882 | 0.004 |
| p.Lys115Glu | substitution | 115 | Lys | Glu | | 0.836 | 0.022 |
| p.Lys115Gly | substitution | 115 | Lys | Gly | | 1.030 | 0.021 |
| p.Lys115His | substitution | 115 | Lys | His | | 0.654 | 0.005 |
| p.Lys115Ile | substitution | 115 | Lys | Ile | | 0.673 | 0.008 |
| p.Lys115Leu | substitution | 115 | Lys | Leu | | 0.931 | 0.024 |
| p.Lys115Phe | substitution | 115 | Lys | Phe | | 0.850 | 0.005 |
| p.Lys115Pro | substitution | 115 | Lys | Pro | | 1.019 | 0.010 |
| p.Lys115Ser | substitution | 115 | Lys | Ser | | 1.101 | 0.032 |
| p.Lys115Ter | nonsense | 115 | Lys | Ter | | 0.069 | 0.012 |
| p.Lys115Thr | substitution | 115 | Lys | Thr | | 0.957 | 0.003 |
| p.Lys115Trp | substitution | 115 | Lys | Trp | | 0.368 | 0.007 |
| p.Lys115Tyr | substitution | 115 | Lys | Tyr | | 0.446 | 0.006 |
| p.Lys115Val | substitution | 115 | Lys | Val | | 0.748 | 0.006 |
| p.Lys128= | synonymous | 128 | Lys | NA | | 1.189 | 0.017 |
| p.Lys128Ala | substitution | 128 | Lys | Ala | | 0.901 | 0.091 |

| | | | | | | | |
|---|---|---|---|---|---|---|---|
| p.Lys128Arg | substitution | 128 | Lys | Arg | | 0.642 | 0.027 |
| p.Lys128Asn | substitution | 128 | Lys | Asn | | 1.104 | 0.049 |
| p.Lys128Asp | substitution | 128 | Lys | Asp | | 0.657 | 0.010 |
| p.Lys128Cys | substitution | 128 | Lys | Cys | | 0.701 | 0.009 |
| p.Lys128Gln | substitution | 128 | Lys | Gln | | 0.377 | 0.030 |
| p.Lys128Glu | substitution | 128 | Lys | Glu | | 1.412 | 0.013 |
| p.Lys128Gly | substitution | 128 | Lys | Gly | | 0.383 | 0.093 |
| p.Lys128His | substitution | 128 | Lys | His | | 0.903 | 0.033 |
| p.Lys128Ile | substitution | 128 | Lys | Ile | | 0.939 | 0.014 |
| p.Lys128Leu | substitution | 128 | Lys | Leu | | 0.902 | 0.041 |
| p.Lys128Met | substitution | 128 | Lys | Met | | 1.244 | 0.006 |
| p.Lys128Phe | substitution | 128 | Lys | Phe | | 0.513 | 0.030 |
| p.Lys128Pro | substitution | 128 | Lys | Pro | | 0.000 | 0.102 |
| p.Lys128Ser | substitution | 128 | Lys | Ser | | 0.757 | 0.025 |
| p.Lys128Ter | nonsense | 128 | Lys | Ter | | 0.467 | 0.152 |
| p.Lys128Thr | substitution | 128 | Lys | Thr | | 1.218 | 0.200 |
| p.Lys128Trp | substitution | 128 | Lys | Trp | | 0.858 | 0.012 |
| p.Lys128Tyr | substitution | 128 | Lys | Tyr | | 0.752 | 0.010 |
| p.Lys128Val | substitution | 128 | Lys | Val | | 0.515 | 0.019 |
| p.Lys132Ala | substitution | 132 | Lys | Ala | | 0.811 | 0.004 |
| p.Lys132Arg | substitution | 132 | Lys | Arg | | 0.899 | 0.002 |
| p.Lys132Asn | substitution | 132 | Lys | Asn | | 1.202 | 0.029 |
| p.Lys132Cys | substitution | 132 | Lys | Cys | | 0.879 | 0.004 |
| p.Lys132Gln | substitution | 132 | Lys | Gln | | 1.139 | 0.006 |
| p.Lys132Glu | substitution | 132 | Lys | Glu | | 0.924 | 0.004 |
| p.Lys132Gly | substitution | 132 | Lys | Gly | | 1.042 | 0.004 |
| p.Lys132His | substitution | 132 | Lys | His | | 0.811 | 0.005 |
| p.Lys132Ile | substitution | 132 | Lys | Ile | | 0.858 | 0.003 |
| p.Lys132Leu | substitution | 132 | Lys | Leu | | 0.812 | 0.006 |
| p.Lys132Met | substitution | 132 | Lys | Met | | 1.090 | 0.003 |
| p.Lys132Phe | substitution | 132 | Lys | Phe | | 1.118 | 0.005 |
| p.Lys132Pro | substitution | 132 | Lys | Pro | | 0.917 | 0.007 |
| p.Lys132Ser | substitution | 132 | Lys | Ser | | 0.966 | 0.002 |
| p.Lys132Ter | nonsense | 132 | Lys | Ter | | 0.375 | 0.007 |
| p.Lys132Thr | substitution | 132 | Lys | Thr | | 0.796 | 0.003 |
| p.Lys132Trp | substitution | 132 | Lys | Trp | | 0.990 | 0.003 |
| p.Lys132Tyr | substitution | 132 | Lys | Tyr | | 0.912 | 0.003 |
| p.Lys132Val | substitution | 132 | Lys | Val | | 0.739 | 0.007 |
| p.Lys140= | synonymous | 140 | Lys | NA | | 1.218 | 0.040 |
| p.Lys140Ala | substitution | 140 | Lys | Ala | | 0.909 | 0.008 |
| p.Lys140Arg | substitution | 140 | Lys | Arg | | 0.856 | 0.042 |
| p.Lys140Asn | substitution | 140 | Lys | Asn | | 0.900 | 0.037 |
| p.Lys140Asp | substitution | 140 | Lys | Asp | | 1.054 | 0.004 |

| | | | | | | |
|---|---|---|---|---|---|---|
| p.Lys140Cys | substitution | 140 | Lys | Cys | 0.814 | 0.011 |
| p.Lys140Glu | substitution | 140 | Lys | Glu | 1.083 | 0.003 |
| p.Lys140Gly | substitution | 140 | Lys | Gly | 0.903 | 0.017 |
| p.Lys140His | substitution | 140 | Lys | His | 0.606 | 0.006 |
| p.Lys140Ile | substitution | 140 | Lys | Ile | 0.592 | 0.004 |
| p.Lys140Leu | substitution | 140 | Lys | Leu | 0.871 | 0.005 |
| p.Lys140Met | substitution | 140 | Lys | Met | 0.816 | 0.003 |
| p.Lys140Phe | substitution | 140 | Lys | Phe | 0.787 | 0.006 |
| p.Lys140Pro | substitution | 140 | Lys | Pro | 0.930 | 0.045 |
| p.Lys140Ser | substitution | 140 | Lys | Ser | 0.996 | 0.002 |
| p.Lys140Ter | nonsense | 140 | Lys | Ter | 0.289 | 0.007 |
| p.Lys140Thr | substitution | 140 | Lys | Thr | 0.801 | 0.006 |
| p.Lys140Trp | substitution | 140 | Lys | Trp | 1.165 | 0.003 |
| p.Lys140Tyr | substitution | 140 | Lys | Tyr | 0.704 | 0.004 |
| p.Lys140Val | substitution | 140 | Lys | Val | 0.780 | 0.063 |
| p.Lys157= | synonymous | 157 | Lys | NA | 1.359 | 0.015 |
| p.Lys157Ala | substitution | 157 | Lys | Ala | 1.031 | 0.004 |
| p.Lys157Arg | substitution | 157 | Lys | Arg | 0.848 | 0.023 |
| p.Lys157Asn | substitution | 157 | Lys | Asn | 1.031 | 0.035 |
| p.Lys157Asp | substitution | 157 | Lys | Asp | 0.653 | 0.008 |
| p.Lys157Cys | substitution | 157 | Lys | Cys | 0.876 | 0.005 |
| p.Lys157Gln | substitution | 157 | Lys | Gln | 0.958 | 0.007 |
| p.Lys157Glu | substitution | 157 | Lys | Glu | 0.486 | 0.017 |
| p.Lys157Gly | substitution | 157 | Lys | Gly | 0.924 | 0.012 |
| p.Lys157His | substitution | 157 | Lys | His | 1.034 | 0.007 |
| p.Lys157Ile | substitution | 157 | Lys | Ile | 0.800 | 0.008 |
| p.Lys157Leu | substitution | 157 | Lys | Leu | 0.948 | 0.003 |
| p.Lys157Met | substitution | 157 | Lys | Met | 0.954 | 0.006 |
| p.Lys157Phe | substitution | 157 | Lys | Phe | 0.816 | 0.006 |
| p.Lys157Pro | substitution | 157 | Lys | Pro | 0.637 | 0.040 |
| p.Lys157Ser | substitution | 157 | Lys | Ser | 0.866 | 0.008 |
| p.Lys157Ter | nonsense | 157 | Lys | Ter | 0.291 | 0.011 |
| p.Lys157Thr | substitution | 157 | Lys | Thr | 0.542 | 0.059 |
| p.Lys157Trp | substitution | 157 | Lys | Trp | 0.303 | 0.007 |
| p.Lys157Tyr | substitution | 157 | Lys | Tyr | 0.943 | 0.005 |
| p.Lys157Val | substitution | 157 | Lys | Val | 0.827 | 0.005 |
| p.Lys176= | synonymous | 176 | Lys | NA | 1.446 | 0.027 |
| p.Lys176Ala | substitution | 176 | Lys | Ala | 0.681 | 0.073 |
| p.Lys176Arg | substitution | 176 | Lys | Arg | 0.892 | 0.008 |
| p.Lys176Asn | substitution | 176 | Lys | Asn | 0.604 | 0.010 |
| p.Lys176Asp | substitution | 176 | Lys | Asp | 0.717 | 0.005 |
| p.Lys176Gln | substitution | 176 | Lys | Gln | 1.057 | 0.008 |
| p.Lys176Glu | substitution | 176 | Lys | Glu | 0.693 | 0.019 |

| | | | | | | |
|---|---|---|---|---|---|---|
| p.Lys176Gly | substitution | 176 | Lys | Gly | 0.592 | 0.139 |
| p.Lys176His | substitution | 176 | Lys | His | 0.697 | 0.005 |
| p.Lys176Ile | substitution | 176 | Lys | Ile | 0.903 | 0.004 |
| p.Lys176Leu | substitution | 176 | Lys | Leu | 0.950 | 0.006 |
| p.Lys176Met | substitution | 176 | Lys | Met | 0.955 | 0.004 |
| p.Lys176Phe | substitution | 176 | Lys | Phe | 1.004 | 0.005 |
| p.Lys176Pro | substitution | 176 | Lys | Pro | 0.000 | 0.054 |
| p.Lys176Ser | substitution | 176 | Lys | Ser | 0.815 | 0.009 |
| p.Lys176Ter | nonsense | 176 | Lys | Ter | 0.404 | 0.013 |
| p.Lys176Thr | substitution | 176 | Lys | Thr | 0.836 | 0.004 |
| p.Lys176Trp | substitution | 176 | Lys | Trp | 0.800 | 0.007 |
| p.Lys176Tyr | substitution | 176 | Lys | Tyr | 0.794 | 0.003 |
| p.Lys176Val | substitution | 176 | Lys | Val | 0.882 | 0.004 |
| p.Lys227Ala | substitution | 227 | Lys | Ala | 1.364 | 0.004 |
| p.Lys227Arg | substitution | 227 | Lys | Arg | 0.611 | 0.024 |
| p.Lys227Asn | substitution | 227 | Lys | Asn | 0.489 | 0.006 |
| p.Lys227Asp | substitution | 227 | Lys | Asp | 1.214 | 0.003 |
| p.Lys227Cys | substitution | 227 | Lys | Cys | 0.000 | 0.160 |
| p.Lys227Gln | substitution | 227 | Lys | Gln | 0.721 | 0.003 |
| p.Lys227Glu | substitution | 227 | Lys | Glu | 1.736 | 0.001 |
| p.Lys227Gly | substitution | 227 | Lys | Gly | 0.147 | 0.008 |
| p.Lys227His | substitution | 227 | Lys | His | 0.264 | 0.010 |
| p.Lys227Ile | substitution | 227 | Lys | Ile | 0.328 | 0.022 |
| p.Lys227Leu | substitution | 227 | Lys | Leu | 1.193 | 0.007 |
| p.Lys227Met | substitution | 227 | Lys | Met | 0.424 | 0.004 |
| p.Lys227Phe | substitution | 227 | Lys | Phe | 0.025 | 0.011 |
| p.Lys227Pro | substitution | 227 | Lys | Pro | 0.000 | 0.214 |
| p.Lys227Ser | substitution | 227 | Lys | Ser | 0.743 | 0.015 |
| p.Lys227Ter | nonsense | 227 | Lys | Ter | 0.427 | 0.135 |
| p.Lys227Thr | substitution | 227 | Lys | Thr | 0.539 | 0.074 |
| p.Lys227Trp | substitution | 227 | Lys | Trp | 0.479 | 0.004 |
| p.Lys227Tyr | substitution | 227 | Lys | Tyr | 1.299 | 0.003 |
| p.Lys227Val | substitution | 227 | Lys | Val | 0.000 | 0.015 |
| p.Lys272= | synonymous | 272 | Lys | NA | 1.425 | 0.014 |
| p.Lys272Arg | substitution | 272 | Lys | Arg | 0.784 | 0.032 |
| p.Lys272Asn | substitution | 272 | Lys | Asn | 1.175 | 0.021 |
| p.Lys272Cys | substitution | 272 | Lys | Cys | 0.413 | 0.005 |
| p.Lys272Gln | substitution | 272 | Lys | Gln | 0.726 | 0.160 |
| p.Lys272Glu | substitution | 272 | Lys | Glu | 0.599 | 0.606 |
| p.Lys272Gly | substitution | 272 | Lys | Gly | 0.000 | 0.101 |
| p.Lys272Leu | substitution | 272 | Lys | Leu | 0.627 | 0.095 |
| p.Lys272Phe | substitution | 272 | Lys | Phe | 0.985 | 0.008 |
| p.Lys272Ser | substitution | 272 | Lys | Ser | 1.115 | 0.036 |

| | | | | | | |
|---|---|---|---|---|---|---|
| p.Lys272Ter | nonsense | 272 | Lys | Ter | 0.544 | 0.880 |
| p.Lys272Thr | substitution | 272 | Lys | Thr | 0.853 | 0.161 |
| p.Lys272Trp | substitution | 272 | Lys | Trp | 1.522 | 0.004 |
| p.Lys272Tyr | substitution | 272 | Lys | Tyr | 0.000 | 0.009 |
| p.Lys272Val | substitution | 272 | Lys | Val | 1.404 | 0.005 |
| p.Lys27= | synonymous | 27 | Lys | NA | 1.033 | 0.006 |
| p.Lys27Ala | substitution | 27 | Lys | Ala | 1.023 | 0.028 |
| p.Lys27Arg | substitution | 27 | Lys | Arg | 1.226 | 0.007 |
| p.Lys27Asn | substitution | 27 | Lys | Asn | 1.139 | 0.001 |
| p.Lys27Asp | substitution | 27 | Lys | Asp | 1.143 | 0.001 |
| p.Lys27Cys | substitution | 27 | Lys | Cys | 0.988 | 0.001 |
| p.Lys27Gln | substitution | 27 | Lys | Gln | 0.649 | 0.001 |
| p.Lys27Glu | substitution | 27 | Lys | Glu | 0.805 | 0.002 |
| p.Lys27Gly | substitution | 27 | Lys | Gly | 0.745 | 0.002 |
| p.Lys27His | substitution | 27 | Lys | His | 0.779 | 0.001 |
| p.Lys27Ile | substitution | 27 | Lys | Ile | 0.778 | 0.000 |
| p.Lys27Leu | substitution | 27 | Lys | Leu | 0.838 | 0.001 |
| p.Lys27Met | substitution | 27 | Lys | Met | 0.883 | 0.000 |
| p.Lys27Phe | substitution | 27 | Lys | Phe | 0.980 | 0.000 |
| p.Lys27Pro | substitution | 27 | Lys | Pro | 0.863 | 0.002 |
| p.Lys27Ser | substitution | 27 | Lys | Ser | 0.992 | 0.002 |
| p.Lys27Ter | nonsense | 27 | Lys | Ter | 0.000 | 0.003 |
| p.Lys27Thr | substitution | 27 | Lys | Thr | 0.727 | 0.046 |
| p.Lys27Trp | substitution | 27 | Lys | Trp | 0.993 | 0.001 |
| p.Lys27Tyr | substitution | 27 | Lys | Tyr | 0.667 | 0.001 |
| p.Lys27Val | substitution | 27 | Lys | Val | 1.126 | 0.013 |
| p.Lys345= | synonymous | 345 | Lys | NA | 1.305 | 0.003 |
| p.Lys345Ala | substitution | 345 | Lys | Ala | 1.071 | 0.158 |
| p.Lys345Arg | substitution | 345 | Lys | Arg | 0.104 | 0.149 |
| p.Lys345Asn | substitution | 345 | Lys | Asn | 0.899 | 0.028 |
| p.Lys345Asp | substitution | 345 | Lys | Asp | 0.281 | 0.015 |
| p.Lys345Cys | substitution | 345 | Lys | Cys | 0.294 | 0.108 |
| p.Lys345Gln | substitution | 345 | Lys | Gln | 0.256 | 0.312 |
| p.Lys345Glu | substitution | 345 | Lys | Glu | 0.000 | 0.250 |
| p.Lys345Gly | substitution | 345 | Lys | Gly | 0.311 | 0.071 |
| p.Lys345Ile | substitution | 345 | Lys | Ile | 1.046 | 0.020 |
| p.Lys345Leu | substitution | 345 | Lys | Leu | 0.878 | 0.124 |
| p.Lys345Met | substitution | 345 | Lys | Met | 1.110 | 0.002 |
| p.Lys345Phe | substitution | 345 | Lys | Phe | 1.437 | 0.003 |
| p.Lys345Pro | substitution | 345 | Lys | Pro | 0.000 | 0.130 |
| p.Lys345Ter | nonsense | 345 | Lys | Ter | 0.229 | 0.051 |
| p.Lys345Thr | substitution | 345 | Lys | Thr | 1.363 | 0.171 |
| p.Lys345Tyr | substitution | 345 | Lys | Tyr | 1.159 | 0.003 |

| | | | | | | | |
|---|---|---|---|---|---|---|---|
| p.Lys345Val | substitution | 345 | Lys | Val | | 0.339 | 0.033 |
| p.Lys348= | synonymous | 348 | Lys | NA | | 0.625 | 0.114 |
| p.Lys348Arg | substitution | 348 | Lys | Arg | | 1.317 | 0.021 |
| p.Lys348Asn | substitution | 348 | Lys | Asn | | 1.329 | 0.020 |
| p.Lys348Asp | substitution | 348 | Lys | Asp | | 0.947 | 0.119 |
| p.Lys348Cys | substitution | 348 | Lys | Cys | | 0.000 | 0.055 |
| p.Lys348Gln | substitution | 348 | Lys | Gln | | 0.732 | 0.077 |
| p.Lys348Glu | substitution | 348 | Lys | Glu | | 1.025 | 0.073 |
| p.Lys348Gly | substitution | 348 | Lys | Gly | | 1.065 | 0.004 |
| p.Lys348His | substitution | 348 | Lys | His | | 1.217 | 0.010 |
| p.Lys348Ile | substitution | 348 | Lys | Ile | | 1.292 | 0.013 |
| p.Lys348Leu | substitution | 348 | Lys | Leu | | 1.166 | 0.125 |
| p.Lys348Met | substitution | 348 | Lys | Met | | 0.356 | 0.006 |
| p.Lys348Phe | substitution | 348 | Lys | Phe | | 0.878 | 0.003 |
| p.Lys348Pro | substitution | 348 | Lys | Pro | | 0.032 | 0.035 |
| p.Lys348Ser | substitution | 348 | Lys | Ser | | 1.127 | 0.016 |
| p.Lys348Ter | nonsense | 348 | Lys | Ter | | 0.184 | 0.089 |
| p.Lys348Thr | substitution | 348 | Lys | Thr | | 0.287 | 0.081 |
| p.Lys348Trp | substitution | 348 | Lys | Trp | | 1.061 | 0.004 |
| p.Lys348Tyr | substitution | 348 | Lys | Tyr | | 0.000 | 0.035 |
| p.Lys348Val | substitution | 348 | Lys | Val | | 0.000 | 0.093 |
| p.Lys43= | synonymous | 43 | Lys | NA | | 0.750 | 0.005 |
| p.Lys43Ala | substitution | 43 | Lys | Ala | | 0.755 | 0.009 |
| p.Lys43Arg | substitution | 43 | Lys | Arg | | 0.797 | 0.010 |
| p.Lys43Asn | substitution | 43 | Lys | Asn | | 0.859 | 0.032 |
| p.Lys43Cys | substitution | 43 | Lys | Cys | | 0.673 | 0.006 |
| p.Lys43Gln | substitution | 43 | Lys | Gln | | 0.855 | 0.047 |
| p.Lys43Glu | substitution | 43 | Lys | Glu | | 0.720 | 0.026 |
| p.Lys43Gly | substitution | 43 | Lys | Gly | | 0.749 | 0.013 |
| p.Lys43His | substitution | 43 | Lys | His | | 0.754 | 0.004 |
| p.Lys43Ile | substitution | 43 | Lys | Ile | | 0.987 | 0.005 |
| p.Lys43Leu | substitution | 43 | Lys | Leu | | 0.669 | 0.016 |
| p.Lys43Met | substitution | 43 | Lys | Met | | 0.691 | 0.002 |
| p.Lys43Phe | substitution | 43 | Lys | Phe | | 0.383 | 0.011 |
| p.Lys43Pro | substitution | 43 | Lys | Pro | | 0.038 | 0.020 |
| p.Lys43Ser | substitution | 43 | Lys | Ser | | 0.817 | 0.008 |
| p.Lys43Ter | nonsense | 43 | Lys | Ter | | 0.132 | 0.152 |
| p.Lys43Thr | substitution | 43 | Lys | Thr | | 0.985 | 0.008 |
| p.Lys43Trp | substitution | 43 | Lys | Trp | | 0.697 | 0.003 |
| p.Lys43Tyr | substitution | 43 | Lys | Tyr | | 0.624 | 0.005 |
| p.Lys43Val | substitution | 43 | Lys | Val | | 1.303 | 0.009 |
| p.Lys62= | synonymous | 62 | Lys | NA | | 0.694 | 0.052 |
| p.Lys62Ala | substitution | 62 | Lys | Ala | | 0.898 | 0.238 |

| | | | | | | | |
|---|---|---|---|---|---|---|---|
| p.Lys62Arg | substitution | 62 | Lys | Arg | | 0.638 | 0.090 |
| p.Lys62Asn | substitution | 62 | Lys | Asn | | 1.263 | 0.007 |
| p.Lys62Asp | substitution | 62 | Lys | Asp | | 1.729 | 0.018 |
| p.Lys62Cys | substitution | 62 | Lys | Cys | | 0.460 | 0.024 |
| p.Lys62Gln | substitution | 62 | Lys | Gln | | 1.364 | 0.040 |
| p.Lys62Glu | substitution | 62 | Lys | Glu | | 0.659 | 0.428 |
| p.Lys62Gly | substitution | 62 | Lys | Gly | | 2.421 | 0.533 |
| p.Lys62His | substitution | 62 | Lys | His | | 0.000 | 0.083 |
| p.Lys62Ile | substitution | 62 | Lys | Ile | | 1.248 | 0.033 |
| p.Lys62Leu | substitution | 62 | Lys | Leu | | 0.275 | 0.056 |
| p.Lys62Met | substitution | 62 | Lys | Met | | 0.818 | 0.006 |
| p.Lys62Phe | substitution | 62 | Lys | Phe | | 0.562 | 0.022 |
| p.Lys62Pro | substitution | 62 | Lys | Pro | | 0.000 | 0.054 |
| p.Lys62Ser | substitution | 62 | Lys | Ser | | 0.846 | 0.062 |
| p.Lys62Ter | nonsense | 62 | Lys | Ter | | 0.784 | 0.006 |
| p.Lys62Thr | substitution | 62 | Lys | Thr | | 0.000 | 0.225 |
| p.Lys62Trp | substitution | 62 | Lys | Trp | | 0.561 | 0.026 |
| p.Lys62Tyr | substitution | 62 | Lys | Tyr | | 0.951 | 0.009 |
| p.Lys62Val | substitution | 62 | Lys | Val | | 1.025 | 0.163 |
| p.Lys70= | synonymous | 70 | Lys | NA | | 1.209 | 0.014 |
| p.Lys70Ala | substitution | 70 | Lys | Ala | | 0.754 | 0.019 |
| p.Lys70Arg | substitution | 70 | Lys | Arg | | 0.631 | 0.046 |
| p.Lys70Asn | substitution | 70 | Lys | Asn | | 1.204 | 0.006 |
| p.Lys70Asp | substitution | 70 | Lys | Asp | | 0.771 | 0.012 |
| p.Lys70Cys | substitution | 70 | Lys | Cys | | 0.657 | 0.007 |
| p.Lys70Gln | substitution | 70 | Lys | Gln | | 0.581 | 0.015 |
| p.Lys70Glu | substitution | 70 | Lys | Glu | | 0.724 | 0.103 |
| p.Lys70Gly | substitution | 70 | Lys | Gly | | 0.872 | 0.005 |
| p.Lys70His | substitution | 70 | Lys | His | | 0.747 | 0.016 |
| p.Lys70Ile | substitution | 70 | Lys | Ile | | 1.039 | 0.006 |
| p.Lys70Leu | substitution | 70 | Lys | Leu | | 0.508 | 0.013 |
| p.Lys70Met | substitution | 70 | Lys | Met | | 1.080 | 0.007 |
| p.Lys70Phe | substitution | 70 | Lys | Phe | | 0.794 | 0.009 |
| p.Lys70Pro | substitution | 70 | Lys | Pro | | 0.686 | 0.023 |
| p.Lys70Ser | substitution | 70 | Lys | Ser | | 0.578 | 0.120 |
| p.Lys70Ter | nonsense | 70 | Lys | Ter | | 0.000 | 0.654 |
| p.Lys70Thr | substitution | 70 | Lys | Thr | | 0.673 | 0.014 |
| p.Lys70Trp | substitution | 70 | Lys | Trp | | 0.724 | 0.005 |
| p.Lys70Tyr | substitution | 70 | Lys | Tyr | | 0.803 | 0.006 |
| p.Lys70Val | substitution | 70 | Lys | Val | | 1.063 | 0.067 |
| p.Lys74= | synonymous | 74 | Lys | NA | | 0.989 | 0.013 |
| p.Lys74Ala | substitution | 74 | Lys | Ala | | 0.780 | 0.005 |
| p.Lys74Arg | substitution | 74 | Lys | Arg | | 0.290 | 0.047 |

| | | | | | | | |
|---|---|---|---|---|---|---|---|
| p.Lys74Asn | substitution | 74 | Lys | Asn | | 1.033 | 0.025 |
| p.Lys74Asp | substitution | 74 | Lys | Asp | | 0.314 | 0.006 |
| p.Lys74Cys | substitution | 74 | Lys | Cys | | 0.309 | 0.009 |
| p.Lys74Gln | substitution | 74 | Lys | Gln | | 1.366 | 0.007 |
| p.Lys74Glu | substitution | 74 | Lys | Glu | | 0.000 | 0.129 |
| p.Lys74Gly | substitution | 74 | Lys | Gly | | 0.724 | 0.062 |
| p.Lys74His | substitution | 74 | Lys | His | | 0.839 | 0.007 |
| p.Lys74Ile | substitution | 74 | Lys | Ile | | 0.543 | 0.049 |
| p.Lys74Leu | substitution | 74 | Lys | Leu | | 0.483 | 0.032 |
| p.Lys74Met | substitution | 74 | Lys | Met | | 0.016 | 0.012 |
| p.Lys74Phe | substitution | 74 | Lys | Phe | | 0.635 | 0.008 |
| p.Lys74Pro | substitution | 74 | Lys | Pro | | 0.476 | 0.010 |
| p.Lys74Ser | substitution | 74 | Lys | Ser | | 0.385 | 0.089 |
| p.Lys74Ter | nonsense | 74 | Lys | Ter | | 0.000 | 0.060 |
| p.Lys74Thr | substitution | 74 | Lys | Thr | | 0.592 | 0.037 |
| p.Lys74Trp | substitution | 74 | Lys | Trp | | 0.000 | 0.069 |
| p.Lys74Tyr | substitution | 74 | Lys | Tyr | | 0.543 | 0.010 |
| p.Lys74Val | substitution | 74 | Lys | Val | | 0.155 | 0.009 |
| p.Lys79= | synonymous | 79 | Lys | NA | | 1.339 | 0.012 |
| p.Lys79Ala | substitution | 79 | Lys | Ala | | 0.796 | 0.018 |
| p.Lys79Arg | substitution | 79 | Lys | Arg | | 0.864 | 0.027 |
| p.Lys79Asn | substitution | 79 | Lys | Asn | | 1.010 | 0.105 |
| p.Lys79Asp | substitution | 79 | Lys | Asp | | 0.841 | 0.004 |
| p.Lys79Cys | substitution | 79 | Lys | Cys | | 0.831 | 0.005 |
| p.Lys79Gln | substitution | 79 | Lys | Gln | | 0.834 | 0.005 |
| p.Lys79Glu | substitution | 79 | Lys | Glu | | 0.933 | 0.010 |
| p.Lys79Gly | substitution | 79 | Lys | Gly | | 0.919 | 0.010 |
| p.Lys79His | substitution | 79 | Lys | His | | 0.958 | 0.009 |
| p.Lys79Ile | substitution | 79 | Lys | Ile | | 0.710 | 0.007 |
| p.Lys79Leu | substitution | 79 | Lys | Leu | | 0.667 | 0.006 |
| p.Lys79Met | substitution | 79 | Lys | Met | | 0.735 | 0.010 |
| p.Lys79Phe | substitution | 79 | Lys | Phe | | 0.354 | 0.008 |
| p.Lys79Pro | substitution | 79 | Lys | Pro | | 0.663 | 0.006 |
| p.Lys79Ser | substitution | 79 | Lys | Ser | | 1.101 | 0.002 |
| p.Lys79Ter | nonsense | 79 | Lys | Ter | | 0.474 | 0.017 |
| p.Lys79Thr | substitution | 79 | Lys | Thr | | 0.773 | 0.007 |
| p.Lys79Trp | substitution | 79 | Lys | Trp | | 0.910 | 0.004 |
| p.Lys79Tyr | substitution | 79 | Lys | Tyr | | 0.976 | 0.004 |
| p.Lys79Val | substitution | 79 | Lys | Val | | 0.456 | 0.039 |
| p.Lys87= | synonymous | 87 | Lys | NA | | 1.035 | 0.017 |
| p.Lys87Ala | substitution | 87 | Lys | Ala | | 0.934 | 0.038 |
| p.Lys87Arg | substitution | 87 | Lys | Arg | | 0.984 | 0.007 |
| p.Lys87Asn | substitution | 87 | Lys | Asn | | 0.944 | 0.904 |

| | | | | | | | |
|---|---|---|---|---|---|---|---|
| p.Lys87Asp | substitution | 87 | Lys | Asp | | 1.042 | 0.004 |
| p.Lys87Cys | substitution | 87 | Lys | Cys | | 0.916 | 0.005 |
| p.Lys87Gln | substitution | 87 | Lys | Gln | | 1.175 | 0.009 |
| p.Lys87Glu | substitution | 87 | Lys | Glu | | 0.784 | 0.006 |
| p.Lys87Gly | substitution | 87 | Lys | Gly | | 0.571 | 0.025 |
| p.Lys87His | substitution | 87 | Lys | His | | 0.607 | 0.021 |
| p.Lys87Ile | substitution | 87 | Lys | Ile | | 1.011 | 0.003 |
| p.Lys87Leu | substitution | 87 | Lys | Leu | | 0.892 | 0.012 |
| p.Lys87Met | substitution | 87 | Lys | Met | | 0.959 | 0.006 |
| p.Lys87Phe | substitution | 87 | Lys | Phe | | 0.692 | 0.004 |
| p.Lys87Pro | substitution | 87 | Lys | Pro | | 0.000 | 0.017 |
| p.Lys87Ser | substitution | 87 | Lys | Ser | | 1.056 | 0.004 |
| p.Lys87Ter | nonsense | 87 | Lys | Ter | | 0.285 | 0.030 |
| p.Lys87Thr | substitution | 87 | Lys | Thr | | 0.962 | 0.006 |
| p.Lys87Trp | substitution | 87 | Lys | Trp | | 0.971 | 0.005 |
| p.Lys87Tyr | substitution | 87 | Lys | Tyr | | 1.266 | 0.003 |
| p.Lys87Val | substitution | 87 | Lys | Val | | 0.504 | 0.011 |
| p.Lys98= | synonymous | 98 | Lys | NA | | 0.842 | 0.028 |
| p.Lys98Ala | substitution | 98 | Lys | Ala | | 0.105 | 0.033 |
| p.Lys98Arg | substitution | 98 | Lys | Arg | | 0.000 | 0.015 |
| p.Lys98Asn | substitution | 98 | Lys | Asn | | 0.466 | 0.011 |
| p.Lys98Asp | substitution | 98 | Lys | Asp | | 0.000 | 0.021 |
| p.Lys98Cys | substitution | 98 | Lys | Cys | | 0.325 | 0.008 |
| p.Lys98Gln | substitution | 98 | Lys | Gln | | 0.000 | 0.012 |
| p.Lys98Glu | substitution | 98 | Lys | Glu | | 0.153 | 0.224 |
| p.Lys98Gly | substitution | 98 | Lys | Gly | | 0.000 | 0.018 |
| p.Lys98His | substitution | 98 | Lys | His | | 0.000 | 0.028 |
| p.Lys98Ile | substitution | 98 | Lys | Ile | | 0.218 | 0.014 |
| p.Lys98Leu | substitution | 98 | Lys | Leu | | 0.000 | 0.007 |
| p.Lys98Met | substitution | 98 | Lys | Met | | 0.000 | 0.017 |
| p.Lys98Phe | substitution | 98 | Lys | Phe | | 0.000 | 0.024 |
| p.Lys98Pro | substitution | 98 | Lys | Pro | | 0.024 | 0.032 |
| p.Lys98Ser | substitution | 98 | Lys | Ser | | 0.000 | 0.015 |
| p.Lys98Ter | nonsense | 98 | Lys | Ter | | 0.000 | 0.046 |
| p.Lys98Thr | substitution | 98 | Lys | Thr | | 0.000 | 0.029 |
| p.Lys98Trp | substitution | 98 | Lys | Trp | | 0.000 | 0.020 |
| p.Lys98Tyr | substitution | 98 | Lys | Tyr | | 0.000 | 0.022 |
| p.Lys98Val | substitution | 98 | Lys | Val | | 0.000 | 0.007 |
| p.Met196Ala | substitution | 196 | Met | Ala | | 0.965 | 0.102 |
| p.Met196Arg | substitution | 196 | Met | Arg | | 0.371 | 0.043 |
| p.Met196Asn | substitution | 196 | Met | Asn | | 0.751 | 0.004 |
| p.Met196Cys | substitution | 196 | Met | Cys | | 1.200 | 0.002 |
| p.Met196Gln | substitution | 196 | Met | Gln | | 1.232 | 0.003 |

| | | | | | | | |
|---|---|---|---|---|---|---|---|
| p.Met196Glu | substitution | 196 | Met | Glu | | 0.233 | 0.006 |
| p.Met196Gly | substitution | 196 | Met | Gly | | 0.730 | 0.031 |
| p.Met196His | substitution | 196 | Met | His | | 0.975 | 0.002 |
| p.Met196Ile | substitution | 196 | Met | Ile | | 1.071 | 0.015 |
| p.Met196Leu | substitution | 196 | Met | Leu | | 0.951 | 0.029 |
| p.Met196Lys | substitution | 196 | Met | Lys | | 0.276 | 0.007 |
| p.Met196Phe | substitution | 196 | Met | Phe | | 1.154 | 0.002 |
| p.Met196Pro | substitution | 196 | Met | Pro | | 0.000 | 0.005 |
| p.Met196Ser | substitution | 196 | Met | Ser | | 0.322 | 0.031 |
| p.Met196Ter | nonsense | 196 | Met | Ter | | 0.000 | 0.017 |
| p.Met196Thr | substitution | 196 | Met | Thr | | 0.409 | 0.007 |
| p.Met196Trp | substitution | 196 | Met | Trp | | 0.268 | 0.002 |
| p.Met196Tyr | substitution | 196 | Met | Tyr | | 0.952 | 0.003 |
| p.Met196Val | substitution | 196 | Met | Val | | 0.864 | 0.007 |
| p.Met212Ala | substitution | 212 | Met | Ala | | 0.000 | 0.082 |
| p.Met212Arg | substitution | 212 | Met | Arg | | 0.000 | 0.026 |
| p.Met212Asn | substitution | 212 | Met | Asn | | 0.000 | 0.031 |
| p.Met212Asp | substitution | 212 | Met | Asp | | 0.000 | 0.063 |
| p.Met212Cys | substitution | 212 | Met | Cys | | 0.000 | 0.171 |
| p.Met212Gln | substitution | 212 | Met | Gln | | 0.019 | 0.029 |
| p.Met212Glu | substitution | 212 | Met | Glu | | 0.296 | 0.140 |
| p.Met212Gly | substitution | 212 | Met | Gly | | 1.195 | 0.022 |
| p.Met212His | substitution | 212 | Met | His | | 1.105 | 0.003 |
| p.Met212Ile | substitution | 212 | Met | Ile | | 1.209 | 0.025 |
| p.Met212Leu | substitution | 212 | Met | Leu | | 1.120 | 0.014 |
| p.Met212Lys | substitution | 212 | Met | Lys | | 0.000 | 0.609 |
| p.Met212Phe | substitution | 212 | Met | Phe | | 2.044 | 0.001 |
| p.Met212Pro | substitution | 212 | Met | Pro | | 0.291 | 0.131 |
| p.Met212Ser | substitution | 212 | Met | Ser | | 0.744 | 0.004 |
| p.Met212Ter | nonsense | 212 | Met | Ter | | 0.000 | 0.013 |
| p.Met212Thr | substitution | 212 | Met | Thr | | 0.847 | 0.030 |
| p.Met212Trp | substitution | 212 | Met | Trp | | 0.000 | 0.013 |
| p.Met212Tyr | substitution | 212 | Met | Tyr | | 0.000 | 0.045 |
| p.Met212Val | substitution | 212 | Met | Val | | 0.165 | 0.252 |
| p.Met271Ala | substitution | 271 | Met | Ala | | 0.076 | 0.013 |
| p.Met271Arg | substitution | 271 | Met | Arg | | 0.567 | 0.124 |
| p.Met271Asn | substitution | 271 | Met | Asn | | 0.601 | 0.007 |
| p.Met271Asp | substitution | 271 | Met | Asp | | 1.269 | 0.008 |
| p.Met271Cys | substitution | 271 | Met | Cys | | 0.454 | 0.004 |
| p.Met271Gln | substitution | 271 | Met | Gln | | 0.706 | 0.010 |
| p.Met271Glu | substitution | 271 | Met | Glu | | 1.447 | 0.004 |
| p.Met271Gly | substitution | 271 | Met | Gly | | 1.017 | 0.007 |
| p.Met271His | substitution | 271 | Met | His | | 1.264 | 0.005 |

| | | | | | | | |
|---|---|---|---|---|---|---|---|
| p.Met271Ile | substitution | 271 | Met | Ile | | 1.282 | 0.013 |
| p.Met271Leu | substitution | 271 | Met | Leu | | 0.979 | 0.067 |
| p.Met271Lys | substitution | 271 | Met | Lys | | 0.823 | 0.344 |
| p.Met271Phe | substitution | 271 | Met | Phe | | 1.056 | 0.004 |
| p.Met271Pro | substitution | 271 | Met | Pro | | 0.000 | 0.060 |
| p.Met271Ser | substitution | 271 | Met | Ser | | 0.961 | 0.023 |
| p.Met271Ter | nonsense | 271 | Met | Ter | | 0.000 | 0.008 |
| p.Met271Thr | substitution | 271 | Met | Thr | | 0.446 | 0.035 |
| p.Met271Trp | substitution | 271 | Met | Trp | | 0.398 | 0.029 |
| p.Met271Tyr | substitution | 271 | Met | Tyr | | 1.225 | 0.005 |
| p.Met271Val | substitution | 271 | Met | Val | | 0.785 | 0.229 |
| p.Met295Ala | substitution | 295 | Met | Ala | | 0.971 | 0.012 |
| p.Met295Arg | substitution | 295 | Met | Arg | | 1.126 | 0.008 |
| p.Met295Asn | substitution | 295 | Met | Asn | | 0.000 | 0.008 |
| p.Met295Asp | substitution | 295 | Met | Asp | | 0.687 | 0.003 |
| p.Met295Cys | substitution | 295 | Met | Cys | | 0.103 | 0.005 |
| p.Met295Gln | substitution | 295 | Met | Gln | | 0.292 | 0.008 |
| p.Met295Glu | substitution | 295 | Met | Glu | | 0.515 | 0.003 |
| p.Met295Gly | substitution | 295 | Met | Gly | | 1.115 | 0.003 |
| p.Met295His | substitution | 295 | Met | His | | 0.055 | 0.021 |
| p.Met295Ile | substitution | 295 | Met | Ile | | 0.916 | 0.571 |
| p.Met295Leu | substitution | 295 | Met | Leu | | 1.083 | 0.013 |
| p.Met295Lys | substitution | 295 | Met | Lys | | 0.932 | 0.011 |
| p.Met295Phe | substitution | 295 | Met | Phe | | 0.525 | 0.005 |
| p.Met295Pro | substitution | 295 | Met | Pro | | 0.000 | 0.017 |
| p.Met295Ser | substitution | 295 | Met | Ser | | 0.173 | 0.005 |
| p.Met295Ter | nonsense | 295 | Met | Ter | | 0.000 | 0.007 |
| p.Met295Thr | substitution | 295 | Met | Thr | | 0.479 | 0.075 |
| p.Met295Trp | substitution | 295 | Met | Trp | | 0.754 | 0.002 |
| p.Met295Tyr | substitution | 295 | Met | Tyr | | 1.367 | 0.008 |
| p.Met295Val | substitution | 295 | Met | Val | | 1.064 | 0.022 |
| p.Met56Ala | substitution | 56 | Met | Ala | | 0.922 | 0.161 |
| p.Met56Arg | substitution | 56 | Met | Arg | | 0.676 | 0.075 |
| p.Met56Asn | substitution | 56 | Met | Asn | | 0.491 | 0.023 |
| p.Met56Asp | substitution | 56 | Met | Asp | | 1.096 | 0.014 |
| p.Met56Cys | substitution | 56 | Met | Cys | | 0.150 | 0.017 |
| p.Met56Gln | substitution | 56 | Met | Gln | | 1.006 | 0.012 |
| p.Met56Glu | substitution | 56 | Met | Glu | | 0.932 | 0.008 |
| p.Met56Gly | substitution | 56 | Met | Gly | | 0.436 | 0.021 |
| p.Met56His | substitution | 56 | Met | His | | 0.971 | 0.037 |
| p.Met56Ile | substitution | 56 | Met | Ile | | 1.166 | 0.033 |
| p.Met56Leu | substitution | 56 | Met | Leu | | 1.076 | 0.104 |
| p.Met56Lys | substitution | 56 | Met | Lys | | 1.055 | 0.052 |

| | | | | | | | |
|---|---|---|---|---|---|---|---|
| p.Met56Phe | substitution | 56 | Met | Phe | | 1.881 | 0.071 |
| p.Met56Pro | substitution | 56 | Met | Pro | | 0.356 | 0.051 |
| p.Met56Ser | substitution | 56 | Met | Ser | | 1.155 | 0.098 |
| p.Met56Ter | nonsense | 56 | Met | Ter | | 0.000 | 0.026 |
| p.Met56Thr | substitution | 56 | Met | Thr | | 0.977 | 0.134 |
| p.Met56Trp | substitution | 56 | Met | Trp | | 0.628 | 0.005 |
| p.Met56Tyr | substitution | 56 | Met | Tyr | | 0.000 | 0.097 |
| p.Met56Val | substitution | 56 | Met | Val | | 0.570 | 0.105 |
| p.Phe108= | synonymous | 108 | Phe | NA | | 1.018 | 0.218 |
| p.Phe108Ala | substitution | 108 | Phe | Ala | | 0.000 | 0.254 |
| p.Phe108Arg | substitution | 108 | Phe | Arg | | 1.094 | 0.049 |
| p.Phe108Asn | substitution | 108 | Phe | Asn | | 0.684 | 0.012 |
| p.Phe108Asp | substitution | 108 | Phe | Asp | | 0.000 | 0.026 |
| p.Phe108Cys | substitution | 108 | Phe | Cys | | 0.784 | 0.075 |
| p.Phe108Gln | substitution | 108 | Phe | Gln | | 1.452 | 0.015 |
| p.Phe108Glu | substitution | 108 | Phe | Glu | | 1.513 | 0.013 |
| p.Phe108Gly | substitution | 108 | Phe | Gly | | 0.538 | 0.129 |
| p.Phe108His | substitution | 108 | Phe | His | | 0.000 | 0.207 |
| p.Phe108Ile | substitution | 108 | Phe | Ile | | 1.127 | 0.007 |
| p.Phe108Leu | substitution | 108 | Phe | Leu | | 1.071 | 0.052 |
| p.Phe108Lys | substitution | 108 | Phe | Lys | | 0.000 | 0.037 |
| p.Phe108Me | substitution | 108 | Phe | Met | | 0.000 | 0.028 |
| p.Phe108Pro | substitution | 108 | Phe | Pro | | 1.183 | 0.161 |
| p.Phe108Ser | substitution | 108 | Phe | Ser | | 0.283 | 0.066 |
| p.Phe108Ter | nonsense | 108 | Phe | Ter | | 0.000 | 0.163 |
| p.Phe108Thr | substitution | 108 | Phe | Thr | | 1.110 | 0.097 |
| p.Phe108Trp | substitution | 108 | Phe | Trp | | 1.139 | 0.006 |
| p.Phe108Tyr | substitution | 108 | Phe | Tyr | | 1.036 | 0.115 |
| p.Phe108Val | substitution | 108 | Phe | Val | | 0.631 | 0.021 |
| p.Phe125= | synonymous | 125 | Phe | NA | | 1.053 | 0.052 |
| p.Phe125Ala | substitution | 125 | Phe | Ala | | 0.361 | 0.021 |
| p.Phe125Arg | substitution | 125 | Phe | Arg | | 0.058 | 0.025 |
| p.Phe125Asn | substitution | 125 | Phe | Asn | | 0.689 | 0.011 |
| p.Phe125Asp | substitution | 125 | Phe | Asp | | 0.000 | 0.042 |
| p.Phe125Cys | substitution | 125 | Phe | Cys | | 0.000 | 0.034 |
| p.Phe125Gln | substitution | 125 | Phe | Gln | | 1.807 | 0.147 |
| p.Phe125Glu | substitution | 125 | Phe | Glu | | 0.517 | 0.013 |
| p.Phe125Gly | substitution | 125 | Phe | Gly | | 0.000 | 0.009 |
| p.Phe125His | substitution | 125 | Phe | His | | 0.112 | 0.104 |
| p.Phe125Ile | substitution | 125 | Phe | Ile | | 1.002 | 0.013 |
| p.Phe125Leu | substitution | 125 | Phe | Leu | | 0.913 | 0.011 |
| p.Phe125Lys | substitution | 125 | Phe | Lys | | 0.000 | 0.028 |
| p.Phe125Me | substitution | 125 | Phe | Met | | 0.900 | 0.008 |

| | | | | | | |
|---|---|---|---|---|---|---|
| p.Phe125Pro | substitution | 125 | Phe | Pro | 0.946 | 0.021 |
| p.Phe125Ser | substitution | 125 | Phe | Ser | 0.168 | 0.065 |
| p.Phe125Ter | nonsense | 125 | Phe | Ter | 0.403 | 0.033 |
| p.Phe125Thr | substitution | 125 | Phe | Thr | 0.109 | 0.068 |
| p.Phe125Trp | substitution | 125 | Phe | Trp | 0.934 | 0.006 |
| p.Phe125Tyr | substitution | 125 | Phe | Tyr | 1.379 | 0.007 |
| p.Phe125Val | substitution | 125 | Phe | Val | 0.772 | 0.024 |
| p.Phe129= | synonymous | 129 | Phe | NA | 0.510 | 0.736 |
| p.Phe129Ala | substitution | 129 | Phe | Ala | 0.524 | 0.022 |
| p.Phe129Arg | substitution | 129 | Phe | Arg | 0.376 | 0.092 |
| p.Phe129Asn | substitution | 129 | Phe | Asn | 0.000 | 0.103 |
| p.Phe129Asp | substitution | 129 | Phe | Asp | 0.660 | 0.012 |
| p.Phe129Cys | substitution | 129 | Phe | Cys | 0.584 | 0.017 |
| p.Phe129Gln | substitution | 129 | Phe | Gln | 1.067 | 0.011 |
| p.Phe129Glu | substitution | 129 | Phe | Glu | 1.588 | 0.023 |
| p.Phe129Gly | substitution | 129 | Phe | Gly | 0.630 | 0.013 |
| p.Phe129His | substitution | 129 | Phe | His | 0.845 | 0.019 |
| p.Phe129Ile | substitution | 129 | Phe | Ile | 0.615 | 0.194 |
| p.Phe129Leu | substitution | 129 | Phe | Leu | 0.955 | 0.009 |
| p.Phe129Lys | substitution | 129 | Phe | Lys | 0.180 | 0.066 |
| p.Phe129Me | substitution | 129 | Phe | Met | 1.274 | 0.015 |
| p.Phe129Pro | substitution | 129 | Phe | Pro | 0.000 | 0.114 |
| p.Phe129Ser | substitution | 129 | Phe | Ser | 0.610 | 0.036 |
| p.Phe129Ter | nonsense | 129 | Phe | Ter | 0.000 | 0.086 |
| p.Phe129Thr | substitution | 129 | Phe | Thr | 0.825 | 0.017 |
| p.Phe129Trp | substitution | 129 | Phe | Trp | 0.000 | 0.073 |
| p.Phe129Tyr | substitution | 129 | Phe | Tyr | 1.009 | 0.018 |
| p.Phe129Val | substitution | 129 | Phe | Val | 0.765 | 0.015 |
| p.Phe158= | synonymous | 158 | Phe | NA | 1.009 | 0.010 |
| p.Phe158Ala | substitution | 158 | Phe | Ala | 0.765 | 0.014 |
| p.Phe158Arg | substitution | 158 | Phe | Arg | 0.734 | 0.029 |
| p.Phe158Asn | substitution | 158 | Phe | Asn | 0.973 | 0.018 |
| p.Phe158Asp | substitution | 158 | Phe | Asp | 0.697 | 0.025 |
| p.Phe158Cys | substitution | 158 | Phe | Cys | 0.968 | 0.016 |
| p.Phe158Gln | substitution | 158 | Phe | Gln | 0.755 | 0.012 |
| p.Phe158Glu | substitution | 158 | Phe | Glu | 0.103 | 0.009 |
| p.Phe158Gly | substitution | 158 | Phe | Gly | 0.666 | 0.013 |
| p.Phe158His | substitution | 158 | Phe | His | 1.161 | 0.009 |
| p.Phe158Ile | substitution | 158 | Phe | Ile | 0.972 | 0.017 |
| p.Phe158Leu | substitution | 158 | Phe | Leu | 0.858 | 0.005 |
| p.Phe158Lys | substitution | 158 | Phe | Lys | 0.431 | 0.014 |
| p.Phe158Pro | substitution | 158 | Phe | Pro | 0.000 | 0.079 |
| p.Phe158Ser | substitution | 158 | Phe | Ser | 0.696 | 0.026 |

| | | | | | | |
|---|---|---|---|---|---|---|
| p.Phe158Ter | nonsense | 158 | Phe | Ter | 0.111 | 0.010 |
| p.Phe158Thr | substitution | 158 | Phe | Thr | 1.083 | 0.009 |
| p.Phe158Trp | substitution | 158 | Phe | Trp | 0.994 | 0.003 |
| p.Phe158Tyr | substitution | 158 | Phe | Tyr | 1.078 | 0.006 |
| p.Phe158Val | substitution | 158 | Phe | Val | 0.355 | 0.060 |
| p.Phe163= | synonymous | 163 | Phe | NA | 1.009 | 0.013 |
| p.Phe163Ala | substitution | 163 | Phe | Ala | 0.949 | 0.008 |
| p.Phe163Arg | substitution | 163 | Phe | Arg | 0.732 | 0.006 |
| p.Phe163Asn | substitution | 163 | Phe | Asn | 1.171 | 0.160 |
| p.Phe163Asp | substitution | 163 | Phe | Asp | 0.131 | 0.056 |
| p.Phe163Cys | substitution | 163 | Phe | Cys | 0.851 | 0.004 |
| p.Phe163Gln | substitution | 163 | Phe | Gln | 0.802 | 0.004 |
| p.Phe163Glu | substitution | 163 | Phe | Glu | 0.603 | 0.004 |
| p.Phe163Gly | substitution | 163 | Phe | Gly | 0.692 | 0.012 |
| p.Phe163His | substitution | 163 | Phe | His | 0.845 | 0.017 |
| p.Phe163Ile | substitution | 163 | Phe | Ile | 1.023 | 0.018 |
| p.Phe163Leu | substitution | 163 | Phe | Leu | 1.110 | 0.017 |
| p.Phe163Me | substitution | 163 | Phe | Met | 0.877 | 0.005 |
| p.Phe163Pro | substitution | 163 | Phe | Pro | 0.900 | 0.006 |
| p.Phe163Ser | substitution | 163 | Phe | Ser | 0.651 | 0.011 |
| p.Phe163Ter | nonsense | 163 | Phe | Ter | 0.000 | 0.010 |
| p.Phe163Thr | substitution | 163 | Phe | Thr | 1.126 | 0.016 |
| p.Phe163Trp | substitution | 163 | Phe | Trp | 1.033 | 0.004 |
| p.Phe163Tyr | substitution | 163 | Phe | Tyr | 0.842 | 0.022 |
| p.Phe163Val | substitution | 163 | Phe | Val | 1.067 | 0.008 |
| p.Phe183= | synonymous | 183 | Phe | NA | 0.975 | 0.035 |
| p.Phe183Cys | substitution | 183 | Phe | Cys | 0.620 | 0.706 |
| p.Phe183Ile | substitution | 183 | Phe | Ile | 1.184 | 0.012 |
| p.Phe183Leu | substitution | 183 | Phe | Leu | 1.128 | 0.047 |
| p.Phe183Ser | substitution | 183 | Phe | Ser | 0.827 | 1.020 |
| p.Phe183Val | substitution | 183 | Phe | Val | 1.098 | 0.119 |
| p.Phe253= | synonymous | 253 | Phe | NA | 1.378 | 0.001 |
| p.Phe253Ala | substitution | 253 | Phe | Ala | 0.062 | 0.020 |
| p.Phe253Arg | substitution | 253 | Phe | Arg | 0.000 | 0.055 |
| p.Phe253Asn | substitution | 253 | Phe | Asn | 0.000 | 0.205 |
| p.Phe253Asp | substitution | 253 | Phe | Asp | 0.000 | 0.059 |
| p.Phe253Cys | substitution | 253 | Phe | Cys | 0.000 | 0.089 |
| p.Phe253Gln | substitution | 253 | Phe | Gln | 0.000 | 0.014 |
| p.Phe253Glu | substitution | 253 | Phe | Glu | 0.000 | 0.045 |
| p.Phe253Gly | substitution | 253 | Phe | Gly | 0.134 | 0.024 |
| p.Phe253His | substitution | 253 | Phe | His | 0.257 | 0.011 |
| p.Phe253Ile | substitution | 253 | Phe | Ile | 1.001 | 0.008 |
| p.Phe253Leu | substitution | 253 | Phe | Leu | 0.770 | 0.006 |

| | | | | | | | |
|---|---|---|---|---|---|---|---|
| p.Phe253Lys | substitution | 253 | Phe | Lys | | 0.000 | 0.049 |
| p.Phe253Me | substitution | 253 | Phe | Met | | 1.135 | 0.002 |
| p.Phe253Pro | substitution | 253 | Phe | Pro | | 0.126 | 0.062 |
| p.Phe253Ser | substitution | 253 | Phe | Ser | | 0.000 | 0.017 |
| p.Phe253Ter | nonsense | 253 | Phe | Ter | | 0.000 | 0.013 |
| p.Phe253Thr | substitution | 253 | Phe | Thr | | 0.205 | 0.022 |
| p.Phe253Trp | substitution | 253 | Phe | Trp | | 0.193 | 0.020 |
| p.Phe253Tyr | substitution | 253 | Phe | Tyr | | 1.006 | 0.006 |
| p.Phe253Val | substitution | 253 | Phe | Val | | 0.304 | 0.088 |
| p.Phe51= | synonymous | 51 | Phe | NA | | 0.346 | 1.140 |
| p.Phe51Ala | substitution | 51 | Phe | Ala | | 0.586 | 0.017 |
| p.Phe51Arg | substitution | 51 | Phe | Arg | | 0.000 | 0.008 |
| p.Phe51Asn | substitution | 51 | Phe | Asn | | 0.318 | 0.013 |
| p.Phe51Asp | substitution | 51 | Phe | Asp | | 0.551 | 0.013 |
| p.Phe51Gln | substitution | 51 | Phe | Gln | | 0.439 | 0.008 |
| p.Phe51Glu | substitution | 51 | Phe | Glu | | 0.126 | 0.015 |
| p.Phe51Gly | substitution | 51 | Phe | Gly | | 0.677 | 0.020 |
| p.Phe51His | substitution | 51 | Phe | His | | 0.735 | 0.012 |
| p.Phe51Ile | substitution | 51 | Phe | Ile | | 0.947 | 0.006 |
| p.Phe51Leu | substitution | 51 | Phe | Leu | | 0.888 | 0.043 |
| p.Phe51Lys | substitution | 51 | Phe | Lys | | 0.317 | 0.013 |
| p.Phe51Met | substitution | 51 | Phe | Met | | 0.750 | 0.010 |
| p.Phe51Pro | substitution | 51 | Phe | Pro | | 0.965 | 0.029 |
| p.Phe51Ser | substitution | 51 | Phe | Ser | | 0.451 | 0.009 |
| p.Phe51Ter | nonsense | 51 | Phe | Ter | | 0.000 | 0.021 |
| p.Phe51Thr | substitution | 51 | Phe | Thr | | 1.032 | 0.047 |
| p.Phe51Trp | substitution | 51 | Phe | Trp | | 0.839 | 0.005 |
| p.Phe51Tyr | substitution | 51 | Phe | Tyr | | 0.626 | 0.015 |
| p.Phe51Val | substitution | 51 | Phe | Val | | 1.130 | 0.010 |
| p.Phe77Ala | substitution | 77 | Phe | Ala | | 0.120 | 0.027 |
| p.Phe77Arg | substitution | 77 | Phe | Arg | | 0.000 | 0.010 |
| p.Phe77Asn | substitution | 77 | Phe | Asn | | 0.000 | 0.013 |
| p.Phe77Asp | substitution | 77 | Phe | Asp | | 0.049 | 0.012 |
| p.Phe77Cys | substitution | 77 | Phe | Cys | | 0.128 | 0.006 |
| p.Phe77Gln | substitution | 77 | Phe | Gln | | 0.100 | 0.011 |
| p.Phe77Glu | substitution | 77 | Phe | Glu | | 0.000 | 0.011 |
| p.Phe77Gly | substitution | 77 | Phe | Gly | | 0.000 | 0.011 |
| p.Phe77His | substitution | 77 | Phe | His | | 0.000 | 0.018 |
| p.Phe77Ile | substitution | 77 | Phe | Ile | | 0.187 | 0.020 |
| p.Phe77Leu | substitution | 77 | Phe | Leu | | 0.000 | 0.019 |
| p.Phe77Lys | substitution | 77 | Phe | Lys | | 0.000 | 0.038 |
| p.Phe77Met | substitution | 77 | Phe | Met | | 0.000 | 0.017 |
| p.Phe77Pro | substitution | 77 | Phe | Pro | | 0.000 | 0.046 |

| | | | | | | |
|---|---|---|---|---|---|---|
| p.Phe77Ser | substitution | 77 | Phe | Ser | 0.094 | 0.057 |
| p.Phe77Ter | nonsense | 77 | Phe | Ter | 0.176 | 0.006 |
| p.Phe77Thr | substitution | 77 | Phe | Thr | 0.000 | 0.032 |
| p.Phe77Trp | substitution | 77 | Phe | Trp | 0.823 | 0.003 |
| p.Phe77Tyr | substitution | 77 | Phe | Tyr | 0.685 | 0.005 |
| p.Phe77Val | substitution | 77 | Phe | Val | 0.000 | 0.010 |
| p.Pro101= | synonymous | 101 | Pro | NA | 0.932 | 0.003 |
| p.Pro101Ala | substitution | 101 | Pro | Ala | 0.965 | 0.020 |
| p.Pro101Arg | substitution | 101 | Pro | Arg | 0.000 | 0.283 |
| p.Pro101Asn | substitution | 101 | Pro | Asn | 0.587 | 0.038 |
| p.Pro101Asp | substitution | 101 | Pro | Asp | 0.000 | 0.099 |
| p.Pro101Cys | substitution | 101 | Pro | Cys | 0.000 | 0.055 |
| p.Pro101Gln | substitution | 101 | Pro | Gln | 0.661 | 0.063 |
| p.Pro101Glu | substitution | 101 | Pro | Glu | 0.658 | 0.009 |
| p.Pro101Gly | substitution | 101 | Pro | Gly | 0.323 | 0.031 |
| p.Pro101His | substitution | 101 | Pro | His | 0.980 | 0.021 |
| p.Pro101Ile | substitution | 101 | Pro | Ile | 0.784 | 0.010 |
| p.Pro101Leu | substitution | 101 | Pro | Leu | 0.490 | 0.079 |
| p.Pro101Lys | substitution | 101 | Pro | Lys | 0.000 | 0.978 |
| p.Pro101Met | substitution | 101 | Pro | Met | 0.341 | 0.097 |
| p.Pro101Phe | substitution | 101 | Pro | Phe | 0.458 | 0.013 |
| p.Pro101Ser | substitution | 101 | Pro | Ser | 0.822 | 0.050 |
| p.Pro101Ter | nonsense | 101 | Pro | Ter | 0.000 | 0.065 |
| p.Pro101Thr | substitution | 101 | Pro | Thr | 0.912 | 0.021 |
| p.Pro101Trp | substitution | 101 | Pro | Trp | 0.000 | 0.030 |
| p.Pro101Tyr | substitution | 101 | Pro | Tyr | 0.000 | 0.115 |
| p.Pro101Val | substitution | 101 | Pro | Val | 0.678 | 0.073 |
| p.Pro105= | synonymous | 105 | Pro | NA | 0.995 | 0.019 |
| p.Pro105Ala | substitution | 105 | Pro | Ala | 0.910 | 0.005 |
| p.Pro105Arg | substitution | 105 | Pro | Arg | 0.802 | 0.011 |
| p.Pro105Asn | substitution | 105 | Pro | Asn | 0.455 | 0.009 |
| p.Pro105Asp | substitution | 105 | Pro | Asp | 0.307 | 0.012 |
| p.Pro105Cys | substitution | 105 | Pro | Cys | 0.369 | 0.011 |
| p.Pro105Gln | substitution | 105 | Pro | Gln | 0.845 | 0.005 |
| p.Pro105Glu | substitution | 105 | Pro | Glu | 0.437 | 0.016 |
| p.Pro105Gly | substitution | 105 | Pro | Gly | 0.952 | 0.008 |
| p.Pro105His | substitution | 105 | Pro | His | 0.885 | 0.004 |
| p.Pro105Ile | substitution | 105 | Pro | Ile | 0.597 | 0.005 |
| p.Pro105Leu | substitution | 105 | Pro | Leu | 0.646 | 0.035 |
| p.Pro105Lys | substitution | 105 | Pro | Lys | 0.891 | 0.019 |
| p.Pro105Met | substitution | 105 | Pro | Met | 0.747 | 0.012 |
| p.Pro105Phe | substitution | 105 | Pro | Phe | 1.145 | 0.003 |
| p.Pro105Ser | substitution | 105 | Pro | Ser | 0.769 | 0.056 |

| | | | | | | | |
|---|---|---|---|---|---|---|---|
| p.Pro105Ter | nonsense | 105 | Pro | Ter | | 0.002 | 0.029 |
| p.Pro105Thr | substitution | 105 | Pro | Thr | | 0.911 | 0.023 |
| p.Pro105Trp | substitution | 105 | Pro | Trp | | 0.669 | 0.020 |
| p.Pro105Tyr | substitution | 105 | Pro | Tyr | | 0.007 | 0.010 |
| p.Pro105Val | substitution | 105 | Pro | Val | | 0.410 | 0.005 |
| p.Pro106= | synonymous | 106 | Pro | NA | | 0.964 | 0.033 |
| p.Pro106Ala | substitution | 106 | Pro | Ala | | 0.925 | 0.010 |
| p.Pro106Arg | substitution | 106 | Pro | Arg | | 0.966 | 0.004 |
| p.Pro106Asn | substitution | 106 | Pro | Asn | | 1.014 | 0.007 |
| p.Pro106Asp | substitution | 106 | Pro | Asp | | 0.889 | 0.005 |
| p.Pro106Cys | substitution | 106 | Pro | Cys | | 1.078 | 0.011 |
| p.Pro106Gln | substitution | 106 | Pro | Gln | | 0.874 | 0.012 |
| p.Pro106Glu | substitution | 106 | Pro | Glu | | 1.140 | 0.008 |
| p.Pro106Gly | substitution | 106 | Pro | Gly | | 0.772 | 0.025 |
| p.Pro106His | substitution | 106 | Pro | His | | 1.005 | 0.005 |
| p.Pro106Ile | substitution | 106 | Pro | Ile | | 0.779 | 0.007 |
| p.Pro106Leu | substitution | 106 | Pro | Leu | | 0.983 | 0.029 |
| p.Pro106Lys | substitution | 106 | Pro | Lys | | 1.082 | 0.009 |
| p.Pro106Met | substitution | 106 | Pro | Met | | 0.381 | 0.018 |
| p.Pro106Phe | substitution | 106 | Pro | Phe | | 1.061 | 0.005 |
| p.Pro106Ser | substitution | 106 | Pro | Ser | | 1.019 | 0.003 |
| p.Pro106Ter | nonsense | 106 | Pro | Ter | | 0.121 | 0.015 |
| p.Pro106Thr | substitution | 106 | Pro | Thr | | 1.009 | 0.003 |
| p.Pro106Trp | substitution | 106 | Pro | Trp | | 0.879 | 0.009 |
| p.Pro106Tyr | substitution | 106 | Pro | Tyr | | 0.486 | 0.010 |
| p.Pro106Val | substitution | 106 | Pro | Val | | 1.160 | 0.012 |
| p.Pro119= | synonymous | 119 | Pro | NA | | 0.836 | 0.069 |
| p.Pro119Ala | substitution | 119 | Pro | Ala | | 0.923 | 0.003 |
| p.Pro119Arg | substitution | 119 | Pro | Arg | | 0.181 | 0.011 |
| p.Pro119Asn | substitution | 119 | Pro | Asn | | 0.388 | 0.008 |
| p.Pro119Asp | substitution | 119 | Pro | Asp | | 0.128 | 0.007 |
| p.Pro119Cys | substitution | 119 | Pro | Cys | | 0.826 | 0.008 |
| p.Pro119Glu | substitution | 119 | Pro | Glu | | 0.451 | 0.034 |
| p.Pro119Gly | substitution | 119 | Pro | Gly | | 0.279 | 0.007 |
| p.Pro119His | substitution | 119 | Pro | His | | 0.782 | 0.002 |
| p.Pro119Ile | substitution | 119 | Pro | Ile | | 0.000 | 0.011 |
| p.Pro119Leu | substitution | 119 | Pro | Leu | | 0.021 | 0.029 |
| p.Pro119Lys | substitution | 119 | Pro | Lys | | 0.110 | 0.061 |
| p.Pro119Met | substitution | 119 | Pro | Met | | 0.000 | 0.014 |
| p.Pro119Phe | substitution | 119 | Pro | Phe | | 0.788 | 0.008 |
| p.Pro119Ser | substitution | 119 | Pro | Ser | | 0.834 | 0.032 |
| p.Pro119Ter | nonsense | 119 | Pro | Ter | | 0.000 | 0.424 |
| p.Pro119Thr | substitution | 119 | Pro | Thr | | 0.981 | 0.033 |

| | | | | | | |
|---|---|---|---|---|---|---|
| p.Pro119Trp | substitution | 119 | Pro | Trp | 0.837 | 0.009 |
| p.Pro119Tyr | substitution | 119 | Pro | Tyr | 0.697 | 0.005 |
| p.Pro119Val | substitution | 119 | Pro | Val | 0.787 | 0.008 |
| p.Pro127= | synonymous | 127 | Pro | NA | 0.965 | 0.075 |
| p.Pro127Ala | substitution | 127 | Pro | Ala | 0.703 | 0.073 |
| p.Pro127Arg | substitution | 127 | Pro | Arg | 1.025 | 0.035 |
| p.Pro127Asn | substitution | 127 | Pro | Asn | 0.000 | 0.875 |
| p.Pro127Asp | substitution | 127 | Pro | Asp | 0.404 | 0.350 |
| p.Pro127Cys | substitution | 127 | Pro | Cys | 0.084 | 0.034 |
| p.Pro127Gln | substitution | 127 | Pro | Gln | 1.174 | 0.037 |
| p.Pro127Gly | substitution | 127 | Pro | Gly | 0.568 | 0.648 |
| p.Pro127His | substitution | 127 | Pro | His | 0.266 | 0.150 |
| p.Pro127Ile | substitution | 127 | Pro | Ile | 1.073 | 0.012 |
| p.Pro127Leu | substitution | 127 | Pro | Leu | 1.155 | 0.068 |
| p.Pro127Lys | substitution | 127 | Pro | Lys | 0.820 | 0.131 |
| p.Pro127Met | substitution | 127 | Pro | Met | 0.067 | 0.044 |
| p.Pro127Phe | substitution | 127 | Pro | Phe | 0.827 | 0.066 |
| p.Pro127Ser | substitution | 127 | Pro | Ser | 0.844 | 0.057 |
| p.Pro127Ter | nonsense | 127 | Pro | Ter | 0.647 | 0.045 |
| p.Pro127Thr | substitution | 127 | Pro | Thr | 0.886 | 0.094 |
| p.Pro127Trp | substitution | 127 | Pro | Trp | 1.135 | 0.021 |
| p.Pro127Tyr | substitution | 127 | Pro | Tyr | 0.000 | 0.898 |
| p.Pro127Val | substitution | 127 | Pro | Val | 0.534 | 0.070 |
| p.Pro138= | synonymous | 138 | Pro | NA | 0.976 | 0.006 |
| p.Pro138Ala | substitution | 138 | Pro | Ala | 1.036 | 0.003 |
| p.Pro138Arg | substitution | 138 | Pro | Arg | 0.772 | 0.022 |
| p.Pro138Asn | substitution | 138 | Pro | Asn | 0.895 | 0.004 |
| p.Pro138Asp | substitution | 138 | Pro | Asp | 0.788 | 0.006 |
| p.Pro138Cys | substitution | 138 | Pro | Cys | 0.816 | 0.004 |
| p.Pro138Gln | substitution | 138 | Pro | Gln | 1.123 | 0.007 |
| p.Pro138Glu | substitution | 138 | Pro | Glu | 0.945 | 0.107 |
| p.Pro138Gly | substitution | 138 | Pro | Gly | 0.628 | 0.008 |
| p.Pro138His | substitution | 138 | Pro | His | 0.754 | 0.004 |
| p.Pro138Ile | substitution | 138 | Pro | Ile | 0.696 | 0.022 |
| p.Pro138Leu | substitution | 138 | Pro | Leu | 0.821 | 0.015 |
| p.Pro138Lys | substitution | 138 | Pro | Lys | 0.836 | 0.030 |
| p.Pro138Met | substitution | 138 | Pro | Met | 0.939 | 0.004 |
| p.Pro138Phe | substitution | 138 | Pro | Phe | 0.582 | 0.005 |
| p.Pro138Ser | substitution | 138 | Pro | Ser | 0.887 | 0.004 |
| p.Pro138Ter | nonsense | 138 | Pro | Ter | 0.121 | 0.049 |
| p.Pro138Thr | substitution | 138 | Pro | Thr | 0.808 | 0.006 |
| p.Pro138Trp | substitution | 138 | Pro | Trp | 0.339 | 0.008 |
| p.Pro138Tyr | substitution | 138 | Pro | Tyr | 0.511 | 0.005 |

| | | | | | | |
|---|---|---|---|---|---|---|
| p.Pro138Val | substitution | 138 | Pro | Val | 0.695 | 0.006 |
| p.Pro159= | synonymous | 159 | Pro | NA | 1.074 | 0.076 |
| p.Pro159Ala | substitution | 159 | Pro | Ala | 0.951 | 0.015 |
| p.Pro159Arg | substitution | 159 | Pro | Arg | 0.805 | 0.009 |
| p.Pro159Asn | substitution | 159 | Pro | Asn | 1.075 | 0.006 |
| p.Pro159Asp | substitution | 159 | Pro | Asp | 0.571 | 0.008 |
| p.Pro159Cys | substitution | 159 | Pro | Cys | 1.001 | 0.005 |
| p.Pro159Gln | substitution | 159 | Pro | Gln | 0.834 | 0.014 |
| p.Pro159Glu | substitution | 159 | Pro | Glu | 0.694 | 0.008 |
| p.Pro159Gly | substitution | 159 | Pro | Gly | 0.726 | 0.006 |
| p.Pro159His | substitution | 159 | Pro | His | 1.015 | 0.006 |
| p.Pro159Ile | substitution | 159 | Pro | Ile | 0.663 | 0.008 |
| p.Pro159Leu | substitution | 159 | Pro | Leu | 0.785 | 0.029 |
| p.Pro159Lys | substitution | 159 | Pro | Lys | 0.786 | 0.008 |
| p.Pro159Met | substitution | 159 | Pro | Met | 0.635 | 0.018 |
| p.Pro159Phe | substitution | 159 | Pro | Phe | 0.652 | 0.030 |
| p.Pro159Ser | substitution | 159 | Pro | Ser | 0.723 | 0.065 |
| p.Pro159Ter | nonsense | 159 | Pro | Ter | 0.348 | 0.018 |
| p.Pro159Thr | substitution | 159 | Pro | Thr | 0.730 | 0.033 |
| p.Pro159Trp | substitution | 159 | Pro | Trp | 0.265 | 0.014 |
| p.Pro159Tyr | substitution | 159 | Pro | Tyr | 0.413 | 0.011 |
| p.Pro159Val | substitution | 159 | Pro | Val | 1.109 | 0.056 |
| p.Pro208= | synonymous | 208 | Pro | NA | 1.252 | 0.014 |
| p.Pro208Ala | substitution | 208 | Pro | Ala | 0.497 | 0.038 |
| p.Pro208Arg | substitution | 208 | Pro | Arg | 0.960 | 0.033 |
| p.Pro208Asn | substitution | 208 | Pro | Asn | 0.000 | 0.050 |
| p.Pro208Asp | substitution | 208 | Pro | Asp | 0.182 | 0.011 |
| p.Pro208Cys | substitution | 208 | Pro | Cys | 0.060 | 0.019 |
| p.Pro208Gln | substitution | 208 | Pro | Gln | 0.781 | 0.006 |
| p.Pro208Glu | substitution | 208 | Pro | Glu | 0.000 | 0.023 |
| p.Pro208Gly | substitution | 208 | Pro | Gly | 1.052 | 0.098 |
| p.Pro208His | substitution | 208 | Pro | His | 0.899 | 0.002 |
| p.Pro208Ile | substitution | 208 | Pro | Ile | 1.058 | 0.004 |
| p.Pro208Leu | substitution | 208 | Pro | Leu | 0.814 | 0.022 |
| p.Pro208Lys | substitution | 208 | Pro | Lys | 0.218 | 0.020 |
| p.Pro208Met | substitution | 208 | Pro | Met | 0.983 | 0.003 |
| p.Pro208Phe | substitution | 208 | Pro | Phe | 1.086 | 0.004 |
| p.Pro208Ser | substitution | 208 | Pro | Ser | 0.785 | 0.047 |
| p.Pro208Ter | nonsense | 208 | Pro | Ter | 0.000 | 0.042 |
| p.Pro208Thr | substitution | 208 | Pro | Thr | 0.649 | 0.003 |
| p.Pro208Tyr | substitution | 208 | Pro | Tyr | 1.171 | 0.004 |
| p.Pro208Val | substitution | 208 | Pro | Val | 0.414 | 0.028 |
| p.Pro241= | synonymous | 241 | Pro | NA | 1.083 | 0.005 |

| | | | | | | |
|---|---|---|---|---|---|---|
| p.Pro241Ala | substitution | 241 | Pro | Ala | 0.467 | 0.050 |
| p.Pro241Arg | substitution | 241 | Pro | Arg | 0.000 | 0.102 |
| p.Pro241Asp | substitution | 241 | Pro | Asp | 0.555 | 0.017 |
| p.Pro241Cys | substitution | 241 | Pro | Cys | 1.079 | 0.011 |
| p.Pro241Gln | substitution | 241 | Pro | Gln | 1.231 | 0.004 |
| p.Pro241His | substitution | 241 | Pro | His | 0.681 | 0.035 |
| p.Pro241Leu | substitution | 241 | Pro | Leu | 0.000 | 0.057 |
| p.Pro241Lys | substitution | 241 | Pro | Lys | 0.000 | 0.017 |
| p.Pro241Met | substitution | 241 | Pro | Met | 1.480 | 0.004 |
| p.Pro241Ser | substitution | 241 | Pro | Ser | 1.136 | 0.091 |
| p.Pro241Ter | nonsense | 241 | Pro | Ter | 0.777 | 0.301 |
| p.Pro241Thr | substitution | 241 | Pro | Thr | 0.601 | 0.084 |
| p.Pro241Trp | substitution | 241 | Pro | Trp | 1.113 | 0.007 |
| p.Pro241Tyr | substitution | 241 | Pro | Tyr | 0.000 | 0.070 |
| p.Pro264= | synonymous | 264 | Pro | NA | 0.488 | 0.755 |
| p.Pro264Ala | substitution | 264 | Pro | Ala | 0.180 | 0.077 |
| p.Pro264Cys | substitution | 264 | Pro | Cys | 0.000 | 0.067 |
| p.Pro264Gln | substitution | 264 | Pro | Gln | 0.764 | 0.033 |
| p.Pro264Glu | substitution | 264 | Pro | Glu | 0.280 | 0.011 |
| p.Pro264Leu | substitution | 264 | Pro | Leu | 0.306 | 0.769 |
| p.Pro264Lys | substitution | 264 | Pro | Lys | 0.000 | 0.131 |
| p.Pro264Met | substitution | 264 | Pro | Met | 0.000 | 0.075 |
| p.Pro264Ser | substitution | 264 | Pro | Ser | 1.315 | 0.071 |
| p.Pro264Ter | nonsense | 264 | Pro | Ter | 0.788 | 0.063 |
| p.Pro264Thr | substitution | 264 | Pro | Thr | 1.610 | 0.062 |
| p.Pro264Trp | substitution | 264 | Pro | Trp | 0.013 | 0.165 |
| p.Pro264Tyr | substitution | 264 | Pro | Tyr | 0.000 | 0.019 |
| p.Pro264Val | substitution | 264 | Pro | Val | 0.000 | 0.124 |
| p.Pro302= | synonymous | 302 | Pro | NA | 0.804 | 0.099 |
| p.Pro302Ala | substitution | 302 | Pro | Ala | 0.945 | 0.109 |
| p.Pro302Arg | substitution | 302 | Pro | Arg | 0.227 | 0.019 |
| p.Pro302Asn | substitution | 302 | Pro | Asn | 0.000 | 0.032 |
| p.Pro302Asp | substitution | 302 | Pro | Asp | 0.923 | 0.005 |
| p.Pro302Cys | substitution | 302 | Pro | Cys | 0.896 | 0.003 |
| p.Pro302Gln | substitution | 302 | Pro | Gln | 0.295 | 0.003 |
| p.Pro302Gly | substitution | 302 | Pro | Gly | 1.414 | 0.004 |
| p.Pro302Ile | substitution | 302 | Pro | Ile | 0.000 | 0.032 |
| p.Pro302Leu | substitution | 302 | Pro | Leu | 0.915 | 0.061 |
| p.Pro302Lys | substitution | 302 | Pro | Lys | 0.332 | 0.018 |
| p.Pro302Met | substitution | 302 | Pro | Met | 1.064 | 0.010 |
| p.Pro302Phe | substitution | 302 | Pro | Phe | 0.227 | 0.019 |
| p.Pro302Ser | substitution | 302 | Pro | Ser | 0.347 | 0.031 |
| p.Pro302Ter | nonsense | 302 | Pro | Ter | 0.000 | 0.133 |

| | | | | | | | |
|---|---|---|---|---|---|---|---|
| p.Pro302Thr | substitution | 302 | Pro | Thr | | 1.129 | 0.026 |
| p.Pro302Trp | substitution | 302 | Pro | Trp | | 0.910 | 0.032 |
| p.Pro302Tyr | substitution | 302 | Pro | Tyr | | 0.655 | 0.150 |
| p.Pro302Val | substitution | 302 | Pro | Val | | 1.084 | 0.227 |
| p.Pro309= | synonymous | 309 | Pro | NA | | 1.150 | 0.011 |
| p.Pro309Ala | substitution | 309 | Pro | Ala | | 1.098 | 0.051 |
| p.Pro309Arg | substitution | 309 | Pro | Arg | | 0.849 | 0.036 |
| p.Pro309Asn | substitution | 309 | Pro | Asn | | 1.349 | 0.001 |
| p.Pro309Cys | substitution | 309 | Pro | Cys | | 0.160 | 0.258 |
| p.Pro309Gln | substitution | 309 | Pro | Gln | | 0.801 | 0.003 |
| p.Pro309Glu | substitution | 309 | Pro | Glu | | 0.066 | 0.006 |
| p.Pro309Gly | substitution | 309 | Pro | Gly | | 1.320 | 0.002 |
| p.Pro309His | substitution | 309 | Pro | His | | 0.829 | 0.002 |
| p.Pro309Ile | substitution | 309 | Pro | Ile | | 0.405 | 0.013 |
| p.Pro309Leu | substitution | 309 | Pro | Leu | | 1.052 | 0.008 |
| p.Pro309Lys | substitution | 309 | Pro | Lys | | 1.052 | 0.004 |
| p.Pro309Met | substitution | 309 | Pro | Met | | 0.000 | 0.020 |
| p.Pro309Phe | substitution | 309 | Pro | Phe | | 1.182 | 0.003 |
| p.Pro309Ser | substitution | 309 | Pro | Ser | | 0.779 | 0.135 |
| p.Pro309Ter | nonsense | 309 | Pro | Ter | | 0.000 | 0.009 |
| p.Pro309Thr | substitution | 309 | Pro | Thr | | 0.592 | 0.013 |
| p.Pro309Trp | substitution | 309 | Pro | Trp | | 0.346 | 0.035 |
| p.Pro309Tyr | substitution | 309 | Pro | Tyr | | 0.000 | 0.028 |
| p.Pro309Val | substitution | 309 | Pro | Val | | 1.268 | 0.016 |
| p.Pro313= | synonymous | 313 | Pro | NA | | 1.152 | 0.031 |
| p.Pro313Ala | substitution | 313 | Pro | Ala | | 1.108 | 0.011 |
| p.Pro313Arg | substitution | 313 | Pro | Arg | | 1.055 | 0.008 |
| p.Pro313Asn | substitution | 313 | Pro | Asn | | 0.665 | 0.004 |
| p.Pro313Asp | substitution | 313 | Pro | Asp | | 0.022 | 0.010 |
| p.Pro313Cys | substitution | 313 | Pro | Cys | | 0.469 | 0.006 |
| p.Pro313Gln | substitution | 313 | Pro | Gln | | 0.115 | 0.118 |
| p.Pro313Gly | substitution | 313 | Pro | Gly | | 0.835 | 0.057 |
| p.Pro313His | substitution | 313 | Pro | His | | 0.389 | 0.031 |
| p.Pro313Ile | substitution | 313 | Pro | Ile | | 1.354 | 0.002 |
| p.Pro313Leu | substitution | 313 | Pro | Leu | | 0.935 | 0.110 |
| p.Pro313Lys | substitution | 313 | Pro | Lys | | 0.677 | 0.018 |
| p.Pro313Phe | substitution | 313 | Pro | Phe | | 0.000 | 0.105 |
| p.Pro313Ser | substitution | 313 | Pro | Ser | | 1.151 | 0.005 |
| p.Pro313Ter | nonsense | 313 | Pro | Ter | | 0.000 | 0.043 |
| p.Pro313Thr | substitution | 313 | Pro | Thr | | 0.583 | 0.122 |
| p.Pro313Trp | substitution | 313 | Pro | Trp | | 0.005 | 0.095 |
| p.Pro313Tyr | substitution | 313 | Pro | Tyr | | 1.273 | 0.002 |
| p.Pro313Val | substitution | 313 | Pro | Val | | 1.030 | 0.013 |

| | | | | | | |
|---|---|---|---|---|---|---|
| p.Pro324= | synonymous | 324 | Pro | NA | 0.950 | 0.020 |
| p.Pro324Ala | substitution | 324 | Pro | Ala | 0.580 | 0.072 |
| p.Pro324Arg | substitution | 324 | Pro | Arg | 1.005 | 0.022 |
| p.Pro324Asp | substitution | 324 | Pro | Asp | 1.402 | 0.004 |
| p.Pro324Cys | substitution | 324 | Pro | Cys | 0.210 | 0.015 |
| p.Pro324Gln | substitution | 324 | Pro | Gln | 0.781 | 0.157 |
| p.Pro324Glu | substitution | 324 | Pro | Glu | 0.054 | 0.014 |
| p.Pro324Gly | substitution | 324 | Pro | Gly | 0.462 | 0.040 |
| p.Pro324His | substitution | 324 | Pro | His | 1.118 | 0.003 |
| p.Pro324Ile | substitution | 324 | Pro | Ile | 0.217 | 0.026 |
| p.Pro324Leu | substitution | 324 | Pro | Leu | 0.923 | 0.018 |
| p.Pro324Lys | substitution | 324 | Pro | Lys | 0.468 | 0.042 |
| p.Pro324Met | substitution | 324 | Pro | Met | 0.037 | 0.026 |
| p.Pro324Phe | substitution | 324 | Pro | Phe | 0.000 | 0.008 |
| p.Pro324Ser | substitution | 324 | Pro | Ser | 0.690 | 0.158 |
| p.Pro324Ter | nonsense | 324 | Pro | Ter | 0.000 | 0.102 |
| p.Pro324Thr | substitution | 324 | Pro | Thr | 1.095 | 0.003 |
| p.Pro324Tyr | substitution | 324 | Pro | Tyr | 0.521 | 0.005 |
| p.Pro324Val | substitution | 324 | Pro | Val | 0.729 | 0.103 |
| p.Pro327= | synonymous | 327 | Pro | NA | 0.888 | 0.015 |
| p.Pro327Arg | substitution | 327 | Pro | Arg | 0.000 | 0.129 |
| p.Pro327Asn | substitution | 327 | Pro | Asn | 0.000 | 0.140 |
| p.Pro327Cys | substitution | 327 | Pro | Cys | 0.000 | 0.035 |
| p.Pro327Gln | substitution | 327 | Pro | Gln | 1.206 | 0.008 |
| p.Pro327His | substitution | 327 | Pro | His | 0.940 | 0.004 |
| p.Pro327Leu | substitution | 327 | Pro | Leu | 0.317 | 0.071 |
| p.Pro327Ser | substitution | 327 | Pro | Ser | 0.318 | 0.621 |
| p.Pro327Thr | substitution | 327 | Pro | Thr | 0.595 | 0.688 |
| p.Pro47= | synonymous | 47 | Pro | NA | 1.147 | 0.032 |
| p.Pro47Ala | substitution | 47 | Pro | Ala | 0.459 | 0.048 |
| p.Pro47Arg | substitution | 47 | Pro | Arg | 0.919 | 0.016 |
| p.Pro47Asn | substitution | 47 | Pro | Asn | 1.003 | 0.004 |
| p.Pro47Asp | substitution | 47 | Pro | Asp | 0.739 | 0.037 |
| p.Pro47Cys | substitution | 47 | Pro | Cys | 0.475 | 0.008 |
| p.Pro47Gln | substitution | 47 | Pro | Gln | 0.893 | 0.004 |
| p.Pro47Glu | substitution | 47 | Pro | Glu | 0.000 | 0.501 |
| p.Pro47Gly | substitution | 47 | Pro | Gly | 0.598 | 0.012 |
| p.Pro47His | substitution | 47 | Pro | His | 0.807 | 0.006 |
| p.Pro47Ile | substitution | 47 | Pro | Ile | 0.136 | 0.014 |
| p.Pro47Leu | substitution | 47 | Pro | Leu | 0.394 | 0.009 |
| p.Pro47Lys | substitution | 47 | Pro | Lys | 1.273 | 0.005 |
| p.Pro47Met | substitution | 47 | Pro | Met | 0.000 | 0.031 |
| p.Pro47Phe | substitution | 47 | Pro | Phe | 0.643 | 0.008 |

| | | | | | | | |
|---|---|---|---|---|---|---|---|
| p.Pro47Ser | substitution | 47 | Pro | Ser | | 0.846 | 0.031 |
| p.Pro47Ter | nonsense | 47 | Pro | Ter | | 0.000 | 0.015 |
| p.Pro47Thr | substitution | 47 | Pro | Thr | | 0.914 | 0.006 |
| p.Pro47Trp | substitution | 47 | Pro | Trp | | 0.318 | 0.024 |
| p.Pro47Tyr | substitution | 47 | Pro | Tyr | | 0.472 | 0.008 |
| p.Pro47Val | substitution | 47 | Pro | Val | | 0.889 | 0.007 |
| p.Ser141= | synonymous | 141 | Ser | NA | | 0.838 | 0.002 |
| p.Ser141Ala | substitution | 141 | Ser | Ala | | 1.032 | 0.007 |
| p.Ser141Arg | substitution | 141 | Ser | Arg | | 0.779 | 0.008 |
| p.Ser141Asn | substitution | 141 | Ser | Asn | | 1.116 | 0.003 |
| p.Ser141Asp | substitution | 141 | Ser | Asp | | 0.589 | 0.005 |
| p.Ser141Gln | substitution | 141 | Ser | Gln | | 0.866 | 0.008 |
| p.Ser141Glu | substitution | 141 | Ser | Glu | | 0.703 | 0.010 |
| p.Ser141Gly | substitution | 141 | Ser | Gly | | 0.880 | 0.024 |
| p.Ser141His | substitution | 141 | Ser | His | | 0.896 | 0.004 |
| p.Ser141Ile | substitution | 141 | Ser | Ile | | 0.399 | 0.004 |
| p.Ser141Leu | substitution | 141 | Ser | Leu | | 0.818 | 0.002 |
| p.Ser141Lys | substitution | 141 | Ser | Lys | | 0.642 | 0.008 |
| p.Ser141Met | substitution | 141 | Ser | Met | | 1.020 | 0.006 |
| p.Ser141Phe | substitution | 141 | Ser | Phe | | 0.646 | 0.007 |
| p.Ser141Pro | substitution | 141 | Ser | Pro | | 0.105 | 0.015 |
| p.Ser141Ter | nonsense | 141 | Ser | Ter | | 0.000 | 0.010 |
| p.Ser141Thr | substitution | 141 | Ser | Thr | | 0.743 | 0.005 |
| p.Ser141Trp | substitution | 141 | Ser | Trp | | 0.399 | 0.007 |
| p.Ser141Tyr | substitution | 141 | Ser | Tyr | | 0.624 | 0.007 |
| p.Ser141Val | substitution | 141 | Ser | Val | | 0.740 | 0.004 |
| p.Ser146= | synonymous | 146 | Ser | NA | | 0.889 | 0.078 |
| p.Ser146Ala | substitution | 146 | Ser | Ala | | 0.722 | 0.040 |
| p.Ser146Arg | substitution | 146 | Ser | Arg | | 0.836 | 0.022 |
| p.Ser146Asn | substitution | 146 | Ser | Asn | | 1.034 | 0.036 |
| p.Ser146Asp | substitution | 146 | Ser | Asp | | 0.282 | 0.041 |
| p.Ser146Cys | substitution | 146 | Ser | Cys | | 0.267 | 0.448 |
| p.Ser146Gln | substitution | 146 | Ser | Gln | | 1.121 | 0.010 |
| p.Ser146Glu | substitution | 146 | Ser | Glu | | 0.000 | 0.011 |
| p.Ser146Gly | substitution | 146 | Ser | Gly | | 0.941 | 0.012 |
| p.Ser146His | substitution | 146 | Ser | His | | 0.445 | 0.159 |
| p.Ser146Ile | substitution | 146 | Ser | Ile | | 0.650 | 0.075 |
| p.Ser146Leu | substitution | 146 | Ser | Leu | | 0.188 | 0.009 |
| p.Ser146Lys | substitution | 146 | Ser | Lys | | 0.191 | 0.015 |
| p.Ser146Met | substitution | 146 | Ser | Met | | 0.494 | 0.008 |
| p.Ser146Phe | substitution | 146 | Ser | Phe | | 0.813 | 0.025 |
| p.Ser146Pro | substitution | 146 | Ser | Pro | | 0.118 | 0.011 |
| p.Ser146Ter | nonsense | 146 | Ser | Ter | | 0.395 | 0.013 |

| | | | | | | | |
|---|---|---|---|---|---|---|---|
| p.Ser146Thr | substitution | 146 | Ser | Thr | | 0.635 | 0.175 |
| p.Ser146Trp | substitution | 146 | Ser | Trp | | 0.832 | 0.013 |
| p.Ser146Tyr | substitution | 146 | Ser | Tyr | | 0.400 | 0.045 |
| p.Ser146Val | substitution | 146 | Ser | Val | | 0.607 | 0.027 |
| p.Ser147= | synonymous | 147 | Ser | NA | | 0.824 | 0.065 |
| p.Ser147Ala | substitution | 147 | Ser | Ala | | 0.000 | 0.641 |
| p.Ser147Arg | substitution | 147 | Ser | Arg | | 0.182 | 0.320 |
| p.Ser147Asn | substitution | 147 | Ser | Asn | | 0.000 | 0.757 |
| p.Ser147Asp | substitution | 147 | Ser | Asp | | 0.483 | 0.027 |
| p.Ser147Cys | substitution | 147 | Ser | Cys | | 0.305 | 0.454 |
| p.Ser147Glu | substitution | 147 | Ser | Glu | | 0.110 | 0.047 |
| p.Ser147Gly | substitution | 147 | Ser | Gly | | 0.000 | 0.610 |
| p.Ser147His | substitution | 147 | Ser | His | | 0.000 | 0.165 |
| p.Ser147Ile | substitution | 147 | Ser | Ile | | 0.685 | 0.466 |
| p.Ser147Leu | substitution | 147 | Ser | Leu | | 0.000 | 0.260 |
| p.Ser147Phe | substitution | 147 | Ser | Phe | | 0.000 | 0.217 |
| p.Ser147Pro | substitution | 147 | Ser | Pro | | 0.000 | 0.812 |
| p.Ser147Ter | nonsense | 147 | Ser | Ter | | 0.000 | 0.812 |
| p.Ser147Thr | substitution | 147 | Ser | Thr | | 0.316 | 0.126 |
| p.Ser147Trp | substitution | 147 | Ser | Trp | | 0.566 | 0.129 |
| p.Ser147Tyr | substitution | 147 | Ser | Tyr | | 0.000 | 0.135 |
| p.Ser147Val | substitution | 147 | Ser | Val | | 0.000 | 0.137 |
| p.Ser165= | synonymous | 165 | Ser | NA | | 0.611 | 0.032 |
| p.Ser165Ala | substitution | 165 | Ser | Ala | | 0.507 | 0.110 |
| p.Ser165Arg | substitution | 165 | Ser | Arg | | 0.737 | 0.015 |
| p.Ser165Asn | substitution | 165 | Ser | Asn | | 0.841 | 0.006 |
| p.Ser165Asp | substitution | 165 | Ser | Asp | | 0.000 | 0.016 |
| p.Ser165Cys | substitution | 165 | Ser | Cys | | 0.959 | 0.009 |
| p.Ser165Gln | substitution | 165 | Ser | Gln | | 0.963 | 0.008 |
| p.Ser165Glu | substitution | 165 | Ser | Glu | | 0.481 | 0.014 |
| p.Ser165Gly | substitution | 165 | Ser | Gly | | 1.008 | 0.038 |
| p.Ser165His | substitution | 165 | Ser | His | | 0.887 | 0.007 |
| p.Ser165Ile | substitution | 165 | Ser | Ile | | 0.697 | 0.008 |
| p.Ser165Leu | substitution | 165 | Ser | Leu | | 0.342 | 0.228 |
| p.Ser165Lys | substitution | 165 | Ser | Lys | | 0.298 | 0.018 |
| p.Ser165Met | substitution | 165 | Ser | Met | | 0.406 | 0.012 |
| p.Ser165Phe | substitution | 165 | Ser | Phe | | 0.579 | 0.013 |
| p.Ser165Pro | substitution | 165 | Ser | Pro | | 1.090 | 0.010 |
| p.Ser165Ter | nonsense | 165 | Ser | Ter | | 0.000 | 0.026 |
| p.Ser165Thr | substitution | 165 | Ser | Thr | | 1.181 | 0.016 |
| p.Ser165Trp | substitution | 165 | Ser | Trp | | 0.312 | 0.010 |
| p.Ser165Tyr | substitution | 165 | Ser | Tyr | | 0.811 | 0.007 |
| p.Ser165Val | substitution | 165 | Ser | Val | | 0.772 | 0.022 |

| | | | | | | | |
|---|---|---|---|---|---|---|---|
| p.Ser184= | synonymous | 184 | Ser | NA | | 1.446 | 0.381 |
| p.Ser184Arg | substitution | 184 | Ser | Arg | | 1.062 | 0.691 |
| p.Ser184Asn | substitution | 184 | Ser | Asn | | 0.999 | 0.016 |
| p.Ser184Cys | substitution | 184 | Ser | Cys | | 1.327 | 0.025 |
| p.Ser184Gly | substitution | 184 | Ser | Gly | | 0.712 | 0.437 |
| p.Ser184Ile | substitution | 184 | Ser | Ile | | 0.889 | 0.033 |
| p.Ser184Thr | substitution | 184 | Ser | Thr | | 1.144 | 0.051 |
| p.Ser262Ala | substitution | 262 | Ser | Ala | | 1.319 | 0.036 |
| p.Ser262Arg | substitution | 262 | Ser | Arg | | 1.248 | 0.094 |
| p.Ser262Asn | substitution | 262 | Ser | Asn | | 1.402 | 0.010 |
| p.Ser262Cys | substitution | 262 | Ser | Cys | | 0.007 | 0.390 |
| p.Ser262Gln | substitution | 262 | Ser | Gln | | 0.575 | 0.023 |
| p.Ser262Glu | substitution | 262 | Ser | Glu | | 1.626 | 0.004 |
| p.Ser262Gly | substitution | 262 | Ser | Gly | | 1.415 | 0.008 |
| p.Ser262His | substitution | 262 | Ser | His | | 0.042 | 0.034 |
| p.Ser262Ile | substitution | 262 | Ser | Ile | | 0.979 | 0.158 |
| p.Ser262Leu | substitution | 262 | Ser | Leu | | 0.000 | 0.095 |
| p.Ser262Met | substitution | 262 | Ser | Met | | 0.000 | 0.176 |
| p.Ser262Phe | substitution | 262 | Ser | Phe | | 0.000 | 0.096 |
| p.Ser262Thr | substitution | 262 | Ser | Thr | | 0.000 | 0.140 |
| p.Ser262Trp | substitution | 262 | Ser | Trp | | 0.000 | 0.097 |
| p.Ser262Val | substitution | 262 | Ser | Val | | 0.746 | 0.047 |
| p.Ser284= | synonymous | 284 | Ser | NA | | 1.377 | 0.023 |
| p.Ser284Ala | substitution | 284 | Ser | Ala | | 1.526 | 0.006 |
| p.Ser284Arg | substitution | 284 | Ser | Arg | | 1.268 | 0.013 |
| p.Ser284Asn | substitution | 284 | Ser | Asn | | 0.807 | 0.053 |
| p.Ser284Asp | substitution | 284 | Ser | Asp | | 0.236 | 0.052 |
| p.Ser284Cys | substitution | 284 | Ser | Cys | | 0.810 | 0.016 |
| p.Ser284Gln | substitution | 284 | Ser | Gln | | 1.625 | 0.010 |
| p.Ser284Glu | substitution | 284 | Ser | Glu | | 0.000 | 0.072 |
| p.Ser284Gly | substitution | 284 | Ser | Gly | | 0.899 | 0.035 |
| p.Ser284Ile | substitution | 284 | Ser | Ile | | 1.235 | 0.011 |
| p.Ser284Leu | substitution | 284 | Ser | Leu | | 0.000 | 0.112 |
| p.Ser284Pro | substitution | 284 | Ser | Pro | | 0.000 | 0.065 |
| p.Ser284Thr | substitution | 284 | Ser | Thr | | 1.205 | 0.106 |
| p.Ser284Trp | substitution | 284 | Ser | Trp | | 0.309 | 0.013 |
| p.Ser284Val | substitution | 284 | Ser | Val | | 0.000 | 0.090 |
| p.Ser288= | synonymous | 288 | Ser | NA | | 0.606 | 0.114 |
| p.Ser288Ala | substitution | 288 | Ser | Ala | | 1.027 | 0.008 |
| p.Ser288Arg | substitution | 288 | Ser | Arg | | 0.002 | 0.018 |
| p.Ser288Asn | substitution | 288 | Ser | Asn | | 0.558 | 0.015 |
| p.Ser288Asp | substitution | 288 | Ser | Asp | | 1.151 | 0.004 |
| p.Ser288Cys | substitution | 288 | Ser | Cys | | 0.869 | 0.004 |

| | | | | | | |
|---|---|---|---|---|---|---|
| p.Ser288Gln | substitution | 288 | Ser | Gln | 1.146 | 0.006 |
| p.Ser288Glu | substitution | 288 | Ser | Glu | 1.174 | 0.040 |
| p.Ser288Gly | substitution | 288 | Ser | Gly | 1.048 | 0.008 |
| p.Ser288His | substitution | 288 | Ser | His | 0.000 | 0.007 |
| p.Ser288Ile | substitution | 288 | Ser | Ile | 1.084 | 0.014 |
| p.Ser288Leu | substitution | 288 | Ser | Leu | 1.048 | 0.012 |
| p.Ser288Lys | substitution | 288 | Ser | Lys | 0.572 | 0.015 |
| p.Ser288Met | substitution | 288 | Ser | Met | 1.308 | 0.002 |
| p.Ser288Phe | substitution | 288 | Ser | Phe | 0.780 | 0.004 |
| p.Ser288Pro | substitution | 288 | Ser | Pro | 0.923 | 0.025 |
| p.Ser288Ter | nonsense | 288 | Ser | Ter | 0.036 | 0.053 |
| p.Ser288Thr | substitution | 288 | Ser | Thr | 0.764 | 0.042 |
| p.Ser288Trp | substitution | 288 | Ser | Trp | 1.142 | 0.004 |
| p.Ser288Tyr | substitution | 288 | Ser | Tyr | 0.467 | 0.010 |
| p.Ser288Val | substitution | 288 | Ser | Val | 0.964 | 0.013 |
| p.Ser28= | synonymous | 28 | Ser | NA | 0.985 | 0.006 |
| p.Ser28Ala | substitution | 28 | Ser | Ala | 0.551 | 0.029 |
| p.Ser28Arg | substitution | 28 | Ser | Arg | 0.000 | 0.015 |
| p.Ser28Asn | substitution | 28 | Ser | Asn | 0.000 | 0.009 |
| p.Ser28Asp | substitution | 28 | Ser | Asp | 0.000 | 0.005 |
| p.Ser28Cys | substitution | 28 | Ser | Cys | 0.112 | 0.167 |
| p.Ser28Gln | substitution | 28 | Ser | Gln | 0.000 | 0.055 |
| p.Ser28Glu | substitution | 28 | Ser | Glu | 0.000 | 0.002 |
| p.Ser28Gly | substitution | 28 | Ser | Gly | 0.901 | 0.007 |
| p.Ser28His | substitution | 28 | Ser | His | 0.000 | 0.061 |
| p.Ser28Ile | substitution | 28 | Ser | Ile | 0.000 | 0.004 |
| p.Ser28Leu | substitution | 28 | Ser | Leu | 0.000 | 0.037 |
| p.Ser28Lys | substitution | 28 | Ser | Lys | 0.000 | 0.003 |
| p.Ser28Met | substitution | 28 | Ser | Met | 0.000 | 0.008 |
| p.Ser28Phe | substitution | 28 | Ser | Phe | 0.000 | 0.005 |
| p.Ser28Pro | substitution | 28 | Ser | Pro | 0.000 | 0.046 |
| p.Ser28Ter | nonsense | 28 | Ser | Ter | 0.000 | 0.014 |
| p.Ser28Thr | substitution | 28 | Ser | Thr | 1.018 | 0.113 |
| p.Ser28Trp | substitution | 28 | Ser | Trp | 0.000 | 0.006 |
| p.Ser28Tyr | substitution | 28 | Ser | Tyr | 0.000 | 0.005 |
| p.Ser28Val | substitution | 28 | Ser | Val | 0.000 | 0.054 |
| p.Ser290= | synonymous | 290 | Ser | NA | 0.000 | 0.098 |
| p.Ser290Ala | substitution | 290 | Ser | Ala | 0.000 | 0.006 |
| p.Ser290Arg | substitution | 290 | Ser | Arg | 0.572 | 0.054 |
| p.Ser290Asn | substitution | 290 | Ser | Asn | 0.704 | 0.104 |
| p.Ser290Asp | substitution | 290 | Ser | Asp | 0.478 | 0.006 |
| p.Ser290Cys | substitution | 290 | Ser | Cys | 0.184 | 0.028 |
| p.Ser290Gln | substitution | 290 | Ser | Gln | 0.000 | 0.088 |

| | | | | | | |
|---|---|---|---|---|---|---|
| p.Ser290Glu | substitution | 290 | Ser | Glu | 2.060 | 0.067 |
| p.Ser290Gly | substitution | 290 | Ser | Gly | 0.930 | 0.013 |
| p.Ser290His | substitution | 290 | Ser | His | 0.000 | 0.036 |
| p.Ser290Ile | substitution | 290 | Ser | Ile | 1.267 | 0.049 |
| p.Ser290Leu | substitution | 290 | Ser | Leu | 0.661 | 0.091 |
| p.Ser290Lys | substitution | 290 | Ser | Lys | 0.528 | 0.006 |
| p.Ser290Met | substitution | 290 | Ser | Met | 1.260 | 0.004 |
| p.Ser290Phe | substitution | 290 | Ser | Phe | 1.014 | 0.077 |
| p.Ser290Pro | substitution | 290 | Ser | Pro | 0.361 | 0.196 |
| p.Ser290Ter | nonsense | 290 | Ser | Ter | 0.000 | 0.008 |
| p.Ser290Thr | substitution | 290 | Ser | Thr | 0.000 | 0.070 |
| p.Ser290Trp | substitution | 290 | Ser | Trp | 1.375 | 0.004 |
| p.Ser290Tyr | substitution | 290 | Ser | Tyr | 0.971 | 0.005 |
| p.Ser290Val | substitution | 290 | Ser | Val | 0.603 | 0.003 |
| p.Ser337= | synonymous | 337 | Ser | NA | 1.184 | 0.011 |
| p.Ser337Arg | substitution | 337 | Ser | Arg | 1.074 | 0.076 |
| p.Ser337Asn | substitution | 337 | Ser | Asn | 1.222 | 0.006 |
| p.Ser337Cys | substitution | 337 | Ser | Cys | 0.869 | 0.188 |
| p.Ser337Gln | substitution | 337 | Ser | Gln | 0.000 | 0.205 |
| p.Ser337Glu | substitution | 337 | Ser | Glu | 0.369 | 0.012 |
| p.Ser337Gly | substitution | 337 | Ser | Gly | 0.906 | 0.084 |
| p.Ser337Ile | substitution | 337 | Ser | Ile | 0.831 | 0.022 |
| p.Ser337Leu | substitution | 337 | Ser | Leu | 1.236 | 0.003 |
| p.Ser337Lys | substitution | 337 | Ser | Lys | 0.370 | 0.092 |
| p.Ser337Met | substitution | 337 | Ser | Met | 0.318 | 0.029 |
| p.Ser337Phe | substitution | 337 | Ser | Phe | 0.000 | 0.098 |
| p.Ser337Pro | substitution | 337 | Ser | Pro | 0.005 | 0.015 |
| p.Ser337Ter | nonsense | 337 | Ser | Ter | 0.282 | 0.214 |
| p.Ser337Thr | substitution | 337 | Ser | Thr | 1.191 | 0.047 |
| p.Ser337Trp | substitution | 337 | Ser | Trp | 0.000 | 0.069 |
| p.Ser337Val | substitution | 337 | Ser | Val | 1.378 | 0.006 |
| p.Ser344= | synonymous | 344 | Ser | NA | 0.425 | 0.049 |
| p.Ser344Ala | substitution | 344 | Ser | Ala | 0.000 | 0.079 |
| p.Ser344Arg | substitution | 344 | Ser | Arg | 1.290 | 0.019 |
| p.Ser344Asn | substitution | 344 | Ser | Asn | 1.151 | 0.134 |
| p.Ser344Asp | substitution | 344 | Ser | Asp | 1.756 | 0.024 |
| p.Ser344Cys | substitution | 344 | Ser | Cys | 0.719 | 0.045 |
| p.Ser344Gln | substitution | 344 | Ser | Gln | 0.474 | 0.069 |
| p.Ser344Glu | substitution | 344 | Ser | Glu | 1.072 | 0.003 |
| p.Ser344Gly | substitution | 344 | Ser | Gly | 1.184 | 0.022 |
| p.Ser344His | substitution | 344 | Ser | His | 1.327 | 0.007 |
| p.Ser344Ile | substitution | 344 | Ser | Ile | 1.188 | 0.004 |
| p.Ser344Leu | substitution | 344 | Ser | Leu | 0.000 | 0.150 |

| | | | | | | |
|---|---|---|---|---|---|---|
| p.Ser344Met | substitution | 344 | Ser | Met | 0.364 | 0.008 |
| p.Ser344Pro | substitution | 344 | Ser | Pro | 1.125 | 0.137 |
| p.Ser344Ter | nonsense | 344 | Ser | Ter | 0.000 | 0.138 |
| p.Ser344Thr | substitution | 344 | Ser | Thr | 1.007 | 0.095 |
| p.Ser344Trp | substitution | 344 | Ser | Trp | 0.000 | 0.128 |
| p.Ser344Tyr | substitution | 344 | Ser | Tyr | 0.608 | 0.073 |
| p.Ser344Val | substitution | 344 | Ser | Val | 0.491 | 0.007 |
| p.Ser37= | synonymous | 37 | Ser | NA | 0.957 | 0.011 |
| p.Ser37Ala | substitution | 37 | Ser | Ala | 1.104 | 0.032 |
| p.Ser37Arg | substitution | 37 | Ser | Arg | 0.865 | 0.027 |
| p.Ser37Asn | substitution | 37 | Ser | Asn | 1.127 | 0.007 |
| p.Ser37Asp | substitution | 37 | Ser | Asp | 0.304 | 0.015 |
| p.Ser37Cys | substitution | 37 | Ser | Cys | 0.908 | 0.004 |
| p.Ser37Glu | substitution | 37 | Ser | Glu | 0.828 | 0.016 |
| p.Ser37Gly | substitution | 37 | Ser | Gly | 0.664 | 0.009 |
| p.Ser37His | substitution | 37 | Ser | His | 0.960 | 0.009 |
| p.Ser37Ile | substitution | 37 | Ser | Ile | 0.982 | 0.004 |
| p.Ser37Leu | substitution | 37 | Ser | Leu | 0.914 | 0.025 |
| p.Ser37Lys | substitution | 37 | Ser | Lys | 0.999 | 0.004 |
| p.Ser37Met | substitution | 37 | Ser | Met | 1.148 | 0.005 |
| p.Ser37Phe | substitution | 37 | Ser | Phe | 0.887 | 0.003 |
| p.Ser37Pro | substitution | 37 | Ser | Pro | 0.439 | 0.011 |
| p.Ser37Ter | nonsense | 37 | Ser | Ter | 0.000 | 0.014 |
| p.Ser37Thr | substitution | 37 | Ser | Thr | 1.155 | 0.005 |
| p.Ser37Trp | substitution | 37 | Ser | Trp | 1.040 | 0.005 |
| p.Ser37Tyr | substitution | 37 | Ser | Tyr | 0.338 | 0.016 |
| p.Ser37Val | substitution | 37 | Ser | Val | 0.660 | 0.006 |
| p.Ser45= | synonymous | 45 | Ser | NA | 1.081 | 0.052 |
| p.Ser45Ala | substitution | 45 | Ser | Ala | 0.969 | 0.004 |
| p.Ser45Arg | substitution | 45 | Ser | Arg | 0.860 | 0.045 |
| p.Ser45Asp | substitution | 45 | Ser | Asp | 0.922 | 0.005 |
| p.Ser45Cys | substitution | 45 | Ser | Cys | 1.237 | 0.003 |
| p.Ser45Gln | substitution | 45 | Ser | Gln | 1.141 | 0.005 |
| p.Ser45Glu | substitution | 45 | Ser | Glu | 0.812 | 0.003 |
| p.Ser45Gly | substitution | 45 | Ser | Gly | 0.594 | 0.004 |
| p.Ser45His | substitution | 45 | Ser | His | 0.657 | 0.008 |
| p.Ser45Ile | substitution | 45 | Ser | Ile | 1.359 | 0.005 |
| p.Ser45Leu | substitution | 45 | Ser | Leu | 0.630 | 0.017 |
| p.Ser45Lys | substitution | 45 | Ser | Lys | 1.275 | 0.006 |
| p.Ser45Met | substitution | 45 | Ser | Met | 0.473 | 0.006 |
| p.Ser45Phe | substitution | 45 | Ser | Phe | 0.471 | 0.005 |
| p.Ser45Pro | substitution | 45 | Ser | Pro | 0.719 | 0.044 |
| p.Ser45Ter | nonsense | 45 | Ser | Ter | 0.326 | 0.006 |

| | | | | | | |
|---|---|---|---|---|---|---|
| p.Ser45Thr | substitution | 45 | Ser | Thr | 1.056 | 0.004 |
| p.Ser45Trp | substitution | 45 | Ser | Trp | 0.837 | 0.004 |
| p.Ser45Tyr | substitution | 45 | Ser | Tyr | 1.015 | 0.007 |
| p.Ser45Val | substitution | 45 | Ser | Val | 0.746 | 0.028 |
| p.Ser57= | synonymous | 57 | Ser | NA | 1.105 | 0.010 |
| p.Ser57Ala | substitution | 57 | Ser | Ala | 0.403 | 0.061 |
| p.Ser57Arg | substitution | 57 | Ser | Arg | 0.868 | 0.036 |
| p.Ser57Asn | substitution | 57 | Ser | Asn | 0.325 | 0.683 |
| p.Ser57Asp | substitution | 57 | Ser | Asp | 1.213 | 0.023 |
| p.Ser57Cys | substitution | 57 | Ser | Cys | 0.790 | 0.008 |
| p.Ser57Glu | substitution | 57 | Ser | Glu | 0.383 | 0.025 |
| p.Ser57Gly | substitution | 57 | Ser | Gly | 0.711 | 0.014 |
| p.Ser57His | substitution | 57 | Ser | His | 0.150 | 0.044 |
| p.Ser57Ile | substitution | 57 | Ser | Ile | 0.000 | 0.394 |
| p.Ser57Leu | substitution | 57 | Ser | Leu | 0.752 | 0.170 |
| p.Ser57Lys | substitution | 57 | Ser | Lys | 0.000 | 0.044 |
| p.Ser57Met | substitution | 57 | Ser | Met | 0.000 | 0.162 |
| p.Ser57Phe | substitution | 57 | Ser | Phe | 1.037 | 0.164 |
| p.Ser57Pro | substitution | 57 | Ser | Pro | 0.408 | 0.287 |
| p.Ser57Ter | nonsense | 57 | Ser | Ter | 0.096 | 0.043 |
| p.Ser57Thr | substitution | 57 | Ser | Thr | 0.586 | 0.408 |
| p.Ser57Trp | substitution | 57 | Ser | Trp | 0.777 | 0.012 |
| p.Ser57Tyr | substitution | 57 | Ser | Tyr | 0.820 | 0.052 |
| p.Ser57Val | substitution | 57 | Ser | Val | 1.060 | 0.009 |
| p.Ser69= | synonymous | 69 | Ser | NA | 0.849 | 0.024 |
| p.Ser69Ala | substitution | 69 | Ser | Ala | 0.666 | 0.010 |
| p.Ser69Arg | substitution | 69 | Ser | Arg | 0.889 | 0.004 |
| p.Ser69Asn | substitution | 69 | Ser | Asn | 0.950 | 0.006 |
| p.Ser69Asp | substitution | 69 | Ser | Asp | 0.699 | 0.008 |
| p.Ser69Cys | substitution | 69 | Ser | Cys | 1.030 | 0.004 |
| p.Ser69Gln | substitution | 69 | Ser | Gln | 0.937 | 0.018 |
| p.Ser69Glu | substitution | 69 | Ser | Glu | 0.982 | 0.011 |
| p.Ser69Gly | substitution | 69 | Ser | Gly | 0.797 | 0.043 |
| p.Ser69His | substitution | 69 | Ser | His | 0.965 | 0.008 |
| p.Ser69Ile | substitution | 69 | Ser | Ile | 0.879 | 0.014 |
| p.Ser69Leu | substitution | 69 | Ser | Leu | 0.695 | 0.003 |
| p.Ser69Lys | substitution | 69 | Ser | Lys | 1.155 | 0.006 |
| p.Ser69Met | substitution | 69 | Ser | Met | 1.150 | 0.004 |
| p.Ser69Phe | substitution | 69 | Ser | Phe | 0.688 | 0.012 |
| p.Ser69Pro | substitution | 69 | Ser | Pro | 0.940 | 0.009 |
| p.Ser69Ter | nonsense | 69 | Ser | Ter | 0.005 | 0.014 |
| p.Ser69Thr | substitution | 69 | Ser | Thr | 0.758 | 0.017 |
| p.Ser69Trp | substitution | 69 | Ser | Trp | 0.742 | 0.005 |

| | | | | | | |
|---|---|---|---|---|---|---|
| p.Ser69Tyr | substitution | 69 | Ser | Tyr | 0.797 | 0.005 |
| p.Ser69Val | substitution | 69 | Ser | Val | 0.797 | 0.004 |
| p.Ser75= | synonymous | 75 | Ser | NA | 0.678 | 0.013 |
| p.Ser75Ala | substitution | 75 | Ser | Ala | 1.045 | 0.011 |
| p.Ser75Arg | substitution | 75 | Ser | Arg | 0.930 | 0.046 |
| p.Ser75Asn | substitution | 75 | Ser | Asn | 1.236 | 0.051 |
| p.Ser75Asp | substitution | 75 | Ser | Asp | 0.460 | 0.036 |
| p.Ser75Cys | substitution | 75 | Ser | Cys | 1.243 | 0.033 |
| p.Ser75Gln | substitution | 75 | Ser | Gln | 0.712 | 0.015 |
| p.Ser75Glu | substitution | 75 | Ser | Glu | 0.279 | 0.049 |
| p.Ser75Gly | substitution | 75 | Ser | Gly | 0.935 | 0.036 |
| p.Ser75His | substitution | 75 | Ser | His | 0.630 | 0.024 |
| p.Ser75Ile | substitution | 75 | Ser | Ile | 0.695 | 0.029 |
| p.Ser75Leu | substitution | 75 | Ser | Leu | 0.844 | 0.097 |
| p.Ser75Lys | substitution | 75 | Ser | Lys | 1.561 | 0.005 |
| p.Ser75Met | substitution | 75 | Ser | Met | 0.470 | 0.014 |
| p.Ser75Phe | substitution | 75 | Ser | Phe | 0.000 | 0.387 |
| p.Ser75Pro | substitution | 75 | Ser | Pro | 0.953 | 0.040 |
| p.Ser75Ter | nonsense | 75 | Ser | Ter | 0.000 | 0.592 |
| p.Ser75Thr | substitution | 75 | Ser | Thr | 1.205 | 0.019 |
| p.Ser75Tyr | substitution | 75 | Ser | Tyr | 0.000 | 0.307 |
| p.Ser75Val | substitution | 75 | Ser | Val | 0.421 | 0.106 |
| p.Ser96= | synonymous | 96 | Ser | NA | 0.992 | 0.020 |
| p.Ser96Ala | substitution | 96 | Ser | Ala | 0.985 | 0.035 |
| p.Ser96Arg | substitution | 96 | Ser | Arg | 0.033 | 0.059 |
| p.Ser96Asn | substitution | 96 | Ser | Asn | 0.000 | 0.085 |
| p.Ser96Asp | substitution | 96 | Ser | Asp | 0.005 | 0.120 |
| p.Ser96Cys | substitution | 96 | Ser | Cys | 0.531 | 0.032 |
| p.Ser96Gln | substitution | 96 | Ser | Gln | 0.000 | 0.062 |
| p.Ser96Gly | substitution | 96 | Ser | Gly | 0.791 | 0.104 |
| p.Ser96His | substitution | 96 | Ser | His | 0.218 | 0.046 |
| p.Ser96Ile | substitution | 96 | Ser | Ile | 0.000 | 0.042 |
| p.Ser96Leu | substitution | 96 | Ser | Leu | 0.000 | 0.078 |
| p.Ser96Lys | substitution | 96 | Ser | Lys | 0.000 | 0.221 |
| p.Ser96Met | substitution | 96 | Ser | Met | 0.311 | 0.060 |
| p.Ser96Phe | substitution | 96 | Ser | Phe | 0.000 | 0.718 |
| p.Ser96Pro | substitution | 96 | Ser | Pro | 0.095 | 0.072 |
| p.Ser96Ter | nonsense | 96 | Ser | Ter | 0.000 | 0.032 |
| p.Ser96Thr | substitution | 96 | Ser | Thr | 0.000 | 0.018 |
| p.Ser96Trp | substitution | 96 | Ser | Trp | 0.000 | 0.582 |
| p.Ser96Tyr | substitution | 96 | Ser | Tyr | 0.128 | 0.052 |
| p.Ser96Val | substitution | 96 | Ser | Val | 0.000 | 0.190 |
| p.Thr102= | synonymous | 102 | Thr | NA | 0.849 | 0.034 |

| | | | | | | |
|---|---|---|---|---|---|---|
| p.Thr102Ala | substitution | 102 | Thr | Ala | 0.921 | 0.015 |
| p.Thr102Arg | substitution | 102 | Thr | Arg | 0.154 | 0.154 |
| p.Thr102Asn | substitution | 102 | Thr | Asn | 0.856 | 0.007 |
| p.Thr102Asp | substitution | 102 | Thr | Asp | 0.370 | 0.097 |
| p.Thr102Cys | substitution | 102 | Thr | Cys | 0.133 | 0.409 |
| p.Thr102Gln | substitution | 102 | Thr | Gln | 0.537 | 0.718 |
| p.Thr102Gly | substitution | 102 | Thr | Gly | 0.706 | 0.011 |
| p.Thr102His | substitution | 102 | Thr | His | 1.210 | 0.008 |
| p.Thr102Ile | substitution | 102 | Thr | Ile | 0.998 | 0.015 |
| p.Thr102Leu | substitution | 102 | Thr | Leu | 0.449 | 1.127 |
| p.Thr102Lys | substitution | 102 | Thr | Lys | 0.000 | 0.359 |
| p.Thr102Met | substitution | 102 | Thr | Met | 0.604 | 0.024 |
| p.Thr102Phe | substitution | 102 | Thr | Phe | 0.072 | 0.124 |
| p.Thr102Pro | substitution | 102 | Thr | Pro | 0.819 | 0.045 |
| p.Thr102Ser | substitution | 102 | Thr | Ser | 1.068 | 0.015 |
| p.Thr102Ter | nonsense | 102 | Thr | Ter | 0.000 | 0.890 |
| p.Thr102Trp | substitution | 102 | Thr | Trp | 0.000 | 0.882 |
| p.Thr102Tyr | substitution | 102 | Thr | Tyr | 1.217 | 0.013 |
| p.Thr102Val | substitution | 102 | Thr | Val | 0.360 | 0.055 |
| p.Thr109= | synonymous | 109 | Thr | NA | 1.119 | 0.003 |
| p.Thr109Ala | substitution | 109 | Thr | Ala | 0.465 | 0.189 |
| p.Thr109Arg | substitution | 109 | Thr | Arg | 0.000 | 0.222 |
| p.Thr109Asn | substitution | 109 | Thr | Asn | 0.392 | 0.108 |
| p.Thr109Asp | substitution | 109 | Thr | Asp | 0.000 | 0.890 |
| p.Thr109Cys | substitution | 109 | Thr | Cys | 0.983 | 0.038 |
| p.Thr109Gln | substitution | 109 | Thr | Gln | 0.555 | 0.075 |
| p.Thr109Glu | substitution | 109 | Thr | Glu | 0.018 | 0.315 |
| p.Thr109Gly | substitution | 109 | Thr | Gly | 0.000 | 0.225 |
| p.Thr109His | substitution | 109 | Thr | His | 0.733 | 0.021 |
| p.Thr109Ile | substitution | 109 | Thr | Ile | 0.519 | 0.066 |
| p.Thr109Leu | substitution | 109 | Thr | Leu | 0.297 | 0.230 |
| p.Thr109Lys | substitution | 109 | Thr | Lys | 1.029 | 0.010 |
| p.Thr109Met | substitution | 109 | Thr | Met | 0.686 | 0.025 |
| p.Thr109Phe | substitution | 109 | Thr | Phe | 0.213 | 0.019 |
| p.Thr109Pro | substitution | 109 | Thr | Pro | 0.000 | 0.148 |
| p.Thr109Ser | substitution | 109 | Thr | Ser | 0.770 | 0.038 |
| p.Thr109Ter | nonsense | 109 | Thr | Ter | 0.000 | 0.185 |
| p.Thr109Trp | substitution | 109 | Thr | Trp | 0.492 | 0.027 |
| p.Thr109Tyr | substitution | 109 | Thr | Tyr | 0.179 | 0.036 |
| p.Thr109Val | substitution | 109 | Thr | Val | 1.229 | 0.199 |
| p.Thr133= | synonymous | 133 | Thr | NA | 1.003 | 0.009 |
| p.Thr133Ala | substitution | 133 | Thr | Ala | 0.582 | 0.064 |
| p.Thr133Arg | substitution | 133 | Thr | Arg | 1.048 | 0.022 |

| | | | | | | |
|---|---|---|---|---|---|---|
| p.Thr133Asn | substitution | 133 | Thr | Asn | 1.075 | 0.022 |
| p.Thr133Asp | substitution | 133 | Thr | Asp | 1.027 | 0.024 |
| p.Thr133Cys | substitution | 133 | Thr | Cys | 0.943 | 0.022 |
| p.Thr133Gln | substitution | 133 | Thr | Gln | 0.435 | 0.013 |
| p.Thr133Glu | substitution | 133 | Thr | Glu | 0.938 | 0.013 |
| p.Thr133Gly | substitution | 133 | Thr | Gly | 0.453 | 0.141 |
| p.Thr133His | substitution | 133 | Thr | His | 0.502 | 0.048 |
| p.Thr133Ile | substitution | 133 | Thr | Ile | 0.747 | 0.017 |
| p.Thr133Leu | substitution | 133 | Thr | Leu | 0.704 | 0.052 |
| p.Thr133Lys | substitution | 133 | Thr | Lys | 0.719 | 0.015 |
| p.Thr133Met | substitution | 133 | Thr | Met | 0.957 | 0.005 |
| p.Thr133Phe | substitution | 133 | Thr | Phe | 0.317 | 0.030 |
| p.Thr133Pro | substitution | 133 | Thr | Pro | 0.368 | 0.101 |
| p.Thr133Ser | substitution | 133 | Thr | Ser | 0.782 | 0.018 |
| p.Thr133Ter | nonsense | 133 | Thr | Ter | 0.000 | 0.015 |
| p.Thr133Trp | substitution | 133 | Thr | Trp | 0.744 | 0.011 |
| p.Thr133Tyr | substitution | 133 | Thr | Tyr | 0.671 | 0.028 |
| p.Thr133Val | substitution | 133 | Thr | Val | 0.670 | 0.019 |
| p.Thr136= | synonymous | 136 | Thr | NA | 0.865 | 0.007 |
| p.Thr136Ala | substitution | 136 | Thr | Ala | 0.617 | 0.094 |
| p.Thr136Arg | substitution | 136 | Thr | Arg | 0.879 | 0.007 |
| p.Thr136Asn | substitution | 136 | Thr | Asn | 0.932 | 0.012 |
| p.Thr136Asp | substitution | 136 | Thr | Asp | 0.950 | 0.025 |
| p.Thr136Cys | substitution | 136 | Thr | Cys | 0.468 | 0.016 |
| p.Thr136Gln | substitution | 136 | Thr | Gln | 0.806 | 0.005 |
| p.Thr136Glu | substitution | 136 | Thr | Glu | 0.908 | 0.003 |
| p.Thr136Gly | substitution | 136 | Thr | Gly | 0.776 | 0.005 |
| p.Thr136His | substitution | 136 | Thr | His | 0.976 | 0.015 |
| p.Thr136Ile | substitution | 136 | Thr | Ile | 1.103 | 0.005 |
| p.Thr136Leu | substitution | 136 | Thr | Leu | 0.741 | 0.007 |
| p.Thr136Lys | substitution | 136 | Thr | Lys | 1.004 | 0.013 |
| p.Thr136Met | substitution | 136 | Thr | Met | 1.075 | 0.004 |
| p.Thr136Phe | substitution | 136 | Thr | Phe | 0.640 | 0.023 |
| p.Thr136Pro | substitution | 136 | Thr | Pro | 0.263 | 0.058 |
| p.Thr136Ser | substitution | 136 | Thr | Ser | 1.084 | 0.036 |
| p.Thr136Ter | nonsense | 136 | Thr | Ter | 0.238 | 0.010 |
| p.Thr136Trp | substitution | 136 | Thr | Trp | 1.082 | 0.004 |
| p.Thr136Tyr | substitution | 136 | Thr | Tyr | 0.875 | 0.015 |
| p.Thr136Val | substitution | 136 | Thr | Val | 0.811 | 0.022 |
| p.Thr145= | synonymous | 145 | Thr | NA | 0.691 | 0.018 |
| p.Thr145Ala | substitution | 145 | Thr | Ala | 1.183 | 0.090 |
| p.Thr145Arg | substitution | 145 | Thr | Arg | 0.301 | 0.037 |
| p.Thr145Asn | substitution | 145 | Thr | Asn | 0.227 | 0.075 |

| | | | | | | |
|---|---|---|---|---|---|---|
| p.Thr145Asp | substitution | 145 | Thr | Asp | 0.000 | 0.033 |
| p.Thr145Cys | substitution | 145 | Thr | Cys | 0.275 | 0.044 |
| p.Thr145Gln | substitution | 145 | Thr | Gln | 0.168 | 0.015 |
| p.Thr145Glu | substitution | 145 | Thr | Glu | 0.000 | 0.022 |
| p.Thr145Gly | substitution | 145 | Thr | Gly | 0.483 | 0.047 |
| p.Thr145His | substitution | 145 | Thr | His | 0.131 | 0.067 |
| p.Thr145Ile | substitution | 145 | Thr | Ile | 0.214 | 0.035 |
| p.Thr145Leu | substitution | 145 | Thr | Leu | 0.596 | 0.019 |
| p.Thr145Lys | substitution | 145 | Thr | Lys | 0.000 | 0.010 |
| p.Thr145Met | substitution | 145 | Thr | Met | 0.562 | 0.009 |
| p.Thr145Phe | substitution | 145 | Thr | Phe | 0.805 | 0.020 |
| p.Thr145Pro | substitution | 145 | Thr | Pro | 0.000 | 0.080 |
| p.Thr145Ser | substitution | 145 | Thr | Ser | 0.869 | 0.020 |
| p.Thr145Ter | nonsense | 145 | Thr | Ter | 0.000 | 0.015 |
| p.Thr145Trp | substitution | 145 | Thr | Trp | 0.413 | 0.009 |
| p.Thr145Tyr | substitution | 145 | Thr | Tyr | 0.000 | 0.086 |
| p.Thr145Val | substitution | 145 | Thr | Val | 0.807 | 0.153 |
| p.Thr172= | synonymous | 172 | Thr | NA | 1.017 | 0.004 |
| p.Thr172Ala | substitution | 172 | Thr | Ala | 1.001 | 0.017 |
| p.Thr172Arg | substitution | 172 | Thr | Arg | 0.781 | 0.004 |
| p.Thr172Asn | substitution | 172 | Thr | Asn | 0.960 | 0.007 |
| p.Thr172Asp | substitution | 172 | Thr | Asp | 0.000 | 0.031 |
| p.Thr172Cys | substitution | 172 | Thr | Cys | 0.650 | 0.008 |
| p.Thr172Gln | substitution | 172 | Thr | Gln | 0.000 | 0.014 |
| p.Thr172Glu | substitution | 172 | Thr | Glu | 0.000 | 0.010 |
| p.Thr172Gly | substitution | 172 | Thr | Gly | 0.763 | 0.011 |
| p.Thr172His | substitution | 172 | Thr | His | 0.471 | 0.012 |
| p.Thr172Ile | substitution | 172 | Thr | Ile | 0.604 | 0.005 |
| p.Thr172Leu | substitution | 172 | Thr | Leu | 0.845 | 0.009 |
| p.Thr172Lys | substitution | 172 | Thr | Lys | 0.691 | 0.014 |
| p.Thr172Met | substitution | 172 | Thr | Met | 0.326 | 0.011 |
| p.Thr172Phe | substitution | 172 | Thr | Phe | 0.772 | 0.012 |
| p.Thr172Pro | substitution | 172 | Thr | Pro | 0.168 | 0.020 |
| p.Thr172Ser | substitution | 172 | Thr | Ser | 0.774 | 0.010 |
| p.Thr172Ter | nonsense | 172 | Thr | Ter | 0.000 | 0.015 |
| p.Thr172Trp | substitution | 172 | Thr | Trp | 0.201 | 0.006 |
| p.Thr172Tyr | substitution | 172 | Thr | Tyr | 0.766 | 0.011 |
| p.Thr172Val | substitution | 172 | Thr | Val | 0.444 | 0.063 |
| p.Thr190= | synonymous | 190 | Thr | NA | 1.090 | 0.014 |
| p.Thr190Ala | substitution | 190 | Thr | Ala | 1.024 | 0.037 |
| p.Thr190Arg | substitution | 190 | Thr | Arg | 0.560 | 0.072 |
| p.Thr190Asn | substitution | 190 | Thr | Asn | 1.316 | 0.002 |
| p.Thr190Asp | substitution | 190 | Thr | Asp | 0.699 | 0.007 |

| | | | | | | |
|---|---|---|---|---|---|---|
| p.Thr190Cys | substitution | 190 | Thr | Cys | 1.129 | 0.012 |
| p.Thr190Gln | substitution | 190 | Thr | Gln | 1.597 | 0.101 |
| p.Thr190Glu | substitution | 190 | Thr | Glu | 0.083 | 0.049 |
| p.Thr190Gly | substitution | 190 | Thr | Gly | 0.879 | 0.007 |
| p.Thr190His | substitution | 190 | Thr | His | 1.215 | 0.003 |
| p.Thr190Ile | substitution | 190 | Thr | Ile | 1.243 | 0.009 |
| p.Thr190Leu | substitution | 190 | Thr | Leu | 1.003 | 0.018 |
| p.Thr190Lys | substitution | 190 | Thr | Lys | 0.839 | 0.005 |
| p.Thr190Met | substitution | 190 | Thr | Met | 0.033 | 0.013 |
| p.Thr190Phe | substitution | 190 | Thr | Phe | 0.000 | 0.171 |
| p.Thr190Pro | substitution | 190 | Thr | Pro | 0.173 | 0.051 |
| p.Thr190Ser | substitution | 190 | Thr | Ser | 0.149 | 0.173 |
| p.Thr190Ter | nonsense | 190 | Thr | Ter | 0.000 | 0.109 |
| p.Thr190Trp | substitution | 190 | Thr | Trp | 1.093 | 0.015 |
| p.Thr190Tyr | substitution | 190 | Thr | Tyr | 0.513 | 0.004 |
| p.Thr190Val | substitution | 190 | Thr | Val | 0.896 | 0.004 |
| p.Thr243= | synonymous | 243 | Thr | NA | 1.235 | 0.040 |
| p.Thr243Ala | substitution | 243 | Thr | Ala | 0.000 | 0.180 |
| p.Thr243Arg | substitution | 243 | Thr | Arg | 0.353 | 0.053 |
| p.Thr243Asn | substitution | 243 | Thr | Asn | 0.799 | 0.006 |
| p.Thr243Cys | substitution | 243 | Thr | Cys | 0.000 | 0.057 |
| p.Thr243Gln | substitution | 243 | Thr | Gln | 1.302 | 0.005 |
| p.Thr243Glu | substitution | 243 | Thr | Glu | 0.312 | 0.017 |
| p.Thr243Gly | substitution | 243 | Thr | Gly | 0.903 | 0.003 |
| p.Thr243His | substitution | 243 | Thr | His | 0.847 | 0.004 |
| p.Thr243Ile | substitution | 243 | Thr | Ile | 0.625 | 0.077 |
| p.Thr243Leu | substitution | 243 | Thr | Leu | 0.000 | 0.048 |
| p.Thr243Lys | substitution | 243 | Thr | Lys | 0.464 | 0.129 |
| p.Thr243Met | substitution | 243 | Thr | Met | 0.000 | 0.119 |
| p.Thr243Phe | substitution | 243 | Thr | Phe | 0.000 | 0.108 |
| p.Thr243Pro | substitution | 243 | Thr | Pro | 0.337 | 0.030 |
| p.Thr243Ser | substitution | 243 | Thr | Ser | 0.862 | 0.082 |
| p.Thr243Ter | nonsense | 243 | Thr | Ter | 0.000 | 0.167 |
| p.Thr243Tyr | substitution | 243 | Thr | Tyr | 0.000 | 0.036 |
| p.Thr243Val | substitution | 243 | Thr | Val | 0.000 | 0.051 |
| p.Thr25= | synonymous | 25 | Thr | NA | 0.859 | 0.003 |
| p.Thr25Ala | substitution | 25 | Thr | Ala | 0.857 | 0.006 |
| p.Thr25Arg | substitution | 25 | Thr | Arg | 0.678 | 0.022 |
| p.Thr25Asn | substitution | 25 | Thr | Asn | 0.385 | 0.005 |
| p.Thr25Asp | substitution | 25 | Thr | Asp | 0.000 | 0.004 |
| p.Thr25Cys | substitution | 25 | Thr | Cys | 0.465 | 0.001 |
| p.Thr25Gln | substitution | 25 | Thr | Gln | 0.032 | 0.003 |
| p.Thr25Glu | substitution | 25 | Thr | Glu | 0.746 | 0.001 |

| | | | | | | |
|---|---|---|---|---|---|---|
| p.Thr25Gly | substitution | 25 | Thr | Gly | 0.304 | 0.005 |
| p.Thr25His | substitution | 25 | Thr | His | 0.843 | 0.001 |
| p.Thr25Ile | substitution | 25 | Thr | Ile | 0.340 | 0.038 |
| p.Thr25Leu | substitution | 25 | Thr | Leu | 0.000 | 0.048 |
| p.Thr25Lys | substitution | 25 | Thr | Lys | 0.320 | 0.001 |
| p.Thr25Met | substitution | 25 | Thr | Met | 0.566 | 0.000 |
| p.Thr25Phe | substitution | 25 | Thr | Phe | 0.236 | 0.001 |
| p.Thr25Pro | substitution | 25 | Thr | Pro | 0.000 | 0.002 |
| p.Thr25Ser | substitution | 25 | Thr | Ser | 0.881 | 0.001 |
| p.Thr25Ter | nonsense | 25 | Thr | Ter | 0.000 | 0.061 |
| p.Thr25Trp | substitution | 25 | Thr | Trp | 0.000 | 0.005 |
| p.Thr25Tyr | substitution | 25 | Thr | Tyr | 0.000 | 0.002 |
| p.Thr25Val | substitution | 25 | Thr | Val | 0.381 | 0.001 |
| p.Thr269= | synonymous | 269 | Thr | NA | 1.124 | 0.028 |
| p.Thr269Ala | substitution | 269 | Thr | Ala | 0.725 | 0.005 |
| p.Thr269Arg | substitution | 269 | Thr | Arg | 0.000 | 0.078 |
| p.Thr269Asn | substitution | 269 | Thr | Asn | 0.454 | 0.007 |
| p.Thr269Cys | substitution | 269 | Thr | Cys | 1.248 | 0.005 |
| p.Thr269Gln | substitution | 269 | Thr | Gln | 0.000 | 0.040 |
| p.Thr269Glu | substitution | 269 | Thr | Glu | 0.000 | 0.027 |
| p.Thr269Gly | substitution | 269 | Thr | Gly | 0.000 | 0.016 |
| p.Thr269Ile | substitution | 269 | Thr | Ile | 0.000 | 0.176 |
| p.Thr269Leu | substitution | 269 | Thr | Leu | 0.022 | 0.021 |
| p.Thr269Lys | substitution | 269 | Thr | Lys | 0.000 | 0.209 |
| p.Thr269Met | substitution | 269 | Thr | Met | 0.261 | 0.006 |
| p.Thr269Phe | substitution | 269 | Thr | Phe | 0.000 | 0.015 |
| p.Thr269Pro | substitution | 269 | Thr | Pro | 0.000 | 0.146 |
| p.Thr269Ser | substitution | 269 | Thr | Ser | 0.094 | 0.022 |
| p.Thr269Ter | nonsense | 269 | Thr | Ter | 0.034 | 0.033 |
| p.Thr269Tyr | substitution | 269 | Thr | Tyr | 0.000 | 0.019 |
| p.Thr269Val | substitution | 269 | Thr | Val | 0.000 | 0.027 |
| p.Thr279= | synonymous | 279 | Thr | NA | 1.101 | 0.028 |
| p.Thr279Ala | substitution | 279 | Thr | Ala | 0.998 | 0.033 |
| p.Thr279Arg | substitution | 279 | Thr | Arg | 0.392 | 0.006 |
| p.Thr279Asn | substitution | 279 | Thr | Asn | 1.075 | 0.005 |
| p.Thr279Asp | substitution | 279 | Thr | Asp | 0.208 | 0.006 |
| p.Thr279Cys | substitution | 279 | Thr | Cys | 0.000 | 0.006 |
| p.Thr279Gln | substitution | 279 | Thr | Gln | 0.691 | 0.005 |
| p.Thr279Glu | substitution | 279 | Thr | Glu | 0.587 | 0.008 |
| p.Thr279Gly | substitution | 279 | Thr | Gly | 0.767 | 0.004 |
| p.Thr279His | substitution | 279 | Thr | His | 0.337 | 0.005 |
| p.Thr279Ile | substitution | 279 | Thr | Ile | 1.016 | 0.006 |
| p.Thr279Leu | substitution | 279 | Thr | Leu | 0.401 | 0.007 |

| | | | | | | | |
|---|---|---|---|---|---|---|---|
| p.Thr279Lys | substitution | 279 | Thr | Lys | | 0.187 | 0.008 |
| p.Thr279Met | substitution | 279 | Thr | Met | | 0.269 | 0.004 |
| p.Thr279Phe | substitution | 279 | Thr | Phe | | 0.749 | 0.004 |
| p.Thr279Pro | substitution | 279 | Thr | Pro | | 0.079 | 0.019 |
| p.Thr279Ser | substitution | 279 | Thr | Ser | | 0.824 | 0.010 |
| p.Thr279Ter | nonsense | 279 | Thr | Ter | | 0.000 | 0.006 |
| p.Thr279Trp | substitution | 279 | Thr | Trp | | 0.237 | 0.014 |
| p.Thr279Tyr | substitution | 279 | Thr | Tyr | | 0.916 | 0.004 |
| p.Thr279Val | substitution | 279 | Thr | Val | | 0.602 | 0.008 |
| p.Thr294= | synonymous | 294 | Thr | NA | | 0.631 | 0.066 |
| p.Thr294Ala | substitution | 294 | Thr | Ala | | 0.545 | 0.141 |
| p.Thr294Arg | substitution | 294 | Thr | Arg | | 0.786 | 0.005 |
| p.Thr294Asn | substitution | 294 | Thr | Asn | | 1.146 | 0.015 |
| p.Thr294Asp | substitution | 294 | Thr | Asp | | 0.000 | 0.110 |
| p.Thr294Cys | substitution | 294 | Thr | Cys | | 0.000 | 0.121 |
| p.Thr294Gln | substitution | 294 | Thr | Gln | | 0.000 | 0.109 |
| p.Thr294Glu | substitution | 294 | Thr | Glu | | 0.000 | 0.075 |
| p.Thr294Gly | substitution | 294 | Thr | Gly | | 1.155 | 0.007 |
| p.Thr294His | substitution | 294 | Thr | His | | 1.547 | 0.011 |
| p.Thr294Ile | substitution | 294 | Thr | Ile | | 1.073 | 0.004 |
| p.Thr294Leu | substitution | 294 | Thr | Leu | | 0.703 | 0.067 |
| p.Thr294Lys | substitution | 294 | Thr | Lys | | 0.011 | 0.003 |
| p.Thr294Phe | substitution | 294 | Thr | Phe | | 0.752 | 0.019 |
| p.Thr294Pro | substitution | 294 | Thr | Pro | | 1.429 | 0.016 |
| p.Thr294Ser | substitution | 294 | Thr | Ser | | 1.060 | 0.029 |
| p.Thr294Ter | nonsense | 294 | Thr | Ter | | 0.000 | 0.059 |
| p.Thr294Tyr | substitution | 294 | Thr | Tyr | | 0.000 | 0.202 |
| p.Thr294Val | substitution | 294 | Thr | Val | | 0.951 | 0.245 |
| p.Thr298= | synonymous | 298 | Thr | NA | | 1.086 | 0.047 |
| p.Thr298Ala | substitution | 298 | Thr | Ala | | 0.676 | 0.155 |
| p.Thr298Arg | substitution | 298 | Thr | Arg | | 1.073 | 0.036 |
| p.Thr298Asn | substitution | 298 | Thr | Asn | | 0.590 | 0.039 |
| p.Thr298Cys | substitution | 298 | Thr | Cys | | 0.607 | 0.009 |
| p.Thr298Gln | substitution | 298 | Thr | Gln | | 0.620 | 0.019 |
| p.Thr298Glu | substitution | 298 | Thr | Glu | | 0.824 | 0.012 |
| p.Thr298Gly | substitution | 298 | Thr | Gly | | 0.047 | 0.052 |
| p.Thr298His | substitution | 298 | Thr | His | | 0.670 | 0.006 |
| p.Thr298Ile | substitution | 298 | Thr | Ile | | 0.193 | 0.136 |
| p.Thr298Leu | substitution | 298 | Thr | Leu | | 0.504 | 0.100 |
| p.Thr298Lys | substitution | 298 | Thr | Lys | | 0.720 | 0.012 |
| p.Thr298Met | substitution | 298 | Thr | Met | | 0.754 | 0.004 |
| p.Thr298Phe | substitution | 298 | Thr | Phe | | 0.480 | 0.057 |
| p.Thr298Pro | substitution | 298 | Thr | Pro | | 0.312 | 0.107 |

| | | | | | | |
|---|---|---|---|---|---|---|
| p.Thr298Ser | substitution | 298 | Thr | Ser | 1.054 | 0.051 |
| p.Thr298Ter | nonsense | 298 | Thr | Ter | 0.000 | 0.101 |
| p.Thr298Trp | substitution | 298 | Thr | Trp | 0.971 | 0.005 |
| p.Thr298Tyr | substitution | 298 | Thr | Tyr | 0.781 | 0.014 |
| p.Thr298Val | substitution | 298 | Thr | Val | 1.011 | 0.035 |
| p.Thr319= | synonymous | 319 | Thr | NA | 0.749 | 0.047 |
| p.Thr319Ala | substitution | 319 | Thr | Ala | 0.663 | 0.031 |
| p.Thr319Arg | substitution | 319 | Thr | Arg | 0.000 | 0.010 |
| p.Thr319Asn | substitution | 319 | Thr | Asn | 0.353 | 0.006 |
| p.Thr319Asp | substitution | 319 | Thr | Asp | 0.542 | 0.006 |
| p.Thr319Cys | substitution | 319 | Thr | Cys | 0.917 | 0.009 |
| p.Thr319Gln | substitution | 319 | Thr | Gln | 0.000 | 0.102 |
| p.Thr319Glu | substitution | 319 | Thr | Glu | 0.174 | 0.017 |
| p.Thr319Gly | substitution | 319 | Thr | Gly | 0.520 | 0.080 |
| p.Thr319His | substitution | 319 | Thr | His | 1.321 | 0.003 |
| p.Thr319Ile | substitution | 319 | Thr | Ile | 0.736 | 0.004 |
| p.Thr319Leu | substitution | 319 | Thr | Leu | 0.000 | 0.027 |
| p.Thr319Lys | substitution | 319 | Thr | Lys | 0.243 | 0.014 |
| p.Thr319Met | substitution | 319 | Thr | Met | 0.000 | 0.019 |
| p.Thr319Phe | substitution | 319 | Thr | Phe | 0.873 | 0.004 |
| p.Thr319Pro | substitution | 319 | Thr | Pro | 0.245 | 0.031 |
| p.Thr319Ser | substitution | 319 | Thr | Ser | 0.870 | 0.062 |
| p.Thr319Ter | nonsense | 319 | Thr | Ter | 0.000 | 0.203 |
| p.Thr319Trp | substitution | 319 | Thr | Trp | 0.000 | 0.016 |
| p.Thr319Tyr | substitution | 319 | Thr | Tyr | 1.194 | 0.007 |
| p.Thr319Val | substitution | 319 | Thr | Val | 0.000 | 0.021 |
| p.Thr35= | synonymous | 35 | Thr | NA | 0.970 | 0.002 |
| p.Thr35Ala | substitution | 35 | Thr | Ala | 0.861 | 0.000 |
| p.Thr35Arg | substitution | 35 | Thr | Arg | 0.000 | 0.001 |
| p.Thr35Asn | substitution | 35 | Thr | Asn | 0.201 | 0.001 |
| p.Thr35Asp | substitution | 35 | Thr | Asp | 0.000 | 0.003 |
| p.Thr35Cys | substitution | 35 | Thr | Cys | 0.624 | 0.000 |
| p.Thr35Gln | substitution | 35 | Thr | Gln | 0.000 | 0.001 |
| p.Thr35Glu | substitution | 35 | Thr | Glu | 0.000 | 0.002 |
| p.Thr35Gly | substitution | 35 | Thr | Gly | 0.838 | 0.001 |
| p.Thr35His | substitution | 35 | Thr | His | 0.000 | 0.005 |
| p.Thr35Ile | substitution | 35 | Thr | Ile | 0.000 | 0.001 |
| p.Thr35Leu | substitution | 35 | Thr | Leu | 0.456 | 0.008 |
| p.Thr35Lys | substitution | 35 | Thr | Lys | 0.000 | 0.021 |
| p.Thr35Met | substitution | 35 | Thr | Met | 0.800 | 0.000 |
| p.Thr35Phe | substitution | 35 | Thr | Phe | 0.000 | 0.002 |
| p.Thr35Pro | substitution | 35 | Thr | Pro | 0.000 | 0.003 |
| p.Thr35Ser | substitution | 35 | Thr | Ser | 0.753 | 0.017 |

| | | | | | | | |
|---|---|---|---|---|---|---|---|
| p.Thr35Ter | nonsense | 35 | Thr | Ter | | 0.000 | 0.002 |
| p.Thr35Trp | substitution | 35 | Thr | Trp | | 0.236 | 0.001 |
| p.Thr35Tyr | substitution | 35 | Thr | Tyr | | 0.000 | 0.002 |
| p.Thr35Val | substitution | 35 | Thr | Val | | 0.762 | 0.002 |
| p.Thr41= | synonymous | 41 | Thr | NA | | 0.891 | 0.024 |
| p.Thr41Ala | substitution | 41 | Thr | Ala | | 0.842 | 0.019 |
| p.Thr41Arg | substitution | 41 | Thr | Arg | | 0.900 | 0.021 |
| p.Thr41Asn | substitution | 41 | Thr | Asn | | 0.741 | 0.004 |
| p.Thr41Asp | substitution | 41 | Thr | Asp | | 0.779 | 0.006 |
| p.Thr41Cys | substitution | 41 | Thr | Cys | | 0.815 | 0.007 |
| p.Thr41Gln | substitution | 41 | Thr | Gln | | 0.593 | 0.038 |
| p.Thr41Glu | substitution | 41 | Thr | Glu | | 0.885 | 0.019 |
| p.Thr41Gly | substitution | 41 | Thr | Gly | | 0.635 | 0.015 |
| p.Thr41His | substitution | 41 | Thr | His | | 0.933 | 0.007 |
| p.Thr41Ile | substitution | 41 | Thr | Ile | | 0.964 | 0.056 |
| p.Thr41Leu | substitution | 41 | Thr | Leu | | 0.735 | 0.008 |
| p.Thr41Lys | substitution | 41 | Thr | Lys | | 0.736 | 0.015 |
| p.Thr41Met | substitution | 41 | Thr | Met | | 1.147 | 0.004 |
| p.Thr41Phe | substitution | 41 | Thr | Phe | | 0.658 | 0.004 |
| p.Thr41Pro | substitution | 41 | Thr | Pro | | 0.434 | 0.022 |
| p.Thr41Ser | substitution | 41 | Thr | Ser | | 0.993 | 0.005 |
| p.Thr41Ter | nonsense | 41 | Thr | Ter | | 0.294 | 0.142 |
| p.Thr41Trp | substitution | 41 | Thr | Trp | | 0.669 | 0.007 |
| p.Thr41Tyr | substitution | 41 | Thr | Tyr | | 0.834 | 0.005 |
| p.Thr41Val | substitution | 41 | Thr | Val | | 0.364 | 0.035 |
| p.Thr58= | synonymous | 58 | Thr | NA | | 1.073 | 0.022 |
| p.Thr58Ala | substitution | 58 | Thr | Ala | | 0.695 | 0.178 |
| p.Thr58Arg | substitution | 58 | Thr | Arg | | 0.000 | 0.019 |
| p.Thr58Asn | substitution | 58 | Thr | Asn | | 0.903 | 0.022 |
| p.Thr58Asp | substitution | 58 | Thr | Asp | | 0.124 | 0.263 |
| p.Thr58Cys | substitution | 58 | Thr | Cys | | 0.974 | 0.016 |
| p.Thr58Gln | substitution | 58 | Thr | Gln | | 0.142 | 0.052 |
| p.Thr58Glu | substitution | 58 | Thr | Glu | | 0.818 | 0.012 |
| p.Thr58Gly | substitution | 58 | Thr | Gly | | 0.000 | 0.343 |
| p.Thr58His | substitution | 58 | Thr | His | | 0.000 | 0.180 |
| p.Thr58Ile | substitution | 58 | Thr | Ile | | 0.338 | 0.124 |
| p.Thr58Leu | substitution | 58 | Thr | Leu | | 0.253 | 0.038 |
| p.Thr58Lys | substitution | 58 | Thr | Lys | | 0.000 | 0.052 |
| p.Thr58Met | substitution | 58 | Thr | Met | | 0.900 | 0.012 |
| p.Thr58Phe | substitution | 58 | Thr | Phe | | 0.000 | 0.141 |
| p.Thr58Pro | substitution | 58 | Thr | Pro | | 0.053 | 0.067 |
| p.Thr58Ser | substitution | 58 | Thr | Ser | | 0.821 | 0.099 |
| p.Thr58Ter | nonsense | 58 | Thr | Ter | | 0.000 | 0.123 |

| | | | | | | | |
|---|---|---|---|---|---|---|---|
| p.Thr58Trp | substitution | 58 | Thr | Trp | | 0.000 | 0.301 |
| p.Thr58Tyr | substitution | 58 | Thr | Tyr | | 0.000 | 0.096 |
| p.Thr58Val | substitution | 58 | Thr | Val | | 0.642 | 0.025 |
| p.Thr59= | synonymous | 59 | Thr | NA | | 1.098 | 0.072 |
| p.Thr59Ala | substitution | 59 | Thr | Ala | | 0.827 | 0.084 |
| p.Thr59Arg | substitution | 59 | Thr | Arg | | 0.619 | 0.044 |
| p.Thr59Asn | substitution | 59 | Thr | Asn | | 0.398 | 0.023 |
| p.Thr59Asp | substitution | 59 | Thr | Asp | | 1.106 | 0.024 |
| p.Thr59Gln | substitution | 59 | Thr | Gln | | 1.014 | 0.032 |
| p.Thr59Gly | substitution | 59 | Thr | Gly | | 0.549 | 0.065 |
| p.Thr59His | substitution | 59 | Thr | His | | 0.825 | 0.017 |
| p.Thr59Ile | substitution | 59 | Thr | Ile | | 0.398 | 0.144 |
| p.Thr59Leu | substitution | 59 | Thr | Leu | | 0.486 | 0.488 |
| p.Thr59Lys | substitution | 59 | Thr | Lys | | 0.599 | 0.122 |
| p.Thr59Met | substitution | 59 | Thr | Met | | 0.986 | 0.014 |
| p.Thr59Phe | substitution | 59 | Thr | Phe | | 1.305 | 0.019 |
| p.Thr59Pro | substitution | 59 | Thr | Pro | | 0.926 | 0.175 |
| p.Thr59Ser | substitution | 59 | Thr | Ser | | 1.118 | 0.017 |
| p.Thr59Ter | nonsense | 59 | Thr | Ter | | 0.307 | 0.783 |
| p.Thr59Trp | substitution | 59 | Thr | Trp | | 0.110 | 0.034 |
| p.Thr59Val | substitution | 59 | Thr | Val | | 0.586 | 0.061 |
| p.Thr66= | synonymous | 66 | Thr | NA | | 0.967 | 0.175 |
| p.Thr66Ala | substitution | 66 | Thr | Ala | | 1.095 | 0.018 |
| p.Thr66Arg | substitution | 66 | Thr | Arg | | 0.761 | 0.006 |
| p.Thr66Asn | substitution | 66 | Thr | Asn | | 0.807 | 0.025 |
| p.Thr66Asp | substitution | 66 | Thr | Asp | | 0.261 | 0.024 |
| p.Thr66Gln | substitution | 66 | Thr | Gln | | 0.463 | 0.026 |
| p.Thr66Glu | substitution | 66 | Thr | Glu | | 1.107 | 0.011 |
| p.Thr66Gly | substitution | 66 | Thr | Gly | | 0.913 | 0.020 |
| p.Thr66His | substitution | 66 | Thr | His | | 0.658 | 0.012 |
| p.Thr66Ile | substitution | 66 | Thr | Ile | | 1.134 | 0.007 |
| p.Thr66Leu | substitution | 66 | Thr | Leu | | 0.759 | 0.024 |
| p.Thr66Lys | substitution | 66 | Thr | Lys | | 0.000 | 0.030 |
| p.Thr66Met | substitution | 66 | Thr | Met | | 0.838 | 0.007 |
| p.Thr66Phe | substitution | 66 | Thr | Phe | | 0.731 | 0.008 |
| p.Thr66Pro | substitution | 66 | Thr | Pro | | 0.962 | 0.051 |
| p.Thr66Ser | substitution | 66 | Thr | Ser | | 1.057 | 0.014 |
| p.Thr66Ter | nonsense | 66 | Thr | Ter | | 0.218 | 0.015 |
| p.Thr66Trp | substitution | 66 | Thr | Trp | | 0.688 | 0.014 |
| p.Thr66Tyr | substitution | 66 | Thr | Tyr | | 0.395 | 0.014 |
| p.Thr66Val | substitution | 66 | Thr | Val | | 1.189 | 0.011 |
| p.Thr78= | synonymous | 78 | Thr | NA | | 0.996 | 0.014 |
| p.Thr78Ala | substitution | 78 | Thr | Ala | | 0.505 | 0.004 |

| | | | | | | |
|---|---|---|---|---|---|---|
| p.Thr78Arg | substitution | 78 | Thr | Arg | 0.000 | 0.012 |
| p.Thr78Asn | substitution | 78 | Thr | Asn | 0.780 | 0.033 |
| p.Thr78Asp | substitution | 78 | Thr | Asp | 0.376 | 0.017 |
| p.Thr78Cys | substitution | 78 | Thr | Cys | 1.101 | 0.014 |
| p.Thr78Gln | substitution | 78 | Thr | Gln | 0.764 | 0.019 |
| p.Thr78Glu | substitution | 78 | Thr | Glu | 0.000 | 0.026 |
| p.Thr78Gly | substitution | 78 | Thr | Gly | 0.901 | 0.005 |
| p.Thr78His | substitution | 78 | Thr | His | 0.263 | 0.056 |
| p.Thr78Ile | substitution | 78 | Thr | Ile | 1.176 | 0.007 |
| p.Thr78Leu | substitution | 78 | Thr | Leu | 0.778 | 0.097 |
| p.Thr78Lys | substitution | 78 | Thr | Lys | 0.961 | 0.014 |
| p.Thr78Met | substitution | 78 | Thr | Met | 0.141 | 0.026 |
| p.Thr78Phe | substitution | 78 | Thr | Phe | 0.000 | 0.093 |
| p.Thr78Pro | substitution | 78 | Thr | Pro | 0.000 | 0.036 |
| p.Thr78Ser | substitution | 78 | Thr | Ser | 0.635 | 0.006 |
| p.Thr78Ter | nonsense | 78 | Thr | Ter | 0.000 | 0.025 |
| p.Thr78Trp | substitution | 78 | Thr | Trp | 0.117 | 0.145 |
| p.Thr78Tyr | substitution | 78 | Thr | Tyr | 0.403 | 0.026 |
| p.Thr78Val | substitution | 78 | Thr | Val | 0.798 | 0.010 |
| p.Trp198Ala | substitution | 198 | Trp | Ala | 1.033 | 0.003 |
| p.Trp198Arg | substitution | 198 | Trp | Arg | 0.677 | 0.010 |
| p.Trp198Asn | substitution | 198 | Trp | Asn | 0.989 | 0.001 |
| p.Trp198Asp | substitution | 198 | Trp | Asp | 0.688 | 0.002 |
| p.Trp198Cys | substitution | 198 | Trp | Cys | 1.038 | 0.013 |
| p.Trp198Gln | substitution | 198 | Trp | Gln | 0.485 | 0.003 |
| p.Trp198Glu | substitution | 198 | Trp | Glu | 0.982 | 0.001 |
| p.Trp198Gly | substitution | 198 | Trp | Gly | 0.653 | 0.018 |
| p.Trp198His | substitution | 198 | Trp | His | 0.851 | 0.003 |
| p.Trp198Ile | substitution | 198 | Trp | Ile | 0.579 | 0.023 |
| p.Trp198Leu | substitution | 198 | Trp | Leu | 0.963 | 0.004 |
| p.Trp198Lys | substitution | 198 | Trp | Lys | 0.887 | 0.002 |
| p.Trp198Met | substitution | 198 | Trp | Met | 1.224 | 0.001 |
| p.Trp198Phe | substitution | 198 | Trp | Phe | 0.882 | 0.002 |
| p.Trp198Pro | substitution | 198 | Trp | Pro | 0.422 | 0.067 |
| p.Trp198Ser | substitution | 198 | Trp | Ser | 0.785 | 0.038 |
| p.Trp198Ter | nonsense | 198 | Trp | Ter | 0.000 | 0.080 |
| p.Trp198Thr | substitution | 198 | Trp | Thr | 0.371 | 0.052 |
| p.Trp198Tyr | substitution | 198 | Trp | Tyr | 0.880 | 0.001 |
| p.Trp198Val | substitution | 198 | Trp | Val | 0.778 | 0.004 |
| p.Trp283Ala | substitution | 283 | Trp | Ala | 0.093 | 0.173 |
| p.Trp283Arg | substitution | 283 | Trp | Arg | 0.568 | 0.022 |
| p.Trp283Asn | substitution | 283 | Trp | Asn | 0.965 | 0.005 |
| p.Trp283Cys | substitution | 283 | Trp | Cys | 0.515 | 0.008 |

| | | | | | | |
|---|---|---|---|---|---|---|
| p.Trp283Gln | substitution | 283 | Trp | Gln | 0.873 | 0.011 |
| p.Trp283Glu | substitution | 283 | Trp | Glu | 0.000 | 0.025 |
| p.Trp283Gly | substitution | 283 | Trp | Gly | 1.003 | 0.079 |
| p.Trp283Ile | substitution | 283 | Trp | Ile | 0.348 | 0.005 |
| p.Trp283Leu | substitution | 283 | Trp | Leu | 0.174 | 0.133 |
| p.Trp283Lys | substitution | 283 | Trp | Lys | 0.000 | 0.014 |
| p.Trp283Met | substitution | 283 | Trp | Met | 0.000 | 0.022 |
| p.Trp283Phe | substitution | 283 | Trp | Phe | 0.000 | 0.006 |
| p.Trp283Pro | substitution | 283 | Trp | Pro | 0.695 | 0.122 |
| p.Trp283Ser | substitution | 283 | Trp | Ser | 1.036 | 0.006 |
| p.Trp283Ter | nonsense | 283 | Trp | Ter | 0.167 | 0.494 |
| p.Trp283Thr | substitution | 283 | Trp | Thr | 0.360 | 0.052 |
| p.Trp283Tyr | substitution | 283 | Trp | Tyr | 1.037 | 0.003 |
| p.Trp283Val | substitution | 283 | Trp | Val | 0.000 | 0.051 |
| p.Tyr213= | synonymous | 213 | Tyr | NA | 1.185 | 0.619 |
| p.Tyr213Ala | substitution | 213 | Tyr | Ala | 0.000 | 0.031 |
| p.Tyr213Arg | substitution | 213 | Tyr | Arg | 0.000 | 0.024 |
| p.Tyr213Asn | substitution | 213 | Tyr | Asn | 0.819 | 0.006 |
| p.Tyr213Asp | substitution | 213 | Tyr | Asp | 0.000 | 0.014 |
| p.Tyr213Cys | substitution | 213 | Tyr | Cys | 0.500 | 0.018 |
| p.Tyr213Gln | substitution | 213 | Tyr | Gln | 0.418 | 0.168 |
| p.Tyr213Glu | substitution | 213 | Tyr | Glu | 1.083 | 0.004 |
| p.Tyr213Gly | substitution | 213 | Tyr | Gly | 0.731 | 0.034 |
| p.Tyr213His | substitution | 213 | Tyr | His | 1.142 | 0.006 |
| p.Tyr213Ile | substitution | 213 | Tyr | Ile | 0.000 | 0.011 |
| p.Tyr213Leu | substitution | 213 | Tyr | Leu | 0.051 | 0.030 |
| p.Tyr213Lys | substitution | 213 | Tyr | Lys | 0.000 | 0.013 |
| p.Tyr213Phe | substitution | 213 | Tyr | Phe | 0.202 | 0.005 |
| p.Tyr213Pro | substitution | 213 | Tyr | Pro | 0.653 | 0.118 |
| p.Tyr213Ser | substitution | 213 | Tyr | Ser | 0.186 | 0.037 |
| p.Tyr213Ter | nonsense | 213 | Tyr | Ter | 0.625 | 0.113 |
| p.Tyr213Thr | substitution | 213 | Tyr | Thr | 0.000 | 0.006 |
| p.Tyr213Val | substitution | 213 | Tyr | Val | 0.000 | 0.006 |
| p.Tyr277= | synonymous | 277 | Tyr | NA | 1.076 | 0.012 |
| p.Tyr277Ala | substitution | 277 | Tyr | Ala | 1.008 | 0.015 |
| p.Tyr277Arg | substitution | 277 | Tyr | Arg | 0.919 | 0.077 |
| p.Tyr277Asn | substitution | 277 | Tyr | Asn | 0.794 | 0.042 |
| p.Tyr277Asp | substitution | 277 | Tyr | Asp | 0.593 | 0.027 |
| p.Tyr277Cys | substitution | 277 | Tyr | Cys | 0.634 | 0.006 |
| p.Tyr277Gln | substitution | 277 | Tyr | Gln | 0.000 | 0.036 |
| p.Tyr277Glu | substitution | 277 | Tyr | Glu | 0.068 | 0.011 |
| p.Tyr277Gly | substitution | 277 | Tyr | Gly | 0.119 | 0.031 |
| p.Tyr277His | substitution | 277 | Tyr | His | 1.419 | 0.726 |

| | | | | | | |
|---|---|---|---|---|---|---|
| p.Tyr277Ile | substitution | 277 | Tyr | Ile | 0.011 | 0.143 |
| p.Tyr277Leu | substitution | 277 | Tyr | Leu | 0.949 | 0.100 |
| p.Tyr277Lys | substitution | 277 | Tyr | Lys | 0.222 | 0.007 |
| p.Tyr277Met | substitution | 277 | Tyr | Met | 0.909 | 0.004 |
| p.Tyr277Phe | substitution | 277 | Tyr | Phe | 1.166 | 0.416 |
| p.Tyr277Pro | substitution | 277 | Tyr | Pro | 0.000 | 0.009 |
| p.Tyr277Ser | substitution | 277 | Tyr | Ser | 0.674 | 0.162 |
| p.Tyr277Ter | nonsense | 277 | Tyr | Ter | 0.110 | 0.034 |
| p.Tyr277Thr | substitution | 277 | Tyr | Thr | 0.396 | 0.054 |
| p.Tyr277Trp | substitution | 277 | Tyr | Trp | 0.000 | 0.024 |
| p.Tyr277Val | substitution | 277 | Tyr | Val | 0.126 | 0.025 |
| p.Tyr46= | synonymous | 46 | Tyr | NA | 1.031 | 0.002 |
| p.Tyr46Ala | substitution | 46 | Tyr | Ala | 0.606 | 0.046 |
| p.Tyr46Arg | substitution | 46 | Tyr | Arg | 0.632 | 0.021 |
| p.Tyr46Asn | substitution | 46 | Tyr | Asn | 1.126 | 0.006 |
| p.Tyr46Asp | substitution | 46 | Tyr | Asp | 0.795 | 0.012 |
| p.Tyr46Cys | substitution | 46 | Tyr | Cys | 0.754 | 0.026 |
| p.Tyr46Gly | substitution | 46 | Tyr | Gly | 0.467 | 0.007 |
| p.Tyr46His | substitution | 46 | Tyr | His | 1.018 | 0.013 |
| p.Tyr46Ile | substitution | 46 | Tyr | Ile | 0.676 | 0.026 |
| p.Tyr46Leu | substitution | 46 | Tyr | Leu | 0.605 | 0.004 |
| p.Tyr46Lys | substitution | 46 | Tyr | Lys | 0.775 | 0.006 |
| p.Tyr46Met | substitution | 46 | Tyr | Met | 1.024 | 0.003 |
| p.Tyr46Phe | substitution | 46 | Tyr | Phe | 0.960 | 0.058 |
| p.Tyr46Pro | substitution | 46 | Tyr | Pro | 0.000 | 0.057 |
| p.Tyr46Ser | substitution | 46 | Tyr | Ser | 0.700 | 0.010 |
| p.Tyr46Ter | nonsense | 46 | Tyr | Ter | 0.471 | 0.024 |
| p.Tyr46Thr | substitution | 46 | Tyr | Thr | 0.784 | 0.014 |
| p.Tyr46Trp | substitution | 46 | Tyr | Trp | 1.016 | 0.002 |
| p.Tyr46Val | substitution | 46 | Tyr | Val | 1.094 | 0.112 |
| p.Val103= | synonymous | 103 | Val | NA | 1.126 | 0.008 |
| p.Val103Ala | substitution | 103 | Val | Ala | 0.578 | 0.044 |
| p.Val103Arg | substitution | 103 | Val | Arg | 0.892 | 0.077 |
| p.Val103Asn | substitution | 103 | Val | Asn | 0.756 | 0.032 |
| p.Val103Asp | substitution | 103 | Val | Asp | 0.784 | 0.013 |
| p.Val103Cys | substitution | 103 | Val | Cys | 1.423 | 0.006 |
| p.Val103Gln | substitution | 103 | Val | Gln | 1.233 | 0.016 |
| p.Val103Glu | substitution | 103 | Val | Glu | 0.762 | 0.008 |
| p.Val103Gly | substitution | 103 | Val | Gly | 0.918 | 0.009 |
| p.Val103His | substitution | 103 | Val | His | 0.000 | 0.323 |
| p.Val103Ile | substitution | 103 | Val | Ile | 0.659 | 0.044 |
| p.Val103Leu | substitution | 103 | Val | Leu | 0.635 | 0.025 |
| p.Val103Lys | substitution | 103 | Val | Lys | 1.016 | 0.011 |

| | | | | | | | |
|---|---|---|---|---|---|---|---|
| p.Val103Met | substitution | 103 | Val | Met | | 0.904 | 0.008 |
| p.Val103Phe | substitution | 103 | Val | Phe | | 0.725 | 0.026 |
| p.Val103Pro | substitution | 103 | Val | Pro | | 0.885 | 0.050 |
| p.Val103Ser | substitution | 103 | Val | Ser | | 0.841 | 0.036 |
| p.Val103Ter | nonsense | 103 | Val | Ter | | 0.000 | 0.023 |
| p.Val103Thr | substitution | 103 | Val | Thr | | 1.224 | 0.279 |
| p.Val103Trp | substitution | 103 | Val | Trp | | 1.203 | 0.016 |
| p.Val103Tyr | substitution | 103 | Val | Tyr | | 0.000 | 0.130 |
| p.Val123= | synonymous | 123 | Val | NA | | 1.050 | 0.018 |
| p.Val123Ala | substitution | 123 | Val | Ala | | 0.885 | 0.007 |
| p.Val123Arg | substitution | 123 | Val | Arg | | 0.000 | 0.217 |
| p.Val123Asn | substitution | 123 | Val | Asn | | 0.000 | 0.145 |
| p.Val123Asp | substitution | 123 | Val | Asp | | 0.338 | 0.342 |
| p.Val123Cys | substitution | 123 | Val | Cys | | 0.523 | 0.016 |
| p.Val123Gln | substitution | 123 | Val | Gln | | 0.036 | 0.038 |
| p.Val123Glu | substitution | 123 | Val | Glu | | 0.000 | 0.019 |
| p.Val123Gly | substitution | 123 | Val | Gly | | 0.540 | 0.015 |
| p.Val123His | substitution | 123 | Val | His | | 0.011 | 0.068 |
| p.Val123Ile | substitution | 123 | Val | Ile | | 1.037 | 0.006 |
| p.Val123Leu | substitution | 123 | Val | Leu | | 0.888 | 0.067 |
| p.Val123Lys | substitution | 123 | Val | Lys | | 0.000 | 0.137 |
| p.Val123Phe | substitution | 123 | Val | Phe | | 0.121 | 0.099 |
| p.Val123Pro | substitution | 123 | Val | Pro | | 0.303 | 0.018 |
| p.Val123Ser | substitution | 123 | Val | Ser | | 0.685 | 0.006 |
| p.Val123Ter | nonsense | 123 | Val | Ter | | 0.000 | 0.039 |
| p.Val123Thr | substitution | 123 | Val | Thr | | 1.094 | 0.008 |
| p.Val123Trp | substitution | 123 | Val | Trp | | 0.000 | 0.113 |
| p.Val123Tyr | substitution | 123 | Val | Tyr | | 0.000 | 0.052 |
| p.Val124= | synonymous | 124 | Val | NA | | 1.053 | 0.004 |
| p.Val124Ala | substitution | 124 | Val | Ala | | 0.919 | 0.008 |
| p.Val124Arg | substitution | 124 | Val | Arg | | 0.073 | 0.011 |
| p.Val124Asn | substitution | 124 | Val | Asn | | 0.090 | 0.017 |
| p.Val124Asp | substitution | 124 | Val | Asp | | 0.000 | 0.023 |
| p.Val124Cys | substitution | 124 | Val | Cys | | 0.696 | 0.010 |
| p.Val124Gln | substitution | 124 | Val | Gln | | 0.140 | 0.030 |
| p.Val124Glu | substitution | 124 | Val | Glu | | 0.161 | 0.006 |
| p.Val124Gly | substitution | 124 | Val | Gly | | 0.299 | 0.028 |
| p.Val124His | substitution | 124 | Val | His | | 0.000 | 0.069 |
| p.Val124Ile | substitution | 124 | Val | Ile | | 0.893 | 0.032 |
| p.Val124Leu | substitution | 124 | Val | Leu | | 0.771 | 0.024 |
| p.Val124Lys | substitution | 124 | Val | Lys | | 0.251 | 0.016 |
| p.Val124Met | substitution | 124 | Val | Met | | 1.006 | 0.004 |
| p.Val124Phe | substitution | 124 | Val | Phe | | 0.493 | 0.008 |

| | | | | | | |
|---|---|---|---|---|---|---|
| p.Val124Pro | substitution | 124 | Val | Pro | 0.011 | 0.010 |
| p.Val124Ser | substitution | 124 | Val | Ser | 0.486 | 0.019 |
| p.Val124Ter | nonsense | 124 | Val | Ter | 0.000 | 0.014 |
| p.Val124Thr | substitution | 124 | Val | Thr | 0.401 | 0.023 |
| p.Val124Trp | substitution | 124 | Val | Trp | 0.273 | 0.013 |
| p.Val124Tyr | substitution | 124 | Val | Tyr | 0.000 | 0.045 |
| p.Val130= | synonymous | 130 | Val | NA | 0.861 | 0.042 |
| p.Val130Ala | substitution | 130 | Val | Ala | 0.906 | 0.019 |
| p.Val130Arg | substitution | 130 | Val | Arg | 1.159 | 0.027 |
| p.Val130Asn | substitution | 130 | Val | Asn | 0.976 | 0.034 |
| p.Val130Asp | substitution | 130 | Val | Asp | 1.432 | 0.018 |
| p.Val130Cys | substitution | 130 | Val | Cys | 1.136 | 0.010 |
| p.Val130Gln | substitution | 130 | Val | Gln | 0.855 | 0.017 |
| p.Val130Glu | substitution | 130 | Val | Glu | 1.632 | 0.081 |
| p.Val130Gly | substitution | 130 | Val | Gly | 0.543 | 0.056 |
| p.Val130His | substitution | 130 | Val | His | 1.011 | 0.019 |
| p.Val130Ile | substitution | 130 | Val | Ile | 1.073 | 0.009 |
| p.Val130Leu | substitution | 130 | Val | Leu | 0.589 | 0.113 |
| p.Val130Lys | substitution | 130 | Val | Lys | 0.515 | 0.103 |
| p.Val130Met | substitution | 130 | Val | Met | 0.719 | 0.155 |
| p.Val130Phe | substitution | 130 | Val | Phe | 0.668 | 0.015 |
| p.Val130Pro | substitution | 130 | Val | Pro | 0.944 | 0.013 |
| p.Val130Ser | substitution | 130 | Val | Ser | 0.199 | 0.078 |
| p.Val130Ter | nonsense | 130 | Val | Ter | 0.025 | 0.099 |
| p.Val130Thr | substitution | 130 | Val | Thr | 0.721 | 0.014 |
| p.Val130Trp | substitution | 130 | Val | Trp | 0.477 | 0.141 |
| p.Val130Tyr | substitution | 130 | Val | Tyr | 1.110 | 0.021 |
| p.Val142= | synonymous | 142 | Val | NA | 1.184 | 0.015 |
| p.Val142Ala | substitution | 142 | Val | Ala | 0.949 | 0.004 |
| p.Val142Arg | substitution | 142 | Val | Arg | 0.966 | 0.002 |
| p.Val142Asn | substitution | 142 | Val | Asn | 0.885 | 0.005 |
| p.Val142Asp | substitution | 142 | Val | Asp | 0.694 | 0.005 |
| p.Val142Cys | substitution | 142 | Val | Cys | 1.002 | 0.003 |
| p.Val142Gln | substitution | 142 | Val | Gln | 0.449 | 0.005 |
| p.Val142Glu | substitution | 142 | Val | Glu | 0.890 | 0.009 |
| p.Val142Gly | substitution | 142 | Val | Gly | 0.772 | 0.003 |
| p.Val142His | substitution | 142 | Val | His | 0.809 | 0.004 |
| p.Val142Leu | substitution | 142 | Val | Leu | 0.825 | 0.003 |
| p.Val142Lys | substitution | 142 | Val | Lys | 0.829 | 0.004 |
| p.Val142Met | substitution | 142 | Val | Met | 0.824 | 0.004 |
| p.Val142Phe | substitution | 142 | Val | Phe | 0.596 | 0.004 |
| p.Val142Pro | substitution | 142 | Val | Pro | 0.474 | 0.006 |
| p.Val142Ser | substitution | 142 | Val | Ser | 0.812 | 0.006 |

| | | | | | | |
|---|---|---|---|---|---|---|
| p.Val142Ter | nonsense | 142 | Val | Ter | 0.000 | 0.016 |
| p.Val142Thr | substitution | 142 | Val | Thr | 0.836 | 0.004 |
| p.Val142Trp | substitution | 142 | Val | Trp | 0.753 | 0.004 |
| p.Val142Tyr | substitution | 142 | Val | Tyr | 0.840 | 0.003 |
| p.Val143= | synonymous | 143 | Val | NA | 1.054 | 0.005 |
| p.Val143Ala | substitution | 143 | Val | Ala | 1.052 | 0.006 |
| p.Val143Arg | substitution | 143 | Val | Arg | 0.000 | 0.005 |
| p.Val143Asn | substitution | 143 | Val | Asn | 0.016 | 0.009 |
| p.Val143Asp | substitution | 143 | Val | Asp | 0.122 | 0.015 |
| p.Val143Cys | substitution | 143 | Val | Cys | 1.180 | 0.004 |
| p.Val143Gln | substitution | 143 | Val | Gln | 0.141 | 0.012 |
| p.Val143Glu | substitution | 143 | Val | Glu | 0.199 | 0.009 |
| p.Val143Gly | substitution | 143 | Val | Gly | 0.143 | 0.005 |
| p.Val143His | substitution | 143 | Val | His | 0.041 | 0.009 |
| p.Val143Ile | substitution | 143 | Val | Ile | 1.000 | 0.004 |
| p.Val143Leu | substitution | 143 | Val | Leu | 0.857 | 0.001 |
| p.Val143Lys | substitution | 143 | Val | Lys | 0.135 | 0.008 |
| p.Val143Met | substitution | 143 | Val | Met | 0.787 | 0.004 |
| p.Val143Pro | substitution | 143 | Val | Pro | 0.055 | 0.006 |
| p.Val143Ser | substitution | 143 | Val | Ser | 0.013 | 0.006 |
| p.Val143Ter | nonsense | 143 | Val | Ter | 0.000 | 0.006 |
| p.Val143Thr | substitution | 143 | Val | Thr | 0.379 | 0.005 |
| p.Val143Trp | substitution | 143 | Val | Trp | 0.006 | 0.005 |
| p.Val143Tyr | substitution | 143 | Val | Tyr | 0.438 | 0.002 |
| p.Val202= | synonymous | 202 | Val | NA | 0.956 | 0.015 |
| p.Val202Ala | substitution | 202 | Val | Ala | 0.796 | 0.051 |
| p.Val202Arg | substitution | 202 | Val | Arg | 0.724 | 0.088 |
| p.Val202Asn | substitution | 202 | Val | Asn | 0.000 | 0.021 |
| p.Val202Asp | substitution | 202 | Val | Asp | 0.764 | 0.084 |
| p.Val202Cys | substitution | 202 | Val | Cys | 0.956 | 0.003 |
| p.Val202Gln | substitution | 202 | Val | Gln | 0.433 | 0.017 |
| p.Val202Glu | substitution | 202 | Val | Glu | 0.841 | 0.003 |
| p.Val202Gly | substitution | 202 | Val | Gly | 0.776 | 0.001 |
| p.Val202His | substitution | 202 | Val | His | 0.000 | 0.022 |
| p.Val202Leu | substitution | 202 | Val | Leu | 0.904 | 0.018 |
| p.Val202Lys | substitution | 202 | Val | Lys | 0.555 | 0.023 |
| p.Val202Met | substitution | 202 | Val | Met | 0.972 | 0.003 |
| p.Val202Phe | substitution | 202 | Val | Phe | 0.000 | 0.067 |
| p.Val202Pro | substitution | 202 | Val | Pro | 1.366 | 0.002 |
| p.Val202Ser | substitution | 202 | Val | Ser | 0.466 | 0.202 |
| p.Val202Ter | nonsense | 202 | Val | Ter | 0.000 | 0.050 |
| p.Val202Thr | substitution | 202 | Val | Thr | 1.164 | 0.085 |
| p.Val202Trp | substitution | 202 | Val | Trp | 0.278 | 0.010 |

| | | | | | | | |
|---|---|---|---|---|---|---|---|
| p.Val202Tyr | substitution | 202 | Val | Tyr | | 0.637 | 0.012 |
| p.Val20= | synonymous | 20 | Val | NA | | 1.249 | 0.001 |
| p.Val20Ala | substitution | 20 | Val | Ala | | 0.751 | 0.011 |
| p.Val20Arg | substitution | 20 | Val | Arg | | 0.919 | 0.008 |
| p.Val20Asp | substitution | 20 | Val | Asp | | 0.283 | 0.001 |
| p.Val20Gln | substitution | 20 | Val | Gln | | 0.945 | 0.002 |
| p.Val20Glu | substitution | 20 | Val | Glu | | 0.478 | 0.002 |
| p.Val20Gly | substitution | 20 | Val | Gly | | 0.589 | 0.001 |
| p.Val20Ile | substitution | 20 | Val | Ile | | 0.821 | 0.002 |
| p.Val20Leu | substitution | 20 | Val | Leu | | 0.627 | 0.013 |
| p.Val20Lys | substitution | 20 | Val | Lys | | 1.197 | 0.001 |
| p.Val20Met | substitution | 20 | Val | Met | | 0.906 | 0.002 |
| p.Val20Phe | substitution | 20 | Val | Phe | | 0.753 | 0.001 |
| p.Val20Pro | substitution | 20 | Val | Pro | | 0.839 | 0.001 |
| p.Val20Ser | substitution | 20 | Val | Ser | | 1.262 | 0.001 |
| p.Val20Ter | nonsense | 20 | Val | Ter | | 0.000 | 0.006 |
| p.Val20Thr | substitution | 20 | Val | Thr | | 0.967 | 0.001 |
| p.Val20Trp | substitution | 20 | Val | Trp | | 0.653 | 0.001 |
| p.Val215= | synonymous | 215 | Val | NA | | 0.910 | 0.014 |
| p.Val215Ala | substitution | 215 | Val | Ala | | 0.734 | 0.129 |
| p.Val215Arg | substitution | 215 | Val | Arg | | 0.185 | 0.054 |
| p.Val215Asn | substitution | 215 | Val | Asn | | 0.000 | 0.015 |
| p.Val215Asp | substitution | 215 | Val | Asp | | 0.000 | 0.007 |
| p.Val215Cys | substitution | 215 | Val | Cys | | 1.331 | 0.003 |
| p.Val215Glu | substitution | 215 | Val | Glu | | 0.647 | 0.664 |
| p.Val215Gly | substitution | 215 | Val | Gly | | 1.192 | 0.002 |
| p.Val215Ile | substitution | 215 | Val | Ile | | 0.152 | 0.026 |
| p.Val215Leu | substitution | 215 | Val | Leu | | 0.512 | 0.105 |
| p.Val215Lys | substitution | 215 | Val | Lys | | 0.000 | 0.036 |
| p.Val215Met | substitution | 215 | Val | Met | | 0.791 | 0.025 |
| p.Val215Phe | substitution | 215 | Val | Phe | | 0.251 | 0.159 |
| p.Val215Pro | substitution | 215 | Val | Pro | | 0.523 | 0.005 |
| p.Val215Ser | substitution | 215 | Val | Ser | | 1.110 | 0.215 |
| p.Val215Trp | substitution | 215 | Val | Trp | | 0.000 | 0.068 |
| p.Val215Tyr | substitution | 215 | Val | Tyr | | 0.000 | 0.124 |
| p.Val222= | synonymous | 222 | Val | NA | | 0.998 | 0.003 |
| p.Val222Ala | substitution | 222 | Val | Ala | | 0.836 | 0.027 |
| p.Val222Arg | substitution | 222 | Val | Arg | | 0.000 | 0.012 |
| p.Val222Asn | substitution | 222 | Val | Asn | | 0.000 | 0.034 |
| p.Val222Asp | substitution | 222 | Val | Asp | | 0.000 | 0.051 |
| p.Val222Cys | substitution | 222 | Val | Cys | | 0.784 | 0.002 |
| p.Val222Gln | substitution | 222 | Val | Gln | | 0.000 | 0.008 |
| p.Val222Glu | substitution | 222 | Val | Glu | | 0.356 | 0.326 |

| | | | | | | |
|---|---|---|---|---|---|---|
| p.Val222Gly | substitution | 222 | Val | Gly | 0.000 | 0.004 |
| p.Val222His | substitution | 222 | Val | His | 0.000 | 0.073 |
| p.Val222Ile | substitution | 222 | Val | Ile | 1.303 | 0.001 |
| p.Val222Leu | substitution | 222 | Val | Leu | 1.165 | 0.010 |
| p.Val222Lys | substitution | 222 | Val | Lys | 0.000 | 0.006 |
| p.Val222Met | substitution | 222 | Val | Met | 0.657 | 0.006 |
| p.Val222Phe | substitution | 222 | Val | Phe | 0.000 | 0.010 |
| p.Val222Pro | substitution | 222 | Val | Pro | 0.000 | 0.035 |
| p.Val222Ser | substitution | 222 | Val | Ser | 0.000 | 0.010 |
| p.Val222Ter | nonsense | 222 | Val | Ter | 0.000 | 0.078 |
| p.Val222Thr | substitution | 222 | Val | Thr | 0.000 | 0.051 |
| p.Val222Trp | substitution | 222 | Val | Trp | 0.000 | 0.015 |
| p.Val222Tyr | substitution | 222 | Val | Tyr | 0.000 | 0.006 |
| p.Val224= | synonymous | 224 | Val | NA | 0.982 | 0.249 |
| p.Val224Ala | substitution | 224 | Val | Ala | 0.598 | 1.015 |
| p.Val224Arg | substitution | 224 | Val | Arg | 0.046 | 0.082 |
| p.Val224Asp | substitution | 224 | Val | Asp | 0.000 | 0.063 |
| p.Val224Cys | substitution | 224 | Val | Cys | 1.094 | 0.002 |
| p.Val224Glu | substitution | 224 | Val | Glu | 0.296 | 0.016 |
| p.Val224Gly | substitution | 224 | Val | Gly | 0.304 | 0.025 |
| p.Val224His | substitution | 224 | Val | His | 0.000 | 0.129 |
| p.Val224Leu | substitution | 224 | Val | Leu | 1.373 | 0.007 |
| p.Val224Met | substitution | 224 | Val | Met | 1.272 | 0.004 |
| p.Val224Phe | substitution | 224 | Val | Phe | 1.091 | 0.071 |
| p.Val224Pro | substitution | 224 | Val | Pro | 0.250 | 0.105 |
| p.Val224Ser | substitution | 224 | Val | Ser | 1.456 | 0.003 |
| p.Val224Ter | nonsense | 224 | Val | Ter | 0.000 | 0.013 |
| p.Val224Thr | substitution | 224 | Val | Thr | 1.381 | 0.003 |
| p.Val224Trp | substitution | 224 | Val | Trp | 0.059 | 0.011 |
| p.Val224Tyr | substitution | 224 | Val | Tyr | 0.121 | 0.021 |
| p.Val235= | synonymous | 235 | Val | NA | 0.942 | 0.023 |
| p.Val235Ala | substitution | 235 | Val | Ala | 0.067 | 0.088 |
| p.Val235Arg | substitution | 235 | Val | Arg | 0.000 | 0.065 |
| p.Val235Asp | substitution | 235 | Val | Asp | 0.000 | 0.014 |
| p.Val235Cys | substitution | 235 | Val | Cys | 0.000 | 0.004 |
| p.Val235Glu | substitution | 235 | Val | Glu | 0.388 | 0.014 |
| p.Val235Gly | substitution | 235 | Val | Gly | 0.160 | 0.035 |
| p.Val235His | substitution | 235 | Val | His | 0.000 | 0.084 |
| p.Val235Ile | substitution | 235 | Val | Ile | 0.013 | 0.024 |
| p.Val235Leu | substitution | 235 | Val | Leu | 0.790 | 0.016 |
| p.Val235Lys | substitution | 235 | Val | Lys | 0.000 | 0.027 |
| p.Val235Met | substitution | 235 | Val | Met | 0.908 | 0.002 |
| p.Val235Phe | substitution | 235 | Val | Phe | 0.000 | 0.099 |

| | | | | | | |
|---|---|---|---|---|---|---|
| p.Val235Pro | substitution | 235 | Val | Pro | 0.000 | 0.035 |
| p.Val235Ser | substitution | 235 | Val | Ser | 0.000 | 0.039 |
| p.Val235Ter | nonsense | 235 | Val | Ter | 0.000 | 0.070 |
| p.Val235Thr | substitution | 235 | Val | Thr | 0.000 | 0.027 |
| p.Val235Trp | substitution | 235 | Val | Trp | 0.000 | 0.007 |
| p.Val235Tyr | substitution | 235 | Val | Tyr | 0.208 | 0.010 |
| p.Val237= | synonymous | 237 | Val | NA | 0.997 | 0.030 |
| p.Val237Ala | substitution | 237 | Val | Ala | 0.000 | 0.167 |
| p.Val237Arg | substitution | 237 | Val | Arg | 0.815 | 0.015 |
| p.Val237Asp | substitution | 237 | Val | Asp | 0.000 | 0.049 |
| p.Val237Cys | substitution | 237 | Val | Cys | 0.126 | 0.074 |
| p.Val237Gln | substitution | 237 | Val | Gln | 0.128 | 0.036 |
| p.Val237Glu | substitution | 237 | Val | Glu | 0.154 | 0.030 |
| p.Val237Gly | substitution | 237 | Val | Gly | 0.253 | 0.030 |
| p.Val237His | substitution | 237 | Val | His | 0.000 | 0.053 |
| p.Val237Leu | substitution | 237 | Val | Leu | 1.052 | 0.012 |
| p.Val237Met | substitution | 237 | Val | Met | 0.655 | 0.039 |
| p.Val237Phe | substitution | 237 | Val | Phe | 0.553 | 0.081 |
| p.Val237Pro | substitution | 237 | Val | Pro | 1.123 | 0.002 |
| p.Val237Ser | substitution | 237 | Val | Ser | 0.900 | 0.065 |
| p.Val237Ter | nonsense | 237 | Val | Ter | 0.180 | 0.023 |
| p.Val237Trp | substitution | 237 | Val | Trp | 0.000 | 0.009 |
| p.Val237Tyr | substitution | 237 | Val | Tyr | 0.209 | 0.024 |
| p.Val23= | synonymous | 23 | Val | NA | 0.967 | 0.001 |
| p.Val23Ala | substitution | 23 | Val | Ala | 1.271 | 0.032 |
| p.Val23Arg | substitution | 23 | Val | Arg | 0.000 | 0.070 |
| p.Val23Asn | substitution | 23 | Val | Asn | 0.000 | 0.005 |
| p.Val23Asp | substitution | 23 | Val | Asp | 0.000 | 0.009 |
| p.Val23Cys | substitution | 23 | Val | Cys | 0.367 | 0.001 |
| p.Val23Glu | substitution | 23 | Val | Glu | 0.000 | 0.027 |
| p.Val23Gly | substitution | 23 | Val | Gly | 0.000 | 0.012 |
| p.Val23His | substitution | 23 | Val | His | 0.193 | 0.011 |
| p.Val23Ile | substitution | 23 | Val | Ile | 0.122 | 0.003 |
| p.Val23Leu | substitution | 23 | Val | Leu | 0.817 | 0.001 |
| p.Val23Lys | substitution | 23 | Val | Lys | 0.000 | 0.005 |
| p.Val23Met | substitution | 23 | Val | Met | 0.949 | 0.001 |
| p.Val23Phe | substitution | 23 | Val | Phe | 1.032 | 0.001 |
| p.Val23Pro | substitution | 23 | Val | Pro | 0.000 | 0.049 |
| p.Val23Ser | substitution | 23 | Val | Ser | 0.000 | 0.028 |
| p.Val23Ter | nonsense | 23 | Val | Ter | 0.000 | 0.093 |
| p.Val23Thr | substitution | 23 | Val | Thr | 0.055 | 0.051 |
| p.Val23Trp | substitution | 23 | Val | Trp | 0.000 | 0.151 |
| p.Val23Tyr | substitution | 23 | Val | Tyr | 0.237 | 0.006 |

| | | | | | | |
|---|---|---|---|---|---|---|
| p.Val263= | synonymous | 263 | Val | NA | 0.814 | 0.011 |
| p.Val263Ala | substitution | 263 | Val | Ala | 1.358 | 0.004 |
| p.Val263Arg | substitution | 263 | Val | Arg | 0.320 | 0.144 |
| p.Val263Asp | substitution | 263 | Val | Asp | 0.079 | 0.110 |
| p.Val263Gln | substitution | 263 | Val | Gln | 1.479 | 0.005 |
| p.Val263Glu | substitution | 263 | Val | Glu | 1.053 | 0.014 |
| p.Val263Gly | substitution | 263 | Val | Gly | 0.000 | 0.197 |
| p.Val263His | substitution | 263 | Val | His | 0.073 | 0.021 |
| p.Val263Leu | substitution | 263 | Val | Leu | 0.948 | 0.070 |
| p.Val263Lys | substitution | 263 | Val | Lys | 1.412 | 0.005 |
| p.Val263Met | substitution | 263 | Val | Met | 0.463 | 0.048 |
| p.Val263Phe | substitution | 263 | Val | Phe | 0.600 | 0.053 |
| p.Val263Ser | substitution | 263 | Val | Ser | 0.693 | 0.008 |
| p.Val263Ter | nonsense | 263 | Val | Ter | 0.010 | 0.019 |
| p.Val263Trp | substitution | 263 | Val | Trp | 0.000 | 0.021 |
| p.Val265= | synonymous | 265 | Val | NA | 1.039 | 0.159 |
| p.Val265Ala | substitution | 265 | Val | Ala | 1.214 | 0.147 |
| p.Val265Arg | substitution | 265 | Val | Arg | 0.000 | 0.035 |
| p.Val265Asn | substitution | 265 | Val | Asn | 0.000 | 0.012 |
| p.Val265Asp | substitution | 265 | Val | Asp | 0.000 | 0.027 |
| p.Val265Gln | substitution | 265 | Val | Gln | 0.000 | 0.039 |
| p.Val265Gly | substitution | 265 | Val | Gly | 0.213 | 0.012 |
| p.Val265His | substitution | 265 | Val | His | 0.000 | 0.083 |
| p.Val265Ile | substitution | 265 | Val | Ile | 0.872 | 0.170 |
| p.Val265Leu | substitution | 265 | Val | Leu | 0.363 | 0.130 |
| p.Val265Met | substitution | 265 | Val | Met | 0.103 | 0.013 |
| p.Val265Phe | substitution | 265 | Val | Phe | 0.000 | 0.027 |
| p.Val265Pro | substitution | 265 | Val | Pro | 0.315 | 0.013 |
| p.Val265Ser | substitution | 265 | Val | Ser | 0.000 | 0.057 |
| p.Val265Ter | nonsense | 265 | Val | Ter | 0.000 | 0.036 |
| p.Val265Thr | substitution | 265 | Val | Thr | 0.255 | 0.028 |
| p.Val267= | synonymous | 267 | Val | NA | 1.589 | 0.036 |
| p.Val267Ala | substitution | 267 | Val | Ala | 0.331 | 0.074 |
| p.Val267Arg | substitution | 267 | Val | Arg | 0.000 | 0.067 |
| p.Val267Asn | substitution | 267 | Val | Asn | 1.290 | 0.028 |
| p.Val267Asp | substitution | 267 | Val | Asp | 0.000 | 0.140 |
| p.Val267Cys | substitution | 267 | Val | Cys | 1.159 | 0.007 |
| p.Val267Gln | substitution | 267 | Val | Gln | 0.000 | 0.094 |
| p.Val267Glu | substitution | 267 | Val | Glu | 0.263 | 0.367 |
| p.Val267Gly | substitution | 267 | Val | Gly | 0.380 | 0.014 |
| p.Val267Ile | substitution | 267 | Val | Ile | 0.229 | 0.077 |
| p.Val267Leu | substitution | 267 | Val | Leu | 0.848 | 0.021 |
| p.Val267Lys | substitution | 267 | Val | Lys | 0.601 | 0.028 |

| | | | | | | |
|---|---|---|---|---|---|---|
| p.Val267Met | substitution | 267 | Val | Met | 1.328 | 0.051 |
| p.Val267Phe | substitution | 267 | Val | Phe | 1.108 | 0.025 |
| p.Val267Pro | substitution | 267 | Val | Pro | 0.000 | 0.037 |
| p.Val267Ser | substitution | 267 | Val | Ser | 0.277 | 0.048 |
| p.Val267Ter | nonsense | 267 | Val | Ter | 0.000 | 0.007 |
| p.Val267Thr | substitution | 267 | Val | Thr | 1.476 | 0.006 |
| p.Val267Trp | substitution | 267 | Val | Trp | 0.000 | 0.013 |
| p.Val267Tyr | substitution | 267 | Val | Tyr | 0.000 | 0.029 |
| p.Val282= | synonymous | 282 | Val | NA | 0.923 | 0.004 |
| p.Val282Ala | substitution | 282 | Val | Ala | 0.757 | 0.086 |
| p.Val282Arg | substitution | 282 | Val | Arg | 0.000 | 0.017 |
| p.Val282Asn | substitution | 282 | Val | Asn | 0.126 | 0.018 |
| p.Val282Asp | substitution | 282 | Val | Asp | 0.114 | 0.095 |
| p.Val282Cys | substitution | 282 | Val | Cys | 0.622 | 0.021 |
| p.Val282Gln | substitution | 282 | Val | Gln | 0.237 | 0.010 |
| p.Val282Glu | substitution | 282 | Val | Glu | 0.000 | 0.011 |
| p.Val282Gly | substitution | 282 | Val | Gly | 0.547 | 0.041 |
| p.Val282His | substitution | 282 | Val | His | 0.249 | 0.015 |
| p.Val282Ile | substitution | 282 | Val | Ile | 0.401 | 0.015 |
| p.Val282Leu | substitution | 282 | Val | Leu | 0.463 | 0.052 |
| p.Val282Lys | substitution | 282 | Val | Lys | 0.177 | 0.009 |
| p.Val282Met | substitution | 282 | Val | Met | 0.650 | 0.004 |
| p.Val282Phe | substitution | 282 | Val | Phe | 0.742 | 0.016 |
| p.Val282Pro | substitution | 282 | Val | Pro | 0.000 | 0.019 |
| p.Val282Ser | substitution | 282 | Val | Ser | 0.304 | 0.030 |
| p.Val282Ter | nonsense | 282 | Val | Ter | 0.000 | 0.055 |
| p.Val282Thr | substitution | 282 | Val | Thr | 0.593 | 0.125 |
| p.Val282Tyr | substitution | 282 | Val | Tyr | 0.041 | 0.026 |
| p.Val301= | synonymous | 301 | Val | NA | 0.945 | 0.046 |
| p.Val301Ala | substitution | 301 | Val | Ala | 0.953 | 0.046 |
| p.Val301Arg | substitution | 301 | Val | Arg | 0.767 | 0.094 |
| p.Val301Asn | substitution | 301 | Val | Asn | 0.000 | 0.023 |
| p.Val301Asp | substitution | 301 | Val | Asp | 0.941 | 0.011 |
| p.Val301Cys | substitution | 301 | Val | Cys | 0.611 | 0.010 |
| p.Val301Gly | substitution | 301 | Val | Gly | 0.979 | 0.080 |
| p.Val301His | substitution | 301 | Val | His | 0.031 | 0.261 |
| p.Val301Ile | substitution | 301 | Val | Ile | 0.999 | 0.337 |
| p.Val301Leu | substitution | 301 | Val | Leu | 0.000 | 0.334 |
| p.Val301Lys | substitution | 301 | Val | Lys | 0.000 | 0.082 |
| p.Val301Met | substitution | 301 | Val | Met | 0.314 | 0.062 |
| p.Val301Phe | substitution | 301 | Val | Phe | 1.005 | 0.010 |
| p.Val301Pro | substitution | 301 | Val | Pro | 0.000 | 0.026 |
| p.Val301Ser | substitution | 301 | Val | Ser | 0.480 | 0.129 |

| | | | | | | |
|---|---|---|---|---|---|---|
| p.Val301Thr | substitution | 301 | Val | Thr | 0.000 | 0.034 |
| p.Val301Tyr | substitution | 301 | Val | Tyr | 0.712 | 0.010 |
| p.Val316= | synonymous | 316 | Val | NA | 0.900 | 0.029 |
| p.Val316Ala | substitution | 316 | Val | Ala | 0.857 | 0.011 |
| p.Val316Arg | substitution | 316 | Val | Arg | 0.987 | 0.184 |
| p.Val316Asn | substitution | 316 | Val | Asn | 0.000 | 0.029 |
| p.Val316Asp | substitution | 316 | Val | Asp | 0.825 | 0.006 |
| p.Val316Gln | substitution | 316 | Val | Gln | 0.000 | 0.069 |
| p.Val316Glu | substitution | 316 | Val | Glu | 0.290 | 0.199 |
| p.Val316Gly | substitution | 316 | Val | Gly | 0.677 | 0.034 |
| p.Val316His | substitution | 316 | Val | His | 0.105 | 0.032 |
| p.Val316Ile | substitution | 316 | Val | Ile | 0.795 | 0.279 |
| p.Val316Leu | substitution | 316 | Val | Leu | 0.994 | 0.029 |
| p.Val316Lys | substitution | 316 | Val | Lys | 0.000 | 0.233 |
| p.Val316Met | substitution | 316 | Val | Met | 0.000 | 0.129 |
| p.Val316Phe | substitution | 316 | Val | Phe | 1.001 | 0.006 |
| p.Val316Pro | substitution | 316 | Val | Pro | 0.386 | 0.026 |
| p.Val316Ser | substitution | 316 | Val | Ser | 0.705 | 0.178 |
| p.Val316Ter | nonsense | 316 | Val | Ter | 0.030 | 0.106 |
| p.Val316Thr | substitution | 316 | Val | Thr | 0.064 | 0.038 |
| p.Val316Tyr | substitution | 316 | Val | Tyr | 0.000 | 0.007 |
| p.Val353= | synonymous | 353 | Val | NA | 1.205 | 0.009 |
| p.Val353Ala | substitution | 353 | Val | Ala | 0.712 | 0.051 |
| p.Val353Arg | substitution | 353 | Val | Arg | 0.476 | 0.017 |
| p.Val353Asn | substitution | 353 | Val | Asn | 1.252 | 0.002 |
| p.Val353Asp | substitution | 353 | Val | Asp | 1.389 | 0.164 |
| p.Val353Cys | substitution | 353 | Val | Cys | 0.000 | 0.072 |
| p.Val353Gln | substitution | 353 | Val | Gln | 1.385 | 0.005 |
| p.Val353Gly | substitution | 353 | Val | Gly | 0.555 | 0.026 |
| p.Val353His | substitution | 353 | Val | His | 1.392 | 0.007 |
| p.Val353Ile | substitution | 353 | Val | Ile | 0.432 | 0.017 |
| p.Val353Leu | substitution | 353 | Val | Leu | 0.463 | 0.040 |
| p.Val353Lys | substitution | 353 | Val | Lys | 0.000 | 0.109 |
| p.Val353Met | substitution | 353 | Val | Met | 1.197 | 0.003 |
| p.Val353Phe | substitution | 353 | Val | Phe | 0.000 | 0.188 |
| p.Val353Pro | substitution | 353 | Val | Pro | 0.613 | 0.231 |
| p.Val353Ser | substitution | 353 | Val | Ser | 0.782 | 0.087 |
| p.Val353Ter | nonsense | 353 | Val | Ter | 0.000 | 0.123 |
| p.Val353Thr | substitution | 353 | Val | Thr | 1.380 | 0.003 |
| p.Val38= | synonymous | 38 | Val | NA | 0.982 | 0.027 |
| p.Val38Ala | substitution | 38 | Val | Ala | 0.691 | 0.035 |
| p.Val38Arg | substitution | 38 | Val | Arg | 0.083 | 0.038 |
| p.Val38Asn | substitution | 38 | Val | Asn | 0.141 | 0.032 |

| | | | | | | |
|---|---|---|---|---|---|---|
| p.Val38Asp | substitution | 38 | Val | Asp | 0.000 | 0.018 |
| p.Val38Cys | substitution | 38 | Val | Cys | 0.700 | 0.009 |
| p.Val38Gln | substitution | 38 | Val | Gln | 0.000 | 0.054 |
| p.Val38Glu | substitution | 38 | Val | Glu | 0.160 | 0.120 |
| p.Val38Gly | substitution | 38 | Val | Gly | 0.000 | 0.012 |
| p.Val38His | substitution | 38 | Val | His | 0.072 | 0.107 |
| p.Val38Ile | substitution | 38 | Val | Ile | 1.199 | 0.008 |
| p.Val38Leu | substitution | 38 | Val | Leu | 0.641 | 0.007 |
| p.Val38Lys | substitution | 38 | Val | Lys | 0.000 | 0.025 |
| p.Val38Met | substitution | 38 | Val | Met | 0.510 | 0.030 |
| p.Val38Phe | substitution | 38 | Val | Phe | 0.334 | 0.011 |
| p.Val38Pro | substitution | 38 | Val | Pro | 0.000 | 0.097 |
| p.Val38Ser | substitution | 38 | Val | Ser | 0.370 | 0.065 |
| p.Val38Ter | nonsense | 38 | Val | Ter | 0.000 | 0.012 |
| p.Val38Thr | substitution | 38 | Val | Thr | 0.970 | 0.005 |
| p.Val38Trp | substitution | 38 | Val | Trp | 0.000 | 0.018 |
| p.Val38Tyr | substitution | 38 | Val | Tyr | 0.000 | 0.078 |
| p.Val39= | synonymous | 39 | Val | NA | 0.874 | 0.019 |
| p.Val39Ala | substitution | 39 | Val | Ala | 1.000 | 0.023 |
| p.Val39Arg | substitution | 39 | Val | Arg | 0.637 | 0.017 |
| p.Val39Asn | substitution | 39 | Val | Asn | 0.681 | 0.009 |
| p.Val39Asp | substitution | 39 | Val | Asp | 0.000 | 0.012 |
| p.Val39Cys | substitution | 39 | Val | Cys | 0.816 | 0.003 |
| p.Val39Gln | substitution | 39 | Val | Gln | 0.603 | 0.006 |
| p.Val39Glu | substitution | 39 | Val | Glu | 0.569 | 0.006 |
| p.Val39Gly | substitution | 39 | Val | Gly | 0.537 | 0.008 |
| p.Val39His | substitution | 39 | Val | His | 0.419 | 0.007 |
| p.Val39Ile | substitution | 39 | Val | Ile | 0.417 | 0.013 |
| p.Val39Leu | substitution | 39 | Val | Leu | 0.726 | 0.003 |
| p.Val39Lys | substitution | 39 | Val | Lys | 0.583 | 0.007 |
| p.Val39Met | substitution | 39 | Val | Met | 0.573 | 0.009 |
| p.Val39Phe | substitution | 39 | Val | Phe | 0.517 | 0.010 |
| p.Val39Pro | substitution | 39 | Val | Pro | 0.000 | 0.013 |
| p.Val39Ser | substitution | 39 | Val | Ser | 0.614 | 0.026 |
| p.Val39Ter | nonsense | 39 | Val | Ter | 0.000 | 0.008 |
| p.Val39Thr | substitution | 39 | Val | Thr | 0.632 | 0.016 |
| p.Val39Trp | substitution | 39 | Val | Trp | 0.445 | 0.005 |
| p.Val39Tyr | substitution | 39 | Val | Tyr | 0.802 | 0.005 |
| p.Val90= | synonymous | 90 | Val | NA | 1.024 | 0.003 |
| p.Val90Ala | substitution | 90 | Val | Ala | 0.510 | 0.015 |
| p.Val90Arg | substitution | 90 | Val | Arg | 0.000 | 0.005 |
| p.Val90Asn | substitution | 90 | Val | Asn | 0.122 | 0.034 |
| p.Val90Asp | substitution | 90 | Val | Asp | 0.000 | 0.022 |

| | | | | | | |
|---|---|---|---|---|---|---|
| p.Val90Cys | substitution | 90 | Val | Cys | 0.862 | 0.004 |
| p.Val90Gln | substitution | 90 | Val | Gln | 0.305 | 0.006 |
| p.Val90Glu | substitution | 90 | Val | Glu | 0.387 | 0.010 |
| p.Val90Gly | substitution | 90 | Val | Gly | 0.032 | 0.016 |
| p.Val90His | substitution | 90 | Val | His | 0.375 | 0.018 |
| p.Val90Ile | substitution | 90 | Val | Ile | 0.863 | 0.005 |
| p.Val90Leu | substitution | 90 | Val | Leu | 0.990 | 0.016 |
| p.Val90Lys | substitution | 90 | Val | Lys | 0.000 | 0.023 |
| p.Val90Met | substitution | 90 | Val | Met | 0.762 | 0.005 |
| p.Val90Phe | substitution | 90 | Val | Phe | 0.286 | 0.014 |
| p.Val90Pro | substitution | 90 | Val | Pro | 0.000 | 0.031 |
| p.Val90Ser | substitution | 90 | Val | Ser | 0.505 | 0.006 |
| p.Val90Ter | nonsense | 90 | Val | Ter | 0.000 | 0.013 |
| p.Val90Thr | substitution | 90 | Val | Thr | 0.549 | 0.005 |
| p.Val90Trp | substitution | 90 | Val | Trp | 0.199 | 0.006 |
| p.Val90Tyr | substitution | 90 | Val | Tyr | 0.346 | 0.019 |
| p.Val93= | synonymous | 93 | Val | NA | 1.140 | 0.004 |
| p.Val93Ala | substitution | 93 | Val | Ala | 0.891 | 0.006 |
| p.Val93Arg | substitution | 93 | Val | Arg | 0.845 | 0.257 |
| p.Val93Asn | substitution | 93 | Val | Asn | 0.657 | 0.012 |
| p.Val93Asp | substitution | 93 | Val | Asp | 0.507 | 0.076 |
| p.Val93Cys | substitution | 93 | Val | Cys | 0.675 | 0.010 |
| p.Val93Glu | substitution | 93 | Val | Glu | 0.000 | 0.059 |
| p.Val93Gly | substitution | 93 | Val | Gly | 0.603 | 0.011 |
| p.Val93His | substitution | 93 | Val | His | 1.020 | 0.008 |
| p.Val93Ile | substitution | 93 | Val | Ile | 1.148 | 0.003 |
| p.Val93Leu | substitution | 93 | Val | Leu | 0.703 | 0.033 |
| p.Val93Lys | substitution | 93 | Val | Lys | 0.000 | 0.986 |
| p.Val93Met | substitution | 93 | Val | Met | 0.727 | 0.048 |
| p.Val93Phe | substitution | 93 | Val | Phe | 0.410 | 0.010 |
| p.Val93Pro | substitution | 93 | Val | Pro | 0.032 | 0.087 |
| p.Val93Ser | substitution | 93 | Val | Ser | 0.172 | 0.177 |
| p.Val93Ter | nonsense | 93 | Val | Ter | 0.000 | 0.029 |
| p.Val93Thr | substitution | 93 | Val | Thr | 0.407 | 0.022 |
| p.Val93Trp | substitution | 93 | Val | Trp | 0.000 | 0.040 |
| p.Val93Tyr | substitution | 93 | Val | Tyr | 0.020 | 0.017 |
| p.Val94= | synonymous | 94 | Val | NA | 0.944 | 0.024 |
| p.Val94Ala | substitution | 94 | Val | Ala | 0.000 | 0.139 |
| p.Val94Arg | substitution | 94 | Val | Arg | 0.000 | 0.059 |
| p.Val94Asn | substitution | 94 | Val | Asn | 0.122 | 0.154 |
| p.Val94Asp | substitution | 94 | Val | Asp | 0.080 | 1.429 |
| p.Val94Cys | substitution | 94 | Val | Cys | 1.100 | 0.007 |
| p.Val94Gln | substitution | 94 | Val | Gln | 0.345 | 0.040 |

| | | | | | | |
|---|---|---|---|---|---|---|
| p.Val94Glu | substitution | 94 | Val | Glu | 0.088 | 0.021 |
| p.Val94Gly | substitution | 94 | Val | Gly | 0.174 | 0.017 |
| p.Val94His | substitution | 94 | Val | His | 0.471 | 0.347 |
| p.Val94Ile | substitution | 94 | Val | Ile | 0.543 | 0.068 |
| p.Val94Leu | substitution | 94 | Val | Leu | 0.606 | 0.061 |
| p.Val94Lys | substitution | 94 | Val | Lys | 0.040 | 0.120 |
| p.Val94Met | substitution | 94 | Val | Met | 0.000 | 0.267 |
| p.Val94Phe | substitution | 94 | Val | Phe | 0.190 | 0.019 |
| p.Val94Pro | substitution | 94 | Val | Pro | 0.363 | 0.128 |
| p.Val94Ser | substitution | 94 | Val | Ser | 0.000 | 0.058 |
| p.Val94Ter | nonsense | 94 | Val | Ter | 0.505 | 0.054 |
| p.Val94Thr | substitution | 94 | Val | Thr | 0.468 | 0.086 |
| p.Val94Trp | substitution | 94 | Val | Trp | 0.000 | 0.037 |
| p.Val94Tyr | substitution | 94 | Val | Tyr | 0.061 | 0.109 |

**Log Likelihood Ratio of Pathogenicity**

| **LLR** | **95% CI** | **Evidence Strength for Variant Class.** |
|---|---|---|
| -1.096 | [-1.099;-1.09 | benign supporting |
| 1.276 | [0.200;5.105 | pathogenic strong |
| 1.276 | [-1.089;5.06 | pathogenic strong |
| 1.116 | [0.804;1.237 | pathogenic moderate |
| -1.107 | [-1.162;-1.08 | benign supporting |
| -0.397 | [-0.451;-0.34 | benign supporting |
| 1.276 | [1.042;1.871 | pathogenic strong |
| -0.462 | [-0.579;-0.37 | benign supporting |
| 1.276 | [1.195;1.389 | pathogenic strong |
| 1.276 | [1.209;1.360 | pathogenic strong |
| -0.463 | [-0.666;-0.28 | benign supporting |
| 1.276 | [1.167;1.443 | pathogenic strong |
| 1.276 | [1.200;1.374 | pathogenic strong |
| 1.276 | [1.199;1.377 | pathogenic strong |
| 0.293 | [-0.206;0.83 | indeterminate |
| -0.693 | [-0.846;-0.60 | benign supporting |
| 0.304 | [0.016;0.513 | indeterminate |
| -0.754 | [-0.860;-0.66 | benign supporting |
| 1.276 | [1.184;1.405 | pathogenic strong |
| 0.764 | [0.469;1.003 | pathogenic moderate |
| -0.345 | [-0.505;-0.11 | benign supporting |
| -1.116 | [-1.168;-1.08 | benign supporting |
| 1.276 | [1.201;1.377 | pathogenic strong |
| -1.015 | [-1.065;-0.96 | benign supporting |
| 0.378 | [0.264;0.480 | pathogenic supporting |
| -0.107 | [-0.207;0.00 | indeterminate |
| 0.547 | [0.262;0.756 | pathogenic supporting |
| -0.977 | [-1.007;-0.94 | benign supporting |
| 0.844 | [0.759;0.936 | pathogenic moderate |
| 1.134 | [1.004;1.213 | pathogenic moderate |
| 0.656 | [0.243;0.940 | pathogenic moderate |
| 1.131 | [0.965;1.214 | pathogenic moderate |
| -0.483 | [-0.554;-0.41 | benign supporting |
| 1.194 | [1.123;1.275 | pathogenic moderate |
| 0.750 | [0.490;0.971 | pathogenic moderate |
| -0.827 | [-1.055;-0.57 | benign supporting |
| 1.276 | [1.209;1.360 | pathogenic strong |
| -0.863 | [-0.913;-0.81 | benign supporting |
| 1.276 | [1.217;1.351 | pathogenic strong |
| 1.162 | [1.127;1.202 | pathogenic moderate |

| | | |
|---|---|---|
| -1.105 | [-1.108;-1.10 | benign supporting |
| -0.852 | [-0.888;-0.81 | benign supporting |
| -1.054 | [-1.070;-1.02 | benign supporting |
| -1.090 | [-1.107;-1.07 | benign supporting |
| -1.091 | [-1.104;-1.08 | benign supporting |
| -0.965 | [-0.999;-0.93 | benign supporting |
| -1.164 | [-1.169;-1.15 | benign supporting |
| 1.276 | [1.109;1.687 | pathogenic strong |
| -1.056 | [-1.072;-1.02 | benign supporting |
| -1.023 | [-1.045;-1.00 | benign supporting |
| -0.003 | [-0.494;1.14( | indeterminate |
| -1.066 | [-1.094;-1.00 | benign supporting |
| -0.538 | [-0.570;-0.50 | benign supporting |
| 1.127 | [0.963;1.207 | pathogenic moderate |
| -0.794 | [-0.906;-0.67 | benign supporting |
| 0.947 | [-0.005;1.27 | pathogenic moderate |
| -0.831 | [-1.065;-0.57 | benign supporting |
| 1.276 | [0.892;5.022 | pathogenic strong |
| -0.564 | [-0.622;-0.50 | benign supporting |
| -1.030 | [-1.059;-1.00 | benign supporting |
| -0.674 | [-0.707;-0.64 | benign supporting |
| -0.966 | [-0.986;-0.94 | benign supporting |
| -1.066 | [-1.084;-1.02 | benign supporting |
| -1.063 | [-1.101;-0.95 | benign supporting |
| -0.370 | [-0.396;-0.34 | benign supporting |
| 0.892 | [-1.165;4.95( | pathogenic moderate |
| -0.888 | [-1.000;-0.74 | benign supporting |
| 0.792 | [0.704;0.885 | pathogenic moderate |
| 0.428 | [0.237;0.585 | pathogenic supporting |
| -0.797 | [-0.896;-0.68 | benign supporting |
| -1.165 | [-1.170;-1.14 | benign supporting |
| -1.108 | [-1.115;-1.09 | benign supporting |
| -1.105 | [-1.115;-1.09 | benign supporting |
| -0.463 | [-0.605;-0.34 | benign supporting |
| -0.852 | [-0.917;-0.77 | benign supporting |
| 1.036 | [0.849;1.137 | pathogenic moderate |
| -0.371 | [-0.494;-0.21 | benign supporting |
| -0.365 | [-0.621;0.01! | benign supporting |
| -1.084 | [-1.121;-1.08 | benign supporting |
| -0.864 | [-0.891;-0.83 | benign supporting |
| -0.639 | [-0.751;-0.56 | benign supporting |
| -1.084 | [-1.115;-1.08 | benign supporting |
| -0.957 | [-1.165;-0.83 | benign supporting |

| | | |
|---|---|---|
| -0.253 | [-0.638;0.867 | indeterminate |
| -0.852 | [-3.763;0.000 | benign supporting |
| -0.050 | [-0.444;1.028 | indeterminate |
| -1.170 | [-1.169;-1.10 | benign supporting |
| -0.720 | [-1.113;-0.04 | benign supporting |
| -1.114 | [-1.165;-0.75 | benign supporting |
| 1.276 | [1.213;1.359 | pathogenic strong |
| 1.235 | [1.165;1.316 | pathogenic moderate |
| 1.276 | [1.199;1.377 | pathogenic strong |
| -0.863 | [-0.898;-0.83 | benign supporting |
| 1.224 | [1.040;1.456 | pathogenic moderate |
| 1.213 | [0.992;1.444 | pathogenic moderate |
| -0.697 | [-0.780;-0.64 | benign supporting |
| 1.276 | [1.228;1.333 | pathogenic strong |
| 1.276 | [0.966;2.770 | pathogenic strong |
| 1.276 | [1.191;1.392 | pathogenic strong |
| 0.339 | [-0.019;0.618 | pathogenic supporting |
| 1.209 | [1.135;1.294 | pathogenic moderate |
| 1.276 | [1.065;1.790 | pathogenic strong |
| 1.170 | [0.686;1.492 | pathogenic moderate |
| -0.557 | [-1.056;-0.11 | benign supporting |
| 0.225 | [-0.659;1.404 | indeterminate |
| 0.187 | [-0.335;1.016 | indeterminate |
| 1.276 | [0.622;5.087 | pathogenic strong |
| 1.276 | [1.224;1.340 | pathogenic strong |
| 1.276 | [1.246;1.309 | pathogenic strong |
| -0.867 | [-1.020;-0.67 | benign supporting |
| 1.276 | [1.129;1.565 | pathogenic strong |
| 0.860 | [0.711;1.019 | pathogenic moderate |
| 0.889 | [0.650;1.102 | pathogenic moderate |
| 1.276 | [-0.239;5.100 | pathogenic strong |
| 1.276 | [1.148;1.496 | pathogenic strong |
| -0.158 | [-0.200;-0.11 | indeterminate |
| 1.276 | [-0.194;5.108 | pathogenic strong |
| 1.276 | [1.190;1.392 | pathogenic strong |
| 1.276 | [1.223;1.340 | pathogenic strong |
| 0.871 | [-0.019;1.225 | pathogenic moderate |
| 1.276 | [1.217;1.349 | pathogenic strong |
| 1.102 | [1.020;1.135 | pathogenic moderate |
| 1.276 | [1.188;1.401 | pathogenic strong |
| -0.562 | [-0.694;-0.42 | benign supporting |
| -0.643 | [-0.660;-0.62 | benign supporting |
| 1.276 | [1.146;1.509 | pathogenic strong |

| | | |
|---|---|---|
| 1.112 | [0.425;1.359 | pathogenic moderate |
| 1.276 | [1.108;1.680 | pathogenic strong |
| -0.637 | [-0.728;-0.57 | benign supporting |
| 1.276 | [1.243;1.314 | pathogenic strong |
| -0.183 | [-0.206;-0.16 | indeterminate |
| -0.187 | [-0.243;-0.12 | indeterminate |
| -0.826 | [-0.854;-0.78 | benign supporting |
| 1.276 | [1.174;1.424 | pathogenic strong |
| 0.657 | [0.613;0.700 | pathogenic moderate |
| -0.592 | [-0.615;-0.56 | benign supporting |
| 1.125 | [1.110;1.141 | pathogenic moderate |
| 1.273 | [1.226;1.327 | pathogenic strong |
| 1.276 | [1.242;1.316 | pathogenic strong |
| -1.164 | [-1.164;-1.16 | benign supporting |
| -0.025 | [-0.085;0.10 | indeterminate |
| -0.617 | [-0.647;-0.58 | benign supporting |
| -0.355 | [-0.542;-0.09 | benign supporting |
| 1.276 | [1.225;1.336 | pathogenic strong |
| -0.486 | [-0.494;-0.47 | benign supporting |
| 1.276 | [1.202;1.373 | pathogenic strong |
| 1.276 | [1.229;1.333 | pathogenic strong |
| 1.276 | [1.255;1.298 | pathogenic strong |
| -0.968 | [-1.028;-0.90 | benign supporting |
| 1.227 | [0.321;5.064 | pathogenic moderate |
| 1.276 | [1.196;1.387 | pathogenic strong |
| 1.276 | [0.909;5.021 | pathogenic strong |
| 1.276 | [1.166;1.441 | pathogenic strong |
| 0.371 | [-0.269;1.04 | pathogenic supporting |
| -0.618 | [-0.998;-0.35 | benign supporting |
| 1.276 | [1.164;1.455 | pathogenic strong |
| 1.276 | [1.246;1.309 | pathogenic strong |
| 1.276 | [-0.156;5.11 | pathogenic strong |
| 1.276 | [0.000;5.107 | pathogenic strong |
| 1.276 | [1.218;1.347 | pathogenic strong |
| -0.633 | [-0.946;-0.42 | benign supporting |
| -0.321 | [-0.368;-0.26 | benign supporting |
| 0.875 | [0.444;1.144 | pathogenic moderate |
| -1.091 | [-1.168;-0.65 | benign supporting |
| -1.090 | [-1.092;-1.08 | benign supporting |
| -0.447 | [-0.497;-0.40 | benign supporting |
| 1.174 | [1.137;1.214 | pathogenic moderate |
| -0.997 | [-1.037;-0.94 | benign supporting |
| -1.091 | [-1.114;-1.07 | benign supporting |

| | | |
|---|---|---|
| -1.103 | [-1.110;-1.09 | benign supporting |
| 0.000 | [0.000;0.000 | indeterminate |
| -1.065 | [-1.079;-1.02 | benign supporting |
| -1.104 | [-1.115;-1.09 | benign supporting |
| -0.413 | [-0.426;-0.40 | benign supporting |
| -0.591 | [-0.666;-0.51 | benign supporting |
| -0.920 | [-3.548;-0.83 | benign supporting |
| -1.018 | [-1.037;-0.99 | benign supporting |
| -0.844 | [-0.891;-0.78 | benign supporting |
| -0.337 | [-0.393;-0.26 | benign supporting |
| -0.573 | [-0.616;-0.52 | benign supporting |
| 1.236 | [1.188;1.289 | pathogenic moderate |
| -0.637 | [-0.870;-0.49 | benign supporting |
| -0.839 | [-0.839;-0.83 | benign supporting |
| -1.105 | [-1.115;-1.02 | benign supporting |
| -1.113 | [-1.115;-1.08 | benign supporting |
| -0.986 | [-1.028;-0.96 | benign supporting |
| 1.276 | [1.213;1.357 | pathogenic strong |
| 1.276 | [1.059;1.835 | pathogenic strong |
| 1.082 | [0.999;1.128 | pathogenic moderate |
| 1.276 | [1.172;1.428 | pathogenic strong |
| 1.234 | [1.043;1.497 | pathogenic moderate |
| 1.276 | [1.248;1.307 | pathogenic strong |
| 1.276 | [1.202;1.373 | pathogenic strong |
| 1.276 | [1.064;1.816 | pathogenic strong |
| 0.867 | [0.503;1.123 | pathogenic moderate |
| 1.002 | [0.848;1.108 | pathogenic moderate |
| 1.276 | [1.167;1.449 | pathogenic strong |
| 1.276 | [1.226;1.336 | pathogenic strong |
| 0.901 | [0.012;1.237 | pathogenic moderate |
| -1.088 | [-1.142;-1.08 | benign supporting |
| 1.276 | [1.020;1.887 | pathogenic strong |
| -1.168 | [-1.169;-1.11 | benign supporting |
| 1.276 | [1.196;1.383 | pathogenic strong |
| 1.276 | [1.234;1.325 | pathogenic strong |
| -0.394 | [-0.454;-0.33 | benign supporting |
| -1.095 | [-1.115;-0.93 | benign supporting |
| 1.159 | [0.743;1.394 | pathogenic moderate |
| 1.276 | [1.241;1.317 | pathogenic strong |
| 1.276 | [0.935;4.832 | pathogenic strong |
| 1.276 | [-0.221;5.10 | pathogenic strong |
| -0.579 | [-0.597;-0.56 | benign supporting |
| -0.165 | [-0.605;1.03 | indeterminate |

| | | |
|---|---|---|
| 1.276 | [-0.031;5.09ε | pathogenic strong |
| 1.192 | [1.128;1.262 | pathogenic moderate |
| 1.276 | [0.948;4.851 | pathogenic strong |
| 1.276 | [1.064;1.747 | pathogenic strong |
| 1.276 | [1.165;1.445 | pathogenic strong |
| -0.971 | [-3.655;0.00( | benign supporting |
| -1.169 | [-1.169;-0.93 | benign supporting |
| 1.276 | [0.902;5.036 | pathogenic strong |
| 1.276 | [1.175;1.420 | pathogenic strong |
| -1.127 | [-1.169;-1.04 | benign supporting |
| -0.956 | [-1.080;-0.74 | benign supporting |
| 1.121 | [1.086;1.145 | pathogenic moderate |
| -0.629 | [-0.729;-0.55 | benign supporting |
| 1.276 | [0.622;5.097 | pathogenic strong |
| 1.276 | [0.864;5.019 | pathogenic strong |
| 0.782 | [0.179;1.129 | pathogenic moderate |
| 0.969 | [0.230;1.238 | pathogenic moderate |
| 1.104 | [-0.392;5.037 | pathogenic moderate |
| 1.206 | [1.111;1.324 | pathogenic moderate |
| -0.514 | [-0.540;-0.48 | benign supporting |
| 1.276 | [0.734;5.084 | pathogenic strong |
| 1.276 | [-0.079;5.10: | pathogenic strong |
| 1.193 | [0.422;4.045 | pathogenic moderate |
| -0.352 | [-1.164;4.81c | benign supporting |
| -0.987 | [-1.167;-0.88 | benign supporting |
| 0.217 | [0.081;0.327 | indeterminate |
| -1.101 | [-1.115;-1.08 | benign supporting |
| 1.152 | [1.118;1.188 | pathogenic moderate |
| -1.094 | [-1.115;-1.00 | benign supporting |
| -1.086 | [-1.089;-1.08 | benign supporting |
| -1.166 | [-1.169;-1.16 | benign supporting |
| -0.115 | [-0.271;0.16: | indeterminate |
| -0.983 | [-1.000;-0.96 | benign supporting |
| -0.409 | [-0.424;-0.39 | benign supporting |
| 1.276 | [1.264;1.290 | pathogenic strong |
| -3.211 | [-3.338;-3.02 | benign strong |
| -1.151 | [-1.169;-1.12 | benign supporting |
| -1.148 | [-1.168;-1.13 | benign supporting |
| 1.276 | [1.136;1.525 | pathogenic strong |
| -0.380 | [-0.985;0.80( | benign supporting |
| -0.209 | [-0.337;-0.03 | indeterminate |
| -0.355 | [-0.393;-0.31 | benign supporting |
| -0.359 | [-0.380;-0.33 | benign supporting |

| | | |
|---|---|---|
| -0.086 | [-0.123;-0.04 | indeterminate |
| -1.105 | [-1.114;-1.09 | benign supporting |
| -1.087 | [-1.101;-1.08 | benign supporting |
| 1.276 | [1.130;1.558 | pathogenic strong |
| 1.276 | [1.022;1.942 | pathogenic strong |
| -0.957 | [-1.113;-0.49 | benign supporting |
| -0.891 | [-0.899;-0.88 | benign supporting |
| 0.698 | [0.579;0.790 | pathogenic moderate |
| 0.573 | [0.432;0.692 | pathogenic supporting |
| -0.899 | [-1.111;-0.42 | benign supporting |
| 1.276 | [0.695;5.085 | pathogenic strong |
| 0.417 | [-0.413;1.21 | pathogenic supporting |
| 1.276 | [1.242;1.314 | pathogenic strong |
| 1.276 | [-0.228;5.10 | pathogenic strong |
| 1.276 | [1.171;1.437 | pathogenic strong |
| 1.276 | [-0.268;5.10 | pathogenic strong |
| -0.105 | [-0.394;0.70 | indeterminate |
| 1.276 | [1.229;1.331 | pathogenic strong |
| -1.079 | [-1.114;-0.96 | benign supporting |
| -1.069 | [-1.077;-1.05 | benign supporting |
| 0.929 | [0.390;1.191 | pathogenic moderate |
| -0.371 | [-1.090;1.14 | benign supporting |
| -0.625 | [-0.659;-0.59 | benign supporting |
| -1.096 | [-1.167;-0.74 | benign supporting |
| -1.000 | [-1.167;-0.89 | benign supporting |
| -0.904 | [-3.594;-0.83 | benign supporting |
| 1.276 | [1.210;1.361 | pathogenic strong |
| -0.837 | [-0.848;-0.82 | benign supporting |
| 1.119 | [0.868;1.221 | pathogenic moderate |
| 1.253 | [1.013;1.676 | pathogenic moderate |
| -1.039 | [-1.049;-1.03 | benign supporting |
| -1.050 | [-1.168;-0.95 | benign supporting |
| -1.145 | [-1.155;-1.13 | benign supporting |
| 1.141 | [-0.375;5.05 | pathogenic moderate |
| -0.837 | [-0.837;-0.83 | benign supporting |
| 1.276 | [1.137;1.529 | pathogenic strong |
| -0.854 | [-1.105;-0.47 | benign supporting |
| -0.682 | [-0.860;-0.57 | benign supporting |
| -1.101 | [-1.115;-1.07 | benign supporting |
| 1.276 | [1.216;1.352 | pathogenic strong |
| -0.163 | [-0.351;0.24 | indeterminate |
| 0.697 | [0.044;1.077 | pathogenic moderate |
| -0.525 | [-0.560;-0.49 | benign supporting |

| | | |
|---|---|---|
| 1.276 | [1.259;1.295 | pathogenic strong |
| 1.276 | [1.253;1.301 | pathogenic strong |
| 1.276 | [1.267;1.286 | pathogenic strong |
| 1.276 | [1.270;1.283 | pathogenic strong |
| 1.276 | [1.251;1.303 | pathogenic strong |
| 1.276 | [1.274;1.278 | pathogenic strong |
| -0.574 | [-0.574;-0.57 | benign supporting |
| 1.276 | [1.263;1.290 | pathogenic strong |
| 1.276 | [1.267;1.286 | pathogenic strong |
| 1.276 | [1.252;1.302 | pathogenic strong |
| 1.276 | [1.258;1.296 | pathogenic strong |
| 1.276 | [1.220;1.346 | pathogenic strong |
| 1.276 | [1.267;1.286 | pathogenic strong |
| 1.276 | [1.238;1.319 | pathogenic strong |
| -0.960 | [-1.007;-0.91 | benign supporting |
| 1.276 | [1.251;1.304 | pathogenic strong |
| 1.276 | [0.932;4.959 | pathogenic strong |
| 1.276 | [1.268;1.285 | pathogenic strong |
| 1.276 | [1.260;1.293 | pathogenic strong |
| 1.276 | [1.235;1.324 | pathogenic strong |
| -0.912 | [-1.015;-0.76 | benign supporting |
| 1.131 | [1.049;1.181 | pathogenic moderate |
| 1.276 | [0.525;5.090 | pathogenic strong |
| -0.663 | [-0.698;-0.63 | benign supporting |
| -0.277 | [-0.301;-0.25 | indeterminate |
| 1.216 | [-0.145;5.08 | pathogenic moderate |
| -1.023 | [-1.030;-1.01 | benign supporting |
| -1.167 | [-1.170;-1.12 | benign supporting |
| 1.276 | [1.154;1.476 | pathogenic strong |
| 1.276 | [1.174;1.426 | pathogenic strong |
| -0.540 | [-1.110;0.99 | benign supporting |
| 0.759 | [0.708;0.811 | pathogenic moderate |
| 0.639 | [-0.628;4.26 | pathogenic moderate |
| 1.263 | [1.227;1.301 | pathogenic moderate |
| 0.988 | [-0.218;1.51 | pathogenic moderate |
| 1.276 | [1.184;1.404 | pathogenic strong |
| 1.276 | [-0.202;5.10 | pathogenic strong |
| -1.102 | [-1.166;-0.61 | benign supporting |
| -1.120 | [-1.153;-1.09 | benign supporting |
| 1.276 | [1.138;1.530 | pathogenic strong |
| -0.928 | [-0.939;-0.91 | benign supporting |
| -0.308 | [-0.584;0.45 | indeterminate |
| -1.086 | [-1.092;-1.08 | benign supporting |

| | | |
|---|---|---|
| -1.084 | [-1.086;-1.08 | benign supporting |
| -1.080 | [-1.088;-1.07 | benign supporting |
| -3.739 | [-3.790;-3.67 | benign strong |
| -0.703 | [-1.165;1.354 | benign supporting |
| 1.243 | [0.986;1.614] | pathogenic moderate |
| 1.276 | [0.837;5.060] | pathogenic strong |
| -0.977 | [-1.168;-0.83 | benign supporting |
| 1.276 | [0.954;2.454] | pathogenic strong |
| -0.392 | [-0.465;-0.31 | benign supporting |
| -0.260 | [-0.408;-0.01 | indeterminate |
| -0.493 | [-1.050;0.215 | benign supporting |
| -0.850 | [-1.113;-0.32 | benign supporting |
| 1.276 | [0.956;4.756] | pathogenic strong |
| -1.169 | [-1.169;-1.13 | benign supporting |
| -0.886 | [-2.612;4.622 | benign supporting |
| 1.276 | [0.245;5.101] | pathogenic strong |
| 1.276 | [0.721;5.096] | pathogenic strong |
| -0.397 | [-1.094;1.131 | benign supporting |
| -1.099 | [-1.107;-1.09 | benign supporting |
| 1.112 | [1.038;1.147] | pathogenic moderate |
| -0.845 | [-3.693;-0.83 | benign supporting |
| 1.276 | [1.196;1.380] | pathogenic strong |
| 1.143 | [1.115;1.173] | pathogenic moderate |
| 1.276 | [1.223;1.340] | pathogenic strong |
| 0.866 | [0.193;1.187] | pathogenic moderate |
| 0.957 | [0.846;1.054] | pathogenic moderate |
| -0.609 | [-1.068;-0.25 | benign supporting |
| 0.603 | [0.062;0.913] | pathogenic supporting |
| -1.069 | [-1.084;-1.03 | benign supporting |
| 1.108 | [0.352;1.367] | pathogenic moderate |
| -0.316 | [-0.402;-0.17 | indeterminate |
| 1.276 | [1.225;1.336] | pathogenic strong |
| 1.276 | [0.799;5.068] | pathogenic strong |
| -1.011 | [-1.068;-0.96 | benign supporting |
| 1.276 | [1.179;1.416] | pathogenic strong |
| 1.276 | [-0.009;5.110 | pathogenic strong |
| 1.276 | [-0.049;5.101 | pathogenic strong |
| -0.366 | [-1.164;4.934 | benign supporting |
| 1.276 | [-0.321;5.094 | pathogenic strong |
| -1.085 | [-1.091;-1.08 | benign supporting |
| 0.934 | [-0.439;4.745 | pathogenic moderate |
| -0.946 | [-1.105;-0.62 | benign supporting |
| -1.168 | [-1.169;-1.10 | benign supporting |

| | | |
|---|---|---|
| -0.899 | [-0.925;-0.86 | benign supporting |
| 1.276 | [0.638;5.090 | pathogenic strong |
| 1.104 | [-0.170;1.833 | pathogenic moderate |
| -1.088 | [-1.167;-1.08 | benign supporting |
| 1.276 | [1.032;1.857 | pathogenic strong |
| 0.651 | [-0.244;1.190 | pathogenic moderate |
| 1.260 | [-0.343;5.096 | pathogenic moderate |
| 1.276 | [-0.354;5.098 | pathogenic strong |
| -0.056 | [-0.295;0.581 | indeterminate |
| 1.243 | [1.004;1.635 | pathogenic moderate |
| -0.899 | [-3.807;0.000 | benign supporting |
| -1.072 | [-1.114;-0.76 | benign supporting |
| -1.165 | [-1.169;-1.03 | benign supporting |
| 1.276 | [1.128;1.579 | pathogenic strong |
| -0.860 | [-0.877;-0.84 | benign supporting |
| 1.258 | [1.177;1.363 | pathogenic moderate |
| -0.839 | [-3.759;0.000 | benign supporting |
| 1.227 | [-0.417;5.095 | pathogenic moderate |
| 1.276 | [-0.080;5.110 | pathogenic strong |
| -1.024 | [-1.038;-1.00 | benign supporting |
| -1.102 | [-1.164;-0.96 | benign supporting |
| -1.070 | [-3.591;0.000 | benign supporting |
| 1.276 | [0.686;5.095 | pathogenic strong |
| 1.117 | [0.967;1.184 | pathogenic moderate |
| -1.084 | [-1.086;-1.08 | benign supporting |
| -1.164 | [-1.169;-1.13 | benign supporting |
| 0.851 | [0.351;1.144 | pathogenic moderate |
| -1.097 | [-1.168;-0.99 | benign supporting |
| 1.276 | [1.033;1.881 | pathogenic strong |
| 0.336 | [-0.202;0.876 | pathogenic supporting |
| -1.164 | [-1.169;-1.08 | benign supporting |
| -0.071 | [-0.303;0.519 | indeterminate |
| 1.276 | [1.082;1.739 | pathogenic strong |
| -1.080 | [-1.100;-1.05 | benign supporting |
| 1.276 | [-0.190;5.098 | pathogenic strong |
| 1.276 | [1.204;1.365 | pathogenic strong |
| -1.114 | [-1.169;-1.02 | benign supporting |
| 0.601 | [-0.422;1.342 | pathogenic supporting |
| -0.381 | [-0.613;-0.04 | benign supporting |
| 1.276 | [1.219;1.346 | pathogenic strong |
| 1.276 | [1.138;1.521 | pathogenic strong |
| 1.276 | [0.650;5.085 | pathogenic strong |
| -1.068 | [-1.096;-1.00 | benign supporting |

| | | |
|---|---|---|
| -1.085 | [-1.168;-0.76 | benign supporting |
| 1.276 | [1.203;1.372] | pathogenic strong |
| -0.994 | [-1.014;-0.97 | benign supporting |
| -1.100 | [-1.115;-1.07 | benign supporting |
| -0.447 | [-0.517;-0.39 | benign supporting |
| -0.408 | [-0.428;-0.39 | benign supporting |
| -1.041 | [-1.065;-1.01 | benign supporting |
| -1.148 | [-1.166;-1.12 | benign supporting |
| -0.454 | [-0.941;0.038 | benign supporting |
| -0.863 | [-0.895;-0.83 | benign supporting |
| -1.089 | [-1.092;-1.08 | benign supporting |
| -0.565 | [-0.581;-0.54 | benign supporting |
| -0.494 | [-0.526;-0.46 | benign supporting |
| -0.674 | [-0.724;-0.63 | benign supporting |
| -1.085 | [-1.092;-1.08 | benign supporting |
| -1.091 | [-1.098;-1.08 | benign supporting |
| -0.403 | [-0.420;-0.38 | benign supporting |
| 1.191 | [0.808;1.480] | pathogenic moderate |
| -0.799 | [-0.881;-0.70 | benign supporting |
| 0.684 | [-0.108;1.145 | pathogenic moderate |
| -1.090 | [-1.115;-1.03 | benign supporting |
| -0.911 | [-0.933;-0.88 | benign supporting |
| -1.098 | [-1.106;-1.09 | benign supporting |
| -1.106 | [-1.115;-1.07 | benign supporting |
| -1.114 | [-1.116;-1.11 | benign supporting |
| -1.100 | [-1.108;-1.09 | benign supporting |
| -1.088 | [-1.098;-1.08 | benign supporting |
| -0.669 | [-0.687;-0.65 | benign supporting |
| 0.186 | [-0.101;0.566 | indeterminate |
| -0.838 | [-0.861;-0.81 | benign supporting |
| -1.110 | [-1.116;-1.10 | benign supporting |
| -1.064 | [-1.075;-1.03 | benign supporting |
| -0.775 | [-0.865;-0.68 | benign supporting |
| -0.735 | [-1.110;-0.21 | benign supporting |
| -0.844 | [-0.898;-0.77 | benign supporting |
| -1.087 | [-1.089;-1.08 | benign supporting |
| 1.130 | [1.107;1.153] | pathogenic moderate |
| -1.115 | [-1.116;-1.09 | benign supporting |
| -0.270 | [-0.319;-0.21 | indeterminate |
| -1.082 | [-1.091;-1.07 | benign supporting |
| 1.276 | [0.995;2.450] | pathogenic strong |
| -1.104 | [-1.115;-1.08 | benign supporting |
| -0.853 | [-0.864;-0.84 | benign supporting |

| | | |
|---|---|---|
| -1.096 | [-1.110;-1.08 | benign supporting |
| -1.084 | [-1.103;-1.08 | benign supporting |
| -1.115 | [-1.168;-1.08 | benign supporting |
| -0.878 | [-0.975;-0.75 | benign supporting |
| -0.459 | [-0.486;-0.43 | benign supporting |
| -1.029 | [-1.040;-1.01 | benign supporting |
| -0.518 | [-0.549;-0.48 | benign supporting |
| -0.908 | [-0.973;-0.83 | benign supporting |
| -0.855 | [-0.861;-0.85 | benign supporting |
| -0.609 | [-0.617;-0.59 | benign supporting |
| -1.105 | [-1.115;-1.09 | benign supporting |
| -0.583 | [-0.600;-0.56 | benign supporting |
| -0.917 | [-0.930;-0.90 | benign supporting |
| -0.443 | [-0.462;-0.42 | benign supporting |
| -0.435 | [-0.460;-0.41 | benign supporting |
| -1.137 | [-1.165;-1.10 | benign supporting |
| -1.049 | [-1.081;-0.99 | benign supporting |
| -1.106 | [-1.115;-1.09 | benign supporting |
| 1.202 | [1.158;1.248] | pathogenic moderate |
| -0.604 | [-0.802;-0.45 | benign supporting |
| -0.599 | [-0.625;-0.56 | benign supporting |
| -1.043 | [-1.063;-1.02 | benign supporting |
| -1.086 | [-1.100;-1.07 | benign supporting |
| -1.166 | [-3.422;1.872 | benign supporting |
| -1.130 | [-1.169;-0.94 | benign supporting |
| 0.753 | [-0.135;1.205 | pathogenic moderate |
| 0.000 | [0.000;0.000] | indeterminate |
| -0.576 | [-0.859;-0.37 | benign supporting |
| 1.276 | [-0.736;5.098 | pathogenic strong |
| -0.751 | [-0.971;-0.57 | benign supporting |
| 1.132 | [-1.006;5.070 | pathogenic moderate |
| -1.041 | [-1.071;-1.01 | benign supporting |
| -0.639 | [-1.112;0.242 | benign supporting |
| -0.841 | [-0.874;-0.83 | benign supporting |
| 1.276 | [1.128;1.583] | pathogenic strong |
| 1.276 | [0.000;5.102] | pathogenic strong |
| 1.141 | [-0.928;5.062 | pathogenic moderate |
| 1.276 | [0.959;3.298] | pathogenic strong |
| 1.276 | [-0.463;5.098 | pathogenic strong |
| -1.150 | [-1.169;-1.09 | benign supporting |
| 1.276 | [1.164;1.450] | pathogenic strong |
| -0.957 | [-1.110;-0.56 | benign supporting |
| -0.902 | [-0.930;-0.86 | benign supporting |

| | | |
|---|---|---|
| -0.063 | [-0.170;0.181 | indeterminate |
| -0.408 | [-0.448;-0.37 | benign supporting |
| -0.530 | [-0.556;-0.50 | benign supporting |
| -0.460 | [-0.519;-0.41 | benign supporting |
| -0.232 | [-0.277;-0.17 | indeterminate |
| -0.340 | [-0.358;-0.32 | benign supporting |
| -0.418 | [-0.470;-0.37 | benign supporting |
| -0.600 | [-0.724;-0.48 | benign supporting |
| 0.011 | [-0.357;0.903 | indeterminate |
| -0.344 | [-0.550;-0.04 | benign supporting |
| -0.595 | [-0.639;-0.54 | benign supporting |
| -1.115 | [-1.168;-1.08 | benign supporting |
| -1.064 | [-1.099;-0.96 | benign supporting |
| -0.233 | [-0.533;0.672 | indeterminate |
| -0.761 | [-0.997;-0.57 | benign supporting |
| 1.200 | [0.840;1.510 | pathogenic moderate |
| -1.048 | [-1.106;-0.91 | benign supporting |
| -0.142 | [-0.246;-0.04 | indeterminate |
| -0.599 | [-1.167;4.633 | benign supporting |
| -0.462 | [-0.517;-0.41 | benign supporting |
| -1.089 | [-1.120;-1.08 | benign supporting |
| -0.509 | [-0.524;-0.49 | benign supporting |
| 1.112 | [1.078;1.131 | pathogenic moderate |
| 0.947 | [0.894;0.999 | pathogenic moderate |
| -0.366 | [-0.378;-0.35 | benign supporting |
| 0.611 | [0.446;0.745 | pathogenic supporting |
| 0.321 | [0.163;0.443 | pathogenic supporting |
| -0.281 | [-0.330;-0.22 | indeterminate |
| 1.061 | [1.000;1.111 | pathogenic moderate |
| 1.276 | [1.260;1.293 | pathogenic strong |
| 1.133 | [1.122;1.145 | pathogenic moderate |
| 1.194 | [1.181;1.207 | pathogenic moderate |
| 1.230 | [1.207;1.255 | pathogenic moderate |
| 1.009 | [0.798;1.130 | pathogenic moderate |
| -0.488 | [-0.553;-0.42 | benign supporting |
| 1.138 | [0.932;1.239 | pathogenic moderate |
| 0.851 | [0.819;0.884 | pathogenic moderate |
| 0.928 | [0.890;0.968 | pathogenic moderate |
| 0.994 | [0.943;1.029 | pathogenic moderate |
| 1.276 | [1.256;1.298 | pathogenic strong |
| -1.106 | [-1.115;-1.07 | benign supporting |
| 1.276 | [1.195;1.388 | pathogenic strong |
| 1.276 | [0.061;5.107 | pathogenic strong |

| | | |
|---|---|---|
| 1.050 | [0.899;1.130 | pathogenic moderate |
| 1.250 | [1.197;1.310 | pathogenic moderate |
| 1.271 | [1.231;1.315 | pathogenic moderate |
| 1.276 | [-0.453;5.09 | pathogenic strong |
| 1.276 | [1.179;1.418 | pathogenic strong |
| -0.203 | [-0.249;-0.15 | indeterminate |
| 1.052 | [-0.328;4.88 | pathogenic moderate |
| 1.276 | [1.194;1.396 | pathogenic strong |
| 1.234 | [1.187;1.287 | pathogenic moderate |
| 1.276 | [0.489;5.097 | pathogenic strong |
| -0.335 | [-0.798;0.73 | benign supporting |
| 1.276 | [1.199;1.381 | pathogenic strong |
| 1.276 | [1.237;1.322 | pathogenic strong |
| 1.276 | [0.854;5.064 | pathogenic strong |
| 1.276 | [1.193;1.392 | pathogenic strong |
| 1.210 | [1.176;1.244 | pathogenic moderate |
| -0.418 | [-1.102;1.13 | benign supporting |
| 1.276 | [0.553;5.100 | pathogenic strong |
| -0.966 | [-1.011;-0.92 | benign supporting |
| 1.276 | [-1.086;5.07 | pathogenic strong |
| -0.213 | [-0.261;-0.15 | indeterminate |
| 1.276 | [0.834;5.045 | pathogenic strong |
| 1.276 | [1.230;1.331 | pathogenic strong |
| 1.012 | [0.449;1.232 | pathogenic moderate |
| 0.597 | [0.272;0.823 | pathogenic supporting |
| 1.276 | [1.156;1.471 | pathogenic strong |
| 1.276 | [1.200;1.377 | pathogenic strong |
| 1.276 | [1.212;1.360 | pathogenic strong |
| -0.055 | [-0.118;0.04 | indeterminate |
| 1.276 | [1.188;1.404 | pathogenic strong |
| 1.175 | [0.046;4.871 | pathogenic moderate |
| 1.154 | [0.965;1.269 | pathogenic moderate |
| 1.231 | [-0.080;5.09 | pathogenic moderate |
| 1.276 | [-0.126;5.10 | pathogenic strong |
| 1.150 | [1.111;1.192 | pathogenic moderate |
| 1.276 | [1.236;1.321 | pathogenic strong |
| -0.113 | [-1.087;1.90 | indeterminate |
| -1.099 | [-1.114;-1.08 | benign supporting |
| -1.092 | [-1.101;-1.08 | benign supporting |
| -1.081 | [-1.106;-1.05 | benign supporting |
| -0.407 | [-0.475;-0.34 | benign supporting |
| -0.666 | [-0.687;-0.64 | benign supporting |
| -1.072 | [-1.096;-1.02 | benign supporting |

| | | |
|---|---|---|
| -0.994 | [-1.016;-0.96 | benign supporting |
| -0.493 | [-0.517;-0.46 | benign supporting |
| -1.085 | [-1.089;-1.08 | benign supporting |
| -1.137 | [-1.169;-1.11 | benign supporting |
| -0.761 | [-0.854;-0.67 | benign supporting |
| -1.107 | [-1.115;-0.99 | benign supporting |
| -1.125 | [-1.149;-1.10 | benign supporting |
| 0.644 | [0.573;0.711 | pathogenic moderate |
| -1.105 | [-1.115;-1.08 | benign supporting |
| -1.085 | [-1.098;-1.08 | benign supporting |
| 1.239 | [1.143;1.361 | pathogenic moderate |
| -1.115 | [-1.115;-1.07 | benign supporting |
| -1.059 | [-1.068;-1.04 | benign supporting |
| -0.049 | [-0.113;0.06 | indeterminate |
| -1.102 | [-1.113;-1.09 | benign supporting |
| -1.120 | [-1.148;-1.10 | benign supporting |
| -1.060 | [-1.082;-1.00 | benign supporting |
| -1.024 | [-1.038;-1.01 | benign supporting |
| -1.107 | [-1.114;-1.10 | benign supporting |
| -0.914 | [-0.928;-0.89 | benign supporting |
| -0.623 | [-0.637;-0.61 | benign supporting |
| -1.134 | [-1.146;-1.12 | benign supporting |
| -0.932 | [-0.945;-0.91 | benign supporting |
| -1.088 | [-1.093;-1.08 | benign supporting |
| -0.715 | [-0.750;-0.68 | benign supporting |
| -0.946 | [-0.964;-0.92 | benign supporting |
| -1.111 | [-1.116;-1.10 | benign supporting |
| -1.074 | [-1.087;-1.05 | benign supporting |
| -0.629 | [-0.642;-0.61 | benign supporting |
| -0.376 | [-0.438;-0.30 | benign supporting |
| -1.037 | [-1.059;-1.01 | benign supporting |
| 1.276 | [1.234;1.325 | pathogenic strong |
| -1.096 | [-1.104;-1.08 | benign supporting |
| -1.059 | [-1.069;-1.04 | benign supporting |
| -0.390 | [-0.405;-0.37 | benign supporting |
| -0.888 | [-0.920;-0.85 | benign supporting |
| -1.085 | [-1.111;-1.08 | benign supporting |
| -0.862 | [-0.887;-0.83 | benign supporting |
| -0.510 | [-0.537;-0.48 | benign supporting |
| 0.495 | [0.402;0.581 | pathogenic supporting |
| 0.777 | [0.733;0.823 | pathogenic moderate |
| -0.683 | [-0.767;-0.62 | benign supporting |
| 1.276 | [1.251;1.303 | pathogenic strong |

| | | |
|---|---|---|
| -0.789 | [-0.993;-0.60 | benign supporting |
| -0.391 | [-0.410;-0.37 | benign supporting |
| -0.604 | [-0.624;-0.58 | benign supporting |
| -0.630 | [-0.650;-0.61 | benign supporting |
| -0.239 | [-0.291;-0.17 | indeterminate |
| -0.378 | [-0.391;-0.36 | benign supporting |
| 0.251 | [0.113;0.360 | indeterminate |
| 1.095 | [1.057;1.113 | pathogenic moderate |
| -0.571 | [-0.607;-0.53 | benign supporting |
| 1.116 | [1.093;1.133 | pathogenic moderate |
| -1.073 | [-1.080;-1.06 | benign supporting |
| -0.364 | [-0.622;0.03 | benign supporting |
| -0.628 | [-0.637;-0.61 | benign supporting |
| -0.618 | [-0.639;-0.59 | benign supporting |
| -1.113 | [-1.166;-0.78 | benign supporting |
| 1.276 | [1.239;1.318 | pathogenic strong |
| 1.208 | [1.184;1.233 | pathogenic moderate |
| 1.276 | [1.154;1.477 | pathogenic strong |
| 1.276 | [1.244;1.312 | pathogenic strong |
| 1.276 | [-0.458;5.09 | pathogenic strong |
| 1.276 | [1.253;1.302 | pathogenic strong |
| 1.188 | [1.158;1.219 | pathogenic moderate |
| 1.276 | [1.211;1.357 | pathogenic strong |
| 1.211 | [1.149;1.278 | pathogenic moderate |
| 1.043 | [0.953;1.111 | pathogenic moderate |
| 1.276 | [1.249;1.307 | pathogenic strong |
| 1.276 | [1.243;1.314 | pathogenic strong |
| 1.276 | [1.216;1.352 | pathogenic strong |
| 1.276 | [1.220;1.343 | pathogenic strong |
| 1.276 | [1.172;1.434 | pathogenic strong |
| 1.276 | [1.193;1.384 | pathogenic strong |
| 1.276 | [0.899;5.019 | pathogenic strong |
| 1.276 | [1.200;1.379 | pathogenic strong |
| 1.276 | [1.245;1.312 | pathogenic strong |
| -1.028 | [-1.065;-0.99 | benign supporting |
| -1.027 | [-1.075;-0.95 | benign supporting |
| -1.010 | [-1.055;-0.95 | benign supporting |
| -0.346 | [-0.464;-0.16 | benign supporting |
| -1.067 | [-1.081;-1.03 | benign supporting |
| -1.085 | [-1.094;-1.08 | benign supporting |
| -0.066 | [-0.103;-0.01 | indeterminate |
| -0.848 | [-0.967;-0.68 | benign supporting |
| 0.105 | [-0.080;0.41 | indeterminate |

| | | |
|---|---|---|
| -0.743 | [-0.805;-0.69 | benign supporting |
| -1.097 | [-1.106;-1.09 | benign supporting |
| -1.090 | [-1.094;-1.08 | benign supporting |
| 1.170 | [1.139;1.203 | pathogenic moderate |
| 1.021 | [-0.034;1.36 | pathogenic moderate |
| -1.016 | [-1.111;-0.69 | benign supporting |
| 1.152 | [1.133;1.173 | pathogenic moderate |
| -0.611 | [-0.804;-0.46 | benign supporting |
| -1.052 | [-1.091;-0.96 | benign supporting |
| -0.192 | [-0.316;-0.04 | indeterminate |
| -1.094 | [-1.114;-1.08 | benign supporting |
| -1.085 | [-1.128;-1.08 | benign supporting |
| 1.142 | [1.094;1.184 | pathogenic moderate |
| 1.276 | [1.231;1.328 | pathogenic strong |
| 1.276 | [-0.305;5.10 | pathogenic strong |
| 0.686 | [-0.205;1.18 | pathogenic moderate |
| 1.045 | [1.002;1.090 | pathogenic moderate |
| 1.128 | [1.063;1.170 | pathogenic moderate |
| 1.115 | [0.840;1.224 | pathogenic moderate |
| -0.827 | [-1.053;-0.57 | benign supporting |
| -1.030 | [-1.168;-0.83 | benign supporting |
| -1.005 | [-1.096;-0.82 | benign supporting |
| 1.268 | [1.188;1.374 | pathogenic moderate |
| 1.276 | [1.177;1.423 | pathogenic strong |
| 0.876 | [0.328;1.165 | pathogenic moderate |
| 1.122 | [1.021;1.175 | pathogenic moderate |
| 1.276 | [0.847;5.046 | pathogenic strong |
| 1.276 | [1.238;1.320 | pathogenic strong |
| 1.276 | [1.223;1.338 | pathogenic strong |
| 0.228 | [0.072;0.351 | indeterminate |
| -0.874 | [-0.930;-0.81 | benign supporting |
| 1.048 | [-0.334;4.59 | pathogenic moderate |
| -0.610 | [-0.610;-0.60 | benign supporting |
| 0.783 | [-1.082;5.04 | pathogenic moderate |
| -1.092 | [-1.164;-1.04 | benign supporting |
| -1.092 | [-1.100;-1.08 | benign supporting |
| -1.096 | [-1.103;-1.09 | benign supporting |
| -0.520 | [-0.521;-0.52 | benign supporting |
| -0.887 | [-1.074;-0.63 | benign supporting |
| -1.091 | [-1.115;-1.08 | benign supporting |
| 1.276 | [1.258;1.296 | pathogenic strong |
| -0.361 | [-0.387;-0.33 | benign supporting |
| -1.095 | [-1.103;-1.08 | benign supporting |

| | | |
|---|---|---|
| -0.388 | [-0.399;-0.37 | benign supporting |
| -0.666 | [-0.731;-0.62 | benign supporting |
| 1.276 | [1.213;1.357 | pathogenic strong |
| -0.587 | [-0.688;-0.48 | benign supporting |
| 1.276 | [1.203;1.371 | pathogenic strong |
| -3.225 | [-3.519;0.00 | benign strong |
| -1.094 | [-1.105;-1.08 | benign supporting |
| -0.739 | [-0.764;-0.71 | benign supporting |
| 1.276 | [0.584;5.097 | pathogenic strong |
| 0.749 | [0.696;0.802 | pathogenic moderate |
| 0.845 | [0.465;1.116 | pathogenic moderate |
| 1.276 | [0.804;5.064 | pathogenic strong |
| 1.215 | [1.112;1.344 | pathogenic moderate |
| -0.288 | [-0.358;-0.18 | indeterminate |
| -0.657 | [-0.672;-0.64 | benign supporting |
| -0.965 | [-0.969;-0.96 | benign supporting |
| -0.858 | [-0.935;-0.76 | benign supporting |
| -1.083 | [-1.111;-1.05 | benign supporting |
| 1.236 | [1.113;1.406 | pathogenic moderate |
| 1.147 | [1.137;1.156 | pathogenic moderate |
| -0.327 | [-0.332;-0.32 | benign supporting |
| -0.989 | [-0.995;-0.98 | benign supporting |
| 1.276 | [0.447;5.096 | pathogenic strong |
| -0.480 | [-0.503;-0.45 | benign supporting |
| 1.276 | [0.951;4.946 | pathogenic strong |
| 0.699 | [-0.412;1.41 | pathogenic moderate |
| -0.746 | [-0.805;-0.69 | benign supporting |
| 1.102 | [0.835;1.197 | pathogenic moderate |
| -1.111 | [-1.113;-1.11 | benign supporting |
| 0.348 | [0.309;0.386 | pathogenic supporting |
| -1.085 | [-1.124;-1.08 | benign supporting |
| -0.523 | [-0.622;-0.42 | benign supporting |
| 1.276 | [0.853;5.064 | pathogenic strong |
| -0.388 | [-0.425;-0.34 | benign supporting |
| -0.645 | [-0.652;-0.63 | benign supporting |
| -0.433 | [-0.485;-0.39 | benign supporting |
| -1.164 | [-1.169;-1.11 | benign supporting |
| -0.940 | [-0.977;-0.90 | benign supporting |
| -0.930 | [-0.960;-0.89 | benign supporting |
| 0.430 | [0.407;0.453 | pathogenic supporting |
| -1.082 | [-1.093;-1.07 | benign supporting |
| -0.946 | [-0.950;-0.94 | benign supporting |
| -0.832 | [-0.841;-0.82 | benign supporting |

| | | |
|---|---|---|
| -0.520 | [-0.530;-0.50 | benign supporting |
| -3.662 | [-3.821;-1.43 | benign strong |
| -1.166 | [-1.168;-1.16 | benign supporting |
| -1.165 | [-1.166;-1.16 | benign supporting |
| -1.100 | [-1.102;-1.09 | benign supporting |
| 0.719 | [0.699;0.739 | pathogenic moderate |
| -0.827 | [-0.835;-0.81 | benign supporting |
| -0.069 | [-0.078;-0.05 | indeterminate |
| -1.070 | [-1.072;-1.06 | benign supporting |
| 1.276 | [1.212;1.358 | pathogenic strong |
| -1.010 | [-1.053;-0.96 | benign supporting |
| -1.144 | [-1.148;-1.14 | benign supporting |
| -0.643 | [-0.662;-0.62 | benign supporting |
| -1.126 | [-1.145;-1.11 | benign supporting |
| -1.084 | [-1.087;-1.08 | benign supporting |
| -1.167 | [-1.169;-1.01 | benign supporting |
| -0.477 | [-0.537;-0.42 | benign supporting |
| 1.276 | [1.227;1.334 | pathogenic strong |
| 0.525 | [0.077;0.796 | pathogenic supporting |
| -1.169 | [-1.170;-1.16 | benign supporting |
| 1.276 | [0.861;5.027 | pathogenic strong |
| -0.872 | [-1.114;-0.34 | benign supporting |
| 0.312 | [0.134;0.452 | indeterminate |
| -0.190 | [-0.308;-0.05 | indeterminate |
| -0.861 | [-0.983;-0.70 | benign supporting |
| -0.128 | [-0.173;-0.08 | indeterminate |
| -0.655 | [-0.660;-0.65 | benign supporting |
| 0.715 | [-0.053;1.135 | pathogenic moderate |
| 0.513 | [-0.065;0.916 | pathogenic supporting |
| -1.104 | [-1.115;-1.06 | benign supporting |
| 1.276 | [1.171;1.425 | pathogenic strong |
| -0.548 | [-0.559;-0.53 | benign supporting |
| -0.088 | [-0.107;-0.06 | indeterminate |
| -0.089 | [-0.150;-0.02 | indeterminate |
| -1.095 | [-1.167;-0.78 | benign supporting |
| -1.060 | [-1.110;-0.92 | benign supporting |
| 1.276 | [-0.166;5.101 | pathogenic strong |
| 0.189 | [-1.095;4.948 | indeterminate |
| 1.276 | [1.240;1.318 | pathogenic strong |
| 0.964 | [-0.519;5.026 | pathogenic moderate |
| 1.276 | [0.721;5.084 | pathogenic strong |
| 1.177 | [1.130;1.229 | pathogenic moderate |
| 1.276 | [1.196;1.384 | pathogenic strong |

| | | |
|---|---|---|
| 1.276 | [1.160;1.463 | pathogenic strong |
| 1.202 | [1.148;1.261 | pathogenic moderate |
| 1.276 | [1.254;1.301 | pathogenic strong |
| -1.095 | [-1.098;-1.09 | benign supporting |
| -0.938 | [-0.961;-0.91 | benign supporting |
| 1.230 | [0.125;5.076 | pathogenic moderate |
| -0.021 | [-0.166;0.363 | indeterminate |
| -0.164 | [-0.452;0.692 | indeterminate |
| 1.276 | [1.244;1.312 | pathogenic strong |
| -0.176 | [-0.439;0.609 | indeterminate |
| -0.924 | [-0.941;-0.90 | benign supporting |
| 1.276 | [1.257;1.297 | pathogenic strong |
| -0.537 | [-1.113;1.038 | benign supporting |
| -0.398 | [-0.599;-0.16 | benign supporting |
| 0.347 | [-0.340;1.121 | pathogenic supporting |
| 1.126 | [1.088;1.155 | pathogenic moderate |
| -1.095 | [-1.113;-1.08 | benign supporting |
| 1.276 | [-0.183;5.105 | pathogenic strong |
| -0.045 | [-0.650;1.230 | indeterminate |
| -1.105 | [-1.108;-1.10 | benign supporting |
| 1.171 | [-0.312;5.077 | pathogenic moderate |
| -1.027 | [-1.096;-0.88 | benign supporting |
| -1.112 | [-1.167;-0.54 | benign supporting |
| -1.071 | [-1.076;-1.06 | benign supporting |
| 1.116 | [0.875;1.212 | pathogenic moderate |
| 1.276 | [1.270;1.283 | pathogenic strong |
| 1.276 | [1.271;1.281 | pathogenic strong |
| 1.276 | [1.272;1.280 | pathogenic strong |
| 1.276 | [1.226;1.337 | pathogenic strong |
| 1.276 | [1.149;1.491 | pathogenic strong |
| 1.276 | [1.215;1.353 | pathogenic strong |
| 1.276 | [1.263;1.290 | pathogenic strong |
| 1.276 | [1.228;1.333 | pathogenic strong |
| 1.276 | [1.264;1.289 | pathogenic strong |
| 1.276 | [1.261;1.292 | pathogenic strong |
| 1.276 | [1.141;1.514 | pathogenic strong |
| 1.276 | [1.261;1.292 | pathogenic strong |
| 1.276 | [1.253;1.301 | pathogenic strong |
| 1.276 | [1.240;1.320 | pathogenic strong |
| 1.276 | [1.264;1.289 | pathogenic strong |
| 1.276 | [1.204;1.368 | pathogenic strong |
| 1.276 | [1.264;1.288 | pathogenic strong |
| 1.276 | [1.258;1.296 | pathogenic strong |

| | | |
|---|---|---|
| 1.276 | [1.204;1.368] | pathogenic strong |
| -1.087 | [-1.126;-1.08 | benign supporting |
| -1.097 | [-1.115;-1.00 | benign supporting |
| -0.197 | [-0.230;-0.16 | indeterminate |
| 0.909 | [0.863;0.960] | pathogenic moderate |
| -0.442 | [-1.164;4.557 | benign supporting |
| -0.221 | [-0.240;-0.20 | indeterminate |
| -1.107 | [-1.115;-1.09 | benign supporting |
| -0.997 | [-1.012;-0.97 | benign supporting |
| -0.907 | [-0.966;-0.83 | benign supporting |
| -1.033 | [-1.051;-1.01 | benign supporting |
| -0.951 | [-0.955;-0.94 | benign supporting |
| -0.384 | [-0.395;-0.37 | benign supporting |
| -1.132 | [-1.134;-1.13 | benign supporting |
| -0.395 | [-1.091;1.135 | benign supporting |
| -1.102 | [-1.115;-1.01 | benign supporting |
| 1.243 | [1.204;1.284] | pathogenic moderate |
| -0.729 | [-1.111;-0.21 | benign supporting |
| 1.276 | [1.262;1.290] | pathogenic strong |
| 1.203 | [1.127;1.286] | pathogenic moderate |
| -0.908 | [-1.093;-0.61 | benign supporting |
| -0.679 | [-0.967;-0.48 | benign supporting |
| 0.448 | [0.254;0.606] | pathogenic supporting |
| -1.084 | [-1.086;-1.08 | benign supporting |
| -0.320 | [-0.449;-0.10 | benign supporting |
| -1.133 | [-1.169;-1.08 | benign supporting |
| -1.032 | [-1.058;-1.00 | benign supporting |
| -1.086 | [-1.088;-1.08 | benign supporting |
| -1.085 | [-1.086;-1.08 | benign supporting |
| -1.081 | [-1.169;-0.96 | benign supporting |
| -1.144 | [-1.169;-1.11 | benign supporting |
| -0.505 | [-0.526;-0.48 | benign supporting |
| 1.276 | [0.174;5.102] | pathogenic strong |
| -1.141 | [-1.145;-1.13 | benign supporting |
| -1.164 | [-1.169;-1.05 | benign supporting |
| -0.666 | [-0.907;-0.51 | benign supporting |
| 1.229 | [1.108;1.396] | pathogenic moderate |
| -0.618 | [-0.652;-0.58 | benign supporting |
| -1.098 | [-1.114;-1.08 | benign supporting |
| -0.420 | [-0.430;-0.41 | benign supporting |
| 1.244 | [1.013;1.577] | pathogenic moderate |
| -1.084 | [-1.086;-1.08 | benign supporting |
| -0.567 | [-0.575;-0.55 | benign supporting |

| | | |
|---|---|---|
| -1.085 | [-1.085;-1.08 | benign supporting |
| 0.610 | [0.599;0.620] | pathogenic supporting |
| -1.067 | [-1.076;-1.04 | benign supporting |
| -1.151 | [-1.152;-1.15 | benign supporting |
| -0.446 | [-0.448;-0.44 | benign supporting |
| -1.050 | [-1.051;-1.04 | benign supporting |
| -0.459 | [-0.472;-0.44 | benign supporting |
| -1.092 | [-1.097;-1.08 | benign supporting |
| -0.613 | [-0.636;-0.58 | benign supporting |
| -0.589 | [-0.592;-0.58 | benign supporting |
| -1.089 | [-1.090;-1.08 | benign supporting |
| -0.405 | [-0.408;-0.40 | benign supporting |
| 1.276 | [1.248;1.307] | pathogenic strong |
| -0.616 | [-0.638;-0.59 | benign supporting |
| 1.276 | [1.271;1.282] | pathogenic strong |
| -0.342 | [-0.358;-0.32 | benign supporting |
| -0.715 | [-0.717;-0.71 | benign supporting |
| -1.157 | [-1.160;-1.15 | benign supporting |
| -0.737 | [-0.803;-0.68 | benign supporting |
| -1.086 | [-1.098;-1.08 | benign supporting |
| 1.137 | [-0.371;5.06( | pathogenic moderate |
| -0.916 | [-1.046;-0.83 | benign supporting |
| 1.276 | [0.585;5.094] | pathogenic strong |
| -1.089 | [-1.093;-1.08 | benign supporting |
| -1.078 | [-3.543;0.00( | benign supporting |
| -0.899 | [-0.935;-0.85 | benign supporting |
| 0.053 | [-0.582;1.232 | indeterminate |
| -0.933 | [-0.937;-0.92 | benign supporting |
| -0.921 | [-2.976;-0.83 | benign supporting |
| -1.046 | [-1.081;-1.01 | benign supporting |
| 0.594 | [-0.387;1.274 | pathogenic supporting |
| -0.950 | [-0.954;-0.94 | benign supporting |
| -0.518 | [-0.559;-0.47 | benign supporting |
| -1.084 | [-1.167;-1.02 | benign supporting |
| -1.114 | [-1.116;-1.10 | benign supporting |
| -1.112 | [-1.149;-1.09 | benign supporting |
| -1.039 | [-1.071;-1.00 | benign supporting |
| -1.066 | [-1.167;-0.99 | benign supporting |
| -1.085 | [-1.093;-1.08 | benign supporting |
| -0.641 | [-0.696;-0.60 | benign supporting |
| -1.166 | [-1.170;-1.14 | benign supporting |
| 0.827 | [0.538;1.078] | pathogenic moderate |
| -1.071 | [-1.114;-0.80 | benign supporting |

| | | |
|---|---|---|
| -1.103 | [-1.143;-0.99 | benign supporting |
| -0.261 | [-0.567;0.614 | indeterminate |
| -0.564 | [-0.629;-0.49 | benign supporting |
| -1.068 | [-1.080;-1.04 | benign supporting |
| -0.809 | [-1.107;-0.38 | benign supporting |
| -0.967 | [-1.086;-0.75 | benign supporting |
| -1.123 | [-1.169;-1.08 | benign supporting |
| 1.025 | [0.891;1.114] | pathogenic moderate |
| -1.164 | [-1.168;-1.16 | benign supporting |
| -0.892 | [-0.932;-0.84 | benign supporting |
| -1.092 | [-1.101;-1.08 | benign supporting |
| -1.089 | [-1.096;-1.08 | benign supporting |
| -0.232 | [-0.281;-0.17 | indeterminate |
| -0.950 | [-0.975;-0.92 | benign supporting |
| -0.285 | [-0.340;-0.21 | indeterminate |
| 0.720 | [0.396;0.966] | pathogenic moderate |
| 0.586 | [0.382;0.742] | pathogenic supporting |
| -0.058 | [-0.089;-0.02 | indeterminate |
| -0.050 | [-0.187;0.304 | indeterminate |
| -1.066 | [-1.105;-0.96 | benign supporting |
| 1.143 | [0.457;1.498] | pathogenic moderate |
| -0.684 | [-0.861;-0.58 | benign supporting |
| -1.045 | [-1.090;-0.94 | benign supporting |
| -0.020 | [-0.366;0.860 | indeterminate |
| 1.276 | [1.099;1.685] | pathogenic strong |
| -1.073 | [-1.086;-1.05 | benign supporting |
| 1.275 | [1.252;1.299] | pathogenic strong |
| -0.588 | [-0.630;-0.54 | benign supporting |
| 0.360 | [0.141;0.529] | pathogenic supporting |
| -0.734 | [-0.966;-0.56 | benign supporting |
| -1.089 | [-1.112;-1.06 | benign supporting |
| -1.087 | [-1.091;-1.08 | benign supporting |
| -0.844 | [-0.916;-0.75 | benign supporting |
| -0.612 | [-0.635;-0.58 | benign supporting |
| -1.165 | [-1.169;-1.03 | benign supporting |
| -1.090 | [-1.103;-1.07 | benign supporting |
| -1.102 | [-1.107;-1.09 | benign supporting |
| -1.005 | [-1.040;-0.96 | benign supporting |
| -0.928 | [-1.081;-0.67 | benign supporting |
| -1.039 | [-1.085;-0.94 | benign supporting |
| -1.109 | [-1.115;-1.09 | benign supporting |
| -1.088 | [-1.102;-1.08 | benign supporting |
| -0.974 | [-1.028;-0.91 | benign supporting |

| | | |
|---|---|---|
| 0.854 | [0.765;0.954 | pathogenic moderate |
| -0.355 | [-0.770;0.61 | benign supporting |
| -1.081 | [-1.088;-1.07 | benign supporting |
| 1.276 | [1.254;1.300 | pathogenic strong |
| -1.073 | [-1.090;-1.04 | benign supporting |
| -1.102 | [-1.115;-1.09 | benign supporting |
| -1.085 | [-1.091;-1.08 | benign supporting |
| -0.399 | [-1.114;1.40 | benign supporting |
| -0.840 | [-0.849;-0.83 | benign supporting |
| -1.070 | [-1.099;-1.01 | benign supporting |
| 1.140 | [1.097;1.183 | pathogenic moderate |
| -1.168 | [-1.169;-1.16 | benign supporting |
| -1.093 | [-1.115;-1.03 | benign supporting |
| 1.276 | [1.113;1.630 | pathogenic strong |
| -0.446 | [-0.454;-0.43 | benign supporting |
| -0.679 | [-0.849;-0.57 | benign supporting |
| -1.166 | [-1.169;-1.03 | benign supporting |
| -1.104 | [-1.115;-1.08 | benign supporting |
| 0.486 | [0.213;0.697 | pathogenic supporting |
| -1.110 | [-1.116;-1.10 | benign supporting |
| -0.942 | [-0.954;-0.93 | benign supporting |
| -1.029 | [-1.052;-1.00 | benign supporting |
| -0.707 | [-1.037;-0.47 | benign supporting |
| 1.276 | [1.237;1.321 | pathogenic strong |
| -1.086 | [-1.167;-0.92 | benign supporting |
| -1.102 | [-1.111;-1.09 | benign supporting |
| -0.588 | [-0.694;-0.47 | benign supporting |
| -0.324 | [-0.410;-0.19 | benign supporting |
| -1.156 | [-1.166;-1.14 | benign supporting |
| 1.276 | [1.076;1.803 | pathogenic strong |
| -1.104 | [-1.167;-0.49 | benign supporting |
| -0.942 | [-3.799;0.00 | benign supporting |
| -1.064 | [-1.102;-1.03 | benign supporting |
| -0.872 | [-0.894;-0.85 | benign supporting |
| -1.097 | [-1.099;-1.09 | benign supporting |
| 1.276 | [1.104;1.660 | pathogenic strong |
| -0.642 | [-1.115;0.57 | benign supporting |
| -0.612 | [-0.810;-0.46 | benign supporting |
| -0.220 | [-0.436;0.41 | indeterminate |
| -0.809 | [-0.870;-0.73 | benign supporting |
| -0.832 | [-0.847;-0.81 | benign supporting |
| -1.119 | [-1.135;-1.10 | benign supporting |
| -0.620 | [-0.641;-0.59 | benign supporting |

| | | |
|---|---|---|
| -0.383 | [-0.479;-0.27 | benign supporting |
| 1.276 | [1.119;1.608] | pathogenic strong |
| -0.359 | [-0.677;0.382 | benign supporting |
| -1.169 | [-1.170;-1.16 | benign supporting |
| -0.592 | [-0.644;-0.53 | benign supporting |
| -1.000 | [-1.025;-0.96 | benign supporting |
| -0.911 | [-0.921;-0.90 | benign supporting |
| 1.276 | [1.112;1.618] | pathogenic strong |
| 1.276 | [1.182;1.411] | pathogenic strong |
| -1.165 | [-1.168;-0.88 | benign supporting |
| -1.081 | [-1.105;-1.06 | benign supporting |
| -1.160 | [-1.169;-1.07 | benign supporting |
| 1.276 | [0.635;5.087] | pathogenic strong |
| 1.276 | [-0.207;5.103 | pathogenic strong |
| 0.406 | [-0.518;1.308 | pathogenic supporting |
| -1.089 | [-1.096;-1.08 | benign supporting |
| 1.088 | [0.966;1.141] | pathogenic moderate |
| -0.511 | [-1.046;-0.01 | benign supporting |
| -1.095 | [-1.166;-0.37 | benign supporting |
| 1.276 | [1.202;1.374] | pathogenic strong |
| -1.061 | [-1.067;-1.05 | benign supporting |
| 1.276 | [0.407;5.095] | pathogenic strong |
| -1.066 | [-1.114;-0.68 | benign supporting |
| 1.276 | [1.177;1.418] | pathogenic strong |
| -1.018 | [-1.030;-1.00 | benign supporting |
| -0.592 | [-0.727;-0.46 | benign supporting |
| -1.112 | [-1.116;-1.10 | benign supporting |
| -0.964 | [-1.081;-0.89 | benign supporting |
| -0.407 | [-0.443;-0.37 | benign supporting |
| 1.276 | [1.219;1.349] | pathogenic strong |
| -0.658 | [-1.164;1.159 | benign supporting |
| -0.852 | [-0.902;-0.83 | benign supporting |
| -0.862 | [-1.164;0.774 | benign supporting |
| -0.876 | [-0.880;-0.87 | benign supporting |
| -1.023 | [-1.110;-0.76 | benign supporting |
| 1.125 | [0.858;1.241] | pathogenic moderate |
| -0.970 | [-0.973;-0.96 | benign supporting |
| -0.959 | [-0.995;-0.93 | benign supporting |
| -1.170 | [-1.169;-0.96 | benign supporting |
| -1.094 | [-1.108;-1.08 | benign supporting |
| -0.275 | [-0.512;0.389 | indeterminate |
| -1.164 | [-1.165;-1.16 | benign supporting |
| -0.764 | [-0.789;-0.74 | benign supporting |

| | | |
|---|---|---|
| -1.056 | [-1.166;0.184 | benign supporting |
| 0.731 | [-0.259;1.262 | pathogenic moderate |
| -0.194 | [-0.291;-0.08 | indeterminate |
| -1.095 | [-1.107;-1.08 | benign supporting |
| -1.085 | [-1.105;-1.08 | benign supporting |
| -1.046 | [-1.069;-1.01 | benign supporting |
| -0.911 | [-1.165;0.629 | benign supporting |
| -0.356 | [-0.748;0.559 | benign supporting |
| -1.104 | [-1.108;-1.10 | benign supporting |
| -1.092 | [-1.102;-1.08 | benign supporting |
| -1.547 | [-3.834;-1.05 | benign strong |
| -0.216 | [-0.288;-0.12 | indeterminate |
| -0.809 | [-1.112;-0.30 | benign supporting |
| -1.103 | [-1.158;-1.08 | benign supporting |
| -1.000 | [-1.051;-0.96 | benign supporting |
| -1.082 | [-1.152;-1.03 | benign supporting |
| 1.276 | [-0.422;5.091 | pathogenic strong |
| 1.118 | [1.040;1.159] | pathogenic moderate |
| -1.007 | [-1.166;-0.91 | benign supporting |
| -0.923 | [-0.927;-0.91 | benign supporting |
| -0.895 | [-0.905;-0.88 | benign supporting |
| -1.040 | [-1.048;-1.03 | benign supporting |
| -1.124 | [-1.169;-0.96 | benign supporting |
| -0.916 | [-0.971;-0.84 | benign supporting |
| -0.928 | [-1.063;-0.83 | benign supporting |
| -1.126 | [-2.892;0.000 | benign supporting |
| -0.877 | [-0.969;-0.76 | benign supporting |
| -1.030 | [-1.061;-1.00 | benign supporting |
| -0.332 | [-0.356;-0.30 | benign supporting |
| -1.084 | [-1.086;-1.08 | benign supporting |
| -0.585 | [-0.607;-0.56 | benign supporting |
| -0.628 | [-0.830;-0.49 | benign supporting |
| -0.950 | [-0.973;-0.92 | benign supporting |
| -1.022 | [-1.031;-1.01 | benign supporting |
| -1.097 | [-1.115;-1.07 | benign supporting |
| -1.135 | [-1.148;-1.12 | benign supporting |
| -1.101 | [-1.108;-1.09 | benign supporting |
| -1.089 | [-1.092;-1.08 | benign supporting |
| -0.397 | [-0.425;-0.36 | benign supporting |
| -1.057 | [-1.073;-1.02 | benign supporting |
| 1.276 | [1.204;1.371] | pathogenic strong |
| -1.086 | [-1.114;-1.08 | benign supporting |
| -1.085 | [-1.087;-1.08 | benign supporting |

| | | |
|---|---|---|
| -1.112 | [-1.115;-1.10 | benign supporting |
| -0.294 | [-0.353;-0.21 | indeterminate |
| 1.276 | [1.251;1.304 | pathogenic strong |
| 1.276 | [1.256;1.297 | pathogenic strong |
| 1.153 | [1.134;1.172 | pathogenic moderate |
| -0.838 | [-0.852;-0.82 | benign supporting |
| 1.276 | [1.229;1.331 | pathogenic strong |
| -0.133 | [-0.304;0.20 | indeterminate |
| 1.109 | [1.064;1.132 | pathogenic moderate |
| 1.263 | [1.232;1.295 | pathogenic moderate |
| 1.276 | [1.256;1.297 | pathogenic strong |
| 1.228 | [1.216;1.239 | pathogenic moderate |
| 1.276 | [1.244;1.313 | pathogenic strong |
| 1.276 | [1.258;1.295 | pathogenic strong |
| 1.276 | [1.249;1.306 | pathogenic strong |
| 1.261 | [1.226;1.297 | pathogenic moderate |
| 1.138 | [1.049;1.196 | pathogenic moderate |
| 1.274 | [1.226;1.329 | pathogenic strong |
| 1.276 | [1.244;1.311 | pathogenic strong |
| 1.276 | [1.251;1.304 | pathogenic strong |
| 1.221 | [1.200;1.243 | pathogenic moderate |
| 1.276 | [1.210;1.360 | pathogenic strong |
| -0.742 | [-0.832;-0.67 | benign supporting |
| -0.336 | [-0.497;-0.09 | benign supporting |
| 1.206 | [1.140;1.277 | pathogenic moderate |
| -0.626 | [-0.700;-0.56 | benign supporting |
| -1.090 | [-1.099;-1.08 | benign supporting |
| -0.098 | [-0.200;0.02 | indeterminate |
| -1.106 | [-1.115;-1.09 | benign supporting |
| -1.106 | [-1.115;-1.08 | benign supporting |
| -1.083 | [-1.093;-1.07 | benign supporting |
| 1.276 | [1.183;1.408 | pathogenic strong |
| -0.941 | [-1.069;-0.76 | benign supporting |
| 0.905 | [0.649;1.114 | pathogenic moderate |
| -1.055 | [-1.069;-1.03 | benign supporting |
| -0.077 | [-0.126;-0.01 | indeterminate |
| 1.276 | [0.000;5.102 | pathogenic strong |
| -0.848 | [-0.958;-0.70 | benign supporting |
| 1.276 | [0.979;2.367 | pathogenic strong |
| -0.412 | [-0.450;-0.37 | benign supporting |
| -0.107 | [-0.161;-0.05 | indeterminate |
| -1.102 | [-1.114;-1.09 | benign supporting |
| -0.944 | [-0.974;-0.91 | benign supporting |

| | | |
|---|---|---|
| -0.698 | [-0.707;-0.69 | benign supporting |
| -1.081 | [-1.165;-0.53 | benign supporting |
| -1.085 | [-1.165;-1.08 | benign supporting |
| -1.084 | [-1.168;-0.67 | benign supporting |
| -1.101 | [-1.106;-1.09 | benign supporting |
| -0.953 | [-0.960;-0.94 | benign supporting |
| -1.044 | [-1.049;-1.04 | benign supporting |
| -1.083 | [-1.132;-0.71 | benign supporting |
| 1.276 | [0.948;3.564 | pathogenic strong |
| 1.276 | [1.256;1.298 | pathogenic strong |
| 1.164 | [1.144;1.187 | pathogenic moderate |
| 1.223 | [1.158;1.295 | pathogenic moderate |
| -0.837 | [-0.891;-0.83 | benign supporting |
| -0.612 | [-1.041;-0.30 | benign supporting |
| 1.276 | [1.238;1.318 | pathogenic strong |
| 1.217 | [1.121;1.332 | pathogenic moderate |
| 1.276 | [-0.040;5.10 | pathogenic strong |
| 1.276 | [1.163;1.443 | pathogenic strong |
| 1.001 | [0.903;1.075 | pathogenic moderate |
| 1.054 | [-1.088;5.04 | pathogenic moderate |
| 0.307 | [-0.052;0.63 | indeterminate |
| -1.106 | [-1.115;-1.09 | benign supporting |
| -0.974 | [-1.128;-0.41 | benign supporting |
| 1.276 | [1.024;2.083 | pathogenic strong |
| -0.943 | [-0.945;-0.94 | benign supporting |
| -1.086 | [-1.089;-1.08 | benign supporting |
| -0.340 | [-0.439;-0.18 | benign supporting |
| 1.276 | [-0.101;5.10 | pathogenic strong |
| -0.977 | [-0.986;-0.97 | benign supporting |
| 1.276 | [1.201;1.373 | pathogenic strong |
| -0.649 | [-0.659;-0.64 | benign supporting |
| -0.874 | [-0.879;-0.86 | benign supporting |
| 1.276 | [0.706;5.079 | pathogenic strong |
| -1.030 | [-1.164;-0.41 | benign supporting |
| -1.087 | [-1.120;-1.08 | benign supporting |
| 0.763 | [0.653;0.863 | pathogenic moderate |
| 1.276 | [1.193;1.389 | pathogenic strong |
| -1.098 | [-1.100;-1.09 | benign supporting |
| -0.350 | [-0.448;-0.21 | benign supporting |
| -1.066 | [-1.084;-1.02 | benign supporting |
| -0.378 | [-0.430;-0.31 | benign supporting |
| -0.617 | [-0.640;-0.59 | benign supporting |
| -1.085 | [-1.086;-1.08 | benign supporting |

| | | |
|---|---|---|
| -0.982 | [-0.995;-0.97 | benign supporting |
| -0.631 | [-0.638;-0.62 | benign supporting |
| -1.067 | [-1.069;-1.06 | benign supporting |
| -0.384 | [-0.407;-0.35 | benign supporting |
| -1.101 | [-1.104;-1.09 | benign supporting |
| -0.662 | [-0.670;-0.65 | benign supporting |
| 1.201 | [1.091;1.324 | pathogenic moderate |
| -0.804 | [-0.827;-0.78 | benign supporting |
| -0.406 | [-0.439;-0.37 | benign supporting |
| -0.087 | [-0.425;0.85 | indeterminate |
| -0.999 | [-1.112;-0.56 | benign supporting |
| 1.276 | [1.258;1.295 | pathogenic strong |
| -0.593 | [-0.600;-0.58 | benign supporting |
| 1.234 | [1.219;1.248 | pathogenic moderate |
| -0.463 | [-0.465;-0.46 | benign supporting |
| -0.413 | [-0.647;-0.14 | benign supporting |
| -0.514 | [-0.870;-0.24 | benign supporting |
| 1.276 | [1.107;1.655 | pathogenic strong |
| -1.027 | [-1.036;-1.01 | benign supporting |
| -1.107 | [-1.110;-1.10 | benign supporting |
| -1.090 | [-1.093;-1.08 | benign supporting |
| -1.041 | [-1.043;-1.03 | benign supporting |
| 1.276 | [1.189;1.394 | pathogenic strong |
| -0.182 | [-0.205;-0.15 | indeterminate |
| 1.276 | [1.183;1.404 | pathogenic strong |
| -0.974 | [-1.015;-0.93 | benign supporting |
| 1.064 | [0.941;1.131 | pathogenic moderate |
| -0.885 | [-0.945;-0.81 | benign supporting |
| 1.100 | [0.751;1.223 | pathogenic moderate |
| 1.276 | [1.179;1.411 | pathogenic strong |
| 1.234 | [0.935;1.610 | pathogenic moderate |
| -0.173 | [-0.212;-0.13 | indeterminate |
| 1.276 | [1.214;1.356 | pathogenic strong |
| -1.086 | [-1.090;-1.08 | benign supporting |
| -1.079 | [-1.166;-0.40 | benign supporting |
| 1.061 | [-0.503;5.04 | pathogenic moderate |
| -1.130 | [-1.169;-1.08 | benign supporting |
| -0.908 | [-0.924;-0.89 | benign supporting |
| -1.064 | [-1.096;-0.97 | benign supporting |
| 1.276 | [-1.089;5.06 | pathogenic strong |
| -0.281 | [-1.109;1.95 | indeterminate |
| -0.073 | [-0.270;0.42 | indeterminate |
| 1.276 | [1.198;1.379 | pathogenic strong |

| | | |
|---|---|---|
| 1.026 | [0.741;1.163 | pathogenic moderate |
| -1.040 | [-1.067;-1.01 | benign supporting |
| -1.076 | [-1.085;-1.06 | benign supporting |
| 0.501 | [0.021;0.786 | pathogenic supporting |
| -1.075 | [-1.081;-1.06 | benign supporting |
| -1.099 | [-1.120;-1.08 | benign supporting |
| -0.453 | [-0.482;-0.42 | benign supporting |
| -1.029 | [-1.042;-1.01 | benign supporting |
| -1.085 | [-1.091;-1.08 | benign supporting |
| -0.407 | [-0.421;-0.39 | benign supporting |
| -0.638 | [-0.716;-0.57 | benign supporting |
| -0.840 | [-0.896;-0.83 | benign supporting |
| -0.562 | [-0.579;-0.54 | benign supporting |
| -0.031 | [-0.066;0.01 | indeterminate |
| -0.342 | [-0.395;-0.27 | benign supporting |
| -0.556 | [-0.839;-0.36 | benign supporting |
| -0.504 | [-0.764;-0.29 | benign supporting |
| 1.276 | [0.904;4.996 | pathogenic strong |
| -0.492 | [-0.589;-0.40 | benign supporting |
| -0.675 | [-0.695;-0.65 | benign supporting |
| 0.764 | [-0.021;1.15 | pathogenic moderate |
| -1.053 | [-1.077;-1.00 | benign supporting |
| -1.086 | [-1.088;-1.08 | benign supporting |
| -0.748 | [-0.891;-0.63 | benign supporting |
| -0.239 | [-0.360;-0.06 | indeterminate |
| 0.942 | [0.802;1.063 | pathogenic moderate |
| -0.916 | [-0.934;-0.89 | benign supporting |
| -0.729 | [-0.751;-0.70 | benign supporting |
| -0.501 | [-0.786;-0.27 | benign supporting |
| -0.932 | [-0.939;-0.92 | benign supporting |
| -0.608 | [-1.113;0.73 | benign supporting |
| 1.046 | [0.977;1.106 | pathogenic moderate |
| -1.089 | [-1.099;-1.08 | benign supporting |
| -0.837 | [-0.906;-0.83 | benign supporting |
| -0.832 | [-1.100;-0.46 | benign supporting |
| 1.276 | [1.240;1.317 | pathogenic strong |
| -1.130 | [-1.169;-0.87 | benign supporting |
| -1.085 | [-1.089;-1.08 | benign supporting |
| -1.001 | [-1.024;-0.96 | benign supporting |
| -1.084 | [-1.144;-1.08 | benign supporting |
| 1.276 | [0.520;5.098 | pathogenic strong |
| 0.465 | [-0.464;1.30 | pathogenic supporting |
| -1.165 | [-1.169;-1.16 | benign supporting |

| | | |
|---|---|---|
| -0.891 | [-0.907;-0.87 | benign supporting |
| 1.061 | [-0.565;5.049 | pathogenic moderate |
| -0.365 | [-0.507;-0.16 | benign supporting |
| -1.115 | [-1.169;-1.03 | benign supporting |
| -0.979 | [-0.987;-0.97 | benign supporting |
| 1.276 | [0.000;5.105] | pathogenic strong |
| 1.276 | [-0.006;5.100 | pathogenic strong |
| 0.000 | [0.000;0.000] | indeterminate |
| -0.880 | [-0.882;-0.87 | benign supporting |
| -0.533 | [-0.586;-0.47 | benign supporting |
| 0.610 | [0.057;0.951] | pathogenic supporting |
| -0.123 | [-0.971;1.353 | indeterminate |
| -0.979 | [-0.984;-0.97 | benign supporting |
| -0.881 | [-0.902;-0.86 | benign supporting |
| -0.586 | [-1.111;0.638 | benign supporting |
| -1.098 | [-1.169;-0.95 | benign supporting |
| -0.933 | [-0.988;-0.87 | benign supporting |
| -1.076 | [-1.079;-1.07 | benign supporting |
| 1.276 | [1.216;1.349] | pathogenic strong |
| -1.015 | [-1.024;-1.00 | benign supporting |
| -1.166 | [-1.170;-1.12 | benign supporting |
| -1.091 | [-1.168;-1.08 | benign supporting |
| -1.134 | [-1.169;-1.10 | benign supporting |
| 1.010 | [0.941;1.064] | pathogenic moderate |
| 0.729 | [0.013;1.117] | pathogenic moderate |
| 1.276 | [1.208;1.359] | pathogenic strong |
| -1.165 | [-1.166;-1.16 | benign supporting |
| 1.259 | [1.217;1.306] | pathogenic moderate |
| -0.634 | [-0.717;-0.57 | benign supporting |
| -0.043 | [-0.193;0.350 | indeterminate |
| 1.276 | [1.237;1.319] | pathogenic strong |
| -0.049 | [-0.198;0.333 | indeterminate |
| -0.599 | [-0.621;-0.57 | benign supporting |
| -0.365 | [-0.378;-0.35 | benign supporting |
| -1.059 | [-1.096;-0.96 | benign supporting |
| -1.110 | [-1.115;-1.10 | benign supporting |
| -1.077 | [-1.115;-0.70 | benign supporting |
| -1.099 | [-1.115;-0.88 | benign supporting |
| -1.065 | [-1.093;-1.04 | benign supporting |
| -1.133 | [-1.169;-1.08 | benign supporting |
| -0.872 | [-0.891;-0.85 | benign supporting |
| -0.749 | [-2.095;4.795 | benign supporting |
| -1.106 | [-1.115;-1.08 | benign supporting |

| | | |
|---|---|---|
| 1.276 | [1.196;1.384 | pathogenic strong |
| -0.142 | [-0.320;0.21 | indeterminate |
| 1.276 | [1.176;1.420 | pathogenic strong |
| -0.485 | [-0.520;-0.45 | benign supporting |
| -0.036 | [-0.442;1.03 | indeterminate |
| 1.230 | [1.050;1.456 | pathogenic moderate |
| 1.276 | [1.180;1.407 | pathogenic strong |
| 1.276 | [1.220;1.343 | pathogenic strong |
| 1.276 | [1.248;1.308 | pathogenic strong |
| -0.022 | [-0.067;0.04 | indeterminate |
| -1.073 | [-1.084;-1.06 | benign supporting |
| -0.186 | [-0.401;0.38 | indeterminate |
| -1.114 | [-1.116;-1.09 | benign supporting |
| -0.935 | [-0.984;-0.88 | benign supporting |
| -0.220 | [-0.275;-0.15 | indeterminate |
| 1.276 | [1.194;1.386 | pathogenic strong |
| -1.005 | [-1.043;-0.95 | benign supporting |
| 0.893 | [0.701;1.079 | pathogenic moderate |
| 1.276 | [-1.096;5.05 | pathogenic strong |
| -1.113 | [-1.127;-1.09 | benign supporting |
| 1.276 | [1.177;1.416 | pathogenic strong |
| -1.083 | [-1.104;-1.06 | benign supporting |
| -0.825 | [-0.841;-0.80 | benign supporting |
| -0.663 | [-0.684;-0.64 | benign supporting |
| -1.166 | [-1.168;-1.16 | benign supporting |
| -1.120 | [-1.169;-1.08 | benign supporting |
| -0.793 | [-0.815;-0.77 | benign supporting |
| -0.855 | [-1.073;-0.59 | benign supporting |
| -0.373 | [-0.663;0.16 | benign supporting |
| -0.676 | [-0.686;-0.66 | benign supporting |
| -1.089 | [-1.093;-1.08 | benign supporting |
| -0.566 | [-0.602;-0.52 | benign supporting |
| -0.837 | [-0.845;-0.83 | benign supporting |
| -0.988 | [-0.997;-0.97 | benign supporting |
| -0.636 | [-0.801;-0.52 | benign supporting |
| -1.114 | [-1.129;-1.10 | benign supporting |
| 0.646 | [-0.091;1.11 | pathogenic moderate |
| -1.029 | [-1.109;-0.83 | benign supporting |
| -1.098 | [-1.115;-1.07 | benign supporting |
| -1.085 | [-1.088;-1.08 | benign supporting |
| -0.182 | [-0.388;0.36 | indeterminate |
| 0.819 | [0.733;0.910 | pathogenic moderate |
| 1.276 | [0.848;5.062 | pathogenic strong |

| | | |
|---|---|---|
| -0.417 | [-0.502;-0.34 | benign supporting |
| 1.206 | [1.153;1.262 | pathogenic moderate |
| -0.976 | [-0.980;-0.97 | benign supporting |
| -1.087 | [-1.092;-1.08 | benign supporting |
| -1.085 | [-1.094;-1.08 | benign supporting |
| 1.166 | [0.250;1.984 | pathogenic moderate |
| -1.079 | [-1.081;-1.07 | benign supporting |
| -0.740 | [-0.878;-0.63 | benign supporting |
| -0.429 | [-0.465;-0.39 | benign supporting |
| -0.940 | [-1.006;-0.86 | benign supporting |
| -0.336 | [-0.341;-0.33 | benign supporting |
| -0.972 | [-0.976;-0.96 | benign supporting |
| -1.004 | [-1.010;-0.99 | benign supporting |
| -1.088 | [-1.095;-1.08 | benign supporting |
| -1.085 | [-1.086;-1.08 | benign supporting |
| -0.371 | [-0.380;-0.36 | benign supporting |
| -0.648 | [-0.850;-0.52 | benign supporting |
| -1.109 | [-1.116;-1.10 | benign supporting |
| -1.164 | [-1.164;-1.16 | benign supporting |
| 1.276 | [1.114;1.612 | pathogenic strong |
| -1.096 | [-1.115;-1.07 | benign supporting |
| 1.078 | [1.039;1.108 | pathogenic moderate |
| -1.062 | [-1.066;-1.05 | benign supporting |
| -0.413 | [-0.419;-0.40 | benign supporting |
| -0.939 | [-0.986;-0.89 | benign supporting |
| -1.112 | [-1.116;-1.10 | benign supporting |
| -1.014 | [-1.098;-0.84 | benign supporting |
| 1.276 | [-1.082;5.078 | pathogenic strong |
| 1.276 | [-1.119;5.004 | pathogenic strong |
| 1.276 | [-1.093;5.056 | pathogenic strong |
| 1.276 | [-1.112;5.026 | pathogenic strong |
| 1.276 | [0.000;5.102 | pathogenic strong |
| -1.093 | [-1.168;-0.47 | benign supporting |
| 1.219 | [-0.194;5.095 | pathogenic moderate |
| 0.647 | [-1.108;5.033 | pathogenic moderate |
| -1.052 | [-1.169;1.243 | benign supporting |
| -0.002 | [-0.234;0.578 | indeterminate |
| 1.276 | [1.173;1.425 | pathogenic strong |
| 1.191 | [-0.384;5.089 | pathogenic moderate |
| 1.276 | [-0.593;5.094 | pathogenic strong |
| 1.276 | [0.000;5.106 | pathogenic strong |
| 1.276 | [0.542;5.099 | pathogenic strong |
| -0.187 | [-1.166;4.914 | indeterminate |

| | | |
|---|---|---|
| 0.165 | [-0.448;1.15 | indeterminate |
| 1.276 | [-0.294;5.10 | pathogenic strong |
| -0.372 | [-0.383;-0.36 | benign supporting |
| -1.110 | [-1.113;-1.10 | benign supporting |
| 0.313 | [0.129;0.450 | indeterminate |
| -0.096 | [-0.131;-0.06 | indeterminate |
| -0.445 | [-0.515;-0.38 | benign supporting |
| -1.143 | [-1.169;-1.09 | benign supporting |
| -0.642 | [-0.706;-0.59 | benign supporting |
| -0.945 | [-0.993;-0.90 | benign supporting |
| -0.543 | [-0.582;-0.50 | benign supporting |
| 0.986 | [0.866;1.076 | pathogenic moderate |
| -1.141 | [-1.156;-1.12 | benign supporting |
| 1.117 | [1.074;1.143 | pathogenic moderate |
| -0.829 | [-0.847;-0.80 | benign supporting |
| -0.756 | [-1.000;-0.56 | benign supporting |
| -0.919 | [-1.027;-0.77 | benign supporting |
| 1.276 | [1.235;1.326 | pathogenic strong |
| -0.763 | [-0.841;-0.68 | benign supporting |
| -0.560 | [-0.575;-0.54 | benign supporting |
| -0.810 | [-0.832;-0.78 | benign supporting |
| -0.776 | [-0.833;-0.71 | benign supporting |
| -0.972 | [-0.975;-0.96 | benign supporting |
| -0.495 | [-0.540;-0.44 | benign supporting |
| 1.110 | [-0.088;1.65 | pathogenic moderate |
| -1.098 | [-1.113;-1.08 | benign supporting |
| 0.394 | [0.077;0.613 | pathogenic supporting |
| -0.271 | [-0.330;-0.19 | indeterminate |
| -1.085 | [-1.152;-1.08 | benign supporting |
| 0.340 | [0.142;0.494 | pathogenic supporting |
| 1.058 | [0.354;1.299 | pathogenic moderate |
| 1.102 | [0.716;1.244 | pathogenic moderate |
| 1.235 | [1.094;1.439 | pathogenic moderate |
| -1.104 | [-1.115;-1.08 | benign supporting |
| 1.192 | [1.162;1.222 | pathogenic moderate |
| 1.276 | [1.154;1.480 | pathogenic strong |
| 0.736 | [-0.118;1.17 | pathogenic moderate |
| -0.644 | [-0.766;-0.56 | benign supporting |
| 1.276 | [1.187;1.396 | pathogenic strong |
| 1.212 | [1.169;1.259 | pathogenic moderate |
| 1.276 | [1.160;1.465 | pathogenic strong |
| 0.683 | [-0.307;1.25 | pathogenic moderate |
| 0.867 | [0.813;0.924 | pathogenic moderate |

| | | |
|---|---|---|
| -1.131 | [-1.168;-1.10 | benign supporting |
| 1.112 | [0.816;1.223 | pathogenic moderate |
| 1.276 | [1.186;1.405 | pathogenic strong |
| 0.898 | [-0.047;1.255 | pathogenic moderate |
| 1.276 | [1.177;1.422 | pathogenic strong |
| 1.276 | [0.762;5.084 | pathogenic strong |
| -0.289 | [-1.131;4.803 | indeterminate |
| 1.106 | [0.706;1.249 | pathogenic moderate |
| -0.341 | [-0.514;-0.08 | benign supporting |
| 1.276 | [0.915;4.967 | pathogenic strong |
| 1.276 | [1.212;1.358 | pathogenic strong |
| 1.276 | [0.000;5.105 | pathogenic strong |
| 0.913 | [0.754;1.061 | pathogenic moderate |
| 1.276 | [0.072;5.100 | pathogenic strong |
| 0.213 | [0.004;0.398 | indeterminate |
| 1.276 | [1.207;1.367 | pathogenic strong |
| 0.019 | [-0.103;0.348 | indeterminate |
| 1.276 | [1.146;1.500 | pathogenic strong |
| 1.276 | [1.237;1.320 | pathogenic strong |
| 1.276 | [1.113;1.644 | pathogenic strong |
| 1.276 | [1.129;1.550 | pathogenic strong |
| -1.088 | [-1.092;-1.08 | benign supporting |
| -0.017 | [-0.370;0.883 | indeterminate |
| -0.670 | [-0.840;-0.57 | benign supporting |
| -0.818 | [-0.927;-0.68 | benign supporting |
| -0.895 | [-0.980;-0.78 | benign supporting |
| -1.098 | [-1.108;-1.08 | benign supporting |
| 1.276 | [1.229;1.331 | pathogenic strong |
| -0.413 | [-0.469;-0.36 | benign supporting |
| 0.582 | [-0.178;1.118 | pathogenic supporting |
| -1.090 | [-1.103;-1.08 | benign supporting |
| -0.853 | [-0.875;-0.83 | benign supporting |
| -0.463 | [-0.514;-0.42 | benign supporting |
| -0.523 | [-0.556;-0.49 | benign supporting |
| -0.939 | [-1.112;-0.47 | benign supporting |
| -1.109 | [-1.115;-1.07 | benign supporting |
| 1.262 | [1.093;1.573 | pathogenic moderate |
| -1.113 | [-1.144;-1.09 | benign supporting |
| -0.604 | [-0.630;-0.57 | benign supporting |
| -0.130 | [-0.597;1.093 | indeterminate |
| -0.086 | [-0.283;0.406 | indeterminate |
| -1.087 | [-1.089;-1.08 | benign supporting |
| -0.959 | [-1.082;-0.74 | benign supporting |

| | | |
|---|---|---|
| 1.276 | [1.138;1.535 | pathogenic strong |
| 1.276 | [1.238;1.321 | pathogenic strong |
| -0.437 | [-0.606;-0.29 | benign supporting |
| -0.945 | [-1.115;-0.41 | benign supporting |
| 1.276 | [1.187;1.397 | pathogenic strong |
| -0.908 | [-0.912;-0.90 | benign supporting |
| -1.009 | [-1.048;-0.97 | benign supporting |
| 1.276 | [1.104;1.667 | pathogenic strong |
| -1.066 | [-1.114;-0.70 | benign supporting |
| 1.196 | [-0.092;5.075 | pathogenic moderate |
| 0.483 | [-0.355;1.196 | pathogenic supporting |
| -0.382 | [-0.635;-0.02 | benign supporting |
| -0.720 | [-1.098;-0.35 | benign supporting |
| -1.081 | [-3.300;1.580 | benign supporting |
| -0.965 | [-0.971;-0.96 | benign supporting |
| -1.020 | [-1.167;-0.92 | benign supporting |
| 1.276 | [1.180;1.413 | pathogenic strong |
| -0.292 | [-0.328;-0.24 | indeterminate |
| 1.276 | [1.126;1.578 | pathogenic strong |
| 0.771 | [0.707;0.837 | pathogenic moderate |
| -0.265 | [-0.418;-0.00 | indeterminate |
| 1.276 | [-0.059;5.105 | pathogenic strong |
| 1.276 | [0.735;5.080 | pathogenic strong |
| 0.612 | [0.376;0.789 | pathogenic supporting |
| 1.069 | [-0.439;5.021 | pathogenic moderate |
| -0.087 | [-0.547;1.110 | indeterminate |
| -0.831 | [-1.050;-0.58 | benign supporting |
| 1.276 | [1.008;2.107 | pathogenic strong |
| -0.139 | [-0.236;-0.04 | indeterminate |
| 1.108 | [1.064;1.129 | pathogenic moderate |
| -0.915 | [-1.105;-0.55 | benign supporting |
| -0.650 | [-0.679;-0.62 | benign supporting |
| -1.001 | [-1.012;-0.98 | benign supporting |
| 1.276 | [1.135;1.527 | pathogenic strong |
| 1.224 | [1.150;1.309 | pathogenic moderate |
| 1.276 | [0.618;5.098 | pathogenic strong |
| 0.561 | [0.159;0.815 | pathogenic supporting |
| 1.196 | [0.330;4.703 | pathogenic moderate |
| 1.276 | [1.236;1.324 | pathogenic strong |
| 0.796 | [-0.953;5.034 | pathogenic moderate |
| 1.276 | [1.203;1.372 | pathogenic strong |
| 1.276 | [1.115;1.630 | pathogenic strong |
| 1.276 | [0.844;5.050 | pathogenic strong |

| | | |
|---|---|---|
| 1.276 | [1.192;1.388] | pathogenic strong |
| 1.276 | [0.789;5.064] | pathogenic strong |
| 0.605 | [0.174;0.878] | pathogenic supporting |
| 1.276 | [0.904;4.956] | pathogenic strong |
| 1.276 | [0.977;2.490] | pathogenic strong |
| 1.192 | [0.660;1.716] | pathogenic moderate |
| -0.466 | [-1.151;1.679 | benign supporting |
| -0.432 | [-1.005;0.538 | benign supporting |
| 1.276 | [1.244;1.312] | pathogenic strong |
| -1.087 | [-1.093;-1.08 | benign supporting |
| 0.552 | [-0.884;4.846 | pathogenic supporting |
| 1.192 | [1.134;1.253] | pathogenic moderate |
| 1.276 | [1.163;1.453] | pathogenic strong |
| 1.276 | [1.182;1.413] | pathogenic strong |
| -0.455 | [-0.498;-0.42 | benign supporting |
| 0.483 | [-0.180;1.045 | pathogenic supporting |
| -0.475 | [-0.495;-0.45 | benign supporting |
| -0.237 | [-0.390;0.017 | indeterminate |
| 1.172 | [1.127;1.222] | pathogenic moderate |
| -0.397 | [-0.453;-0.34 | benign supporting |
| 1.114 | [0.982;1.174] | pathogenic moderate |
| -0.401 | [-0.422;-0.38 | benign supporting |
| 1.276 | [-0.483;5.100 | pathogenic strong |
| -1.030 | [-1.070;-0.97 | benign supporting |
| -0.585 | [-0.608;-0.56 | benign supporting |
| 1.157 | [-0.622;5.086 | pathogenic moderate |
| 0.577 | [-0.381;1.275 | pathogenic supporting |
| -1.114 | [-1.116;-1.09 | benign supporting |
| -0.089 | [-0.921;1.363 | indeterminate |
| -0.929 | [-0.964;-0.89 | benign supporting |
| -1.113 | [-1.122;-1.10 | benign supporting |
| -1.022 | [-1.057;-0.98 | benign supporting |
| -0.327 | [-0.364;-0.28 | benign supporting |
| -1.090 | [-1.167;-0.74 | benign supporting |
| -0.893 | [-0.922;-0.86 | benign supporting |
| -0.991 | [-1.008;-0.96 | benign supporting |
| -1.100 | [-1.113;-1.08 | benign supporting |
| -1.049 | [-1.062;-1.03 | benign supporting |
| -0.881 | [-0.924;-0.83 | benign supporting |
| -1.090 | [-1.095;-1.08 | benign supporting |
| -0.999 | [-1.014;-0.97 | benign supporting |
| -1.021 | [-1.052;-0.99 | benign supporting |
| -0.857 | [-1.034;-0.63 | benign supporting |

| | | |
|---|---|---|
| -1.084 | [-1.090;-1.08 | benign supporting |
| 1.276 | [1.201;1.375 | pathogenic strong |
| -0.989 | [-1.015;-0.95 | benign supporting |
| -0.988 | [-1.007;-0.96 | benign supporting |
| -0.634 | [-0.655;-0.61 | benign supporting |
| -0.869 | [-0.895;-0.84 | benign supporting |
| -0.917 | [-0.973;-0.85 | benign supporting |
| -1.170 | [-1.170;-1.14 | benign supporting |
| -1.085 | [-1.090;-1.08 | benign supporting |
| -0.852 | [-0.869;-0.83 | benign supporting |
| -0.507 | [-0.567;-0.44 | benign supporting |
| -0.997 | [-1.018;-0.97 | benign supporting |
| -0.874 | [-0.904;-0.84 | benign supporting |
| -1.071 | [-1.111;-0.97 | benign supporting |
| 1.148 | [1.120;1.176 | pathogenic moderate |
| -1.070 | [-1.093;-1.01 | benign supporting |
| -1.086 | [-1.092;-1.08 | benign supporting |
| -0.876 | [-0.926;-0.82 | benign supporting |
| 1.183 | [1.139;1.230 | pathogenic moderate |
| -0.723 | [-0.789;-0.67 | benign supporting |
| -1.030 | [-1.112;-0.67 | benign supporting |
| 0.592 | [0.391;0.753 | pathogenic supporting |
| -0.938 | [-0.958;-0.91 | benign supporting |
| -0.207 | [-0.286;-0.11 | indeterminate |
| -1.052 | [-1.080;-0.99 | benign supporting |
| -0.684 | [-0.731;-0.64 | benign supporting |
| -0.567 | [-0.786;-0.40 | benign supporting |
| -0.451 | [-0.552;-0.37 | benign supporting |
| -1.090 | [-1.106;-1.08 | benign supporting |
| -0.195 | [-0.236;-0.15 | indeterminate |
| -0.493 | [-0.519;-0.46 | benign supporting |
| 1.276 | [1.153;1.485 | pathogenic strong |
| -1.166 | [-1.168;-1.16 | benign supporting |
| -1.023 | [-1.113;-0.64 | benign supporting |
| -0.661 | [-0.704;-0.62 | benign supporting |
| -0.460 | [-0.482;-0.44 | benign supporting |
| -0.687 | [-0.758;-0.63 | benign supporting |
| -1.069 | [-1.081;-1.04 | benign supporting |
| -0.869 | [-0.899;-0.84 | benign supporting |
| -0.866 | [-0.894;-0.84 | benign supporting |
| -0.056 | [-0.241;0.41 | indeterminate |
| -0.865 | [-0.928;-0.79 | benign supporting |
| 1.199 | [1.133;1.272 | pathogenic moderate |

| | | |
|---|---|---|
| -1.095 | [-1.100;-1.09 | benign supporting |
| -0.878 | [-0.913;-0.84 | benign supporting |
| -1.165 | [-1.168;-1.16 | benign supporting |
| -0.224 | [-0.396;0.20 | indeterminate |
| -1.118 | [-1.148;-1.09 | benign supporting |
| -1.103 | [-1.115;-1.08 | benign supporting |
| -1.085 | [-1.168;-0.99 | benign supporting |
| -0.259 | [-0.365;-0.10 | indeterminate |
| -0.865 | [-0.886;-0.84 | benign supporting |
| 1.162 | [1.120;1.209 | pathogenic moderate |
| -0.092 | [-0.125;-0.05 | indeterminate |
| -0.058 | [-0.113;0.01 | indeterminate |
| 0.025 | [-0.158;0.50 | indeterminate |
| 1.276 | [-0.312;5.10 | pathogenic strong |
| -0.781 | [-0.958;-0.62 | benign supporting |
| -1.103 | [-1.106;-1.09 | benign supporting |
| -0.955 | [-0.962;-0.94 | benign supporting |
| 1.276 | [1.179;1.421 | pathogenic strong |
| -0.816 | [-1.137;-0.11 | benign supporting |
| 1.118 | [1.030;1.163 | pathogenic moderate |
| 0.893 | [-1.151;4.99 | pathogenic moderate |
| 1.129 | [0.900;1.234 | pathogenic moderate |
| 0.611 | [-0.004;1.00 | pathogenic supporting |
| -1.126 | [-1.169;-0.99 | benign supporting |
| -0.929 | [-1.059;-0.75 | benign supporting |
| 1.060 | [0.899;1.141 | pathogenic moderate |
| -0.662 | [-0.943;-0.47 | benign supporting |
| 1.276 | [1.111;1.670 | pathogenic strong |
| 1.214 | [1.180;1.251 | pathogenic moderate |
| 1.276 | [1.176;1.429 | pathogenic strong |
| -0.711 | [-1.110;-0.20 | benign supporting |
| -0.350 | [-1.033;1.10 | benign supporting |
| -1.110 | [-1.114;-1.10 | benign supporting |
| 1.276 | [1.182;1.409 | pathogenic strong |
| -1.166 | [-1.169;-1.04 | benign supporting |
| -0.446 | [-0.462;-0.43 | benign supporting |
| 1.276 | [0.765;5.075 | pathogenic strong |
| -0.537 | [-0.569;-0.50 | benign supporting |
| -0.845 | [-3.784;0.00 | benign supporting |
| 0.376 | [0.234;0.496 | pathogenic supporting |
| 1.276 | [1.141;1.519 | pathogenic strong |
| 1.276 | [1.253;1.302 | pathogenic strong |
| -0.516 | [-0.531;-0.50 | benign supporting |

| | | |
|---|---|---|
| -1.105 | [-1.111;-1.10 | benign supporting |
| -0.086 | [-1.165;4.967 | indeterminate |
| -0.186 | [-0.506;0.746 | indeterminate |
| 1.276 | [1.020;2.030 | pathogenic strong |
| 1.276 | [1.173;1.430 | pathogenic strong |
| 1.276 | [-0.354;5.096 | pathogenic strong |
| 1.227 | [1.179;1.278 | pathogenic moderate |
| 1.119 | [1.016;1.171 | pathogenic moderate |
| 1.276 | [1.215;1.357 | pathogenic strong |
| -0.102 | [-0.133;-0.07 | indeterminate |
| 1.226 | [1.205;1.247 | pathogenic moderate |
| 1.276 | [1.105;1.667 | pathogenic strong |
| 0.809 | [0.691;0.935 | pathogenic moderate |
| 1.276 | [1.190;1.395 | pathogenic strong |
| 1.276 | [1.143;1.517 | pathogenic strong |
| -0.444 | [-1.025;0.513 | benign supporting |
| -0.645 | [-1.111;0.251 | benign supporting |
| 1.113 | [-0.409;5.038 | pathogenic moderate |
| 1.276 | [1.246;1.310 | pathogenic strong |
| 1.276 | [1.252;1.303 | pathogenic strong |
| -0.223 | [-0.518;0.676 | indeterminate |
| 1.276 | [1.230;1.332 | pathogenic strong |
| -0.335 | [-0.415;-0.22 | benign supporting |
| -1.093 | [-1.114;-1.07 | benign supporting |
| -1.088 | [-1.089;-1.08 | benign supporting |
| -1.087 | [-1.088;-1.08 | benign supporting |
| -1.167 | [-1.170;-1.16 | benign supporting |
| -1.084 | [-1.085;-1.08 | benign supporting |
| -1.102 | [-1.127;-1.08 | benign supporting |
| -1.069 | [-1.071;-1.06 | benign supporting |
| 0.666 | [0.372;0.878 | pathogenic moderate |
| -0.638 | [-0.652;-0.62 | benign supporting |
| -0.546 | [-0.557;-0.53 | benign supporting |
| -1.076 | [-1.078;-1.07 | benign supporting |
| -1.086 | [-1.087;-1.08 | benign supporting |
| -0.336 | [-0.542;-0.00 | benign supporting |
| 1.164 | [1.062;1.250 | pathogenic moderate |
| 1.276 | [-0.402;5.089 | pathogenic strong |
| -0.439 | [-0.585;-0.32 | benign supporting |
| -1.091 | [-1.093;-1.09 | benign supporting |
| -0.077 | [-0.169;0.077 | indeterminate |
| -0.743 | [-1.164;1.076 | benign supporting |
| 1.276 | [1.209;1.361 | pathogenic strong |

| | | |
|---|---|---|
| -1.091 | [-1.167;-1.08 | benign supporting |
| 1.163 | [1.146;1.181 | pathogenic moderate |
| 1.264 | [1.138;1.467 | pathogenic moderate |
| 1.276 | [1.186;1.402 | pathogenic strong |
| 0.391 | [-0.244;1.01 | pathogenic supporting |
| -0.837 | [-0.838;-0.83 | benign supporting |
| -1.170 | [-1.169;-1.00 | benign supporting |
| 0.419 | [-0.303;1.12 | pathogenic supporting |
| 1.276 | [1.139;1.526 | pathogenic strong |
| -1.108 | [-1.115;-1.08 | benign supporting |
| -0.909 | [-0.914;-0.90 | benign supporting |
| -0.122 | [-0.190;-0.06 | indeterminate |
| -0.469 | [-1.105;1.08 | benign supporting |
| 1.276 | [1.250;1.305 | pathogenic strong |
| 1.276 | [1.068;1.783 | pathogenic strong |
| 1.276 | [1.230;1.333 | pathogenic strong |
| 1.276 | [1.258;1.296 | pathogenic strong |
| 1.276 | [1.074;1.798 | pathogenic strong |
| -1.103 | [-1.164;-0.95 | benign supporting |
| -1.059 | [-1.070;-1.04 | benign supporting |
| -0.504 | [-0.636;-0.38 | benign supporting |
| -0.757 | [-0.884;-0.64 | benign supporting |
| -0.970 | [-0.985;-0.95 | benign supporting |
| -0.842 | [-0.885;-0.78 | benign supporting |
| -1.094 | [-1.100;-1.08 | benign supporting |
| -1.114 | [-1.161;-1.06 | benign supporting |
| -0.558 | [-0.606;-0.50 | benign supporting |
| -1.169 | [-1.170;-1.12 | benign supporting |
| -0.900 | [-0.904;-0.89 | benign supporting |
| -1.102 | [-1.112;-1.09 | benign supporting |
| 0.413 | [0.384;0.442 | pathogenic supporting |
| -1.089 | [-1.093;-1.08 | benign supporting |
| -0.634 | [-0.645;-0.62 | benign supporting |
| -0.359 | [-0.373;-0.34 | benign supporting |
| -1.142 | [-1.169;-1.08 | benign supporting |
| -0.079 | [-0.338;0.59 | indeterminate |
| -0.898 | [-1.016;-0.73 | benign supporting |
| -0.294 | [-0.311;-0.27 | indeterminate |
| -0.099 | [-0.119;-0.07 | indeterminate |
| -0.402 | [-0.446;-0.36 | benign supporting |
| -1.085 | [-1.164;-1.08 | benign supporting |
| 1.276 | [1.248;1.309 | pathogenic strong |
| -1.084 | [-1.085;-1.08 | benign supporting |

| | | |
|---|---|---|
| -1.087 | [-1.094;-1.08 | benign supporting |
| 1.115 | [1.079;1.137 | pathogenic moderate |
| 1.276 | [1.246;1.311 | pathogenic strong |
| -1.000 | [-1.011;-0.98 | benign supporting |
| -1.111 | [-1.116;-1.10 | benign supporting |
| -1.110 | [-1.169;-1.08 | benign supporting |
| -0.342 | [-0.360;-0.32 | benign supporting |
| -1.094 | [-1.119;-1.07 | benign supporting |
| -1.098 | [-1.113;-1.08 | benign supporting |
| 1.276 | [1.253;1.302 | pathogenic strong |
| -0.431 | [-0.559;-0.33 | benign supporting |
| -0.521 | [-0.616;-0.42 | benign supporting |
| 1.042 | [-0.983;5.06 | pathogenic moderate |
| -0.034 | [-0.311;0.68 | indeterminate |
| -0.396 | [-0.420;-0.37 | benign supporting |
| -0.897 | [-0.945;-0.84 | benign supporting |
| -1.109 | [-1.116;-1.09 | benign supporting |
| -1.170 | [-1.170;-1.16 | benign supporting |
| -1.109 | [-1.116;-1.09 | benign supporting |
| -1.085 | [-1.085;-1.08 | benign supporting |
| -0.959 | [-0.960;-0.95 | benign supporting |
| -1.090 | [-1.091;-1.08 | benign supporting |
| -1.161 | [-1.163;-1.15 | benign supporting |
| -0.348 | [-0.353;-0.34 | benign supporting |
| -1.166 | [-1.166;-1.16 | benign supporting |
| -0.907 | [-0.911;-0.90 | benign supporting |
| -0.337 | [-0.340;-0.33 | benign supporting |
| -1.014 | [-1.019;-1.00 | benign supporting |
| -1.094 | [-1.095;-1.09 | benign supporting |
| -1.091 | [-1.092;-1.09 | benign supporting |
| -0.331 | [-0.339;-0.32 | benign supporting |
| -0.922 | [-0.927;-0.91 | benign supporting |
| -1.115 | [-1.116;-1.10 | benign supporting |
| 1.276 | [1.253;1.300 | pathogenic strong |
| -0.784 | [-0.818;-0.75 | benign supporting |
| -0.969 | [-0.970;-0.96 | benign supporting |
| -0.580 | [-0.584;-0.57 | benign supporting |
| -0.364 | [-0.366;-0.36 | benign supporting |
| -1.167 | [-1.169;-1.16 | benign supporting |
| -1.042 | [-1.085;-1.00 | benign supporting |
| -1.094 | [-1.102;-1.08 | benign supporting |
| -0.988 | [-1.160;-0.91 | benign supporting |
| 1.276 | [1.207;1.361 | pathogenic strong |

| | | |
|---|---|---|
| -1.088 | [-1.089;-1.08 | benign supporting |
| -1.086 | [-1.087;-1.08 | benign supporting |
| -0.248 | [-0.387;-0.03 | indeterminate |
| -0.610 | [-0.697;-0.52 | benign supporting |
| 1.202 | [-0.026;5.074 | pathogenic moderate |
| 0.587 | [-0.180;1.132 | pathogenic supporting |
| -1.141 | [-1.148;-1.13 | benign supporting |
| -0.283 | [-0.304;-0.26 | indeterminate |
| -0.062 | [-0.388;0.808 | indeterminate |
| -1.095 | [-1.169;-1.06 | benign supporting |
| 1.255 | [0.658;5.080] | pathogenic moderate |
| -0.360 | [-0.383;-0.33 | benign supporting |
| -0.260 | [-0.273;-0.24 | indeterminate |
| 1.276 | [1.125;1.558] | pathogenic strong |
| -1.062 | [-3.588;0.000 | benign supporting |
| 0.986 | [0.879;1.065] | pathogenic moderate |
| 1.276 | [1.219;1.345] | pathogenic strong |
| -1.152 | [-1.170;-1.13 | benign supporting |
| 1.276 | [1.226;1.335] | pathogenic strong |
| -1.097 | [-1.099;-1.09 | benign supporting |
| -1.110 | [-1.117;-1.10 | benign supporting |
| 1.276 | [1.205;1.370] | pathogenic strong |
| -1.090 | [-1.115;-1.08 | benign supporting |
| -0.332 | [-0.662;0.551 | benign supporting |
| -1.102 | [-1.115;-1.08 | benign supporting |
| 1.276 | [0.000;5.100] | pathogenic strong |
| -1.085 | [-1.086;-1.08 | benign supporting |
| -0.977 | [-1.111;-0.58 | benign supporting |
| 0.328 | [-1.107;4.992 | pathogenic supporting |
| 0.839 | [0.680;1.007] | pathogenic moderate |
| -0.884 | [-1.164;-0.05 | benign supporting |
| -0.886 | [-0.899;-0.87 | benign supporting |
| 1.276 | [1.231;1.330] | pathogenic strong |
| -0.968 | [-1.047;-0.87 | benign supporting |
| -0.970 | [-3.626;0.000 | benign supporting |
| -0.267 | [-0.294;-0.24 | indeterminate |
| 1.276 | [-1.114;5.004 | pathogenic strong |
| 1.276 | [-0.253;5.104 | pathogenic strong |
| -1.086 | [-1.092;-1.08 | benign supporting |
| -0.627 | [-0.634;-0.62 | benign supporting |
| -1.069 | [-1.112;-0.93 | benign supporting |
| -0.615 | [-0.950;-0.39 | benign supporting |
| -1.087 | [-1.089;-1.08 | benign supporting |

| | | |
|---|---|---|
| -1.006 | [-1.016;-0.99 | benign supporting |
| -0.882 | [-0.958;-0.79 | benign supporting |
| -0.849 | [-3.754;0.000 | benign supporting |
| 1.176 | [1.138;1.216] | pathogenic moderate |
| -1.142 | [-1.160;-1.12 | benign supporting |
| 0.399 | [-0.146;0.891 | pathogenic supporting |
| 1.276 | [-1.150;4.927 | pathogenic strong |
| -0.983 | [-1.165;-0.27 | benign supporting |
| -0.253 | [-0.381;-0.06 | indeterminate |
| -0.443 | [-0.584;-0.32 | benign supporting |
| 1.276 | [-0.223;5.101 | pathogenic strong |
| -1.114 | [-1.133;-0.92 | benign supporting |
| -0.521 | [-0.561;-0.48 | benign supporting |
| -1.062 | [-1.142;-1.01 | benign supporting |
| 1.276 | [1.012;2.063] | pathogenic strong |
| -0.570 | [-1.101;0.332 | benign supporting |
| -1.078 | [-1.085;-1.07 | benign supporting |
| -0.951 | [-0.953;-0.94 | benign supporting |
| 0.576 | [0.508;0.636] | pathogenic supporting |
| -0.016 | [-0.332;0.767 | indeterminate |
| -1.013 | [-1.167;-0.91 | benign supporting |
| 1.116 | [-0.726;5.064 | pathogenic moderate |
| -1.084 | [-1.168;-0.94 | benign supporting |
| -1.084 | [-1.087;-1.08 | benign supporting |
| -1.087 | [-1.091;-1.08 | benign supporting |
| -1.162 | [-1.169;-1.09 | benign supporting |
| -0.506 | [-0.583;-0.43 | benign supporting |
| 1.276 | [1.272;1.280] | pathogenic strong |
| -0.738 | [-0.741;-0.73 | benign supporting |
| 1.158 | [1.156;1.161] | pathogenic moderate |
| -0.087 | [-0.092;-0.08 | indeterminate |
| 1.276 | [1.271;1.281] | pathogenic strong |
| -0.421 | [-0.423;-0.41 | benign supporting |
| 0.189 | [0.166;0.213] | indeterminate |
| -0.178 | [-0.183;-0.17 | indeterminate |
| 0.521 | [0.502;0.541] | pathogenic supporting |
| 1.276 | [1.247;1.309] | pathogenic strong |
| -0.493 | [-0.495;-0.49 | benign supporting |
| 0.934 | [0.929;0.939] | pathogenic moderate |
| 1.276 | [1.114;1.629] | pathogenic strong |
| -0.920 | [-0.925;-0.91 | benign supporting |
| 1.276 | [1.207;1.364] | pathogenic strong |
| 0.176 | [0.145;0.206] | indeterminate |

| | | |
|---|---|---|
| 1.276 | [1.269;1.284 | pathogenic strong |
| 1.168 | [1.165;1.170 | pathogenic moderate |
| -0.166 | [-0.249;-0.08 | indeterminate |
| -0.837 | [-0.839;-0.83 | benign supporting |
| 1.276 | [-0.098;5.103 | pathogenic strong |
| -1.087 | [-1.168;-0.60 | benign supporting |
| -1.086 | [-1.110;-1.08 | benign supporting |
| 1.276 | [-0.320;5.092 | pathogenic strong |
| -0.791 | [-1.110;-0.31 | benign supporting |
| -1.063 | [-1.164;-0.47 | benign supporting |
| 1.276 | [0.682;5.088 | pathogenic strong |
| 1.162 | [0.735;1.416 | pathogenic moderate |
| -0.883 | [-0.887;-0.87 | benign supporting |
| -0.586 | [-0.632;-0.53 | benign supporting |
| 1.224 | [0.875;1.620 | pathogenic moderate |
| 0.029 | [-0.251;0.699 | indeterminate |
| -0.969 | [-0.995;-0.95 | benign supporting |
| -0.975 | [-1.110;-0.60 | benign supporting |
| 1.214 | [-0.242;5.092 | pathogenic moderate |
| 1.276 | [0.000;5.104 | pathogenic strong |
| -0.956 | [-0.961;-0.95 | benign supporting |
| -0.057 | [-0.433;0.952 | indeterminate |
| -1.095 | [-1.115;-0.90 | benign supporting |
| -0.602 | [-0.939;-0.36 | benign supporting |
| -1.098 | [-1.112;-1.08 | benign supporting |
| -1.166 | [-1.169;-1.04 | benign supporting |
| -1.046 | [-1.112;-0.99 | benign supporting |
| -0.568 | [-0.658;-0.47 | benign supporting |
| -0.999 | [-1.167;-0.90 | benign supporting |
| 1.276 | [-0.433;5.097 | pathogenic strong |
| 1.152 | [1.131;1.176 | pathogenic moderate |
| -0.830 | [-1.059;-0.57 | benign supporting |
| 1.276 | [0.000;5.102 | pathogenic strong |
| -1.111 | [-1.115;-1.07 | benign supporting |
| 1.276 | [0.636;5.093 | pathogenic strong |
| -0.897 | [-0.964;-0.81 | benign supporting |
| -0.351 | [-0.570;-0.03 | benign supporting |
| -0.340 | [-0.488;-0.11 | benign supporting |
| -0.094 | [-0.235;0.177 | indeterminate |
| -1.109 | [-1.115;-1.05 | benign supporting |
| -1.091 | [-1.097;-1.08 | benign supporting |
| -0.767 | [-1.106;-0.34 | benign supporting |
| -1.051 | [-1.074;-1.01 | benign supporting |

| | | |
|---|---|---|
| -1.016 | [-1.062;-0.96 | benign supporting |
| -0.966 | [-0.983;-0.95 | benign supporting |
| -0.826 | [-0.934;-0.68 | benign supporting |
| -0.904 | [-1.164;-0.00 | benign supporting |
| -0.241 | [-0.312;-0.14 | indeterminate |
| -1.104 | [-1.115;-1.08 | benign supporting |
| 1.276 | [-0.282;5.10 | pathogenic strong |
| -0.127 | [-0.281;0.13 | indeterminate |
| 1.174 | [1.104;1.253 | pathogenic moderate |
| -0.170 | [-0.253;-0.08 | indeterminate |
| -1.034 | [-1.103;-0.88 | benign supporting |
| -1.049 | [-1.076;-1.00 | benign supporting |
| -1.108 | [-1.169;-1.08 | benign supporting |
| -0.935 | [-0.956;-0.91 | benign supporting |
| -1.086 | [-1.093;-1.08 | benign supporting |
| -1.087 | [-1.101;-1.08 | benign supporting |
| -0.484 | [-0.504;-0.46 | benign supporting |
| -1.071 | [-1.085;-1.04 | benign supporting |
| -0.653 | [-0.679;-0.63 | benign supporting |
| -1.119 | [-1.169;-1.08 | benign supporting |
| -0.636 | [-0.710;-0.58 | benign supporting |
| 1.103 | [0.969;1.154 | pathogenic moderate |
| -1.012 | [-1.061;-0.95 | benign supporting |
| -1.107 | [-1.115;-1.07 | benign supporting |
| 0.769 | [0.595;0.933 | pathogenic moderate |
| -1.114 | [-1.132;-1.10 | benign supporting |
| -1.009 | [-1.023;-0.99 | benign supporting |
| -1.000 | [-1.047;-0.93 | benign supporting |
| -0.518 | [-0.562;-0.47 | benign supporting |
| 0.693 | [0.040;1.063 | pathogenic moderate |
| -0.954 | [-1.111;-0.55 | benign supporting |
| -1.026 | [-1.047;-1.00 | benign supporting |
| -0.655 | [-0.680;-0.63 | benign supporting |
| -1.110 | [-1.118;-1.10 | benign supporting |
| 1.261 | [-1.116;1.55 | pathogenic moderate |
| -1.084 | [-1.086;-1.08 | benign supporting |
| -0.513 | [-0.578;-0.44 | benign supporting |
| -1.088 | [-1.093;-1.08 | benign supporting |
| -1.086 | [-1.108;-1.08 | benign supporting |
| -1.091 | [-1.096;-1.08 | benign supporting |
| -1.054 | [-1.080;-1.03 | benign supporting |
| -0.901 | [-0.914;-0.88 | benign supporting |
| -1.164 | [-1.169;-1.12 | benign supporting |

| | | |
|---|---|---|
| -1.164 | [-1.166;-1.16 | benign supporting |
| -1.126 | [-1.153;-1.10 | benign supporting |
| -1.133 | [-1.151;-1.11 | benign supporting |
| -0.573 | [-0.590;-0.55 | benign supporting |
| -1.094 | [-1.100;-1.09 | benign supporting |
| 1.276 | [0.888;4.986 | pathogenic strong |
| -0.733 | [-0.755;-0.71 | benign supporting |
| -0.174 | [-0.214;-0.13 | indeterminate |
| -0.492 | [-0.536;-0.44 | benign supporting |
| -1.152 | [-1.169;-1.13 | benign supporting |
| -1.113 | [-1.116;-1.10 | benign supporting |
| -0.503 | [-0.520;-0.48 | benign supporting |
| -1.164 | [-1.169;-1.11 | benign supporting |
| -0.411 | [-0.472;-0.35 | benign supporting |
| -1.079 | [-1.114;-0.82 | benign supporting |
| 0.064 | [-0.073;0.347 | indeterminate |
| -0.951 | [-1.040;-0.84 | benign supporting |
| -0.939 | [-0.958;-0.92 | benign supporting |
| -0.336 | [-0.465;-0.13 | benign supporting |
| -0.620 | [-1.102;-0.06 | benign supporting |
| -0.103 | [-0.884;1.315 | indeterminate |
| -1.070 | [-1.081;-1.04 | benign supporting |
| -1.112 | [-1.115;-1.06 | benign supporting |
| -1.118 | [-1.167;-1.09 | benign supporting |
| -1.035 | [-1.071;-0.99 | benign supporting |
| -1.164 | [-1.169;-1.16 | benign supporting |
| 1.276 | [-0.370;5.095 | pathogenic strong |
| -0.140 | [-0.285;0.048 | indeterminate |
| -0.388 | [-0.437;-0.33 | benign supporting |
| 0.726 | [-0.969;4.987 | pathogenic moderate |
| 1.192 | [1.148;1.240 | pathogenic moderate |
| -0.954 | [-0.962;-0.94 | benign supporting |
| -1.097 | [-1.114;-1.08 | benign supporting |
| -1.168 | [-1.169;-1.09 | benign supporting |
| -1.096 | [-1.162;-1.08 | benign supporting |
| -0.662 | [-0.848;-0.55 | benign supporting |
| -1.091 | [-1.100;-1.08 | benign supporting |
| -0.407 | [-0.510;-0.31 | benign supporting |
| -0.889 | [-0.915;-0.86 | benign supporting |
| -1.097 | [-1.107;-1.08 | benign supporting |
| -0.490 | [-0.516;-0.46 | benign supporting |
| -0.655 | [-0.688;-0.62 | benign supporting |
| -1.045 | [-1.062;-1.02 | benign supporting |

| | | |
|---|---|---|
| -0.634 | [-0.721;-0.57 | benign supporting |
| -0.616 | [-0.724;-0.52 | benign supporting |
| -0.853 | [-0.890;-0.81 | benign supporting |
| -0.374 | [-0.401;-0.34 | benign supporting |
| -0.261 | [-0.325;-0.17 | indeterminate |
| -1.001 | [-1.105;-0.75 | benign supporting |
| -0.022 | [-0.097;0.158 | indeterminate |
| -0.998 | [-1.030;-0.95 | benign supporting |
| -0.747 | [-0.794;-0.70 | benign supporting |
| -0.891 | [-0.938;-0.83 | benign supporting |
| -1.087 | [-1.092;-1.08 | benign supporting |
| -1.067 | [-1.095;-1.00 | benign supporting |
| -1.100 | [-1.115;-1.08 | benign supporting |
| -0.658 | [-1.029;-0.41 | benign supporting |
| -0.837 | [-0.897;-0.83 | benign supporting |
| -0.346 | [-1.164;4.906 | benign supporting |
| 0.180 | [-0.278;0.868 | indeterminate |
| -0.905 | [-3.701;0.000 | benign supporting |
| 1.122 | [-0.536;5.058 | pathogenic moderate |
| 1.242 | [0.258;5.089 | pathogenic moderate |
| -0.719 | [-1.115;-0.04 | benign supporting |
| 1.276 | [1.250;1.304 | pathogenic strong |
| -0.729 | [-0.956;-0.56 | benign supporting |
| -1.110 | [-1.125;-1.09 | benign supporting |
| -1.043 | [-1.070;-1.01 | benign supporting |
| -0.474 | [-0.858;-0.14 | benign supporting |
| -0.771 | [-0.812;-0.73 | benign supporting |
| 1.276 | [0.446;5.086 | pathogenic strong |
| 0.562 | [-0.266;1.155 | pathogenic supporting |
| 0.314 | [-1.072;4.928 | indeterminate |
| -0.987 | [-0.990;-0.98 | benign supporting |
| -0.898 | [-0.922;-0.87 | benign supporting |
| -0.018 | [-0.350;0.816 | indeterminate |
| 1.126 | [1.058;1.167 | pathogenic moderate |
| 1.189 | [0.984;1.360 | pathogenic moderate |
| 1.276 | [1.247;1.308 | pathogenic strong |
| -1.108 | [-1.169;-1.08 | benign supporting |
| 1.188 | [1.084;1.297 | pathogenic moderate |
| -0.646 | [-1.147;0.871 | benign supporting |
| -0.448 | [-0.629;-0.29 | benign supporting |
| -1.101 | [-1.115;-1.08 | benign supporting |
| -1.106 | [-1.115;-1.09 | benign supporting |
| -0.630 | [-1.067;-0.32 | benign supporting |

| | | |
|---|---|---|
| -0.977 | [-0.980;-0.97 | benign supporting |
| 0.167 | [-0.160;0.657 | indeterminate |
| 1.021 | [0.301;1.273 | pathogenic moderate |
| 1.276 | [1.115;1.630 | pathogenic strong |
| -0.872 | [-0.917;-0.85 | benign supporting |
| -1.085 | [-1.096;-1.08 | benign supporting |
| 1.276 | [1.243;1.313 | pathogenic strong |
| 1.276 | [1.233;1.326 | pathogenic strong |
| 1.276 | [1.233;1.324 | pathogenic strong |
| 0.608 | [0.322;0.820 | pathogenic supporting |
| 0.504 | [-0.320;1.173 | pathogenic supporting |
| -0.465 | [-0.959;0.023 | benign supporting |
| -0.986 | [-0.997;-0.97 | benign supporting |
| 0.203 | [-0.292;0.917 | indeterminate |
| -1.067 | [-1.072;-1.06 | benign supporting |
| -0.339 | [-1.156;4.949 | benign supporting |
| 1.177 | [0.178;4.070 | pathogenic moderate |
| 1.276 | [1.155;1.473 | pathogenic strong |
| -1.007 | [-1.022;-0.99 | benign supporting |
| -0.459 | [-1.158;1.954 | benign supporting |
| -1.145 | [-1.169;-1.11 | benign supporting |
| 1.201 | [-0.519;5.084 | pathogenic moderate |
| -0.421 | [-0.761;-0.03 | benign supporting |
| -0.269 | [-0.494;0.344 | indeterminate |
| 1.201 | [1.105;1.318 | pathogenic moderate |
| -0.933 | [-1.111;-0.50 | benign supporting |
| 1.276 | [-0.219;5.100 | pathogenic strong |
| 1.276 | [-0.086;5.105 | pathogenic strong |
| 1.276 | [1.065;1.797 | pathogenic strong |
| 1.087 | [0.780;1.203 | pathogenic moderate |
| 1.183 | [1.058;1.297 | pathogenic moderate |
| -0.247 | [-0.922;1.173 | indeterminate |
| -0.422 | [-0.943;0.422 | benign supporting |
| 1.276 | [1.240;1.318 | pathogenic strong |
| 1.276 | [1.240;1.318 | pathogenic strong |
| 0.530 | [-0.525;1.422 | pathogenic supporting |
| -1.101 | [-1.165;-1.08 | benign supporting |
| -0.673 | [-1.097;-0.30 | benign supporting |
| 1.276 | [0.000;5.102 | pathogenic strong |
| -1.100 | [-1.103;-1.09 | benign supporting |
| 1.263 | [1.006;1.830 | pathogenic moderate |
| -0.849 | [-0.866;-0.83 | benign supporting |
| 1.239 | [1.177;1.311 | pathogenic moderate |

| | | |
|---|---|---|
| -0.971 | [-0.975;-0.96 | benign supporting |
| 1.276 | [1.246;1.309] | pathogenic strong |
| -0.982 | [-1.166;0.758 | benign supporting |
| -0.937 | [-0.951;-0.92 | benign supporting |
| 0.403 | [0.345;0.459] | pathogenic supporting |
| -0.873 | [-0.886;-0.85 | benign supporting |
| -1.088 | [-1.168;-0.52 | benign supporting |
| 1.101 | [0.921;1.163] | pathogenic moderate |
| 0.565 | [0.514;0.608] | pathogenic supporting |
| -0.479 | [-1.076;0.433 | benign supporting |
| 1.276 | [1.201;1.377] | pathogenic strong |
| -1.107 | [-1.119;-1.09 | benign supporting |
| -1.010 | [-1.016;-1.00 | benign supporting |
| 1.276 | [-0.100;5.102 | pathogenic strong |
| 1.276 | [1.227;1.336] | pathogenic strong |
| -0.967 | [-3.647;0.000 | benign supporting |
| 1.276 | [1.225;1.340] | pathogenic strong |
| -0.698 | [-0.796;-0.63 | benign supporting |
| 1.276 | [1.238;1.319] | pathogenic strong |
| 1.276 | [1.224;1.338] | pathogenic strong |
| 1.276 | [1.170;1.436] | pathogenic strong |
| 1.276 | [1.170;1.436] | pathogenic strong |
| 1.250 | [-0.195;5.100 | pathogenic moderate |
| 1.276 | [1.187;1.401] | pathogenic strong |
| 1.276 | [0.993;2.208] | pathogenic strong |
| 1.276 | [1.191;1.393] | pathogenic strong |
| 1.276 | [1.212;1.357] | pathogenic strong |
| 1.276 | [1.106;1.647] | pathogenic strong |
| 1.182 | [0.856;1.404] | pathogenic moderate |
| 1.276 | [1.252;1.302] | pathogenic strong |
| 1.276 | [1.163;1.448] | pathogenic strong |
| 1.276 | [1.107;1.640] | pathogenic strong |
| -0.632 | [-0.652;-0.61 | benign supporting |
| 1.119 | [0.658;1.301] | pathogenic moderate |
| 1.276 | [0.362;5.104] | pathogenic strong |
| 1.276 | [1.214;1.355] | pathogenic strong |
| -1.050 | [-3.580;0.000 | benign supporting |
| -0.977 | [-0.981;-0.97 | benign supporting |
| -1.003 | [-1.148;-0.93 | benign supporting |
| -1.100 | [-1.109;-1.09 | benign supporting |
| 1.238 | [0.987;1.591] | pathogenic moderate |
| -1.164 | [-1.169;-0.92 | benign supporting |
| 1.265 | [1.238;1.295] | pathogenic moderate |

| | | |
|---|---|---|
| -0.292 | [-0.334;-0.24 | indeterminate |
| -0.096 | [-0.275;0.322 | indeterminate |
| -0.438 | [-0.580;-0.32 | benign supporting |
| 0.336 | [-0.283;1.042 | pathogenic supporting |
| 1.276 | [1.188;1.402 | pathogenic strong |
| 1.191 | [-0.365;5.087 | pathogenic moderate |
| -1.084 | [-1.086;-1.08 | benign supporting |
| -0.976 | [-3.501;0.000 | benign supporting |
| -0.923 | [-3.534;-0.83 | benign supporting |
| -0.986 | [-1.009;-0.95 | benign supporting |
| -0.424 | [-0.490;-0.37 | benign supporting |
| -0.611 | [-0.620;-0.60 | benign supporting |
| -1.117 | [-1.158;-1.09 | benign supporting |
| -0.462 | [-0.471;-0.45 | benign supporting |
| -1.151 | [-1.170;-1.12 | benign supporting |
| -1.100 | [-1.108;-1.09 | benign supporting |
| -1.084 | [-1.086;-1.08 | benign supporting |
| -0.674 | [-0.691;-0.65 | benign supporting |
| -0.233 | [-0.301;-0.14 | indeterminate |
| -1.038 | [-1.054;-1.02 | benign supporting |
| -0.449 | [-0.461;-0.43 | benign supporting |
| 1.276 | [1.233;1.326 | pathogenic strong |
| -1.087 | [-1.099;-1.07 | benign supporting |
| 0.851 | [0.773;0.935 | pathogenic moderate |
| 1.168 | [1.045;1.269 | pathogenic moderate |
| -1.145 | [-1.158;-1.13 | benign supporting |
| -0.387 | [-0.398;-0.37 | benign supporting |
| -0.381 | [-0.393;-0.36 | benign supporting |
| -1.127 | [-1.134;-1.12 | benign supporting |
| -1.091 | [-1.110;-1.07 | benign supporting |
| -1.094 | [-1.114;-1.07 | benign supporting |
| -0.892 | [-0.895;-0.88 | benign supporting |
| -0.366 | [-0.374;-0.35 | benign supporting |
| 1.099 | [1.008;1.136 | pathogenic moderate |
| -1.100 | [-1.103;-1.09 | benign supporting |
| -0.330 | [-0.487;-0.08 | benign supporting |
| -1.095 | [-1.103;-1.08 | benign supporting |
| -1.069 | [-1.076;-1.05 | benign supporting |
| -1.114 | [-1.168;-1.08 | benign supporting |
| -0.974 | [-1.007;-0.95 | benign supporting |
| 0.980 | [0.917;1.028 | pathogenic moderate |
| -0.904 | [-0.908;-0.90 | benign supporting |
| -0.366 | [-0.558;-0.10 | benign supporting |

| | | |
|---|---|---|
| -0.955 | [-1.147;-0.36 | benign supporting |
| -0.264 | [-0.444;0.153 | indeterminate |
| 1.229 | [1.205;1.255] | pathogenic moderate |
| -1.062 | [-1.067;-1.05 | benign supporting |
| 0.372 | [0.188;0.524] | pathogenic supporting |
| -1.140 | [-1.159;-1.12 | benign supporting |
| -1.085 | [-1.168;-0.92 | benign supporting |
| -1.094 | [-1.115;-1.08 | benign supporting |
| 1.276 | [1.153;1.486] | pathogenic strong |
| -0.137 | [-0.380;0.487 | indeterminate |
| -1.049 | [-1.055;-1.04 | benign supporting |
| -1.100 | [-1.115;-0.96 | benign supporting |
| -0.887 | [-0.889;-0.88 | benign supporting |
| 1.276 | [1.225;1.338] | pathogenic strong |
| -0.802 | [-1.030;-0.57 | benign supporting |
| -0.624 | [-0.637;-0.61 | benign supporting |
| -0.956 | [-0.966;-0.94 | benign supporting |
| 1.276 | [1.158;1.459] | pathogenic strong |
| 1.184 | [1.113;1.264] | pathogenic moderate |
| 0.656 | [-0.567;1.768 | pathogenic moderate |
| -0.183 | [-0.202;-0.16 | indeterminate |
| 1.276 | [-0.492;5.094 | pathogenic strong |
| 1.157 | [1.137;1.179] | pathogenic moderate |
| 1.276 | [1.219;1.344] | pathogenic strong |
| -0.648 | [-2.208;4.863 | benign supporting |
| -0.368 | [-0.441;-0.28 | benign supporting |
| -1.089 | [-1.165;-1.08 | benign supporting |
| -1.109 | [-1.115;-1.06 | benign supporting |
| 0.930 | [0.820;1.030] | pathogenic moderate |
| -1.064 | [-1.068;-1.05 | benign supporting |
| -1.086 | [-1.092;-1.08 | benign supporting |
| 1.276 | [-0.429;5.097 | pathogenic strong |
| -1.074 | [-1.089;-1.05 | benign supporting |
| 0.594 | [-0.078;1.039 | pathogenic supporting |
| -1.108 | [-1.115;-1.09 | benign supporting |
| 0.630 | [0.016;1.001] | pathogenic supporting |
| 0.099 | [-0.103;0.466 | indeterminate |
| 1.276 | [1.208;1.361] | pathogenic strong |
| -1.043 | [-1.088;-0.95 | benign supporting |
| -0.412 | [-0.621;-0.17 | benign supporting |
| -1.068 | [-1.115;-0.63 | benign supporting |
| 1.276 | [1.089;1.703] | pathogenic strong |
| -1.020 | [-1.089;-0.89 | benign supporting |

| | | |
|---|---|---|
| 0.761 | [0.635;0.872 | pathogenic moderate |
| -0.869 | [-0.955;-0.76 | benign supporting |
| -1.090 | [-1.114;-0.96 | benign supporting |
| -0.006 | [-1.166;4.731 | indeterminate |
| -0.683 | [-0.706;-0.66 | benign supporting |
| -0.984 | [-1.005;-0.96 | benign supporting |
| -0.331 | [-0.376;-0.27 | benign supporting |
| -0.994 | [-3.398;1.818 | benign supporting |
| -0.398 | [-0.448;-0.34 | benign supporting |
| -0.657 | [-0.735;-0.60 | benign supporting |
| -0.281 | [-0.312;-0.24 | indeterminate |
| -1.017 | [-1.032;-1.00 | benign supporting |
| -0.356 | [-0.395;-0.31 | benign supporting |
| -0.315 | [-0.778;0.826 | indeterminate |
| -0.819 | [-0.868;-0.74 | benign supporting |
| -0.554 | [-0.580;-0.52 | benign supporting |
| -0.311 | [-0.333;-0.28 | indeterminate |
| -0.462 | [-0.505;-0.42 | benign supporting |
| -1.067 | [-1.088;-1.01 | benign supporting |
| 0.731 | [0.659;0.798 | pathogenic moderate |
| -1.164 | [-1.165;-1.16 | benign supporting |
| 0.919 | [0.844;0.998 | pathogenic moderate |
| -0.909 | [-0.928;-0.88 | benign supporting |
| -0.672 | [-0.764;-0.61 | benign supporting |
| -0.519 | [-0.574;-0.46 | benign supporting |
| -1.006 | [-1.012;-1.00 | benign supporting |
| -0.677 | [-0.691;-0.66 | benign supporting |
| -0.216 | [-0.246;-0.18 | indeterminate |
| -1.133 | [-1.168;-1.10 | benign supporting |
| -1.044 | [-1.051;-1.03 | benign supporting |
| -0.791 | [-0.809;-0.77 | benign supporting |
| -0.549 | [-0.556;-0.54 | benign supporting |
| -0.422 | [-0.470;-0.38 | benign supporting |
| -0.632 | [-0.646;-0.61 | benign supporting |
| -0.317 | [-0.344;-0.28 | indeterminate |
| -0.696 | [-0.716;-0.68 | benign supporting |
| -0.901 | [-0.914;-0.88 | benign supporting |
| 1.276 | [1.162;1.454 | pathogenic strong |
| -0.510 | [-0.525;-0.49 | benign supporting |
| -0.745 | [-0.831;-0.67 | benign supporting |
| 1.254 | [0.014;5.098 | pathogenic moderate |
| -1.053 | [-1.091;-0.96 | benign supporting |
| -0.442 | [-0.451;-0.43 | benign supporting |

| | | |
|---|---|---|
| 1.276 | [1.250;1.305 | pathogenic strong |
| -0.445 | [-0.450;-0.44 | benign supporting |
| -1.048 | [-1.077;-1.00 | benign supporting |
| -1.080 | [-1.166;-0.30 | benign supporting |
| -0.509 | [-0.582;-0.43 | benign supporting |
| -0.872 | [-0.946;-0.77 | benign supporting |
| -0.907 | [-1.020;-0.74 | benign supporting |
| -0.300 | [-0.355;-0.22 | indeterminate |
| -0.264 | [-0.373;-0.10 | indeterminate |
| 1.103 | [0.520;1.304 | pathogenic moderate |
| 1.192 | [1.140;1.249 | pathogenic moderate |
| 1.276 | [1.223;1.342 | pathogenic strong |
| -0.473 | [-0.578;-0.38 | benign supporting |
| -0.252 | [-0.316;-0.17 | indeterminate |
| 1.276 | [1.216;1.354 | pathogenic strong |
| 0.000 | [0.000;0.000 | indeterminate |
| 1.276 | [1.187;1.403 | pathogenic strong |
| -0.913 | [-1.065;-0.68 | benign supporting |
| 0.424 | [-0.246;1.05( | pathogenic supporting |
| -0.470 | [-1.106;1.10{ | benign supporting |
| -0.150 | [-0.219;-0.08 | indeterminate |
| 1.047 | [0.932;1.119 | pathogenic moderate |
| -0.161 | [-0.483;0.77{ | indeterminate |
| -0.921 | [-0.946;-0.89 | benign supporting |
| -1.014 | [-1.070;-0.93 | benign supporting |
| -1.093 | [-1.103;-1.08 | benign supporting |
| -0.183 | [-0.239;-0.12 | indeterminate |
| -1.108 | [-1.115;-1.09 | benign supporting |
| -1.071 | [-1.082;-1.05 | benign supporting |
| -0.491 | [-0.523;-0.45 | benign supporting |
| -1.087 | [-1.092;-1.08 | benign supporting |
| -1.097 | [-1.103;-1.09 | benign supporting |
| -0.878 | [-1.009;-0.70 | benign supporting |
| -1.134 | [-1.169;-1.10 | benign supporting |
| -0.832 | [-0.901;-0.73 | benign supporting |
| -0.619 | [-0.655;-0.58 | benign supporting |
| 1.276 | [1.131;1.567 | pathogenic strong |
| -1.029 | [-1.091;-0.91 | benign supporting |
| -0.307 | [-0.331;-0.27 | indeterminate |
| -0.876 | [-0.964;-0.76 | benign supporting |
| 0.916 | [0.778;1.047 | pathogenic moderate |
| -0.837 | [-0.839;-0.83 | benign supporting |
| -0.691 | [-0.806;-0.62 | benign supporting |

| | | |
|---|---|---|
| -0.823 | [-0.838;-0.79 | benign supporting |
| -0.935 | [-0.960;-0.91 | benign supporting |
| -1.086 | [-1.090;-1.08 | benign supporting |
| -1.102 | [-1.105;-1.10 | benign supporting |
| -1.090 | [-1.114;-1.06 | benign supporting |
| -0.577 | [-0.587;-0.56 | benign supporting |
| -1.127 | [-1.169;-1.08 | benign supporting |
| -0.599 | [-0.635;-0.55 | benign supporting |
| -0.614 | [-0.624;-0.60 | benign supporting |
| -1.089 | [-1.096;-1.08 | benign supporting |
| -0.923 | [-0.947;-0.89 | benign supporting |
| -0.694 | [-0.728;-0.66 | benign supporting |
| -0.612 | [-0.626;-0.59 | benign supporting |
| -1.107 | [-1.111;-1.10 | benign supporting |
| 1.276 | [1.202;1.374] | pathogenic strong |
| -1.091 | [-1.099;-1.08 | benign supporting |
| 0.190 | [-0.161;0.68 | indeterminate |
| -0.927 | [-0.953;-0.89 | benign supporting |
| -0.520 | [-0.533;-0.50 | benign supporting |
| -1.094 | [-1.096;-1.09 | benign supporting |
| -1.065 | [-1.089;-1.00 | benign supporting |
| -1.099 | [-1.105;-1.09 | benign supporting |
| -0.736 | [-0.875;-0.63 | benign supporting |
| -0.736 | [-0.885;-0.62 | benign supporting |
| -0.415 | [-0.475;-0.36 | benign supporting |
| -1.091 | [-1.114;-0.92 | benign supporting |
| -1.102 | [-1.114;-1.09 | benign supporting |
| -1.123 | [-3.450;1.091 | benign supporting |
| -0.881 | [-0.965;-0.77 | benign supporting |
| -0.607 | [-0.834;-0.43 | benign supporting |
| 1.276 | [-0.520;5.09 | pathogenic strong |
| -0.690 | [-1.009;-0.46 | benign supporting |
| -0.426 | [-0.471;-0.38 | benign supporting |
| -1.085 | [-1.095;-1.08 | benign supporting |
| -1.085 | [-1.090;-1.08 | benign supporting |
| 1.232 | [-0.341;5.09 | pathogenic moderate |
| -0.911 | [-0.988;-0.82 | benign supporting |
| 1.276 | [1.216;1.348] | pathogenic strong |
| 1.276 | [0.000;5.108] | pathogenic strong |
| -0.955 | [-0.963;-0.94 | benign supporting |
| 1.276 | [1.117;1.595] | pathogenic strong |
| -0.669 | [-0.724;-0.62 | benign supporting |
| -0.863 | [-0.933;-0.77 | benign supporting |

| | | |
|---|---|---|
| -0.025 | [-0.191;0.418 | indeterminate |
| 1.254 | [1.152;1.401] | pathogenic moderate |
| 1.276 | [1.245;1.312] | pathogenic strong |
| 1.260 | [1.227;1.294] | pathogenic moderate |
| 1.276 | [1.253;1.301] | pathogenic strong |
| 1.276 | [1.164;1.456] | pathogenic strong |
| 1.188 | [1.168;1.208] | pathogenic moderate |
| 1.276 | [1.216;1.349] | pathogenic strong |
| 1.276 | [1.136;1.549] | pathogenic strong |
| 1.276 | [1.230;1.333] | pathogenic strong |
| 0.937 | [0.374;1.191] | pathogenic moderate |
| 1.276 | [1.235;1.324] | pathogenic strong |
| 1.157 | [1.122;1.196] | pathogenic moderate |
| 1.276 | [1.187;1.398] | pathogenic strong |
| -0.400 | [-0.415;-0.38 | benign supporting |
| 1.118 | [0.997;1.178] | pathogenic moderate |
| 1.224 | [1.187;1.262] | pathogenic moderate |
| 1.276 | [1.251;1.303] | pathogenic strong |
| 1.276 | [1.211;1.359] | pathogenic strong |
| 0.951 | [0.894;1.005] | pathogenic moderate |
| -1.075 | [-1.113;-0.96 | benign supporting |
| -1.097 | [-1.114;-1.08 | benign supporting |
| -0.913 | [-0.932;-0.88 | benign supporting |
| -0.725 | [-0.770;-0.68 | benign supporting |
| -0.833 | [-0.861;-0.79 | benign supporting |
| -0.341 | [-0.367;-0.31 | benign supporting |
| -1.094 | [-1.110;-1.08 | benign supporting |
| -0.668 | [-0.722;-0.62 | benign supporting |
| -1.084 | [-1.088;-1.08 | benign supporting |
| 0.757 | [0.704;0.808] | pathogenic moderate |
| -0.525 | [-0.548;-0.50 | benign supporting |
| -0.579 | [-0.621;-0.53 | benign supporting |
| -0.875 | [-0.903;-0.84 | benign supporting |
| -0.158 | [-0.258;-0.05 | indeterminate |
| -1.018 | [-1.053;-0.98 | benign supporting |
| 1.276 | [1.246;1.309] | pathogenic strong |
| -0.589 | [-0.829;-0.42 | benign supporting |
| -1.081 | [-1.085;-1.07 | benign supporting |
| -0.960 | [-0.995;-0.93 | benign supporting |
| -1.081 | [-1.089;-1.07 | benign supporting |
| -1.112 | [-1.116;-1.10 | benign supporting |
| -1.037 | [-1.107;-0.87 | benign supporting |
| -0.399 | [-0.481;-0.32 | benign supporting |

| | | |
|---|---|---|
| 1.276 | [1.159;1.453 | pathogenic strong |
| 1.131 | [0.826;1.265 | pathogenic moderate |
| -0.535 | [-0.589;-0.47 | benign supporting |
| 1.276 | [1.106;1.680 | pathogenic strong |
| 0.631 | [0.509;0.733 | pathogenic supporting |
| 1.276 | [1.157;1.460 | pathogenic strong |
| 1.276 | [1.202;1.373 | pathogenic strong |
| -0.112 | [-0.238;0.07 | indeterminate |
| -0.599 | [-0.638;-0.55 | benign supporting |
| -1.081 | [-1.091;-1.07 | benign supporting |
| 1.276 | [1.223;1.340 | pathogenic strong |
| 1.276 | [0.852;5.048 | pathogenic strong |
| -0.137 | [-0.182;-0.09 | indeterminate |
| -0.122 | [-0.462;0.82 | indeterminate |
| -0.759 | [-0.849;-0.67 | benign supporting |
| 1.140 | [1.126;1.155 | pathogenic moderate |
| 0.062 | [-0.042;0.27 | indeterminate |
| -1.060 | [-1.092;-0.98 | benign supporting |
| 1.276 | [1.176;1.426 | pathogenic strong |
| 1.276 | [1.214;1.355 | pathogenic strong |
| 1.090 | [0.997;1.134 | pathogenic moderate |
| 1.276 | [1.250;1.304 | pathogenic strong |
| 1.276 | [1.242;1.314 | pathogenic strong |
| 1.276 | [1.205;1.365 | pathogenic strong |
| 1.276 | [1.181;1.412 | pathogenic strong |
| 1.242 | [1.215;1.270 | pathogenic moderate |
| 1.276 | [1.163;1.451 | pathogenic strong |
| 1.130 | [1.105;1.155 | pathogenic moderate |
| 1.276 | [1.134;1.546 | pathogenic strong |
| 1.276 | [1.211;1.361 | pathogenic strong |
| 1.276 | [1.234;1.326 | pathogenic strong |
| 1.276 | [1.194;1.389 | pathogenic strong |
| 1.276 | [1.231;1.327 | pathogenic strong |
| 0.985 | [0.600;1.174 | pathogenic moderate |
| 1.276 | [1.140;1.528 | pathogenic strong |
| 1.276 | [1.253;1.300 | pathogenic strong |
| 1.276 | [1.253;1.302 | pathogenic strong |
| 1.276 | [1.135;1.543 | pathogenic strong |
| -1.085 | [-1.098;-1.08 | benign supporting |
| -0.062 | [-0.173;0.19 | indeterminate |
| 1.276 | [0.743;5.081 | pathogenic strong |
| 1.276 | [1.120;1.581 | pathogenic strong |
| 1.121 | [0.843;1.235 | pathogenic moderate |

| | | |
|---|---|---|
| -1.031 | [-1.166;-0.11 | benign supporting |
| 1.276 | [0.026;5.102 | pathogenic strong |
| 0.507 | [-0.505;1.379 | pathogenic supporting |
| 1.276 | [0.634;5.085 | pathogenic strong |
| 1.276 | [-1.123;5.037 | pathogenic strong |
| 0.349 | [-0.653;1.485 | pathogenic supporting |
| 1.276 | [0.000;5.107 | pathogenic strong |
| -1.091 | [-2.939;1.352 | benign supporting |
| 1.276 | [-0.217;5.102 | pathogenic strong |
| 1.051 | [0.893;1.135 | pathogenic moderate |
| 1.276 | [1.177;1.425 | pathogenic strong |
| -0.343 | [-0.598;0.177 | benign supporting |
| -0.358 | [-0.367;-0.34 | benign supporting |
| 0.799 | [0.544;1.027 | pathogenic moderate |
| -0.801 | [-1.048;-0.56 | benign supporting |
| 0.962 | [0.849;1.056 | pathogenic moderate |
| 1.240 | [1.102;1.453 | pathogenic moderate |
| -1.142 | [-1.169;-1.10 | benign supporting |
| 1.276 | [1.153;1.484 | pathogenic strong |
| 0.635 | [0.604;0.666 | pathogenic supporting |
| 1.276 | [1.258;1.296 | pathogenic strong |
| 1.276 | [0.837;5.061 | pathogenic strong |
| 0.966 | [0.444;1.201 | pathogenic moderate |
| 1.178 | [1.096;1.269 | pathogenic moderate |
| 1.276 | [1.195;1.384 | pathogenic strong |
| -0.369 | [-0.421;-0.30 | benign supporting |
| 1.065 | [-0.196;1.652 | pathogenic moderate |
| -1.113 | [-1.165;-0.83 | benign supporting |
| 1.276 | [1.238;1.321 | pathogenic strong |
| 1.143 | [-0.260;5.039 | pathogenic moderate |
| 0.756 | [0.726;0.788 | pathogenic moderate |
| -0.339 | [-0.421;-0.22 | benign supporting |
| -0.221 | [-0.305;-0.11 | indeterminate |
| -0.985 | [-1.029;-0.95 | benign supporting |
| -1.093 | [-1.167;-0.54 | benign supporting |
| 0.995 | [-0.341;2.112 | pathogenic moderate |
| -1.084 | [-1.168;-0.99 | benign supporting |
| 0.553 | [-1.150;5.018 | pathogenic supporting |
| 1.276 | [0.000;5.106 | pathogenic strong |
| 1.276 | [1.222;1.342 | pathogenic strong |
| 1.276 | [1.139;1.530 | pathogenic strong |
| 1.276 | [1.193;1.387 | pathogenic strong |
| 1.276 | [1.192;1.391 | pathogenic strong |

| | | |
|---|---|---|
| 1.276 | [1.042;1.881 | pathogenic strong |
| -0.965 | [-0.988;-0.94 | benign supporting |
| 1.276 | [-0.175;5.10 | pathogenic strong |
| 1.276 | [0.905;5.033 | pathogenic strong |
| -1.155 | [-1.161;-1.14 | benign supporting |
| -1.121 | [-1.130;-1.11 | benign supporting |
| -1.036 | [-1.053;-1.02 | benign supporting |
| 0.931 | [-0.209;1.43 | pathogenic moderate |
| 1.276 | [1.094;1.702 | pathogenic strong |
| 1.276 | [0.411;5.096 | pathogenic strong |
| -1.165 | [-1.168;-1.16 | benign supporting |
| 1.276 | [-0.241;5.10 | pathogenic strong |
| 1.276 | [0.649;5.098 | pathogenic strong |
| 1.276 | [-0.245;5.10 | pathogenic strong |
| 1.276 | [1.077;1.727 | pathogenic strong |
| 1.276 | [1.139;1.529 | pathogenic strong |
| 1.276 | [1.165;1.446 | pathogenic strong |
| 1.219 | [1.058;1.400 | pathogenic moderate |
| 0.550 | [0.452;0.628 | pathogenic supporting |
| 1.276 | [0.922;5.025 | pathogenic strong |
| -1.087 | [-1.164;-0.63 | benign supporting |
| 1.276 | [0.983;2.618 | pathogenic strong |
| 1.276 | [1.207;1.367 | pathogenic strong |
| 1.276 | [1.231;1.329 | pathogenic strong |
| 0.127 | [-1.029;2.22 | indeterminate |
| 1.276 | [1.054;1.851 | pathogenic strong |
| -0.688 | [-1.091;-0.35 | benign supporting |
| -0.014 | [-0.285;0.67 | indeterminate |
| 1.276 | [1.229;1.333 | pathogenic strong |
| 1.276 | [1.181;1.408 | pathogenic strong |
| 1.276 | [1.232;1.327 | pathogenic strong |
| 1.276 | [0.487;5.092 | pathogenic strong |
| 1.276 | [1.198;1.381 | pathogenic strong |
| 1.115 | [0.818;1.229 | pathogenic moderate |
| 1.276 | [1.227;1.336 | pathogenic strong |
| 1.276 | [1.104;1.688 | pathogenic strong |
| 1.079 | [0.762;1.200 | pathogenic moderate |
| 1.276 | [1.110;1.660 | pathogenic strong |
| 1.276 | [-0.198;5.10 | pathogenic strong |
| 1.276 | [0.000;5.109 | pathogenic strong |
| 1.276 | [0.791;5.079 | pathogenic strong |
| 1.276 | [1.223;1.342 | pathogenic strong |
| 0.889 | [0.031;1.220 | pathogenic moderate |

| | | |
|---|---|---|
| 1.276 | [1.060;1.832 | pathogenic strong |
| 1.149 | [1.135;1.163 | pathogenic moderate |
| 1.276 | [1.246;1.309 | pathogenic strong |
| 0.690 | [0.468;0.852 | pathogenic moderate |
| -0.345 | [-0.497;-0.11 | benign supporting |
| -0.389 | [-0.609;-0.11 | benign supporting |
| 1.276 | [1.246;1.310 | pathogenic strong |
| 1.276 | [1.057;1.837 | pathogenic strong |
| 1.276 | [1.205;1.369 | pathogenic strong |
| -0.948 | [-1.123;-0.38 | benign supporting |
| 1.276 | [1.231;1.330 | pathogenic strong |
| -0.071 | [-0.129;0.00 | indeterminate |
| -1.085 | [-1.087;-1.08 | benign supporting |
| 0.748 | [0.593;0.882 | pathogenic moderate |
| 1.276 | [1.099;1.713 | pathogenic strong |
| 1.276 | [1.223;1.342 | pathogenic strong |
| 1.177 | [0.921;1.345 | pathogenic moderate |
| 0.858 | [0.507;1.120 | pathogenic moderate |
| -0.416 | [-0.621;-0.21 | benign supporting |
| 1.276 | [0.862;5.051 | pathogenic strong |
| -0.305 | [-0.332;-0.27 | indeterminate |
| 1.276 | [1.238;1.320 | pathogenic strong |
| 0.705 | [0.491;0.870 | pathogenic moderate |
| 1.197 | [1.090;1.316 | pathogenic moderate |
| -1.047 | [-1.055;-1.04 | benign supporting |
| -1.121 | [-1.169;-0.44 | benign supporting |
| -1.090 | [-1.167;-0.58 | benign supporting |
| 1.276 | [-0.303;5.10 | pathogenic strong |
| -1.057 | [-1.072;-1.04 | benign supporting |
| -0.948 | [-0.950;-0.94 | benign supporting |
| 1.276 | [1.150;1.476 | pathogenic strong |
| 0.760 | [0.257;1.106 | pathogenic moderate |
| 1.276 | [-0.125;5.10 | pathogenic strong |
| 1.276 | [1.248;1.307 | pathogenic strong |
| -1.093 | [-1.105;-1.08 | benign supporting |
| 1.276 | [1.241;1.316 | pathogenic strong |
| -0.784 | [-1.153;0.44 | benign supporting |
| 1.276 | [1.191;1.394 | pathogenic strong |
| 1.276 | [1.106;1.669 | pathogenic strong |
| 1.107 | [0.648;1.280 | pathogenic moderate |
| -1.104 | [-1.104;-1.10 | benign supporting |
| -0.788 | [-0.802;-0.77 | benign supporting |
| 1.276 | [1.268;1.285 | pathogenic strong |

| | | |
|---|---|---|
| 1.276 | [1.265;1.288] | pathogenic strong |
| 1.276 | [1.197;1.380] | pathogenic strong |
| -0.636 | [-0.637;-0.63 | benign supporting |
| 1.276 | [1.269;1.283] | pathogenic strong |
| 1.276 | [1.273;1.279] | pathogenic strong |
| 1.276 | [1.271;1.282] | pathogenic strong |
| 1.026 | [1.017;1.035] | pathogenic moderate |
| -0.726 | [-0.739;-0.71 | benign supporting |
| 1.276 | [1.270;1.283] | pathogenic strong |
| -0.309 | [-0.313;-0.30 | indeterminate |
| 1.276 | [1.271;1.281] | pathogenic strong |
| 1.276 | [1.220;1.345] | pathogenic strong |
| -0.264 | [-0.296;-0.23 | indeterminate |
| 1.276 | [1.272;1.281] | pathogenic strong |
| 1.166 | [1.158;1.174] | pathogenic moderate |
| 1.276 | [1.275;1.278] | pathogenic strong |
| 1.276 | [1.273;1.280] | pathogenic strong |
| -0.617 | [-0.623;-0.61 | benign supporting |
| -1.077 | [-1.114;-0.90 | benign supporting |
| -0.968 | [-1.065;-0.85 | benign supporting |
| -0.236 | [-0.423;0.241 | indeterminate |
| 1.276 | [0.496;5.096] | pathogenic strong |
| -1.048 | [-1.065;-1.03 | benign supporting |
| -0.819 | [-0.870;-0.74 | benign supporting |
| 1.276 | [1.203;1.371] | pathogenic strong |
| -0.967 | [-1.004;-0.93 | benign supporting |
| 1.276 | [1.193;1.391] | pathogenic strong |
| -1.076 | [-1.114;-0.90 | benign supporting |
| -1.088 | [-1.100;-1.07 | benign supporting |
| -1.056 | [-1.069;-1.03 | benign supporting |
| 1.276 | [0.435;5.102] | pathogenic strong |
| -0.999 | [-1.091;-0.82 | benign supporting |
| -0.401 | [-1.107;1.247 | benign supporting |
| -1.084 | [-1.166;-1.08 | benign supporting |
| 1.276 | [0.915;4.968] | pathogenic strong |
| 0.209 | [0.026;0.356] | indeterminate |
| 0.006 | [-0.379;0.958 | indeterminate |
| -0.191 | [-0.350;0.063 | indeterminate |
| 0.826 | [0.426;1.114] | pathogenic moderate |
| 1.276 | [0.751;5.081] | pathogenic strong |
| -0.939 | [-0.942;-0.93 | benign supporting |
| -1.056 | [-1.067;-1.04 | benign supporting |
| 1.276 | [1.002;2.079] | pathogenic strong |

| | | |
|---|---|---|
| 0.645 | [-0.566;1.64( | pathogenic moderate |
| -1.141 | [-3.443;0.00( | benign supporting |
| -1.164 | [-1.169;-1.10 | benign supporting |
| -0.839 | [-3.762;0.00( | benign supporting |
| 0.564 | [-0.367;1.23( | pathogenic supporting |
| 1.276 | [1.245;1.312 | pathogenic strong |
| 0.901 | [0.496;1.143 | pathogenic moderate |
| -3.347 | [-3.805;0.00( | benign strong |
| 0.538 | [0.367;0.677 | pathogenic supporting |
| 1.276 | [1.127;1.569 | pathogenic strong |
| -0.641 | [-0.841;-0.51 | benign supporting |
| -0.556 | [-0.803;-0.36 | benign supporting |
| 1.276 | [1.059;1.850 | pathogenic strong |
| 1.276 | [1.182;1.409 | pathogenic strong |
| 1.276 | [0.000;5.105 | pathogenic strong |
| -0.828 | [-0.865;-0.77 | benign supporting |
| 1.276 | [1.198;1.382 | pathogenic strong |
| -1.109 | [-1.115;-1.06 | benign supporting |
| -0.418 | [-0.612;-0.22 | benign supporting |
| -1.164 | [-1.166;-1.16 | benign supporting |
| 1.188 | [0.801;1.467 | pathogenic moderate |
| 1.276 | [1.258;1.294 | pathogenic strong |
| 1.085 | [0.935;1.146 | pathogenic moderate |
| 1.276 | [1.006;2.065 | pathogenic strong |
| 1.276 | [1.249;1.306 | pathogenic strong |
| 1.276 | [0.892;5.002 | pathogenic strong |
| 1.276 | [0.600;5.094 | pathogenic strong |
| 1.172 | [1.036;1.284 | pathogenic moderate |
| -0.649 | [-0.692;-0.61 | benign supporting |
| 1.276 | [1.253;1.302 | pathogenic strong |
| 1.276 | [1.254;1.300 | pathogenic strong |
| 1.276 | [1.179;1.416 | pathogenic strong |
| 0.927 | [0.604;1.138 | pathogenic moderate |
| 1.154 | [0.747;1.366 | pathogenic moderate |
| 1.276 | [1.238;1.318 | pathogenic strong |
| 1.276 | [1.256;1.299 | pathogenic strong |
| 1.276 | [1.244;1.314 | pathogenic strong |
| -0.352 | [-0.518;-0.11 | benign supporting |
| -1.108 | [-1.168;-1.04 | benign supporting |
| -1.079 | [-1.136;-0.65 | benign supporting |
| -0.598 | [-1.085;-0.17 | benign supporting |
| 1.276 | [1.221;1.345 | pathogenic strong |
| 1.276 | [1.129;1.551 | pathogenic strong |

| | | |
|---|---|---|
| -0.852 | [-1.101;-0.49 | benign supporting |
| -0.853 | [-0.925;-0.84 | benign supporting |
| 0.672 | [0.454;0.833] | pathogenic moderate |
| 1.276 | [0.487;5.096] | pathogenic strong |
| -0.510 | [-0.840;-0.25 | benign supporting |
| 1.276 | [1.211;1.358] | pathogenic strong |
| 1.276 | [1.232;1.327] | pathogenic strong |
| 1.276 | [1.195;1.390] | pathogenic strong |
| 0.000 | [0.000;0.000] | indeterminate |
| 0.392 | [-0.208;0.977 | pathogenic supporting |
| -0.902 | [-1.078;-0.63 | benign supporting |
| 1.276 | [1.125;1.591] | pathogenic strong |
| -0.845 | [-1.074;-0.57 | benign supporting |
| -1.090 | [-1.094;-1.08 | benign supporting |
| 0.181 | [-0.273;0.865 | indeterminate |
| -0.338 | [-0.568;0.083 | benign supporting |
| 1.235 | [1.194;1.278] | pathogenic moderate |
| -0.271 | [-0.314;-0.22 | indeterminate |
| 0.071 | [-0.048;0.302 | indeterminate |
| 1.276 | [1.188;1.396] | pathogenic strong |
| 1.276 | [1.236;1.323] | pathogenic strong |
| 0.628 | [-0.260;1.200 | pathogenic supporting |
| 0.672 | [0.353;0.902] | pathogenic moderate |
| 0.048 | [-0.295;0.804 | indeterminate |
| 1.276 | [1.052;1.854] | pathogenic strong |
| -1.096 | [-1.104;-1.09 | benign supporting |
| 0.834 | [0.371;1.131] | pathogenic moderate |
| 1.276 | [1.239;1.317] | pathogenic strong |
| -0.429 | [-0.615;-0.25 | benign supporting |
| 1.178 | [0.998;1.315] | pathogenic moderate |
| 1.276 | [1.210;1.356] | pathogenic strong |
| 1.276 | [1.015;2.163] | pathogenic strong |
| -0.115 | [-0.360;0.530 | indeterminate |
| -1.017 | [-1.054;-0.98 | benign supporting |
| -0.814 | [-0.985;-0.62 | benign supporting |
| -1.065 | [-1.074;-1.03 | benign supporting |
| -0.420 | [-0.454;-0.38 | benign supporting |
| -1.087 | [-1.091;-1.08 | benign supporting |
| -1.085 | [-1.112;-1.08 | benign supporting |
| -0.936 | [-0.947;-0.92 | benign supporting |
| -1.016 | [-1.023;-1.00 | benign supporting |
| -1.166 | [-1.169;-1.15 | benign supporting |
| -1.039 | [-1.070;-1.00 | benign supporting |

| | | |
|---|---|---|
| 1.276 | [1.149;1.492 | pathogenic strong |
| 1.145 | [1.129;1.161 | pathogenic moderate |
| 1.276 | [-0.034;5.104 | pathogenic strong |
| 0.318 | [-0.082;0.688 | pathogenic supporting |
| -0.766 | [-1.061;-0.51 | benign supporting |
| 1.136 | [1.116;1.155 | pathogenic moderate |
| -0.564 | [-0.581;-0.54 | benign supporting |
| -1.048 | [-1.052;-1.04 | benign supporting |
| 1.276 | [0.842;5.057 | pathogenic strong |
| -1.095 | [-1.101;-1.09 | benign supporting |
| -1.097 | [-1.114;-1.08 | benign supporting |
| 1.276 | [0.979;2.124 | pathogenic strong |
| 1.167 | [1.012;1.281 | pathogenic moderate |
| 0.439 | [-0.414;1.232 | pathogenic supporting |
| -1.166 | [-1.170;-1.12 | benign supporting |
| 1.276 | [1.226;1.337 | pathogenic strong |
| 1.276 | [1.117;1.596 | pathogenic strong |
| 1.276 | [1.189;1.397 | pathogenic strong |
| -0.676 | [-0.916;-0.53 | benign supporting |
| 1.253 | [1.191;1.326 | pathogenic moderate |
| 1.276 | [0.108;5.104 | pathogenic strong |
| -1.057 | [-1.060;-1.05 | benign supporting |
| 1.276 | [1.043;1.895 | pathogenic strong |
| -0.964 | [-1.013;-0.91 | benign supporting |
| 0.974 | [0.470;1.197 | pathogenic moderate |
| 1.276 | [-0.049;5.106 | pathogenic strong |
| -0.376 | [-0.523;-0.18 | benign supporting |
| -1.084 | [-1.088;-1.08 | benign supporting |
| -0.359 | [-0.417;-0.28 | benign supporting |
| -0.257 | [-0.421;0.055 | indeterminate |
| 0.377 | [-1.153;5.015 | pathogenic supporting |
| -1.047 | [-1.056;-1.03 | benign supporting |
| -0.558 | [-0.707;-0.41 | benign supporting |
| -1.087 | [-1.167;-0.95 | benign supporting |
| 1.276 | [0.981;2.247 | pathogenic strong |
| -1.086 | [-1.088;-1.08 | benign supporting |
| 0.718 | [-0.296;1.281 | pathogenic moderate |
| 1.276 | [1.036;1.929 | pathogenic strong |
| -1.057 | [-1.112;-0.85 | benign supporting |
| -0.999 | [-1.078;-0.86 | benign supporting |
| 1.201 | [-0.219;5.094 | pathogenic moderate |
| -0.861 | [-0.875;-0.84 | benign supporting |
| -1.088 | [-1.114;-1.03 | benign supporting |

| | | |
|---|---|---|
| -0.832 | [-0.844;-0.81 | benign supporting |
| -1.105 | [-1.115;-0.98 | benign supporting |
| 1.242 | [-0.069;5.103 | pathogenic moderate |
| 1.276 | [0.591;5.090 | pathogenic strong |
| -0.200 | [-0.403;0.352 | indeterminate |
| -0.420 | [-0.996;0.571 | benign supporting |
| 0.378 | [-0.841;2.226 | pathogenic supporting |
| 1.224 | [0.419;5.056 | pathogenic moderate |
| 1.276 | [-0.030;5.104 | pathogenic strong |
| 1.276 | [1.192;1.395 | pathogenic strong |
| 1.247 | [1.112;1.463 | pathogenic moderate |
| 0.895 | [0.563;1.131 | pathogenic moderate |
| -0.868 | [-1.053;-0.63 | benign supporting |
| 1.276 | [-0.353;5.105 | pathogenic strong |
| 1.276 | [1.106;1.634 | pathogenic strong |
| -0.216 | [-0.384;0.139 | indeterminate |
| 1.127 | [1.083;1.160 | pathogenic moderate |
| 1.187 | [1.157;1.217 | pathogenic moderate |
| 1.276 | [1.047;1.899 | pathogenic strong |
| 1.276 | [0.984;2.346 | pathogenic strong |
| 1.224 | [-1.084;5.065 | pathogenic moderate |
| -0.013 | [-0.285;0.679 | indeterminate |
| -0.997 | [-1.055;-0.92 | benign supporting |
| -0.433 | [-1.092;0.992 | benign supporting |
| 1.201 | [0.435;4.963 | pathogenic moderate |
| -1.046 | [-1.054;-1.03 | benign supporting |
| 1.276 | [1.106;1.671 | pathogenic strong |
| -0.972 | [-1.142;-0.89 | benign supporting |
| 0.577 | [0.311;0.770 | pathogenic supporting |
| -0.632 | [-1.108;-0.02 | benign supporting |
| 1.276 | [1.217;1.350 | pathogenic strong |
| -0.349 | [-0.361;-0.33 | benign supporting |
| 1.276 | [0.781;5.081 | pathogenic strong |
| 1.276 | [1.065;1.839 | pathogenic strong |
| -0.963 | [-1.019;-0.90 | benign supporting |
| -0.578 | [-0.612;-0.54 | benign supporting |
| 1.276 | [1.129;1.550 | pathogenic strong |
| -0.456 | [-0.541;-0.39 | benign supporting |
| -1.112 | [-1.116;-1.09 | benign supporting |
| -1.102 | [-1.115;-1.08 | benign supporting |
| -0.982 | [-1.046;-0.90 | benign supporting |
| -0.977 | [-0.999;-0.96 | benign supporting |
| -1.099 | [-1.103;-1.09 | benign supporting |

| | | |
|---|---|---|
| -1.062 | [-1.074;-1.03 | benign supporting |
| -1.167 | [-1.170;-1.15 | benign supporting |
| -1.106 | [-1.110;-1.10 | benign supporting |
| 1.008 | [0.179;1.282 | pathogenic moderate |
| -0.626 | [-1.071;-0.30 | benign supporting |
| -1.101 | [-1.115;-1.08 | benign supporting |
| -1.101 | [-1.120;-1.09 | benign supporting |
| -0.321 | [-0.340;-0.29 | benign supporting |
| -0.454 | [-0.630;-0.30 | benign supporting |
| -0.037 | [-0.172;0.318 | indeterminate |
| -1.070 | [-1.097;-1.01 | benign supporting |
| 1.276 | [1.232;1.329 | pathogenic strong |
| -1.091 | [-1.114;-1.06 | benign supporting |
| -0.906 | [-0.924;-0.88 | benign supporting |
| -0.660 | [-0.728;-0.61 | benign supporting |
| -0.956 | [-1.021;-0.88 | benign supporting |
| -1.116 | [-1.169;-0.97 | benign supporting |
| 0.847 | [-0.286;1.398 | pathogenic moderate |
| 0.747 | [0.111;1.117 | pathogenic moderate |
| -0.522 | [-0.642;-0.40 | benign supporting |
| 1.143 | [0.457;1.499 | pathogenic moderate |
| -0.457 | [-0.627;-0.30 | benign supporting |
| 1.276 | [-1.101;5.061 | pathogenic strong |
| 1.219 | [-1.084;5.074 | pathogenic moderate |
| 1.276 | [0.000;5.100 | pathogenic strong |
| 1.276 | [0.693;5.089 | pathogenic strong |
| 1.276 | [1.118;1.585 | pathogenic strong |
| 1.276 | [0.777;5.069 | pathogenic strong |
| 1.276 | [-1.141;4.993 | pathogenic strong |
| 1.276 | [1.011;2.012 | pathogenic strong |
| 1.276 | [1.047;1.884 | pathogenic strong |
| 1.276 | [-1.125;5.007 | pathogenic strong |
| -0.390 | [-0.485;-0.28 | benign supporting |
| -0.375 | [-0.407;-0.33 | benign supporting |
| 1.276 | [-0.957;5.079 | pathogenic strong |
| -1.009 | [-1.032;-0.98 | benign supporting |
| -1.116 | [-1.115;-1.07 | benign supporting |
| -0.914 | [-2.831;4.754 | benign supporting |
| -0.484 | [-0.642;-0.34 | benign supporting |
| -0.328 | [-0.370;-0.27 | benign supporting |
| -0.122 | [-0.253;0.034 | indeterminate |
| 1.042 | [0.871;1.135 | pathogenic moderate |
| -1.096 | [-1.102;-1.09 | benign supporting |

| | | |
|---|---|---|
| -0.640 | [-0.671;-0.61 | benign supporting |
| -1.086 | [-1.129;-1.08 | benign supporting |
| -0.440 | [-0.497;-0.39 | benign supporting |
| 1.157 | [1.061;1.234 | pathogenic moderate |
| -0.740 | [-0.826;-0.67 | benign supporting |
| -1.162 | [-1.170;-1.14 | benign supporting |
| -1.084 | [-1.167;-1.08 | benign supporting |
| -1.094 | [-1.108;-1.08 | benign supporting |
| -0.631 | [-0.813;-0.51 | benign supporting |
| -0.651 | [-0.690;-0.62 | benign supporting |
| -0.069 | [-0.128;0.00 | indeterminate |
| -0.943 | [-0.955;-0.93 | benign supporting |
| -0.702 | [-0.752;-0.66 | benign supporting |
| 0.374 | [-0.347;1.12 | pathogenic supporting |
| -1.097 | [-1.104;-1.09 | benign supporting |
| -1.104 | [-1.109;-1.10 | benign supporting |
| -0.821 | [-0.838;-0.79 | benign supporting |
| -1.103 | [-1.107;-1.09 | benign supporting |
| -0.644 | [-0.851;-0.52 | benign supporting |
| -1.093 | [-1.099;-1.09 | benign supporting |
| -0.702 | [-0.729;-0.68 | benign supporting |
| -0.108 | [-0.139;-0.07 | indeterminate |
| -1.084 | [-1.088;-1.08 | benign supporting |
| -1.109 | [-1.114;-1.10 | benign supporting |
| -1.090 | [-1.102;-1.07 | benign supporting |
| 1.117 | [1.101;1.132 | pathogenic moderate |
| -1.106 | [-1.114;-1.09 | benign supporting |
| 1.211 | [1.176;1.246 | pathogenic moderate |
| -1.166 | [-1.170;-1.15 | benign supporting |
| -1.104 | [-1.108;-1.10 | benign supporting |
| -1.084 | [-1.086;-1.08 | benign supporting |
| -0.628 | [-0.664;-0.59 | benign supporting |
| -0.673 | [-0.769;-0.61 | benign supporting |
| -0.881 | [-0.940;-0.82 | benign supporting |
| -0.553 | [-0.616;-0.47 | benign supporting |
| -0.981 | [-1.102;-0.71 | benign supporting |
| -0.778 | [-1.053;-0.53 | benign supporting |
| -1.089 | [-1.110;-1.08 | benign supporting |
| -1.098 | [-1.109;-1.08 | benign supporting |
| -1.085 | [-1.168;-0.61 | benign supporting |
| -1.020 | [-1.053;-0.98 | benign supporting |
| -0.995 | [-1.037;-0.96 | benign supporting |
| -0.584 | [-0.723;-0.45 | benign supporting |

| | | |
|---|---|---|
| -1.104 | [-1.115;-1.08 | benign supporting |
| -0.480 | [-0.503;-0.45 | benign supporting |
| -0.470 | [-0.502;-0.44 | benign supporting |
| -1.064 | [-1.105;-0.95 | benign supporting |
| -1.101 | [-1.115;-1.09 | benign supporting |
| 0.997 | [0.818;1.114 | pathogenic moderate |
| -0.597 | [-0.729;-0.47 | benign supporting |
| 0.115 | [-0.052;0.362 | indeterminate |
| -0.109 | [-0.297;0.327 | indeterminate |
| -0.391 | [-0.429;-0.35 | benign supporting |
| -0.005 | [-0.182;0.459 | indeterminate |
| -1.098 | [-1.122;-1.08 | benign supporting |
| -1.111 | [-1.116;-1.10 | benign supporting |
| -1.097 | [-1.109;-1.08 | benign supporting |
| -1.085 | [-1.088;-1.08 | benign supporting |
| -1.115 | [-1.116;-1.10 | benign supporting |
| -1.042 | [-1.082;-0.96 | benign supporting |
| -0.914 | [-0.936;-0.88 | benign supporting |
| -0.946 | [-0.966;-0.92 | benign supporting |
| -1.128 | [-1.144;-1.11 | benign supporting |
| -1.166 | [-1.169;-1.15 | benign supporting |
| -0.937 | [-0.975;-0.89 | benign supporting |
| -0.921 | [-1.041;-0.75 | benign supporting |
| 1.276 | [1.238;1.323 | pathogenic strong |
| -1.087 | [-1.105;-1.08 | benign supporting |
| -0.866 | [-0.909;-0.82 | benign supporting |
| 0.635 | [0.507;0.740 | pathogenic supporting |
| -0.973 | [-0.979;-0.96 | benign supporting |
| -0.872 | [-1.055;-0.63 | benign supporting |
| -1.056 | [-1.098;-0.95 | benign supporting |
| -1.121 | [-1.126;-1.11 | benign supporting |
| -0.902 | [-0.948;-0.84 | benign supporting |
| -0.088 | [-0.142;-0.02 | indeterminate |
| -0.835 | [-0.852;-0.81 | benign supporting |
| -0.771 | [-0.879;-0.66 | benign supporting |
| -1.109 | [-1.115;-1.08 | benign supporting |
| -0.738 | [-0.845;-0.65 | benign supporting |
| -1.090 | [-1.098;-1.08 | benign supporting |
| -1.084 | [-1.085;-1.08 | benign supporting |
| -1.037 | [-1.047;-1.02 | benign supporting |
| -1.050 | [-1.068;-1.03 | benign supporting |
| -1.084 | [-1.134;-1.08 | benign supporting |
| 0.472 | [-0.709;2.102 | pathogenic supporting |

| | | |
|---|---|---|
| -0.161 | [-0.245;-0.07 | indeterminate |
| -0.879 | [-0.918;-0.84 | benign supporting |
| -1.088 | [-1.090;-1.08 | benign supporting |
| -1.130 | [-1.142;-1.11 | benign supporting |
| 0.840 | [-0.018;1.205 | pathogenic moderate |
| -1.092 | [-1.169;-0.99 | benign supporting |
| -0.496 | [-0.917;-0.16 | benign supporting |
| 0.059 | [-0.002;0.165 | indeterminate |
| 0.324 | [-0.483;1.259 | pathogenic supporting |
| -1.058 | [-1.074;-1.02 | benign supporting |
| -1.112 | [-1.116;-1.09 | benign supporting |
| -1.140 | [-1.148;-1.13 | benign supporting |
| -0.966 | [-1.016;-0.91 | benign supporting |
| -0.367 | [-0.397;-0.33 | benign supporting |
| -0.979 | [-1.018;-0.93 | benign supporting |
| -1.087 | [-1.088;-1.08 | benign supporting |
| -0.502 | [-0.550;-0.45 | benign supporting |
| -1.065 | [-1.106;-0.95 | benign supporting |
| -1.088 | [-1.108;-1.08 | benign supporting |
| 1.224 | [1.136;1.327] | pathogenic moderate |
| -1.098 | [-1.105;-1.09 | benign supporting |
| -1.126 | [-1.136;-1.11 | benign supporting |
| -1.169 | [-1.169;-1.08 | benign supporting |
| -0.881 | [-0.925;-0.83 | benign supporting |
| -1.088 | [-1.093;-1.08 | benign supporting |
| -1.115 | [-1.116;-1.10 | benign supporting |
| 1.276 | [1.185;1.403] | pathogenic strong |
| -0.354 | [-0.423;-0.26 | benign supporting |
| -1.092 | [-1.164;-1.08 | benign supporting |
| -0.848 | [-0.877;-0.82 | benign supporting |
| -0.446 | [-0.493;-0.40 | benign supporting |
| 0.676 | [0.603;0.737] | pathogenic moderate |
| 1.276 | [1.187;1.398] | pathogenic strong |
| -0.898 | [-0.973;-0.81 | benign supporting |
| 0.025 | [-0.080;0.308 | indeterminate |
| -0.031 | [-0.065;0.011 | indeterminate |
| 0.336 | [0.007;0.585] | pathogenic supporting |
| 1.276 | [1.182;1.411] | pathogenic strong |
| -0.362 | [-1.088;1.183 | benign supporting |
| -1.081 | [-1.114;-0.91 | benign supporting |
| 1.276 | [1.235;1.325] | pathogenic strong |
| -1.105 | [-1.169;-1.01 | benign supporting |
| 1.276 | [-0.229;5.103 | pathogenic strong |

| | | |
|---|---|---|
| -0.088 | [-0.336;0.565 | indeterminate |
| 1.161 | [1.059;1.248 | pathogenic moderate |
| -1.065 | [-1.072;-1.04 | benign supporting |
| -0.929 | [-0.940;-0.91 | benign supporting |
| -1.052 | [-1.073;-1.01 | benign supporting |
| 0.564 | [0.411;0.688 | pathogenic supporting |
| 1.173 | [0.889;1.362 | pathogenic moderate |
| -1.084 | [-1.092;-1.07 | benign supporting |
| 0.250 | [0.019;0.427 | indeterminate |
| -0.948 | [-0.951;-0.94 | benign supporting |
| -1.062 | [-3.229;-0.33 | benign supporting |
| 1.009 | [-0.423;4.921 | pathogenic moderate |
| -0.736 | [-0.990;-0.55 | benign supporting |
| -0.048 | [-0.077;-0.01 | indeterminate |
| -1.138 | [-1.162;-1.11 | benign supporting |
| 0.554 | [0.051;0.851 | pathogenic supporting |
| -1.072 | [-1.166;-0.33 | benign supporting |
| 1.276 | [1.177;1.423 | pathogenic strong |
| -1.092 | [-1.134;-1.08 | benign supporting |
| -1.070 | [-1.166;-1.01 | benign supporting |
| 0.010 | [-0.380;0.965 | indeterminate |
| -0.680 | [-0.738;-0.63 | benign supporting |
| -1.088 | [-1.166;-1.08 | benign supporting |
| -0.948 | [-0.954;-0.94 | benign supporting |
| -1.105 | [-1.169;-0.87 | benign supporting |
| -0.773 | [-1.164;0.706 | benign supporting |
| -0.431 | [-0.494;-0.38 | benign supporting |
| -1.169 | [-1.169;-1.15 | benign supporting |
| -1.082 | [-1.107;-1.06 | benign supporting |
| 1.276 | [1.230;1.331 | pathogenic strong |
| 1.157 | [1.086;1.225 | pathogenic moderate |
| -0.627 | [-1.099;-0.11 | benign supporting |
| -0.612 | [-0.900;-0.41 | benign supporting |
| -1.010 | [-1.028;-0.99 | benign supporting |
| -1.089 | [-1.093;-1.08 | benign supporting |
| 1.008 | [0.633;1.182 | pathogenic moderate |
| 1.010 | [0.768;1.140 | pathogenic moderate |
| 1.276 | [1.221;1.342 | pathogenic strong |
| -1.100 | [-1.121;-1.08 | benign supporting |
| -1.167 | [-1.169;-0.50 | benign supporting |
| 1.276 | [0.599;5.095 | pathogenic strong |
| 0.540 | [0.024;0.837 | pathogenic supporting |
| -1.105 | [-1.111;-1.09 | benign supporting |

| | | |
|---|---|---|
| -0.920 | [-0.922;-0.91 | benign supporting |
| -1.089 | [-1.092;-1.08 | benign supporting |
| -0.973 | [-1.071;-0.84 | benign supporting |
| -1.069 | [-1.089;-1.02 | benign supporting |
| -1.104 | [-1.111;-1.09 | benign supporting |
| -0.949 | [-1.094;-0.67 | benign supporting |
| -1.083 | [-1.114;-0.98 | benign supporting |
| -1.163 | [-1.170;-1.15 | benign supporting |
| 1.276 | [1.163;1.458 | pathogenic strong |
| -1.165 | [-1.168;-1.16 | benign supporting |
| -0.903 | [-1.077;-0.64 | benign supporting |
| -0.484 | [-0.845;-0.19 | benign supporting |
| 1.276 | [1.225;1.336 | pathogenic strong |
| -1.163 | [-1.169;-1.15 | benign supporting |
| -1.098 | [-1.168;-0.40 | benign supporting |
| -1.164 | [-3.453;1.357 | benign supporting |
| 1.276 | [1.217;1.351 | pathogenic strong |
| 0.352 | [-0.174;0.863 | pathogenic supporting |
| -1.092 | [-1.094;-1.09 | benign supporting |
| -1.129 | [-1.134;-1.12 | benign supporting |
| -1.138 | [-1.148;-1.12 | benign supporting |
| -1.092 | [-1.116;-1.08 | benign supporting |
| -1.012 | [-1.022;-1.00 | benign supporting |
| -1.090 | [-1.099;-1.08 | benign supporting |
| -1.105 | [-1.108;-1.10 | benign supporting |
| -0.938 | [-1.134;-0.36 | benign supporting |
| 1.222 | [1.183;1.262 | pathogenic moderate |
| -0.397 | [-0.404;-0.39 | benign supporting |
| 1.276 | [1.220;1.346 | pathogenic strong |
| 1.276 | [1.233;1.327 | pathogenic strong |
| -0.216 | [-0.434;0.412 | indeterminate |
| 1.271 | [1.119;1.581 | pathogenic moderate |
| 1.276 | [0.711;5.096 | pathogenic strong |
| -0.639 | [-0.653;-0.62 | benign supporting |
| -1.155 | [-1.166;-1.14 | benign supporting |
| -0.843 | [-1.096;-0.50 | benign supporting |
| 0.000 | [0.000;0.000 | indeterminate |
| -1.116 | [-1.116;-1.10 | benign supporting |
| -1.091 | [-1.092;-1.09 | benign supporting |
| -1.101 | [-1.104;-1.09 | benign supporting |
| -1.109 | [-1.114;-1.10 | benign supporting |
| -1.085 | [-1.087;-1.08 | benign supporting |
| -1.084 | [-1.087;-1.08 | benign supporting |

| | | |
|---|---|---|
| -1.096 | [-1.099;-1.09 | benign supporting |
| -0.885 | [-0.898;-0.87 | benign supporting |
| -0.614 | [-0.627;-0.59 | benign supporting |
| -1.129 | [-1.146;-1.11 | benign supporting |
| -1.074 | [-1.088;-1.05 | benign supporting |
| -1.112 | [-1.116;-1.10 | benign supporting |
| -1.066 | [-1.073;-1.04 | benign supporting |
| -0.085 | [-0.107;-0.06 | indeterminate |
| -1.102 | [-1.114;-1.08 | benign supporting |
| 1.219 | [1.176;1.264 | pathogenic moderate |
| -1.101 | [-1.115;-1.08 | benign supporting |
| -1.088 | [-1.090;-1.08 | benign supporting |
| -1.134 | [-1.143;-1.12 | benign supporting |
| -0.963 | [-0.972;-0.95 | benign supporting |
| -1.066 | [-1.070;-1.05 | benign supporting |
| -0.627 | [-0.662;-0.59 | benign supporting |
| 1.276 | [1.225;1.336 | pathogenic strong |
| -1.085 | [-1.087;-1.08 | benign supporting |
| 1.068 | [0.522;1.263 | pathogenic moderate |
| -0.966 | [-1.094;-0.71 | benign supporting |
| 1.154 | [1.108;1.203 | pathogenic moderate |
| 0.820 | [0.726;0.921 | pathogenic moderate |
| 1.276 | [1.103;1.702 | pathogenic strong |
| -0.633 | [-0.745;-0.55 | benign supporting |
| 1.276 | [1.155;1.470 | pathogenic strong |
| -0.176 | [-0.213;-0.13 | indeterminate |
| -1.091 | [-1.115;-1.05 | benign supporting |
| 1.276 | [1.233;1.327 | pathogenic strong |
| 0.505 | [0.302;0.673 | pathogenic supporting |
| 0.953 | [0.888;1.013 | pathogenic moderate |
| -0.367 | [-1.102;1.28 | benign supporting |
| 1.276 | [1.239;1.318 | pathogenic strong |
| 1.239 | [1.105;1.440 | pathogenic moderate |
| -0.695 | [-0.843;-0.61 | benign supporting |
| -1.105 | [-1.115;-1.09 | benign supporting |
| 1.276 | [1.141;1.512 | pathogenic strong |
| 1.276 | [1.229;1.333 | pathogenic strong |
| 0.822 | [0.658;0.994 | pathogenic moderate |
| 0.969 | [0.644;1.151 | pathogenic moderate |
| -0.673 | [-0.717;-0.63 | benign supporting |
| 1.276 | [1.193;1.391 | pathogenic strong |
| 1.112 | [0.957;1.177 | pathogenic moderate |
| 1.276 | [1.070;1.776 | pathogenic strong |

| | | |
|---|---|---|
| 1.023 | [0.904;1.106 | pathogenic moderate |
| -0.788 | [-0.853;-0.71 | benign supporting |
| 0.933 | [0.745;1.103 | pathogenic moderate |
| 0.360 | [-0.538;1.328 | pathogenic supporting |
| 1.276 | [1.156;1.469 | pathogenic strong |
| 1.276 | [1.211;1.359 | pathogenic strong |
| 1.276 | [1.237;1.323 | pathogenic strong |
| 0.347 | [0.076;0.546 | pathogenic supporting |
| 1.276 | [1.227;1.332 | pathogenic strong |
| 1.148 | [1.023;1.234 | pathogenic moderate |
| -0.919 | [-1.093;-0.62 | benign supporting |
| -1.092 | [-1.108;-1.08 | benign supporting |
| -0.750 | [-1.092;-0.41 | benign supporting |
| 1.276 | [0.663;5.089 | pathogenic strong |
| 1.276 | [1.142;1.519 | pathogenic strong |
| 0.921 | [0.619;1.130 | pathogenic moderate |
| -1.002 | [-1.071;-0.90 | benign supporting |
| 0.733 | [-0.247;1.252 | pathogenic moderate |
| 1.276 | [-1.124;5.007 | pathogenic strong |
| 1.175 | [0.360;2.076 | pathogenic moderate |
| 1.276 | [-1.093;5.048 | pathogenic strong |
| -1.084 | [-1.095;-1.08 | benign supporting |
| 1.276 | [1.216;1.350 | pathogenic strong |
| 1.276 | [1.138;1.522 | pathogenic strong |
| 1.276 | [-0.308;5.104 | pathogenic strong |
| 1.276 | [-1.115;5.006 | pathogenic strong |
| 1.172 | [0.974;1.309 | pathogenic moderate |
| 1.276 | [0.821;5.077 | pathogenic strong |
| -0.284 | [-0.620;0.732 | indeterminate |
| 0.272 | [-1.084;4.850 | indeterminate |
| 1.276 | [1.170;1.435 | pathogenic strong |
| 1.276 | [0.000;5.103 | pathogenic strong |
| -1.160 | [-1.169;4.550 | benign supporting |
| -0.583 | [-0.961;-0.31 | benign supporting |
| 0.820 | [0.612;1.023 | pathogenic moderate |
| -0.519 | [-0.560;-0.47 | benign supporting |
| 1.276 | [1.235;1.323 | pathogenic strong |
| -1.109 | [-1.115;-1.10 | benign supporting |
| -0.066 | [-0.196;0.250 | indeterminate |
| 1.276 | [1.210;1.358 | pathogenic strong |
| 0.824 | [0.758;0.895 | pathogenic moderate |
| -0.653 | [-0.678;-0.63 | benign supporting |
| -0.991 | [-1.045;-0.92 | benign supporting |

| | | |
|---|---|---|
| 1.097 | [1.029;1.126 | pathogenic moderate |
| -1.084 | [-1.087;-1.08 | benign supporting |
| -1.096 | [-1.103;-1.09 | benign supporting |
| 1.276 | [1.227;1.334 | pathogenic strong |
| -0.362 | [-0.400;-0.32 | benign supporting |
| 1.205 | [1.165;1.246 | pathogenic moderate |
| -0.163 | [-0.350;0.252 | indeterminate |
| -1.097 | [-1.114;-1.08 | benign supporting |
| -0.413 | [-0.455;-0.37 | benign supporting |
| -0.305 | [-0.554;0.310 | indeterminate |
| -1.063 | [-1.066;-1.05 | benign supporting |
| -1.019 | [-1.066;-0.96 | benign supporting |
| 1.276 | [1.223;1.341 | pathogenic strong |
| -0.172 | [-0.180;-0.16 | indeterminate |
| 0.868 | [0.005;1.214 | pathogenic moderate |
| -0.688 | [-0.712;-0.66 | benign supporting |
| -1.088 | [-1.091;-1.08 | benign supporting |
| -1.081 | [-1.084;-1.07 | benign supporting |
| -0.011 | [-0.137;0.340 | indeterminate |
| -0.429 | [-0.465;-0.39 | benign supporting |
| -0.612 | [-0.647;-0.57 | benign supporting |
| 1.160 | [1.126;1.197 | pathogenic moderate |
| -0.460 | [-0.465;-0.45 | benign supporting |
| 1.276 | [1.242;1.316 | pathogenic strong |
| -0.126 | [-0.197;-0.06 | indeterminate |
| 1.138 | [1.046;1.198 | pathogenic moderate |
| -1.098 | [-1.107;-1.09 | benign supporting |
| 1.276 | [1.254;1.300 | pathogenic strong |
| 1.174 | [1.144;1.208 | pathogenic moderate |
| -1.025 | [-1.051;-1.00 | benign supporting |
| -0.972 | [-0.989;-0.96 | benign supporting |
| -0.530 | [-0.552;-0.50 | benign supporting |
| 1.276 | [1.200;1.371 | pathogenic strong |
| 0.315 | [-0.030;0.599 | indeterminate |
| 1.276 | [1.142;1.509 | pathogenic strong |
| -0.252 | [-0.302;-0.18 | indeterminate |
| 1.066 | [1.006;1.112 | pathogenic moderate |
| 1.258 | [1.242;1.275 | pathogenic moderate |
| -0.310 | [-0.324;-0.29 | indeterminate |
| 1.276 | [1.222;1.344 | pathogenic strong |
| -0.673 | [-0.720;-0.63 | benign supporting |
| 1.276 | [1.260;1.294 | pathogenic strong |
| -0.544 | [-0.549;-0.54 | benign supporting |

| | | |
|---|---|---|
| -0.715 | [-0.756;-0.68 | benign supporting |
| 1.276 | [1.253;1.301 | pathogenic strong |
| -0.343 | [-0.394;-0.27 | benign supporting |
| 1.270 | [1.256;1.285 | pathogenic moderate |
| 0.579 | [-0.394;1.288 | pathogenic supporting |
| 1.276 | [1.248;1.307 | pathogenic strong |
| 1.276 | [1.225;1.338 | pathogenic strong |
| -0.678 | [-0.753;-0.62 | benign supporting |
| -0.998 | [-1.015;-0.98 | benign supporting |
| -0.531 | [-0.651;-0.40 | benign supporting |
| -0.375 | [-0.540;-0.15 | benign supporting |
| -0.827 | [-0.937;-0.68 | benign supporting |
| -1.084 | [-1.088;-1.08 | benign supporting |
| -1.090 | [-1.097;-1.08 | benign supporting |
| -1.120 | [-1.167;-1.07 | benign supporting |
| -1.165 | [-1.166;-1.16 | benign supporting |
| -1.107 | [-1.113;-1.10 | benign supporting |
| -1.115 | [-1.167;-1.08 | benign supporting |
| -1.115 | [-1.168;-0.89 | benign supporting |
| -0.774 | [-0.786;-0.76 | benign supporting |
| 1.276 | [-0.197;5.111 | pathogenic strong |
| 1.276 | [1.115;1.628 | pathogenic strong |
| -0.246 | [-0.472;0.420 | indeterminate |
| 0.579 | [0.408;0.721 | pathogenic supporting |
| -0.965 | [-1.094;-0.70 | benign supporting |
| 1.276 | [-0.301;5.105 | pathogenic strong |
| -0.881 | [-0.890;-0.87 | benign supporting |
| -1.097 | [-1.103;-1.09 | benign supporting |
| -1.089 | [-1.090;-1.08 | benign supporting |
| -0.129 | [-0.132;-0.12 | indeterminate |
| 1.252 | [1.238;1.265 | pathogenic moderate |
| 1.276 | [1.258;1.295 | pathogenic strong |
| 0.436 | [0.422;0.449 | pathogenic supporting |
| 1.276 | [1.263;1.290 | pathogenic strong |
| 1.276 | [1.216;1.347 | pathogenic strong |
| 1.276 | [1.173;1.431 | pathogenic strong |
| -0.420 | [-0.423;-0.41 | benign supporting |
| -1.052 | [-1.056;-1.04 | benign supporting |
| 1.276 | [1.271;1.281 | pathogenic strong |
| -1.095 | [-1.096;-1.09 | benign supporting |
| -0.738 | [-0.742;-0.73 | benign supporting |
| 0.530 | [0.508;0.551 | pathogenic supporting |
| 1.276 | [1.052;1.868 | pathogenic strong |

| | | |
|---|---|---|
| 1.276 | [1.124;1.595] | pathogenic strong |
| -0.629 | [-0.641;-0.61 | benign supporting |
| -0.909 | [-0.911;-0.90 | benign supporting |
| -0.716 | [-0.719;-0.71 | benign supporting |
| -1.098 | [-1.100;-1.09 | benign supporting |
| -0.728 | [-0.781;-0.68 | benign supporting |
| -0.968 | [-1.168;-0.83 | benign supporting |
| 1.017 | [0.643;1.186] | pathogenic moderate |
| -0.355 | [-0.563;-0.04 | benign supporting |
| -1.104 | [-1.112;-1.09 | benign supporting |
| 1.270 | [1.070;1.723] | pathogenic moderate |
| 1.276 | [1.172;1.428] | pathogenic strong |
| 1.276 | [1.234;1.327] | pathogenic strong |
| -0.288 | [-0.341;-0.22 | indeterminate |
| 1.276 | [1.221;1.344] | pathogenic strong |
| 1.069 | [0.928;1.138] | pathogenic moderate |
| 0.511 | [0.411;0.602] | pathogenic supporting |
| -1.086 | [-1.096;-1.08 | benign supporting |
| 1.276 | [1.173;1.431] | pathogenic strong |
| 1.276 | [1.218;1.346] | pathogenic strong |
| -0.836 | [-0.946;-0.69 | benign supporting |
| 1.276 | [1.081;1.737] | pathogenic strong |
| -0.463 | [-0.575;-0.37 | benign supporting |
| -0.425 | [-0.433;-0.41 | benign supporting |
| -1.084 | [-1.084;-1.08 | benign supporting |
| -1.016 | [-1.059;-0.96 | benign supporting |
| 1.230 | [1.117;1.379] | pathogenic moderate |
| 1.002 | [0.877;1.097] | pathogenic moderate |
| -0.256 | [-0.329;-0.15 | indeterminate |
| 1.276 | [1.180;1.411] | pathogenic strong |
| 1.276 | [-0.338;5.09 | pathogenic strong |
| 1.276 | [0.886;5.013] | pathogenic strong |
| 1.276 | [0.984;2.382] | pathogenic strong |
| -0.998 | [-1.164;-0.36 | benign supporting |
| 1.276 | [0.056;5.100] | pathogenic strong |
| 0.615 | [-0.704;2.71 | pathogenic supporting |
| -1.036 | [-1.164;-0.38 | benign supporting |
| 1.276 | [0.725;5.084] | pathogenic strong |
| -0.515 | [-1.105;0.79 | benign supporting |
| 1.276 | [-0.100;5.10 | pathogenic strong |
| 0.571 | [-0.263;1.161 | pathogenic supporting |
| 0.488 | [0.017;0.783] | pathogenic supporting |
| 1.276 | [1.208;1.365] | pathogenic strong |

| | | |
|---|---|---|
| -0.433 | [-0.641;-0.22 | benign supporting |
| -1.104 | [-1.115;-1.09 | benign supporting |
| -1.095 | [-1.102;-1.08 | benign supporting |
| -0.940 | [-0.958;-0.92 | benign supporting |
| -0.708 | [-0.759;-0.66 | benign supporting |
| -1.097 | [-1.104;-1.09 | benign supporting |
| -1.097 | [-1.105;-1.09 | benign supporting |
| -0.452 | [-0.518;-0.39 | benign supporting |
| -0.859 | [-0.883;-0.83 | benign supporting |
| -1.016 | [-1.048;-0.98 | benign supporting |
| -1.084 | [-1.118;-1.08 | benign supporting |
| -1.085 | [-1.088;-1.08 | benign supporting |
| -0.728 | [-0.775;-0.68 | benign supporting |
| -0.406 | [-0.474;-0.33 | benign supporting |
| -0.282 | [-0.368;-0.15 | indeterminate |
| -0.637 | [-0.648;-0.62 | benign supporting |
| -0.554 | [-0.564;-0.54 | benign supporting |
| -0.386 | [-0.418;-0.34 | benign supporting |
| -0.957 | [-1.077;-0.76 | benign supporting |
| -1.059 | [-1.114;-0.70 | benign supporting |
| -0.518 | [-0.539;-0.49 | benign supporting |
| -0.566 | [-0.651;-0.47 | benign supporting |
| -1.087 | [-1.102;-1.07 | benign supporting |
| -0.976 | [-0.984;-0.97 | benign supporting |
| -1.014 | [-1.028;-1.00 | benign supporting |
| 1.276 | [1.185;1.407 | pathogenic strong |
| -0.781 | [-0.810;-0.75 | benign supporting |
| -0.853 | [-1.106;-0.45 | benign supporting |
| 0.092 | [0.021;0.169 | indeterminate |
| -1.030 | [-1.041;-1.01 | benign supporting |
| -1.136 | [-1.169;-1.10 | benign supporting |
| -0.300 | [-0.619;0.57 | indeterminate |
| 1.276 | [1.247;1.309 | pathogenic strong |
| -1.123 | [-1.145;-1.10 | benign supporting |
| 0.779 | [0.528;1.001 | pathogenic moderate |
| -1.106 | [-1.115;-1.09 | benign supporting |
| -1.101 | [-1.164;-0.69 | benign supporting |
| -1.091 | [-1.124;-1.08 | benign supporting |
| 1.276 | [0.000;5.102 | pathogenic strong |
| 0.526 | [-0.307;1.17 | pathogenic supporting |
| -0.230 | [-0.540;0.71 | indeterminate |
| 1.104 | [0.932;1.170 | pathogenic moderate |
| 1.276 | [0.478;5.096 | pathogenic strong |

| | | |
|---|---|---|
| 0.695 | [-0.416;1.411 | pathogenic moderate |
| -0.434 | [-0.474;-0.40 | benign supporting |
| 0.012 | [-0.127;0.394 | indeterminate |
| -0.912 | [-0.916;-0.90 | benign supporting |
| -0.565 | [-0.593;-0.53 | benign supporting |
| -1.126 | [-1.169;-1.07 | benign supporting |
| 1.276 | [0.567;5.095 | pathogenic strong |
| -0.955 | [-1.108;-0.62 | benign supporting |
| -0.630 | [-0.871;-0.47 | benign supporting |
| -0.353 | [-0.480;-0.17 | benign supporting |
| 1.276 | [1.245;1.311 | pathogenic strong |
| -0.939 | [-0.956;-0.92 | benign supporting |
| -1.169 | [-1.169;-1.10 | benign supporting |
| -1.159 | [-1.168;-0.83 | benign supporting |
| 1.171 | [1.150;1.193 | pathogenic moderate |
| -0.416 | [-0.807;0.014 | benign supporting |
| -0.674 | [-0.681;-0.66 | benign supporting |
| -0.397 | [-0.405;-0.38 | benign supporting |
| -1.027 | [-1.035;-1.02 | benign supporting |
| -0.971 | [-0.976;-0.96 | benign supporting |
| -0.275 | [-0.296;-0.25 | indeterminate |
| 1.107 | [1.019;1.146 | pathogenic moderate |
| -0.439 | [-0.444;-0.43 | benign supporting |
| -1.166 | [-1.169;-1.11 | benign supporting |
| 1.253 | [1.052;1.575 | pathogenic moderate |
| 0.605 | [0.566;0.642 | pathogenic supporting |
| -0.887 | [-0.895;-0.87 | benign supporting |
| 1.276 | [1.156;1.466 | pathogenic strong |
| -0.890 | [-0.992;-0.75 | benign supporting |
| 0.757 | [0.481;0.992 | pathogenic moderate |
| -0.562 | [-0.574;-0.55 | benign supporting |
| -0.911 | [-0.920;-0.89 | benign supporting |
| -1.060 | [-1.066;-1.05 | benign supporting |
| -1.114 | [-1.126;-1.10 | benign supporting |
| -0.455 | [-0.467;-0.44 | benign supporting |
| -0.961 | [-0.967;-0.95 | benign supporting |
| 1.276 | [0.677;5.091 | pathogenic strong |
| 1.276 | [1.176;1.422 | pathogenic strong |
| 1.276 | [1.080;1.704 | pathogenic strong |
| 0.574 | [0.429;0.700 | pathogenic supporting |
| 1.276 | [1.175;1.435 | pathogenic strong |
| 0.752 | [0.606;0.877 | pathogenic moderate |
| -0.905 | [-0.925;-0.88 | benign supporting |

| | | |
|---|---|---|
| -0.986 | [-1.020;-0.96 | benign supporting |
| 1.276 | [0.016;5.100 | pathogenic strong |
| 1.276 | [1.158;1.463 | pathogenic strong |
| -1.090 | [-1.096;-1.08 | benign supporting |
| 1.276 | [0.892;5.030 | pathogenic strong |
| -0.018 | [-0.216;0.50 | indeterminate |
| -0.060 | [-0.077;-0.04 | indeterminate |
| -1.166 | [-1.167;-1.16 | benign supporting |
| -0.919 | [-0.932;-0.90 | benign supporting |
| -1.002 | [-1.019;-0.98 | benign supporting |
| -1.045 | [-1.052;-1.03 | benign supporting |
| -0.573 | [-0.578;-0.56 | benign supporting |
| 1.251 | [1.229;1.274 | pathogenic moderate |
| -0.440 | [-0.513;-0.38 | benign supporting |
| -0.396 | [-0.405;-0.38 | benign supporting |
| -1.002 | [-1.036;-0.95 | benign supporting |
| 1.276 | [1.223;1.340 | pathogenic strong |
| -0.533 | [-0.546;-0.52 | benign supporting |
| -0.278 | [-0.282;-0.27 | indeterminate |
| 1.276 | [1.236;1.326 | pathogenic strong |
| -1.015 | [-1.112;-0.68 | benign supporting |
| 1.276 | [1.242;1.312 | pathogenic strong |
| -1.097 | [-1.115;-1.07 | benign supporting |
| -1.089 | [-1.098;-1.08 | benign supporting |
| -0.973 | [-0.974;-0.97 | benign supporting |
| -0.596 | [-0.600;-0.59 | benign supporting |
| -1.105 | [-1.114;-1.09 | benign supporting |
| 0.710 | [0.073;1.085 | pathogenic moderate |
| -0.349 | [-1.087;1.17 | benign supporting |
| -0.101 | [-0.635;1.16 | indeterminate |
| 0.913 | [0.353;1.184 | pathogenic moderate |
| -0.089 | [-0.220;0.15 | indeterminate |
| 0.946 | [0.404;1.198 | pathogenic moderate |
| 1.276 | [-0.122;5.10 | pathogenic strong |
| 1.276 | [-0.065;5.09 | pathogenic strong |
| -0.885 | [-0.890;-0.88 | benign supporting |
| 1.106 | [0.751;1.235 | pathogenic moderate |
| 1.276 | [0.757;5.074 | pathogenic strong |
| -0.643 | [-1.131;0.81 | benign supporting |
| -1.167 | [-1.169;-1.03 | benign supporting |
| 1.276 | [1.027;2.010 | pathogenic strong |
| 1.076 | [0.893;1.156 | pathogenic moderate |
| 1.276 | [1.216;1.351 | pathogenic strong |

| | | |
|---|---|---|
| 1.276 | [1.193;1.392 | pathogenic strong |
| -0.966 | [-0.976;-0.95 | benign supporting |
| -0.839 | [-0.840;-0.83 | benign supporting |
| -1.084 | [-1.167;-1.08 | benign supporting |
| -1.074 | [-1.088;-1.05 | benign supporting |
| -0.262 | [-0.337;-0.16 | indeterminate |
| -0.577 | [-0.910;-0.35 | benign supporting |
| -0.245 | [-0.389;-0.01 | indeterminate |
| -0.784 | [-0.936;-0.63 | benign supporting |
| -0.464 | [-0.502;-0.43 | benign supporting |
| -0.483 | [-1.164;3.567 | benign supporting |
| -0.435 | [-0.514;-0.37 | benign supporting |
| 0.656 | [0.561;0.740 | pathogenic moderate |
| 1.276 | [0.917;4.993 | pathogenic strong |
| -1.079 | [-1.104;-1.04 | benign supporting |
| -0.997 | [-1.016;-0.96 | benign supporting |
| -0.612 | [-0.626;-0.59 | benign supporting |
| -1.092 | [-1.114;-1.08 | benign supporting |
| -1.150 | [-1.169;-0.96 | benign supporting |
| -0.538 | [-0.572;-0.50 | benign supporting |
| -0.255 | [-0.297;-0.20 | indeterminate |
| -0.428 | [-0.457;-0.40 | benign supporting |
| 0.014 | [-0.051;0.174 | indeterminate |
| -1.084 | [-1.086;-1.08 | benign supporting |
| -1.055 | [-1.112;-0.82 | benign supporting |
| -1.087 | [-2.744;4.629 | benign supporting |
| -0.385 | [-0.444;-0.31 | benign supporting |
| -0.637 | [-0.652;-0.62 | benign supporting |
| -0.694 | [-0.741;-0.66 | benign supporting |
| -0.191 | [-0.238;-0.14 | indeterminate |
| 1.192 | [1.137;1.252 | pathogenic moderate |
| 1.232 | [1.194;1.272 | pathogenic moderate |
| -1.127 | [-1.138;-1.11 | benign supporting |
| -0.077 | [-0.117;-0.02 | indeterminate |
| 0.411 | [0.299;0.511 | pathogenic supporting |
| -0.871 | [-0.905;-0.83 | benign supporting |
| -1.067 | [-1.076;-1.04 | benign supporting |
| -0.641 | [-0.661;-0.62 | benign supporting |
| -1.115 | [-1.116;-1.10 | benign supporting |
| 1.276 | [1.242;1.316 | pathogenic strong |
| -0.460 | [-0.482;-0.44 | benign supporting |
| 1.276 | [1.230;1.331 | pathogenic strong |
| -0.656 | [-0.693;-0.62 | benign supporting |

| | | |
|---|---|---|
| 0.834 | [0.771;0.902 | pathogenic moderate |
| -1.064 | [-1.071;-1.04 | benign supporting |
| -0.834 | [-0.858;-0.80 | benign supporting |
| -1.084 | [-1.122;-1.08 | benign supporting |
| -0.823 | [-1.165;1.10 | benign supporting |
| -1.090 | [-1.122;-1.08 | benign supporting |
| 1.276 | [0.971;2.806 | pathogenic strong |
| -0.966 | [-1.088;-0.73 | benign supporting |
| -0.111 | [-0.312;0.36 | indeterminate |
| 1.276 | [0.000;5.104 | pathogenic strong |
| -1.067 | [-1.090;-1.01 | benign supporting |
| -0.120 | [-0.446;0.80 | indeterminate |
| -1.090 | [-1.113;-1.08 | benign supporting |
| 0.889 | [-1.099;5.04 | pathogenic moderate |
| -1.101 | [-1.112;-1.09 | benign supporting |
| -1.101 | [-1.114;-1.08 | benign supporting |
| -1.083 | [-1.166;-0.26 | benign supporting |
| -0.831 | [-1.107;-0.41 | benign supporting |
| 1.276 | [-1.124;4.96 | pathogenic strong |
| 1.276 | [-0.382;5.09 | pathogenic strong |
| -0.608 | [-0.700;-0.51 | benign supporting |
| -0.050 | [-0.292;0.58 | indeterminate |
| -0.523 | [-1.065;-0.03 | benign supporting |
| -0.988 | [-1.168;-0.83 | benign supporting |
| 1.276 | [1.159;1.467 | pathogenic strong |
| 0.477 | [-0.109;0.95 | pathogenic supporting |
| -0.914 | [-0.947;-0.87 | benign supporting |
| -0.396 | [-0.454;-0.33 | benign supporting |
| -0.781 | [-0.833;-0.72 | benign supporting |
| -0.838 | [-0.844;-0.83 | benign supporting |
| -0.659 | [-0.695;-0.62 | benign supporting |
| -0.563 | [-0.599;-0.52 | benign supporting |
| -1.115 | [-1.116;-1.09 | benign supporting |
| -1.079 | [-1.099;-1.05 | benign supporting |
| -1.068 | [-1.093;-1.05 | benign supporting |
| -0.993 | [-1.059;-0.91 | benign supporting |
| -0.584 | [-0.627;-0.53 | benign supporting |
| -0.710 | [-0.760;-0.67 | benign supporting |
| 1.276 | [0.513;5.101 | pathogenic strong |
| -0.634 | [-0.745;-0.55 | benign supporting |
| -0.340 | [-0.378;-0.29 | benign supporting |
| -0.968 | [-1.001;-0.93 | benign supporting |
| -1.040 | [-1.113;-0.69 | benign supporting |

| | | |
|---|---|---|
| -1.112 | [-1.116;-1.09 | benign supporting |
| 1.138 | [1.079;1.185] | pathogenic moderate |
| 1.276 | [1.131;1.540] | pathogenic strong |
| 1.276 | [1.163;1.462] | pathogenic strong |
| 1.276 | [1.127;1.561] | pathogenic strong |
| -0.918 | [-0.925;-0.91 | benign supporting |
| -0.689 | [-1.096;-0.34 | benign supporting |
| -1.095 | [-1.109;-1.08 | benign supporting |
| 1.276 | [1.193;1.385] | pathogenic strong |
| 1.276 | [1.146;1.501] | pathogenic strong |
| -1.160 | [-1.169;-1.09 | benign supporting |
| 1.276 | [1.117;1.619] | pathogenic strong |
| -0.761 | [-0.787;-0.73 | benign supporting |
| 1.276 | [1.186;1.397] | pathogenic strong |
| 0.398 | [-1.114;5.017 | pathogenic supporting |
| -0.261 | [-0.629;0.790 | indeterminate |
| 1.276 | [0.910;4.895] | pathogenic strong |
| -0.313 | [-0.584;0.363 | indeterminate |
| 0.392 | [0.110;0.607] | pathogenic supporting |
| -1.109 | [-1.115;-1.07 | benign supporting |
| -0.531 | [-0.883;-0.27 | benign supporting |
| -0.627 | [-0.644;-0.61 | benign supporting |
| -0.607 | [-0.959;-0.36 | benign supporting |
| -0.383 | [-0.484;-0.26 | benign supporting |
| 1.243 | [1.186;1.305] | pathogenic moderate |
| 0.254 | [0.053;0.401] | indeterminate |
| -0.635 | [-0.654;-0.62 | benign supporting |
| -0.526 | [-0.548;-0.50 | benign supporting |
| 0.522 | [0.388;0.635] | pathogenic supporting |
| -0.552 | [-0.605;-0.49 | benign supporting |
| -0.882 | [-0.918;-0.84 | benign supporting |
| -1.100 | [-1.102;-1.09 | benign supporting |
| 1.276 | [0.978;2.214] | pathogenic strong |
| -1.084 | [-1.086;-1.08 | benign supporting |
| -0.825 | [-0.837;-0.80 | benign supporting |
| -0.574 | [-0.616;-0.53 | benign supporting |
| -0.820 | [-0.881;-0.73 | benign supporting |
| 1.276 | [1.235;1.323] | pathogenic strong |
| -0.133 | [-0.191;-0.08 | indeterminate |
| -0.736 | [-0.779;-0.69 | benign supporting |
| -0.767 | [-0.816;-0.72 | benign supporting |
| -1.111 | [-1.116;-1.09 | benign supporting |
| -0.468 | [-0.639;-0.32 | benign supporting |

| | | |
|---|---|---|
| 1.217 | [1.175;1.261 | pathogenic moderate |
| 1.276 | [1.214;1.356 | pathogenic strong |
| 1.276 | [1.216;1.353 | pathogenic strong |
| -0.472 | [-0.498;-0.45 | benign supporting |
| 1.276 | [1.001;2.359 | pathogenic strong |
| 1.137 | [0.860;1.270 | pathogenic moderate |
| 0.911 | [0.605;1.129 | pathogenic moderate |
| 1.276 | [1.243;1.315 | pathogenic strong |
| -0.625 | [-0.664;-0.58 | benign supporting |
| 1.276 | [1.119;1.598 | pathogenic strong |
| -0.822 | [-0.849;-0.78 | benign supporting |
| -0.374 | [-0.386;-0.36 | benign supporting |
| 1.017 | [0.582;1.202 | pathogenic moderate |
| 1.276 | [1.185;1.403 | pathogenic strong |
| 0.252 | [-0.148;0.72 | indeterminate |
| 0.020 | [-0.114;0.38 | indeterminate |
| -0.138 | [-0.170;-0.10 | indeterminate |
| 1.276 | [1.229;1.332 | pathogenic strong |
| -0.656 | [-1.168;4.07 | benign supporting |
| -1.109 | [-1.115;-1.09 | benign supporting |
| -0.154 | [-0.262;-0.04 | indeterminate |
| 0.623 | [0.280;0.860 | pathogenic supporting |
| 0.825 | [0.782;0.870 | pathogenic moderate |
| 1.175 | [1.142;1.210 | pathogenic moderate |
| -0.263 | [-0.274;-0.25 | indeterminate |
| -0.023 | [-0.049;0.00 | indeterminate |
| 1.262 | [1.235;1.291 | pathogenic moderate |
| 1.216 | [1.210;1.222 | pathogenic moderate |
| 0.330 | [0.201;0.430 | pathogenic supporting |
| -1.165 | [-1.166;-1.16 | benign supporting |
| 1.276 | [1.250;1.305 | pathogenic strong |
| -1.047 | [-1.052;-1.04 | benign supporting |
| -0.403 | [-0.410;-0.39 | benign supporting |
| 1.276 | [1.248;1.307 | pathogenic strong |
| -0.311 | [-0.328;-0.29 | indeterminate |
| 1.276 | [1.260;1.293 | pathogenic strong |
| 0.814 | [0.767;0.864 | pathogenic moderate |
| 1.276 | [1.262;1.292 | pathogenic strong |
| 0.829 | [0.791;0.869 | pathogenic moderate |
| -1.008 | [-1.013;-1.00 | benign supporting |
| -1.088 | [-1.092;-1.08 | benign supporting |
| 1.276 | [0.035;5.107 | pathogenic strong |
| -0.704 | [-0.956;-0.53 | benign supporting |

| | | |
|---|---|---|
| 1.276 | [-1.116;4.993 | pathogenic strong |
| -0.318 | [-0.379;-0.24 | benign supporting |
| -0.659 | [-0.725;-0.61 | benign supporting |
| -1.044 | [-1.071;-1.00 | benign supporting |
| 0.656 | [-0.106;1.129 | pathogenic moderate |
| -1.106 | [-1.121;-1.09 | benign supporting |
| 0.945 | [0.763;1.103 | pathogenic moderate |
| -1.084 | [-1.089;-1.08 | benign supporting |
| -1.150 | [-1.170;-1.12 | benign supporting |
| -1.169 | [-1.169;-1.14 | benign supporting |
| -1.104 | [-1.128;-1.08 | benign supporting |
| -0.839 | [-3.721;-0.83 | benign supporting |
| -0.321 | [-0.720;0.751 | benign supporting |
| 1.276 | [1.229;1.334 | pathogenic strong |
| -0.360 | [-1.099;1.230 | benign supporting |
| -0.261 | [-0.305;-0.20 | indeterminate |
| 1.276 | [1.237;1.322 | pathogenic strong |
| -1.065 | [-1.070;-1.05 | benign supporting |
| -1.111 | [-1.116;-1.09 | benign supporting |
| -0.191 | [-0.416;0.451 | indeterminate |
| 0.874 | [0.785;0.974 | pathogenic moderate |
| 0.831 | [0.760;0.907 | pathogenic moderate |
| 0.399 | [0.066;0.634 | pathogenic supporting |
| -1.110 | [-1.115;-1.10 | benign supporting |
| -0.621 | [-0.656;-0.58 | benign supporting |
| 1.276 | [1.257;1.298 | pathogenic strong |
| -0.871 | [-0.887;-0.85 | benign supporting |
| -0.731 | [-0.763;-0.70 | benign supporting |
| 0.168 | [-0.058;0.452 | indeterminate |
| 0.533 | [0.429;0.624 | pathogenic supporting |
| -0.766 | [-0.828;-0.70 | benign supporting |
| -0.371 | [-0.389;-0.35 | benign supporting |
| 1.276 | [1.154;1.484 | pathogenic strong |
| -0.261 | [-0.464;0.283 | indeterminate |
| 1.276 | [1.193;1.385 | pathogenic strong |
| -0.479 | [-0.539;-0.42 | benign supporting |
| -0.351 | [-0.369;-0.33 | benign supporting |
| -0.316 | [-0.337;-0.29 | indeterminate |
| -0.878 | [-0.921;-0.83 | benign supporting |
| -1.095 | [-1.112;-1.08 | benign supporting |
| 0.039 | [-0.052;0.265 | indeterminate |
| 1.276 | [1.189;1.396 | pathogenic strong |
| 1.105 | [0.868;1.190 | pathogenic moderate |

| | | |
|---|---|---|
| 1.276 | [1.157;1.473 | pathogenic strong |
| -1.116 | [-1.116;-1.10 | benign supporting |
| -1.115 | [-1.116;-1.09 | benign supporting |
| 1.276 | [1.223;1.341 | pathogenic strong |
| 1.002 | [0.927;1.056 | pathogenic moderate |
| -1.011 | [-1.031;-0.99 | benign supporting |
| -0.722 | [-0.774;-0.68 | benign supporting |
| -0.622 | [-0.642;-0.60 | benign supporting |
| -1.094 | [-1.106;-1.08 | benign supporting |
| -0.402 | [-0.426;-0.38 | benign supporting |
| 1.276 | [1.159;1.457 | pathogenic strong |
| -0.293 | [-0.331;-0.24 | indeterminate |
| 1.240 | [1.213;1.270 | pathogenic moderate |
| -0.562 | [-0.627;-0.49 | benign supporting |
| -0.359 | [-0.378;-0.34 | benign supporting |
| -1.061 | [-1.070;-1.04 | benign supporting |
| -0.422 | [-0.443;-0.40 | benign supporting |
| -1.073 | [-1.082;-1.06 | benign supporting |
| -0.400 | [-0.508;-0.29 | benign supporting |
| 1.276 | [1.225;1.338 | pathogenic strong |
| -0.823 | [-0.908;-0.70 | benign supporting |
| 1.276 | [1.157;1.474 | pathogenic strong |
| -0.615 | [-0.655;-0.57 | benign supporting |
| -1.098 | [-1.110;-1.09 | benign supporting |
| -0.371 | [-0.418;-0.31 | benign supporting |
| -1.099 | [-1.115;-1.05 | benign supporting |
| -0.658 | [-0.861;-0.53 | benign supporting |
| -0.840 | [-1.013;-0.63 | benign supporting |
| 1.223 | [1.154;1.298 | pathogenic moderate |
| -1.087 | [-1.092;-1.08 | benign supporting |
| -1.064 | [-1.074;-1.03 | benign supporting |
| 1.276 | [1.236;1.322 | pathogenic strong |
| -0.360 | [-0.445;-0.24 | benign supporting |
| 1.217 | [1.194;1.240 | pathogenic moderate |
| -0.450 | [-0.549;-0.37 | benign supporting |
| 1.276 | [1.226;1.337 | pathogenic strong |
| -0.972 | [-0.996;-0.95 | benign supporting |
| -0.964 | [-1.008;-0.92 | benign supporting |
| -1.110 | [-1.116;-1.10 | benign supporting |
| -0.251 | [-0.319;-0.15 | indeterminate |
| -0.567 | [-0.623;-0.50 | benign supporting |
| 0.833 | [0.782;0.886 | pathogenic moderate |
| 1.276 | [1.257;1.298 | pathogenic strong |

| | | |
|---|---|---|
| -1.101 | [-1.108;-1.09 | benign supporting |
| -0.103 | [-0.147;-0.06 | indeterminate |
| 1.076 | [1.000;1.122 | pathogenic moderate |
| 0.806 | [0.708;0.908 | pathogenic moderate |
| -0.611 | [-0.628;-0.59 | benign supporting |
| -1.089 | [-1.090;-1.08 | benign supporting |
| -1.131 | [-1.145;-1.11 | benign supporting |
| -1.164 | [-1.166;-1.16 | benign supporting |
| 1.276 | [1.190;1.394 | pathogenic strong |
| -0.155 | [-0.189;-0.12 | indeterminate |
| 1.276 | [1.242;1.316 | pathogenic strong |
| -0.449 | [-0.605;-0.32 | benign supporting |
| -0.126 | [-0.177;-0.08 | indeterminate |
| -0.621 | [-0.641;-0.60 | benign supporting |
| -0.895 | [-0.961;-0.82 | benign supporting |
| -1.117 | [-1.159;-1.09 | benign supporting |
| 0.451 | [0.372;0.527 | pathogenic supporting |
| 1.116 | [1.099;1.132 | pathogenic moderate |
| -0.404 | [-0.432;-0.37 | benign supporting |
| 0.992 | [0.723;1.141 | pathogenic moderate |
| -0.345 | [-0.363;-0.32 | benign supporting |
| 0.506 | [0.398;0.601 | pathogenic supporting |
| 1.276 | [1.251;1.303 | pathogenic strong |
| 1.276 | [1.253;1.301 | pathogenic strong |
| 1.276 | [1.248;1.306 | pathogenic strong |
| -0.370 | [-0.400;-0.33 | benign supporting |
| 1.276 | [1.235;1.325 | pathogenic strong |
| -1.110 | [-1.115;-1.10 | benign supporting |
| 0.013 | [-0.025;0.08 | indeterminate |
| 1.276 | [-1.111;5.02 | pathogenic strong |
| 1.276 | [1.266;1.287 | pathogenic strong |
| 1.276 | [1.219;1.346 | pathogenic strong |
| -0.303 | [-0.332;-0.27 | indeterminate |
| 1.276 | [1.213;1.354 | pathogenic strong |
| 1.276 | [1.238;1.320 | pathogenic strong |
| -1.095 | [-1.115;-1.05 | benign supporting |
| -1.110 | [-1.115;-1.09 | benign supporting |
| -1.011 | [-1.058;-0.95 | benign supporting |
| 1.276 | [1.268;1.285 | pathogenic strong |
| -0.244 | [-0.358;-0.07 | indeterminate |
| 1.276 | [1.229;1.333 | pathogenic strong |
| -1.045 | [-1.050;-1.04 | benign supporting |
| -0.327 | [-0.352;-0.29 | benign supporting |

| | | |
|---|---|---|
| 1.276 | [1.249;1.306 | pathogenic strong |
| -0.676 | [-0.841;-0.58 | benign supporting |
| -1.098 | [-1.100;-1.09 | benign supporting |
| 1.189 | [1.174;1.204 | pathogenic moderate |
| 1.276 | [1.240;1.317 | pathogenic strong |
| -0.587 | [-0.602;-0.57 | benign supporting |
| -0.551 | [-0.564;-0.53 | benign supporting |
| 0.717 | [0.553;0.849 | pathogenic moderate |
| -0.181 | [-0.401;0.41 | indeterminate |
| 0.751 | [0.711;0.790 | pathogenic moderate |
| -0.957 | [-1.075;-0.77 | benign supporting |
| 1.276 | [1.253;1.300 | pathogenic strong |
| 1.276 | [1.225;1.337 | pathogenic strong |
| 1.276 | [1.237;1.321 | pathogenic strong |
| -1.082 | [-1.104;-1.06 | benign supporting |
| 1.276 | [1.115;1.613 | pathogenic strong |
| 1.276 | [1.264;1.289 | pathogenic strong |
| -0.246 | [-0.305;-0.17 | indeterminate |
| 1.276 | [1.244;1.310 | pathogenic strong |
| 1.276 | [1.224;1.338 | pathogenic strong |
| -0.364 | [-0.380;-0.34 | benign supporting |
| 1.276 | [1.220;1.346 | pathogenic strong |
| -1.090 | [-1.091;-1.08 | benign supporting |
| 1.114 | [1.084;1.134 | pathogenic moderate |
| 0.461 | [0.342;0.573 | pathogenic supporting |
| 1.276 | [1.218;1.346 | pathogenic strong |
| -1.095 | [-1.098;-1.09 | benign supporting |
| -0.179 | [-0.227;-0.13 | indeterminate |
| -0.505 | [-1.088;0.51 | benign supporting |
| -0.530 | [-0.554;-0.50 | benign supporting |
| 1.228 | [1.076;1.434 | pathogenic moderate |
| -0.013 | [-0.226;0.53 | indeterminate |
| 1.276 | [1.218;1.346 | pathogenic strong |
| 1.276 | [1.176;1.425 | pathogenic strong |
| 0.570 | [-0.027;0.95 | pathogenic supporting |
| -0.638 | [-0.704;-0.58 | benign supporting |
| 1.276 | [1.232;1.327 | pathogenic strong |
| 1.230 | [1.224;1.236 | pathogenic moderate |
| 1.276 | [1.111;1.613 | pathogenic strong |
| 0.566 | [0.518;0.607 | pathogenic supporting |
| -0.841 | [-0.847;-0.83 | benign supporting |
| 1.194 | [1.158;1.231 | pathogenic moderate |
| 1.276 | [1.259;1.294 | pathogenic strong |

| | | |
|---|---|---|
| -0.242 | [-0.302;-0.16 | indeterminate |
| 1.276 | [0.864;5.015 | pathogenic strong |
| 1.237 | [1.217;1.260 | pathogenic moderate |
| 1.276 | [1.257;1.297 | pathogenic strong |
| -1.135 | [-1.140;-1.13 | benign supporting |
| -1.094 | [-1.095;-1.09 | benign supporting |
| 0.498 | [-0.355;1.19 | pathogenic supporting |
| 0.964 | [0.817;1.084 | pathogenic moderate |
| 1.276 | [1.215;1.351 | pathogenic strong |
| -0.200 | [-0.890;1.19 | indeterminate |
| 0.838 | [0.821;0.855 | pathogenic moderate |
| 1.276 | [1.237;1.321 | pathogenic strong |
| -0.332 | [-0.853;0.87 | benign supporting |
| -0.737 | [-0.922;-0.60 | benign supporting |
| -1.128 | [-1.168;-1.09 | benign supporting |
| 1.217 | [1.181;1.255 | pathogenic moderate |
| 1.270 | [1.218;1.331 | pathogenic moderate |
| 1.054 | [0.831;1.154 | pathogenic moderate |
| -0.760 | [-0.784;-0.73 | benign supporting |
| 1.276 | [1.252;1.302 | pathogenic strong |
| 1.276 | [1.251;1.303 | pathogenic strong |
| 1.156 | [0.935;1.289 | pathogenic moderate |
| 1.194 | [1.171;1.218 | pathogenic moderate |
| 1.276 | [1.120;1.594 | pathogenic strong |
| -1.103 | [-1.106;-1.10 | benign supporting |
| -0.291 | [-0.332;-0.24 | indeterminate |
| 1.276 | [1.067;1.816 | pathogenic strong |
| -0.114 | [-0.591;1.11 | indeterminate |
| 1.276 | [1.237;1.321 | pathogenic strong |
| -1.025 | [-1.087;-0.91 | benign supporting |
| 0.286 | [0.183;0.369 | indeterminate |
| -0.660 | [-0.678;-0.64 | benign supporting |
| -1.033 | [-1.067;-0.99 | benign supporting |
| -0.906 | [-0.908;-0.90 | benign supporting |
| -0.482 | [-0.617;-0.36 | benign supporting |
| 0.797 | [0.200;1.136 | pathogenic moderate |
| -1.085 | [-1.086;-1.08 | benign supporting |
| 0.010 | [-0.045;0.13 | indeterminate |
| -1.038 | [-1.049;-1.02 | benign supporting |
| 1.276 | [1.240;1.318 | pathogenic strong |
| 0.930 | [0.850;1.009 | pathogenic moderate |
| -1.100 | [-1.105;-1.09 | benign supporting |
| -0.348 | [-0.362;-0.33 | benign supporting |

| | | |
|---|---|---|
| -1.101 | [-1.108;-1.09 | benign supporting |
| -1.104 | [-1.107;-1.10 | benign supporting |
| -1.067 | [-1.071;-1.06 | benign supporting |
| -1.056 | [-1.072;-1.02 | benign supporting |
| -0.103 | [-0.231;0.083 | indeterminate |
| 1.083 | [0.698;1.222 | pathogenic moderate |
| -0.079 | [-0.127;-0.01 | indeterminate |
| -1.115 | [-1.116;-1.11 | benign supporting |
| -0.543 | [-0.551;-0.53 | benign supporting |
| 0.540 | [-0.069;0.970 | pathogenic supporting |
| -1.140 | [-1.169;-1.08 | benign supporting |
| -1.102 | [-1.115;-1.08 | benign supporting |
| -1.143 | [-1.149;-1.13 | benign supporting |
| -0.423 | [-1.099;1.104 | benign supporting |
| -1.165 | [-1.166;-1.16 | benign supporting |
| 0.769 | [0.619;0.908 | pathogenic moderate |
| -1.067 | [-1.070;-1.06 | benign supporting |
| -0.575 | [-0.596;-0.55 | benign supporting |
| 1.276 | [1.248;1.307 | pathogenic strong |
| 1.130 | [1.103;1.155 | pathogenic moderate |
| -0.483 | [-0.521;-0.44 | benign supporting |
| -1.086 | [-1.091;-1.08 | benign supporting |
| -0.334 | [-0.346;-0.32 | benign supporting |
| -0.457 | [-0.466;-0.44 | benign supporting |
| 1.276 | [1.104;1.692 | pathogenic strong |
| 1.210 | [1.142;1.288 | pathogenic moderate |
| 0.696 | [-0.093;1.142 | pathogenic moderate |
| -0.212 | [-0.425;0.384 | indeterminate |
| -1.046 | [-1.052;-1.04 | benign supporting |
| 1.276 | [1.106;1.685 | pathogenic strong |
| -1.093 | [-1.125;-1.08 | benign supporting |
| -1.073 | [-1.083;-1.06 | benign supporting |
| 1.276 | [1.048;1.807 | pathogenic strong |
| 1.276 | [1.218;1.347 | pathogenic strong |
| 1.276 | [0.743;5.079 | pathogenic strong |
| -1.028 | [-1.108;-0.83 | benign supporting |
| 1.144 | [-0.403;5.079 | pathogenic moderate |
| -0.418 | [-0.549;-0.30 | benign supporting |
| 1.209 | [1.171;1.247 | pathogenic moderate |
| 1.276 | [0.921;4.971 | pathogenic strong |
| 1.276 | [0.508;5.102 | pathogenic strong |
| 1.276 | [-0.341;5.108 | pathogenic strong |
| -1.088 | [-1.094;-1.08 | benign supporting |

| | | |
|---|---|---|
| 1.276 | [1.235;1.323] | pathogenic strong |
| -0.405 | [-0.609;-0.17 | benign supporting |
| 1.109 | [1.045;1.140] | pathogenic moderate |
| 1.276 | [0.693;5.093] | pathogenic strong |
| 1.276 | [0.357;5.100] | pathogenic strong |
| 1.145 | [1.122;1.166] | pathogenic moderate |
| -0.442 | [-0.590;-0.32 | benign supporting |
| -1.108 | [-1.115;-1.09 | benign supporting |
| 1.114 | [0.931;1.192] | pathogenic moderate |
| 1.156 | [1.140;1.174] | pathogenic moderate |
| 0.423 | [0.230;0.580] | pathogenic supporting |
| 1.276 | [-0.102;5.108 | pathogenic strong |
| -1.108 | [-1.112;-1.10 | benign supporting |
| -0.922 | [-0.926;-0.91 | benign supporting |
| 1.276 | [1.039;1.927] | pathogenic strong |
| 1.276 | [1.168;1.438] | pathogenic strong |
| 1.163 | [1.136;1.194] | pathogenic moderate |
| 1.276 | [1.246;1.310] | pathogenic strong |
| 1.276 | [1.218;1.350] | pathogenic strong |
| -0.656 | [-0.676;-0.63 | benign supporting |
| 1.276 | [1.258;1.295] | pathogenic strong |
| 1.074 | [0.826;1.175] | pathogenic moderate |
| -1.138 | [-1.168;-0.91 | benign supporting |
| 0.564 | [-0.058;0.976 | pathogenic supporting |
| -1.106 | [-1.168;-0.85 | benign supporting |
| 1.276 | [1.237;1.321] | pathogenic strong |
| -1.165 | [-1.166;-1.16 | benign supporting |
| -0.718 | [-1.111;-0.12 | benign supporting |
| -1.006 | [-1.107;-0.75 | benign supporting |
| -1.139 | [-1.169;-1.08 | benign supporting |
| 1.231 | [1.164;1.310] | pathogenic moderate |
| 1.276 | [1.133;1.536] | pathogenic strong |
| -0.961 | [-0.975;-0.94 | benign supporting |
| -0.670 | [-0.679;-0.66 | benign supporting |
| 1.160 | [1.150;1.171] | pathogenic moderate |
| 0.766 | [0.012;1.150] | pathogenic moderate |
| -0.657 | [-0.852;-0.54 | benign supporting |
| -1.103 | [-1.106;-1.10 | benign supporting |
| 1.072 | [0.991;1.122] | pathogenic moderate |
| -1.166 | [-1.170;-1.16 | benign supporting |
| -0.219 | [-0.435;0.415 | indeterminate |
| 1.276 | [0.657;5.090] | pathogenic strong |
| 1.276 | [1.193;1.388] | pathogenic strong |

| | | |
|---|---|---|
| 1.276 | [1.207;1.367 | pathogenic strong |
| -0.880 | [-0.896;-0.86 | benign supporting |
| -1.166 | [-1.169;-1.13 | benign supporting |
| -0.736 | [-1.067;-0.47 | benign supporting |
| -1.087 | [-1.115;-0.85 | benign supporting |
| 1.276 | [1.236;1.324 | pathogenic strong |
| 1.180 | [-0.350;5.078 | pathogenic moderate |
| 1.276 | [1.171;1.431 | pathogenic strong |
| -1.106 | [-1.115;-1.09 | benign supporting |
| 1.276 | [1.022;1.965 | pathogenic strong |
| 1.276 | [1.208;1.364 | pathogenic strong |
| 1.276 | [1.094;1.748 | pathogenic strong |
| -0.658 | [-0.689;-0.63 | benign supporting |
| 1.276 | [0.940;3.376 | pathogenic strong |
| -0.200 | [-1.161;4.975 | indeterminate |
| -0.233 | [-0.358;-0.05 | indeterminate |
| 1.276 | [0.591;5.091 | pathogenic strong |
| -1.014 | [-1.018;-1.01 | benign supporting |
| 1.276 | [0.000;5.107 | pathogenic strong |
| -0.958 | [-1.135;-0.36 | benign supporting |
| 0.000 | [0.000;0.000 | indeterminate |
| -0.967 | [-1.169;-0.83 | benign supporting |
| -1.103 | [-1.111;-1.09 | benign supporting |
| -1.134 | [-1.169;-1.08 | benign supporting |
| -1.083 | [-1.114;-1.00 | benign supporting |
| 0.892 | [0.707;1.073 | pathogenic moderate |
| 1.276 | [1.229;1.333 | pathogenic strong |
| 1.276 | [0.404;5.104 | pathogenic strong |
| 1.221 | [1.196;1.245 | pathogenic moderate |
| -0.974 | [-1.001;-0.94 | benign supporting |
| 0.836 | [-0.611;4.973 | pathogenic moderate |
| 1.265 | [1.221;1.313 | pathogenic moderate |
| 0.373 | [-0.283;1.067 | pathogenic supporting |
| 1.276 | [1.227;1.333 | pathogenic strong |
| 0.316 | [-0.109;0.739 | indeterminate |
| 1.276 | [1.241;1.316 | pathogenic strong |
| -1.167 | [-1.169;-1.16 | benign supporting |
| 1.030 | [1.002;1.059 | pathogenic moderate |
| 1.276 | [1.003;2.725 | pathogenic strong |
| 1.256 | [1.224;1.291 | pathogenic moderate |
| 1.276 | [1.247;1.308 | pathogenic strong |
| 1.276 | [1.011;2.022 | pathogenic strong |
| 1.276 | [1.219;1.347 | pathogenic strong |

| | | |
|---|---|---|
| -1.089 | [-1.092;-1.08 | benign supporting |
| -1.085 | [-1.087;-1.08 | benign supporting |
| -1.086 | [-1.096;-1.08 | benign supporting |
| 0.402 | [0.317;0.482 | pathogenic supporting |
| 1.149 | [1.127;1.172 | pathogenic moderate |
| 1.276 | [1.231;1.329 | pathogenic strong |
| 1.276 | [1.118;1.596 | pathogenic strong |
| -1.107 | [-1.114;-1.09 | benign supporting |
| -0.948 | [-0.961;-0.93 | benign supporting |
| 1.276 | [1.233;1.327 | pathogenic strong |
| -0.710 | [-1.037;-0.46 | benign supporting |
| 1.241 | [-0.205;5.105 | pathogenic moderate |
| 1.276 | [1.250;1.305 | pathogenic strong |
| -0.241 | [-0.278;-0.19 | indeterminate |
| -0.768 | [-0.839;-0.69 | benign supporting |
| -0.447 | [-0.807;-0.10 | benign supporting |
| -0.390 | [-0.426;-0.35 | benign supporting |
| 1.276 | [1.245;1.311 | pathogenic strong |
| -0.306 | [-0.639;0.645 | indeterminate |
| 1.015 | [0.966;1.056 | pathogenic moderate |
| 1.276 | [1.228;1.335 | pathogenic strong |
| -0.073 | [-0.261;0.412 | indeterminate |
| -1.081 | [-1.097;-1.06 | benign supporting |
| -1.101 | [-1.113;-1.09 | benign supporting |
| -0.868 | [-0.918;-0.81 | benign supporting |
| -0.919 | [-0.923;-0.91 | benign supporting |
| -1.164 | [-1.166;-1.16 | benign supporting |
| -0.283 | [-0.297;-0.27 | indeterminate |
| -0.459 | [-0.591;-0.35 | benign supporting |
| 0.725 | [0.073;1.108 | pathogenic moderate |
| -1.141 | [-1.166;-1.11 | benign supporting |
| -0.550 | [-0.561;-0.53 | benign supporting |
| -0.772 | [-0.837;-0.70 | benign supporting |
| -0.697 | [-0.913;-0.56 | benign supporting |
| -1.102 | [-1.108;-1.09 | benign supporting |
| -1.088 | [-1.088;-1.08 | benign supporting |
| -1.164 | [-1.165;-1.16 | benign supporting |
| 1.276 | [1.147;1.487 | pathogenic strong |
| 1.141 | [0.498;1.467 | pathogenic moderate |
| -0.849 | [-0.984;-0.68 | benign supporting |
| -0.502 | [-0.517;-0.48 | benign supporting |
| -1.131 | [-1.145;-1.11 | benign supporting |
| -0.812 | [-0.872;-0.73 | benign supporting |

| | | |
|---|---|---|
| -1.083 | [-1.084;-1.08 | benign supporting |
| 1.276 | [1.272;1.281 | pathogenic strong |
| 1.276 | [1.273;1.279 | pathogenic strong |
| 1.276 | [1.269;1.283 | pathogenic strong |
| 1.276 | [1.268;1.285 | pathogenic strong |
| -0.953 | [-0.954;-0.95 | benign supporting |
| 1.276 | [1.264;1.289 | pathogenic strong |
| 1.276 | [1.273;1.280 | pathogenic strong |
| 1.276 | [1.272;1.280 | pathogenic strong |
| 1.276 | [1.269;1.283 | pathogenic strong |
| 1.276 | [1.274;1.278 | pathogenic strong |
| 1.276 | [1.272;1.281 | pathogenic strong |
| -0.424 | [-0.425;-0.42 | benign supporting |
| 1.276 | [1.270;1.282 | pathogenic strong |
| 1.276 | [1.266;1.287 | pathogenic strong |
| 1.276 | [1.263;1.290 | pathogenic strong |
| 1.276 | [1.268;1.285 | pathogenic strong |
| 1.276 | [1.272;1.280 | pathogenic strong |
| 1.276 | [1.273;1.279 | pathogenic strong |
| 0.835 | [0.829;0.841 | pathogenic moderate |
| -0.495 | [-0.506;-0.48 | benign supporting |
| -1.077 | [-1.114;-0.87 | benign supporting |
| -0.336 | [-1.069;1.129 | benign supporting |
| 1.016 | [-0.773;5.053 | pathogenic moderate |
| 0.850 | [0.718;0.997 | pathogenic moderate |
| 1.131 | [1.106;1.157 | pathogenic moderate |
| -0.913 | [-0.918;-0.90 | benign supporting |
| 1.181 | [1.085;1.277 | pathogenic moderate |
| 1.276 | [1.211;1.360 | pathogenic strong |
| 0.608 | [-0.278;1.186 | pathogenic supporting |
| 1.276 | [1.213;1.357 | pathogenic strong |
| -0.408 | [-0.456;-0.36 | benign supporting |
| -0.680 | [-0.708;-0.65 | benign supporting |
| -1.061 | [-1.065;-1.05 | benign supporting |
| -0.736 | [-1.158;0.548 | benign supporting |
| -1.103 | [-1.168;-0.89 | benign supporting |
| 0.794 | [-0.315;1.364 | pathogenic moderate |
| 1.234 | [-0.115;5.094 | pathogenic moderate |
| -0.892 | [-0.914;-0.87 | benign supporting |
| 1.276 | [1.181;1.412 | pathogenic strong |
| 1.167 | [0.226;2.151 | pathogenic moderate |
| -0.994 | [-1.167;-0.88 | benign supporting |
| -0.941 | [-0.972;-0.91 | benign supporting |

| | | |
|---|---|---|
| -0.734 | [-1.074;-0.44 | benign supporting |
| 1.276 | [1.033;1.962 | pathogenic strong |
| -1.126 | [-1.155;-1.09 | benign supporting |
| 1.276 | [1.231;1.330 | pathogenic strong |
| -1.110 | [-1.167;-0.49 | benign supporting |
| -0.866 | [-0.918;-0.83 | benign supporting |
| -1.040 | [-1.047;-1.03 | benign supporting |
| 1.276 | [0.000;5.104 | pathogenic strong |
| -1.098 | [-1.115;-1.08 | benign supporting |
| -1.082 | [-1.167;0.10 | benign supporting |
| -1.096 | [-3.546;0.00 | benign supporting |
| 0.706 | [0.438;0.909 | pathogenic moderate |
| 0.043 | [-0.070;0.31 | indeterminate |
| -0.952 | [-0.988;-0.92 | benign supporting |
| -0.906 | [-0.914;-0.89 | benign supporting |
| 1.276 | [1.187;1.401 | pathogenic strong |
| 0.000 | [0.000;0.000 | indeterminate |
| -1.086 | [-1.089;-1.08 | benign supporting |
| 1.276 | [1.223;1.340 | pathogenic strong |
| -1.011 | [-1.036;-0.98 | benign supporting |
| 1.276 | [1.224;1.341 | pathogenic strong |
| 1.276 | [1.231;1.331 | pathogenic strong |
| -1.084 | [-1.086;-1.08 | benign supporting |
| -0.914 | [-1.040;-0.73 | benign supporting |
| -0.346 | [-1.086;1.18 | benign supporting |
| -0.381 | [-0.414;-0.34 | benign supporting |
| 1.276 | [0.420;5.101 | pathogenic strong |
| -0.474 | [-0.516;-0.43 | benign supporting |
| 0.118 | [-0.070;0.40 | indeterminate |
| -0.837 | [-0.842;-0.83 | benign supporting |
| -1.092 | [-1.112;-1.08 | benign supporting |
| -1.139 | [-1.159;-1.12 | benign supporting |
| 1.142 | [0.641;1.378 | pathogenic moderate |
| 1.276 | [0.821;5.062 | pathogenic strong |
| -0.521 | [-0.556;-0.48 | benign supporting |
| 1.107 | [-0.433;5.04 | pathogenic moderate |
| 1.276 | [1.216;1.351 | pathogenic strong |
| 1.276 | [1.183;1.411 | pathogenic strong |
| 1.203 | [0.681;1.820 | pathogenic moderate |
| -0.056 | [-0.578;1.15 | indeterminate |
| -1.116 | [-1.116;-1.10 | benign supporting |
| 1.276 | [0.756;5.087 | pathogenic strong |
| -0.785 | [-1.112;-0.25 | benign supporting |

| | | |
|---|---|---|
| -1.072 | [-1.076;-1.06 | benign supporting |
| 1.276 | [0.541;5.101 | pathogenic strong |
| -0.669 | [-0.723;-0.62 | benign supporting |
| 1.276 | [1.009;2.123 | pathogenic strong |
| 1.276 | [0.000;5.109 | pathogenic strong |
| 1.276 | [0.455;5.102 | pathogenic strong |
| -0.856 | [-1.000;-0.67 | benign supporting |
| 1.276 | [-0.356;5.101 | pathogenic strong |
| -0.472 | [-0.623;-0.34 | benign supporting |
| -1.158 | [-1.169;-1.14 | benign supporting |
| -0.374 | [-0.382;-0.36 | benign supporting |
| -1.012 | [-1.036;-0.98 | benign supporting |
| -0.589 | [-0.856;-0.40 | benign supporting |
| -0.186 | [-0.227;-0.14 | indeterminate |
| 1.276 | [0.841;5.059 | pathogenic strong |
| -1.107 | [-1.111;-1.10 | benign supporting |
| -0.968 | [-1.000;-0.93 | benign supporting |
| -0.365 | [-0.400;-0.32 | benign supporting |
| 1.097 | [0.702;1.234 | pathogenic moderate |
| 0.021 | [-0.466;1.140 | indeterminate |
| 0.138 | [-0.063;0.419 | indeterminate |
| -1.097 | [-1.115;-1.08 | benign supporting |
| -0.557 | [-0.564;-0.54 | benign supporting |
| -0.424 | [-0.495;-0.36 | benign supporting |
| -0.933 | [-1.107;-0.57 | benign supporting |
| -1.086 | [-1.090;-1.08 | benign supporting |
| 1.276 | [1.255;1.299 | pathogenic strong |
| 1.276 | [1.255;1.299 | pathogenic strong |
| 0.368 | [0.330;0.406 | pathogenic supporting |
| -0.537 | [-0.559;-0.51 | benign supporting |
| 1.271 | [1.251;1.293 | pathogenic moderate |
| -0.056 | [-0.100;0.003 | indeterminate |
| -1.005 | [-1.017;-0.99 | benign supporting |
| -0.326 | [-0.341;-0.30 | benign supporting |
| -0.972 | [-1.169;-0.83 | benign supporting |
| 1.276 | [1.221;1.342 | pathogenic strong |
| -0.518 | [-0.561;-0.47 | benign supporting |
| 0.972 | [0.895;1.034 | pathogenic moderate |
| -0.461 | [-0.516;-0.41 | benign supporting |
| 1.223 | [1.211;1.236 | pathogenic moderate |
| -1.096 | [-1.098;-1.09 | benign supporting |
| -0.901 | [-0.924;-0.87 | benign supporting |
| -1.085 | [-1.094;-1.08 | benign supporting |

| | | |
|---|---|---|
| -0.566 | [-0.960;-0.27 | benign supporting |
| -1.103 | [-1.124;-1.09 | benign supporting |
| -1.125 | [-1.133;-1.11 | benign supporting |
| -1.165 | [-1.167;-1.16 | benign supporting |
| -0.262 | [-0.283;-0.24 | indeterminate |
| -0.361 | [-0.383;-0.33 | benign supporting |
| -0.414 | [-0.432;-0.39 | benign supporting |
| -0.832 | [-0.851;-0.80 | benign supporting |
| -1.089 | [-1.091;-1.08 | benign supporting |
| 1.276 | [1.244;1.312 | pathogenic strong |
| 0.396 | [0.201;0.557 | pathogenic supporting |
| -1.126 | [-1.147;-1.10 | benign supporting |
| -0.925 | [-0.934;-0.91 | benign supporting |
| 0.411 | [-0.415;1.224 | pathogenic supporting |
| -0.242 | [-0.381;-0.02 | indeterminate |
| 1.276 | [0.818;5.070 | pathogenic strong |
| -0.759 | [-1.147;0.583 | benign supporting |
| -0.947 | [-0.952;-0.94 | benign supporting |
| 1.276 | [1.193;1.388 | pathogenic strong |
| -0.950 | [-0.963;-0.93 | benign supporting |
| 0.707 | [-0.041;1.123 | pathogenic moderate |
| 1.276 | [0.280;5.102 | pathogenic strong |
| -0.351 | [-0.494;-0.14 | benign supporting |
| 1.276 | [-0.326;5.106 | pathogenic strong |
| -1.100 | [-1.127;-1.08 | benign supporting |
| -0.919 | [-0.924;-0.91 | benign supporting |
| -1.013 | [-1.013;-1.01 | benign supporting |
| -1.129 | [-1.136;-1.12 | benign supporting |
| 1.276 | [0.257;5.110 | pathogenic strong |
| -1.029 | [-1.070;-0.97 | benign supporting |
| -0.585 | [-0.613;-0.55 | benign supporting |
| 1.276 | [1.146;1.501 | pathogenic strong |
| -1.145 | [-1.157;-1.13 | benign supporting |
| -0.712 | [-0.809;-0.65 | benign supporting |
| -0.359 | [-1.089;1.190 | benign supporting |
| 1.189 | [1.138;1.244 | pathogenic moderate |
| -1.094 | [-1.167;-0.69 | benign supporting |
| 0.123 | [0.009;0.254 | indeterminate |
| -1.070 | [-1.166;-0.09 | benign supporting |
| 0.033 | [-0.194;0.599 | indeterminate |
| -1.045 | [-3.555;0.000 | benign supporting |
| -1.091 | [-1.168;-0.40 | benign supporting |
| 1.276 | [0.912;4.956 | pathogenic strong |

| | | |
|---|---|---|
| -0.873 | [-0.922;-0.83 | benign supporting |
| -1.112 | [-1.168;-0.96 | benign supporting |
| -0.910 | [-0.970;-0.83 | benign supporting |
| -1.166 | [-1.169;-1.16 | benign supporting |
| -0.945 | [-0.982;-0.91 | benign supporting |
| 1.276 | [-0.163;5.10 | pathogenic strong |
| -1.000 | [-1.013;-0.98 | benign supporting |
| -0.853 | [-1.109;-0.40 | benign supporting |
| -1.060 | [-1.067;-1.04 | benign supporting |
| -0.239 | [-0.266;-0.20 | indeterminate |
| 1.276 | [1.251;1.304 | pathogenic strong |
| 1.276 | [1.222;1.340 | pathogenic strong |
| 1.276 | [1.251;1.304 | pathogenic strong |
| -0.487 | [-0.498;-0.47 | benign supporting |
| 1.276 | [1.215;1.351 | pathogenic strong |
| 1.222 | [1.189;1.255 | pathogenic moderate |
| 1.276 | [1.260;1.294 | pathogenic strong |
| 1.276 | [1.234;1.324 | pathogenic strong |
| -0.804 | [-0.827;-0.78 | benign supporting |
| 1.276 | [1.241;1.317 | pathogenic strong |
| -0.594 | [-0.608;-0.58 | benign supporting |
| -0.638 | [-0.648;-0.63 | benign supporting |
| 1.276 | [1.253;1.301 | pathogenic strong |
| 1.276 | [1.255;1.298 | pathogenic strong |
| 1.131 | [1.092;1.164 | pathogenic moderate |
| 0.786 | [0.727;0.848 | pathogenic moderate |
| 0.939 | [0.894;0.986 | pathogenic moderate |
| -0.365 | [-0.376;-0.35 | benign supporting |
| -1.085 | [-1.088;-1.08 | benign supporting |
| -1.111 | [-1.167;-1.08 | benign supporting |
| -0.353 | [-0.603;0.05 | benign supporting |
| 1.193 | [-1.111;5.04 | pathogenic moderate |
| -0.538 | [-1.069;-0.03 | benign supporting |
| 1.276 | [-0.389;5.10 | pathogenic strong |
| -1.094 | [-1.107;-1.08 | benign supporting |
| 1.276 | [-1.115;5.01 | pathogenic strong |
| -1.077 | [-1.165;-0.53 | benign supporting |
| -0.935 | [-0.984;-0.88 | benign supporting |
| -0.366 | [-0.509;-0.17 | benign supporting |
| -0.439 | [-0.507;-0.38 | benign supporting |
| -0.987 | [-1.014;-0.95 | benign supporting |
| 0.747 | [-0.282;1.27 | pathogenic moderate |
| -1.049 | [-1.113;-0.70 | benign supporting |

| | | |
|---|---|---|
| -1.065 | [-1.081;-1.02 | benign supporting |
| 0.826 | [-1.144;5.026 | pathogenic moderate |
| -1.100 | [-1.115;-1.08 | benign supporting |
| -0.499 | [-0.532;-0.46 | benign supporting |
| -3.812 | [-3.827;-0.86 | benign strong |
| -1.086 | [-1.115;-1.08 | benign supporting |
| -1.056 | [-1.114;-0.63 | benign supporting |
| -3.228 | [-3.515;-1.25 | benign strong |
| -0.677 | [-1.027;-0.43 | benign supporting |
| -0.538 | [-0.586;-0.48 | benign supporting |
| -1.104 | [-1.115;-1.06 | benign supporting |
| -0.562 | [-0.603;-0.51 | benign supporting |
| 1.276 | [0.000;5.100] | pathogenic strong |
| -1.094 | [-1.168;-1.01 | benign supporting |
| -1.006 | [-1.033;-0.97 | benign supporting |
| -0.987 | [-1.015;-0.95 | benign supporting |
| -1.105 | [-1.115;-1.08 | benign supporting |
| -0.610 | [-0.646;-0.56 | benign supporting |
| -0.751 | [-0.789;-0.71 | benign supporting |
| -1.115 | [-1.115;-1.06 | benign supporting |
| -0.300 | [-0.470;0.003 | indeterminate |
| 1.276 | [-1.089;5.075 | pathogenic strong |
| -1.082 | [-1.166;-0.36 | benign supporting |
| -1.019 | [-1.047;-0.99 | benign supporting |
| -1.104 | [-1.123;-1.09 | benign supporting |
| 0.323 | [-0.348;1.111 | pathogenic supporting |
| -1.052 | [-1.067;-1.03 | benign supporting |
| -0.959 | [-1.137;-0.41 | benign supporting |
| -1.077 | [-1.114;-0.96 | benign supporting |
| -0.166 | [-0.228;-0.10 | indeterminate |
| -0.653 | [-0.702;-0.61 | benign supporting |
| -0.759 | [-0.846;-0.68 | benign supporting |
| -0.098 | [-0.175;-0.01 | indeterminate |
| -0.401 | [-0.443;-0.36 | benign supporting |
| -0.313 | [-0.372;-0.23 | indeterminate |
| 0.604 | [-0.182;1.138 | pathogenic supporting |
| -1.097 | [-1.114;-1.08 | benign supporting |
| -1.087 | [-1.092;-1.08 | benign supporting |
| -1.092 | [-1.109;-1.08 | benign supporting |
| -0.054 | [-0.245;0.413 | indeterminate |
| -0.579 | [-0.603;-0.55 | benign supporting |
| -0.836 | [-0.895;-0.75 | benign supporting |
| 0.382 | [0.112;0.583] | pathogenic supporting |

| | | |
|---|---|---|
| -0.057 | [-0.148;0.140 | indeterminate |
| -0.449 | [-0.499;-0.40 | benign supporting |
| -0.973 | [-1.015;-0.92 | benign supporting |
| -1.069 | [-1.108;-0.97 | benign supporting |
| -1.084 | [-1.097;-1.08 | benign supporting |
| -0.185 | [-0.253;-0.11 | indeterminate |
| -1.065 | [-1.072;-1.04 | benign supporting |
| -1.077 | [-1.092;-1.06 | benign supporting |
| -1.071 | [-1.084;-1.05 | benign supporting |
| -1.166 | [-1.169;-1.16 | benign supporting |
| -0.882 | [-0.913;-0.85 | benign supporting |
| 0.024 | [-0.041;0.187 | indeterminate |
| -0.786 | [-1.065;-0.52 | benign supporting |
| -0.200 | [-0.292;-0.09 | indeterminate |
| -1.113 | [-1.121;-1.10 | benign supporting |
| -1.112 | [-1.115;-1.10 | benign supporting |
| -0.392 | [-0.410;-0.37 | benign supporting |
| -0.806 | [-0.849;-0.75 | benign supporting |
| -0.968 | [-1.087;-0.74 | benign supporting |
| 1.276 | [1.237;1.320 | pathogenic strong |
| -0.920 | [-0.992;-0.84 | benign supporting |
| -0.802 | [-0.824;-0.78 | benign supporting |
| -0.420 | [-0.461;-0.38 | benign supporting |
| -1.085 | [-1.089;-1.08 | benign supporting |
| -1.069 | [-1.115;-0.68 | benign supporting |
| 0.992 | [0.841;1.100 | pathogenic moderate |
| 1.276 | [1.191;1.398 | pathogenic strong |
| 1.276 | [1.188;1.395 | pathogenic strong |
| 1.132 | [1.103;1.162 | pathogenic moderate |
| -0.575 | [-0.600;-0.55 | benign supporting |
| 1.276 | [1.226;1.338 | pathogenic strong |
| 1.226 | [1.182;1.274 | pathogenic moderate |
| -0.455 | [-0.492;-0.42 | benign supporting |
| -0.977 | [-1.016;-0.93 | benign supporting |
| -1.093 | [-1.097;-1.08 | benign supporting |
| 1.276 | [1.241;1.317 | pathogenic strong |
| -0.499 | [-0.512;-0.48 | benign supporting |
| -1.110 | [-1.113;-1.10 | benign supporting |
| -0.360 | [-0.413;-0.29 | benign supporting |
| 1.220 | [1.186;1.258 | pathogenic moderate |
| 1.276 | [1.241;1.315 | pathogenic strong |
| -0.812 | [-0.872;-0.73 | benign supporting |
| -0.665 | [-0.688;-0.64 | benign supporting |

| | | |
|---|---|---|
| 1.142 | [1.084;1.193 | pathogenic moderate |
| -0.560 | [-0.579;-0.53 | benign supporting |
| -1.129 | [-1.161;-1.10 | benign supporting |
| -0.311 | [-0.396;-0.18 | indeterminate |
| 1.276 | [1.233;1.325 | pathogenic strong |
| 1.276 | [1.162;1.455 | pathogenic strong |
| 1.276 | [1.178;1.420 | pathogenic strong |
| -0.998 | [-1.010;-0.98 | benign supporting |
| -0.211 | [-0.237;-0.18 | indeterminate |
| 1.246 | [1.225;1.267 | pathogenic moderate |
| 1.276 | [1.223;1.340 | pathogenic strong |
| 1.276 | [1.244;1.313 | pathogenic strong |
| -0.406 | [-0.435;-0.37 | benign supporting |
| 1.184 | [1.124;1.249 | pathogenic moderate |
| -0.969 | [-0.984;-0.95 | benign supporting |
| 0.935 | [0.883;0.990 | pathogenic moderate |
| 1.276 | [1.119;1.623 | pathogenic strong |
| 1.276 | [1.259;1.294 | pathogenic strong |
| 0.535 | [0.354;0.682 | pathogenic supporting |
| -0.106 | [-0.179;-0.03 | indeterminate |
| 1.276 | [1.244;1.312 | pathogenic strong |
| 1.276 | [1.213;1.356 | pathogenic strong |
| -0.819 | [-0.831;-0.80 | benign supporting |
| -1.087 | [-1.093;-1.08 | benign supporting |
| -0.843 | [-0.885;-0.79 | benign supporting |
| 1.276 | [1.234;1.325 | pathogenic strong |
| -1.113 | [-1.115;-1.00 | benign supporting |
| 1.276 | [1.103;1.678 | pathogenic strong |
| -1.110 | [-1.115;-1.09 | benign supporting |
| 1.102 | [0.662;1.256 | pathogenic moderate |
| 1.276 | [1.112;1.624 | pathogenic strong |
| -0.194 | [-0.262;-0.11 | indeterminate |
| 0.572 | [0.484;0.647 | pathogenic supporting |
| -1.081 | [-1.091;-1.07 | benign supporting |
| 1.239 | [1.198;1.282 | pathogenic moderate |
| -1.092 | [-1.097;-1.08 | benign supporting |
| -0.848 | [-0.885;-0.81 | benign supporting |
| 1.276 | [0.402;5.094 | pathogenic strong |
| 0.808 | [0.763;0.855 | pathogenic moderate |
| 0.209 | [-0.009;0.417 | indeterminate |
| -0.467 | [-0.905;-0.08 | benign supporting |
| 1.276 | [1.168;1.431 | pathogenic strong |
| 1.276 | [1.234;1.326 | pathogenic strong |

| | | |
|---|---|---|
| -0.355 | [-0.927;0.843 | benign supporting |
| -1.088 | [-1.096;-1.08 | benign supporting |
| -0.937 | [-0.956;-0.92 | benign supporting |
| 1.276 | [1.211;1.363 | pathogenic strong |
| 0.903 | [0.841;0.969 | pathogenic moderate |
| 1.276 | [1.241;1.316 | pathogenic strong |
| -0.370 | [-0.407;-0.32 | benign supporting |
| 1.276 | [1.237;1.322 | pathogenic strong |
| 1.103 | [1.040;1.127 | pathogenic moderate |
| 1.276 | [1.249;1.305 | pathogenic strong |
| 1.276 | [1.232;1.326 | pathogenic strong |
| -0.526 | [-0.544;-0.50 | benign supporting |
| 0.932 | [0.860;1.008 | pathogenic moderate |
| -0.548 | [-0.585;-0.50 | benign supporting |
| -0.853 | [-0.879;-0.82 | benign supporting |
| 1.276 | [1.203;1.375 | pathogenic strong |
| 0.017 | [-0.236;0.625 | indeterminate |
| -0.079 | [-0.278;0.406 | indeterminate |
| -1.018 | [-1.034;-1.00 | benign supporting |
| 1.276 | [1.208;1.364 | pathogenic strong |
| 1.276 | [1.227;1.336 | pathogenic strong |
| -1.111 | [-1.115;-1.08 | benign supporting |
| -1.090 | [-1.136;-1.08 | benign supporting |
| -0.365 | [-0.416;-0.30 | benign supporting |
| -0.971 | [-1.017;-0.92 | benign supporting |
| -1.086 | [-1.090;-1.08 | benign supporting |
| -0.519 | [-0.559;-0.48 | benign supporting |
| -1.025 | [-1.038;-1.01 | benign supporting |
| -0.940 | [-1.021;-0.84 | benign supporting |
| -1.090 | [-1.109;-1.08 | benign supporting |
| -0.559 | [-0.574;-0.54 | benign supporting |
| -0.588 | [-0.613;-0.56 | benign supporting |
| -1.080 | [-1.111;-1.03 | benign supporting |
| -0.968 | [-0.990;-0.94 | benign supporting |
| -1.094 | [-1.104;-1.08 | benign supporting |
| -1.102 | [-1.168;-1.08 | benign supporting |
| 1.180 | [1.151;1.211 | pathogenic moderate |
| -1.099 | [-1.103;-1.09 | benign supporting |
| -0.036 | [-0.076;0.015 | indeterminate |
| -0.234 | [-0.259;-0.20 | indeterminate |
| -0.718 | [-0.751;-0.69 | benign supporting |
| -1.167 | [-1.169;-1.11 | benign supporting |
| -1.059 | [-1.114;-0.66 | benign supporting |

| | | |
|---|---|---|
| -0.539 | [-0.620;-0.45 | benign supporting |
| -1.104 | [-1.169;-1.08 | benign supporting |
| -0.563 | [-0.593;-0.52 | benign supporting |
| -0.628 | [-0.657;-0.60 | benign supporting |
| -0.063 | [-0.208;0.295 | indeterminate |
| -0.923 | [-0.939;-0.90 | benign supporting |
| -0.078 | [-0.428;0.867 | indeterminate |
| -1.062 | [-1.106;-0.94 | benign supporting |
| -1.085 | [-1.107;-1.06 | benign supporting |
| -1.061 | [-1.110;-0.91 | benign supporting |
| -1.074 | [-1.099;-1.05 | benign supporting |
| -0.352 | [-0.426;-0.25 | benign supporting |
| 1.276 | [0.860;5.024] | pathogenic strong |
| -0.744 | [-0.873;-0.63 | benign supporting |
| -0.277 | [-0.767;0.992 | indeterminate |
| -1.126 | [-1.169;-0.83 | benign supporting |
| -0.986 | [-1.025;-0.93 | benign supporting |
| -0.729 | [-0.787;-0.68 | benign supporting |
| -0.354 | [-0.403;-0.29 | benign supporting |
| -0.889 | [-0.910;-0.86 | benign supporting |
| -1.055 | [-1.061;-1.04 | benign supporting |
| -1.161 | [-1.169;-1.05 | benign supporting |
| -1.022 | [-1.033;-1.01 | benign supporting |
| -1.150 | [-1.169;-1.13 | benign supporting |
| -1.076 | [-1.081;-1.07 | benign supporting |
| -1.087 | [-1.089;-1.08 | benign supporting |
| -0.889 | [-0.914;-0.86 | benign supporting |
| -0.985 | [-0.999;-0.97 | benign supporting |
| -0.892 | [-0.919;-0.86 | benign supporting |
| -1.093 | [-1.098;-1.09 | benign supporting |
| -1.119 | [-1.132;-1.10 | benign supporting |
| -1.072 | [-1.079;-1.06 | benign supporting |
| -1.106 | [-1.110;-1.10 | benign supporting |
| -0.058 | [-0.094;-0.01 | indeterminate |
| -0.853 | [-0.866;-0.84 | benign supporting |
| -1.108 | [-1.112;-1.10 | benign supporting |
| -1.069 | [-1.073;-1.06 | benign supporting |
| -0.697 | [-0.731;-0.67 | benign supporting |
| -1.126 | [-1.169;-1.01 | benign supporting |
| -1.067 | [-1.076;-1.04 | benign supporting |
| -0.981 | [-1.085;-0.80 | benign supporting |
| -1.057 | [-1.108;-0.92 | benign supporting |
| -1.084 | [-1.086;-1.08 | benign supporting |

| | | |
|---|---|---|
| -0.895 | [-0.938;-0.84 | benign supporting |
| -1.090 | [-1.092;-1.08 | benign supporting |
| -1.062 | [-1.084;-1.00 | benign supporting |
| -0.477 | [-0.497;-0.45 | benign supporting |
| -0.455 | [-0.467;-0.44 | benign supporting |
| -1.009 | [-1.023;-0.99 | benign supporting |
| -0.901 | [-0.915;-0.88 | benign supporting |
| -0.835 | [-0.861;-0.80 | benign supporting |
| -1.080 | [-1.114;-0.95 | benign supporting |
| -1.105 | [-1.108;-1.10 | benign supporting |
| 0.480 | [0.404;0.555] | pathogenic supporting |
| -0.865 | [-0.892;-0.84 | benign supporting |
| -1.164 | [-1.166;-1.16 | benign supporting |
| -0.632 | [-0.642;-0.62 | benign supporting |
| -0.819 | [-1.066;-0.56 | benign supporting |
| -0.955 | [-0.975;-0.93 | benign supporting |
| -1.090 | [-1.092;-1.08 | benign supporting |
| -0.964 | [-1.046;-0.87 | benign supporting |
| -1.090 | [-1.114;-1.08 | benign supporting |
| -0.557 | [-0.581;-0.52 | benign supporting |
| -1.017 | [-1.034;-1.00 | benign supporting |
| -1.099 | [-1.110;-1.09 | benign supporting |
| -0.311 | [-0.361;-0.24 | indeterminate |
| -1.076 | [-1.090;-1.05 | benign supporting |
| -1.089 | [-1.093;-1.08 | benign supporting |
| -0.863 | [-0.903;-0.82 | benign supporting |
| -1.092 | [-1.096;-1.08 | benign supporting |
| -1.097 | [-1.106;-1.08 | benign supporting |
| -0.900 | [-0.925;-0.87 | benign supporting |
| -0.530 | [-0.648;-0.41 | benign supporting |
| -1.002 | [-1.027;-0.96 | benign supporting |
| 0.466 | [0.337;0.585] | pathogenic supporting |
| -0.390 | [-0.558;-0.18 | benign supporting |
| 0.396 | [0.306;0.479] | pathogenic supporting |
| -1.088 | [-1.096;-1.08 | benign supporting |
| -0.923 | [-0.942;-0.90 | benign supporting |
| -0.901 | [-0.935;-0.85 | benign supporting |
| -0.601 | [-0.910;-0.39 | benign supporting |
| -1.043 | [-1.066;-1.01 | benign supporting |
| -0.474 | [-0.507;-0.44 | benign supporting |
| -0.652 | [-0.670;-0.63 | benign supporting |
| -1.084 | [-1.087;-1.08 | benign supporting |
| -0.618 | [-0.677;-0.56 | benign supporting |

| | | |
|---|---|---|
| -0.455 | [-1.005;0.287 | benign supporting |
| -0.624 | [-0.638;-0.61 | benign supporting |
| -1.061 | [-1.068;-1.04 | benign supporting |
| -1.093 | [-1.103;-1.08 | benign supporting |
| -1.097 | [-1.103;-1.09 | benign supporting |
| -1.101 | [-1.106;-1.09 | benign supporting |
| 1.276 | [1.137;1.548 | pathogenic strong |
| -0.898 | [-0.932;-0.85 | benign supporting |
| -0.127 | [-0.195;-0.06 | indeterminate |
| -0.940 | [-0.955;-0.92 | benign supporting |
| -0.862 | [-0.896;-0.83 | benign supporting |
| -0.849 | [-0.861;-0.83 | benign supporting |
| -1.027 | [-1.040;-1.01 | benign supporting |
| -0.953 | [-0.958;-0.94 | benign supporting |
| -0.488 | [-0.566;-0.41 | benign supporting |
| -0.315 | [-0.334;-0.29 | indeterminate |
| -1.135 | [-1.149;-1.12 | benign supporting |
| 1.276 | [0.000;5.096 | pathogenic strong |
| -0.660 | [-0.669;-0.65 | benign supporting |
| -3.481 | [-3.497;-3.46 | benign strong |
| 1.060 | [1.012;1.105 | pathogenic moderate |
| 0.610 | [0.512;0.702 | pathogenic supporting |
| 0.225 | [-0.046;0.507 | indeterminate |
| -1.168 | [-1.169;-1.14 | benign supporting |
| -0.178 | [-0.201;-0.15 | indeterminate |
| 1.240 | [1.211;1.271 | pathogenic moderate |
| 1.276 | [-0.161;5.102 | pathogenic strong |
| -0.705 | [-0.791;-0.64 | benign supporting |
| -0.188 | [-0.619;1.005 | indeterminate |
| -0.386 | [-0.604;-0.09 | benign supporting |
| -0.299 | [-0.312;-0.28 | indeterminate |
| -1.008 | [-1.014;-1.00 | benign supporting |
| 1.276 | [1.234;1.326 | pathogenic strong |
| -0.915 | [-0.932;-0.89 | benign supporting |
| -0.829 | [-0.963;-0.65 | benign supporting |
| -1.164 | [-1.169;-1.12 | benign supporting |
| -0.151 | [-0.178;-0.12 | indeterminate |
| -0.671 | [-1.106;-0.16 | benign supporting |
| -0.465 | [-1.166;4.799 | benign supporting |
| 1.276 | [0.875;5.035 | pathogenic strong |
| -0.514 | [-0.899;-0.22 | benign supporting |
| -1.112 | [-1.116;-1.10 | benign supporting |
| -1.116 | [-1.169;-1.08 | benign supporting |

| | | |
|---|---|---|
| -0.392 | [-1.168;4.932 | benign supporting |
| -0.974 | [-1.152;-0.38 | benign supporting |
| -0.837 | [-0.845;-0.83 | benign supporting |
| 1.276 | [1.251;1.303] | pathogenic strong |
| -0.928 | [-0.933;-0.92 | benign supporting |
| -1.089 | [-1.093;-1.08 | benign supporting |
| -1.093 | [-1.114;-1.08 | benign supporting |
| -1.109 | [-1.139;-1.07 | benign supporting |
| -1.150 | [-1.152;-1.14 | benign supporting |
| -1.157 | [-1.161;-1.15 | benign supporting |
| -1.110 | [-1.111;-1.10 | benign supporting |
| -0.552 | [-0.555;-0.55 | benign supporting |
| -0.874 | [-0.885;-0.86 | benign supporting |
| -0.710 | [-0.721;-0.69 | benign supporting |
| -0.814 | [-0.823;-0.80 | benign supporting |
| -0.811 | [-0.814;-0.80 | benign supporting |
| -0.945 | [-0.949;-0.94 | benign supporting |
| -1.028 | [-1.029;-1.02 | benign supporting |
| -1.115 | [-1.115;-1.11 | benign supporting |
| -0.997 | [-1.002;-0.99 | benign supporting |
| -1.107 | [-1.110;-1.10 | benign supporting |
| 1.276 | [1.266;1.287] | pathogenic strong |
| -0.673 | [-0.901;-0.53 | benign supporting |
| -1.107 | [-1.108;-1.10 | benign supporting |
| -0.579 | [-0.581;-0.57 | benign supporting |
| -1.129 | [-1.168;-1.10 | benign supporting |
| -1.000 | [-1.007;-0.99 | benign supporting |
| -1.086 | [-1.168;-0.73 | benign supporting |
| 1.138 | [-0.107;4.346 | pathogenic moderate |
| -1.055 | [-1.095;-0.95 | benign supporting |
| 0.522 | [0.352;0.664] | pathogenic supporting |
| 0.449 | [-0.340;1.157 | pathogenic supporting |
| 0.647 | [-1.004;4.993 | pathogenic moderate |
| 1.276 | [-0.319;5.102 | pathogenic strong |
| 0.346 | [-0.242;0.990 | pathogenic supporting |
| -1.086 | [-1.100;-1.08 | benign supporting |
| -1.020 | [-1.116;-0.52 | benign supporting |
| -1.110 | [-1.115;-1.10 | benign supporting |
| -0.907 | [-0.911;-0.90 | benign supporting |
| 1.276 | [0.630;5.097] | pathogenic strong |
| 0.756 | [0.187;1.106] | pathogenic moderate |
| -0.953 | [-2.865;-0.83 | benign supporting |
| -1.166 | [-1.169;-1.16 | benign supporting |

| | | |
|---|---|---|
| 0.133 | [-0.123;0.564 | indeterminate |
| -0.510 | [-0.961;-0.12 | benign supporting |
| -0.986 | [-1.034;-0.95 | benign supporting |
| -0.974 | [-1.018;-0.95 | benign supporting |
| -1.091 | [-1.164;-0.63 | benign supporting |
| 1.276 | [1.133;1.540] | pathogenic strong |
| -0.681 | [-1.028;-0.44 | benign supporting |
| -1.092 | [-1.165;-1.02 | benign supporting |
| -1.085 | [-1.087;-1.08 | benign supporting |
| -1.129 | [-1.168;-1.08 | benign supporting |
| -1.016 | [-1.047;-0.98 | benign supporting |
| -1.164 | [-1.168;-0.92 | benign supporting |
| 0.003 | [-0.036;0.077 | indeterminate |
| -1.020 | [-1.030;-1.01 | benign supporting |
| 1.231 | [1.142;1.341] | pathogenic moderate |
| -1.131 | [-1.169;-1.09 | benign supporting |
| 0.928 | [0.002;1.266] | pathogenic moderate |
| 0.487 | [-0.237;1.108 | pathogenic supporting |
| -1.084 | [-1.086;-1.08 | benign supporting |
| 1.276 | [1.183;1.412] | pathogenic strong |
| 1.276 | [0.935;4.763] | pathogenic strong |
| -0.725 | [-0.752;-0.70 | benign supporting |
| -0.737 | [-0.791;-0.69 | benign supporting |
| -0.856 | [-0.903;-0.80 | benign supporting |
| -0.990 | [-1.075;-0.85 | benign supporting |
| -0.588 | [-0.608;-0.56 | benign supporting |
| -0.979 | [-1.092;-0.77 | benign supporting |
| -0.659 | [-0.793;-0.57 | benign supporting |
| -0.722 | [-0.803;-0.66 | benign supporting |
| -0.736 | [-0.762;-0.71 | benign supporting |
| -1.110 | [-1.115;-1.10 | benign supporting |
| -0.582 | [-0.629;-0.53 | benign supporting |
| -0.615 | [-0.621;-0.61 | benign supporting |
| -0.079 | [-0.131;-0.01 | indeterminate |
| 1.222 | [1.170;1.279] | pathogenic moderate |
| -0.903 | [-0.934;-0.86 | benign supporting |
| 1.103 | [-0.198;2.056 | pathogenic moderate |
| -1.112 | [-1.116;-1.10 | benign supporting |
| -0.624 | [-0.632;-0.61 | benign supporting |
| -0.509 | [-0.526;-0.49 | benign supporting |
| -1.003 | [-1.023;-0.98 | benign supporting |
| -0.619 | [-0.846;-0.45 | benign supporting |
| -1.052 | [-1.166;-0.21 | benign supporting |

| | | |
|---|---|---|
| -0.531 | [-0.890;-0.25 | benign supporting |
| -1.050 | [-1.065;-1.03 | benign supporting |
| -3.544 | [-3.768;-2.91 | benign strong |
| -0.262 | [-0.342;-0.14 | indeterminate |
| -0.953 | [-1.024;-0.90 | benign supporting |
| -0.566 | [-1.164;1.657 | benign supporting |
| 0.000 | [-3.061;0.000 | indeterminate |
| 1.276 | [1.018;2.049] | pathogenic strong |
| -1.068 | [-1.167;-0.99 | benign supporting |
| 0.559 | [-0.078;1.006 | pathogenic supporting |
| -0.905 | [-0.929;-0.87 | benign supporting |
| -0.414 | [-0.476;-0.35 | benign supporting |
| 1.276 | [1.137;1.530] | pathogenic strong |
| -0.960 | [-1.103;-0.66 | benign supporting |
| -0.829 | [-0.853;-0.79 | benign supporting |
| 1.276 | [-0.205;5.108 | pathogenic strong |
| -0.413 | [-0.490;-0.34 | benign supporting |
| -1.094 | [-1.108;-1.08 | benign supporting |
| -1.092 | [-1.167;-0.63 | benign supporting |
| -1.146 | [-1.169;-1.08 | benign supporting |
| -0.735 | [-0.844;-0.65 | benign supporting |
| -0.521 | [-0.662;-0.38 | benign supporting |
| -1.156 | [-1.169;-1.13 | benign supporting |
| -0.787 | [-0.850;-0.71 | benign supporting |
| -0.563 | [-0.583;-0.54 | benign supporting |
| -0.438 | [-0.483;-0.40 | benign supporting |
| -0.666 | [-1.077;-0.36 | benign supporting |
| -1.011 | [-1.027;-0.99 | benign supporting |
| -0.716 | [-0.812;-0.64 | benign supporting |
| -1.087 | [-1.091;-1.08 | benign supporting |
| -0.344 | [-0.378;-0.30 | benign supporting |
| -1.089 | [-1.096;-1.08 | benign supporting |
| -0.849 | [-0.890;-0.80 | benign supporting |
| -0.609 | [-0.682;-0.53 | benign supporting |
| -0.434 | [-0.899;0.119 | benign supporting |
| 1.276 | [-1.103;5.047 | pathogenic strong |
| -0.588 | [-0.628;-0.54 | benign supporting |
| -0.666 | [-0.686;-0.64 | benign supporting |
| -0.869 | [-0.896;-0.84 | benign supporting |
| -1.085 | [-1.168;-1.07 | benign supporting |
| -1.109 | [-1.115;-1.09 | benign supporting |
| -0.818 | [-0.840;-0.78 | benign supporting |
| 0.470 | [-0.076;0.876 | pathogenic supporting |

| | | |
|---|---|---|
| -1.089 | [-1.111;-1.08 | benign supporting |
| 0.327 | [0.242;0.399] | pathogenic supporting |
| 0.361 | [0.247;0.459] | pathogenic supporting |
| -0.951 | [-0.959;-0.94 | benign supporting |
| 1.276 | [0.608;5.087] | pathogenic strong |
| -0.666 | [-0.955;-0.46 | benign supporting |
| -0.946 | [-0.974;-0.92 | benign supporting |
| -0.391 | [-0.535;-0.23 | benign supporting |
| -0.306 | [-0.394;-0.17 | indeterminate |
| 1.252 | [1.220;1.287] | pathogenic moderate |
| -0.527 | [-0.552;-0.50 | benign supporting |
| -0.294 | [-0.326;-0.25 | indeterminate |
| -0.083 | [-0.411;0.848 | indeterminate |
| 1.276 | [1.119;1.594] | pathogenic strong |
| -0.455 | [-0.574;-0.36 | benign supporting |
| 1.276 | [1.097;1.708] | pathogenic strong |
| -0.391 | [-0.416;-0.36 | benign supporting |
| 1.035 | [0.979;1.090] | pathogenic moderate |
| -0.969 | [-0.987;-0.95 | benign supporting |
| -0.853 | [-0.932;-0.75 | benign supporting |
| -0.999 | [-1.072;-0.88 | benign supporting |
| -1.098 | [-1.167;-0.86 | benign supporting |
| -0.950 | [-0.965;-0.93 | benign supporting |
| -0.931 | [-0.950;-0.91 | benign supporting |
| -0.937 | [-0.956;-0.91 | benign supporting |
| -1.081 | [-1.095;-1.06 | benign supporting |
| -1.073 | [-1.085;-1.05 | benign supporting |
| -1.099 | [-1.113;-1.08 | benign supporting |
| -0.641 | [-0.667;-0.62 | benign supporting |
| -0.578 | [-0.599;-0.56 | benign supporting |
| -0.689 | [-0.741;-0.65 | benign supporting |
| 0.012 | [-0.043;0.141 | indeterminate |
| -0.573 | [-0.591;-0.55 | benign supporting |
| -1.102 | [-1.105;-1.09 | benign supporting |
| -0.290 | [-0.342;-0.22 | indeterminate |
| -0.793 | [-0.833;-0.74 | benign supporting |
| -1.068 | [-1.072;-1.06 | benign supporting |
| -1.114 | [-1.116;-1.10 | benign supporting |
| -0.256 | [-0.377;-0.06 | indeterminate |
| -1.089 | [-1.103;-1.08 | benign supporting |
| -1.082 | [-1.114;-0.98 | benign supporting |
| -1.112 | [-1.116;-1.10 | benign supporting |
| -1.089 | [-3.077;4.500 | benign supporting |

| | | |
|---|---|---|
| -1.087 | [-1.089;-1.08 | benign supporting |
| -1.071 | [-1.077;-1.06 | benign supporting |
| -1.164 | [-1.168;-1.16 | benign supporting |
| -0.829 | [-0.852;-0.79 | benign supporting |
| -0.425 | [-0.502;-0.36 | benign supporting |
| -0.479 | [-0.549;-0.41 | benign supporting |
| -1.098 | [-1.100;-1.09 | benign supporting |
| -1.042 | [-1.071;-1.00 | benign supporting |
| -1.100 | [-1.109;-1.09 | benign supporting |
| -0.617 | [-0.627;-0.60 | benign supporting |
| 1.276 | [1.227;1.335 | pathogenic strong |
| -1.084 | [-1.085;-1.08 | benign supporting |
| 0.502 | [0.091;0.774 | pathogenic supporting |
| -1.102 | [-1.111;-1.09 | benign supporting |
| -1.110 | [-1.116;-1.10 | benign supporting |
| -1.046 | [-1.053;-1.04 | benign supporting |
| -0.338 | [-0.370;-0.30 | benign supporting |
| -0.953 | [-1.052;-0.83 | benign supporting |
| 1.136 | [0.988;1.221 | pathogenic moderate |
| 1.276 | [1.235;1.326 | pathogenic strong |
| -0.274 | [-0.313;-0.23 | indeterminate |
| 1.276 | [1.217;1.347 | pathogenic strong |
| 0.248 | [0.109;0.357 | indeterminate |
| 1.276 | [1.241;1.315 | pathogenic strong |
| 1.041 | [-0.459;5.02( | pathogenic moderate |
| 1.276 | [1.225;1.337 | pathogenic strong |
| 1.276 | [1.200;1.379 | pathogenic strong |
| 0.797 | [0.694;0.906 | pathogenic moderate |
| 1.276 | [1.254;1.298 | pathogenic strong |
| 1.276 | [1.229;1.331 | pathogenic strong |
| 1.276 | [1.210;1.357 | pathogenic strong |
| 1.242 | [1.159;1.342 | pathogenic moderate |
| 1.276 | [1.233;1.327 | pathogenic strong |
| 1.276 | [1.153;1.482 | pathogenic strong |
| 1.276 | [1.196;1.383 | pathogenic strong |
| 1.276 | [1.219;1.347 | pathogenic strong |
| 1.276 | [1.214;1.355 | pathogenic strong |
| 1.276 | [1.256;1.297 | pathogenic strong |
| -1.105 | [-1.164;-0.76 | benign supporting |
| -0.045 | [-0.251;0.50( | indeterminate |
| -0.728 | [-0.752;-0.70 | benign supporting |
| -1.164 | [-1.169;-1.15 | benign supporting |
| -1.097 | [-1.108;-1.08 | benign supporting |

| | | |
|---|---|---|
| 0.741 | [0.697;0.784 | pathogenic moderate |
| -0.677 | [-0.837;-0.58 | benign supporting |
| -1.113 | [-1.116;-1.11 | benign supporting |
| -1.086 | [-1.102;-1.08 | benign supporting |
| -1.094 | [-1.115;-1.04 | benign supporting |
| 0.551 | [0.469;0.616 | pathogenic supporting |
| -1.168 | [-1.169;-1.16 | benign supporting |
| 1.276 | [1.263;1.290 | pathogenic strong |
| 0.271 | [-0.076;0.624 | indeterminate |
| 1.276 | [1.229;1.331 | pathogenic strong |
| -0.140 | [-0.178;-0.10 | indeterminate |
| 0.590 | [0.572;0.607 | pathogenic supporting |
| -1.095 | [-1.099;-1.09 | benign supporting |
| -0.999 | [-1.020;-0.96 | benign supporting |
| 1.276 | [1.016;2.113 | pathogenic strong |
| 1.276 | [1.207;1.367 | pathogenic strong |
| 1.276 | [1.192;1.390 | pathogenic strong |
| 1.276 | [1.114;1.614 | pathogenic strong |
| 1.276 | [0.000;5.103 | pathogenic strong |
| 1.248 | [1.171;1.340 | pathogenic moderate |
| 0.439 | [-0.420;1.248 | pathogenic supporting |
| -1.169 | [-1.169;-1.08 | benign supporting |
| -1.105 | [-1.111;-1.09 | benign supporting |
| -1.146 | [-1.169;-1.05 | benign supporting |
| -1.122 | [-1.164;-1.09 | benign supporting |
| 1.276 | [-1.098;5.041 | pathogenic strong |
| 0.000 | [0.000;0.000 | indeterminate |
| 0.466 | [-0.401;1.233 | pathogenic supporting |
| -0.709 | [-0.732;-0.68 | benign supporting |
| 1.276 | [1.240;1.319 | pathogenic strong |
| -0.963 | [-1.066;-0.84 | benign supporting |
| 1.276 | [1.241;1.317 | pathogenic strong |
| 1.276 | [1.156;1.467 | pathogenic strong |
| 1.004 | [-0.567;5.027 | pathogenic moderate |
| 1.171 | [1.139;1.205 | pathogenic moderate |
| -0.420 | [-0.884;0.212 | benign supporting |
| -0.469 | [-0.495;-0.44 | benign supporting |
| -1.043 | [-1.062;-1.02 | benign supporting |
| -0.251 | [-0.268;-0.23 | indeterminate |
| -0.636 | [-0.667;-0.61 | benign supporting |
| -0.901 | [-0.906;-0.89 | benign supporting |
| -1.095 | [-1.101;-1.09 | benign supporting |
| -1.049 | [-1.061;-1.03 | benign supporting |

| | | |
|---|---|---|
| -1.028 | [-1.059;-0.99 | benign supporting |
| -1.115 | [-1.116;-0.96 | benign supporting |
| -0.917 | [-1.166;1.046 | benign supporting |
| -1.084 | [-1.085;-1.08 | benign supporting |
| 1.276 | [1.121;1.590 | pathogenic strong |
| -1.101 | [-1.115;-1.07 | benign supporting |
| 1.276 | [1.253;1.302 | pathogenic strong |
| -0.236 | [-0.353;-0.06 | indeterminate |
| -0.113 | [-0.254;0.102 | indeterminate |
| -1.112 | [-1.132;-1.09 | benign supporting |
| -0.830 | [-1.160;0.149 | benign supporting |
| -1.109 | [-1.116;-1.09 | benign supporting |
| -1.129 | [-1.152;-1.11 | benign supporting |
| 1.276 | [1.252;1.303 | pathogenic strong |
| -0.610 | [-0.618;-0.60 | benign supporting |
| 1.138 | [1.126;1.149 | pathogenic moderate |
| 0.462 | [0.373;0.549 | pathogenic supporting |
| -0.354 | [-0.362;-0.34 | benign supporting |
| -1.116 | [-1.124;-1.10 | benign supporting |
| 1.199 | [1.146;1.257 | pathogenic moderate |
| -1.071 | [-2.993;1.507 | benign supporting |
| -1.090 | [-1.110;-1.08 | benign supporting |
| -1.081 | [-1.096;-1.06 | benign supporting |
| -0.368 | [-0.383;-0.35 | benign supporting |
| 1.276 | [1.228;1.333 | pathogenic strong |
| 0.978 | [0.935;1.011 | pathogenic moderate |
| 1.276 | [1.255;1.298 | pathogenic strong |
| -0.300 | [-0.516;0.197 | indeterminate |
| -0.734 | [-0.745;-0.72 | benign supporting |
| -0.950 | [-0.961;-0.94 | benign supporting |
| -1.085 | [-1.106;-1.08 | benign supporting |
| -1.074 | [-1.165;-0.48 | benign supporting |
| -0.592 | [-0.921;-0.37 | benign supporting |
| -0.319 | [-0.384;-0.23 | benign supporting |
| -1.097 | [-1.127;-1.08 | benign supporting |
| 1.049 | [0.933;1.121 | pathogenic moderate |
| -1.100 | [-1.112;-1.09 | benign supporting |
| -1.081 | [-1.091;-1.07 | benign supporting |
| -0.209 | [-0.296;-0.10 | indeterminate |
| -1.109 | [-1.115;-1.05 | benign supporting |
| -1.164 | [-1.169;-1.09 | benign supporting |
| -1.088 | [-1.168;-0.97 | benign supporting |
| -1.084 | [-1.165;-1.08 | benign supporting |

| | | |
|---|---|---|
| 0.000 | [-3.388;0.00 | indeterminate |
| 0.005 | [-0.256;0.64 | indeterminate |
| -1.168 | [-1.169;-0.96 | benign supporting |
| 1.276 | [1.205;1.371 | pathogenic strong |
| -1.114 | [-1.166;-0.64 | benign supporting |
| -0.516 | [-0.534;-0.49 | benign supporting |
| 1.276 | [0.894;4.992 | pathogenic strong |
| -0.424 | [-0.823;-0.02 | benign supporting |
| -1.095 | [-1.167;-0.45 | benign supporting |
| 1.276 | [-0.330;5.10 | pathogenic strong |
| -1.095 | [-1.168;-1.08 | benign supporting |
| -0.607 | [-0.639;-0.56 | benign supporting |
| 1.276 | [1.202;1.369 | pathogenic strong |
| -0.828 | [-1.082;-0.53 | benign supporting |
| -0.897 | [-0.916;-0.87 | benign supporting |
| -0.847 | [-0.870;-0.83 | benign supporting |
| -0.385 | [-0.855;0.49 | benign supporting |
| 1.276 | [-0.145;5.10 | pathogenic strong |
| -1.131 | [-1.154;-1.11 | benign supporting |
| -1.086 | [-1.167;-1.08 | benign supporting |
| 1.276 | [1.177;1.418 | pathogenic strong |
| 1.276 | [1.198;1.382 | pathogenic strong |
| -1.166 | [-1.168;-0.84 | benign supporting |
| 0.509 | [-0.153;1.03 | pathogenic supporting |
| 1.276 | [0.000;5.100 | pathogenic strong |
| -1.111 | [-1.168;-0.99 | benign supporting |
| -1.150 | [-1.169;-1.13 | benign supporting |
| -1.088 | [-1.167;-0.90 | benign supporting |
| -0.521 | [-0.586;-0.45 | benign supporting |
| -1.084 | [-1.164;-1.08 | benign supporting |
| -0.015 | [-0.126;0.28 | indeterminate |
| 1.195 | [1.134;1.262 | pathogenic moderate |
| -0.612 | [-0.642;-0.57 | benign supporting |
| 1.276 | [1.166;1.451 | pathogenic strong |
| 1.276 | [1.184;1.407 | pathogenic strong |
| 0.000 | [-3.779;0.00 | indeterminate |
| -0.357 | [-0.390;-0.32 | benign supporting |
| 1.276 | [1.249;1.305 | pathogenic strong |
| 1.128 | [0.312;1.492 | pathogenic moderate |
| -1.102 | [-1.114;-1.09 | benign supporting |
| -1.069 | [-1.083;-1.04 | benign supporting |
| 1.276 | [1.200;1.373 | pathogenic strong |
| -1.057 | [-1.070;-1.03 | benign supporting |

| | | |
|---|---|---|
| -1.090 | [-1.114;-1.06 | benign supporting |
| 0.996 | [0.434;1.215 | pathogenic moderate |
| -0.123 | [-0.277;0.135 | indeterminate |
| 1.131 | [0.702;1.310 | pathogenic moderate |
| -1.083 | [-1.090;-1.07 | benign supporting |
| -0.943 | [-0.952;-0.93 | benign supporting |
| -0.791 | [-0.907;-0.66 | benign supporting |
| -0.346 | [-1.165;4.931 | benign supporting |
| -0.366 | [-0.420;-0.30 | benign supporting |
| -0.059 | [-0.404;0.886 | indeterminate |
| 1.276 | [0.846;5.039 | pathogenic strong |
| -0.567 | [-0.604;-0.52 | benign supporting |
| -0.443 | [-0.497;-0.40 | benign supporting |
| -1.085 | [-1.092;-1.08 | benign supporting |
| -0.844 | [-0.883;-0.83 | benign supporting |
| -0.518 | [-0.561;-0.47 | benign supporting |
| -0.957 | [-1.025;-0.88 | benign supporting |
| -0.494 | [-1.095;0.753 | benign supporting |
| -1.097 | [-1.111;-1.08 | benign supporting |
| 0.947 | [0.352;1.202 | pathogenic moderate |
| -1.036 | [-1.072;-1.00 | benign supporting |
| 1.276 | [0.755;5.079 | pathogenic strong |
| -0.486 | [-0.602;-0.38 | benign supporting |
| 1.276 | [0.987;2.255 | pathogenic strong |
| -0.920 | [-0.989;-0.84 | benign supporting |
| 1.276 | [1.075;1.760 | pathogenic strong |
| -1.099 | [-1.114;-1.08 | benign supporting |
| -0.768 | [-0.851;-0.68 | benign supporting |
| -1.098 | [-1.109;-1.09 | benign supporting |
| -0.768 | [-0.848;-0.69 | benign supporting |
| -0.686 | [-0.844;-0.59 | benign supporting |
| -1.111 | [-1.115;-1.08 | benign supporting |
| -0.623 | [-0.712;-0.54 | benign supporting |
| -1.107 | [-1.115;-1.08 | benign supporting |
| -0.739 | [-0.814;-0.68 | benign supporting |
| 1.139 | [1.117;1.160 | pathogenic moderate |
| -0.577 | [-0.616;-0.53 | benign supporting |
| -1.165 | [-1.170;-1.16 | benign supporting |
| -1.110 | [-1.115;-1.08 | benign supporting |
| -0.985 | [-1.005;-0.96 | benign supporting |
| -0.196 | [-0.257;-0.12 | indeterminate |
| 1.276 | [1.045;1.930 | pathogenic strong |
| -0.621 | [-0.715;-0.54 | benign supporting |

| | | |
|---|---|---|
| 1.129 | [1.106;1.151] | pathogenic moderate |
| -1.090 | [-1.102;-1.08 | benign supporting |
| -1.106 | [-1.110;-1.10 | benign supporting |
| -1.088 | [-1.092;-1.08 | benign supporting |
| 0.010 | [-0.288;0.72( | indeterminate |
| -1.099 | [-1.112;-1.08 | benign supporting |
| -1.093 | [-1.105;-1.08 | benign supporting |
| -0.682 | [-0.707;-0.66 | benign supporting |
| -1.164 | [-1.168;-0.85 | benign supporting |
| 1.105 | [0.711;1.244] | pathogenic moderate |
| -0.970 | [-0.989;-0.95 | benign supporting |
| -0.867 | [-0.886;-0.84 | benign supporting |
| -0.473 | [-0.489;-0.45 | benign supporting |
| -0.617 | [-0.650;-0.58 | benign supporting |
| -0.957 | [-1.019;-0.89 | benign supporting |
| -1.092 | [-1.110;-1.08 | benign supporting |
| -1.110 | [-1.155;-1.08 | benign supporting |
| -1.017 | [-1.032;-1.00 | benign supporting |
| -1.056 | [-1.068;-1.03 | benign supporting |
| -0.554 | [-0.586;-0.51 | benign supporting |
| 1.276 | [1.247;1.309] | pathogenic strong |
| -1.130 | [-1.169;-1.09 | benign supporting |
| -1.089 | [-1.092;-1.08 | benign supporting |
| -0.951 | [-1.030;-0.86 | benign supporting |
| -1.085 | [-1.090;-1.08 | benign supporting |
| -1.113 | [-1.115;-1.06 | benign supporting |
| -0.502 | [-1.168;4.87: | benign supporting |
| -1.166 | [-1.169;-1.15 | benign supporting |
| -1.133 | [-1.169;-1.08 | benign supporting |
| -0.924 | [-2.829;4.74: | benign supporting |
| -1.099 | [-1.168;-0.94 | benign supporting |
| -0.944 | [-0.945;-0.94 | benign supporting |
| 1.190 | [1.141;1.243] | pathogenic moderate |
| 1.276 | [1.134;1.537] | pathogenic strong |
| 1.276 | [-0.148;5.10 | pathogenic strong |
| 1.276 | [1.120;1.598] | pathogenic strong |
| 1.276 | [0.966;2.482] | pathogenic strong |
| 1.276 | [1.236;1.321] | pathogenic strong |
| 1.276 | [1.155;1.473] | pathogenic strong |
| 1.101 | [0.948;1.157] | pathogenic moderate |
| 0.640 | [0.538;0.729] | pathogenic moderate |
| -1.102 | [-1.111;-1.09 | benign supporting |
| -0.783 | [-0.822;-0.74 | benign supporting |

| | | |
|---|---|---|
| 1.276 | [1.149;1.486 | pathogenic strong |
| -1.143 | [-1.151;-1.13 | benign supporting |
| 1.110 | [0.687;1.267 | pathogenic moderate |
| 1.276 | [1.228;1.333 | pathogenic strong |
| 1.276 | [1.241;1.317 | pathogenic strong |
| 0.844 | [0.678;1.014 | pathogenic moderate |
| 0.890 | [0.747;1.033 | pathogenic moderate |
| -1.100 | [-1.106;-1.09 | benign supporting |
| 0.392 | [-0.288;1.10 | pathogenic supporting |
| 0.059 | [-1.166;4.94 | indeterminate |
| -0.446 | [-0.501;-0.40 | benign supporting |
| 1.276 | [1.253;1.302 | pathogenic strong |
| 0.302 | [0.095;0.453 | indeterminate |
| -0.400 | [-0.433;-0.37 | benign supporting |
| -0.217 | [-0.254;-0.17 | indeterminate |
| 1.111 | [1.035;1.146 | pathogenic moderate |
| -0.594 | [-0.653;-0.53 | benign supporting |
| -0.687 | [-0.750;-0.64 | benign supporting |
| -1.091 | [-1.099;-1.08 | benign supporting |
| -1.036 | [-1.107;-0.87 | benign supporting |
| 0.312 | [0.099;0.470 | indeterminate |
| -0.724 | [-0.784;-0.67 | benign supporting |
| -1.104 | [-1.115;-1.06 | benign supporting |
| -0.246 | [-0.280;-0.20 | indeterminate |
| 1.276 | [1.218;1.347 | pathogenic strong |
| -1.089 | [-1.128;-1.08 | benign supporting |
| -0.947 | [-0.965;-0.93 | benign supporting |
| -0.511 | [-0.561;-0.46 | benign supporting |
| -1.135 | [-1.167;-1.11 | benign supporting |
| 1.118 | [0.967;1.186 | pathogenic moderate |
| 1.276 | [1.249;1.307 | pathogenic strong |
| 1.276 | [1.239;1.318 | pathogenic strong |
| 1.208 | [1.178;1.239 | pathogenic moderate |
| 1.109 | [1.084;1.122 | pathogenic moderate |
| 1.142 | [1.117;1.166 | pathogenic moderate |
| 1.276 | [1.245;1.311 | pathogenic strong |
| 1.276 | [1.245;1.311 | pathogenic strong |
| 1.276 | [1.227;1.335 | pathogenic strong |
| 0.917 | [0.761;1.059 | pathogenic moderate |
| 1.276 | [1.223;1.340 | pathogenic strong |
| 1.276 | [1.172;1.428 | pathogenic strong |
| 1.276 | [1.228;1.335 | pathogenic strong |
| 1.276 | [1.153;1.473 | pathogenic strong |

| | | |
|---|---|---|
| 1.149 | [0.829;1.303 | pathogenic moderate |
| 0.963 | [0.913;1.007 | pathogenic moderate |
| 1.276 | [1.189;1.397 | pathogenic strong |
| -0.916 | [-0.925;-0.90 | benign supporting |
| -0.608 | [-0.621;-0.59 | benign supporting |
| 1.276 | [1.246;1.310 | pathogenic strong |
| -1.081 | [-1.086;-1.07 | benign supporting |
| -1.104 | [-1.115;-1.07 | benign supporting |
| 1.276 | [-0.410;5.097 | pathogenic strong |
| -0.448 | [-0.575;-0.35 | benign supporting |
| 1.276 | [0.891;5.029 | pathogenic strong |
| 1.276 | [1.130;1.547 | pathogenic strong |
| -0.570 | [-0.825;-0.38 | benign supporting |
| -0.565 | [-0.592;-0.53 | benign supporting |
| 0.266 | [-0.082;0.622 | indeterminate |
| -1.115 | [-1.115;-1.08 | benign supporting |
| -0.828 | [-0.869;-0.76 | benign supporting |
| -0.318 | [-0.541;0.145 | indeterminate |
| 1.276 | [-1.137;4.988 | pathogenic strong |
| 0.109 | [-0.375;1.042 | indeterminate |
| -0.260 | [-0.309;-0.20 | indeterminate |
| -0.916 | [-1.073;-0.66 | benign supporting |
| 1.276 | [1.107;1.654 | pathogenic strong |
| -1.069 | [-1.096;-1.00 | benign supporting |
| 1.276 | [1.191;1.389 | pathogenic strong |
| 1.276 | [0.766;5.078 | pathogenic strong |
| -0.597 | [-0.917;-0.38 | benign supporting |
| -1.106 | [-1.115;-1.08 | benign supporting |
| -1.068 | [-1.073;-1.05 | benign supporting |
| -0.867 | [-0.919;-0.81 | benign supporting |
| -0.253 | [-0.289;-0.21 | indeterminate |
| 0.371 | [0.207;0.512 | pathogenic supporting |
| -0.039 | [-0.093;0.047 | indeterminate |
| -0.958 | [-0.978;-0.93 | benign supporting |
| -0.212 | [-0.276;-0.13 | indeterminate |
| -1.094 | [-1.106;-1.08 | benign supporting |
| -1.032 | [-1.045;-1.01 | benign supporting |
| -0.461 | [-0.477;-0.44 | benign supporting |
| -0.546 | [-0.647;-0.43 | benign supporting |
| -1.042 | [-1.079;-0.97 | benign supporting |
| -0.716 | [-0.790;-0.66 | benign supporting |
| -1.161 | [-1.170;-1.15 | benign supporting |
| -0.782 | [-1.022;-0.56 | benign supporting |

| | | |
|---|---|---|
| 1.273 | [1.195;1.379 | pathogenic strong |
| -1.068 | [-1.098;-1.00 | benign supporting |
| -0.581 | [-0.639;-0.51 | benign supporting |
| 1.265 | [1.236;1.296 | pathogenic moderate |
| -0.142 | [-0.169;-0.11 | indeterminate |
| -1.104 | [-1.115;-1.05 | benign supporting |
| -1.077 | [-1.090;-1.06 | benign supporting |
| -1.106 | [-1.112;-1.10 | benign supporting |
| -1.096 | [-1.102;-1.09 | benign supporting |
| -1.038 | [-1.056;-1.02 | benign supporting |
| -1.088 | [-1.100;-1.08 | benign supporting |
| -1.013 | [-1.051;-0.96 | benign supporting |
| -1.152 | [-1.170;-1.12 | benign supporting |
| -0.791 | [-0.911;-0.66 | benign supporting |
| -1.100 | [-1.105;-1.09 | benign supporting |
| -0.814 | [-0.848;-0.76 | benign supporting |
| -1.113 | [-1.115;-1.07 | benign supporting |
| -1.090 | [-1.101;-1.08 | benign supporting |
| -0.072 | [-0.160;0.064 | indeterminate |
| -1.084 | [-1.086;-1.08 | benign supporting |
| -1.094 | [-1.096;-1.09 | benign supporting |
| 1.116 | [1.048;1.152 | pathogenic moderate |
| -1.098 | [-1.101;-1.09 | benign supporting |
| -1.021 | [-1.051;-0.99 | benign supporting |
| -0.311 | [-0.340;-0.27 | indeterminate |
| -1.166 | [-1.170;-1.14 | benign supporting |
| -0.939 | [-1.104;-0.62 | benign supporting |
| -1.075 | [-1.080;-1.07 | benign supporting |
| 0.942 | [0.856;1.021 | pathogenic moderate |
| -0.090 | [-0.127;-0.05 | indeterminate |
| 1.109 | [1.078;1.125 | pathogenic moderate |
| -0.922 | [-0.951;-0.88 | benign supporting |
| -0.245 | [-0.355;-0.08 | indeterminate |
| 0.536 | [0.459;0.603 | pathogenic supporting |
| -0.825 | [-0.833;-0.81 | benign supporting |
| 1.276 | [1.245;1.311 | pathogenic strong |
| 1.245 | [1.167;1.336 | pathogenic moderate |
| 1.130 | [0.755;1.295 | pathogenic moderate |
| 1.276 | [1.237;1.320 | pathogenic strong |
| -0.837 | [-0.871;-0.79 | benign supporting |
| -0.936 | [-1.053;-0.79 | benign supporting |
| 1.276 | [-0.901;5.081 | pathogenic strong |
| -1.114 | [-1.115;-1.07 | benign supporting |

| | | |
|---|---|---|
| -0.942 | [-0.979;-0.90 | benign supporting |
| -0.623 | [-0.637;-0.61 | benign supporting |
| -0.834 | [-0.868;-0.78 | benign supporting |
| -1.104 | [-1.116;-0.90 | benign supporting |
| -0.631 | [-0.949;-0.41 | benign supporting |
| -1.092 | [-1.115;-1.08 | benign supporting |
| 1.276 | [-1.130;4.992 | pathogenic strong |
| -0.127 | [-1.104;4.702 | indeterminate |
| 1.161 | [1.054;1.251] | pathogenic moderate |
| -1.164 | [-1.169;-1.07 | benign supporting |
| -0.421 | [-1.166;4.885 | benign supporting |
| 0.600 | [-0.413;1.317 | pathogenic supporting |
| -1.087 | [-1.098;-1.08 | benign supporting |
| -1.168 | [-1.169;-1.02 | benign supporting |
| -0.912 | [-1.112;-0.41 | benign supporting |
| 1.183 | [1.045;1.306] | pathogenic moderate |
| -0.924 | [-1.095;-0.62 | benign supporting |
| -0.957 | [-1.096;-0.68 | benign supporting |
| -0.547 | [-0.687;-0.41 | benign supporting |
| -1.032 | [-1.113;-0.63 | benign supporting |
| -1.143 | [-1.169;-1.09 | benign supporting |
| 1.276 | [-1.136;5.007 | pathogenic strong |
| -0.381 | [-0.586;-0.11 | benign supporting |
| -1.114 | [-1.116;-1.10 | benign supporting |
| -1.088 | [-1.090;-1.08 | benign supporting |
| -0.790 | [-0.897;-0.67 | benign supporting |
| -1.047 | [-1.063;-1.03 | benign supporting |
| -0.836 | [-0.863;-0.80 | benign supporting |
| -0.900 | [-0.918;-0.88 | benign supporting |
| -1.126 | [-1.146;-1.11 | benign supporting |
| -1.089 | [-1.156;-0.68 | benign supporting |
| -0.515 | [-0.542;-0.49 | benign supporting |
| -0.737 | [-0.758;-0.71 | benign supporting |
| -0.621 | [-0.697;-0.55 | benign supporting |
| -0.914 | [-0.971;-0.84 | benign supporting |
| -0.940 | [-1.046;-0.80 | benign supporting |
| -1.086 | [-1.091;-1.08 | benign supporting |
| -0.441 | [-0.456;-0.42 | benign supporting |
| -1.035 | [-1.049;-1.02 | benign supporting |
| 1.117 | [0.794;1.238] | pathogenic moderate |
| -0.881 | [-0.909;-0.85 | benign supporting |
| 0.126 | [0.009;0.261] | indeterminate |
| -0.349 | [-0.362;-0.33 | benign supporting |

| | | |
|---|---|---|
| -0.621 | [-0.638;-0.60 | benign supporting |
| -1.087 | [-1.168;-1.06 | benign supporting |
| -1.094 | [-1.114;-1.07 | benign supporting |
| -0.874 | [-0.915;-0.83 | benign supporting |
| -1.087 | [-1.092;-1.08 | benign supporting |
| -0.426 | [-0.449;-0.40 | benign supporting |
| -1.102 | [-1.108;-1.09 | benign supporting |
| -0.937 | [-0.996;-0.87 | benign supporting |
| -0.619 | [-0.640;-0.59 | benign supporting |
| -0.670 | [-0.694;-0.64 | benign supporting |
| -1.096 | [-1.102;-1.09 | benign supporting |
| -0.572 | [-0.597;-0.54 | benign supporting |
| -0.830 | [-0.949;-0.67 | benign supporting |
| -0.833 | [-0.866;-0.78 | benign supporting |
| -0.526 | [-0.583;-0.46 | benign supporting |
| -0.556 | [-0.643;-0.45 | benign supporting |
| -0.663 | [-0.966;-0.46 | benign supporting |
| 0.046 | [-0.078;0.33 | indeterminate |
| -0.678 | [-0.851;-0.57 | benign supporting |
| 0.604 | [0.458;0.725 | pathogenic supporting |
| -0.149 | [-0.205;-0.09 | indeterminate |
| -1.110 | [-1.169;-1.08 | benign supporting |
| -1.063 | [-1.112;-1.03 | benign supporting |
| -0.329 | [-0.425;-0.18 | benign supporting |
| -1.100 | [-1.115;-1.04 | benign supporting |
| 1.276 | [1.145;1.517 | pathogenic strong |
| 0.937 | [0.850;1.017 | pathogenic moderate |
| 1.193 | [1.146;1.243 | pathogenic moderate |
| -0.822 | [-0.845;-0.78 | benign supporting |
| 1.276 | [1.213;1.356 | pathogenic strong |
| -1.084 | [-1.168;-0.98 | benign supporting |
| -1.056 | [-1.064;-1.04 | benign supporting |
| -1.084 | [-1.085;-1.08 | benign supporting |
| -0.896 | [-0.986;-0.78 | benign supporting |
| 0.798 | [0.641;0.945 | pathogenic moderate |
| -1.113 | [-1.116;-1.10 | benign supporting |
| -1.091 | [-1.096;-1.08 | benign supporting |
| -0.832 | [-1.023;-0.61 | benign supporting |
| 1.276 | [1.163;1.454 | pathogenic strong |
| -0.551 | [-0.558;-0.54 | benign supporting |
| -1.164 | [-1.165;-1.16 | benign supporting |
| -0.153 | [-0.280;-0.01 | indeterminate |
| -1.090 | [-1.095;-1.08 | benign supporting |

| | | |
|---|---|---|
| -0.276 | [-0.419;-0.04 | indeterminate |
| 1.276 | [0.856;5.030] | pathogenic strong |
| -0.405 | [-0.449;-0.36 | benign supporting |
| -1.089 | [-1.102;-1.08 | benign supporting |
| -1.098 | [-1.113;-1.08 | benign supporting |
| -0.602 | [-0.726;-0.49 | benign supporting |
| 1.276 | [1.130;1.576] | pathogenic strong |
| 1.276 | [1.228;1.334] | pathogenic strong |
| -0.877 | [-0.883;-0.87 | benign supporting |
| -1.145 | [-1.169;-0.98 | benign supporting |
| -0.806 | [-1.164;0.878 | benign supporting |
| -0.468 | [-0.775;-0.21 | benign supporting |
| -1.114 | [-1.132;-1.10 | benign supporting |
| 1.276 | [1.083;1.709] | pathogenic strong |
| -0.313 | [-1.166;4.956 | indeterminate |
| 0.945 | [0.174;1.236] | pathogenic moderate |
| 1.276 | [1.102;1.677] | pathogenic strong |
| -0.766 | [-0.928;-0.62 | benign supporting |
| 0.527 | [0.399;0.635] | pathogenic supporting |
| 0.376 | [-1.157;4.993 | pathogenic supporting |
| 1.276 | [0.574;5.090] | pathogenic strong |
| 1.276 | [1.052;1.810] | pathogenic strong |
| -0.988 | [-1.166;-0.89 | benign supporting |
| -0.836 | [-1.068;-0.57 | benign supporting |
| -0.855 | [-3.805;-0.83 | benign supporting |
| 1.257 | [0.000;5.102] | pathogenic moderate |
| 1.276 | [1.224;1.337] | pathogenic strong |
| 1.276 | [0.666;5.083] | pathogenic strong |
| -0.871 | [-1.106;-0.48 | benign supporting |
| -1.090 | [-1.157;-0.68 | benign supporting |
| 0.763 | [0.608;0.908] | pathogenic moderate |
| 1.276 | [1.188;1.393] | pathogenic strong |
| -1.075 | [-1.081;-1.07 | benign supporting |
| -1.049 | [-1.061;-1.03 | benign supporting |
| 0.441 | [0.402;0.481] | pathogenic supporting |
| -0.922 | [-0.927;-0.91 | benign supporting |
| 1.276 | [1.190;1.397] | pathogenic strong |
| -1.070 | [-1.114;-0.85 | benign supporting |
| 0.192 | [-0.034;0.431 | indeterminate |
| -1.085 | [-1.090;-1.08 | benign supporting |
| 0.765 | [0.612;0.907] | pathogenic moderate |
| 0.058 | [-0.135;0.497 | indeterminate |
| 1.276 | [0.578;5.101] | pathogenic strong |

| | | |
|---|---|---|
| -1.134 | [-1.169;-1.08 | benign supporting |
| -1.068 | [-1.109;-0.96 | benign supporting |
| -0.561 | [-1.087;0.008 | benign supporting |
| -1.090 | [-1.168;-0.54 | benign supporting |
| -1.170 | [-1.170;-1.13 | benign supporting |
| -1.099 | [-1.168;-1.08 | benign supporting |
| -0.967 | [-1.073;-0.81 | benign supporting |
| -0.962 | [-0.963;-0.96 | benign supporting |
| 1.018 | [-0.607;5.045 | pathogenic moderate |
| -0.865 | [-0.880;-0.85 | benign supporting |
| 1.185 | [1.169;1.200] | pathogenic moderate |
| -0.982 | [-0.986;-0.97 | benign supporting |
| -0.927 | [-0.936;-0.91 | benign supporting |
| -0.129 | [-0.196;-0.07 | indeterminate |
| -1.084 | [-1.088;-1.08 | benign supporting |
| -1.084 | [-1.086;-1.08 | benign supporting |
| 1.276 | [1.219;1.346] | pathogenic strong |
| -1.165 | [-1.166;-1.16 | benign supporting |
| -0.815 | [-1.109;-0.35 | benign supporting |
| 1.276 | [1.249;1.305] | pathogenic strong |
| -0.454 | [-0.496;-0.41 | benign supporting |
| 0.060 | [-0.153;0.538 | indeterminate |
| 1.276 | [1.199;1.379] | pathogenic strong |
| -1.044 | [-1.090;-1.00 | benign supporting |
| -1.169 | [-1.169;-1.09 | benign supporting |
| -1.109 | [-1.137;-1.09 | benign supporting |
| -1.084 | [-1.087;-1.08 | benign supporting |
| -0.576 | [-0.588;-0.56 | benign supporting |
| 1.244 | [1.216;1.272] | pathogenic moderate |
| -0.280 | [-0.301;-0.25 | indeterminate |
| 1.124 | [0.028;1.610] | pathogenic moderate |
| -0.937 | [-1.090;-0.66 | benign supporting |
| -0.091 | [-0.243;0.229 | indeterminate |
| -0.959 | [-0.961;-0.95 | benign supporting |
| -1.083 | [-1.147;-0.66 | benign supporting |
| -0.595 | [-0.646;-0.53 | benign supporting |
| 1.276 | [0.853;5.068] | pathogenic strong |
| -1.169 | [-1.170;-1.15 | benign supporting |
| 1.276 | [1.164;1.450] | pathogenic strong |
| -0.442 | [-0.914;0.096 | benign supporting |
| 1.268 | [0.882;3.053] | pathogenic moderate |
| -1.038 | [-1.043;-1.03 | benign supporting |
| -1.090 | [-1.100;-1.08 | benign supporting |

| | | |
|---|---|---|
| -1.094 | [-1.115;-1.06 | benign supporting |
| -0.437 | [-0.656;-0.21 | benign supporting |
| -1.100 | [-1.115;-1.08 | benign supporting |
| -0.929 | [-0.934;-0.92 | benign supporting |
| 0.828 | [0.718;0.948] | pathogenic moderate |
| -0.822 | [-1.113;-0.28 | benign supporting |
| 1.201 | [1.166;1.238] | pathogenic moderate |
| -0.266 | [-0.388;-0.07 | indeterminate |
| -1.120 | [-1.127;-1.11 | benign supporting |
| 0.801 | [0.596;1.000] | pathogenic moderate |
| -1.075 | [-1.100;-1.03 | benign supporting |
| -0.278 | [-0.400;-0.08 | indeterminate |
| 1.224 | [1.156;1.302] | pathogenic moderate |
| 1.276 | [1.253;1.301] | pathogenic strong |
| -0.614 | [-1.101;-0.06 | benign supporting |
| 1.276 | [0.850;5.039] | pathogenic strong |
| -1.096 | [-1.101;-1.09 | benign supporting |
| -0.363 | [-0.376;-0.35 | benign supporting |
| -0.675 | [-1.081;-0.37 | benign supporting |
| -1.035 | [-1.071;-0.98 | benign supporting |
| 1.276 | [0.585;5.088] | pathogenic strong |
| 1.276 | [0.445;5.101] | pathogenic strong |
| 1.276 | [1.179;1.415] | pathogenic strong |
| -1.152 | [-1.169;-1.11 | benign supporting |
| -1.086 | [-1.092;-1.08 | benign supporting |
| 0.306 | [-0.253;0.96 | indeterminate |
| 0.305 | [-1.143;4.97 | indeterminate |
| -0.459 | [-1.167;4.87 | benign supporting |
| -1.165 | [-1.169;-1.09 | benign supporting |
| -0.261 | [-0.404;-0.02 | indeterminate |
| -1.073 | [-1.094;-1.03 | benign supporting |
| -1.101 | [-1.106;-1.09 | benign supporting |
| -0.696 | [-0.891;-0.57 | benign supporting |
| -0.291 | [-0.319;-0.26 | indeterminate |
| -1.044 | [-1.057;-1.03 | benign supporting |
| 1.276 | [-1.083;5.07 | pathogenic strong |
| -0.464 | [-0.506;-0.43 | benign supporting |
| -0.878 | [-0.908;-0.85 | benign supporting |
| 1.093 | [1.009;1.130] | pathogenic moderate |
| -0.104 | [-0.149;-0.06 | indeterminate |
| -1.038 | [-1.049;-1.02 | benign supporting |
| 1.276 | [1.191;1.389] | pathogenic strong |
| -0.540 | [-0.564;-0.51 | benign supporting |

| | | |
|---|---|---|
| -0.961 | [-1.066;-0.83 | benign supporting |
| 1.276 | [1.235;1.323] | pathogenic strong |
| -1.070 | [-1.077;-1.06 | benign supporting |
| 0.305 | [-0.026;0.57ε | indeterminate |
| -0.286 | [-0.314;-0.25 | indeterminate |
| -1.037 | [-1.061;-1.01 | benign supporting |
| -0.944 | [-0.954;-0.93 | benign supporting |
| -1.090 | [-1.094;-1.08 | benign supporting |
| -0.813 | [-0.848;-0.76 | benign supporting |
| -1.117 | [-1.125;-1.11 | benign supporting |
| -0.450 | [-0.466;-0.43 | benign supporting |
| -1.001 | [-1.026;-0.96 | benign supporting |
| -0.632 | [-0.666;-0.60 | benign supporting |
| -1.023 | [-1.078;-0.93 | benign supporting |
| -1.049 | [-1.063;-1.03 | benign supporting |
| -0.115 | [-0.137;-0.09 | indeterminate |
| -0.906 | [-0.916;-0.89 | benign supporting |
| -0.539 | [-0.564;-0.51 | benign supporting |
| -1.094 | [-1.099;-1.08 | benign supporting |
| -0.547 | [-0.567;-0.52 | benign supporting |
| 1.136 | [1.099;1.172] | pathogenic moderate |
| 1.276 | [1.249;1.307] | pathogenic strong |
| -0.706 | [-0.734;-0.68 | benign supporting |
| -0.115 | [-0.149;-0.08 | indeterminate |
| -0.508 | [-0.530;-0.48 | benign supporting |
| -0.697 | [-0.715;-0.68 | benign supporting |
| -1.038 | [-1.113;-0.70 | benign supporting |
| -0.662 | [-0.864;-0.54 | benign supporting |
| -0.941 | [-1.022;-0.84 | benign supporting |
| -1.089 | [-1.115;-1.08 | benign supporting |
| 0.517 | [-0.019;0.84ε | pathogenic supporting |
| 0.594 | [-1.100;5.03( | pathogenic supporting |
| -1.124 | [-1.154;-1.10 | benign supporting |
| 1.276 | [1.243;1.313] | pathogenic strong |
| -1.087 | [-1.104;-1.07 | benign supporting |
| -0.232 | [-0.727;1.09! | indeterminate |
| -0.554 | [-0.849;-0.34 | benign supporting |
| 0.914 | [0.842;0.991] | pathogenic moderate |
| 0.899 | [0.788;1.014] | pathogenic moderate |
| -0.324 | [-0.346;-0.29 | benign supporting |
| -0.893 | [-0.996;-0.76 | benign supporting |
| 1.120 | [1.083;1.145] | pathogenic moderate |
| -0.106 | [-0.171;-0.04 | indeterminate |

| | | |
|---|---|---|
| -0.526 | [-1.094;0.442 | benign supporting |
| -0.933 | [-0.984;-0.87 | benign supporting |
| -0.116 | [-0.314;0.347 | indeterminate |
| -0.479 | [-0.569;-0.40 | benign supporting |
| -0.919 | [-1.095;-0.62 | benign supporting |
| 1.276 | [-1.102;5.044 | pathogenic strong |
| 0.936 | [-0.881;5.043 | pathogenic moderate |
| 1.276 | [-1.112;5.028 | pathogenic strong |
| -0.306 | [-0.383;-0.18 | indeterminate |
| 0.384 | [-1.109;5.022 | pathogenic supporting |
| 1.130 | [0.860;1.249] | pathogenic moderate |
| 1.276 | [-1.100;5.067 | pathogenic strong |
| 1.276 | [0.000;5.103] | pathogenic strong |
| -0.608 | [-1.166;1.677 | benign supporting |
| 1.276 | [-0.337;5.101 | pathogenic strong |
| 1.276 | [-0.192;5.103 | pathogenic strong |
| 1.276 | [-1.119;5.003 | pathogenic strong |
| 1.276 | [-1.115;5.028 | pathogenic strong |
| 0.312 | [-0.419;1.182 | indeterminate |
| -0.419 | [-0.903;0.340 | benign supporting |
| 1.276 | [0.548;5.100] | pathogenic strong |
| 1.276 | [0.521;5.095] | pathogenic strong |
| -0.488 | [-0.590;-0.39 | benign supporting |
| -0.343 | [-0.666;0.472 | benign supporting |
| -0.692 | [-0.775;-0.63 | benign supporting |
| -0.949 | [-0.973;-0.92 | benign supporting |
| 1.276 | [1.232;1.326] | pathogenic strong |
| -1.100 | [-1.114;-1.08 | benign supporting |
| -1.103 | [-1.115;-1.09 | benign supporting |
| -0.303 | [-0.345;-0.24 | indeterminate |
| -1.099 | [-1.115;-1.08 | benign supporting |
| -1.034 | [-1.058;-1.01 | benign supporting |
| -0.623 | [-0.645;-0.60 | benign supporting |
| 0.097 | [-0.848;1.497 | indeterminate |
| 0.427 | [0.186;0.614] | pathogenic supporting |
| -0.133 | [-0.192;-0.08 | indeterminate |
| -0.436 | [-0.473;-0.40 | benign supporting |
| -1.093 | [-1.110;-1.08 | benign supporting |
| 1.276 | [1.205;1.368] | pathogenic strong |
| -1.165 | [-1.169;-1.13 | benign supporting |
| 0.342 | [0.199;0.461] | pathogenic supporting |
| -0.889 | [-0.921;-0.85 | benign supporting |
| -0.790 | [-0.899;-0.67 | benign supporting |

| | | |
|---|---|---|
| -0.901 | [-3.712;0.00( | benign supporting |
| -1.084 | [-3.364;1.53( | benign supporting |
| -1.104 | [-1.115;-1.09 | benign supporting |
| -0.976 | [-1.032;-0.94 | benign supporting |
| -0.645 | [-1.166;1.42( | benign supporting |
| -1.037 | [-1.096;-0.91 | benign supporting |
| -1.159 | [-1.169;-1.07 | benign supporting |
| -0.984 | [-1.067;-0.93 | benign supporting |
| -1.068 | [-1.168;-0.91 | benign supporting |
| -0.929 | [-0.941;-0.91 | benign supporting |
| 1.266 | [-0.769;5.08 | pathogenic moderate |
| -0.430 | [-0.500;-0.37 | benign supporting |
| -0.870 | [-0.882;-0.86 | benign supporting |
| -0.921 | [-0.930;-0.91 | benign supporting |
| 1.217 | [1.131;1.316 | pathogenic moderate |
| -1.115 | [-1.167;-0.58 | benign supporting |
| 1.276 | [0.913;4.953 | pathogenic strong |
| 1.276 | [0.000;5.104 | pathogenic strong |
| 1.276 | [0.920;4.843 | pathogenic strong |
| 1.276 | [0.467;5.099 | pathogenic strong |
| 1.276 | [0.919;4.754 | pathogenic strong |
| -0.714 | [-0.943;-0.55 | benign supporting |
| -0.945 | [-0.973;-0.91 | benign supporting |
| -0.837 | [-0.846;-0.83 | benign supporting |
| -1.044 | [-1.077;-1.01 | benign supporting |
| -0.878 | [-1.069;-0.63 | benign supporting |
| 0.730 | [0.145;1.101 | pathogenic moderate |
| -0.887 | [-0.950;-0.82 | benign supporting |
| -0.869 | [-0.941;-0.85 | benign supporting |
| 1.276 | [1.074;1.772 | pathogenic strong |
| -1.054 | [-1.105;-0.93 | benign supporting |
| -1.091 | [-1.135;-1.05 | benign supporting |
| 1.276 | [0.789;5.066 | pathogenic strong |
| 1.276 | [1.109;1.666 | pathogenic strong |
| -1.154 | [-1.169;-0.92 | benign supporting |
| 0.360 | [0.162;0.514 | pathogenic supporting |
| 1.276 | [0.947;2.464 | pathogenic strong |
| -0.477 | [-0.927;-0.07 | benign supporting |
| -1.091 | [-1.097;-1.08 | benign supporting |
| 1.274 | [1.223;1.334 | pathogenic strong |
| -0.409 | [-0.448;-0.37 | benign supporting |
| -1.169 | [-1.170;-1.15 | benign supporting |
| -1.006 | [-1.017;-0.99 | benign supporting |

| | | |
|---|---|---|
| -1.163 | [-1.170;-1.14 | benign supporting |
| -1.164 | [-1.169;-1.06 | benign supporting |
| -1.085 | [-1.090;-1.08 | benign supporting |
| 1.276 | [1.256;1.298 | pathogenic strong |
| -1.090 | [-1.112;-1.08 | benign supporting |
| -1.085 | [-1.092;-1.08 | benign supporting |
| -0.426 | [-0.468;-0.39 | benign supporting |
| -0.997 | [-1.003;-0.99 | benign supporting |
| -0.817 | [-0.837;-0.78 | benign supporting |
| -1.075 | [-1.109;-1.01 | benign supporting |
| 1.224 | [1.072;1.411 | pathogenic moderate |
| -0.766 | [-0.962;-0.60 | benign supporting |
| -1.155 | [-1.169;-1.14 | benign supporting |
| -0.276 | [-0.312;-0.23 | indeterminate |
| -1.103 | [-1.115;-1.08 | benign supporting |
| -1.112 | [-1.116;-1.10 | benign supporting |
| -0.400 | [-0.483;-0.32 | benign supporting |
| 1.276 | [1.235;1.324 | pathogenic strong |
| 1.276 | [1.252;1.303 | pathogenic strong |
| 1.276 | [1.261;1.292 | pathogenic strong |
| 1.128 | [-0.233;4.96( | pathogenic moderate |
| 1.276 | [1.134;1.536 | pathogenic strong |
| 1.276 | [1.269;1.283 | pathogenic strong |
| -1.058 | [-1.070;-1.03 | benign supporting |
| 1.276 | [1.116;1.602 | pathogenic strong |
| 1.276 | [1.264;1.289 | pathogenic strong |
| 1.276 | [1.178;1.417 | pathogenic strong |
| 1.276 | [1.268;1.285 | pathogenic strong |
| 1.276 | [1.255;1.299 | pathogenic strong |
| 1.276 | [1.261;1.292 | pathogenic strong |
| 1.276 | [1.153;1.468 | pathogenic strong |
| 1.276 | [1.238;1.321 | pathogenic strong |
| -1.094 | [-1.167;-0.85 | benign supporting |
| 1.276 | [1.258;1.296 | pathogenic strong |
| 1.276 | [1.260;1.293 | pathogenic strong |
| 1.276 | [1.132;1.539 | pathogenic strong |
| 1.276 | [0.875;5.000 | pathogenic strong |
| 1.276 | [1.258;1.296 | pathogenic strong |
| -0.426 | [-0.597;-0.27 | benign supporting |
| -0.633 | [-1.070;-0.33 | benign supporting |
| -0.297 | [-0.317;-0.27 | indeterminate |
| 0.927 | [0.718;1.107 | pathogenic moderate |
| 1.276 | [0.963;2.481 | pathogenic strong |

| | | |
|---|---|---|
| 0.000 | [0.000;0.000 | indeterminate |
| -1.080 | [-1.097;-1.06 | benign supporting |
| 1.276 | [1.179;1.422 | pathogenic strong |
| -1.045 | [-1.167;-0.95 | benign supporting |
| -0.570 | [-0.947;-0.30 | benign supporting |
| -0.373 | [-0.389;-0.35 | benign supporting |
| -1.053 | [-1.063;-1.04 | benign supporting |
| -1.096 | [-1.165;-0.99 | benign supporting |
| -0.013 | [-0.697;1.305 | indeterminate |
| 1.276 | [1.252;1.302 | pathogenic strong |
| 1.276 | [1.077;1.729 | pathogenic strong |
| -0.945 | [-0.950;-0.94 | benign supporting |
| -1.110 | [-1.115;-1.10 | benign supporting |
| -0.472 | [-0.483;-0.46 | benign supporting |
| -1.166 | [-1.169;-1.15 | benign supporting |
| -1.087 | [-1.168;-1.06 | benign supporting |
| -1.119 | [-1.142;-1.09 | benign supporting |
| -1.006 | [-1.164;-0.33 | benign supporting |
| 1.276 | [-0.149;5.103 | pathogenic strong |
| -0.042 | [-0.101;0.063 | indeterminate |
| -1.066 | [-1.114;-0.69 | benign supporting |
| -0.931 | [-1.013;-0.83 | benign supporting |
| -1.088 | [-1.101;-1.07 | benign supporting |
| -0.044 | [-0.399;0.916 | indeterminate |
| 0.305 | [-0.053;0.630 | indeterminate |
| 1.276 | [0.908;5.025 | pathogenic strong |
| 1.268 | [1.227;1.314 | pathogenic moderate |
| 0.515 | [-0.636;1.683 | pathogenic supporting |
| -1.167 | [-1.169;-1.02 | benign supporting |
| 1.276 | [1.092;1.711 | pathogenic strong |
| -0.944 | [-0.951;-0.93 | benign supporting |
| -0.180 | [-0.363;0.234 | indeterminate |
| 1.276 | [1.020;1.962 | pathogenic strong |
| -1.017 | [-1.063;-0.97 | benign supporting |
| -1.169 | [-1.168;-0.91 | benign supporting |
| -3.197 | [-3.682;0.000 | benign strong |
| -0.657 | [-0.875;-0.51 | benign supporting |
| -0.289 | [-0.476;0.146 | indeterminate |
| -1.087 | [-1.088;-1.08 | benign supporting |
| -1.166 | [-1.169;-1.10 | benign supporting |
| -0.976 | [-0.991;-0.96 | benign supporting |
| -1.167 | [-1.169;-1.16 | benign supporting |
| 1.276 | [0.223;5.098 | pathogenic strong |

| | | |
|---|---|---|
| -0.024 | [-0.070;0.04( | indeterminate |
| -1.128 | [-1.168;-0.90 | benign supporting |
| 1.276 | [0.440;5.099] | pathogenic strong |
| -1.099 | [-1.166;-0.91 | benign supporting |
| 1.276 | [0.615;5.094] | pathogenic strong |
| -0.482 | [-0.724;-0.27 | benign supporting |
| -0.319 | [-0.340;-0.29 | benign supporting |
| -1.098 | [-1.114;-1.08 | benign supporting |
| -1.104 | [-1.168;-1.08 | benign supporting |
| -1.000 | [-1.071;-0.88 | benign supporting |
| -1.132 | [-1.155;-1.11 | benign supporting |
| 0.390 | [0.182;0.558] | pathogenic supporting |
| -1.066 | [-1.071;-1.05 | benign supporting |
| -0.926 | [-0.992;-0.85 | benign supporting |
| -0.574 | [-0.603;-0.54 | benign supporting |
| -1.101 | [-1.114;-1.08 | benign supporting |
| -1.114 | [-1.116;-1.10 | benign supporting |
| -1.070 | [-1.103;-1.00 | benign supporting |
| -1.104 | [-1.108;-1.09 | benign supporting |
| -1.168 | [-1.170;-1.15 | benign supporting |
| -1.034 | [-1.045;-1.02 | benign supporting |
| -0.217 | [-0.265;-0.16 | indeterminate |
| 1.276 | [1.237;1.322] | pathogenic strong |
| -1.168 | [-1.170;-1.16 | benign supporting |
| -1.087 | [-1.090;-1.08 | benign supporting |
| 0.139 | [-0.044;0.37( | indeterminate |
| -0.568 | [-0.587;-0.54 | benign supporting |
| -1.089 | [-1.168;-1.08 | benign supporting |
| -1.108 | [-1.115;-1.10 | benign supporting |
| -0.991 | [-1.091;-0.79 | benign supporting |
| -1.075 | [-1.081;-1.06 | benign supporting |
| -1.085 | [-1.098;-1.07 | benign supporting |
| -1.154 | [-1.170;-1.13 | benign supporting |
| -0.892 | [-0.906;-0.87 | benign supporting |
| -0.457 | [-0.469;-0.44 | benign supporting |
| -0.563 | [-0.587;-0.53 | benign supporting |
| -0.956 | [-0.962;-0.94 | benign supporting |
| -0.518 | [-0.573;-0.46 | benign supporting |
| -1.036 | [-1.050;-1.02 | benign supporting |
| -0.288 | [-0.309;-0.26 | indeterminate |
| -0.284 | [-0.303;-0.26 | indeterminate |
| -0.656 | [-0.875;-0.52 | benign supporting |
| 0.240 | [0.130;0.330] | indeterminate |

| | | |
|---|---|---|
| -1.084 | [-1.085;-1.08 | benign supporting |
| -0.941 | [-0.959;-0.92 | benign supporting |
| -1.096 | [-1.102;-1.09 | benign supporting |
| -0.715 | [-0.865;-0.61 | benign supporting |
| -1.105 | [-1.128;-1.09 | benign supporting |
| -0.124 | [-0.365;0.514 | indeterminate |
| -1.005 | [-1.086;-0.85 | benign supporting |
| 0.250 | [-1.160;5.000 | indeterminate |
| -1.136 | [-1.169;-1.05 | benign supporting |
| -0.841 | [-0.878;-0.79 | benign supporting |
| -0.077 | [-0.200;0.178 | indeterminate |
| -0.643 | [-0.694;-0.60 | benign supporting |
| 1.050 | [0.731;1.188] | pathogenic moderate |
| 1.276 | [-0.799;5.083 | pathogenic strong |
| -0.730 | [-1.112;-0.16 | benign supporting |
| 1.276 | [1.158;1.465] | pathogenic strong |
| 1.276 | [0.000;5.105] | pathogenic strong |
| -1.088 | [-1.168;-0.64 | benign supporting |
| -0.137 | [-1.088;1.685 | indeterminate |
| 1.147 | [0.942;1.253] | pathogenic moderate |
| -0.445 | [-1.157;1.790 | benign supporting |
| -0.807 | [-0.860;-0.73 | benign supporting |
| -0.911 | [-1.074;-0.65 | benign supporting |
| -1.084 | [-1.089;-1.08 | benign supporting |
| -0.967 | [-1.051;-0.87 | benign supporting |
| -0.577 | [-0.609;-0.54 | benign supporting |
| -1.038 | [-1.052;-1.02 | benign supporting |
| -1.093 | [-1.102;-1.08 | benign supporting |
| -0.626 | [-0.649;-0.60 | benign supporting |
| -1.090 | [-1.093;-1.08 | benign supporting |
| -1.084 | [-1.110;-1.06 | benign supporting |
| -1.114 | [-1.116;-1.09 | benign supporting |
| -0.856 | [-1.026;-0.64 | benign supporting |
| -1.104 | [-1.115;-1.09 | benign supporting |
| -1.021 | [-1.065;-0.97 | benign supporting |
| -0.621 | [-0.629;-0.61 | benign supporting |
| -1.168 | [-1.170;-1.15 | benign supporting |
| -1.170 | [-1.170;-1.15 | benign supporting |
| -0.611 | [-0.644;-0.57 | benign supporting |
| -1.087 | [-1.099;-1.07 | benign supporting |
| 1.268 | [1.230;1.310] | pathogenic moderate |
| -0.746 | [-0.843;-0.66 | benign supporting |
| -0.703 | [-0.732;-0.68 | benign supporting |

| | | |
|---|---|---|
| -0.856 | [-0.877;-0.83 | benign supporting |
| -0.857 | [-0.876;-0.83 | benign supporting |
| -0.597 | [-0.634;-0.55 | benign supporting |
| -1.086 | [-1.092;-1.08 | benign supporting |
| -1.079 | [-1.114;-0.94 | benign supporting |
| -1.089 | [-1.169;-0.97 | benign supporting |
| -0.262 | [-0.375;-0.09 | indeterminate |
| -1.076 | [-1.168;-0.99 | benign supporting |
| -0.644 | [-0.700;-0.60 | benign supporting |
| 0.538 | [-0.059;0.937 | pathogenic supporting |
| -1.083 | [-1.114;-0.99 | benign supporting |
| -0.517 | [-0.593;-0.44 | benign supporting |
| -0.621 | [-0.732;-0.53 | benign supporting |
| -0.956 | [-1.111;-0.55 | benign supporting |
| -0.838 | [-0.840;-0.83 | benign supporting |
| -0.282 | [-0.330;-0.22 | indeterminate |
| 1.276 | [-0.817;5.077 | pathogenic strong |
| -1.095 | [-1.115;-1.01 | benign supporting |
| 1.276 | [-1.096;5.055 | pathogenic strong |
| -1.155 | [-1.169;-1.07 | benign supporting |
| 1.276 | [-0.451;5.095 | pathogenic strong |
| -0.171 | [-0.514;0.819 | indeterminate |
| -1.108 | [-1.115;-1.08 | benign supporting |
| -1.111 | [-1.115;-1.07 | benign supporting |
| 1.228 | [1.054;1.453] | pathogenic moderate |
| 1.276 | [0.999;2.086] | pathogenic strong |
| 1.268 | [0.709;5.078] | pathogenic moderate |
| -0.376 | [-0.455;-0.28 | benign supporting |
| 1.276 | [1.114;1.613] | pathogenic strong |
| -0.843 | [-1.104;-0.44 | benign supporting |
| 0.798 | [0.350;1.105] | pathogenic moderate |
| 1.276 | [1.162;1.455] | pathogenic strong |
| 1.276 | [1.049;1.907] | pathogenic strong |
| 1.276 | [-0.204;5.107 | pathogenic strong |
| 0.345 | [-0.197;0.890 | pathogenic supporting |
| 1.276 | [-1.109;5.046 | pathogenic strong |
| 1.148 | [0.746;1.359] | pathogenic moderate |
| 1.276 | [1.187;1.397] | pathogenic strong |
| 1.276 | [1.225;1.338] | pathogenic strong |
| 1.276 | [-1.096;5.059 | pathogenic strong |
| 1.108 | [0.751;1.236] | pathogenic moderate |
| 1.276 | [-0.040;5.102 | pathogenic strong |
| -0.965 | [-1.070;-0.82 | benign supporting |

| | | |
|---|---|---|
| -1.074 | [-1.093;-1.04 | benign supporting |
| 1.036 | [-0.256;1.827 | pathogenic moderate |
| -0.982 | [-1.008;-0.95 | benign supporting |
| -0.044 | [-0.407;0.943 | indeterminate |
| 1.102 | [-1.065;5.070 | pathogenic moderate |
| -0.384 | [-1.166;4.855 | benign supporting |
| -0.636 | [-0.673;-0.60 | benign supporting |
| -1.143 | [-1.169;-1.11 | benign supporting |
| -1.104 | [-1.115;-1.09 | benign supporting |
| -0.240 | [-1.167;4.921 | indeterminate |
| 1.276 | [-0.632;5.093 | pathogenic strong |
| -0.474 | [-0.556;-0.40 | benign supporting |
| 1.177 | [0.319;2.235] | pathogenic moderate |
| -0.908 | [-1.065;-0.68 | benign supporting |
| -1.086 | [-1.099;-1.08 | benign supporting |
| 1.276 | [-1.130;5.019 | pathogenic strong |
| 1.276 | [-1.129;4.980 | pathogenic strong |
| -1.129 | [-1.169;-1.07 | benign supporting |
| -0.008 | [-0.278;0.668 | indeterminate |
| -1.121 | [-1.130;-1.11 | benign supporting |
| -0.272 | [-0.939;1.160 | indeterminate |
| 1.276 | [-0.216;5.107 | pathogenic strong |
| -0.098 | [-0.468;0.941 | indeterminate |
| 1.276 | [-1.123;4.983 | pathogenic strong |
| -1.113 | [-1.115;-1.06 | benign supporting |
| -0.406 | [-0.632;-0.13 | benign supporting |
| 1.250 | [-0.549;5.091 | pathogenic moderate |
| 1.276 | [-0.229;5.108 | pathogenic strong |
| -0.683 | [-0.801;-0.61 | benign supporting |
| -0.359 | [-0.548;-0.09 | benign supporting |
| 0.434 | [-0.728;1.851 | pathogenic supporting |
| -1.090 | [-1.099;-1.08 | benign supporting |
| -0.609 | [-0.691;-0.52 | benign supporting |
| 0.814 | [0.674;0.964] | pathogenic moderate |
| 1.276 | [0.340;5.098] | pathogenic strong |
| -0.783 | [-0.957;-0.61 | benign supporting |
| 1.276 | [-0.030;5.101 | pathogenic strong |
| -0.320 | [-0.394;-0.21 | benign supporting |
| 0.950 | [0.683;1.131] | pathogenic moderate |
| -1.103 | [-1.169;-0.83 | benign supporting |
| -1.101 | [-1.111;-1.09 | benign supporting |
| -0.441 | [-0.643;-0.25 | benign supporting |
| -1.085 | [-1.102;-1.08 | benign supporting |

| | | |
|---|---|---|
| -1.087 | [-1.121;-1.08 | benign supporting |
| -1.091 | [-1.112;-1.08 | benign supporting |
| -1.088 | [-1.114;-1.05 | benign supporting |
| -0.208 | [-0.265;-0.14 | indeterminate |
| -1.085 | [-1.104;-1.06 | benign supporting |
| -0.249 | [-0.669;0.952 | indeterminate |
| -0.336 | [-0.459;-0.14 | benign supporting |
| -0.715 | [-0.814;-0.64 | benign supporting |
| -0.633 | [-0.869;-0.47 | benign supporting |
| -0.656 | [-0.720;-0.61 | benign supporting |
| -1.098 | [-1.105;-1.09 | benign supporting |
| 0.309 | [-0.059;0.634 | indeterminate |
| -0.038 | [-0.428;1.000 | indeterminate |
| -0.825 | [-0.907;-0.71 | benign supporting |
| 1.276 | [1.235;1.325] | pathogenic strong |
| -0.708 | [-0.769;-0.66 | benign supporting |
| -0.585 | [-0.670;-0.49 | benign supporting |
| -0.583 | [-0.636;-0.52 | benign supporting |
| -1.000 | [-1.019;-0.97 | benign supporting |
| -0.498 | [-0.873;-0.18 | benign supporting |
| -1.021 | [-1.044;-1.00 | benign supporting |
| -1.081 | [-1.098;-1.06 | benign supporting |
| -1.093 | [-1.115;-1.05 | benign supporting |
| -0.279 | [-0.332;-0.21 | indeterminate |
| -0.877 | [-0.902;-0.85 | benign supporting |
| -1.067 | [-1.070;-1.06 | benign supporting |
| -0.803 | [-0.832;-0.77 | benign supporting |
| -1.113 | [-1.116;-1.09 | benign supporting |
| -1.104 | [-1.115;-1.09 | benign supporting |
| -0.700 | [-0.736;-0.67 | benign supporting |
| -1.101 | [-1.114;-1.09 | benign supporting |
| -1.087 | [-1.090;-1.08 | benign supporting |
| -0.536 | [-0.609;-0.45 | benign supporting |
| 0.615 | [-0.065;1.055 | pathogenic supporting |
| -1.090 | [-1.166;-1.08 | benign supporting |
| 0.725 | [0.641;0.797] | pathogenic moderate |
| -1.090 | [-1.093;-1.08 | benign supporting |
| -1.015 | [-1.065;-0.95 | benign supporting |
| -0.889 | [-0.975;-0.77 | benign supporting |
| -0.616 | [-0.670;-0.56 | benign supporting |
| -1.166 | [-1.169;-0.95 | benign supporting |
| 0.409 | [-0.046;0.759 | pathogenic supporting |
| 0.764 | [-0.044;1.167 | pathogenic moderate |

| | | |
|---|---|---|
| 1.276 | [1.185;1.402 | pathogenic strong |
| 0.558 | [-0.018;0.915 | pathogenic supporting |
| 0.995 | [0.876;1.081 | pathogenic moderate |
| 1.276 | [1.216;1.351 | pathogenic strong |
| -0.307 | [-0.437;-0.09 | indeterminate |
| 1.104 | [0.604;1.272 | pathogenic moderate |
| 0.812 | [0.520;1.078 | pathogenic moderate |
| -0.461 | [-0.524;-0.41 | benign supporting |
| 1.276 | [1.248;1.307 | pathogenic strong |
| -0.414 | [-0.436;-0.39 | benign supporting |
| -0.875 | [-0.955;-0.77 | benign supporting |
| 1.276 | [1.032;1.896 | pathogenic strong |
| -1.006 | [-1.067;-0.92 | benign supporting |
| 1.276 | [1.234;1.326 | pathogenic strong |
| -0.152 | [-0.196;-0.10 | indeterminate |
| 1.276 | [0.978;2.240 | pathogenic strong |
| -0.879 | [-1.114;-0.34 | benign supporting |
| -1.095 | [-1.098;-1.09 | benign supporting |
| -1.102 | [-1.115;-1.08 | benign supporting |
| -0.822 | [-0.839;-0.79 | benign supporting |
| -1.101 | [-1.112;-1.09 | benign supporting |
| 1.276 | [1.192;1.389 | pathogenic strong |
| -0.553 | [-0.575;-0.52 | benign supporting |
| 1.276 | [1.237;1.321 | pathogenic strong |
| 1.276 | [1.247;1.308 | pathogenic strong |
| -0.763 | [-0.830;-0.70 | benign supporting |
| -0.284 | [-0.327;-0.23 | indeterminate |
| -0.474 | [-0.492;-0.45 | benign supporting |
| -0.959 | [-0.998;-0.92 | benign supporting |
| -0.616 | [-0.657;-0.57 | benign supporting |
| 0.242 | [0.058;0.383 | indeterminate |
| -0.791 | [-0.854;-0.71 | benign supporting |
| 0.995 | [0.834;1.106 | pathogenic moderate |
| -0.797 | [-0.850;-0.73 | benign supporting |
| 1.276 | [1.235;1.323 | pathogenic strong |
| 0.860 | [0.812;0.909 | pathogenic moderate |
| -0.771 | [-0.838;-0.70 | benign supporting |
| -0.229 | [-0.419;0.298 | indeterminate |
| -1.093 | [-1.120;-1.08 | benign supporting |
| -1.092 | [-1.114;-1.08 | benign supporting |
| -0.412 | [-0.628;-0.16 | benign supporting |
| -0.987 | [-0.991;-0.98 | benign supporting |
| -0.626 | [-0.645;-0.60 | benign supporting |

| | | |
|---|---|---|
| -1.134 | [-1.169;-1.10 | benign supporting |
| -0.847 | [-3.797;0.00 | benign supporting |
| 1.162 | [0.950;1.294 | pathogenic moderate |
| -1.021 | [-1.045;-0.99 | benign supporting |
| -1.133 | [-1.144;-1.12 | benign supporting |
| -1.076 | [-1.113;-1.05 | benign supporting |
| -1.101 | [-1.115;-1.08 | benign supporting |
| -0.945 | [-0.966;-0.92 | benign supporting |
| 1.229 | [1.195;1.265 | pathogenic moderate |
| 1.276 | [0.000;5.103 | pathogenic strong |
| 0.974 | [0.563;1.173 | pathogenic moderate |
| 1.054 | [-0.321;2.87 | pathogenic moderate |
| 1.276 | [0.818;5.064 | pathogenic strong |
| -1.095 | [-1.124;-1.08 | benign supporting |
| -0.351 | [-0.363;-0.34 | benign supporting |
| -1.050 | [-1.063;-1.03 | benign supporting |
| -1.091 | [-1.168;-0.99 | benign supporting |
| 1.276 | [-0.002;5.10 | pathogenic strong |
| 0.016 | [-0.257;0.68 | indeterminate |
| -0.861 | [-0.888;-0.83 | benign supporting |
| 1.276 | [1.130;1.565 | pathogenic strong |
| -1.004 | [-1.014;-0.99 | benign supporting |
| 0.343 | [0.068;0.536 | pathogenic supporting |
| -1.062 | [-1.067;-1.05 | benign supporting |
| -0.962 | [-0.978;-0.94 | benign supporting |
| -0.510 | [-0.804;-0.29 | benign supporting |
| 1.276 | [1.150;1.490 | pathogenic strong |
| -0.270 | [-0.646;0.81 | indeterminate |
| 1.276 | [0.726;5.092 | pathogenic strong |
| 1.276 | [0.818;5.065 | pathogenic strong |
| 0.149 | [-0.109;0.54 | indeterminate |
| -0.996 | [-1.111;-0.62 | benign supporting |
| 1.276 | [0.000;5.105 | pathogenic strong |
| 1.276 | [1.180;1.416 | pathogenic strong |
| 1.276 | [1.140;1.511 | pathogenic strong |
| -0.989 | [-1.000;-0.97 | benign supporting |
| -0.985 | [-1.005;-0.96 | benign supporting |
| -0.597 | [-0.660;-0.52 | benign supporting |
| -0.082 | [-0.106;-0.05 | indeterminate |
| 1.276 | [1.266;1.287 | pathogenic strong |
| -0.272 | [-0.275;-0.26 | indeterminate |
| 1.231 | [1.224;1.238 | pathogenic moderate |
| -0.713 | [-0.716;-0.71 | benign supporting |

| | | |
|---|---|---|
| 0.391 | [0.334;0.445 | pathogenic supporting |
| -0.954 | [-0.958;-0.95 | benign supporting |
| 0.122 | [-0.155;0.602 | indeterminate |
| 1.276 | [1.148;1.486 | pathogenic strong |
| 0.289 | [0.271;0.306 | indeterminate |
| -0.419 | [-0.420;-0.41 | benign supporting |
| 0.731 | [0.723;0.738 | pathogenic moderate |
| 1.276 | [1.270;1.283 | pathogenic strong |
| -1.025 | [-1.029;-1.02 | benign supporting |
| 1.276 | [1.118;1.598 | pathogenic strong |
| 1.276 | [1.261;1.293 | pathogenic strong |
| 1.276 | [1.272;1.281 | pathogenic strong |
| -0.073 | [-0.077;-0.06 | indeterminate |
| -1.127 | [-1.169;-1.08 | benign supporting |
| -0.668 | [-0.686;-0.65 | benign supporting |
| 1.276 | [1.038;1.883 | pathogenic strong |
| -0.250 | [-0.277;-0.21 | indeterminate |
| -1.068 | [-1.085;-1.05 | benign supporting |
| 1.276 | [1.169;1.442 | pathogenic strong |
| 1.276 | [1.200;1.376 | pathogenic strong |
| 1.276 | [1.232;1.327 | pathogenic strong |
| 1.276 | [0.000;5.106 | pathogenic strong |
| 1.244 | [1.188;1.307 | pathogenic moderate |
| 1.276 | [-0.136;5.109 | pathogenic strong |
| 0.624 | [0.566;0.681 | pathogenic supporting |
| 1.276 | [1.233;1.325 | pathogenic strong |
| 1.276 | [0.406;5.096 | pathogenic strong |
| 1.149 | [1.091;1.205 | pathogenic moderate |
| 1.228 | [1.145;1.330 | pathogenic moderate |
| 1.276 | [1.220;1.341 | pathogenic strong |
| 1.276 | [1.205;1.370 | pathogenic strong |
| -1.102 | [-1.167;-1.08 | benign supporting |
| -1.104 | [-1.115;-1.08 | benign supporting |
| -0.098 | [-0.127;-0.06 | indeterminate |
| -1.087 | [-1.091;-1.08 | benign supporting |
| 0.833 | [0.794;0.876 | pathogenic moderate |
| 1.276 | [1.259;1.294 | pathogenic strong |
| -0.616 | [-0.628;-0.60 | benign supporting |
| -0.447 | [-0.472;-0.42 | benign supporting |
| -0.776 | [-0.802;-0.75 | benign supporting |
| 0.150 | [0.055;0.239 | indeterminate |
| -1.095 | [-1.100;-1.09 | benign supporting |
| -0.119 | [-0.153;-0.08 | indeterminate |

| | | |
|---|---|---|
| 0.914 | [0.855;0.978] | pathogenic moderate |
| 0.584 | [0.542;0.625] | pathogenic supporting |
| -0.722 | [-0.746;-0.70 | benign supporting |
| 1.167 | [1.122;1.217] | pathogenic moderate |
| -0.918 | [-0.954;-0.87 | benign supporting |
| 1.276 | [1.259;1.295] | pathogenic strong |
| 0.726 | [0.606;0.829] | pathogenic moderate |
| -1.071 | [-1.075;-1.06 | benign supporting |
| -0.470 | [-0.499;-0.44 | benign supporting |
| -0.520 | [-0.757;-0.33 | benign supporting |
| -0.394 | [-0.915;0.603 | benign supporting |
| -0.832 | [-0.854;-0.80 | benign supporting |
| -1.164 | [-1.170;-1.11 | benign supporting |
| 1.276 | [0.818;5.063] | pathogenic strong |
| 1.276 | [0.713;5.084] | pathogenic strong |
| 1.276 | [0.815;5.043] | pathogenic strong |
| 1.276 | [1.059;1.771] | pathogenic strong |
| -1.168 | [-1.170;-1.15 | benign supporting |
| -0.837 | [-0.840;-0.83 | benign supporting |
| -1.087 | [-1.089;-1.08 | benign supporting |
| -0.631 | [-0.936;-0.42 | benign supporting |
| 1.260 | [1.252;1.269] | pathogenic moderate |
| -0.731 | [-0.842;-0.64 | benign supporting |
| -0.912 | [-0.932;-0.89 | benign supporting |
| -1.084 | [-1.120;-1.08 | benign supporting |
| 1.276 | [1.126;1.568] | pathogenic strong |
| 1.276 | [-0.117;5.103 | pathogenic strong |
| -1.094 | [-1.167;-0.28 | benign supporting |
| -1.091 | [-1.168;-1.08 | benign supporting |
| -0.592 | [-1.096;-0.01 | benign supporting |
| -1.087 | [-1.159;-1.08 | benign supporting |
| -0.451 | [-0.577;-0.35 | benign supporting |
| -0.480 | [-0.509;-0.45 | benign supporting |
| -0.501 | [-0.563;-0.44 | benign supporting |
| -0.918 | [-0.962;-0.86 | benign supporting |
| 1.209 | [1.054;1.377] | pathogenic moderate |
| -0.583 | [-0.604;-0.56 | benign supporting |
| 0.890 | [-0.261;1.424 | pathogenic moderate |
| -0.339 | [-0.627;0.345 | benign supporting |
| -0.658 | [-0.705;-0.62 | benign supporting |
| -0.736 | [-0.761;-0.71 | benign supporting |
| -0.300 | [-0.458;-0.04 | indeterminate |
| 0.342 | [-0.366;1.143 | pathogenic supporting |

| | | |
|---|---|---|
| -1.084 | [-1.165;-1.08 | benign supporting |
| 1.276 | [0.871;5.013 | pathogenic strong |
| -1.109 | [-1.115;-1.10 | benign supporting |
| -0.823 | [-0.886;-0.73 | benign supporting |
| -1.098 | [-1.115;-1.08 | benign supporting |
| -0.723 | [-0.947;-0.56 | benign supporting |
| -0.572 | [-0.665;-0.47 | benign supporting |
| 1.276 | [1.246;1.310 | pathogenic strong |
| 0.016 | [-0.029;0.115 | indeterminate |
| -0.390 | [-0.403;-0.37 | benign supporting |
| -1.072 | [-1.083;-1.05 | benign supporting |
| 1.276 | [0.847;5.025 | pathogenic strong |
| 0.970 | [0.836;1.076 | pathogenic moderate |
| -0.361 | [-0.593;-0.02 | benign supporting |
| -0.981 | [-0.987;-0.97 | benign supporting |
| -0.689 | [-0.707;-0.67 | benign supporting |
| 1.276 | [1.202;1.374 | pathogenic strong |
| 0.707 | [0.582;0.806 | pathogenic moderate |
| 1.276 | [1.222;1.342 | pathogenic strong |
| -1.013 | [-1.024;-1.00 | benign supporting |
| 0.697 | [0.379;0.925 | pathogenic moderate |
| -1.008 | [-1.109;-0.71 | benign supporting |
| 1.276 | [-0.113;5.103 | pathogenic strong |
| 1.276 | [1.230;1.328 | pathogenic strong |
| -1.168 | [-1.169;-1.15 | benign supporting |
| 1.276 | [1.219;1.346 | pathogenic strong |
| -1.109 | [-1.112;-1.10 | benign supporting |
| -0.994 | [-0.994;-0.99 | benign supporting |
| 1.276 | [1.272;1.280 | pathogenic strong |
| 0.860 | [0.850;0.871 | pathogenic moderate |
| 1.276 | [1.269;1.284 | pathogenic strong |
| -0.508 | [-0.509;-0.50 | benign supporting |
| 1.276 | [1.274;1.278 | pathogenic strong |
| 1.276 | [1.271;1.282 | pathogenic strong |
| -0.944 | [-0.948;-0.94 | benign supporting |
| 1.276 | [1.263;1.290 | pathogenic strong |
| 1.276 | [1.272;1.280 | pathogenic strong |
| -0.255 | [-0.287;-0.21 | indeterminate |
| 1.276 | [1.219;1.346 | pathogenic strong |
| -0.863 | [-0.865;-0.86 | benign supporting |
| 1.276 | [1.269;1.283 | pathogenic strong |
| 1.276 | [1.267;1.286 | pathogenic strong |
| -0.732 | [-0.831;-0.65 | benign supporting |

| | | |
|---|---|---|
| 1.276 | [1.272;1.281] | pathogenic strong |
| 0.730 | [0.722;0.738] | pathogenic moderate |
| 1.276 | [1.269;1.283] | pathogenic strong |
| -0.758 | [-0.769;-0.74 | benign supporting |
| -1.041 | [-1.085;-0.95 | benign supporting |
| -0.952 | [-1.020;-0.87 | benign supporting |
| -1.057 | [-1.086;-0.99 | benign supporting |
| -0.700 | [-0.721;-0.68 | benign supporting |
| -0.814 | [-0.840;-0.77 | benign supporting |
| -0.898 | [-0.927;-0.86 | benign supporting |
| -0.457 | [-0.577;-0.36 | benign supporting |
| -1.030 | [-1.075;-0.96 | benign supporting |
| -0.526 | [-0.572;-0.47 | benign supporting |
| -1.081 | [-1.090;-1.07 | benign supporting |
| -1.104 | [-1.115;-0.97 | benign supporting |
| -0.688 | [-0.724;-0.65 | benign supporting |
| -0.690 | [-0.772;-0.63 | benign supporting |
| -1.165 | [-1.170;-1.15 | benign supporting |
| -0.564 | [-0.577;-0.55 | benign supporting |
| -0.204 | [-0.294;-0.09 | indeterminate |
| -1.107 | [-1.113;-1.10 | benign supporting |
| 0.449 | [-0.424;1.249 | pathogenic supporting |
| -0.581 | [-0.604;-0.56 | benign supporting |
| -0.936 | [-0.954;-0.91 | benign supporting |
| -0.024 | [-0.201;0.445 | indeterminate |
| -1.087 | [-1.117;-1.08 | benign supporting |
| -0.621 | [-1.108;0.006 | benign supporting |
| 1.276 | [1.222;1.341] | pathogenic strong |
| -1.063 | [-1.090;-0.99 | benign supporting |
| 1.113 | [-0.527;5.064 | pathogenic moderate |
| -1.112 | [-1.115;-1.08 | benign supporting |
| 1.076 | [0.705;1.217] | pathogenic moderate |
| -0.907 | [-0.953;-0.85 | benign supporting |
| 1.276 | [-0.587;5.095 | pathogenic strong |
| 1.276 | [0.000;5.104] | pathogenic strong |
| 0.135 | [-0.442;1.148 | indeterminate |
| 0.661 | [0.224;0.957] | pathogenic moderate |
| 1.276 | [1.139;1.525] | pathogenic strong |
| -1.057 | [-1.076;-1.01 | benign supporting |
| 1.276 | [0.462;5.103] | pathogenic strong |
| 1.202 | [0.929;1.448] | pathogenic moderate |
| -0.913 | [-1.108;-0.51 | benign supporting |
| 1.276 | [0.678;5.083] | pathogenic strong |

| | | |
|---|---|---|
| 1.276 | [-0.431;5.091 | pathogenic strong |
| 1.276 | [0.901;4.861 | pathogenic strong |
| -0.540 | [-0.615;-0.45 | benign supporting |
| -1.099 | [-1.169;-1.07 | benign supporting |
| -0.924 | [-1.106;-0.57 | benign supporting |
| -0.500 | [-0.631;-0.38 | benign supporting |
| -0.113 | [-0.229;0.017 | indeterminate |
| -1.107 | [-1.167;-1.08 | benign supporting |
| -1.096 | [-1.115;-1.08 | benign supporting |
| -0.399 | [-0.594;-0.17 | benign supporting |
| -0.921 | [-0.986;-0.84 | benign supporting |
| -0.113 | [-0.588;1.126 | indeterminate |
| -0.310 | [-1.148;4.893 | indeterminate |
| -0.466 | [-0.937;-0.00 | benign supporting |
| -1.111 | [-1.115;-1.09 | benign supporting |
| -1.000 | [-1.044;-0.96 | benign supporting |
| -1.077 | [-1.165;-0.45 | benign supporting |
| -1.119 | [-1.167;-1.09 | benign supporting |
| 0.372 | [-1.164;4.981 | pathogenic supporting |
| 1.130 | [0.962;1.214 | pathogenic moderate |
| -0.446 | [-0.637;-0.27 | benign supporting |
| -1.106 | [-1.167;-0.50 | benign supporting |
| -1.097 | [-1.135;-1.08 | benign supporting |
| -0.755 | [-0.794;-0.71 | benign supporting |
| -0.878 | [-0.983;-0.74 | benign supporting |
| 0.624 | [0.367;0.814 | pathogenic supporting |
| -0.268 | [-0.350;-0.14 | indeterminate |
| -1.107 | [-1.133;-1.09 | benign supporting |
| -1.069 | [-1.095;-1.01 | benign supporting |
| -0.565 | [-0.601;-0.52 | benign supporting |
| -1.141 | [-1.166;-1.12 | benign supporting |
| -0.751 | [-0.878;-0.64 | benign supporting |
| 1.276 | [1.193;1.389 | pathogenic strong |
| -0.945 | [-0.974;-0.91 | benign supporting |
| -0.679 | [-0.716;-0.64 | benign supporting |
| -1.102 | [-1.115;-0.99 | benign supporting |
| -1.084 | [-1.092;-1.08 | benign supporting |
| 0.798 | [0.693;0.909 | pathogenic moderate |
| -0.612 | [-0.650;-0.57 | benign supporting |
| -0.106 | [-0.176;-0.03 | indeterminate |
| -1.167 | [-1.169;-1.14 | benign supporting |
| -1.105 | [-1.115;-1.09 | benign supporting |
| -0.341 | [-0.352;-0.32 | benign supporting |

| | | |
|---|---|---|
| 1.276 | [1.241;1.316 | pathogenic strong |
| -0.819 | [-0.958;-0.65 | benign supporting |
| -0.060 | [-0.136;0.08 | indeterminate |
| -1.101 | [-1.134;-1.08 | benign supporting |
| -0.764 | [-0.867;-0.66 | benign supporting |
| 1.276 | [1.203;1.371 | pathogenic strong |
| -1.058 | [-1.068;-1.04 | benign supporting |
| 0.612 | [-0.053;1.04 | pathogenic supporting |
| -1.164 | [-1.167;-1.16 | benign supporting |
| -0.813 | [-1.098;-0.44 | benign supporting |
| -1.102 | [-1.115;-1.08 | benign supporting |
| 1.078 | [0.898;1.153 | pathogenic moderate |
| 1.276 | [0.942;4.931 | pathogenic strong |
| 1.276 | [1.180;1.415 | pathogenic strong |
| -0.527 | [-0.547;-0.50 | benign supporting |
| 1.276 | [1.208;1.363 | pathogenic strong |
| 1.121 | [-0.117;2.12 | pathogenic moderate |
| -0.125 | [-0.252;0.01 | indeterminate |
| -0.858 | [-0.907;-0.80 | benign supporting |
| -1.089 | [-1.091;-1.08 | benign supporting |
| -0.595 | [-0.622;-0.56 | benign supporting |
| -1.109 | [-1.111;-1.10 | benign supporting |
| -0.611 | [-0.617;-0.60 | benign supporting |
| -1.088 | [-1.097;-1.08 | benign supporting |
| -0.310 | [-0.319;-0.30 | indeterminate |
| -1.113 | [-1.115;-1.11 | benign supporting |
| -0.558 | [-0.611;-0.50 | benign supporting |
| -0.970 | [-0.983;-0.95 | benign supporting |
| -0.437 | [-0.511;-0.38 | benign supporting |
| -1.103 | [-1.109;-1.09 | benign supporting |
| -1.034 | [-1.040;-1.02 | benign supporting |
| -1.115 | [-1.119;-1.11 | benign supporting |
| -1.027 | [-1.032;-1.02 | benign supporting |
| -0.173 | [-0.402;0.45 | indeterminate |
| -0.830 | [-0.990;-0.64 | benign supporting |
| 1.276 | [1.034;1.953 | pathogenic strong |
| -0.045 | [-0.288;0.58 | indeterminate |
| -1.024 | [-1.027;-1.02 | benign supporting |
| -0.811 | [-0.831;-0.78 | benign supporting |
| 1.151 | [-0.198;5.03 | pathogenic moderate |
| -0.422 | [-0.487;-0.36 | benign supporting |
| -1.105 | [-1.112;-1.09 | benign supporting |
| -0.353 | [-0.374;-0.33 | benign supporting |

| | | |
|---|---|---|
| -1.011 | [-1.047;-0.96 | benign supporting |
| 1.276 | [1.208;1.365] | pathogenic strong |
| -1.102 | [-1.164;-0.95 | benign supporting |
| 0.047 | [-0.002;0.14[illegible] | indeterminate |
| 0.972 | [-0.205;1.45[illegible] | pathogenic moderate |
| 1.276 | [1.235;1.322] | pathogenic strong |
| 1.276 | [1.214;1.356] | pathogenic strong |
| 1.276 | [1.259;1.294] | pathogenic strong |
| -0.620 | [-1.083;-0.25 | benign supporting |
| -1.088 | [-1.092;-1.08 | benign supporting |
| 0.998 | [-1.101;5.05[illegible] | pathogenic moderate |
| -0.012 | [-0.265;0.63[illegible] | indeterminate |
| -1.088 | [-1.090;-1.08 | benign supporting |
| 1.276 | [1.142;1.520] | pathogenic strong |
| -1.166 | [-3.481;1.22[illegible] | benign supporting |
| 1.276 | [1.192;1.390] | pathogenic strong |
| 1.276 | [1.208;1.364] | pathogenic strong |
| -0.909 | [-0.932;-0.87 | benign supporting |
| 1.276 | [1.237;1.322] | pathogenic strong |
| -0.334 | [-0.382;-0.27 | benign supporting |
| -0.163 | [-0.716;1.15[illegible] | indeterminate |
| -1.090 | [-1.094;-1.08 | benign supporting |
| -0.679 | [-0.855;-0.57 | benign supporting |
| -1.156 | [-1.170;-1.13 | benign supporting |
| 1.276 | [1.245;1.312] | pathogenic strong |
| 1.204 | [1.130;1.288] | pathogenic moderate |
| 1.276 | [1.240;1.317] | pathogenic strong |
| 0.856 | [0.817;0.898] | pathogenic moderate |
| -0.557 | [-1.026;-0.17 | benign supporting |
| 0.920 | [0.627;1.125] | pathogenic moderate |
| -0.509 | [-0.962;-0.13 | benign supporting |
| 1.276 | [1.259;1.295] | pathogenic strong |
| 1.276 | [1.260;1.294] | pathogenic strong |
| -1.088 | [-1.100;-1.08 | benign supporting |
| -1.099 | [-1.114;-1.08 | benign supporting |
| -1.073 | [-1.114;-0.77 | benign supporting |
| -0.849 | [-1.016;-0.64 | benign supporting |
| -0.456 | [-0.545;-0.38 | benign supporting |
| -0.525 | [-0.546;-0.50 | benign supporting |
| 1.276 | [1.178;1.418] | pathogenic strong |
| 1.181 | [1.154;1.210] | pathogenic moderate |
| 1.119 | [0.946;1.197] | pathogenic moderate |
| -0.919 | [-3.551;1.18[illegible] | benign supporting |

| | | |
|---|---|---|
| 1.260 | [0.380;5.091] | pathogenic moderate |
| -1.093 | [-1.146;-0.74 | benign supporting |
| 0.781 | [0.729;0.835] | pathogenic moderate |
| -1.067 | [-1.072;-1.05 | benign supporting |
| -1.164 | [-3.476;0.01[illegible] | benign supporting |
| 1.276 | [1.250;1.305] | pathogenic strong |
| -0.589 | [-1.099;0.07[illegible] | benign supporting |
| 1.130 | [0.952;1.213] | pathogenic moderate |
| -0.109 | [-0.341;0.47[illegible] | indeterminate |
| 1.276 | [1.209;1.364] | pathogenic strong |
| 1.111 | [0.967;1.171] | pathogenic moderate |
| -1.090 | [-1.091;-1.08 | benign supporting |
| -0.478 | [-0.624;-0.35 | benign supporting |
| -0.522 | [-0.586;-0.45 | benign supporting |
| -1.130 | [-1.147;-1.11 | benign supporting |
| -0.852 | [-0.908;-0.78 | benign supporting |
| -0.737 | [-0.876;-0.63 | benign supporting |
| -0.277 | [-0.302;-0.25 | indeterminate |
| -1.095 | [-1.107;-1.08 | benign supporting |
| -0.593 | [-0.674;-0.50 | benign supporting |
| -0.475 | [-0.491;-0.46 | benign supporting |
| -0.800 | [-0.833;-0.76 | benign supporting |
| -1.092 | [-1.095;-1.09 | benign supporting |
| -1.101 | [-1.115;-0.96 | benign supporting |
| 1.276 | [1.128;1.570] | pathogenic strong |
| -0.628 | [-0.659;-0.60 | benign supporting |
| -0.284 | [-0.358;-0.17 | indeterminate |
| -0.829 | [-0.892;-0.74 | benign supporting |
| -1.095 | [-1.097;-1.09 | benign supporting |
| -1.095 | [-1.168;-0.96 | benign supporting |
| -1.130 | [-1.153;-1.11 | benign supporting |
| -0.435 | [-0.571;-0.31 | benign supporting |
| -1.043 | [-1.113;-0.70 | benign supporting |
| -0.742 | [-0.909;-0.61 | benign supporting |
| -0.828 | [-0.882;-0.75 | benign supporting |
| -0.916 | [-0.923;-0.90 | benign supporting |
| -1.096 | [-1.164;-1.04 | benign supporting |
| -0.760 | [-0.808;-0.71 | benign supporting |
| -1.073 | [-1.083;-1.06 | benign supporting |
| 1.276 | [-0.539;5.09[illegible] | pathogenic strong |
| -0.566 | [-0.707;-0.43 | benign supporting |
| -0.527 | [-0.606;-0.44 | benign supporting |
| -1.096 | [-1.106;-1.08 | benign supporting |

| | | |
|---|---|---|
| -1.064 | [-1.073;-1.03 | benign supporting |
| -0.668 | [-0.809;-0.58 | benign supporting |
| -1.030 | [-1.110;-0.83 | benign supporting |
| -0.949 | [-1.068;-0.78 | benign supporting |
| 1.276 | [1.212;1.356 | pathogenic strong |
| -1.113 | [-3.182;-0.22 | benign supporting |
| -1.159 | [-1.169;-1.09 | benign supporting |
| 1.276 | [0.625;5.093 | pathogenic strong |
| -1.085 | [-1.097;-1.08 | benign supporting |
| -1.031 | [-1.056;-1.00 | benign supporting |
| 1.276 | [-0.186;5.10 | pathogenic strong |
| 1.276 | [0.347;5.104 | pathogenic strong |
| 0.141 | [-1.092;4.92 | indeterminate |
| -0.365 | [-0.406;-0.31 | benign supporting |
| 1.224 | [1.131;1.341 | pathogenic moderate |
| 1.276 | [1.224;1.339 | pathogenic strong |
| -0.388 | [-0.423;-0.34 | benign supporting |
| 1.260 | [1.078;1.612 | pathogenic moderate |
| -1.088 | [-1.091;-1.08 | benign supporting |
| -1.037 | [-1.112;-0.74 | benign supporting |
| 1.276 | [0.540;5.094 | pathogenic strong |
| 1.116 | [0.351;1.420 | pathogenic moderate |
| 0.396 | [0.136;0.592 | pathogenic supporting |
| -0.608 | [-0.622;-0.58 | benign supporting |
| 1.276 | [1.171;1.434 | pathogenic strong |
| -1.096 | [-1.109;-1.08 | benign supporting |
| 1.276 | [0.765;5.092 | pathogenic strong |
| 1.276 | [1.138;1.523 | pathogenic strong |
| -1.084 | [-1.086;-1.08 | benign supporting |
| -1.073 | [-1.082;-1.06 | benign supporting |
| 1.175 | [1.150;1.203 | pathogenic moderate |
| 1.153 | [1.113;1.198 | pathogenic moderate |
| 1.276 | [1.212;1.358 | pathogenic strong |
| -0.621 | [-0.652;-0.59 | benign supporting |
| 1.080 | [0.873;1.167 | pathogenic moderate |
| 1.017 | [0.980;1.053 | pathogenic moderate |
| 0.421 | [0.009;0.705 | pathogenic supporting |
| 1.276 | [1.092;1.717 | pathogenic strong |
| -1.045 | [-1.097;-0.93 | benign supporting |
| -0.786 | [-0.904;-0.66 | benign supporting |
| 0.672 | [0.526;0.789 | pathogenic moderate |
| -1.100 | [-1.104;-1.09 | benign supporting |
| -0.322 | [-0.344;-0.29 | benign supporting |

| | | |
|---|---|---|
| 1.260 | [1.233;1.289 | pathogenic moderate |
| -0.310 | [-0.365;-0.24 | indeterminate |
| 1.276 | [1.236;1.322 | pathogenic strong |
| -0.119 | [-0.235;0.007 | indeterminate |
| 0.566 | [0.423;0.687 | pathogenic supporting |
| 1.276 | [1.156;1.476 | pathogenic strong |
| -0.995 | [-1.089;-0.82 | benign supporting |
| -1.065 | [-1.088;-1.00 | benign supporting |
| -1.166 | [-1.169;-1.10 | benign supporting |
| -1.114 | [-1.115;-1.06 | benign supporting |
| -0.910 | [-0.932;-0.88 | benign supporting |
| -1.146 | [-1.170;-1.11 | benign supporting |
| -0.980 | [-1.036;-0.91 | benign supporting |
| -0.879 | [-3.811;0.000 | benign supporting |
| -0.391 | [-0.559;-0.20 | benign supporting |
| -1.098 | [-1.114;-1.08 | benign supporting |
| -1.087 | [-1.093;-1.08 | benign supporting |
| -0.450 | [-0.898;-0.04 | benign supporting |
| -0.354 | [-0.657;0.338 | benign supporting |
| -0.656 | [-1.106;-0.16 | benign supporting |
| -0.580 | [-0.624;-0.53 | benign supporting |
| -1.089 | [-1.108;-1.07 | benign supporting |
| 0.870 | [0.026;1.210 | pathogenic moderate |
| 1.240 | [0.793;2.206 | pathogenic moderate |
| -0.659 | [-0.719;-0.61 | benign supporting |
| -0.295 | [-0.744;0.852 | indeterminate |
| -1.110 | [-1.166;-1.08 | benign supporting |
| -1.166 | [-1.169;-1.13 | benign supporting |
| -1.092 | [-1.099;-1.08 | benign supporting |
| -1.105 | [-1.108;-1.10 | benign supporting |
| -1.031 | [-1.048;-1.01 | benign supporting |
| -0.620 | [-0.634;-0.60 | benign supporting |
| -1.102 | [-1.105;-1.09 | benign supporting |
| -0.241 | [-0.260;-0.21 | indeterminate |
| -1.040 | [-1.067;-1.01 | benign supporting |
| -0.791 | [-0.808;-0.77 | benign supporting |
| -0.884 | [-0.902;-0.86 | benign supporting |
| -0.920 | [-0.931;-0.90 | benign supporting |
| -0.928 | [-0.942;-0.91 | benign supporting |
| -0.918 | [-0.930;-0.90 | benign supporting |
| -0.461 | [-0.476;-0.44 | benign supporting |
| -0.289 | [-0.310;-0.26 | indeterminate |
| -0.892 | [-0.920;-0.86 | benign supporting |

| | | |
|---|---|---|
| 1.276 | [1.231;1.329] | pathogenic strong |
| -0.941 | [-0.958;-0.92 | benign supporting |
| -0.734 | [-0.755;-0.71 | benign supporting |
| -0.947 | [-0.959;-0.93 | benign supporting |
| -1.084 | [-1.086;-1.08 | benign supporting |
| -1.084 | [-1.087;-1.08 | benign supporting |
| 1.276 | [1.262;1.291] | pathogenic strong |
| 1.253 | [1.228;1.281] | pathogenic moderate |
| 1.115 | [1.045;1.150] | pathogenic moderate |
| -1.165 | [-1.166;-1.16 | benign supporting |
| 1.078 | [1.004;1.122] | pathogenic moderate |
| 0.869 | [0.806;0.937] | pathogenic moderate |
| 1.070 | [1.041;1.100] | pathogenic moderate |
| 1.218 | [1.195;1.242] | pathogenic moderate |
| -1.103 | [-1.107;-1.09 | benign supporting |
| -0.983 | [-0.990;-0.97 | benign supporting |
| 1.099 | [1.047;1.119] | pathogenic moderate |
| -0.835 | [-0.851;-0.81 | benign supporting |
| 1.199 | [1.184;1.214] | pathogenic moderate |
| 1.257 | [1.240;1.276] | pathogenic moderate |
| 1.276 | [1.260;1.293] | pathogenic strong |
| -0.067 | [-0.089;-0.04 | indeterminate |
| 1.267 | [1.252;1.282] | pathogenic moderate |
| -0.216 | [-0.227;-0.20 | indeterminate |
| -1.097 | [-1.115;-1.07 | benign supporting |
| -0.855 | [-1.054;-0.62 | benign supporting |
| -0.666 | [-1.054;-0.40 | benign supporting |
| 1.276 | [1.219;1.348] | pathogenic strong |
| -0.766 | [-1.077;-0.46 | benign supporting |
| -1.098 | [-1.102;-1.09 | benign supporting |
| -0.201 | [-0.271;-0.11 | indeterminate |
| -0.950 | [-0.962;-0.93 | benign supporting |
| -0.804 | [-0.813;-0.79 | benign supporting |
| 1.276 | [1.215;1.354] | pathogenic strong |
| -1.064 | [-1.085;-1.00 | benign supporting |
| -0.405 | [-0.466;-0.34 | benign supporting |
| -1.110 | [-1.115;-1.10 | benign supporting |
| 1.276 | [1.108;1.684] | pathogenic strong |
| -0.952 | [-0.954;-0.94 | benign supporting |
| -0.275 | [-0.985;1.18( | indeterminate |
| 1.276 | [1.143;1.507] | pathogenic strong |
| -1.164 | [-1.169;-0.97 | benign supporting |
| 0.541 | [0.430;0.633] | pathogenic supporting |

| | | |
|---|---|---|
| -0.531 | [-0.568;-0.49 | benign supporting |
| -1.066 | [-1.069;-1.06 | benign supporting |
| -0.726 | [-0.789;-0.67 | benign supporting |
| -1.073 | [-1.083;-1.06 | benign supporting |
| 0.512 | [0.496;0.528 | pathogenic supporting |
| -1.090 | [-1.093;-1.08 | benign supporting |
| -0.298 | [-0.304;-0.29 | indeterminate |
| -0.451 | [-0.453;-0.44 | benign supporting |
| -0.914 | [-0.920;-0.90 | benign supporting |
| -0.513 | [-0.557;-0.46 | benign supporting |
| -1.169 | [-1.170;-1.16 | benign supporting |
| -1.066 | [-1.068;-1.06 | benign supporting |
| -0.734 | [-0.741;-0.72 | benign supporting |
| -0.947 | [-0.952;-0.94 | benign supporting |
| -1.051 | [-1.054;-1.04 | benign supporting |
| 1.276 | [1.259;1.294 | pathogenic strong |
| -1.106 | [-1.108;-1.10 | benign supporting |
| -0.558 | [-0.560;-0.55 | benign supporting |
| -1.068 | [-1.085;-1.02 | benign supporting |
| -0.685 | [-1.100;-0.29 | benign supporting |
| 0.926 | [0.481;1.166 | pathogenic moderate |
| 1.276 | [1.232;1.327 | pathogenic strong |
| 1.276 | [1.256;1.298 | pathogenic strong |
| -0.973 | [-0.977;-0.97 | benign supporting |
| -0.548 | [-1.168;4.758 | benign supporting |
| -1.168 | [-1.169;-1.16 | benign supporting |
| 1.042 | [0.847;1.142 | pathogenic moderate |
| -0.350 | [-0.649;0.389 | benign supporting |
| 1.276 | [1.178;1.408 | pathogenic strong |
| -0.844 | [-0.948;-0.70 | benign supporting |
| 0.670 | [-0.425;1.391 | pathogenic moderate |
| -0.366 | [-0.380;-0.35 | benign supporting |
| -1.110 | [-1.168;-0.61 | benign supporting |
| 1.276 | [1.106;1.694 | pathogenic strong |
| 1.276 | [0.685;5.091 | pathogenic strong |
| -1.104 | [-1.107;-1.10 | benign supporting |
| -0.940 | [-1.036;-0.82 | benign supporting |
| 1.276 | [1.243;1.313 | pathogenic strong |
| 1.276 | [1.184;1.403 | pathogenic strong |
| 1.276 | [1.143;1.509 | pathogenic strong |
| -0.829 | [-0.838;-0.81 | benign supporting |
| 1.276 | [1.253;1.300 | pathogenic strong |
| 0.003 | [-1.090;4.819 | indeterminate |

| | | |
|---|---|---|
| 1.276 | [1.266;1.287 | pathogenic strong |
| 1.276 | [1.066;1.774 | pathogenic strong |
| -1.002 | [-1.005;-0.99 | benign supporting |
| -1.164 | [-1.169;-1.16 | benign supporting |
| 1.276 | [1.258;1.296 | pathogenic strong |
| -0.564 | [-0.583;-0.54 | benign supporting |
| 1.276 | [1.247;1.308 | pathogenic strong |
| 1.276 | [1.180;1.407 | pathogenic strong |
| 1.276 | [1.247;1.309 | pathogenic strong |
| 1.276 | [1.044;1.936 | pathogenic strong |
| 1.276 | [1.141;1.511 | pathogenic strong |
| 1.276 | [1.233;1.325 | pathogenic strong |
| 1.276 | [1.258;1.296 | pathogenic strong |
| -1.113 | [-1.167;-0.32 | benign supporting |
| -0.463 | [-1.262;4.89 | benign supporting |
| 1.212 | [0.829;1.585 | pathogenic moderate |
| 1.276 | [1.112;1.631 | pathogenic strong |
| -1.095 | [-1.099;-1.09 | benign supporting |
| 0.435 | [0.236;0.601 | pathogenic supporting |
| 0.389 | [0.017;0.650 | pathogenic supporting |
| 1.276 | [0.647;5.091 | pathogenic strong |
| -0.947 | [-0.956;-0.93 | benign supporting |
| -1.039 | [-1.049;-1.02 | benign supporting |
| -1.093 | [-1.168;-1.07 | benign supporting |
| 0.674 | [-0.262;1.21 | pathogenic moderate |
| -0.894 | [-0.899;-0.88 | benign supporting |
| 1.276 | [1.239;1.319 | pathogenic strong |
| -0.942 | [-0.945;-0.93 | benign supporting |
| 1.193 | [1.166;1.222 | pathogenic moderate |
| 1.116 | [1.016;1.164 | pathogenic moderate |
| -1.088 | [-1.114;-1.05 | benign supporting |
| 1.183 | [0.718;1.532 | pathogenic moderate |
| 1.276 | [1.105;1.666 | pathogenic strong |
| 1.276 | [1.238;1.321 | pathogenic strong |
| 1.276 | [1.263;1.289 | pathogenic strong |
| -0.090 | [-0.155;-0.01 | indeterminate |
| 1.019 | [0.758;1.153 | pathogenic moderate |
| 1.276 | [0.996;2.188 | pathogenic strong |
| 1.257 | [1.192;1.337 | pathogenic moderate |
| -0.840 | [-0.916;-0.74 | benign supporting |
| 1.276 | [1.200;1.376 | pathogenic strong |
| -1.067 | [-1.069;-1.06 | benign supporting |
| 1.276 | [0.898;5.004 | pathogenic strong |

| | | |
|---|---|---|
| 1.276 | [1.182;1.411 | pathogenic strong |
| 1.276 | [1.171;1.436 | pathogenic strong |
| 1.276 | [1.088;1.728 | pathogenic strong |
| 1.276 | [1.200;1.376 | pathogenic strong |
| 1.276 | [1.256;1.297 | pathogenic strong |
| 0.833 | [0.763;0.907 | pathogenic moderate |
| -1.105 | [-1.115;-1.08 | benign supporting |
| 1.276 | [0.000;5.103 | pathogenic strong |
| -0.898 | [-0.956;-0.83 | benign supporting |
| 1.276 | [1.149;1.488 | pathogenic strong |
| 1.111 | [0.557;1.310 | pathogenic moderate |
| 1.108 | [0.880;1.193 | pathogenic moderate |
| 1.039 | [0.819;1.149 | pathogenic moderate |
| 0.659 | [0.338;0.881 | pathogenic moderate |
| 1.276 | [1.135;1.527 | pathogenic strong |
| -1.084 | [-1.091;-1.08 | benign supporting |
| -0.561 | [-0.678;-0.43 | benign supporting |
| -0.402 | [-0.641;-0.09 | benign supporting |
| -1.126 | [-1.132;-1.12 | benign supporting |
| -1.057 | [-1.113;-0.79 | benign supporting |
| 0.945 | [0.768;1.103 | pathogenic moderate |
| 1.276 | [1.251;1.305 | pathogenic strong |
| 0.831 | [0.652;1.017 | pathogenic moderate |
| -1.106 | [-1.107;-1.10 | benign supporting |
| -1.040 | [-1.146;-0.97 | benign supporting |
| 1.276 | [1.103;1.764 | pathogenic strong |
| 1.276 | [1.262;1.292 | pathogenic strong |
| 1.276 | [1.252;1.303 | pathogenic strong |
| -0.034 | [-0.040;-0.02 | indeterminate |
| 1.276 | [1.203;1.369 | pathogenic strong |
| 1.276 | [1.240;1.316 | pathogenic strong |
| 0.890 | [0.810;0.978 | pathogenic moderate |
| 1.115 | [1.108;1.123 | pathogenic moderate |
| -0.905 | [-0.911;-0.89 | benign supporting |
| 1.276 | [1.261;1.293 | pathogenic strong |
| -1.093 | [-1.094;-1.09 | benign supporting |
| -1.089 | [-1.090;-1.08 | benign supporting |
| 1.276 | [1.144;1.495 | pathogenic strong |
| 1.276 | [1.202;1.377 | pathogenic strong |
| 1.276 | [0.929;2.920 | pathogenic strong |
| 1.199 | [1.043;1.355 | pathogenic moderate |
| 1.276 | [0.117;5.098 | pathogenic strong |
| 0.728 | [0.681;0.771 | pathogenic moderate |

| | | |
|---|---|---|
| -0.896 | [-0.940;-0.84 | benign supporting |
| -0.956 | [-0.962;-0.95 | benign supporting |
| 0.286 | [-0.481;1.232 | indeterminate |
| 1.167 | [0.458;1.724] | pathogenic moderate |
| -0.877 | [-0.886;-0.86 | benign supporting |
| -1.084 | [-1.092;-1.08 | benign supporting |
| 1.276 | [-0.092;5.105 | pathogenic strong |
| 1.175 | [1.127;1.230] | pathogenic moderate |
| -1.092 | [-1.115;-0.89 | benign supporting |
| -0.923 | [-0.929;-0.91 | benign supporting |
| -0.268 | [-0.409;-0.03 | indeterminate |
| -0.467 | [-0.635;-0.32 | benign supporting |
| -0.618 | [-0.640;-0.59 | benign supporting |
| 1.261 | [1.208;1.322] | pathogenic moderate |
| 1.276 | [1.217;1.347] | pathogenic strong |
| -1.087 | [-1.168;-0.68 | benign supporting |
| -1.136 | [-1.168;-0.86 | benign supporting |
| 1.276 | [1.180;1.412] | pathogenic strong |
| 1.276 | [1.242;1.314] | pathogenic strong |
| 1.276 | [1.202;1.373] | pathogenic strong |
| 1.276 | [1.172;1.435] | pathogenic strong |
| 0.814 | [0.728;0.907] | pathogenic moderate |
| 1.276 | [1.022;2.087] | pathogenic strong |
| -1.011 | [-1.164;-0.38 | benign supporting |
| -0.021 | [-0.506;1.127 | indeterminate |
| 1.138 | [1.106;1.171] | pathogenic moderate |
| 1.276 | [1.201;1.374] | pathogenic strong |
| 0.319 | [0.118;0.474] | pathogenic supporting |
| 1.276 | [1.127;1.566] | pathogenic strong |
| 1.276 | [1.180;1.414] | pathogenic strong |
| 0.650 | [0.352;0.864] | pathogenic moderate |
| -0.844 | [-1.085;-0.83 | benign supporting |
| 0.204 | [-0.293;0.921 | indeterminate |
| 1.276 | [1.103;1.687] | pathogenic strong |
| -1.017 | [-1.090;-0.96 | benign supporting |
| 1.276 | [0.464;5.097] | pathogenic strong |
| -1.166 | [-1.170;-1.16 | benign supporting |
| 1.276 | [0.931;4.749] | pathogenic strong |
| 0.615 | [-1.085;5.014 | pathogenic supporting |
| -0.072 | [-0.136;0.011 | indeterminate |
| 0.756 | [-0.069;1.165 | pathogenic moderate |
| -0.965 | [-1.039;-0.88 | benign supporting |
| -0.468 | [-0.562;-0.39 | benign supporting |

| | | |
|---|---|---|
| -0.976 | [-1.108;-0.91 | benign supporting |
| -1.108 | [-1.167;-1.08 | benign supporting |
| 1.276 | [1.176;1.432 | pathogenic strong |
| 0.544 | [-0.051;0.939 | pathogenic supporting |
| 1.276 | [1.255;1.299 | pathogenic strong |
| -0.880 | [-0.888;-0.87 | benign supporting |
| 1.276 | [1.240;1.317 | pathogenic strong |
| 1.276 | [1.199;1.382 | pathogenic strong |
| -1.075 | [-1.081;-1.07 | benign supporting |
| -0.744 | [-1.075;-0.44 | benign supporting |
| 1.276 | [1.227;1.334 | pathogenic strong |
| 1.110 | [1.018;1.152 | pathogenic moderate |
| 1.125 | [0.428;1.425 | pathogenic moderate |
| -0.506 | [-0.573;-0.43 | benign supporting |
| 0.727 | [0.645;0.796 | pathogenic moderate |
| 1.276 | [1.246;1.310 | pathogenic strong |
| -0.395 | [-0.514;-0.27 | benign supporting |
| 0.677 | [0.536;0.791 | pathogenic moderate |
| -0.118 | [-0.192;-0.04 | indeterminate |
| -0.268 | [-0.417;-0.00 | indeterminate |
| 0.959 | [0.889;1.020 | pathogenic moderate |
| -0.553 | [-0.564;-0.53 | benign supporting |
| -0.703 | [-0.796;-0.64 | benign supporting |
| 1.276 | [1.222;1.342 | pathogenic strong |
| 0.388 | [-0.020;0.703 | pathogenic supporting |
| 1.276 | [1.131;1.546 | pathogenic strong |
| -0.456 | [-0.940;0.077 | benign supporting |
| 1.218 | [1.149;1.291 | pathogenic moderate |
| -1.090 | [-1.114;-0.98 | benign supporting |
| -1.095 | [-1.115;-1.00 | benign supporting |
| -0.776 | [-1.090;-0.44 | benign supporting |
| 1.276 | [1.212;1.354 | pathogenic strong |
| -1.087 | [-1.103;-1.07 | benign supporting |
| -0.488 | [-0.520;-0.45 | benign supporting |
| -1.115 | [-1.134;-0.92 | benign supporting |
| 1.231 | [-0.383;5.095 | pathogenic moderate |
| -1.104 | [-1.168;0.165 | benign supporting |
| 1.276 | [-0.563;5.094 | pathogenic strong |
| 1.276 | [1.024;2.043 | pathogenic strong |
| 0.331 | [-0.218;0.891 | pathogenic supporting |
| -1.100 | [-1.111;-1.09 | benign supporting |
| 1.276 | [1.205;1.375 | pathogenic strong |
| -0.302 | [-0.685;0.765 | indeterminate |

| | | |
|---|---|---|
| 1.276 | [1.184;1.401 | pathogenic strong |
| -0.644 | [-0.682;-0.61 | benign supporting |
| -1.058 | [-1.100;-0.95 | benign supporting |
| -0.983 | [-1.019;-0.94 | benign supporting |
| -1.110 | [-1.167;-0.51 | benign supporting |
| 1.276 | [1.197;1.380 | pathogenic strong |
| -0.920 | [-0.940;-0.89 | benign supporting |
| 1.276 | [1.103;1.709 | pathogenic strong |
| 0.470 | [-0.602;1.50 | pathogenic supporting |
| -0.594 | [-0.705;-0.48 | benign supporting |
| 1.136 | [1.000;1.215 | pathogenic moderate |
| -0.851 | [-1.164;0.68 | benign supporting |
| -1.106 | [-1.115;-1.08 | benign supporting |
| 1.276 | [-0.231;5.09 | pathogenic strong |
| 1.276 | [0.626;5.080 | pathogenic strong |
| -1.102 | [-1.109;-1.09 | benign supporting |
| -0.086 | [-0.208;0.16 | indeterminate |
| -0.634 | [-1.108;0.02 | benign supporting |
| 1.233 | [0.727;2.624 | pathogenic moderate |
| 1.187 | [1.090;1.294 | pathogenic moderate |
| 1.276 | [1.256;1.299 | pathogenic strong |
| -1.154 | [-1.169;-1.11 | benign supporting |
| -0.645 | [-0.889;-0.49 | benign supporting |
| -0.294 | [-0.348;-0.22 | indeterminate |
| -1.063 | [-1.068;-1.05 | benign supporting |
| -0.937 | [-3.467;-0.83 | benign supporting |
| 1.276 | [1.087;1.748 | pathogenic strong |
| -0.940 | [-0.945;-0.93 | benign supporting |
| -0.405 | [-0.477;-0.33 | benign supporting |
| -0.935 | [-0.943;-0.92 | benign supporting |
| -0.199 | [-0.273;-0.11 | indeterminate |
| -0.268 | [-0.389;-0.08 | indeterminate |
| 1.276 | [0.799;5.066 | pathogenic strong |
| -1.169 | [-1.170;-1.15 | benign supporting |
| 1.276 | [-0.055;5.10 | pathogenic strong |
| -0.491 | [-1.107;1.00 | benign supporting |
| -0.825 | [-1.094;-0.48 | benign supporting |
| 1.276 | [0.673;5.078 | pathogenic strong |
| -0.943 | [-0.946;-0.94 | benign supporting |
| -1.113 | [-1.115;-1.07 | benign supporting |
| -0.616 | [-0.748;-0.50 | benign supporting |
| 1.162 | [1.025;1.266 | pathogenic moderate |
| 1.079 | [0.852;1.168 | pathogenic moderate |

| | | |
|---|---|---|
| 1.276 | [1.225;1.336 | pathogenic strong |
| -0.627 | [-0.654;-0.60 | benign supporting |
| 1.276 | [1.131;1.530 | pathogenic strong |
| 1.019 | [-0.102;1.421 | pathogenic moderate |
| 1.276 | [1.241;1.316 | pathogenic strong |
| 1.177 | [0.525;1.734 | pathogenic moderate |
| -1.167 | [-1.169;-1.13 | benign supporting |
| -0.537 | [-0.561;-0.51 | benign supporting |
| 1.276 | [1.207;1.368 | pathogenic strong |
| -0.347 | [-0.421;-0.24 | benign supporting |
| 0.174 | [0.009;0.336 | indeterminate |
| 1.276 | [0.902;4.958 | pathogenic strong |
| -0.045 | [-0.333;0.69 | indeterminate |
| 1.276 | [1.242;1.316 | pathogenic strong |
| -1.109 | [-1.115;-1.10 | benign supporting |
| 1.276 | [1.226;1.335 | pathogenic strong |
| 1.276 | [1.038;1.925 | pathogenic strong |
| -1.014 | [-1.068;-0.94 | benign supporting |
| -1.103 | [-1.115;-1.08 | benign supporting |
| -0.530 | [-0.584;-0.47 | benign supporting |
| -0.601 | [-0.626;-0.57 | benign supporting |
| 1.276 | [1.241;1.314 | pathogenic strong |
| -0.902 | [-0.915;-0.88 | benign supporting |
| -0.471 | [-0.493;-0.45 | benign supporting |
| -0.423 | [-0.440;-0.40 | benign supporting |
| -0.384 | [-0.403;-0.36 | benign supporting |
| -0.167 | [-0.201;-0.13 | indeterminate |
| -0.161 | [-0.229;-0.09 | indeterminate |
| -0.669 | [-0.679;-0.66 | benign supporting |
| -0.441 | [-0.461;-0.42 | benign supporting |
| -0.429 | [-0.453;-0.40 | benign supporting |
| -0.356 | [-0.383;-0.32 | benign supporting |
| 1.276 | [1.238;1.320 | pathogenic strong |
| -0.493 | [-0.575;-0.41 | benign supporting |
| 1.276 | [1.254;1.300 | pathogenic strong |
| -0.522 | [-0.573;-0.46 | benign supporting |
| -0.232 | [-0.252;-0.20 | indeterminate |
| -0.867 | [-0.889;-0.84 | benign supporting |
| -1.092 | [-1.094;-1.09 | benign supporting |
| -0.347 | [-0.387;-0.30 | benign supporting |
| 1.276 | [1.262;1.291 | pathogenic strong |
| 1.115 | [0.901;1.201 | pathogenic moderate |
| 1.276 | [1.216;1.351 | pathogenic strong |

| | | |
|---|---|---|
| -0.996 | [-1.007;-0.97 | benign supporting |
| 0.382 | [0.309;0.451 | pathogenic supporting |
| -0.088 | [-0.131;-0.03 | indeterminate |
| 1.231 | [1.189;1.275 | pathogenic moderate |
| -0.058 | [-0.147;0.115 | indeterminate |
| -0.997 | [-1.012;-0.97 | benign supporting |
| -1.109 | [-1.115;-1.09 | benign supporting |
| 1.276 | [1.212;1.356 | pathogenic strong |
| -0.760 | [-0.793;-0.73 | benign supporting |
| 0.493 | [0.331;0.633 | pathogenic supporting |
| 1.276 | [1.192;1.393 | pathogenic strong |
| -0.340 | [-0.356;-0.32 | benign supporting |
| 1.276 | [1.241;1.317 | pathogenic strong |
| -0.398 | [-0.409;-0.38 | benign supporting |
| 0.868 | [0.822;0.916 | pathogenic moderate |
| 0.068 | [-0.077;0.361 | indeterminate |
| -1.151 | [-1.168;-1.13 | benign supporting |
| -1.041 | [-1.063;-1.02 | benign supporting |
| -0.957 | [-1.165;0.067 | benign supporting |
| -0.563 | [-0.601;-0.52 | benign supporting |
| -0.342 | [-0.563;-0.00 | benign supporting |
| -0.592 | [-0.619;-0.56 | benign supporting |
| 1.276 | [1.123;1.568 | pathogenic strong |
| -0.472 | [-0.509;-0.44 | benign supporting |
| -1.094 | [-1.100;-1.08 | benign supporting |
| -1.167 | [-1.170;-1.15 | benign supporting |
| -0.631 | [-0.773;-0.53 | benign supporting |
| 1.276 | [-1.148;4.983 | pathogenic strong |
| -0.672 | [-0.915;-0.52 | benign supporting |
| -0.142 | [-0.195;-0.09 | indeterminate |
| 1.230 | [0.870;1.766 | pathogenic moderate |
| 0.981 | [-0.353;2.541 | pathogenic moderate |
| 1.276 | [1.196;1.382 | pathogenic strong |
| -0.135 | [-0.243;-0.02 | indeterminate |
| 1.276 | [1.170;1.437 | pathogenic strong |
| 1.247 | [1.200;1.298 | pathogenic moderate |
| -1.089 | [-1.114;-1.05 | benign supporting |
| 1.276 | [0.452;5.098 | pathogenic strong |
| 1.276 | [1.122;1.575 | pathogenic strong |
| 1.115 | [-0.171;2.963 | pathogenic moderate |
| 1.167 | [-1.164;4.838 | pathogenic moderate |
| -1.101 | [-1.116;-1.09 | benign supporting |
| 0.068 | [-0.178;0.587 | indeterminate |

| | | |
|---|---|---|
| 1.156 | [1.107;1.211 | pathogenic moderate |
| 0.972 | [0.837;1.083 | pathogenic moderate |
| -0.285 | [-1.109;1.951 | indeterminate |
| -0.391 | [-0.592;-0.14 | benign supporting |
| -0.478 | [-0.671;-0.31 | benign supporting |
| 1.219 | [0.555;4.963 | pathogenic moderate |
| 1.276 | [-0.359;5.101 | pathogenic strong |
| 0.905 | [0.765;1.040 | pathogenic moderate |
| -0.019 | [-0.488;1.12( | indeterminate |
| 1.276 | [1.125;1.564 | pathogenic strong |
| -0.340 | [-0.483;-0.11 | benign supporting |
| -0.278 | [-0.527;0.41( | indeterminate |
| 1.276 | [1.177;1.415 | pathogenic strong |
| 1.191 | [0.583;1.898 | pathogenic moderate |
