## Supplemental Table S9 for "Systematically testing human HMBS missense variants to reveal mechanism and pathogenic variation"

**PopCode mutagenesis - degenerate oligo sequences**

| | |
|---|---|
| uHMBS_S1X | ACAAAAAAGTTGGCATGNNKGGTAACGGCAATGC |
| uHMBS_G2X | AAAAGTTGGCATGTCTNNKAACGGCAATGCGG |
| uHMBS_N3X | GTTGGCATGTCTGGTNNKGGCAATGCGGCTG |
| uHMBS_G4X | GGCATGTCTGGTAACNNKAATGCGGCTGCAA |
| uHMBS_N5X | ATGTCTGGTAACGGCNNKGCGGCTGCAACG |
| uHMBS_A6X | TCTGGTAACGGCAATNNKGCTGCAACGGCG |
| uHMBS_A7X | GTAACGGCAATGCGNNKGCAACGGCGGAAG |
| uHMBS_A8X | GGCAATGCGGCTNNKACGGCGGAAGAAAA |
| uHMBS_T9X | CAATGCGGCTGCANNKGCGGAAGAAAACAGC |
| uHMBS_A10X | GCGGCTGCAACGNNKGAAGAAAACAGCCCAA |
| uHMBS_E11X | GCTGCAACGGCGNNKGAAAACAGCCCAAAGA |
| uHMBS_E12X | TGCAACGGCGGAANNKAACAGCCCAAAGATGA |
| uHMBS_N13X | CAACGGCGGAAGAANNKAGCCCAAAGATGAGAG |
| uHMBS_S14X | CGGCGGAAGAAAACNNKCCAAAGATGAGAGTGAT |
| uHMBS_P15X | GCGGAAGAAAACAGCNNKAAGATGAGAGTGATTCG |
| uHMBS_K16X | GAAGAAAACAGCCCANNKATGAGAGTGATTCGCG |
| uHMBS_M17X | AGAAAACAGCCCAAAGNNKAGAGTGATTCGCGTG |
| uHMBS_R18X | AAACAGCCCAAAGATGNNKGTGATTCGCGTGGG |
| uHMBS_V19X | AGCCCAAAGATGAGANNKATTCGCGTGGGTAC |
| uHMBS_I20X | CCCAAAGATGAGAGTGNNKCGCGTGGGTACCC |
| uHMBS_R21X | CCAAAGATGAGAGTGATTNNKGTGGGTACCCGCA |
| uHMBS_V22X | GATGAGAGTGATTCGCNNKGGTACCCGCAAGAG |
| uHMBS_G23X | AGAGTGATTCGCGTGNNKACCCGCAAGAGCC |
| uHMBS_T24X | GATTCGCGTGGGTNNKCGCAAGAGCCAGC |
| uHMBS_R25X | TCGCGTGGGTACCNNKAAGAGCCAGCTTGC |
| uHMBS_K26X | CGTGGGTACCCGCNNKAGCCAGCTTGCTC |
| uHMBS_S27X | TGGGTACCCGCAAGNNKCAGCTTGCTCGCA |
| uHMBS_Q28X | GTACCCGCAAGAGCNNKCTTGCTCGCATACAG |
| uHMBS_L29X | CCGCAAGAGCCAGNNKGCTCGCATACAGACG |
| uHMBS_A30X | GCAAGAGCCAGCTTNNKCGCATACAGACGGA |
| uHMBS_R31X | AGAGCCAGCTTGCTNNKATACAGACGGACAGTG |
| uHMBS_I32X | CCAGCTTGCTCGCNNKCAGACGGACAGTGTG |
| uHMBS_Q33X | AGCTTGCTCGCATANNKACGGACAGTGTGGT |
| uHMBS_T34X | CTTGCTCGCATACAGNNKGACAGTGTGGTGGC |
| uHMBS_D35X | GCTCGCATACAGACGNNKAGTGTGGTGGCAAC |
| uHMBS_S36X | CGCATACAGACGGACNNKGTGGTGGCAACATTG |
| uHMBS_V37X | CATACAGACGGACAGTNNKGTGGCAACATTGAAAG |
| uHMBS_V38X | AGACGGACAGTGTGNNKGCAACATTGAAAGCCT |
| uHMBS_A39X | CGGACAGTGTGGTGNNKACATTGAAAGCCTCG |
| uHMBS_T40X | ACAGTGTGGTGGCANNKTTGAAAGCCTCGTACC |
| uHMBS_L41X | GTGTGGTGGCAACANNKAAAGCCTCGTACCCT |
| uHMBS_K42X | GTGGTGGCAACATTGNNKGCCTCGTACCCTGG |
| uHMBS_A43X | GGTGGCAACATTGAAANNKTCGTACCCTGGCC |
| uHMBS_S44X | GCAACATTGAAAGCCNNKTACCCTGGCCTGC |

| | |
|---|---|
| uHMBS_Y45X | AACATTGAAAGCCTCGNNKCCTGGCCTGCAGT |
| uHMBS_P46X | ATTGAAAGCCTCGTACNNKGGCCTGCAGTTTGA |
| uHMBS_G47X | AAGCCTCGTACCCTNNKCTGCAGTTTGAAATCAT |
| uHMBS_L48X | CCTCGTACCCTGGCNNKCAGTTTGAAATCATTGCT |
| uHMBS_Q49X | CGTACCCTGGCCTGNNKTTTGAAATCATTGCTATGT |
| uHMBS_F50X | CCCTGGCCTGCAGNNKGAAATCATTGCTATGTCC |
| uHMBS_E51X | CTGGCCTGCAGTTTNNKATCATTGCTATGTCCAC |
| uHMBS_I52X | GGCCTGCAGTTTGAANNKATTGCTATGTCCACCA |
| uHMBS_I53X | CCTGCAGTTTGAAATCNNKGCTATGTCCACCACA |
| uHMBS_A54X | CTGCAGTTTGAAATCATTNNKATGTCCACCACAGG |
| uHMBS_M55X | CAGTTTGAAATCATTGCTNNKTCCACCACAGGGG |
| uHMBS_S56X | GTTTGAAATCATTGCTATGNNKACCACAGGGGACAA |
| uHMBS_T57X | GAAATCATTGCTATGTCCNNKACAGGGGACAAGATTC |
| uHMBS_T58X | CATTGCTATGTCCACCNNKGGGGACAAGATTCTTG |
| uHMBS_G59X | GCTATGTCCACCACANNKGACAAGATTCTTGATACTG |
| uHMBS_D60X | TGTCCACCACAGGGNNKAAGATTCTTGATACTGCA |
| uHMBS_K61X | CCACCACAGGGGACNNKATTCTTGATACTGCACT |
| uHMBS_I62X | CCACAGGGGACAAGNNKCTTGATACTGCACTCTC |
| uHMBS_L63X | CACAGGGGACAAGATTNNKGATACTGCACTCTCTAAG |
| uHMBS_D64X | AGGGGACAAGATTCTTNNKACTGCACTCTCTAAGAT |
| uHMBS_T65X | GGGACAAGATTCTTGATNNKGCACTCTCTAAGATTGG |
| uHMBS_A66X | GACAAGATTCTTGATACTNNKCTCTCTAAGATTGGAGAG |
| uHMBS_L67X | AAGATTCTTGATACTGCANNKTCTAAGATTGGAGAGAAAA |
| uHMBS_S68X | TTCTTGATACTGCACTCNNKAAGATTGGAGAGAAAAGC |
| uHMBS_K69X | TGATACTGCACTCTCTNNKATTGGAGAGAAAAGCC |
| uHMBS_I70X | GATACTGCACTCTCTAAGNNKGGAGAGAAAAGCCTGT |
| uHMBS_G71X | CTGCACTCTCTAAGATTNNKGAGAAAAGCCTGTTTAC |
| uHMBS_E72X | CACTCTCTAAGATTGGANNKAAAAGCCTGTTTACCAA |
| uHMBS_K73X | CTCTCTAAGATTGGAGAGNNKAGCCTGTTTACCAAGG |
| uHMBS_S74X | TCTAAGATTGGAGAGAAANNKCTGTTTACCAAGGAGC |
| uHMBS_L75X | AGATTGGAGAGAAAAGCNNKTTTACCAAGGAGCTTG |
| uHMBS_F76X | TGGAGAGAAAAGCCTGNNKACCAAGGAGCTTGAA |
| uHMBS_T77X | GAGAGAAAAGCCTGTTTNNKAAGGAGCTTGAACATG |
| uHMBS_K78X | AGAAAAGCCTGTTTACCNNKGAGCTTGAACATGCC |
| uHMBS_E79X | AAGCCTGTTTACCAAGNNKCTTGAACATGCCCTG |
| uHMBS_L80X | CCTGTTTACCAAGGAGNNKGAACATGCCCTGGA |
| uHMBS_E81X | GTTTACCAAGGAGCTTNNKCATGCCCTGGAGAAG |
| uHMBS_H82X | TACCAAGGAGCTTGAANNKGCCCTGGAGAAGAAT |
| uHMBS_A83X | CAAGGAGCTTGAACATNNKCTGGAGAAGAATGAAGT |
| uHMBS_L84X | GAGCTTGAACATGCCNNKGAGAAGAATGAAGTGGA |
| uHMBS_E85X | CTTGAACATGCCCTGNNKAAGAATGAAGTGGACC |
| uHMBS_K86X | AACATGCCCTGGAGNNKAATGAAGTGGACCTGG |
| uHMBS_N87X | CATGCCCTGGAGAAGNNKGAAGTGGACCTGGTT |
| uHMBS_E88X | GCCCTGGAGAAGAATNNKGTGGACCTGGTTGTT |
| uHMBS_V89X | CCTGGAGAAGAATGAANNKGACCTGGTTGTTCAC |

| | |
|---|---|
| uHMBS_D90X | TGGAGAAGAATGAAGTGNNKCTGGTTGTTCACTCCT |
| uHMBS_L91X | AGAAGAATGAAGTGGACNNKGTTGTTCACTCCTTGAA |
| uHMBS_V92X | GAATGAAGTGGACCTGNNKGTTCACTCCTTGAAGG |
| uHMBS_V93X | GAAGTGGACCTGGTTNNKCACTCCTTGAAGGACC |
| uHMBS_H94X | GTGGACCTGGTTGTTNNKTCCTTGAAGGACCTG |
| uHMBS_S95X | GACCTGGTTGTTCACNNKTTGAAGGACCTGCC |
| uHMBS_L96X | CTGGTTGTTCACTCCNNKAAGGACCTGCCCAC |
| uHMBS_K97X | GGTTGTTCACTCCTTGNNKGACCTGCCCACTGT |
| uHMBS_D98X | TTGTTCACTCCTTGAAGNNKCTGCCCACTGTGC |
| uHMBS_L99X | TCACTCCTTGAAGGACNNKCCCACTGTGCTTCC |
| uHMBS_P100X | TCCTTGAAGGACCTGNNKACTGTGCTTCCTCCT |
| uHMBS_T101X | TGAAGGACCTGCCCNNKGTGCTTCCTCCTGG |
| uHMBS_V102X | GGACCTGCCCACTNNKCTTCCTCCTGGCTTC |
| uHMBS_L103X | CCTGCCCACTGTGNNKCCTCCTGGCTTCAC |
| uHMBS_P104X | GCCCACTGTGCTTNNKCCTGGCTTCACCATC |
| uHMBS_P105X | CCACTGTGCTTCCTNNKGGCTTCACCATCGG |
| uHMBS_G106X | ACTGTGCTTCCTCCTNNKTTCACCATCGGAGC |
| uHMBS_F107X | TGCTTCCTCCTGGCNNKACCATCGGAGCCAT |
| uHMBS_T108X | CTTCCTCCTGGCTTCNNKATCGGAGCCATCTG |
| uHMBS_I109X | CTCCTGGCTTCACCNNKGGAGCCATCTGCAA |
| uHMBS_G110X | CCTGGCTTCACCATCNNKGCCATCTGCAAGCG |
| uHMBS_A111X | GCTTCACCATCGGANNKATCTGCAAGCGGGA |
| uHMBS_I112X | TCACCATCGGAGCCNNKTGCAAGCGGGAAAA |
| uHMBS_C113X | CCATCGGAGCCATCNNKAAGCGGGAAAACCC |
| uHMBS_K114X | CGGAGCCATCTGCNNKCGGGAAAACCCTCAT |
| uHMBS_R115X | GGAGCCATCTGCAAGNNKGAAAACCCTCATGATGC |
| uHMBS_E116X | CCATCTGCAAGCGGNNKAACCCTCATGATGCT |
| uHMBS_N117X | CTGCAAGCGGGAANNKCCTCATGATGCTGTTG |
| uHMBS_P118X | GCAAGCGGGAAAACNNKCATGATGCTGTTGTCT |
| uHMBS_H119X | AGCGGGAAAACCCTNNKGATGCTGTTGTCTTTCA |
| uHMBS_D120X | CGGGAAAACCCTCATNNKGCTGTTGTCTTTCACC |
| uHMBS_A121X | GGAAAACCCTCATGATNNKGTTGTCTTTCACCCAA |
| uHMBS_V122X | AAACCCTCATGATGCTNNKGTCTTTCACCCAAAATTT |
| uHMBS_V123X | CCCTCATGATGCTGTTNNKTTTCACCCAAAATTTGTT |
| uHMBS_F124X | TCATGATGCTGTTGTCNNKCACCCAAAATTTGTTGG |
| uHMBS_H125X | ATGATGCTGTTGTCTTTNNKCCAAAATTTGTTGGGAA |
| uHMBS_P126X | TGCTGTTGTCTTTCACNNKAAATTTGTTGGGAAGAC |
| uHMBS_K127X | TGTTGTCTTTCACCCANNKTTTGTTGGGAAGACCC |
| uHMBS_F128X | TTGTCTTTCACCCAAAANNKGTTGGGAAGACCCTAG |
| uHMBS_V129X | GTCTTTCACCCAAAATTTNNKGGGAAGACCCTAGAAA |
| uHMBS_G130X | TTTCACCCAAAATTTGTTNNKAAGACCCTAGAAACCC |
| uHMBS_K131X | CCCAAAATTTGTTGGGNNKACCCTAGAAACCCTG |
| uHMBS_T132X | CAAAATTTGTTGGGAAGNNKCTAGAAACCCTGCCA |
| uHMBS_L133X | ATTTGTTGGGAAGACCNNKGAAACCCTGCCAGAG |
| uHMBS_E134X | GTTGGGAAGACCCTANNKACCCTGCCAGAGAA |

| | |
|---|---|
| uHMBS_T135X | TGGGAAGACCCTAGAANNKCTGCCAGAGAAGAGT |
| uHMBS_L136X | GAAGACCCTAGAAACCNNKCCAGAGAAGAGTGTGG |
| uHMBS_P137X | GACCCTAGAAACCCTGNNKGAGAAGAGTGTGGTGG |
| uHMBS_E138X | CTAGAAACCCTGCCANNKAAGAGTGTGGTGGGA |
| uHMBS_K139X | GAAACCCTGCCAGAGNNKAGTGTGGTGGGAAC |
| uHMBS_S140X | CCCTGCCAGAGAAGNNKGTGGTGGGAACCAG |
| uHMBS_V141X | CTGCCAGAGAAGAGTNNKGTGGGAACCAGCTC |
| uHMBS_V142X | CCAGAGAAGAGTGTGNNKGGAACCAGCTCCCT |
| uHMBS_G143X | AGAGAAGAGTGTGGTGNNKACCAGCTCCCTGC |
| uHMBS_T144X | AGAGTGTGGTGGGANNKAGCTCCCTGCGAA |
| uHMBS_S145X | GTGTGGTGGGAACCNNKTCCCTGCGAAGAGC |
| uHMBS_S146X | GGTGGGAACCAGCNNKCTGCGAAGAGCAGC |
| uHMBS_L147X | TGGGAACCAGCTCCNNKCGAAGAGCAGCCC |
| uHMBS_R148X | GAACCAGCTCCCTGNNKAGAGCAGCCCAGC |
| uHMBS_R149X | CAGCTCCCTGCGANNKGCAGCCCAGCTG |
| uHMBS_A150X | GCTCCCTGCGAAGANNKGCCCAGCTGCAGA |
| uHMBS_A151X | CCTGCGAAGAGCANNKCAGCTGCAGAGAAAG |
| uHMBS_Q152X | GCGAAGAGCAGCCNNKCTGCAGAGAAAGTTCC |
| uHMBS_L153X | GAAGAGCAGCCCAGNNKCAGAGAAAGTTCCCG |
| uHMBS_Q154X | AGCAGCCCAGCTGNNKAGAAAGTTCCCGCAT |
| uHMBS_R155X | AGCCCAGCTGCAGNNKAAGTTCCCGCATCTG |
| uHMBS_K156X | CCCAGCTGCAGAGANNKTTCCCGCATCTGGA |
| uHMBS_F157X | CAGCTGCAGAGAAAGNNKCCGCATCTGGAGTT |
| uHMBS_P158X | GCTGCAGAGAAAGTTCNNKCATCTGGAGTTCAGGA |
| uHMBS_H159X | CAGAGAAAGTTCCCGNNKCTGGAGTTCAGGAGTA |
| uHMBS_L160X | AGAAAGTTCCCGCATNNKGAGTTCAGGAGTATTCG |
| uHMBS_E161X | AGTTCCCGCATCTGNNKTTCAGGAGTATTCGGG |
| uHMBS_F162X | TCCCGCATCTGGAGNNKAGGAGTATTCGGGGA |
| uHMBS_R163X | CCGCATCTGGAGTTCNNKAGTATTCGGGGAAACC |
| uHMBS_S164X | CATCTGGAGTTCAGGNNKATTCGGGGAAACCTC |
| uHMBS_I165X | CTGGAGTTCAGGAGTNNKCGGGGAAACCTCAA |
| uHMBS_R166X | TGGAGTTCAGGAGTATTNNKGGAAACCTCAACACC |
| uHMBS_G167X | GTTCAGGAGTATTCGGNNKAACCTCAACACCCG |
| uHMBS_N168X | AGGAGTATTCGGGGANNKCTCAACACCCGGC |
| uHMBS_L169X | GAGTATTCGGGGAAACNNKAACACCCGGCTTC |
| uHMBS_N170X | ATTCGGGGAAACCTCNNKACCCGGCTTCGG |
| uHMBS_T171X | CGGGGAAACCTCAACNNKCGGCTTCGGAAGC |
| uHMBS_R172X | GGAAACCTCAACACCNNKCTTCGGAAGCTGGAC |
| uHMBS_L173X | ACCTCAACACCCGGNNKCGGAAGCTGGACG |
| uHMBS_R174X | TCAACACCCGGCTTNNKAAGCTGGACGAGCA |
| uHMBS_K175X | ACCCGGCTTCGGNNKCTGGACGAGCAGC |
| uHMBS_L176X | CCGGCTTCGGAAGNNKGACGAGCAGCAGG |
| uHMBS_D177X | GGCTTCGGAAGCTGNNKGAGCAGCAGGAGTT |
| uHMBS_E178X | TTCGGAAGCTGGACNNKCAGCAGGAGTTCAGT |
| uHMBS_Q179X | GGAAGCTGGACGAGNNKCAGGAGTTCAGTGCC |

| | |
|---|---|
| uHMBS_Q180X | GCTGGACGAGCAGNNKGAGTTCAGTGCCATC |
| uHMBS_E181X | TGGACGAGCAGCAGNNKTTCAGTGCCATCATCC |
| uHMBS_F182X | ACGAGCAGCAGGAGNNKAGTGCCATCATCCTG |
| uHMBS_S183X | GAGCAGCAGGAGTTCNNKGCCATCATCCTGGC |
| uHMBS_A184X | CAGCAGGAGTTCAGTNNKATCATCCTGGCAACAG |
| uHMBS_I185X | CAGGAGTTCAGTGCCNNKATCCTGGCAACAGCT |
| uHMBS_I186X | GGAGTTCAGTGCCATCNNKCTGGCAACAGCTGG |
| uHMBS_L187X | AGTTCAGTGCCATCATCNNKGCAACAGCTGGCC |
| uHMBS_A188X | CAGTGCCATCATCCTGNNKACAGCTGGCCTGC |
| uHMBS_T189X | GCCATCATCCTGGCANNKGCTGGCCTGCAGC |
| uHMBS_A190X | CATCATCCTGGCAACANNKGGCCTGCAGCGC |
| uHMBS_G191X | ATCCTGGCAACAGCTNNKCTGCAGCGCATGG |
| uHMBS_L192X | GGCAACAGCTGGCNNKCAGCGCATGGGCT |
| uHMBS_Q193X | CAACAGCTGGCCTGNNKCGCATGGGCTGGC |
| uHMBS_R194X | AGCTGGCCTGCAGNNKATGGGCTGGCACAA |
| uHMBS_M195X | GGCCTGCAGCGCNNKGGCTGGCACAACC |
| uHMBS_G196X | CCTGCAGCGCATGNNKTGGCACAACCGGG |
| uHMBS_W197X | GCAGCGCATGGGCNNKCACAACCGGGTGGG |
| uHMBS_H198X | GCGCATGGGCTGGNNKAACCGGGTGGGGC |
| uHMBS_N199X | CATGGGCTGGCACNNKCGGGTGGGGCAGA |
| uHMBS_R200X | TGGGCTGGCACAACNNKGTGGGGCAGATCCT |
| uHMBS_V201X | CTGGCACAACCGGNNKGGGCAGATCCTGCA |
| uHMBS_G202X | GCACAACCGGGTGNNKCAGATCCTGCACCCT |
| uHMBS_Q203X | CAACCGGGTGGGGNNKATCCTGCACCCTGAG |
| uHMBS_I204X | CGGGTGGGGCAGNNKCTGCACCCTGAGGAA |
| uHMBS_L205X | GGGTGGGGCAGATCNNKCACCCTGAGGAATGC |
| uHMBS_H206X | TGGGGCAGATCCTGNNKCCTGAGGAATGCATGT |
| uHMBS_P207X | GGCAGATCCTGCACNNKGAGGAATGCATGTATGC |
| uHMBS_E208X | CAGATCCTGCACCCTNNKGAATGCATGTATGCTGT |
| uHMBS_E209X | ATCCTGCACCCTGAGNNKTGCATGTATGCTGTGG |
| uHMBS_C210X | CTGCACCCTGAGGAANNKATGTATGCTGTGGGC |
| uHMBS_M211X | CACCCTGAGGAATGCNNKTATGCTGTGGGCCA |
| uHMBS_Y212X | CCCTGAGGAATGCATGNNKGCTGTGGGCCAGG |
| uHMBS_A213X | CCTGAGGAATGCATGTATNNKGTGGGCCAGGGGG |
| uHMBS_V214X | AGGAATGCATGTATGCTNNKGGCCAGGGGGCC |
| uHMBS_G215X | AATGCATGTATGCTGTGNNKCAGGGGGCCTTGG |
| uHMBS_Q216X | ATGTATGCTGTGGGCNNKGGGGCCTTGGGC |
| uHMBS_G217X | ATGCTGTGGGCCAGNNKGCCTTGGGCGTGG |
| uHMBS_A218X | TGTGGGCCAGGGGNNKTTGGGCGTGGAAGT |
| uHMBS_L219X | GGCCAGGGGGCCNNKGGCGTGGAAGTGC |
| uHMBS_G220X | CCAGGGGGCCTTGNNKGTGGAAGTGCGAGC |
| uHMBS_V221X | GGGGCCTTGGGCNNKGAAGTGCGAGCCAAG |
| uHMBS_E222X | GGCCTTGGGCGTGNNKGTGCGAGCCAAGGA |
| uHMBS_V223X | CCTTGGGCGTGGAANNKCGAGCCAAGGACCA |
| uHMBS_R224X | TGGGCGTGGAAGTGNNKGCCAAGGACCAGGA |

| | |
|---|---|
| uHMBS_A225X | GCGTGGAAGTGCGANNKAAGGACCAGGACATCT |
| uHMBS_K226X | TGGAAGTGCGAGCCNNKGACCAGGACATCTTGG |
| uHMBS_D227X | GAAGTGCGAGCCAAGNNKCAGGACATCTTGGATCT |
| uHMBS_Q228X | TGCGAGCCAAGGACNNKGACATCTTGGATCTGGT |
| uHMBS_D229X | CGAGCCAAGGACCAGNNKATCTTGGATCTGGTGG |
| uHMBS_I230X | GCCAAGGACCAGGACNNKTTGGATCTGGTGGGT |
| uHMBS_L231X | CAAGGACCAGGACATCNNKGATCTGGTGGGTGTG |
| uHMBS_D232X | GGACCAGGACATCTTGNNKCTGGTGGGTGTGCT |
| uHMBS_L233X | CCAGGACATCTTGGATNNKGTGGGTGTGCTGCA |
| uHMBS_V234X | AGGACATCTTGGATCTGNNKGGTGTGCTGCACG |
| uHMBS_G235X | ACATCTTGGATCTGGTGNNKGTGCTGCACGATCC |
| uHMBS_V236X | TTGGATCTGGTGGGTNNKCTGCACGATCCCGA |
| uHMBS_L237X | GATCTGGTGGGTGTGNNKCACGATCCCGAGACT |
| uHMBS_H238X | TGGTGGGTGTGCTGNNKGATCCCGAGACTCTG |
| uHMBS_D239X | TGGGTGTGCTGCACNNKCCCGAGACTCTGCTT |
| uHMBS_P240X | GTGTGCTGCACGATNNKGAGACTCTGCTTCGC |
| uHMBS_E241X | TGCTGCACGATCCCNNKACTCTGCTTCGCTG |
| uHMBS_T242X | TGCACGATCCCGAGNNKCTGCTTCGCTGCAT |
| uHMBS_L243X | CACGATCCCGAGACTNNKCTTCGCTGCATCGC |
| uHMBS_L244X | CGATCCCGAGACTCTGNNKCGCTGCATCGCTG |
| uHMBS_R245X | CCCGAGACTCTGCTTNNKTGCATCGCTGAAAGG |
| uHMBS_C246X | GAGACTCTGCTTCGCNNKATCGCTGAAAGGGC |
| uHMBS_I247X | CTCTGCTTCGCTGCNNKGCTGAAAGGGCCTTC |
| uHMBS_A248X | TGCTTCGCTGCATCNNKGAAAGGGCCTTCCTG |
| uHMBS_E249X | TTCGCTGCATCGCTNNKAGGGCCTTCCTGAG |
| uHMBS_R250X | GCTGCATCGCTGAANNKGCCTTCCTGAGGCA |
| uHMBS_A251X | TGCATCGCTGAAAGGNNKTTCCTGAGGCACCTG |
| uHMBS_F252X | TCGCTGAAAGGGCCNNKCTGAGGCACCTGGAA |
| uHMBS_L253X | GCTGAAAGGGCCTTCNNKAGGCACCTGGAAGG |
| uHMBS_R254X | GAAAGGGCCTTCCTGNNKCACCTGGAAGGAGGC |
| uHMBS_H255X | GGGCCTTCCTGAGGNNKCTGGAAGGAGGCTGC |
| uHMBS_L256X | GCCTTCCTGAGGCACNNKGAAGGAGGCTGCAGT |
| uHMBS_E257X | TTCCTGAGGCACCTGNNKGGAGGCTGCAGTGT |
| uHMBS_G258X | CTGAGGCACCTGGAANNKGGCTGCAGTGTGC |
| uHMBS_G259X | GGCACCTGGAAGGANNKTGCAGTGTGCCAGT |
| uHMBS_C260X | ACCTGGAAGGAGGCNNKAGTGTGCCAGTAGCC |
| uHMBS_S261X | TGGAAGGAGGCTGCNNKGTGCCAGTAGCCGT |
| uHMBS_V262X | GAAGGAGGCTGCAGTNNKCCAGTAGCCGTGCA |
| uHMBS_P263X | GAGGCTGCAGTGTGNNKGTAGCCGTGCATACA |
| uHMBS_V264X | GCTGCAGTGTGCCANNKGCCGTGCATACAGC |
| uHMBS_A265X | TGCAGTGTGCCAGTANNKGTGCATACAGCTATGAAG |
| uHMBS_V266X | AGTGTGCCAGTAGCCNNKCATACAGCTATGAAGGATG |
| uHMBS_H267X | TGCCAGTAGCCGTGNNKACAGCTATGAAGGATGG |
| uHMBS_T268X | CCAGTAGCCGTGCATNNKGCTATGAAGGATGGGC |
| uHMBS_A269X | GTAGCCGTGCATACANNKATGAAGGATGGGCAAC |

| | |
|---|---|
| uHMBS_M270X | CCGTGCATACAGCTNNKAAGGATGGGCAACTG |
| uHMBS_K271X | CGTGCATACAGCTATGNNKGATGGGCAACTGTACC |
| uHMBS_D272X | GTGCATACAGCTATGAAGNNKGGGCAACTGTACCTG |
| uHMBS_G273X | CATACAGCTATGAAGGATNNKCAACTGTACCTGACTGG |
| uHMBS_Q274X | CAGCTATGAAGGATGGGNNKCTGTACCTGACTGGAG |
| uHMBS_L275X | CTATGAAGGATGGGCAANNKTACCTGACTGGAGGAG |
| uHMBS_Y276X | GAAGGATGGGCAACTGNNKCTGACTGGAGGAGTCT |
| uHMBS_L277X | GGATGGGCAACTGTACNNKACTGGAGGAGTCTGG |
| uHMBS_T278X | GGGCAACTGTACCTGNNKGGAGGAGTCTGGAGT |
| uHMBS_G279X | GCAACTGTACCTGACTNNKGGAGTCTGGAGTCTAGA |
| uHMBS_G280X | ACTGTACCTGACTGGANNKGTCTGGAGTCTAGACG |
| uHMBS_V281X | GTACCTGACTGGAGGANNKTGGAGTCTAGACGGC |
| uHMBS_W282X | CCTGACTGGAGGAGTCNNKAGTCTAGACGGCTCAG |
| uHMBS_S283X | ACTGGAGGAGTCTGGNNKCTAGACGGCTCAGATAG |
| uHMBS_L284X | GGAGGAGTCTGGAGTNNKGACGGCTCAGATAGC |
| uHMBS_D285X | GAGGAGTCTGGAGTCTANNKGGCTCAGATAGCATACA |
| uHMBS_G286X | GAGTCTGGAGTCTAGACNNKTCAGATAGCATACAAGAGA |
| uHMBS_S287X | TGGAGTCTAGACGGCNNKGATAGCATACAAGAGACC |
| uHMBS_D288X | GAGTCTAGACGGCTCANNKAGCATACAAGAGACCAT |
| uHMBS_S289X | TCTAGACGGCTCAGATNNKATACAAGAGACCATGCA |
| uHMBS_I290X | GACGGCTCAGATAGCNNKCAAGAGACCATGCAGG |
| uHMBS_Q291X | CGGCTCAGATAGCATANNKGAGACCATGCAGGC |
| uHMBS_E292X | GGCTCAGATAGCATACAANNKACCATGCAGGCTACC |
| uHMBS_T293X | CTCAGATAGCATACAAGAGNNKATGCAGGCTACCATC |
| uHMBS_M294X | GATAGCATACAAGAGACCNNKCAGGCTACCATCCATG |
| uHMBS_Q295X | GCATACAAGAGACCATGNNKGCTACCATCCATGTCC |
| uHMBS_A296X | ACAAGAGACCATGCAGNNKACCATCCATGTCCCT |
| uHMBS_T297X | GAGACCATGCAGGCTNNKATCCATGTCCCTGCC |
| uHMBS_I298X | ACCATGCAGGCTACCNNKCATGTCCCTGCCCA |
| uHMBS_H299X | CATGCAGGCTACCATCNNKGTCCCTGCCCAGC |
| uHMBS_V300X | GCAGGCTACCATCCATNNKCCTGCCCAGCATGA |
| uHMBS_P301X | GGCTACCATCCATGTCNNKGCCCAGCATGAAGAT |
| uHMBS_A302X | TACCATCCATGTCCCTNNKCAGCATGAAGATGGCC |
| uHMBS_Q303X | ATCCATGTCCCTGCCNNKCATGAAGATGGCCCTG |
| uHMBS_H304X | ATGTCCCTGCCCAGNNKGAAGATGGCCCTGAG |
| uHMBS_E305X | TCCCTGCCCAGCATNNKGATGGCCCTGAGGAT |
| uHMBS_D306X | CCTGCCCAGCATGAANNKGGCCCTGAGGATGA |
| uHMBS_G307X | GCCCAGCATGAAGATNNKCCTGAGGATGACCCA |
| uHMBS_P308X | CCAGCATGAAGATGGCNNKGAGGATGACCCACAGT |
| uHMBS_E309X | CATGAAGATGGCCCTNNKGATGACCCACAGTTGG |
| uHMBS_D310X | GAAGATGGCCCTGAGNNKGACCCACAGTTGGTAG |
| uHMBS_D311X | GATGGCCCTGAGGATNNKCCACAGTTGGTAGGC |
| uHMBS_P312X | GGCCCTGAGGATGACNNKCAGTTGGTAGGCATCA |
| uHMBS_Q313X | CCTGAGGATGACCCANNKTTGGTAGGCATCACTG |
| uHMBS_L314X | TGAGGATGACCCACAGNNKGTAGGCATCACTGCTC |

| | |
|---|---|
| uHMBS_V315X | GGATGACCCACAGTTGNNKGGCATCACTGCTCG |
| uHMBS_G316X | TGACCCACAGTTGGTANNKATCACTGCTCGTAACAT |
| uHMBS_I317X | CCACAGTTGGTAGGCNNKACTGCTCGTAACATTCC |
| uHMBS_T318X | ACAGTTGGTAGGCATCNNKGCTCGTAACATTCCAC |
| uHMBS_A319X | GTTGGTAGGCATCACTNNKCGTAACATTCCACGAG |
| uHMBS_R320X | GTAGGCATCACTGCTNNKAACATTCCACGAGGG |
| uHMBS_N321X | GCATCACTGCTCGTNNKATTCCACGAGGGCC |
| uHMBS_I322X | CATCACTGCTCGTAACNNKCCACGAGGGCCC |
| uHMBS_P323X | TCACTGCTCGTAACATTNNKCGAGGGCCCCAGT |
| uHMBS_R324X | TGCTCGTAACATTCCANNKGGGCCCCAGTTGG |
| uHMBS_G325X | CTCGTAACATTCCACGANNKCCCCAGTTGGCTGC |
| uHMBS_P326X | TAACATTCCACGAGGGNNKCAGTTGGCTGCCCA |
| uHMBS_Q327X | TCCACGAGGGCCCNNKTTGGCTGCCCAGAA |
| uHMBS_L328X | ACGAGGGCCCCAGNNKGCTGCCCAGAACTTG |
| uHMBS_A329X | GAGGGCCCCAGTTGNNKGCCCAGAACTTGGG |
| uHMBS_A330X | GCCCCAGTTGGCTNNKCAGAACTTGGGCATCA |
| uHMBS_Q331X | CCCAGTTGGCTGCCNNKAACTTGGGCATCAGC |
| uHMBS_N332X | AGTTGGCTGCCCAGNNKTTGGGCATCAGCCT |
| uHMBS_L333X | TGGCTGCCCAGAACNNKGGCATCAGCCTGGC |
| uHMBS_G334X | GCTGCCCAGAACTTGNNKATCAGCCTGGCCAA |
| uHMBS_I335X | CCCAGAACTTGGGCNNKAGCCTGGCCAACTT |
| uHMBS_S336X | CCAGAACTTGGGCATCNNKCTGGCCAACTTGTTG |
| uHMBS_L337X | AACTTGGGCATCAGCNNKGCCAACTTGTTGCTG |
| uHMBS_A338X | TGGGCATCAGCCTGNNKAACTTGTTGCTGAGCA |
| uHMBS_N339X | GCATCAGCCTGGCCNNKTTGTTGCTGAGCAAAG |
| uHMBS_L340X | TCAGCCTGGCCAACNNKTTGCTGAGCAAAGGAG |
| uHMBS_L341X | GCCTGGCCAACTTGNNKCTGAGCAAAGGAGCC |
| uHMBS_L342X | CTGGCCAACTTGTTGNNKAGCAAAGGAGCCAAAA |
| uHMBS_S343X | GCCAACTTGTTGCTGNNKAAAGGAGCCAAAAACAT |
| uHMBS_K344X | CAACTTGTTGCTGAGCNNKGGAGCCAAAAACATCC |
| uHMBS_G345X | CTTGTTGCTGAGCAAANNKGCCAAAAACATCCTGG |
| uHMBS_A346X | GTTGCTGAGCAAAGGANNKAAAAACATCCTGGATGTT |
| uHMBS_K347X | CTGAGCAAAGGAGCCNNKAACATCCTGGATGTTGC |
| uHMBS_N348X | GAGCAAAGGAGCCAAANNKATCCTGGATGTTGCAC |
| uHMBS_I349X | GCAAAGGAGCCAAAAACNNKCTGGATGTTGCACGG |
| uHMBS_L350X | AAGGAGCCAAAAACATCNNKGATGTTGCACGGCA |
| uHMBS_D351X | GAGCCAAAAACATCCTGNNKGTTGCACGGCAGC |
| uHMBS_V352X | CCAAAAACATCCTGGATNNKGCACGGCAGCTTAA |
| uHMBS_A353X | AAAAACATCCTGGATGTTNNKCGGCAGCTTAACGAT |
| uHMBS_R354X | CATCCTGGATGTTGCANNKCAGCTTAACGATGCCC |
| uHMBS_Q355X | CTGGATGTTGCACGGNNKCTTAACGATGCCCATTAA |
| uHMBS_L356X | ATGTTGCACGGCAGNNKAACGATGCCCATTAAC |
| uHMBS_N357X | TGCACGGCAGCTTNNKGATGCCCATTAACCAAC |
| uHMBS_D358X | GCACGGCAGCTTAACNNKGCCCATTAACCAACTTT |
| uHMBS_A359X | CGGCAGCTTAACGATNNKCATTAACCAACTTTCTTGTA |

uHMBS_H360X CAGCTTAACGATGCCNNKTAACCAACTTTCTTGTACA

**TileSeq primer sequences**

| | |
|---|---|
| HMBS_1F | TACACGACGCTCTTCCGATCTCAAGTTTGTACAAAAAAGTTGGCATG |
| HMBS_1R | AGACGTGTGCTCTTCCGATCTTTCAATGTTGCCACCACACT |
| HMBS_2F | TACACGACGCTCTTCCGATCTGCTCGCATACAGACGGAC |
| HMBS_2R | AGACGTGTGCTCTTCCGATCTGCTCCTTGGTAAACAGGCTT |
| HMBS_3F | TACACGACGCTCTTCCGATCTCTGCACTCTCTAAGATTGGAGAG |
| HMBS_3R | AGACGTGTGCTCTTCCGATCTCGCTTGCAGATGGCTCCG |
| HMBS_4F | TACACGACGCTCTTCCGATCTCTTCCTCCTGGCTTCACC |
| HMBS_4R | AGACGTGTGCTCTTCCGATCTGCTGGGCTGCTCTTCGCA |
| HMBS_5F | TACACGACGCTCTTCCGATCTGTGGTGGGAACCAGCTCC |
| HMBS_5R | AGACGTGTGCTCTTCCGATCTCAGCTGTTGCCAGGATGAT |
| HMBS_6F | TACACGACGCTCTTCCGATCTCAGCAGGAGTTCAGTGCC |
| HMBS_6R | AGACGTGTGCTCTTCCGATCTCCTGGTCCTTGGCTCGCA |
| HMBS_7F | TACACGACGCTCTTCCGATCTGGGGCCTTGGGCGTGGAA |
| HMBS_7R | AGACGTGTGCTCTTCCGATCTGCACGGCTACTGGCACAC |
| HMBS_8F | TACACGACGCTCTTCCGATCTCACCTGGAAGGAGGCTGC |
| HMBS_8R | AGACGTGTGCTCTTCCGATCTCATGCTGGGCAGGGACAT |
| HMBS_9F | TACACGACGCTCTTCCGATCTGACCATGCAGGCTACCATC |
| HMBS_9R | AGACGTGTGCTCTTCCGATCTCAGCAACAAGTTGGCCAG |
| HMBS_10F | TACACGACGCTCTTCCGATCTCAGAACTTGGGCATCAGC |
| HMBS_10R | AGACGTGTGCTCTTCCGATCTAGAGCTCGACGTCTTACTTACTTA |

**TileSeq ancil**

uHMBS_stop
eHMBS_stop
HMBS_stop_
M13F:
M13Fext:
M13Rext:
BP_F1:
BP_NoStop_

**lary primers**

GGGGACAACTTTGTACAAAAAAGTTGGCATGAGAGTGATTCGCGTGGG
GGGGACAACTTTGTACAAAAAAGTTGGCATGTCTGGTAACGGCAATGCG
GGGGACAACTTTGTACAAGAAAGTTGGTTAATGGGCATCGTTAAGCTGCCGTG
GTAAAACGACGGCCAGT
GTAAAACGACGGCCAGTCTTAA
CAGGAAACAGCTATGACCATGT
GGGGACAACTTTGTACAAAAAAGTTGGC
GGGGACAACTTTGTACAAGAAAGTTGG
